# Activation of programmed cell death and counter-defense functions of phage accessory genes

**DOI:** 10.1101/2023.04.06.535777

**Authors:** Sukrit Silas, Héloïse Carion, Kira S. Makarova, Eric Laderman, David Sanchez Godinez, Matthew Johnson, Andrea Fossati, Danielle Swaney, Michael Bocek, Eugene V. Koonin, Joseph Bondy-Denomy

## Abstract

Viruses have been evolving host-modifying factors for billions of years. Genomes of bacterial and archaeal viruses are replete with fast-evolving, uncharacterized accessory genes (AGs), most of which likely antagonize host defenses or other viruses^1, 2^. Systematic investigation of AGs could uncover a multitude of biological mechanisms involved in virus-host competition, but AG identification in genomic databases remains a challenge. We developed an integrated computational and high-throughput discovery platform to identify AGs in virus genomes and assay their functions in complementary phage infection-dependent and -independent contexts. Our approach showcases how phages interact with the principal layers of antiviral immunity, including cell surface modifications, restriction systems, and abortive infection (Abi) mechanisms, which operate simultaneously in the same host. We discovered multiple Enterobacteriophage AGs associated with counter-defense functions that activate rather than inhibit antiviral immunity in cells, including the surprising finding that anti-restriction AGs elicit programmed cell death (PCD) activity of some restriction-modification (R-M) systems. We propose that counter-defense AGs that trigger PCD create a conundrum for phages whereby keeping the AGs causes PCD but losing them exposes the phage to restriction by bacteria. Strategies employed by viruses to avoid this double jeopardy could be an important factor in virus evolution that remains to be explored.

## Introduction

To a large extent, the evolution of life is a story of the virus-host arms race^3^. As hosts evolve elaborate antiviral defense mechanisms, viruses retort with equally versatile counter-defenses. Signatures of this arms-race are evident in prokaryotic genomes, where diverse defense systems tend to be encoded within ‘defense islands’^4^, and in virus genomes, where hotspots known as accessory regions (ARs) contain clusters of highly variable anti-defense genes^1, 2^ (Figure 1a). These accessory genes (AGs) are at the forefront of virus-host co-evolution, and many of them are known or predicted to interface with host defenses^2^. Bacteriophage AGs are typically non-essential, undergo frequent horizontal transfer, and diversify rapidly under enhanced evolutionary pressure^1^. The characterized AGs are involved in antagonizing other phages or bacterial defense systems such as restriction-modification (R-M)^5^ and CRISPR-Cas^6^, but a vast pool of AGs in rapidly growing (meta)genomic sequence databases remains unexplored^7^. Determining the identity and functions of AGs systematically remains an important and unaddressed problem.

**Figure 1.**
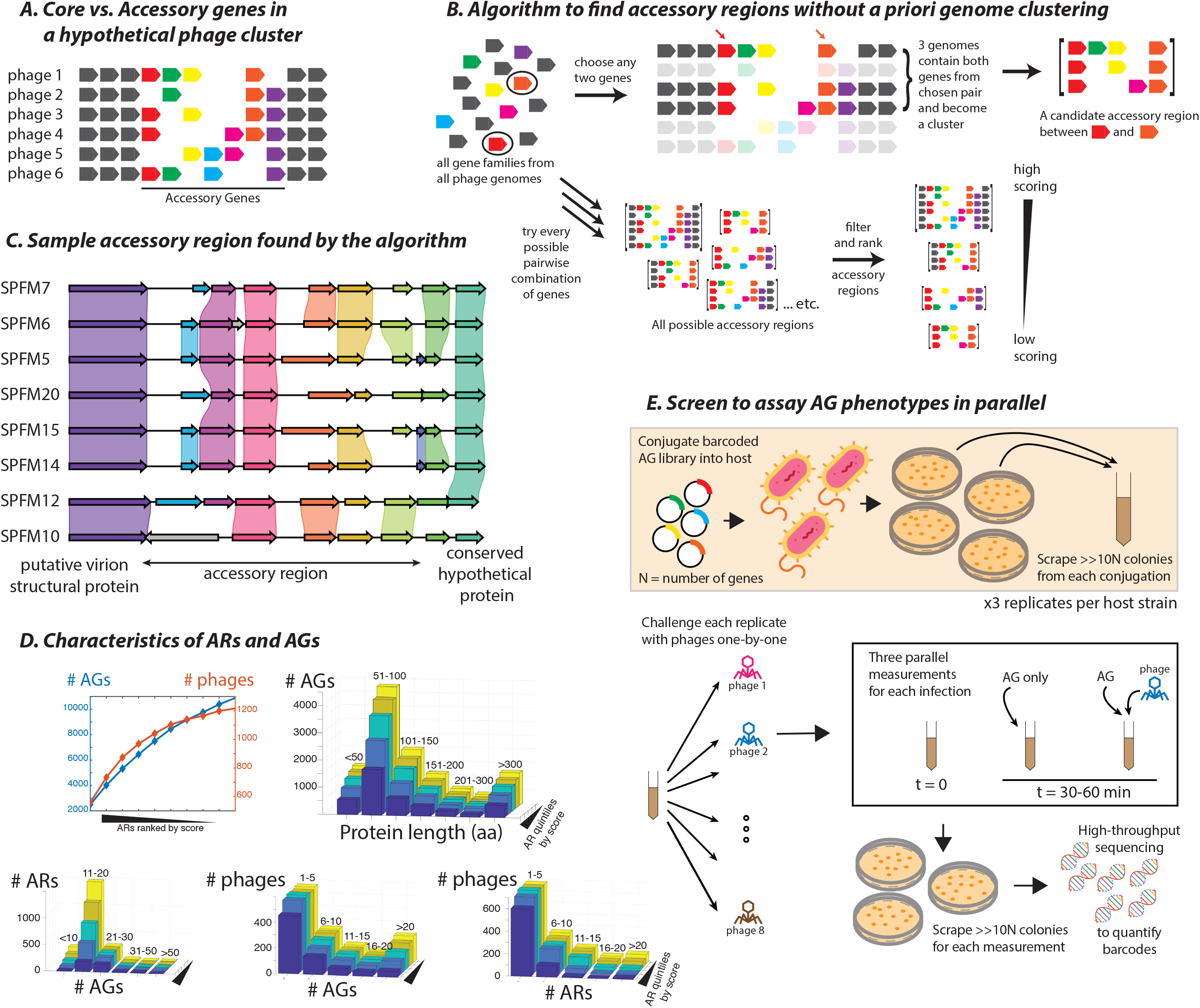
Platform for identifying and testing phage AGs. **(A)** Schematic of a genomic region showing core (shown in grey) and accessory (in various colors) genes in a hypothetical phage genome cluster. **(B)** Schematic of algorithm to exhaustively identify accessory regions using pairwise combinations of genes, without *a priori* genome clustering. **(C)** A sample high-scoring AR in a family of *Salmonella* phages. Purple shading denotes >80% and green denotes >90% nucleotide identity. **(D)** Statistical features of ARs and AGs. 2D chart depicts saturation curves (cumulative distribution functions) of unique AGs (left axis) and phage genomes (right axis) represented in ∼2000 non-redundant ARs ranked by score on the X-axis. 3D bar graphs show various distributions (length of AGs, number of AGs per AR, number of AGs per phage, and number of ARs per phage). Color-coded histograms in each bar graph are cumulative i.e., the first histogram (dark blue) shows the top 20% highest-scoring ARs, the next (light blue) represents top 40% and so on. **(E)** Schematic of the AG screen.

Comparative genomic approaches can exploit the highly organized architecture of virus genomes to identify evolutionary hotspots. However, ARs have so far only been characterized in isolated phage-families^1, 8–^<u>^1^</u>^2^, and AGs have not been studied at scale. We therefore developed a scalable, high-throughput platform to identify and functionally test AGs from large datasets of phage genomes. Here, we focus on AGs that can inhibit, and in some cases, also activate bacterial immunity. We uncover a multi-layered defense architecture and extensive co-evolution of defense and counter-defense mechanisms, including an R-M system that can trigger PCD in addition to its canonical defense functions and another R-M locus providing decoy immunity previously only known in eukaryotes^13^. On the phage side, we discover proteins that alter host metabolism and modify the cell surface, or inhibit antiviral restriction. Moreover, we identify several counter-defense AGs that trigger abortive infection (Abi) responses and propose that bacteria routinely exploit phage AGs to sense infection.

## Results and Discussion

### A high throughput screening platform for identifying and testing virus accessory genes

Given that AGs are usually defined within a cluster of closely-related viruses^8–11^, AG detection is highly sensitive to the genome clustering strategy. We therefore sought to identify ARs without *a priori* sequence comparison or clustering. All pairwise combinations of all phage genes were evaluated sequentially (Figure 1b) to determine if any pair of conserved genes bounds an AR (Supplementary File 1). We also did not rely on any gene ontology to identify AGs (eg. genes similar to or encoded next to known counter-defense factors). Starting with an initial set of 1706 Enterobacteriophage genomes, 2014 non-redundant ARs containing 10888 putative AGs were identified in 1217 phages. All ARs were then scored and ranked by diversity (See Methods). A sample AR is shown in Figure 1c. A typical AR tallied 11-30 unique AGs cumulatively across multiple related phages (we discarded ARs encountered in <6 genomes). AGs were mostly of ∼50-100 aa in length. A typical phage genome contained <5 ARs and <10 AGs, while some phages encompassed many more AGs (Figure 1d).

We manually inspected the 200 highest-scoring ARs and chose multiple (2-5) AGs from 62 of these (see Supplementary File 2 for detailed notes on each AR and rationale for AG selection). We synthesized 200 AGs, of which 54 were from ARs containing known counter-defense factors (Supplementary File 3), whereas 135 AGs were from ARs consisting entirely of genes of unknown function. Eleven phage genes with previously elucidated functions were included as controls (Abc2, ArdA, Arn, Cor, Dmd, Imm, Ip1, Ocr, Ral, SieA, Stp). To identify AG function(s), the genes were barcoded and chromosomally integrated under an IPTG-inducible promoter in a subset of the *E. coli* Reference (ECOR) strains^14^ (one AG per cell) in pools. Fitness was measured in triplicate by next-generation sequencing of AG-specific barcodes upon AG-expression (Figure 1e, Supplementary File 4).

### Phage AGs trigger programmed cell death responses in bacteria

Several AGs produced “conditional lethal” phenotypes, imposing a severe fitness burden in a minority of strains, suggestive of PCD induction (Figure 2a). Although most known triggers of abortive defense systems are conserved phage proteins^15–18^, we hypothesized that these strains harbor systems that sense variable phage proteins encoded by AGs. Surprisingly, several of these trigger AGs were known anti-restriction factors. Anti-restriction-induced cell death mechanisms might serve as a backup to thwart phage-encoded counter-defense strategies. PCD in these wild strains is triggered by some R-M inhibitors but not others, even discriminating between the two DNA mimics Ocr^19^ and ArdA^20^ (Supplementary Figure 1a). A novel AG, *orf7* (hereafter, *aop1* for Activator Of PCD) also triggers PCD in ECOR55 and 66 similarly to Ocr and ArdA, but does not inhibit Type I R-M in MG1655 (Supplementary Figure 1b). These observations suggest a diversity of PCD mechanisms.

**Figure 2.**
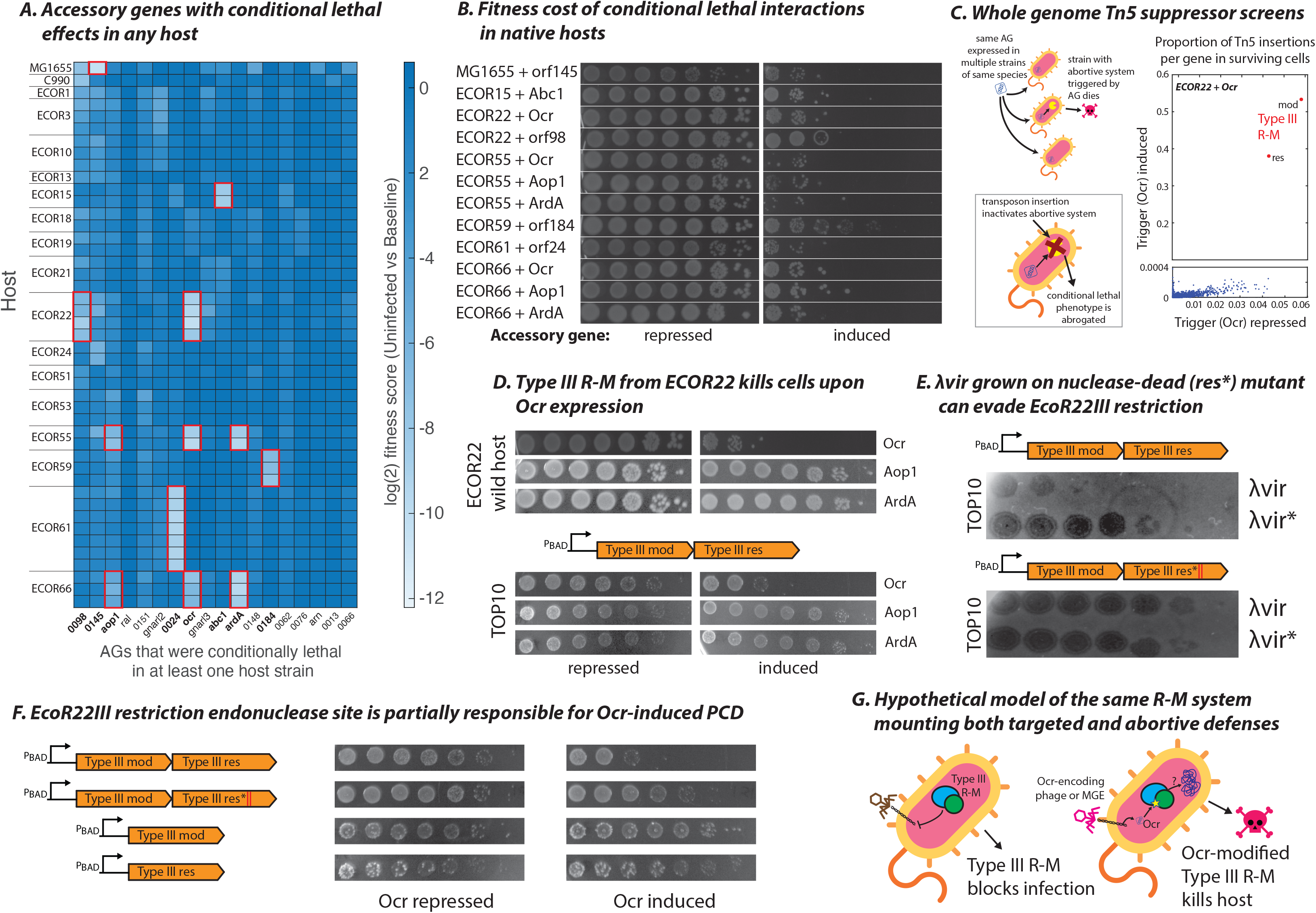
AGs that trigger programmed cell death. **(A)** Heatmap of log2-transformed fitness scores comparing induced and uninduced samples without phage infection. Of 200 AGs tested, only AGs that exhibited lethality in some (but not all) hosts are shown. Red boxes/bold names highlight conditional-lethal AGs selected for further study. Named AGs are *orf7:aop1, orf87:ral, orf63:gnarl2, orf1:ocr, orf92:gnarl3, orf116:abc1, orf169:ardA* and *orf2:arn*. **(B)** Host-AG combinations in red boxes from (C) tested individually for AG-induced lethality (serial tenfold dilutions of saturated culture; AGs repressed: left, induced: right). **(C)** Schematic depicts hypothetical strain-specific Abi system being triggered by AGs, and inactivated in follow-up transposon knockout screens (inset). Data from a representative screen with a Tn5 library constructed from ECOR22 with chromosomally encoded *ocr*. Proportion of transposon insertions per gene in surviving cells with *ocr* repressed is plotted on the X-axis, and with *ocr* induced on the Y-axis. Y-axis interrupted to resolve genes other than Type III R-M *mod*, *res*. **(D)** Abortive phenotype of EcoR22I in its native host, and reconstituted when cloned onto a plasmid and expressed in TOP10 (AGs repressed: left, induced: right)**. (E)** Plaque assay showing R-M phenotype of EcoR22I against methylated and unmethylated λvir in TOP10. λvir* were methylated by passaging in a TOP10 strain with nuclease-dead EcoR22I (res* double mutant; D1024A, D1043A) and tested for their ability to evade restriction by wildtype or res* EcoR22I. **(F)** As in (D), PCD activity of EcoR22I variants in TOP10 (wildtype, nuclease dead (res*), and *mod*, *res* genes expressed individually). **(G)** Hypothetical model of EcoR22I R-M and PCD activities. EcoR22I kills its host in the presence of Ocr, partially relying on its nuclease activity. “pBAD” indicates the construct was overexpressed from the plasmid in (F-H).

We transformed the respective wild hosts with the trigger AGs and confirmed that they induce PCD (Figure 2b). We then performed genome-wide transposon suppressor screens to map the loci causing PCD (Figure 2c). Transposon insertions in the causal system would be expected to disable PCD and allow survival of the bacteria despite heterologous expression of a trigger AG. We identified putative PCD systems in all tested hosts, except for ECOR22 with *orf98* and ECOR66 with *orf1*, *orf7*, or *orf169*, where no suppressor mutants could be obtained (Supplementary File 5).

### Restriction-modification systems can elicit PCD

Surprisingly, a Type III R-M system in ECOR22 (hereafter, EcoR22I) was required for cell-death in response to Ocr (Figure 2c), the DNA mimic that blocks both Type I and Type III R-M^21^. R-M defenses are ubiquitous in bacteria, and so are anti-restriction factors in phages^22^. In response, bacterial anti-anti-restriction mechanisms such as retrons^23^, PARIS^24^, and PrrC^25^ provide backup defense. Our results suggest that in ECOR22, the frontline (R-M) and backup (PCD) defense functions coalesced into an integrated defense system. To test this hypothesis directly, we reconstituted the PCD and R-M activities of EcoR22I in a model *E. coli* strain.

In its native host, the *mod* and *res* genes of EcoR22I are associated with an SNF2-like helicase related to the ancillary DISARM helicase, *drmD*^26^, and all 3 genes are encoded in a P4 satellite prophage hotspot^24^. However, transposon insertions that suppressed Ocr-toxicity in ECOR22 were found only in *mod* and *res* (Figure 2c, Supplementary Figure 2). We overexpressed only the *mod* and *res* genes of EcoR22I in a lab strain of *E. coli* and found that EcoR22I caused PCD upon Ocr induction but not upon induction of ArdA or Aop1, consistent with the behavior of the wild host (Figure 2d). Ocr was not toxic in the lab strain on its own (see Figure 3a). An F54V Ocr mutant, previously reported to evade detection by PARIS^24^, also lost the ability to trigger EcoR22I PCD, as did various other Ocr mutants^19^ (Supplementary Figure 1c). Because EcoR22I is the only R-M system in the lab strain and in ECOR22, EcoR22I likely detects Ocr directly.

**Figure 3.**
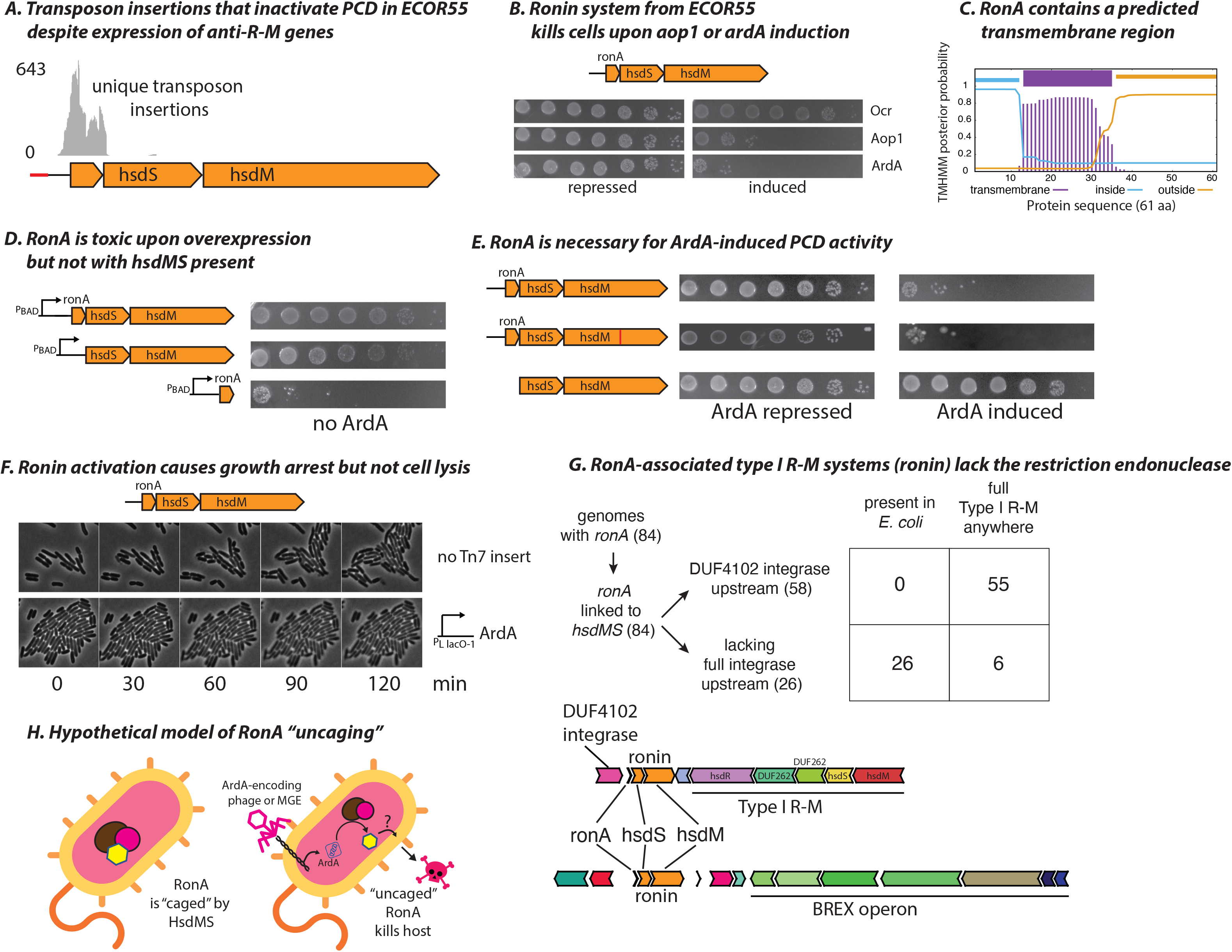
Decoy immunity in bacteria. **(A)** Distribution of unique transposon insertions recovered from survivors in ECOR55 Tn5 libraries upon *aop1* induction, mapped to Ronin. **(B)** Schematic of Ronin system from ECOR55. Abortive phenotype reconstituted in TOP10 upon expression of Ronin from a plasmid (serial tenfold dilutions of saturated culture; AGs repressed: left, induced: right). **(C)** Prediction of transmembrane character along the 61 aa RonA sequence. **(D)** Lethality of Ronin components tested when overexpressed (indicated as “pBAD”) in TOP10 without AGs. **(E)** As in (B), genetic requirements for the Abi phenotype of Ronin. Red bar in *hsdM* (second row) denotes the F287G active-site mutation. **(F)** Time-lapse imaging of TOP10 cells expressing Ronin with (bottom) or without (top) chromosomally-integrated ArdA. **(G)** Two prevalent architectures of *ronA*-linked Type I R-M-like Abi system (ronin), based on the presence of an upstream DUF4102 integrase. Number of (*E. coli* and other) genomes with a full Type I R-M system (with *hsdR, hsdM, hsdS* genes) anywhere in the genome is shown for each Ronin architecture. **(H)** Schematic depicting hypothetical RonA “uncaging” when ArdA is present.

Next, we tested whether the restriction activity of EcoR22I could block phage infection. We constructed an active-site mutant of the PD-(D/E)xK restriction nuclease (res*: D1024A, D1043A double mutant) and compared immunity conferred by wild type and res* EcoR22I against a panel of phages. EcoR22I but not the catalytic site mutant blocked λvir (Figure 2e). Furthermore, λvir passaged through the res* host (and thus modified by the EcoR22I methyltransferase) regained the ability to infect cells carrying wildtype EcoR22I, indicating that EcoR22I indeed restricts only unmodified λvir DNA.

We then tested if the immune (R-M) and PCD functions of EcoR22I could be uncoupled. EcoR22I *mod* and *res* individually did not induce PCD when co-expressed with Ocr, indicating that PCD induction is a property of the complex (Figure 2f). The res* nuclease-dead mutation of EcoR22I that completely abrogated λvir restriction also decreased PCD by 100-fold, although it did not fully abolish Ocr-induced toxicity (Figure 2f). These results show that the EcoR22I PD-(D/E)xK nuclease motif is involved in both R-M and PCD but does not appear to be fully responsible for the latter.

Thus, EcoR22I mounts both a targeted antiviral defense and a PCD response in the presence of certain phage AG products (Figure 2g). Coupling restriction and PCD could allow cells to respond dynamically depending on the progress of the infection^27^.

### Decoy immunity in bacteria

A second R-M-like system also elicited PCD in the presence of anti-R-M AGs. This locus in ECOR55 contains *hsdS* and *hsdM* genes typical of Type I R-M but lacks the nuclease (*hsdR*), and instead has a tightly linked gene that encodes a 61aa protein. This small gene was identified in our suppressor screens as the PCD-inducing effector activated by Ocr, Aop1, and ArdA (Figure 3a). We named this system Ronin, after the master-less samurai in the 1962 Japanese film, *Harakiri*. Ronin induces cell suicide in ECOR55 when Ocr, ArdA, or Aop1 are expressed (Figure 2b), but only responds to ArdA and Aop1 when cloned into a lab strain (Figure 3b).

Several lines of evidence suggest that the 61aa protein (hereafter, RonA) in Ronin is the PCD effector. The presence of a predicted transmembrane helix suggests RonA might disrupt membrane integrity (Figure 3c). Over-expression of *ronA* alone is lethal, but this effect is suppressed by HsdMS co-expression (Figure 3d). Ronin activity is independent of the methyltransferase function of HsdMS because PCD was not abrogated by an F287G mutation in motif IV (NPPF) of the *hsdM* catalytic domain^28^ (Figure 3e). However, removal of *ronA* inactivates Ronin, such that anti-restriction proteins no longer trigger PCD (Figure 3e). The 235-bp non-coding region upstream of *ronA* also contributes an unknown, essential functionality (Supplementary Figure 3). Live cell imaging showed that Ronin activation arrested growth without lysis, consistent with RonA-mediated inner-membrane disruption as a PCD mechanism (Figure 3f). As *ronA* is not lethal when co-expressed with HsdM and HsdS (hereafter, HsdMS), we surmise that its activity is sequestered or suppressed by HsdMS. This model is reminiscent of PrrC activation upon blocking of EcoPrrI restriction by T4-encoded peptide Stp. Similarly to Stp binding EcoPrrI and releasing the Abi anticodon nuclease PrrC^25^, an anti-restriction protein could bind Ronin HsdMS to release RonA, the PCD effector.

Examination of the gene neighborhoods of *ronA* homologs showed that they are always linked to *hsdS* and *hsdM* genes and occur in two predominant genetic architectures. Ronin is either linked to an atypical Type I/Type IV composite R-M system or to a BREX system (Figure 3g). The BREX-linked architecture is exclusive to *E. coli*. Most (77%) Ronin-encoding *E. coli* genomes contain no Type I R-M systems anywhere in the genome, and therefore lack *hsdR* genes (Supplementary File 6). The predominantly nuclease-lacking configuration in *E. coli* suggests that PCD is Ronin’s primary function. This is reminiscent of decoy immunity (especially common in plants), where a host factor mimics the target of a pathogen effector. The effector then binds the decoy instead of its actual target, and the decoy alerts the host to the presence of the pathogen^13^.

Thus, Ronin appears to be a Type I R-M derivative that lost the defense nuclease but gained the ability to sense anti-restriction effectors and trigger PCD. Ronin encodes the canonical target of Ocr/ArdA, the Type I R-M HsdMS complex, and its activation likely depends on these anti-restriction proteins binding HsdMS (Figure 3h). Consistent with this, Ronin could still sense the Ocr F54V mutant that evades PARIS and EcoR22I but retains anti-restriction activity^24^. Surprisingly, however, Ronin also responded to various Ocr mutants that lost their anti-restriction activity (Supplementary Figure 1c). Because it almost always appears linked to an R-M or BREX system, both of which can be blocked by Ocr^29^, Ronin might drive evolution of Ocr-encoding phages to a BREX/R-M susceptible state through the loss of Ocr. Decoy immunity could thus be a powerful host strategy to constrain virus escape from restriction systems in bacteria.

### Counter-defense-associated AGs trigger Abi mechanisms encoded in prophages

We identified three prophage-encoded Abi loci that induced PCD in response to counter-defense-associated AGs (Supplementary File 5). First, we found that heterologous *orf116* expression triggered PCD in ECOR15. Transposon-insertions identified the P4 satellite prophage (especially the ∼100-bp intergenic region marked by a star in Figure 4a) as the Abi locus. *orf116* is the poorly characterized *abc1* gene of phage P22 that is adjacent to the RecBCD inhibitor *abc2* (Figure 4a)^30^. We cloned the P4 satellite prophage into a plasmid and reconstituted its PCD effect in a lab strain. P4 cannot be induced without a co-resident P2 prophage (which was absent in the lab strain), ruling out prophage-induction as the cause of PCD. Abc1 contains a helix-turn-helix (HTH) domain typical of DNA-binding proteins (Supplementary File 7). Thus, Abc1 might trigger PCD by interfering with P4 transcriptional regulation, perhaps upregulating the lethal gene *kil*.

**Figure 4.**
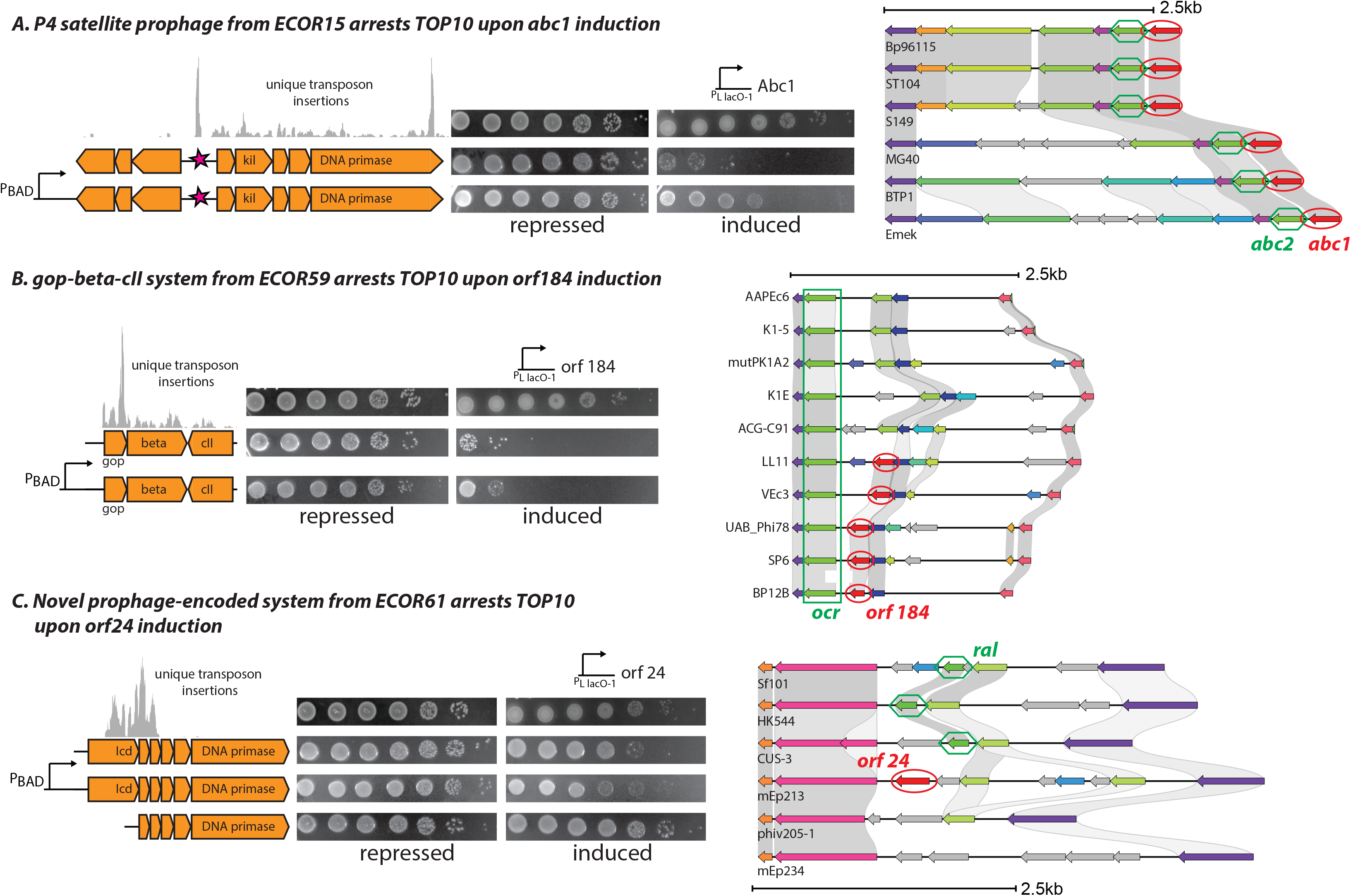
AGs that trigger prophage-encoded PCD mechanisms. **(A)** Abortive phenotype of P4 satellite prophage from ECOR15 cloned onto a plasmid and expressed in TOP10. Abc1 is expressed from a single-copy chromosomal insertion (serial tenfold dilutions of saturated culture; AGs repressed: left, induced: right). Star marks an intergenic region that was strongly targeted in transposon suppressor screens. **(B)** As in (A), PCD phenotype of gop-beta-cII system from ECOR59 upon expression of *orf184*. **(C)** As in (A), PCD phenotype of novel prophage-encoded superinfection exclusion system from ECOR61 upon expression of *orf24*. “pBAD” indicates where the constructs were overexpressed in (A-C). Distribution of unique transposon insertions recovered from survivors in Tn5 libraries upon trigger AG induction are mapped to each PCD locus in (A-C). Accessory Regions where trigger AGs in (A-C) were found (*orf116*:*abc1*, *orf184*, *orf24*; highlighted with red ovals), with co-occurring counter-defense AGs (anti-RecBCD *abc2*, anti-R-M *ocr*, anti-R-M *ral*; indicated in green boxes) are shown in the respective panels.

Second, the gop-beta-cII Abi system is a toxin-antitoxin-like locus located in the P4 hotspot^24, 31^ with apparent anti-phage activity^24^. Our suppressor screens showed that gop-beta-cII toxicity is activated by *orf184*, which is widely distributed among coliphages and is present in the same AR with the anti-restriction gene *ral* (Figure 4b). We cloned gop-beta-cII into a lab strain and confirmed that *orf184* triggered PCD (Figure 4b).

Finally, a distinct prophage-encoded system in ECOR61 inhibited growth upon expression of *orf24*, which is narrowly distributed and present in the same AR with the anti-restriction gene *ocr* (Figure 4c). We found suppressor transposon-insertions in three prophage genes, including an Icd-like cell-division-inhibitor with a long N-terminal extension (Superfamily cl41269). However, expression of all 6 linked genes (including a DNA primase) was necessary to reconstitute PCD in a lab strain, still yielding a muted effect (Figure 4c) compared to that in the native host (Figure 2b). The Icd-like protein is the likely effector because PCD induction was abrogated upon its removal (Figure 4c). These findings demonstrate that prophages can activate superinfection exclusion mechanisms in response to highly variable phage AGs.

### Phage AGs disable host defenses

In the previous sections, we show that phage AGs can activate immune pathways that lead to cell death or dormancy. Next, we wondered how AGs might affect the progress of phage infections in ECOR strains. We layered phage infection onto our AG screens, measuring fitness in the presence or absence of infection by 8 model phages (Figure 1e). Forty-five phage-host combinations were selected where phage replicative titers were highly attenuated compared to growth on a sensitive lab strain (Supplementary Figure 4a). This phenotype suggests that an antiviral mechanism was blocking phage replication, such that expressing an AG that inhibits this defense system could alleviate restriction. Although 8 model phages were used in our infection-based screens, the AGs were taken from hundreds of diverse phage genomes (see Figure 1).

To identify AGs with counter-defense phenotypes, we compared infected samples with uninfected controls and identified several host-phage-AG combinations that were depleted from the infected pools (Figure 5a). Each AG that sensitized any wild strain to infection was then individually validated in that strain against the entire phage set. In these validation experiments, some additional phage-host combinations that were omitted in the initial screen also displayed AG counter-defense phenotypes (Supplementary Figure 4b). In all, increased plaquing of several phages was observed upon expression of 3 known Type I R-M inhibitors (Ocr, Ral, ArdA), 6 novel AGs, and the T4 internal protein 2 (IpII) in a total of 28 phage-host combinations. These AGs likely antagonize host factors that inhibit phage reproduction.

**Figure 5.**
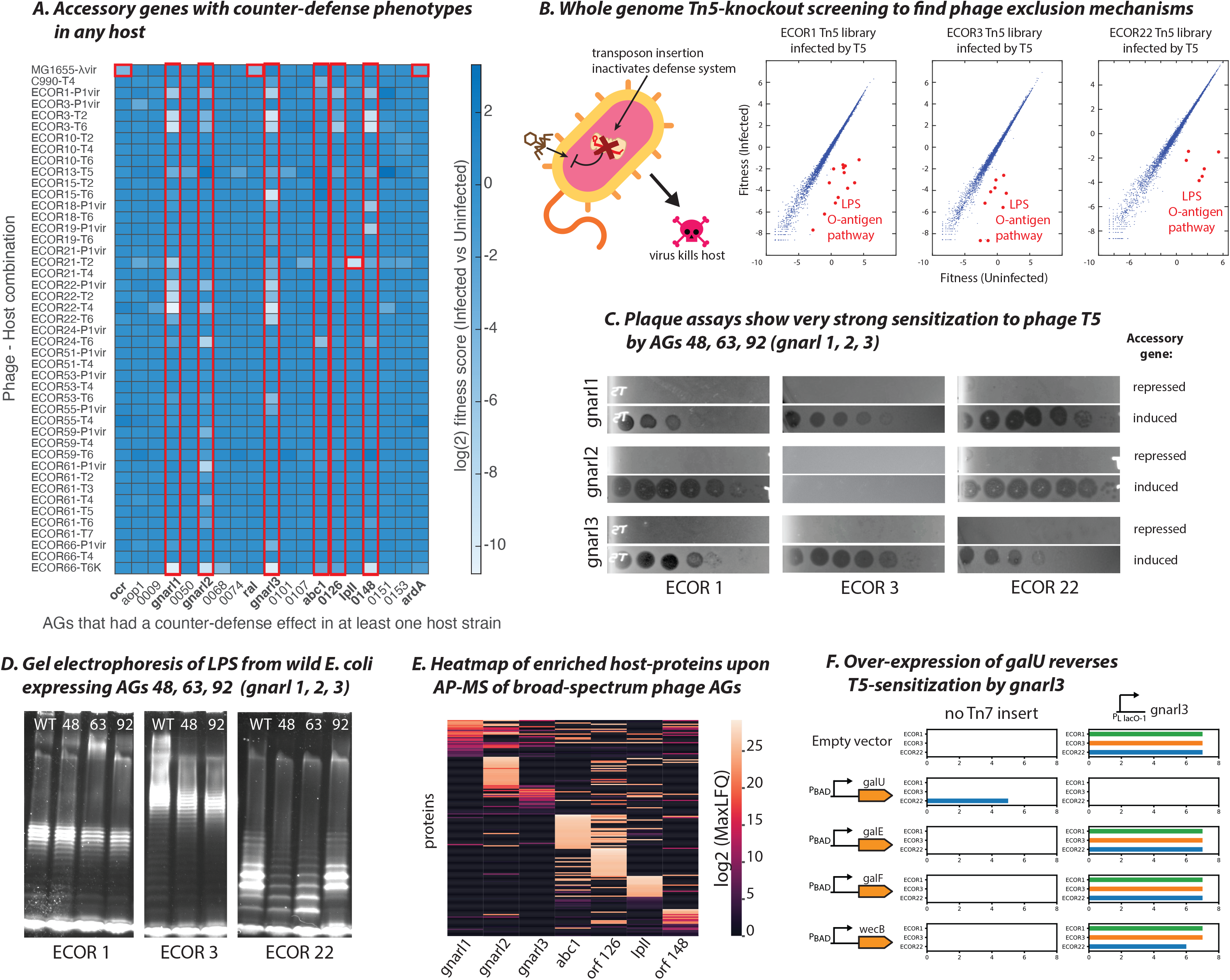
AGs that sensitize hosts to infection by modifying barrier defenses. **(A)** Heatmap of log2-transformed fitness scores comparing infected and uninfected samples. Of 200 AGs tested, only AGs with a host-sensitizing phenotype are shown on the X-axis. Red boxes/bold names highlight counter-defense AGs selected for further study. Named AGs are *orf1:ocr, orf7:aop1, orf48:gnarl1, orf63:gnarl2, orf87:ral, orf92:gnarl3, orf116:abc1, orf143:IpII* and *orf169:ardA*. **(B)** Schematic of follow-up transposon knockout screens, performed in the native host without AGs. Graphs show fitness of transposon-mediated knockouts in infected (Y-axis) and uninfected (X-axis) Tn5-libraries from three representative hosts challenged by T5. Red dots are gene-disruptions that lower bacterial fitness upon infection (e.g., disruptions in O-antigen biosynthesis). **(C)** Plaque assays with phage T5 and hosts from (D) with *gnarl* genes repressed (top) or induced (bottom). **(D)** Lipopolysaccharide (LPS) visualized by electrophoresis from host-AG combinations in (E) where AGs produced a phage-sensitizing effect. **(E)** Log2-transformed protein representation scores (MaxLFQ) for the top 30 enriched host proteins upon expression of AGs. **(F)** log10-transformed T5 infection scores in wild hosts ECOR1 (green), ECOR3 (orange), ECOR22 (blue) with or without UDP-glucose biosynthesis pathway genes *galU, galE, galF,* and ECA precursor *wecB* cloned onto a plasmid and overexpressed.

To identify antiviral mechanisms inhibited by AGs, we employed two unbiased approaches: (1) transposon-knockout screens to pinpoint gene disruptions in hosts that phenocopy AG expression (i.e. increased phage infection) (Supplementary File 8) and (2) affinity-purification and mass spectrometry (AP-MS) of AGs to identify binding partners in the wild strains (Supplementary File 9). We performed these assays for every host found to be sensitized to infection by any AG.

### AGs antagonize conserved host pathways and barrier defenses

Six novel AGs – *orf48*, *orf63*, *orf92*, *orf116* (*abc1*, see above), *orf126*, and *orf148* – all exhibited counter-defense phenotypes in multiple host strains. Because the broad-spectrum effects of these AGs did not correlate with the variable defense repertoires of these strains (Supplementary Figure 4c), we sought to determine whether they targeted conserved host antiviral pathways.

Genome-wide knockout screens indicated that disrupting the host O-antigen or capsule genes phenocopied the phage-sensitization effects of all 6 broad-spectrum counter-defense AGs with almost all tested phages (Supplementary File 8). Some of the most pronounced phenotypes we observed were that transposon-mediated O-antigen disruption (Figure 5b) or expression of *orf48*, *orf63*, or *orf92* (Figure 5c) both converged on the same effect, namely, enhanced infection by phage T5 (which does not natively encode these AGs). We hypothesized that T5 adsorption was blocked by the wild-type O-antigen structures in these hosts, but heterologous AG expression enabled infection by modifying or removing the O-antigen. We therefore examined LPS preparations from every host where any AG exhibited a counter-defense phenotype. Hosts expressing *orf48*, *orf63*, or *orf92* exhibited subtle downshifts in the O-antigen banding pattern (Figure 5d), as observed previously in some seroconverting lysogens as a means to exclude superinfecting phages^32^. Specifically, the following host-AG combinations showed visible downshifts in O-antigen bands: *orf63* and *orf92* in ECOR1, *orf48* and *orf92* in ECOR3, and *orf48* and *orf63* in ECOR22. These downshifts correlate with the host-AG pairs where T5 infection was especially strongly (10^5–6^-fold) enhanced (Figure 5c). We reasoned that the AGs might interfere with O-antigen biosynthesis, and tentatively named *orf48*, *orf63* and *orf92*, *gnarl1, 2,* and *3* respectively, after the skin-eating demon in the popular TV series *Buffy the Vampire Slayer*.

All known seroconverting prophages modify O-antigens using enzymes such as glycosyltransferases and acetyltransferases)^32–36^. *Gnarl-*encoding phages in our genomic dataset do not encode any such seroconverting enzymes. Gnarl proteins (40aa – 90aa) are also much smaller than typical enzymes, hinting that these AGs might inhibit O-antigen-modifying enzymes, constituting a distinct mode of host seroconversion. *Gnarl* genes are not restricted to either lytic or lysogenic phages; *gnarl1* and *gnarl3* are present in lytic phages (Supplementary File 10), whereas *gnarl2* is present in the classical temperate phage Mu (gene E6/Mup07). Seroconversion could occur in lysogenic infections to exclude superinfecting phages, or in lytic infections as receptor-masking to prevent newly synthesized virions from unproductively binding lysed cell fragments^37^.

To identify the cellular targets of gnarl proteins, we analyzed AP-MS data for all novel counter-defense AGs. Each accessory gene product bound distinct host proteins, although the profiles of *orf116* (*abc1*) and *orf126* were somewhat correlated (Figure 5e). Gene Ontology pathway analysis identified UDP-glucose biosynthesis factors as significantly enriched in the *gnarl3* set, suggesting that *gnarl3* inhibits a step in this pathway. Among these, GalU catalyzes the formation of UDP-glucose^38^ to build the LPS outer core^39^; GalF regulates the levels of UDP-glucose^40^; and GalE reversibly converts UDP-glucose to UDP-galactose (also an LPS outer core component)^41^. Overexpression of *galU,* but not *galE*, *galF*, or *wecB,* completely reversed the *gnarl3* T5-sensitization phenotype in ECOR1, ECOR3, and ECOR22 (Figure 5f). suggesting a specific interaction between *galU* and *gnarl3*. The same effect was observed with the unrelated phage T4 (Supplementary Figure 5), indicating the *gnarl3-galU* interaction was phage-agnostic. Overexpressing *galU* without *gnarl3* did not affect phage sensitivity except in ECOR22, where it partially potentiated infection for unknown reasons. Coupled with LPS electrophoresis and AP-MS data, this observation suggests a model where *gnarl3* inhibits GalU, reducing the availability of UDP-glucose and resulting in a modification of the cell envelope that potentiates phage adsorption. More broadly, it is notable that the most prevalent and potent “counter-defense” phenotype for multiple, distinct AGs appears to hinge on removing barrier defenses through diverse mechanisms.

### Barrier defenses and restriction systems provide layered immunity

The existence of broad-spectrum counter-defense AGs underscores the importance of barrier defenses. In strain ECOR21, transposon insertions within the O121-specific O-antigen pathway^42^ indeed sensitized the strain to multiple phages, but not T2 (Figure 6a). We wondered if an additional defense mechanism in ECOR21 complemented barrier defenses by restricting T2 infection (Figure 6b). In the AG screen, internal-protein IpII (*orf143*) partially sensitized ECOR21 to T2 (Figure 5a), suggesting that T2 is blocked by an IpII-inhibited system in addition to O121 O-antigen. Interestingly, although phage T2 itself does not encode IpII, phage T4 encodes IpII as well as the only internal-protein from T-even phages with a known function, IpI. IpI is packaged into the phage capsid and disables canonical Type IV R-M (consisting of two enzymes, GmrS and GmrD) upon co-injection with phage DNA^43^.

**Figure 6.**
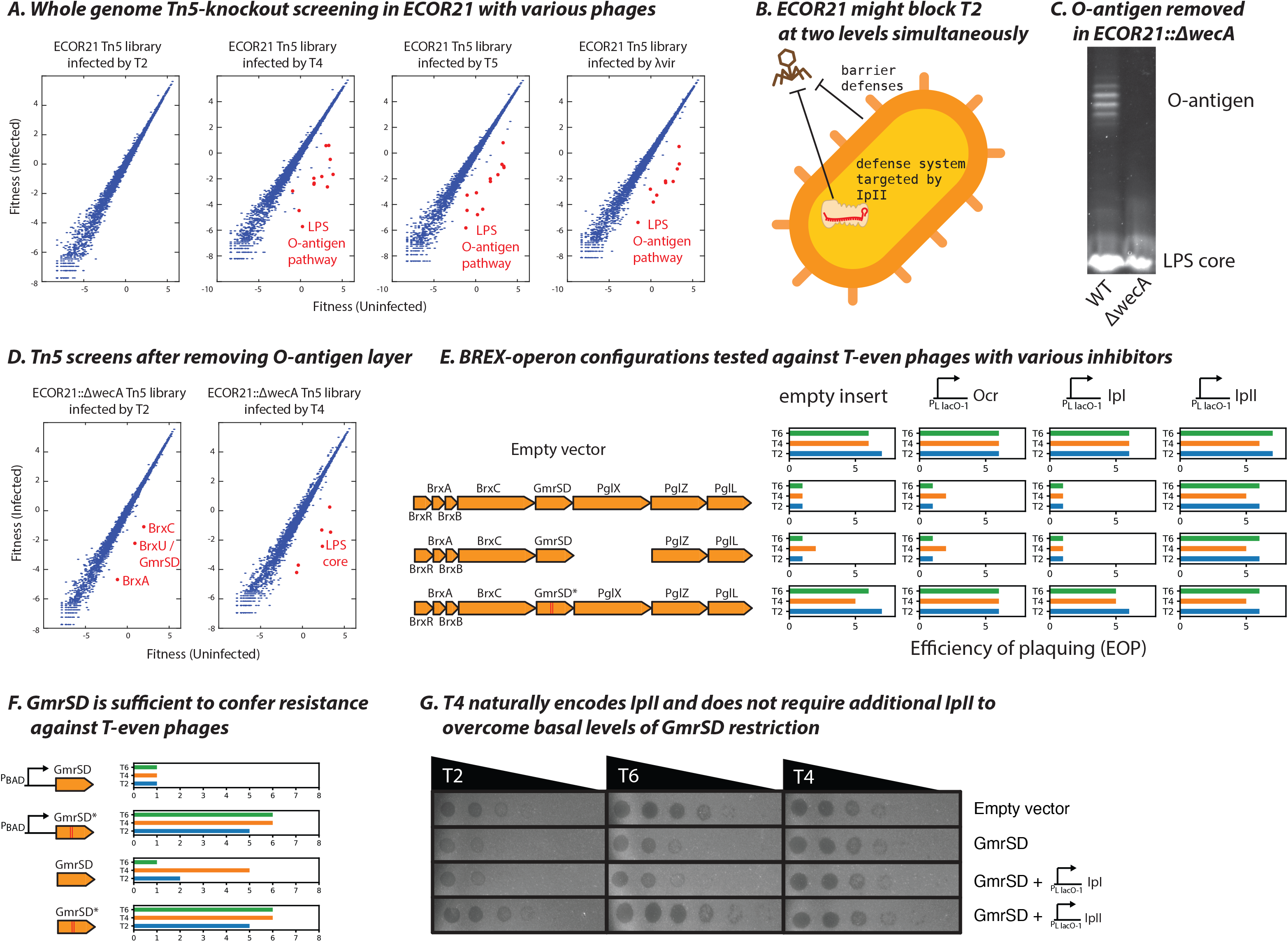
AGs that block canonical antiviral defenses. **(A)** Fitness of transposon-mediated knockouts in infected and uninfected Tn5-libraries from ECOR21 challenged by various phages. **(B)** Schematic of ECOR21 defense against T2 through both barrier defenses and a defense system which is putatively blocked by *orf143*/IpII. **(C)** LPS visualized from ECOR21 with the O-antigen biosynthesis initiator *wecA* deleted by allelic exchange. **(D)** Whole-genome transposon-knockout screens as in (A) but with the ECOR21::Δ*wecA*. **(E)** Variants of the BREX/GmrSD operon cloned onto a plasmid along with their native promoters and tested in a lab strain of *E. coli* (TOP10) against T-even phages (T2, T4, T6). Removal of the PglX gene inactivates BREX. Active-site mutations that disable GmrSD are shown as two parallel red bars (GmrSD* double mutant; D474A, H475A). Ocr, IpI and IpII are expressed from single-copy chromosomal insertions. Bar graphs show log10 infection scores. **(F)** GmrSD and its inactive variant GmrSD* cloned onto a plasmid and tested in TOP10 for restriction activity against T-even phages. “pBAD” indicates the defense construct was overexpressed from the plasmid. **(G)** Plaque assays with TOP10 cells with uninduced GmrSD (i.e., not overexpressed) challenged by T-even phages.

To identify the IpII-inhibited system, we eliminated ECOR21 O-antigen by deleting *wecA*^44^ (Figure 6c) and repeated the transposon screen, looking for mutants in the ECOR21Δ*wecA* background that now became sensitive to T2. We obtained several transposon insertions in an atypical BREX system (Figure 6d) in which a fused Type IV R-M enzyme GmrSD (also known as BrxU) is embedded^45^. Type IV R-M systems target glucosyl-hmC modified DNA of T-even phages (T2, T4, T6)^43^. The entire ECOR21 BREX-GmrSD system cloned into a lab strain and expressed from its native promoter blocked T2, but an active-site mutant in GmrSD (GmrSD*) did not (Figure 6e). Inactivation of the BREX system (Δ*pglX*) had no impact on T2, confirming that GmrSD causes T2 inhibition and is the likely target of IpII. Consistent with this conclusion, expression of IpII, but neither Ocr (anti-Type I R-M and anti-BREX^29^) nor IpI (which inhibits canonical GmrS-GmrD two component Type IV R-M, but not the fused GmrSD^37, 43^), abolished GmrSD defense (Figure 6e).

To test sufficiency, we expressed GmrSD or the catalytic mutant GmrSD* alone. Overexpressing GmrSD was sufficient to block all T-even phages, while the catalytic mutant had no effect (Figure 6f). Leaky expression of GmrSD from the uninduced plasmid still inhibited replication of T2 and T6 (which do not encode IpII), but not the IpII-encoding phage T4. When IpII was additionally expressed, T2 and T6 plaquing was restored, whereas plaquing of T4 did not improve further (Figure 6g). Of the many described T-even phage internal proteins^12^, IpII is only the second (after IpI) with an identified target. The observed redundancy between barrier defenses and restriction systems likely obscures phenotypes of other anti-restriction AGs as well.

## Conclusion

Our strategy for identifying AGs and their interactions with host defenses can tease apart the layers of bacterial anti-phage immunity, providing a holistic view of the most salient defenses across multiple strains. Deploying this platform in pathogenic bacteria will help tailor phage therapy approaches to overcome species-and strain-specific antiviral mechanisms. In *E. coli*, we discovered a diverse array of AG functions that interact with multilayered host immunity at every level. AG-encoded proteins modify the cell surface to block viral entry, disable antiviral restriction systems inside the cell, and trigger PCD mechanisms in the host and its prophages, while cellular decoy immunity guards the host against phage counter-defense strategies. We show here how restriction systems work in tandem with barrier defenses on the cell surface, and with Abi/decoy guardians within the cell to protect the host.

We report surprising exaptation of well-studied R-M systems that are triggered by anti-restriction proteins to induce PCD. Our results further expand the conceptual framework of layered anti-phage defense in bacteria, whereby anti-anti-restriction mechanisms such as retrons^23^, PARIS^24^, and PrrC^25^ provide backup immunity via Abi. In particular, EcoR22I combines both R-M and PCD functions, and Ronin is a PCD mechanism that mimics an R-M system to thwart phage anti-R-M AGs under the decoy immunity strategy previously identified only in eukaryotes. Thus, experimentally assaying AGs in wild bacterial strains can uncover defense functions that cannot be identified by comparison with known defense or counter-defense factors. For instance, EcoR22I does not have any domains currently known to be involved in PCD. Similarly, *ronA*-like toxins have no previously characterized links to R-M.

Assaying AGs individually in diverse hosts also directly identifies otherwise elusive triggers of defense systems. Anti-phage defense systems have been shown to detect highly conserved features of phages (nucleic acids, capsids, terminases, or portal proteins)^15–18^. We show that a variety of poorly conserved phage AGs can also trigger defense mechanisms in the host and its prophages. Non-essential gene products might seem ill-suited for this role because the virus could simply lose the gene to escape detection. However, all AGs we found to trigger PCD are associated with common counter-defense functions, trapping the virus into a Scylla vs Charybdis catch where a counter-defense AG triggers PCD upon infection, but losing that AG renders the virus vulnerable to immunity. The strategies viruses employ to navigate this treacherous narrows are currently unknown and might be an important force in virus evolution. The host counterpoints to virus AGs were likely overlooked because many viruses encoding inhibitors and triggers of specialized host defenses can be difficult to procure and culture. Advanced technologies for DNA synthesis combined with rapidly growing metagenomic sequence databases are starting to paint a more complete picture of virus-host coevolution.

## Methods

### Algorithm to identify phage accessory regions

1706 non-redundant Enterobacteriophage genomes were obtained (with help from Dr. Andrew Millard, Leicester) from public databases. These phages were both lytic and lysogenic, but were sequenced and assembled from cultured isolates (i.e. prophages and metagenomic samples were not included). All annotated protein records from all the phages were assigned unique numeric IDs and clustered by MMseqs2^46^. ORF IDs in phage genomes were then replaced by their MMseqs2 cluster representative IDs in each genome. Thus, each phage genome can be represented as an ordered list of IDs of clustered proteins. The gene-neighborhood of every ORF (up to 10 genes upstream and downstream) was extracted from every virus. Given that phage genomic termini can be difficult to identify, every genome was considered potentially circularly permuted. Allowing for circularization also ensures all gene-neighborhoods were the same size and putative ARs towards ends of genomes would not be missed. A 2D matrix of all possible pairwise combinations of genes was constructed, with every matrix position containing lists of (a) the phages that each gene appears in, (b) the 10 upstream, and (c) 10 downstream neighboring ORFs of each gene in every genome. We then traversed the matrix and assessed every gene pair to identify all possible ARs, defined as the region between a pair of genes that are both present in at least 6 different phage genomes but are no more than 10 genes apart in any genome. Hence, every virus genome that contained both genes of the gene pair was grouped into a possible virus cluster, and genes encoded between the genes of the query pair were considered members of a possible AR. Regions trivially containing the same genes in each participating genome were discarded. Genomes involved in a putative AR were then compared pairwise at the nucleotide level using FastANI^47^, and ARs with divergent genomes were removed. In practice, this was done by pre-computing an all-vs-all matrix of genome ANI and then traversing subsets of the matrix containing all the genomes participating in the putative AR to ensure no cells were empty (FastANI produces no output when very divergent genomes are compared). Various parameters were extracted from surviving ARs. *N* is the number of unique genes in the AR. The vector *c* contains conservation numbers (the number of different genomes that each gene appears in; also indicated on top of each gene) for all the unique genes in this AR. The vector *l* contains the lengths (in numbers of genes) of the AR in each genome. *CV* denotes coefficient of variation. *GCdev* is a vector containing one minus the absolute deviation in GC content of each accessory gene from its resident genome. A scaling factor is also included to normalize the score by the number of phage genomes in the group that have at least one gene in the AR. ARs were scored according to the formula below.

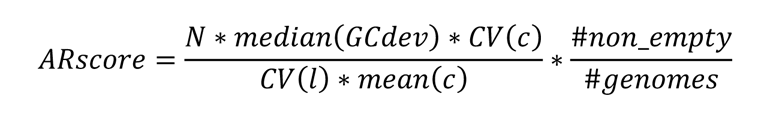

To filter out ARs that were subsets of other ARs, we successively removed redundant, low scoring ARs with at least 90% of genes that had already been encountered in higher scoring ARs. The top 200 ARs were examined manually and 1-15 AGs were selected from 62 ARs (Supplementary Files 2,3). A worked example of the code is provided in Supplementary File 1.

### Strains and Vectors

The *E. Coli* Reference (ECOR) collection of wild *E. coli* strains was obtained from the STEC Center (Michigan State University). Lab strains were obtained as follows: MG1655 (from Dr. Carol Gross, UCSF), C990 (from Dr. Ry Young, Texas A&M), DH5α (NEB), DH10ß/TOP10 (Life Technologies), BW25113 and BW25141 (from Dr. Vivek Mutalik, LBNL), and WM6026 (from Dr. Jason Peters, UW-Madison). All strains were cultured at 37°C with shaking (180-225 rpm) in lysogeny broth (LB) (10 g/l tryptone, 5 g/l yeast extract, 10 g/l NaCl with 15 g/l agar for plates) supplemented with 100 µg/ml of ampicillin/carbenicillin, 10-20 µg/ml gentamicin (except WM6026, which yielded more consistent colony sizes with 50 µg/ml), 50 µg/ml kanamycin, and 0.5% w/v glucose, 0.1% w/v L-arabinose, and 1 mM IPTG (isopropyl-b-D-thiogalactopyranoside) as needed. The WM6026 strain required diaminopimelic acid (DAP) for growth, which was supplied at 300 µM. All strains were stored in 20% glycerol at −80°C for long term storage.

R6K-origin pTn7 plasmids (from Dr. Jason Peters, UW-Madison) – the pJMP1039/pTn7C1 helper plasmid (carbenicillin) and the pJMP1360/pTn7C185 transposon vector (gentamicin, carbenicillin) – were used for integration of phage AGs into the *E. coli* genome. All host defense systems identified in the various screens were placed under the araBAD promoter (arabinose inducible) in the pBAD Myc/His A vector (Invitrogen). Conjugative allelic exchange was carried out using R6K-origin pKEK2201 (kanamycin) (from Karl E. Klose, UT San Antonio). All plasmids were constructed by Gibson Assembly.

R6K-origin plasmids were maintained in BW25141 (routine cloning) and WM6026 (conjugative donor strain). All other plasmids were maintained in DH5α or TOP10. All plasmid sequences were verified by Sanger or whole-plasmid sequencing (Plasmidsaurus, Primordium) and are available upon request. Frequent plasmid dimerization during the Gibson Assembly procedure (especially when using the NEBuilder reagent) was ameliorated by using 10-fold less of one of the insert DNA fragments in a reaction involving at least 2 insert fragments (not including the backbone).

### AG conjugation into host strains

AGs were delivered to *E. coli* hosts by Tn7 transposition via conjugation performed by triparental mating as described previously^48^. Briefly, the transposon and helper plasmids were delivered to the recipient from WM6026 donor strains by overnight incubation on LB agar plates supplemented with 300 µM DAP and 0.5% glucose to repress AG expression. The next day, cells were scraped and resuspended in 1 ml LB, and 1000X-20000X dilutions of this conjugation mixture were plated on LB plates (lacking DAP) supplemented with the appropriate antibiotics and 0.5% glucose. All strains were maintained under strict glucose repression of the pLlacO-1 promoter at every step until screens were performed. Thus, all AGs were maintained as single-copy chromosomal inserts at the same attTn7 site in the genomic region between *glmS* and *pstS* genes.

Efficient Tn7 transposition is necessary to construct an AG library in any given *E. coli* host. Tn7 conjugation-transposition efficiencies were determined in 44 ECOR stains containing an intact *glmS-pstS* region (we note that other strains can support Tn7 integration as well). Tn7 transposons carried only a barcode sequence and the gentamicin-resistance gene to minimize payload-dependent variation in conjugation efficiency between strains. Efficiency of conjugation was determined by streaking the conjugation mixture on two different gentamicin concentrations (25 µg/ml and 10 µg/ml) to account for possible differences in natural antibiotic resistance between the wild strains. The barcode sequence was verified by Sanger sequencing and both flanks were checked by PCR to ensure proper integration of the Tn7 transposon at the expected attTn7 site. Strains that supported Tn7 transposition at high levels and were amenable to agar-overlay plaque assays were selected for further experiments.

### DNA extraction + Nextera Genomic Sequencing + Analysis

Genomic DNA was extracted using a modified SDS–proteinase K method: Briefly, cells pelleted from 50 µl of saturated culture (or 5 µl of the cell mixture scraped from selective plates during AG library construction) were re-suspended in 200 µl of lysis buffer (10 mM tris, 10 mM EDTA, 400 mg/ml proteinase K, 0.5% SDS) and incubated at 50-55°C for 1 hour. Subsequently, the temperature was lowered to 37°C and RNase A (Thermo Scientific) was added to a concentration of 1 mg/ml. After 30-60 min of incubation, the digests subsequently purified using the Genomic DNA Clean & Concentrator Kit (Zymo Research). DNA was prepared for whole-genome sequencing using the Nextera Flex Library Prep Kit (now, the Illumina DNA Prep Kit) according to manufacturer’s instructions, except that we lowered the volumes of all tagmentation reactions 5-fold, and used a custom dual-indexed primer set JSW-SS-22:33 (CAAGCAGAAGACGGCATACGAGAT NNNNNNNN GTCTCGTGGGCTCGG) and JSW-SS-34:41 (AATGATACGGCGACCACCGAGATCTACAC NNNNNNNN TCGTCGGCAGCGTC) to amplify libraries for 11-13 cycles using Phusion High Fidelity PCR Master Mix (NEB) instead of the Illumina-supplied PCR reagents. The N_8_ sequences correspond to reverse-complemented Nextera DNA indexes N701 to N712 and N501 to N508, respectively (Illumina). Libraries were resolved by agarose gel electrophoresis and DNA was excised in the 300-400 bp range. Gel slices were purified using the Zymo Gel DNA Recovery Kit and sequenced on the Illumina MiSeq using 150-cycle v3 or 600-cycle v2 reagent kits.

Sequencing adaptors were trimmed from reads using cutadapt, mapped to reference genomes using bowtie2, and visualized using IGV. Reference genomic contigs of ECOR strains were obtained from the USDA^49^. SNPs, indels, and large structural variants were analysed using a mapping-free variant finder^50^.

### Phage propagation and plaque assays

High-titer phage stocks were generated by growth in liquid culture. All phage infections were performed at 37°C in LB supplemented with 10 mM MgSO_4_ and 5 mM CaCl_2_. *E. coli* BW25113 or C990 were grown to an optical density (at 600 nm; OD_600_) ∼ 0.5 (1-5 X 10^8^ cfu/ml) and infected with the desired phage at MOI ∼ 5. Infections were allowed to proceed at 37°C with agitation for 5-6 hours until the cultures were clear. Lysates were clarified by centrifugation (8000 X g for 2 min) and sterile-filtered using a 0.45 µm SFCA syringe filter (Millipore). Phage lysates were stored at 4°C.

Bacteriophages T2, T3, T4, T5, T6, T7, P1vir were obtained from Dr. Vivek Mutalik (LBNL), and λvir was obtained from Dr. Ry Young (Texas A&M). To the extent possible, all phages were grown on *E. coli* C990 which lacks restriction-modification. T4 and P1vir could not be grown on C990 and were instead grown on *E. coli* BW25113. To ensure that our phage lysates were free from ancestral Type I R-M methylation marks or other host-passage derived DNA modifications, we serially passaged all phages in the desired hosts 3 times. All phage genomes were subsequently verified by Nextera high-throughput sequencing.

Phage titers were determined against various hosts by the agar-overlay method with LB top agar containing 0.7% agar, 10 mM MgSO_4_ and 5 mM CaCl_2_. Phage spot-titration assays were performed with 10-fold serial dilutions of phage lysates (3-5 µl of each dilution pipetted onto bacteria immobilized in top-agar overlay using a multichannel pipette).

### AG Library Preparation + NGS verification + testing post-conjugation evenness

200 AGs with unique 10 bp DNA barcodes were synthesized (Twist and IDT) and cloned individually under pLlacO-1 control (IPTG inducible, repressed by glucose) into a modified pTn7C185 vector (with mobile CRISPRi components sgRNA and dCas9 removed). Plasmid assembly reactions were performed in 96-well format using the 2X Gibson assembly master mix (NEB) and transformed into chemically-competent BW25141 cells in 96-well format. Transformations were individually plated on LB agar supplemented with 20 µg/ml gentamicin and 0.5% glucose and one colony was initially checked by Sanger sequencing from each plate. Where a successful transformant could not be obtained on the first attempt, more colonies were sequenced in successive rounds until a plasmid with no mutations in the region spanning the pLlacO-1 promoter, the AG ORF, and the barcode sequence was obtained. Verified vectors were miniprepped using the ZR Plasmid miniprep – Classic kit (Zymo) and transformed into chemically-competent WM6026 cells in 96-well format, and each transformation was plated individually on LB agar supplemented with 50 µg/ml gentamicin, 300 µM DAP, and 0.5% glucose.

One colony from each of the 200 WM6026 transformants was grown in 2 ml square-well 96-well plates with shaking overnight, and 100 ul of each culture was then pooled together to yield ∼20 ml of the donor mixture. This mixture was concentrated down to 10 ml by centrifugation at 6000 g for 10 min, and 5 ml of 60% glycerol (20% final) was added to yield the donor library. The 15 ml library was divided into single-use 50 ul aliquots (each containing >50M cells, i.e. >300K of each donor strain) and stored at −80°C.

Genomic DNA was extracted from one of the library aliquots and all constructs in the final pool were verified by Nextera high-throughput sequencing. This allowed us to assess library evenness as well as verify the fidelity of our cloning, colony picking, and strain pooling process. Overall, 195 AGs were cloned successfully. One AG (*orf88*) never yielded any colonies in multiple cloning attempts, possibly due to its cytotoxicity. Three AGs (*orf36, orf52, orf94*) were dropouts during the WM6026 growth step, and another two (*orf25, orf84*) were discovered to contain mutations upon high-throughput sequencing that had been missed by Sanger sequencing. *Orf25* had previously presented difficulties in cloning, and did not yield a mutation-free construct even after checking >12 colonies (whereas most transformations yielded a correct construct on the first attempt). The *orf84* construct contained a mixed base at the last position of the second repeat of the Lac-operator sequence upstream of the ORF, which would not be expected to affect any downstream assays significantly.

The donor library was conjugated into MG1655 *E. coli* according to the Tn7 transposition procedure (above) and a 1000X dilution of the conjugation mixture was plated on LB agar supplemented with 20 µg/ml gentamicin and 0.5% glucose. We routinely use multiple 150 mm X 15 mm bacteriological petri dishes (Corning) to obtain sufficient numbers of transconjugant colonies to assemble libraries. The next day, >>2,000-20,000 transconjugant colonies (10-100X the number of AGs in the library) were scraped and combined thoroughly, and the thick cell mixture was directly frozen at −80°C in 20% glycerol. Genomic DNA was extracted from this mixture and the distribution of DNA barcodes corresponding to individual AGs was measured in both this post-conjugation strain mixture as well as the genomic DNA extracted from the WM6026 donor library aliquot.

To amplify barcodes for high-throughput sequencing, genomic DNA was subjected to two successive rounds of PCR. The DNA barcodes were amplified from genomic DNA using universal primers JSW-SS-170 (CGACGCTCTTCCGATCTNNNNN <u>TGATGTCGTTGTTGCCATCG</u>) and JSW-SS-171 (ACTGACGCTAGTGCATCA <u>CTTTCTGAGCCAGTGTTGCT</u>) and the Q5 Hot Start High-Fidelity 2X PCR master mix (NEB) according to manufacturer’s instructions. Sequencing adaptors were attached in a second round of PCR using amplicons from the first PCR round as the templates, with dual-indexed primer sets JSW-SS-42:53 (CAAGCAGAAGACGGCATACGAGAT NNNNNNNN GTGACTGGAGTTCAGACGTGTGCTCTTCCGATCT <u>ACTGACGCTAGTGCATCA</u>) and JSW-SS-54:61 (AATGATACGGCGACCACCGAGATCTACAC NNNNNN ACACTCTTTCCCTACA<u>CGACGCTCTTCCGATCT</u>), where the N_8_ sequences correspond to reverse-complemented TruSeq HT indexes D701 to D712 and D501 to D508, respectively (Illumina). Template-matching regions in the primers are underlined. Cycling conditions for round 1 were as follows: one cycle at 98°C for 1 min; two cycles at 98°C for 10s, 66°C for 30s, and 72°C for 10s; 22 cycles at 98°C for 10s, and 72°C for 20s; and one cycle at 72°C for 2 min. Conditions for round 2 were one cycle at 98°C for 1 min; two cycles at 98°C for 10 s, 64°C for 30 s, and 72°C for 10 s; 4 cycles at 98°C for 10 s, and 72°C for 20 s; and one cycle at 72°C for 2 min. 0.5-1 ul of unpurified 1^st^ round reaction product was used as template for the 2^nd^ round of PCR. Amplicons from the 2^nd^ round were gel-purified by electrophoresis (3% agarose gel, 4.2 V/cm, 2 hours) and quantified using the 1X dsDNA HS kit with the Qubit 4.0 Fluorometer (Invitrogen). Amplicons were sequenced on a MiSeq using the 150-cycle v3 reagent kit (Illumina) in single-end format (100 bp) with two (8 bp) index reads. DNA barcodes were trimmed from the reads and the proportions of various AGs in the mixtures were determined using the prevalence of the corresponding barcode sequence.

Concordance between the Nextera whole-genome sequencing and the barcode sequencing of the WM6026 donor mixture showed that barcode representation was a suitable proxy for the true distribution of AGs in the mixture and that amplification biases during library preparation were not a significant source of noise. Concordance between barcode sequencing in the WM6026 donor and MG1655 post-conjugation recipient mixtures indicated that the Tn7 transposition and colony scraping process does not meaningfully bottleneck AG distribution.

The AG library was then conjugated (in triplicate) into ECOR strains that were previously confirmed to support robust Tn7 transposition. Strain mixtures were prepared in a procedure analogous to library construction in MG1655, except higher dilutions of the ECOR+AG conjugation mixtures (10,000-20,000X) were plated and the gentamicin concentration on selective plates was lowered to 10 µg/ml to account for higher steady state cell densities and higher gentamicin sensitivity of the wild *E. coli* strains. As with the MG1655 library, thick cell mixtures comprised of >>2,000-20,000 colonies scraped from each replicate conjugation were directly stored at −80°C in 20% glycerol and used to seed cultures for the AG screens (below). Genomic DNA was extracted from each mixture and barcode sequencing was performed to measure the AG distribution in every replicate of all ECOR strain derived libraries. The replicates were very highly correlated, and AG distributions in all libraries were confirmed to be free of significant bottlenecking. These datasets are the reference libraries that serve as the baseline for AG induction and phage infection experiments.

### MOI calculation for liquid killing assays

Phage adsorption to host cells may be inefficient in very dilute cultures (10^6–7^ cfu/ml). Therefore, the apparent multiplicity of infection (MOI-a) might be lower than the expected MOI (MOI-e). We added 2x serial dilutions of a λvir lysate to a 1:100 dilution of a saturated culture of *E. coli* BW25113 that had been allowed to acclimate at 37°C with shaking for 15 min. The proportion of surviving cells was determined (relative to uninfected controls) after a 60-75 min incubation (one phage replication cycle) by plating on LB agar. An apparent MOI value was calculated using the Poisson distribution function with zero occurrences *MOIa* = *In* (*MOIe*) for each phage dilution, assuming that all infected cells would be killed by the virus (i.e. the proportion of surviving cells would be the zero class in the Poisson distribution). The apparent and expected MOI values were plotted against each other and the inverse of the slope (1/m) of the line (MOI-a = m*MOI-e + 0) fit through the data was determined as the fold-change reduction in MOI-a as compared to MOI-e for these experimental conditions. This fold-change reduction was determined to be ∼26x. Hence, all phage infection screens were targeted to an MOI-a value of ∼100 to ensure efficient phage adsorption and infection within the timeframe of a single phage replication cycle.

### AG Screen + Plaque assay confirmation + Data analysis

Thick cell mixtures from each of the three replicates of the various ECOR+AG libraries were thawed on ice and diluted into LB supplemented with Mg^2+^, Ca^2+^, and 10 µg/ml gentamicin. Typically, 1-5 µl of the library mixture was diluted in 2 ml broth to yield an initial OD_600_ of 0.1-0.15. The cultures were immediately split x2, with one tube used to monitor OD_600_ values as the experiments progressed, and the other left untouched for the screen. AG expression was induced by the addition of 1 mM IPTG after acclimatization for 15 min at 37°C in a shaking incubator and OD_600_ values were measured again for each ECOR+AG library at time of induction. Libraries were then grown for ∼80-120 min depending on the relative growth rates of the various ECOR strains, to OD_600_ ∼ 1.2-1.5. A 10^-5^ dilution of each late-log phase culture was then plated on LB agar (with 10 µg/ml gentamicin and 0.5% glucose to repress AGs and avoid fitness costs) to obtain a count of viable colony forming units (cfu). Cultures were immediately diluted ∼1:20 in fresh LB (with IPTG, Mg^2+^, Ca^2+^, and 10 µg/ml gentamicin) and 1 ml aliquots were gently mixed with phage lysates at an MOI-e of ∼30-100 (the actual MOI-e values were recorded the next day when cfu counts were obtained). Parallel uninfected controls were maintained for each experiment, even when the same ECOR+AG library was being tested against different phages and the uninfected controls would be interchangeable. Infections were allowed to proceed for 30-35 min for T-phages and 60-70 min for P1vir and λvir, i.e. one replication cycle. AG induction was limited to 1-2 hrs and subsequent infections were only allowed to proceed for ∼1 phage replication cycle to prevent prolonged induction of various AGs from biasing the results. This setup also avoids biases resulting from epigenetic modification of phages by widespread restriction-modification (R-M) defenses in *E. coli*. However, we note that allowing sufficient time for the phages to complete cell lysis was necessary as fast-growing cells were sometimes able to form colonies on LB agar despite having been infected. These colonies had an uneven size distribution and a “splotchy” appearance, perhaps due to ongoing infection on the plate that was nevertheless not able to outpace colony formation. After the infection period was complete, the cultures were vortexed, diluted 1:100, vortexed again, and 50 µl of the dilution was plated on 2 large (150 mm X 15 mm) LB agar plates (with 10 µg/ml gentamicin and 0.5% glucose to repress AGs and avoid fitness costs). The next day >>2,000-20,000 colonies were scraped from each infected and uninfected set of plates. 5 µl of the thick resulting cell mixtures was reserved for genomic DNA extraction, and the rest was frozen at −80°C in 20% glycerol as a backup. A full technical record of our screens is provided in Supplementary File 4.

AG barcodes were amplified and sequenced as with the WM6026 donor library, with a slight modification. The three biological replicate sets of ECOR+AG experiments were amplified using JSW-SS-171 and different versions of primer JSW-SS-170 containing extra N’s at the start of the sequencing read. The first replicate was amplified with JSW-SS-170 (containing 5 N’s), the second replicate with JBD-SS-254 (CGACGCTCTTCCGATCTNNNNNN TGATGTCGTTGTTGCCATCG) with 6 N’s, and the third replicate with JBD-SS-255 (CGACGCTCTTCCGATCTNNNNNNN TGATGTCGTTGTTGCCATCG) with 7 N’s. This ensured that reads from the three sets would not cross contaminate analyses of replicate experiments. Amplicons were sequenced to the same depth as the baseline libraries on Illumina MiSeq and NextSeq instruments in single-end format (100 bp) with two (8 bp) index reads.

We required exact matches for the 20 bp primer-derived invariant sequences flanking the barcodes. These sequences were trimmed and the proportions of various AG-encoding strains in the mixtures were determined using the prevalence of the corresponding 10 bp barcode sequences. Barcode counts per AG were normalized by dividing each count by the median of each experiment, which would represent neutral AG fitness. A log2 transformation was subsequently applied to obtain fitness scores. Care was taken to sequence each library to similar depth such that fitness scores across experiments were distributed in the same range. Poorly represented AGs were identified in the baseline libraries as well as the infected and uninfected experimental samples using a low read cutoff of 100, or if the log2 normalized average read counts across the three replicates for any AG were >3x the median absolute deviation below the median for that experiment. Reproducibility was determined using T-tests across the three replicates, and we enforced a P-value cutoff of 0.2. The magnitude of the AG fitness effect was determined by (a) subtracting the AG fitness scores in the uninfected library from the infected libraries to assess whether AGs changed host sensitivity to phage infection, and (b) subtracting the AG fitness scores in the baseline library from the uninfected libraries to assess whether AGs change host fitness directly. AGs were selected for further study if they were not poorly represented to begin with, produced reproducible effects across replicates, and changed host fitness by at least 4-fold. Although we had targeted our experiments to phage-host combinations where the host was mostly resistant to the phage in order to find counter-defense AGs, a few hosts turned out to be more sensitive to certain phages in liquid infection than suggested by agar-overlay plaque assays (perhaps due to abortive antiviral defenses). In these cases, our platform was also able to robustly identify AGs that conferred a protective advantage to the host (e.g. Imm and Cor, which both block phage DNA entry ^51, 52^). In practice however, these effects can be very variable and the reproducibility filters typically needed to be relaxed in order to find such superinfection exclusion AGs.

All counter-defense AGs were subsequently conjugated into the wild ECOR hosts where they demonstrated an effect, and tested against all 8 phages (not just the ones that were used in the screen with that host). In several hosts, the AG was found to additionally sensitize the host to phages that were omitted in the screen. Not all the screen phenotypes could be confirmed in agar-overlay plaque assays, possibly due to variations in cell growth between liquid and solid media, or due to the requirement for successive infection cycles for plaque formation that would render plaque assays less sensitive than single-infection cycle dynamics in our screen. Nevertheless, further experiments were performed only in phage-host-AG combinations that could be confirmed by plaque assay due to ease of follow up testing.

Conditional lethal AGs were identified directly from the heatmap of fitness scores, with a subset selected for further experimentation using an arbitrary fitness cutoff of ∼6-7 (∼100-fold).

### Tn5 Dropout Screen + Data analysis to find phage exclusion systems

Tn5 whole-genome knockout strain libraries were constructed using the RB-TnSeq donor library (pKMW7-derived strain APA766 from Dr. Adam Deutschbauer) conjugated with the recipient ECOR strain in a biparental mating on LB plates supplements with 300 µM DAP. The next day, cell mixtures were scraped and resuspended in 1 ml LB and 1000X dilutions of this conjugation mixture were plated on 10 large (150 mm X 15 mm) LB plates (lacking DAP) supplemented with 50 µg/ml kanamycin. Approximately 0.5-1 million colonies were scraped taking care to break colonies apart on the surface of the plates, mixed very thoroughly, aliquoted and stored at −80°C with 20% glycerol, with ∼5-10 µl reserved for genomic DNA extraction and reference library preparation.

RB-TnSeq allows the same mutant library to be challenged under various conditions such as challenge by various phages, and simplifies the downstream sequencing-library preparation by substituting barcode sequencing for all experimental conditions after initial characterization of the reference library. To characterize the Tn5-knockout reference libraries in the various ECOR strains, genomic DNA was extracted from 5-10 µl of the cell scrape and prepared for sequencing using the NEBNext Ultra II FS DNA Library Prep Kit (NEB) with the following modifications. 500 ng DNA was fragmented for 25 min according to manufacturer’s instructions. The adaptor ligation step was performed as directed albeit with a custom pre-annealed oligo adaptor comprising of YL001 (/5Phos/GATCGGAAGAG/3ddC/) annealed to one of YL002:005 (AGCGGCAATTTCACACAGGACAAGCAGAAGACGGCATACGAGATNNNNNNNN<u>AGTG</u>GTG ACTGGAGTTCAGACGTGTGCTCTTCCGATC*T) where the 4-base underlined sequence serves as a variable barcode unique to each oligo. The USER Enzyme steps post-ligation were skipped and the bead-cleanup was performed using 40 µl and 20 µl of Sample Purification Beads successively as instructed for library size selection in the 150-250 bp range. The PCR amplification of transposon-genome junctions was performed using the cycling parameters described in the kit for 11 cycles with Q5 Ultra II FS Master Mix (NEB) using primers YL006 (AGCGGCAATTTCACACAGGA) and JBD-SS-272/oAD493 (ACACTGGCAGAGCATTACGCCCT) instead of the kit-supplied primers. A second nested-PCR enrichment step was performed for 11 cycles using the same cycling parameters as before, using the entire bead-purified reaction product from the first step as a template with primers YL009 (CAAGCAGAAGACGGCATACGAG) and JBD-SS-273:280 (AATGATACGGCGACCACCGAGATCTACAC TATAGCCT ACACTCTTTCCCTACACGACGCTCTTCCGATCT NNNNNNN GCAGGGATGTCCACGAGGTCTCT) where N_8_ barcodes correspond to reverse-complemented Illumina TruSeq HT D501-508 indexes. Bead-purified amplicons were quantified using the 1X dsDNA HS kit with the Qubit 4.0 Fluorometer (Invitrogen) and sequenced on Illumina MiSeq and NovaSeq instruments in single-end format (150 bp) with two (8 bp) index reads.

The first 100 bp of these reads contain transposon derived sequences and a DNA barcode that uniquely identifies a particular Tn5 insertion. The last 50 bp of these reads contain host-genome derived sequence that help determine the location of the transposon insertion and hence construct a map between the Tn5 barcode and the insertion site (IS) in the genome. Briefly, 20 bp barcodes and 16-50 bp IS sequences were trimmed from the sequencing reads. To isolate the barcode and IS, we allowed 3 mismatches in the 23 bp invariant sequence upstream and up to 6 mismatches in the 50 bp downstream of the barcode using a Hamming distance function to compare the expected and actual sequences. Any reads failing these sequence-fidelity requirements were discarded. Reads containing an IS that matched the Tn5 transposon vector in the donor strain were also discarded. The 4 Ns adjacent to the underlined index sequences in YL002:005 were also sequenced as part of one 8 bp index read and served as a tag to remove PCR duplicates. This allowed us to distinguish between sequencing “reads” and DNA “fragments” present in the sample at the adaptor ligation step. Thus, reads containing unique combinations of barcode, PCR tag, and IS were entered into the barcode/IS map for each Tn5 library, each representing a distinct DNA fragment (but also machine errors in sequencing, chimeric PCR amplicons, 3’ truncations etc.). The filtered ISs were then mapped to genes within their respective host genomes. As described previously^53^, it is possible for transposon barcodes to map to the genomes ambiguously. Barcodes associated with multiple distinct ISs were retained only if their most frequently represented IS accounted for over 90% of the fragments containing that barcode. We then counted the number of DNA fragments recovered per gene and used this as a reference library for subsequent BarSeq experiments to assay depletion of strains with mutations in specific genes upon phage infection.

Tn5 mutant strain libraries of the various ECOR strains were then challenged by various phages in a procedure analogous to the ECOR+AG screens. Cultures of Tn5 libraries were initiated with 100 µl of thick cell scrape mixture in 100 ml LB supplemented with 50 µg/ml kanamycin, Mg^2+^, and Ca^2+^, to an initial OD_600_ value of 0.1-0.15. Cultures were grown at 37°C with shaking for 80-90 min to an OD_600_ of 0.7-1. A 10^-5^ dilution of each late-log phase culture was then plated on LB agar (with 50 µg/ml kanamycin) to obtain a count of viable colony forming units (cfu). Cultures were immediately diluted ∼1:10 in fresh LB (with Mg^2+^, Ca^2+^, and 50 µg/ml kanamycin) and 1 ml aliquots were gently mixed with lysates of the appropriate phages at an MOI-e of ∼30-100 (the actual MOI-e values were recorded the next day when cfu counts were obtained). Since the parental ECOR strains were typically quite resistant to phage infection to begin with, only mutants with transposon-inactivated defense systems would be expected to drop out upon infection. In practice however, some strains exhibited more pronounced mortality upon prolonged phage infection in the Tn5 screens. In particular, in experiments with T2 and the Tn5 library constructed in the *ΔwecA* strain of ECOR21 (which was generally more susceptible to infection by T2 and other phages), MOI was titrated in three parallel infection screens (with varying amounts of phage lysate added: 350 µl, 100 µl, and 25 µl) to empirically ensure specific depletion of only the phage-sensitive mutants. Parallel uninfected controls were maintained for each experiment, and each set of experiments was performed in biological duplicates. Infections were allowed to proceed for 5 min at the benchtop without shaking, and 75 min at 37°C with shaking. Since an interaction between the host and phage had already been identified in the earlier AG screen, infections were not restriction to a single phage replication cycle, but allowed to proceed further to achieve full depletion of the phage-sensitive Tn5-knockout mutants in the pool. After the infection period was complete, the cultures were vortexed, diluted 1:10, vortexed again, and 100 µl of the dilution was plated on 2 large (150 mm X 15 mm) LB agar plates (with 50 µg/ml kanamycin). We note that this dilution yields bacterial lawns and in our experience this preserves diversity of surviving colonies without degrading the signal of dropouts by phage-killing in the experiment. The next day, bacterial lawns were scraped from each infected and uninfected set of plates. 5 µl of the thick resulting cell mixtures was reserved for genomic DNA extraction, and the rest was frozen at −80°C in 20% glycerol as a backup. A full technical record of our screens is provided in Supplementary File 8.

To amplify barcodes from the post-selection infected and uninfected Tn5 libraries, we followed a modified BarSeq98 method^53^, with two successive rounds of PCR. The transposon barcode sequence was amplified by PCR using Q5 Ultra II FS Master Mix (NEB) with primers JBD-SS-310 (CGACGCTCTTCCGATCTNNNNNNN <u>GATGTCCACGAGGTCTCT</u>) and JBD-SS-311 (ACTGACGCTAGTGCATCA <u>GTCGACCTGCAGCGTACG</u>). Template-matching regions in the primers are underlined. Sequencing adaptors were attached in a second round of PCR using amplicons from the first PCR round as the templates, with dual-indexed primer sets JSW-SS-42:53 and JSW-SS-54:61 (sequences already provided in section describing AG screening). Cycling conditions for round 1 were as follows: one cycle at 98°C for 4 min; two cycles at 98°C for 30s, 63°C for 30s, and 72°C for 30s; 22 cycles at 98°C for 30s, and 72°C for 30s; and one cycle at 72°C for 4 min. Conditions for round 2 were one cycle at 98°C for 1 min; two cycles at 98°C for 10 s, 64°C for 30 s, and 72°C for 10 s; 4 cycles at 98°C for 10 s, and 72°C for 20 s; and one cycle at 72°C for 2 min. 0.5-1 ul of unpurified 1^st^ round reaction product was used as template for the 2^nd^ round of PCR. Amplicons from the 2^nd^ round were gel-purified by electrophoresis (3% agarose gel, 4.2 V/cm, 2 hours) and quantified using the 1X dsDNA HS kit with the Qubit 4.0 Fluorometer (Invitrogen). Amplicons were sequenced on Illumina MiSeq and NovaSeq platforms in single-end format (150 bp) with two (8 bp) index reads.

DNA barcodes were trimmed from the reads and the proportions of various Tn5-knockout mutants in the mixtures were determined using the prevalence of the corresponding barcode sequence. Briefly, we allowed 3 mismatches in the 18 bp invariant sequence upstream, and 4 mismatches in the 18 bp downstream of the transposon barcodes using a Hamming distance function to compare the expected and actual sequences. The frequency of occurrence of each barcode (representation number) was counted. Transposon barcodes were matched to those found in the reference libraries, inferred to represent knockouts in the genes containing the corresponding ISs, and the various barcode representation numbers were added together for each gene. Cumulative transposon insertions per gene were then compared between the infected and uninfected samples (with the reference library as a baseline to filter out poorly represented, presumably essential genes) using the same data analysis procedure as for the AG screen, albeit with two biological replicates instead of three.

### Tn5 Suppressor Screen + Data analysis to find abortive loci

Tn5 whole-genome knockout strain libraries were constructed using the RB-TnSeq donor conjugated with the recipient ECOR strain as before, with modifications. No phages are involved. The recipient ECOR strain was equipped with the AG (as a single copy chromosomal Tn7 insertion as usual) which triggers PCD, presumably by activating an Abi system. The trigger AG was inserted prior to Tn5 library construction in order to minimize passages of the library prior to screening and avoid potential bottlenecking due to fitness effects of the various Tn5 knockouts. Libraries were constructed as above, except that conjugation was carried out under strict glucose repression to prevent PCD activation at this stage, and the conjugation mixture scraped from the LB+DAP+glucose plates was then plated directly on LB plates containing 10 µg/ml gentamicin and 1 mM IPTG in addition to 50 µg/ml kanamycin. We typically needed to plate the entire conjugation mixture scraped from five 100 mm LB+DAP+glucose plate onto five 150 mm LB+IPTG+Kan+Gm plate in order to obtain sufficient numbers of suppressor colonies for characterization by high-throughput sequencing. Control samples were obtained by plating 1000-2000X dilutions on LB+glucose+Kan+Gm plates. Suitable dilutions were empirically determined for the various ECOR hosts. The next day, suppressor colonies were scraped from the large plates and stored at −80°C with 20% glycerol, with ∼5-10 µl reserved for genomic DNA extraction and sequencing. Genomic DNA was extracted and libraries were prepared and sequenced akin to the treatment of the RB-TnSeq/NEBNext reference libraries detailed above. There was no need for barcode sequencing since the screen identifies survivors instead of dropouts, and each library was only characterized once after IPTG induction/selection for suppressor mutants. Tn5 reference libraries were constructed from the data using the same procedures for the baseline libraries constructed in the various naïve (without AGs) ECOR strains above. The number of transposon insertions (i.e. the number of uniquely mapping ISs from distinct DNA fragments) recovered per gene were used directly as a measure of survival of mutants despite expression of PCD-triggering AGs. In practice, no statistical analysis was necessary to identify the genetic loci responsible for PCD, as the signal-to-noise ratio was sufficiently high to allow visual identification of hits when transposon insertion frequency was plotted in control vs experimental samples (i.e. when the trigger AG was repressed by plating on glucose-containing media, vs induced with IPTG).

### LPS gel electrophoresis

Lipopolysaccharide was prepared from liquid overnight cultures of wild *E. coli* strains expressing the relevant AGs (LB + IPTG). Briefly, 50-100 µl of saturated culture was spun down and cells were thoroughly resuspended in 100-200 µl of lysing buffer (62.5 mM Tris-HCl, pH6.8, 10% glycerol, 2% SDS, 4% ß-mercaptoethanol). We routinely supplement 1X Laemmli Buffer (BioRad) with SDS and ßME to make lysing buffer. Samples were boiled at 100°C for 15 min, and allowed to rest at room temperature for 15 min. 100 µg of Protease K were added to each sample and protein was digested at 60°C for at least 1 hr. 15 µl of each sample were run onto a 12% polyacrylamide TGX Mini-Protean Precast gel (BioRad) at 200V for 35 min, and visualized using the Pro-Q Emerald 300 LPS stain kit (Thermo Fisher) precisely according to manufacturer’s instructions.

### Allelic Exchange for wecA deletion

The R6K-origin pKEK2201 vector (from Prof. Karl E. Klose) carrying the desired genetic modification (eg. a *wecA* deletion) flanked by 1000 bp homology arms was prepared by Gibson assembly, and delivered as a suicide vector into wild *E. coli* strains via conjugation from the WM6026 auxotrophic donor. Conjugation was performed using the same method used to deliver AGs, albeit with biparental instead of triparental mating. Successful transconjugants (containing a chromosomally integrated plasmid after the first homologous recombination event HR1) were selected by incubation on LB plates supplemented with kanamycin. The second recombination event HR2 to complete the allelic exchange was selected via *sacB*-mediated counterselection on LB plates supplemented with 300 mM sucrose after overnight growth of HR1 transconjugants. Successful allele exchange was verified by PCR, and by LPS electrophoresis to assay for the absence of O-antigen in the case of the *wecA* deletion.

### Live cell Microscopy

Cells were prepared for microscopy, immobilized under an LB agarose pad, and imaged as described previously^54^. Where necessary, AGs were pre-induced with 1 mM IPTG for 30 min immediately before harvesting for microscopy.

### Verification of abortive loci activity

Abortive/PCD systems identified by transposon screening of conditionally lethal AGs were cloned by Gibson assembly using the NEBuilder HiFi DNA assembly master mix (NEB) onto pBAD Myc/His A vectors along with their native promoters wherever possible. AGs that trigger PCD systems (or appropriate controls) were conjugated into DH10ß/TOP10 strains carrying pBAD-PCD plasmids using the Tn7 transposition method (above). Where the restriction/PCD systems prevented conjugation of AGs, the order of transformations was reversed – the AG was conjugated into naïve DH10ß/TOP10 cells first, and subsequently these strains were transformed with the pBAD-PCD vectors. Successful transconjugants were grown overnight with glucose to repress trigger AGs, and viability was tested the next day by spotting 10-fold serial dilutions of saturated cultures on LB agar plates with four additives: 1) Glucose repression to assess background toxicity, (2) Arabinose induction of PCD systems alone to assess fitness costs of over-expression of abortive components without the trigger AGs, (3) IPTG to assess ability of AGs to trigger PCD without over-expression of abortive components (i.e. under leaky expression conditions, or when expressed from the native promoters where present), and (4) IPTG and Arabinose to assess ability of AGs to trigger PCD when abortive components were also over-expressed.

### Affinity purification of AGs from wild E. coli hosts

FLAG-tagged AGs were conjugated into ECOR strains by Tn7-transposition (above) according to Supplementary File 9. Each strain was grown overnight at 37°C in 2 mL of LB supplemented with .5% glucose and 10 mg/mL gentamicin, then diluted 1:100 into 100 mL LB supplemented with 10 mg/mL gentamicin and grown at 37°C for 30 minutes. The media was then supplemented with 1 mM IPTG and grown for another 2.5 hours. “Popcorn” pellets were collected by centrifugation for 30 minutes at 4,000 X g, resuspension in 100 µL of ice-cooled lysis buffer [50 mM Tris pH 7.4, 150 mM NaCl, 1 mM EDTA, 1 mM MgCl_2_, 0.5% NP40 (Research Products International), 125 U/mL Benzonase (Millipore), 1x protease inhibitor cocktail (Roche, cOmplete ULTRA tablets, mini, EDTA-free), 0.5 mg/ml lysozyme (Fisher Scientific)], followed by dropwise addition into liquid nitrogen and stored at −80°C. Popcorn pellets were lysed by 10 cycles of cryomilling for 2 minutes at 12 counts per second in a SPEX SamplePrep 6870D Freezer/Mill and stored at −80°C.

For each phage-host combination, 3 replicate lysates were reconstituted using 200 mg of the cryomilled powder resuspended in 750 µL of lysis buffer. Lysates were cleared by centrifugation at 16,000 X g at 4°C for 15 minutes. Anti-FLAG magnetic agarose beads (Pierce) were washed on a magnet with 1 mL of ice-cooled IP buffer (50 mM Tris pH 7.4, 150 mM NaCl, 1 mM EDTA, 1mM MgCl_2_) twice and resuspended to their original volume in IP buffer. Cleared lysates were incubated with 30 µL of FLAG beads on an end-over-end tube rotator at 10 rpm at 4°C overnight. The supernatant was separated from the beads on a magnet. Beads were washed once with 700 µL of .05% NP-40 and 1x protease inhibitor cocktail in IP, once with 700 µL of .05% NP-40 in IP, and 3 times with 700 µL of IP. Proteins were eluted by incubation with 25 µL of 100 µg/ml 3xFLAG peptide (Sigma) in 0.05% RapiGest (Waters Corp) in IP buffer on an end-over-end tube rotator at 10 rpm at 4°C overnight. The eluate was separated from the beads on a magnet. The elution step was then repeated with the same beads and fresh elution buffer at room temperature for 1 hour. The eluates were combined and stored at −20°C.

### Mass spectrometry sample preparation and data acquisition

Samples were prepared for Mass Spectrometry (MS) as described previously in the Filter-Aided Sample Preparation Protocol^55^ with the following modifications. Before transferring to an AcroPrep Advance 10k Omega plate, each eluate was resuspended in 100 µL of 8M Urea (Thermo Scientific) dissolved in 50 mM ammonium bicarbonate (MP Biomedicals) and 5 mM TCEP (Aldrich), followed by centrifugation to dryness. During alkylation of cysteine residues, 10 mM chloroacetamide (Thermo Scientific) was used instead of 10 mM iodoacetamide. Wash steps prior to proteolysis were carried out with 200 μL of 20 mM ammonium bicarbonate (instead of 100 μL). Proteolysis was performed without the addition of lysyl endopeptidase.

Proteolysis was quenched by addition of 10% trifluoroacetic acid (Thermo Scientific) to reach ≈ pH 2. Peptides were then purified over a BioPureSPE Mini 96-Well Plate (Nest Group). Briefly, C18 columns were activated by washing once with 400 µL of acetonitrile (Fisher Scientific) and twice with 400 µL of 0.1% formic acid (Thermo Scientific) in water solution, LC–MS grade (Thermo Scientific). Samples were loaded and centrifuged at 1500 X g for 2 minutes. The bound peptides were washed 3 times with 400 µL of 1% formic acid before elution with 150 µL of 50% acetonitrile in 0.1% trifluoroacetic acid. Peptides were vacuum dried and stored at −80°C.

Samples were resuspended in 15 µl of 0.1% formic acid prior to loading for MS. The MS acquisition was performed as described^56^ with same MS parameters and LC configuration.

### Mass Spectrometry data analysis and statistical analysis

Raw files were processed in FragPipe^57^ using the LFQ-MBR workflow with minor modifications. The files were searched against a combined database of protein sequences from ECOR1, ECOR3, ECOR15, ECOR21, ECOR22, ECOR66^49^, and MG1655 with duplicate entries removed and common contaminants added. Decoys were generated by pseudo-inversion. Protein and peptide FDR was fixed to 1% and MBR was disabled. Cysteine carbamylation was set as fixed modification while N-term acetylation, methionine oxidation and pyro-glu formation were set as variable modification with a maximum of 3 modifications per peptide.

MaxLFQ intensities were used for statistical analysis^58^. Each triplicate set of host-AG samples was quantile normalized against a triplicate set of samples from the same host expressing *orf74*. For quantile normalization, all proteins in each sample were ranked according to their MaxLFQ intensity. Next, the mean value of the intensities of proteins occupying a given rank was calculated, then each protein’s value was replaced with the mean value according to its rank. Fold changes were calculated by taking the log2 ratio of average normalized values between the test set and the control set. For each host-AG sample, all positively enriched proteins were ranked according to their fold change. For each AG, proteins were assigned “summed rank” values by calculating the sum of the ranks of each protein across all hosts where that AG was tested. P-values were calculated by student’s t-test on the normalized sets of replicates from the test and control samples. Top enriched candidates for each AG were selected based on their summed ranks and p-values.

### Ronin systems phylogenetic analysis

The *ronA* nucleotide sequence in the ECOR55 Ronin system was used to identify similar genes by BLASTN. *RonA*-encoding bacterial genomes were downloaded using the entrez direct e-utilities command line tools. Of 100 complete genomes retrieved, *ronA* homologs had been annotated in 84. *hsdR* genes and other Type I R-M components were identified using phmmer and Defense Finder^59^ respectively. Ronin-adjacent genetic architectures were visualized using Clinker^60^. All ronin-adjacent architectures are shown in HTML files with mouse-over annotations for genes.

## Acknowledgements

S.S. was supported by the Damon Runyon Fellowship Award (DRG 2352-19). Experiments were funded by awards to J.B.D by the Vallee, Searle, and Kleberg Foundations. K.S.M. and E.V.K. are supported by intramural funds of the US Department of Health and Human Services (National Institutes of Health, National Library of Medicine). We thank Jason Peters for technical discussions, Andy Millard for the Enterobacteriophage genome dataset, and Ry Young, Andrew Fire, Carol Gross and Jonathan Weissman for guidance.

## Author Contributions

Conceptualization, S.S., J.B.D.; Methodology, S.S., E.L.; Resources, A.F., D.S.; Software, S.S.; Investigation, S.S., H.C., D.S.G.; Formal Analysis, S.S., H.C., M.J., K.S.M., E.V.K.; Visualization, S.S., H.C., M.J., K.S.M.; Writing – Original Draft, S.S., H.C.; Writing – Reviewing & Editing, S.S., H.C., M.B., E.V.K., J.B.D.; Supervision, S.S., J.B.D.; Funding Acquisition, J.B.D.

## Additional Information

Supplementary Information is available for this paper. Mass Spectrometry data available at PRIDE https://www.ebi.ac.uk/pride/login with Username: and Password: 83OmeYpD. High-throughput sequencing data available at SRA with accession PRJNA952709. Correspondence and requests for materials should be addressed to. Reprints and permissions information is available at www.nature.com/reprints.

**Supplementary Figure 1.**
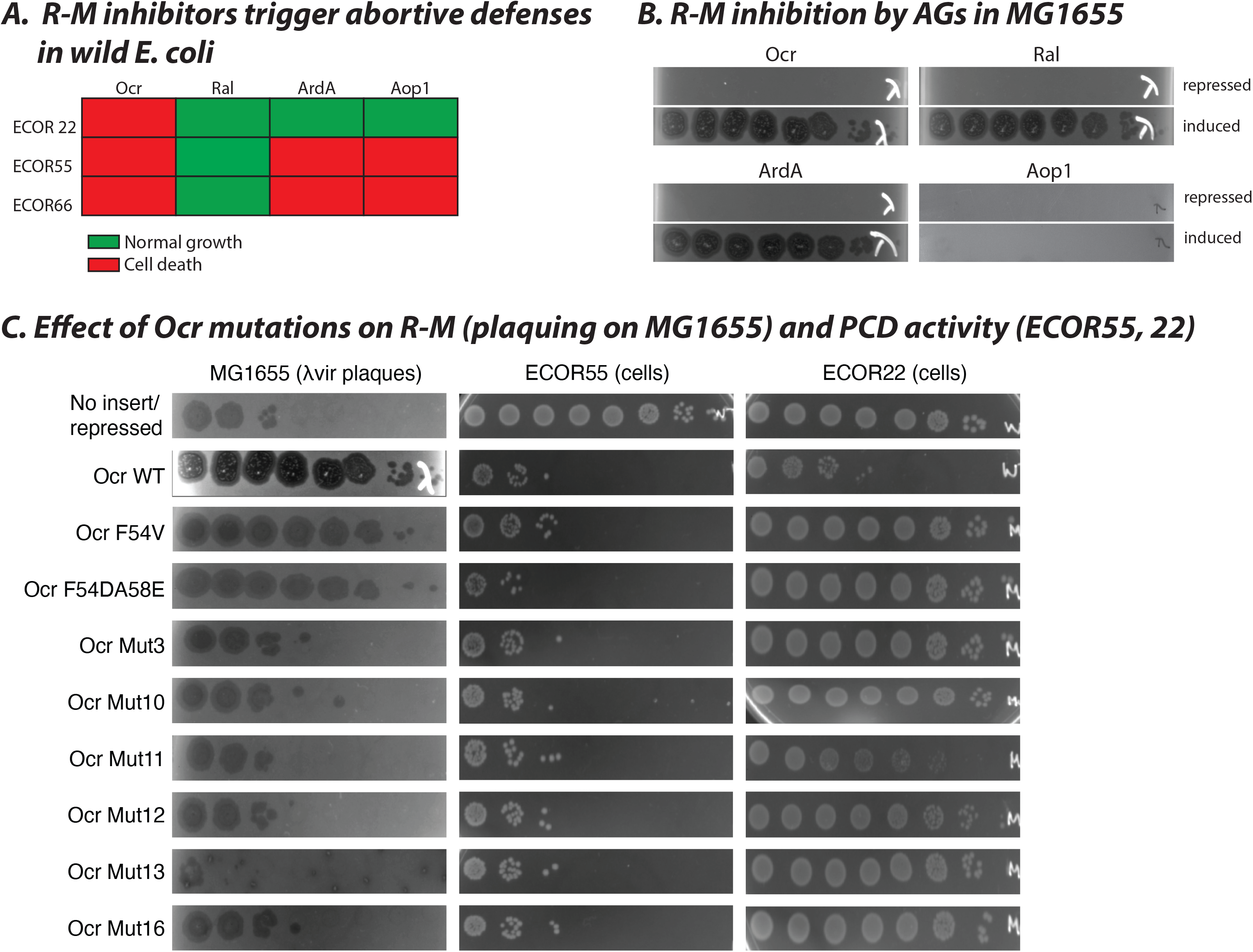
Conditional-lethal phenotypes of known R-M inhibitors in wild *E. coli.* **(A)** Schematic representation of conditional-lethality data from Figure 1d, showing only known R-M inhibitors and an uncharacterized AG, *orf7*:*aop1*. **(B)** AGs from (A) tested for their ability to block EcoKI restriction of phage λvir in plaque assays with restriction-competent *E. coli* MG1655. **(C)** Various mutants of Ocr tested in MG1655 for their ability to block EcoKI restriction of λvir, and in ECOR22 and ECOR55 for their ability to trigger PCD activity of EcoR22I and Ronin respectively. F54V mutant of Ocr (known to evade a previously described ARIA, PARIS) still triggers ARIA activity in ECOR55 but not ECOR22. Numbered Ocr mutants were described elsewhere^19^. Notably, Ral, which operates by allosterically hyperactivating the methyltransferase^61^ (unlike Ocr and ArdA) does not trigger PCD in any strains. Conversely, Aop1 elicits PCD from the same systems triggered by Ocr and ArdA in (A) but does not inhibit Type I R-M activity in (B). The predicted structure of Aop1 does not resemble a DNA mimic (Supplementary File 7), and the protein is not negatively charged (pI ∼4 for Ocr/ArdA, but ∼8 for *aop1*), suggesting a distinct mechanism of R-M inhibition. Thus, depending on their mechanism, some R-M inhibitors trigger PCD in these strains, whereas others do not.

**Supplementary Figure 2.**
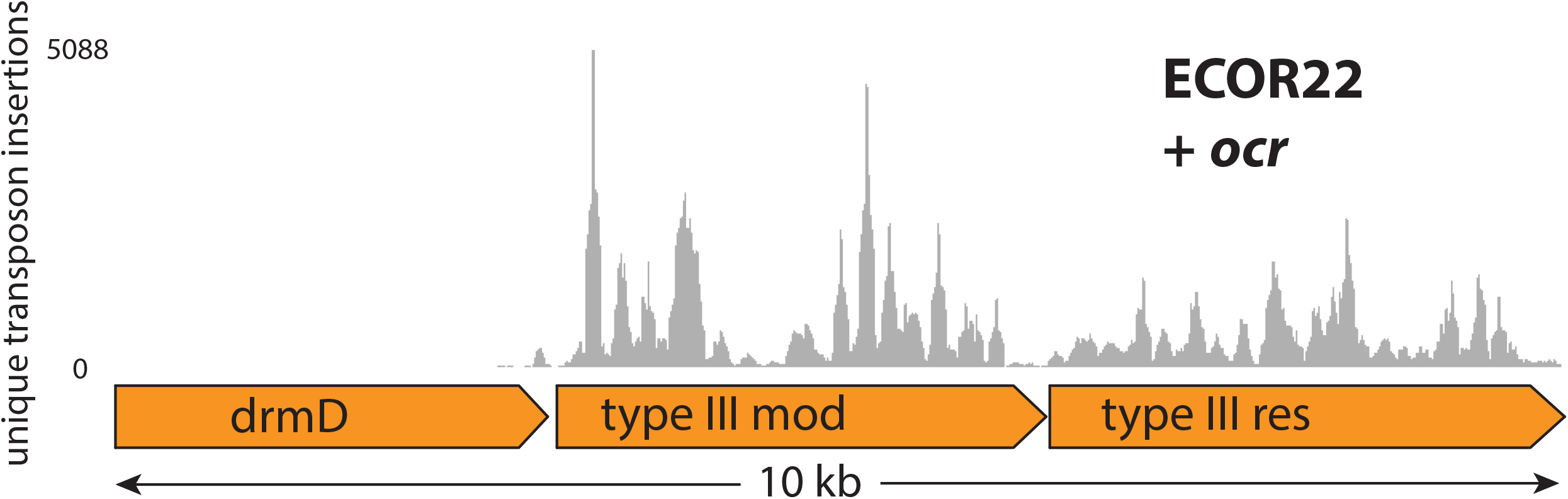
Transposon insertions that suppress AG-induced conditional-lethal phenotype in ECOR22. Distribution of unique transposon insertions recovered from ECOR22 survivors from Tn5 libraries upon *ocr* expression, mapped to the EcoR22I locus.

**Supplementary Figure 3.**
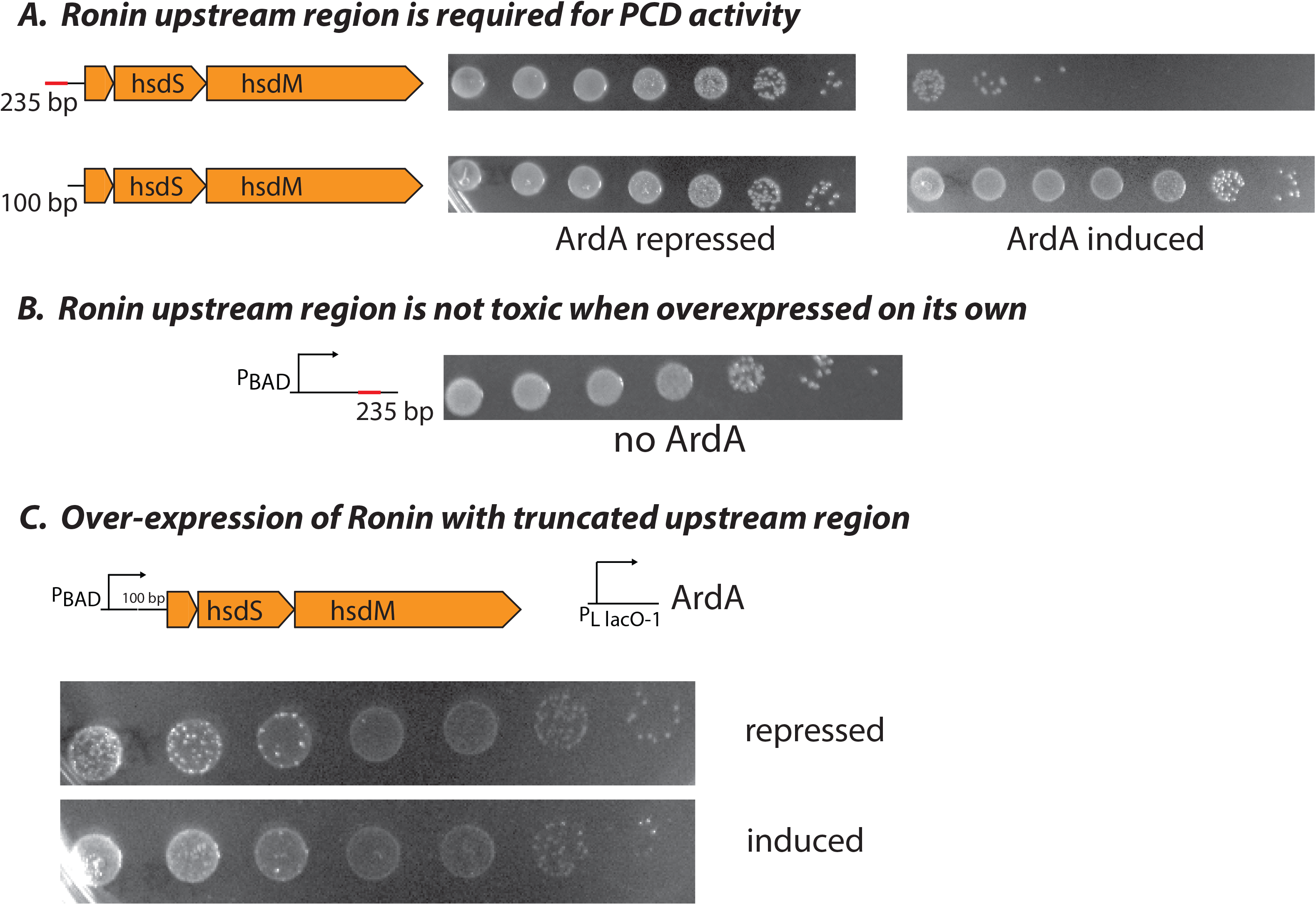
Upstream non-coding region is required for Ronin PCD activity. **(A)** The first 135-bp of the 235-bp upstream non-coding region (marked by a thick red line) are required for activity as truncation down to 100-bp causes inactivation. ArdA is chromosomally integrated under pLlacO-1 control and strains in the left (ArdA repressed) and right (ArdA induced) panels are isogenic. **(B)** Over-expression of 235-bp upstream region is not toxic in TOP10 indicating that it is not the PCD effector. **(C)** As in (A), PCD phenotype of truncated Ronin system when overexpressed (indicated as “pBAD”). Over-expression of truncated Ronin did not rescue the PCD defect indicating that the upstream region contributes something in addition to the promoter.

**Supplementary Figure 4.**
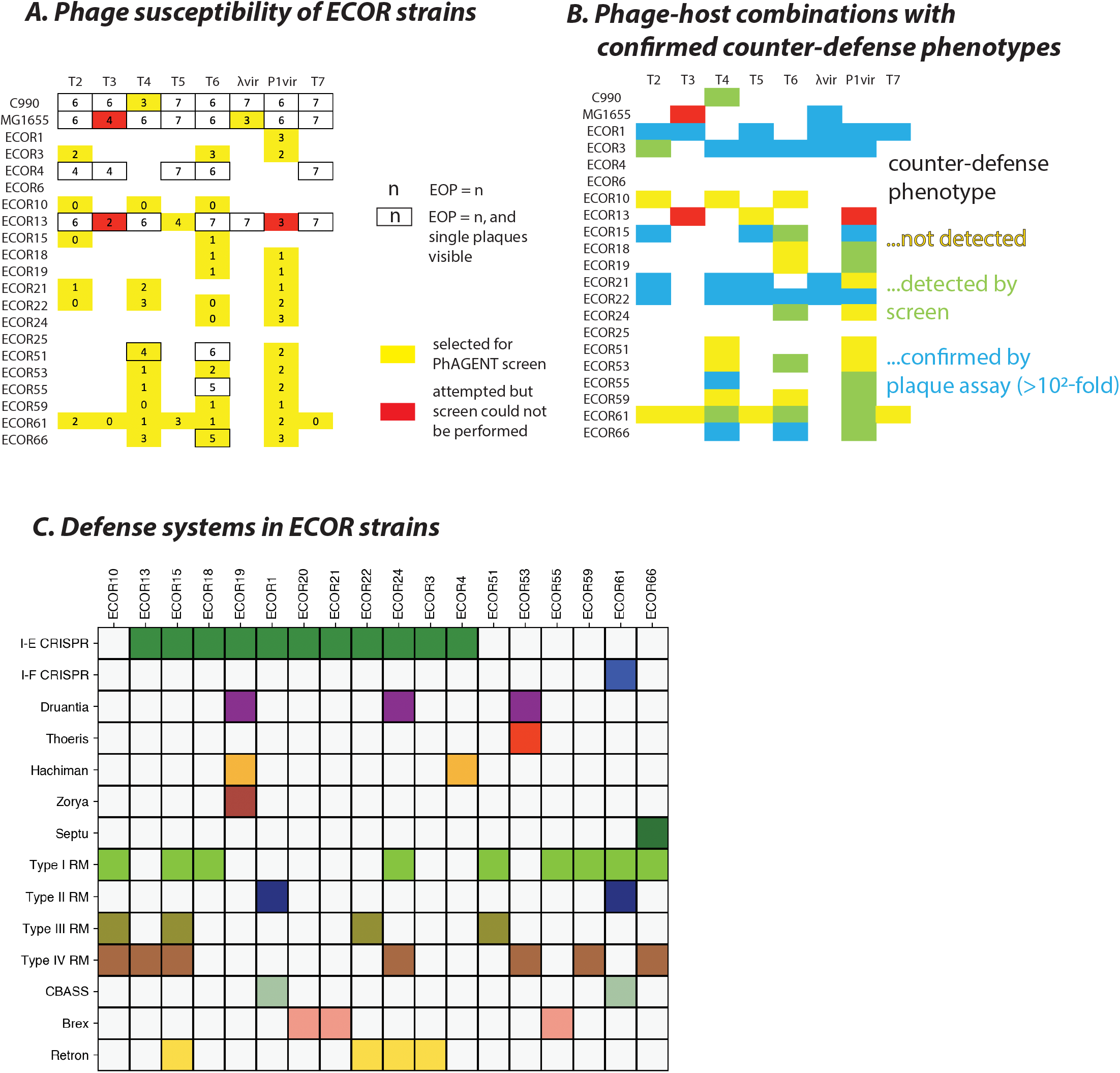
Phage-host combinations included in AG screens. **(A)** Phage susceptibility of select wild ECOR strains. Numbers in boxes are log10-transformed infection scores (7 = 10^7^-fold higher plaquing than 0). Borders indicate that individual plaques were visible in that phage-host combination. Combinations with attenuated phage infection that were selected for the AG screen are in yellow. Combinations with attenuated infection that were attempted but yielded no survivors in liquid infection experiments (perhaps due to Abi phenotypes) are in red. **(B)** Phage-host combinations where counter-defense phenotypes were confirmed for any AGs. Red denotes combinations where the screen could not be completed (as in (A)). Yellow: combinations that were tested where none of the 200 AGs produced counter-defense effects. Green: combinations where phage-sensitizing effects could not be verified by plaque assays. Blue: combinations where counter-defense phenotypes were confirmed by plaque assay (at least 100X higher EOP upon AG expression). **(C)** Defense repertoire of selected ECOR strains. Computational prediction of the presence or absence of various defense systems in ECOR strains used in the AG screen. Predictions were generated using the ISLAND software suite^62^.

**Supplementary Figure 5.**
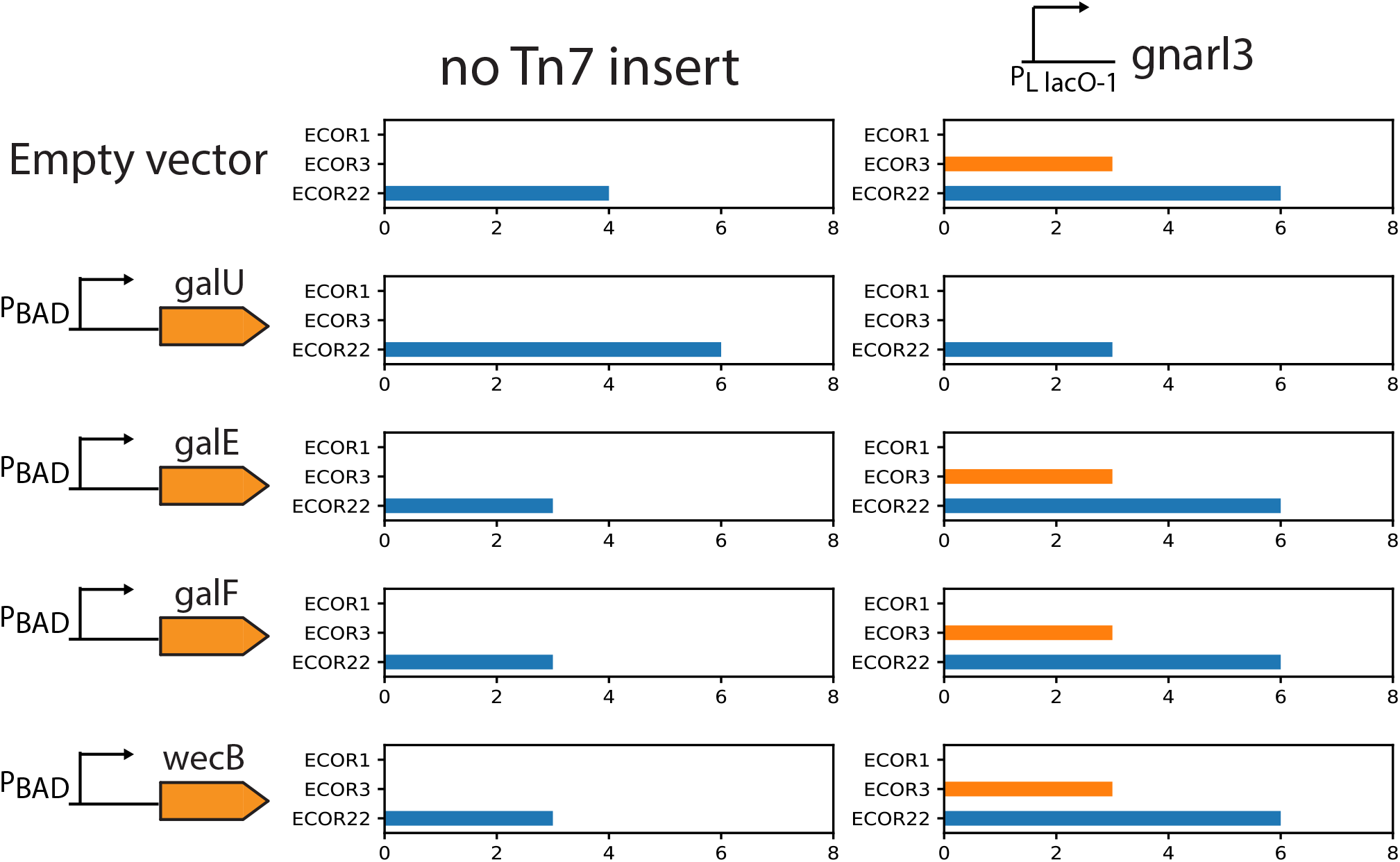
Over-expression of *galU* reverses T4 sensitization by *gnarl3*. log10-transformed T4 infection scores in wild hosts ECOR1 (green), ECOR3 (orange), ECOR22 (blue) with or without UDP-glucose biosynthesis pathway genes *galU, galE, galF,* and ECA precursor *wecB* cloned onto a plasmid and overexpressed.

**Supplementary File 1. Worked example of AG-finding algorithm.** Readme.txt describes how to run the code. Sample database of Salmonella phage genomes is included. All ARs from this phage dataset are reported in the output file SPFMphages_accRegions_nr_all.txt. Any AR can be extracted using the supplied code and then visualized using Clinker^60^. The AR shown in Figure 1d is extracted by default.

**Supplementary File 2. Manually inspected accessory regions from Enterobacteriophage genomes.** Visualizations of all ∼200 accessory regions (ARs) that were manually inspected. The first slide in the file shows the layout for the rest, with the AR score calculated according to the formula in Methods, followed by a unique identifier for each hypothetical phage group (“phamily”). Each AR was also given a short, memorable name (top right). Slides with green backgrounds indicate ARs from which genes were selected for the AG screen. Authors’ impressions upon AR inspection are included in the Notes section.

**Supplementary File 3. Phage accessory genes selected for AG screens.** 200 accessory genes (AGs) selected for screening. Genomic locations, protein and phage genome identifiers, and protein descriptions are indicated. IDs denoted by “UDP” correspond to the orf/AG numbers used throughout the manuscript.

**Supplementary File 4. Experimental record of AG screens.** A detailed record of all AG screens. The first sheet contains phage-host combinations, dates of the experiment, and preliminary results. The next three sheets each correspond to one replicate each of AG screening. Empirically measured multiplicity at time of infection, and dilutions used before plating are also recorded. Screens that had to be repeated for technical reasons are color-coded.

**Supplementary File 5. Hits from transposon-mediated PCD suppressor screens.** Follow-up screens to find inactivating transposon insertions in putative Abi mechanisms that might trigger PCD in response to conditional-lethal AGs. These screens are performed with RB-TnSeq libraries prepared with ECOR hosts already containing IPTG-inducible trigger-AGs (no phages present). Genes inactivated by transposon insertions that were overrepresented in the AG-on condition (IPTG-induced) relative to the AG-off condition (Glucose-repressed) are reported as hits.

**Supplementary File 6. Genomic architectures of all ECOR55-like Ronin systems.** Spreadsheet contains accession numbers and genomic locations of all *ronA* homologs. Grouping variables indicate whether a Type I R-M system (containing *hsdR, hsdM,* and *hsdS*) was found anywhere in the genome, and also whether a full (>400 aa) DUF4102 integrase was present immediately upstream of Ronin. HTML files split Ronin systems into two groups based on the presence of the integrase, and gene annotations can be viewed by mouse-over.

**Supplementary File 7. AlphaFold predictions for all AGs that produced phenotypes in AG screens.** PDB files containing predicted structures of *orf7* (*aop1*; triggers PCD by Ronin), *orf24* (triggers PCD in ECOR61), *orf48*, *orf63*, *orf92*, (*gnarl1-3*; putative O-antigen modifiers), *orf98* (triggers PCD in ECOR22), *orf116* (*abc1*; triggers PCD by P4 prophage), *orf126* (broad-spectrum counter-defense), *orf143* (IpII; inhibitor of GmrSD), *orf145* (triggers PCD in MG1655), *orf148* (broad-spectrum counter-defense), and *orf184* (triggers PCD by gop-beta-cII).

**Supplementary File 8. Experimental record of follow-up transposon whole-genome knockout screens.** The first sheet contains a detailed record of all follow-up transposon screens designed to phenocopy counter-defense phenotypes of AGs. These screens are performed with RB-TnSeq libraries prepared with naïve ECOR hosts (no AGs present). Each library is challenged by various phages one at a time. The timeline of each experiment indicates library growth characteristics. Empirically measured multiplicity at time of infection, and dilutions used before plating are also recorded. Screens were performed in two stages: first, a pilot with one phage for each RB-TnSeq library, and then a larger experiment with all other phages that showed enhanced infection upon expression of a counter-defense AG. The second sheet lists all hits recovered from the screens.

**Supplementary File 9. Host-AG combinations for affinity-purification/mass-spectrometry.** Matrix of all host-AG combinations where potential binding partners of AGs were affinity-purified from the native host expressing counter-defense AGs, and identified by mass-spectrometry. Experiments were performed in triplicate. *Orf74* produced no phenotypes during AG screening and was used as a control for all experiments in the relevant hosts.

**Supplementary File 10. Accessory regions of AGs selected for further study.** Visualizations of ARs which contain AGs that produced counter-defense or conditional-lethal phenotypes in AG screens.

