## Supplementary File 1 for "Activation of programmed cell death and counter-defense functions of phage accessory genes": accRegionMatrixConstructor.py

```
# Construct matrix of connected genes to find accessory regions exhaustively
# python accRegionMatrixConstructor.py SPFMphages

import sys
dataset = sys.argv[1]
import pickle
clustered = pickle.load(open(dataset+'.clustered.p', 'rb'))
neighbourhood_size = 10 	# Max length of accessory region						
# How many genes to the left and how many genes to the right?

orfdict = {}
for virus in clustered:
	#print virus
	#print clustered[virus]
	for i,orf in enumerate(clustered[virus]):
		genome_len = len(clustered[virus])
		orfname = orf[0]
		cid = orfname.split('__')[0].split('_')[0]
		uid = orfname.split('__')[0].split('_')[-1]
		pro = orf[1]
		start = orf[2]
		stop = orf[3]
		strand = orf[4]
		seq = orf[5]
		gc_bias = orf[6]
		
		# Add virus genome to each gene
		# orfdict has gene cid as keys (the clustered id, not the unique id)
		# orfdict[cid] is a dictionary of genome names
		# orfdict[cid][virus] is a dictionary of gene neighbors' cids
		if cid not in orfdict:
			orfdict[cid] = {}
		
		# Get indexes of gene neighbours 
		# MODULUS OPERATOR ALLOWS YOU TO CIRCLE AROUND THE LIST! EVEN HANDLES NEGATIVE INDEXES CORRECTLY!
		neighbourhood = range(i-neighbourhood_size,i+neighbourhood_size+1)
		neighbourhood_names = [clustered[virus][k % genome_len][0].split('__')[0] for k in neighbourhood] #.split('_')[0]
		#print neighbourhood_cids
		
		# If there is a duplicated gene, it will just overwrite the previous iteration, and this is a hit we'll have to take
		orfdict[cid][virus] = (orfname,{}) # dicts of genes to the left and to the right
		for neighbour,k in zip(neighbourhood_names,neighbourhood):
			orfdict[cid][virus][1][neighbour] = k % genome_len # This will be an index to find the gene in that virus

pickle.dump(orfdict, open(dataset+'_geneNeighbourhoodMatrix.p','wb'))
```
