## Supplementary File 1 for "Activation of programmed cell death and counter-defense functions of phage accessory genes": accRfromMatrix.py

```
# Code to identify accessory regions from gene neighbourhood matrix
# python accRfromMatrix.py SPFMphages

accR_size = 6	# Min number of genomes that must be involved in accR
import sys, pickle
dataset = sys.argv[1]
orfdict = pickle.load(open(dataset+'_geneNeighbourhoodMatrix.p', 'rb'))

# orfdict has gene cid as keys (the clustered id, not the unique id)
# orfdict[cid] is a dictionary of genome names
# orfdict[cid][virus] is a dictionary of gene neighbors' cids with values corresponding to the index of that neighbour
cid_list = sorted(orfdict, key=lambda k: len(orfdict[k]), reverse=False)
too_small = {}
anchor_dict = {}

# Start from the smallest regions first to populate the too_small list
for i,cid1 in enumerate(cid_list):
	if cid1 in too_small:	continue
	gene_penetrance = len(orfdict[cid1])
	if gene_penetrance < accR_size:
		too_small[cid1] = gene_penetrance
		continue
	genome_set1 = set(orfdict[cid1].keys())
	
	for cid2 in cid_list[i+1:]: # This way no need to worry about the i==j test case
		if cid2 in too_small:	continue
		genome_set2 = set(orfdict[cid2].keys())
		common_vir = genome_set1.intersection(genome_set2)
		
		# If the two genes are both present in at least N genomes
		if len(common_vir) >= accR_size:
			barrier = len(common_vir)
			# If the two genes are present in each other's neighborhoods in ALL the genomes
			for vir in common_vir:
				neighbourhood1 = orfdict[cid1][vir][1]
				orfname1 = orfdict[cid1][vir][0].split('__')[0]
				neighbourhood2 = orfdict[cid2][vir][1]
				orfname2 = orfdict[cid2][vir][0].split('__')[0]
				#print vir
				#print orfname1
				#print orfname2
				#print neighbourhood1
				#print neighbourhood2
				
				if (orfname1 in neighbourhood2) and (orfname2 in neighbourhood1):
					barrier = barrier - 1
				else:
					break
					
			if barrier == 0:
				key = tuple(sorted([cid1,cid2]))
				if key in anchor_dict:
					print('Collision of anchor pairs!')
				anchor_dict[key] = common_vir
	#break
print(str(len(anchor_dict))+' accessory regions found')

import pandas as pd
import numpy as np
vir_too_diverse = {}
df = pickle.load(open(dataset+'SimilarityMatrix.p','rb'))
fastaniMatrix = df.replace('',np.nan)
with open(dataset+'_viralDiversityFilter.log', mode='w') as g:
	for entry in anchor_dict:
		virset = anchor_dict[entry]
		subdf = fastaniMatrix.loc[virset,virset]
		if subdf.isnull().values.any():
			vir_too_diverse[entry] = virset
			g.write(str(entry)+' omitted because at least one phage is too diverged\n')
			continue
		if subdf.mean().mean() < float(85):
			vir_too_diverse[entry] = virset
			g.write(str(entry)+' omitted because overall mean ANI is below 85%\n')
			continue
		g.write(str(entry)+' passes viral diversity filter\n')
		#break

new_anchor_dict = {}
for entry in anchor_dict:
	if entry not in vir_too_diverse:
		new_anchor_dict[entry] = anchor_dict[entry]

print(str(len(new_anchor_dict))+' accessory regions passed genome similarity filter')
pickle.dump(new_anchor_dict, open(dataset+'_anchorDict.p','wb'))
```
