## Supplementary File 1 for "Activation of programmed cell death and counter-defense functions of phage accessory genes": accRScorerAndPrinter.py

```
# Score regions of low conservation in viral genome clusters
# python accRScorerAndPrinter.py SPFMphages

print('Importing data...')
import sys, os
dataset = sys.argv[1]

import pickle
original = pickle.load(open(dataset+'.p','rb'))
clustered = pickle.load(open(dataset+'.clustered.p','rb'))
anchor_dict = pickle.load(open(dataset+'_anchorDict.p','rb'))
neighbourhood_dict = pickle.load(open(dataset+'_geneNeighbourhoodMatrix.p','rb'))

def geneCounter(d): #Input is a dictionary of lists
	e = {}
	dg = {}
	for v in d:
		dg[v] = {}
		for exg in d[v][0]:
			# Make a mirror dict with just the gene cluster representatives, this will speed up counting later
			if exg.split('_is_')[0] not in dg[v]:
				dg[v][exg.split('_is_')[0]] = ''
			if exg not in e:
				e[exg] = 0
	# Going virus by virus will not allow duplicated genes to be double counted
	for exg in e:
		for v in dg:
			if exg.split('_is_')[0] in dg[v]:
				e[exg]+=1
	#Output is a dictionary of gene IDs with counts as values			
	return e

print('Finding accessory regions in clusters of viruses...')
# Read in all clusters into a dict of dicts
clusters = {}

regions = {}
id = 0
# Go through the largest phamily first
# Give unique names to each phamily even if the same viruses are present to preserve anchor indexing calculations
# Otherwise a bunch of different a_pairs end up in the same phamily and I can't calculate geneCounts over entire genomes...
# Get all the region information here, but you will have to assemble them first before getting gene counts
for ap in sorted(anchor_dict, key=lambda x: len(anchor_dict[x]), reverse=True):
	id+=1
	vir_list = anchor_dict[ap]
	n = 'phamily'+str(id)
	clusters[n] = {}
	regions[n] = {}
	for virus in vir_list:
		g_list = [x[0].split('__')[0] for x in clustered[virus]]
		anchor1 = neighbourhood_dict[ap[0]][virus][0].split('__')[0]
		anchor2 = neighbourhood_dict[ap[1]][virus][0].split('__')[0]
		i1 = neighbourhood_dict[ap[0]][virus][1][anchor1]
		i2 = neighbourhood_dict[ap[1]][virus][1][anchor2]
		a_list = sorted([i1,i2])
		r = []
		circle = False
		
		# Non-circular region
		lr = list(range(a_list[0]+1,a_list[1]))
		#print lr
		
		# Circular region
		temp = list(range(0,len(g_list)))
		cr = temp[a_list[1]+1:] + temp[:a_list[0]]
		#print cr
		
		if (len(lr)>10) and (len(cr)>10):
			print('Error! '+str(ap)+' has no accessory region smaller than the maximum length in phage '+ virus)
			print(a_list)
			print(lr)
			print(cr)
			print(g_list)
			quit()
		if (len(lr)<10) and (len(cr)>10):
			r = lr
			circle = False
		if (len(lr)>10) and (len(cr)<10):
			r = cr
			circle = True
		if (len(lr)<10) and (len(cr)<10):
			#print('Warning! '+str(ap)+' has both circular and linear accR in '+virus+'. Picking the smaller one')
			r = min(lr,cr)
			if r == cr:
				circle = True
		
		sub_g_list = [g_list[x] for x in r]
			
		pair_gene_counts = {}
		pair_distance = {}
		pair_names = {}
		pair_gene_orfs = {}
		pair_gene_gcs = {}
		pair_indexes = {}
		pair_genes = {}
		if ap not in regions[n]:
			regions[n][ap] = [pair_gene_counts,pair_distance,pair_names,pair_gene_orfs,pair_gene_gcs,pair_indexes,pair_genes]
		
		# If region is not empty, append the pair_distance (genome count is the length of this list)
		# Also record the explicit names of the pair_genes for each virus
		if len(r) > 0:
			#regions[n][a_pair][2]+=1
			regions[n][ap][1][virus] = len(r) #.append(len(r))
			regions[n][ap][2][virus] = (tuple(sorted([ anchor1, anchor2 ])), circle)
			regions[n][ap][5][virus] = r
			
		# This has to be there whether or not the region is empty to avoid later keyerrors
		regions[n][ap][6][virus] = sub_g_list
		
		# Some troubleshooting	
		#print '*'*60
		#print virus
		#print a_list
		#print ap
		#print (tuple(sorted([ anchor1, anchor2 ])))
		#print g_list
		#print r
		#print sub_g_list
		#print circle
		
		clusters[n][virus] = (sub_g_list, g_list)

# Sanity check
for n in regions:
	if len(regions[n]) > 1:
		print('Why are there multiple a_pairs in phamily '+n)
print(str(len(regions))+' accessory regions loaded. Starting the gene counting process...')
		
# Make a dict of in situ gene counts to match the dict of dicts
counts = {}
# Obtain total gene counts (i.e. how many genomes contain each gene) for each cluster
for n in clusters:
	counts[n] = {}
	geneCounts = geneCounter(clusters[n])
	
	# Replace genes with gene counts for each virus in cluster
	for virus in clusters[n]:
		sub_g_list = clusters[n][virus][0]
		c_list = []
		for j,gene in enumerate(sub_g_list):
			c_list.append(geneCounts[gene])
		counts[n][virus] = c_list

# Update gene counts in regions dict
for n in clusters:
	for virus in clusters[n]:
		ap = regions[n].keys()[0]
		#print ap
		c_list = counts[n][virus]
		g_list = regions[n][ap][6][virus]
		
		# Go through the genes within to update the conservation count
		for i,g in enumerate(g_list):
			if g not in regions[n][ap][0]:
				regions[n][ap][0][g] = c_list[i]

def listExtract(l,pn,pd,c): # inclusive list
	out = []
	i1 = l.index(pn[0])
	i2 = l.index(pn[1])
	if i1 > i2:
		dummy = i1
		i1 = i2
		i2 = dummy
	for i,n in enumerate(l):
		if i>i2:	continue
		if i>=i1:	out.append(n)
	
	# Allow for circularization - also check that the region is the right length (either c or a length mismatch will trigger this)
	if (len(out)-2 != pd) or c: # -2 because the extracted list still has the bookends (i.e. list is inclusive)
		out = []
		for i,n in enumerate(l):
			if i<=i1:	out.append(n)
			if i>=i2:	out.append(n)
	return out

print('Formatting results...')

toWrite = {} # This will get sorted by score at the end

allProteins = {}
# Extract the regions out of the .clustered.seq files, use the header to mark the genomes
for cluster in regions:
	toWrite[cluster] = {}
	#if cluster != 'cluster1115':	continue #Debugging mode
	allProteins[cluster] = {}
	for a_pair in regions[cluster]:
		if regions[cluster][a_pair][1] == {}:	continue # These are anchor genes that always appear together
		geneCounts = regions[cluster][a_pair][0] # dict of gene counts in the region
		pairDistances = regions[cluster][a_pair][1] # dict of expected pair distances in each genome
		pairNames = regions[cluster][a_pair][2] # dict of explicit names of anchor genes in each genome
		#pairGeneSeq = regions[cluster][a_pair][3] # (still empty) dict where orfs should be stored
		#pairGeneGCs = regions[cluster][a_pair][4] # (still empty) dict where GC ratios should be stored
		
		alignment = ''
		proteins = ''
		allProteins[cluster][a_pair] = {}
		# Now go through every genome in that cluster and extract region
		for virus in clusters[cluster]:
			#print virus
			#print original[virus]
			genome = clusters[cluster][virus][1]
			#print genome
			subgenomeString = ''
			
			if virus not in pairDistances.keys():
				subgenomeString = ''
				alignment = alignment + ('%90s\t%s\n' % (virus, subgenomeString))
				continue
			
			pd = pairDistances[virus] # Expected length of region
			pn = pairNames[virus][0] # Explicit names of the anchor genes
			c = pairNames[virus][1] # Is this region a circle?
			
			subgenome = listExtract(genome,pn,pd,c)
			
			# Sanity check
			sub_g_list = clusters[cluster][virus][0]
			if set(list(pn)+sub_g_list).symmetric_difference(set(subgenome)) != set([]):
				print('Subgenome and Sub_g_list do not match')
				print(subgenome)
				print(pn[0]+sub_g_list+pn[1])
				
			
			proteins = proteins + '\n' + virus + '\n'
			# Use original dict to get protein sequences here
			realGeneList = [x[0].split('__')[0] for x in original[virus]] # original[virus] is a list of tuples made by: orfs.append((pname,pro,sta,sto,strand))
				
			for exgene in subgenome:
				# Get the protein sequences of the actual genes in the alignment
				realGene = exgene.split('_is_')[1] # This is the uid of the gene that is specific to the virus, not the protein cluster representative
				
				realIndex = realGeneList.index(realGene)
				realSeq = original[virus][realIndex][1]
				realName = original[virus][realIndex][0]
				fakeGene = exgene.split('_is_')[0]
				gc_gene = original[virus][realIndex][-1]
				
				# Add protein sequences to the accessory regions dictionary
				if fakeGene not in a_pair: # Cluster representative will be in a_pair, not the unique gene ID
					regions[cluster][a_pair][3][realGene] = realSeq
					regions[cluster][a_pair][4][realGene] = gc_gene
					
				# Add protein sequences to the overall protein compendium
				#if fakeGene not in a_pair:
				if fakeGene not in allProteins:
					allProteins[cluster][a_pair][fakeGene] = [realSeq, exgene + '__' + realName.split('__')[1], 0]
				# Replace with annotated version if available
				#if len(realSeq) > len(allProteins[cluster][a_pair][fakeGene][0]):
				if 'ypothetical' not in realName.split('desc:')[-1]:	
					allProteins[cluster][a_pair][fakeGene] = [realSeq, exgene + '__' + realName.split('__')[1], 0]
				
				proteins = proteins + '>' + exgene + '__' + realName.split('__')[1] + '\n' + realSeq + '\n'	

				# Format the alignment so that the gene counts are immediately visible in the alignment itself
				if exgene in pn:
					#subgenomeString = subgenomeString+gene+', ' # Don't write the anchors, it's confusing
					continue
				try:
					subgenomeString = subgenomeString+exgene.split('_is_')[0]+':'+str(geneCounts[exgene])+', '
				except KeyError:
					print('\nERROR: A duplicated gene is being elevated to the level of ordinal, or your circularization detector was flummoxed by a rearrangement!')
					print(cluster)
					print(virus)
					print(exgene)
					print(a_pair)
					print(pn)
					print(pd)
					print(subgenome)
					print(genome)
					#print geneCounts
					#print pairNames
					#print pairDistances
			
			alignment = alignment + ('%90s\t%s\n' % (virus, subgenomeString))
			
		toWrite[cluster][a_pair] = cluster + '\nAnchors: ' + str(a_pair).replace("'","") + '\n' + alignment + proteins + '*'*180 + '\n'# #!!!!!!!! 

print('Calculating scores and ranking clusters...')
import numpy as np
# Score regions
scores = {}		

for n in regions:
	for ap in regions[n]:
		if regions[n][ap][1] == {}:	continue # These are anchor genes that always appear together
		
		# Get total number of viruses in each cluster
		cs = len(clusters[n])
		
		# Get Conservation and diversity stats
		conservation_uid = regions[n][ap][0]
		conservation_cid = {}
		for key in conservation_uid:
			cid_key = key.split('_is_')[0]
			if cid_key not in conservation_cid:
				conservation_cid[cid_key] = conservation_uid[key]
		
		# These hurt a region
		pd = regions[n][ap][1].values() # list of all distances in the cluster for given pair
		pc = conservation_cid.values() # list of all gene counts in the cluster for given pair 
		porf = [len(o) for o in regions[n][ap][3].values()] # list of all lengths of orfs in region
		median_pd = np.median(pd) # How long is this region in each virus?
		cv_pd = (np.std(pd)/np.mean(pd))+1 # How varied are the region lengths in each virus?
		mean_pc = np.mean(pc) # How conserved are the genes?
		den = cv_pd*mean_pc#*median_pd*np.median(porf)
		
		# These help a region
		pgc = len(pd) # number of non-empty regions in the cluster for that given pair
		gc = len(pc) # number of genes between this pair in all genomes in the cluster
		pggc = [abs(1-r) for r in regions[n][ap][4].values()] # Magnitude of deviation from 1 i.e. perfect match to GC content of genome
		num_genes = float(gc) # How many unique genes in this region?
		median_gcdev = np.median(pggc)
		cv_pc = (np.std(pc)/np.mean(pc))+1 # How varied are the gene conservation counts?
		genome_frac = float(pgc)/cs # How many genomes share this region (i.e. non-empty)?
		num = num_genes*median_gcdev*cv_pc*genome_frac # Normalize pgc by cluster size to avoid preferring larger clusters by default
		#num = float(gc)*pcc*pgc/cs # This version takes cross-cluster gene spread into account, and severely penalizes all clusters that don't have such genes (score = 0)
		
		score_pieces = ['num_genes',num_genes, 'median_gcdev',median_gcdev, 'cv_conservation',cv_pc, 'genome_frac',genome_frac, 'cv_regionLen',cv_pd, 'mean_conservation',mean_pc]
		# These are anchor genes that have no variation between them
		if all([x==pgc for x in pc]):	
		#	print toWrite[n][ap]
			continue 
		
		scores[(n,ap)] = (num/den, score_pieces)
		
		for fg in allProteins[n][ap]:
			allProteins[n][ap][fg][-1] = num/den

# Write in sorted order using scores
x = 0
content_filter = {}	
with open(dataset+'_accRegions_nr_all.txt', mode='w') as g:
	with open(dataset+'_accProteins_nr_all.txt', mode='w') as h:
		with open(dataset+'_accRegions_Redundant_all.txt', mode='w') as i:
			for region in sorted(scores.keys(), key=lambda k: scores[k][0], reverse=True):
			
				# Subset output if desired --
				#allowed_clusters = {'cluster100':1,'cluster203':1,'cluster303':1,'cluster334':1,'cluster338':1,'cluster437':1,'cluster439':1,'cluster455':1}
				#if region[0] not in allowed_clusters:	continue # Select a particular cluster of interest
				#if scores[region] < 0.1:	continue
				#if 'scherichia' not in toWrite[region[0]][region[1]]:	continue
				
				sc = "{:.3f}".format(scores[region][0])
				sp = str(scores[region][1])
				
				# REDUNDANCY FILTER
				rrr = regions[region[0]][region[1]] #same as regions[n][a_pair] above
				gene_cids = [y.split('_')[0] for y in rrr[0].keys()]
				red_score = 0
				limit = float(0.9) # If 90% of the genes have already been seen, skip
				for cid in gene_cids:
					if cid in content_filter:
						red_score = red_score + 1
				red_score_norm = float(red_score)/len(gene_cids)
				# Once score is calculated, add all encountered genes to redundancy dict to make filter better
				for cid in gene_cids:
					content_filter[cid] = 0
				if red_score_norm > limit:
					i.write(sp+'\n')
					i.write(sc)
					i.write(toWrite[region[0]][region[1]])
					continue
				
				# WRITE MAIN FILES
				g.write(sp+'\n')
				g.write(sc)
				g.write(toWrite[region[0]][region[1]])
			
				for fg in allProteins[region[0]][region[1]]:
					AccName = allProteins[region[0]][region[1]][fg][1]+'__'+sc+str(region[0])
					AccSeq = allProteins[region[0]][region[1]][fg][0]
					h.write('>'+AccName+'\n'+AccSeq+'\n')
				
				x+=1
				
				# LIMIT OUTPUT TO TOP 1000 REGIONS TO CONTROL FILE SIZES IF NEEDED
				#if x >= 1000:
				#	break
			
```
