## Supplementary File 1 for "Activation of programmed cell death and counter-defense functions of phage accessory genes": gbk2dict.py

```
# Code to convert gb files to fasta records
# python gbk2dict.py genbankFile

import sys, os
gbfile = sys.argv[1]
from Bio import SeqIO
from Bio.SeqUtils import GC

d = {}
out = {}
c = 0
gene_id = 0

with open(gbfile, mode='rU') as f:
	
	# Go through all viruses
	for record in SeqIO.parse(f, "genbank"):
		id = record.id+'_'+record.annotations['source'].replace(' ','_').split(',')[0].replace('/','_')
		genome = str(record.seq)
		gc_genome = GC(genome) # float GC content
		hash = str(record.features) # This is just a context-independent listing of features and should only be identical for identical viruses
		#print hash
		c+=1
		# Redundancy filter and master dict
		# Don't filter by seq because the annotations might be different
		if hash not in d:
			d[hash] = id
			#print id
		else:
			print('Duplicate entries found: '+id+' and '+d[hash])
			continue
			
		orfs = []
		for feat in record.features:
			if feat.type == 'CDS':
				sta = str(feat.location.start).strip('.<>')
				sto = str(feat.location.end).strip('.<>')
				seq = genome[int(sta):int(sto)]
				gc_seq = GC(seq)/gc_genome
				#print gc_seq
				try:
					pid = feat.qualifiers['protein_id'][0]
				except KeyError:
					try:
						pid = feat.qualifiers['locus_tag'][0]
					except KeyError:
						pid = ''
				
				try:
					product = feat.qualifiers['product'][0].replace(' ','-').replace(',-','-')
				except KeyError:
					product = ''
				
				try:
					pro = feat.qualifiers['translation'][0]
				except KeyError:
					try:
						pseudo = feat.qualifiers['pseudo']
						continue  # Pseudogenes are skipped. If an exception occurs, this "continue" will be skipped and the gene will be written
					except KeyError:
						print('CDS without translation that is not a pseudogene. Empty translation record written.')
						print(feat)
						pro = ''
				
				#print seq
				#print pro
				gene_id+=1
				pname = str(gene_id)+'__'+pid+'_in_'+id+'_at_'+sta+'..'+sto+'_desc:'+product
				orfs.append((pname.replace(' ',''),pro,sta,sto,feat.strand,seq,gc_seq))
		
		if len(orfs) == 0:	continue	
		if id in out:
			print('Collision in virus names! '+id)	
		out[id] = orfs # already in sorted order in gb file
		
		#break # Stop after the first record

print(str(c)+' records processed')
print(str(len(out))+' unique records imported')

# Pickle dict
import pickle
pickle.dump(out, open(gbfile.split('.')[0]+'.p','wb'))

with open(gbfile.split('.')[0]+'.faa', mode='w') as g:
	for virus in out:
		#print(out[k])
		for orf in out[virus]:
			g.write('>'+orf[0]+'\n'+orf[1]+'\n')
		#break # This break writes only one genome for debugging purposes

with open(gbfile.split('.')[0]+'.seq', mode='w') as h:
	for virus in out:
		h.write('>'+virus+'\n')		
		for orf in out[virus]:
			h.write(orf[0].split('__')[0]+',')
		h.write('\n')
```
