## Supplementary File 1 for "Activation of programmed cell death and counter-defense functions of phage accessory genes": gbkPrinterByCluster.py

```
# Code to print individual genbank files of viruses of interest
# python 2020_03_23_gbkPrinterByCluster.py matrixcoli anchors-(eg "(148535,150715)")
# No space in anchors!!

import sys, os, pickle
dataset = sys.argv[1]
gbfile = dataset+'.gb'
clusterfile = pickle.load(open(dataset+'_anchorDict.p', 'rb'))
from Bio import SeqIO

anchors = sys.argv[2]
(a1,a2)= anchors.strip('()').split(',')
akey = tuple(sorted([a1.strip(),a2.strip()]))
viruses = clusterfile[akey]
v_dict = {i:0 for i in viruses}
	
'''
with open(clusterfile, mode='rU') as c:
	viruses = c.readline().strip().split('##HEADER: ')[1].split(',')
v_dict = {i:0 for i in viruses}
'''

redundant = {}
lastName = ''
with open(gbfile, mode='rU') as f:	
	#with open('output_gbk/runBlastForACT.sh', mode='w') as j:	
	# Go through all viruses
	for record in SeqIO.parse(f, "genbank"):
		id = record.id+'_'+record.annotations['source'].replace(' ','_').split(',')[0].replace('/','_')
		seq = str(record.seq)
		#hash = str(record.features) # This is just a context-independent listing of features and should only be identical for identical viruses
		
		if id in v_dict:
		#if 'eudomonas' in id:
			if seq not in redundant:
				redundant[seq] = id
				name = str(id).replace('/','-').replace(':','')
				print(name)
				'''
				j.write('makeblastdb -in '+name+'.fasta'+' -out '+name.split('.')[0]+' -dbtype nucl\n')
				if lastName != '':
					j.write('blastn -query '+name+'.fasta'+' -db '+lastName.split('.')[0]+' -evalue 1 -task megablast -outfmt 6 > '+name+'_vs_'+lastName.split('.')[0]+'.crunch\n')
				lastName = str(id).replace('/','-').replace(':','')
				'''
					
				with open('output_gbk/'+name+'.gbk', mode='w') as g:
					SeqIO.write(record, g, "genbank")
				#with open('output_gbk/'+name+'.fasta', mode='w') as h:
				#	SeqIO.write(record, h, "fasta")
				
```
