## Supplementary File 1 for "Activation of programmed cell death and counter-defense functions of phage accessory genes": gbSubSelector.py

```
# Code to split genbank files according to coordinates
# python 2020_04_28_gbSubSelector.py dir-with-gb accRegionFile accRegionName(eg.0.75cluster393)

import sys, os
gbdir = sys.argv[1]
accFile = sys.argv[2]
accName = sys.argv[3]

phageDict = {}
# Import phage names of interest
for a,b,f in os.walk(gbdir):
	phageDict = {i.split('.gb')[0]:0 for i in f}
#print phageDict

# Figure out acc region coordinates for each phage
began = False
finished = False
currentPhage = 'Dummy Variable - you should never see this'
a1 = 'Dummy Variable - you should never see this'
a2 = 'Dummy Variable - you should never see this'
coords = {}
accLen = {}
deduplicated =  False
with open(accFile, mode='rU') as a:
	for line in a:
		if line.strip() == accName:
			began = True
		if began and not finished:
			#print line
			if 'Anchors: ' in line:
				(a1,a2)= line.strip().strip('Anchors: ').strip('()').split(',')
				a1 = '>'+a1.strip()+'_'
				a2 = '>'+a2.strip()+'_'
				#print(a1,a2)
			if line.strip().replace('/','-').replace(':','') in phageDict:
				currentPhage = line.strip().replace('/','-').replace(':','')
				deduplicated = True
				if currentPhage not in coords:
					coords[currentPhage] = [0,0]
					accLen[currentPhage] = 0
				else:
					print('Phage collisions in coordinate dictionary! Does your phage cluster have duplicate entries?')
				#print(currentPhage)
			if a1 in line and deduplicated:
				c1 = int(line.split('..')[0].split('_')[-1])
				c2 = int(line.split('..')[1].split('_')[0])
			if a2 in line and deduplicated:
				c3 = int(line.split('..')[0].split('_')[-1])
				c4 = int(line.split('..')[1].split('_')[0])
			if ('c1' in locals()) and ('c2' in locals()) and ('c3' in locals()) and ('c4' in locals()) and deduplicated:
				#print line.strip()
				coords[currentPhage][0] = min(c1,c2,c3,c4)
				coords[currentPhage][1] = max(c1,c2,c3,c4)
				accLen[currentPhage] = coords[currentPhage][1]-coords[currentPhage][0]
				deduplicated = False # Don't use the same coordinates more than once
				#print coords
				#print '\n'
				del c1, c2, c3, c4, currentPhage
			if '**********' in line:
				finished = True
				#print coords	
					
	if not began:
		print('Accessory region not found. Are you sure you supplied the correct name?')

# Can't handle circles
from numpy import median
lenMed = median(accLen.values())
excludeCirc = {}
for phage in accLen:
	if accLen[phage] > 3*lenMed:
		excludeCirc[phage] = accLen[phage]
#print excludeCirc

accRename = accName.replace('.','-')
from Bio import SeqIO
# Splice genbank files
for a,b,f in os.walk(gbdir):
	for fname in f:
		if 'accR' in fname:	continue
		if '._' in fname: continue
		if fname.split('.gb')[0] in excludeCirc:
			print('Omitting circular accRegion in '+fname)
			continue
		for record in SeqIO.parse(a+fname, "genbank"):
			seq = record.seq
			feat = record.features
			#print(len(feat))
		with open(a+fname, mode='rU') as i:
			with open(a+fname.split('.gb')[0]+'.'+accRename+'.accR.gb', mode='w') as j:
				id = fname.split('.gb')[0]
				start = coords[id][0]
				end = coords[id][1]
				regionSeq = str(seq)[start:end]
				firstFeat = ''
				firstDone = False
				lastFeat  = ''
				lastDone  = False
				if coords[id] == [0,0]:
					print('Omitting empty accRegion in '+fname)
					continue
				print(id)
				print(coords[id])
				
				
				#print regionSeq
				for feature in feat:
					#print feature
					if feature.type == 'CDS':
						fsta = str(feature.location.start).strip('.<>')
						fsto = str(feature.location.end).strip('.<>')
						if (int(fsta) == int(start)) and not firstDone:
							firstFeat = str(int(fsta)+1)+'..'+fsto #This should be the first feature
							firstDone = True
							#print fsta
							#print start
							#print firstFeat
						if (int(fsto) > int(end)) and firstDone and not lastDone: # 07/09/20
							lastFeat =  str(int(fsta)+1)+'..'+fsto #This should be last+1
							lastDone = True
							#print fsto
							#print end
							#print lastFeat
							break
				
				writing = True
				tot_len = str(len(seq))
				new_len = str(len(regionSeq))
				for line in i:
					#if 'CDS' == line.strip().split()[0]:
					#	print line.strip().split()
					
					if ('..' in line) and ('=' not in line) and (len(line.strip().split()) > 1): # 07/09/20
						#print line
						#print len(line.strip().split())				
						if ('..' in line) and (firstFeat in line.strip().split()[1]): #'CDS' == line.strip().split()[0]
							#print line
							writing = True
						if ('..' in line) and (lastFeat in line.strip().split()[1]): #'CDS' == line.strip().split()[0]
							#print line
							writing = False
						
					# Correct the coordinates
					if ('..' in line) and writing and ('=' not in line) and (len(line.strip().split()) > 1):  # 07/09/20
						coo = line.strip().split()[-1].split('..')
						#print coo
						correction = start
						old_fsta = int(coo[0].split('(')[-1].strip('.<>,')) # 07/09/20
						old_fsto = int(coo[-1].split(')')[0].strip('.<>,')) # 07/09/20
						new_fsta = old_fsta - correction
						new_fsto = old_fsto - correction
						#print(str(seq)[old_fsta:old_fsto]+'\n')
						#print(regionSeq[new_fsta:new_fsto]+'\n')
						line = line.replace(str(old_fsta),str(new_fsta)).replace(str(old_fsto),str(new_fsto))
						if line.count('..') > 1: # 07/09/20
							line = '..'.join([line.split('..')[0], line.split('..')[-1]]).replace('join(','').replace('))',')') # 07/09/20
							
					if writing:
						# Correct total length
						if 'LOCUS' == line.strip().split()[0]:
							line = line.replace(tot_len,new_len)
						if 'REFERENCE' == line.strip().split()[0]:
							line = line.replace(tot_len,new_len)
						j.write(line)
					
					if 'FEATURES' == line.strip().split()[0]:
						writing = False
						
				#  Write the sequence
				toWrite = 'ORIGIN\n'
				for k in range(1,len(regionSeq),60):
					#print k
					subseqString = str(regionSeq[k-1:k+60-1])
					subseqString = ' '.join([subseqString[i:i+10] for i in range(0, len(subseqString), 10)])
					toWrite = toWrite + ('%9s %s\n' % (str(k), subseqString))
				j.write(toWrite+'//\n')
				
				
				
```
