## Supplementary File 1 for "Activation of programmed cell death and counter-defense functions of phage accessory genes": makeFastANI.sh

```
mkdir fastANImatrix
mv fastANI fastANImatrix/
mv splitGenbanksIntoFastaFiles.py fastANImatrix/
mv parseFastANIresults.py fastANImatrix/
cd fastANImatrix/
python splitGenbanksIntoFastaFiles.py ../SPFMphages.gb 
ls *.fa | sort > SPFMphagesANIlist.txt
./fastANI --ql SPFMphagesANIlist.txt --rl SPFMphagesANIlist.txt -o SPFMphagesANIout.txt
python parseFastANIresults.py SPFMphagesANIout.txt
mv SPFMphagesSimilarityMatrix.p ../
mkdir phageFasta/
mv *.fa phageFasta/
cd ..
```
