## Supplementary File 1 for "Activation of programmed cell death and counter-defense functions of phage accessory genes": mmseqsCluster.sh

```
mmseqs createdb SPFMphages.faa SPFMphages_CDS_mmdb  # Tested with and without shuffling and it made almost no difference to number of clusters produced
nice mmseqs cluster SPFMphages_CDS_mmdb SPFMphages_CDS_clustered temp --threads 100
mmseqs createtsv SPFMphages_CDS_mmdb SPFMphages_CDS_mmdb SPFMphages_CDS_clustered SPFMphages_CDS_clustered.tsv --full-header
```
