## Supplementary File 1 for "Activation of programmed cell death and counter-defense functions of phage accessory genes": parseFastANIresults.py

```
# Code to parse fastANI results into a pandas dataframe
# python 2020_09_12_parseFastANIresults.py matrixcoliANIout.txt

import sys
infile = sys.argv[1]
name = infile.strip().split('ANIout.txt')[0]
d = {}

with open(infile, mode='rU') as f:
	for line in f:
		a,b,c,e,g = line.strip().split()
		phageA = str(a).split('/')[-1].split('.fa')[0]
		phageB = str(b).split('/')[-1].split('.fa')[0]
		score = c
		if phageA not in d:
			d[phageA] = {}
		if phageB not in d[phageA]:
			d[phageA][phageB] = float(score)
		else:	print('Duplicates! '+a+' and '+b)

import pandas as pd
df = pd.DataFrame(d)		
df.to_csv(name+'SimilarityMatrix.tsv', sep='\t')
import pickle
pickle.dump(df, open(name+'SimilarityMatrix.p','wb'))
```
