## Supplementary File 1 for "Activation of programmed cell death and counter-defense functions of phage accessory genes": Readme.txt

This code will find accessory regions from the Salmonella phages in SPFMphages.gb

You must have MMseqs2 installed before running this script (see References)

Run the pipeline by entering the following code on the Unix command prompt sh RunMe.sh

This will execute all the steps for finding accessory regions in SPFMphages.gb in order

Non-redundant accessory regions will be in file: SPFMphages_accRegions_nr_all.txt

The last program in RunMe.sh (subsetAccRegionMatrix.sh) subsets an individual accessory
region and extracts it from the various genome genbank files

This program can be modified and used to extract any accessory region in
SPFMphages_accRegions_nr_all.txt, like so --

sh subsetAccRegionMatrix.sh "(3058,5668)" 1.172phamily2571

By default, RunMe.sh automatically extracts the accessory region 1.172phamily2571 between
anchors 3058,5668. This generates the accessory region in Figure 1

You only need to run RunMe.sh once. To extract other accessory regions, run only the
subsetAccRegionMatrix.sh program again with the phamily name and the anchors substituted
appropriately with the desired accessory region from SPFMphages_accRegions_nr_all.txt

To visualize the accessory regions, upload the extracted genbank files into Clinker
https://github.com/gamcil/clinker
