## Supplementary File 1 for "Activation of programmed cell death and counter-defense functions of phage accessory genes": replaceGeneNamesByCluster.py

```
# Read clustering info and replace orf IDs with cluster IDs
# python replaceGeneNamesByCluster.py out_dict

import sys
genomes = sys.argv[1]
import pickle
out = pickle.load(open(genomes,'rb'))

# AFTER CLUSTERING
locusDict = {}
with open(genomes.split('.')[0]+'_CDS_clustered.tsv', mode='rU') as f: #HARDCODED NAME BE CAREFUL
	for line in f:
		#print line.split()
		locus1 = line.split()[0].split('__')[0].strip('"')
		locus2 = line.split()[-1].split('__')[0].strip('"')
		if locus2 not in locusDict:
			locusDict[locus2] = locus1

with open(genomes.split('.')[0]+'.explicitlyClustered.seq', mode='w') as j:
	for virus in out:
		j.write('>'+virus+'\n')
		for orf in out[virus]:
			l = locusDict[orf[0].split('__')[0]] + '_is_' + orf[0].split('__')[0]
			j.write(l+',')
		j.write('\n')

with open(genomes.split('.')[0]+'.clustered.seq', mode='w') as j:
	for virus in out:
		j.write('>'+virus+'\n')
		for orf in out[virus]:
			l = locusDict[orf[0].split('__')[0]]
			j.write(l+',')
		j.write('\n')

# Replace dict values
clustered = {}
for virus in out:
	clustered[virus] = []
	for orf in out[virus]:
		pname = orf[0].split('__')
		old_pid = pname[0]
		new_pid = locusDict[old_pid]
		new_pname = new_pid+'_is_'+orf[0]
		clustered[virus].append((new_pname,orf[1],orf[2],orf[3],orf[4],orf[5],orf[6]))
		
# Pickle dict
import pickle
pickle.dump(clustered, open(genomes.split('.')[0]+'.clustered.p','wb')) # Change names depending on clustering
```
