## Supplementary File 1 for "Activation of programmed cell death and counter-defense functions of phage accessory genes": RunMe.sh

```
#1) Parse in the genbank file of phage genomes
python gbk2dict.py SPFMphages.gb > gbProcessing.log

#2) Cluster proteins in .faa file -- MUST HAVE MMSEQS2 INSTALLED
sh mmseqsCluster.sh > mmseqs.log

#2.5) Make folders to stash away mmseqs output and recover .tsv file
sh mmSeqsCleanup.sh

#3) Replace names of genes in .seq file with names of cluster representatives
python replaceGeneNamesByCluster.py SPFMphages.p

#4) Construct gene matrix to identify accessory regions exhaustively (NEIGHBOURHOOD SIZE MAGIC CONSTANT 10)
python accRegionMatrixConstructor.py SPFMphages

#4.5) Construct fastANI phage genome similarity matrix
sh makeFastANI.sh

#5) Identify accessory regions (PHAGE FAMILY SIZE MAGIC CONSTANT 6)(VIRAL DIVERSITY CUTOFF MAGIC CONSTANT 85%)
python accRfromMatrix.py SPFMphages

#6) Score and print accessory regions (ACC-R REDUNDANCY CUTOFF MAGIC CONSTANT 90%)
python accRScorerAndPrinter.py SPFMphages

#7) Run shell script for plotting -- NO SPACE BETWEEN ANCHORS IN QUOTES!!
mkdir output_gbk
sh subsetAccRegionMatrix.sh "(3058,5668)" 1.172phamily2571

# RUN THIS SCRIPT ONLY ONCE
# USE THE LAST COMMAND WITH VARIOUS PHAMILIES TO GENERATE SUB-GBK FILES FOR VARIOUS ACC-REGIONS OF INTEREST
```
