## Supplementary File 1 for "Activation of programmed cell death and counter-defense functions of phage accessory genes": splitGenbanksIntoFastaFiles.py

```
# Code to take a large fasta file and break it into individual files by name
# python 2020_09_12_splitFastasIntoFiles.py input.fna
    
import sys
infile = sys.argv[1]

from Bio import SeqIO

for record in SeqIO.parse(infile, "genbank"):
    id = str(record.id+'_'+record.annotations['source'].replace(' ','_').split(',')[0].replace('/','_'))
    print(id)
    seq = str(record.seq)
    with open(id+'.fa', mode='w') as f:
    	f.write('>'+id+'\n')
    	f.write(seq+'\n')
    	
```
