## Supplementary File 1 for "Activation of programmed cell death and counter-defense functions of phage accessory genes": subsetAccRegionMatrix.sh

```
# Run with "(125120,36701)" as $1 and 4.319phamily22448 as $2
# eg. sh subsetAccRegionMatrix.sh "(125120,36701)" 4.319phamily22448
python gbkPrinterByCluster.py SPFMphages $1
python gbSubSelector.py output_gbk/ SPFMphages_accRegions_nr_all.txt $2
mkdir $2
cd output_gbk/
mv *accR.gb ../$2/
rm *.gbk
cd ..
```
