## Supplementary File 2 for "Activation of programmed cell death and counter-defense functions of phage accessory genes"

### Slide 1
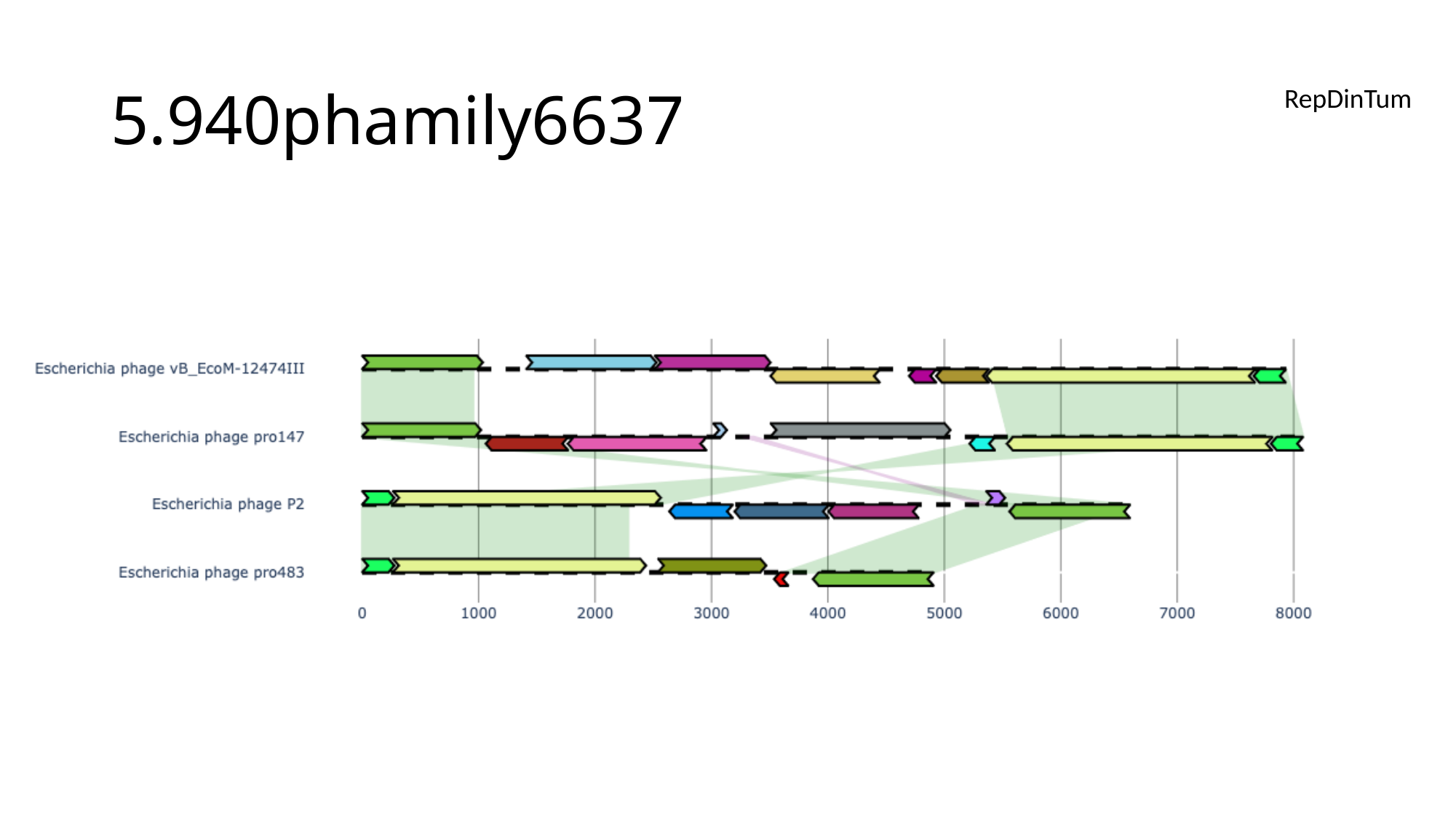

#### 5.940phamily6637
RepDinTum

### Slide 2
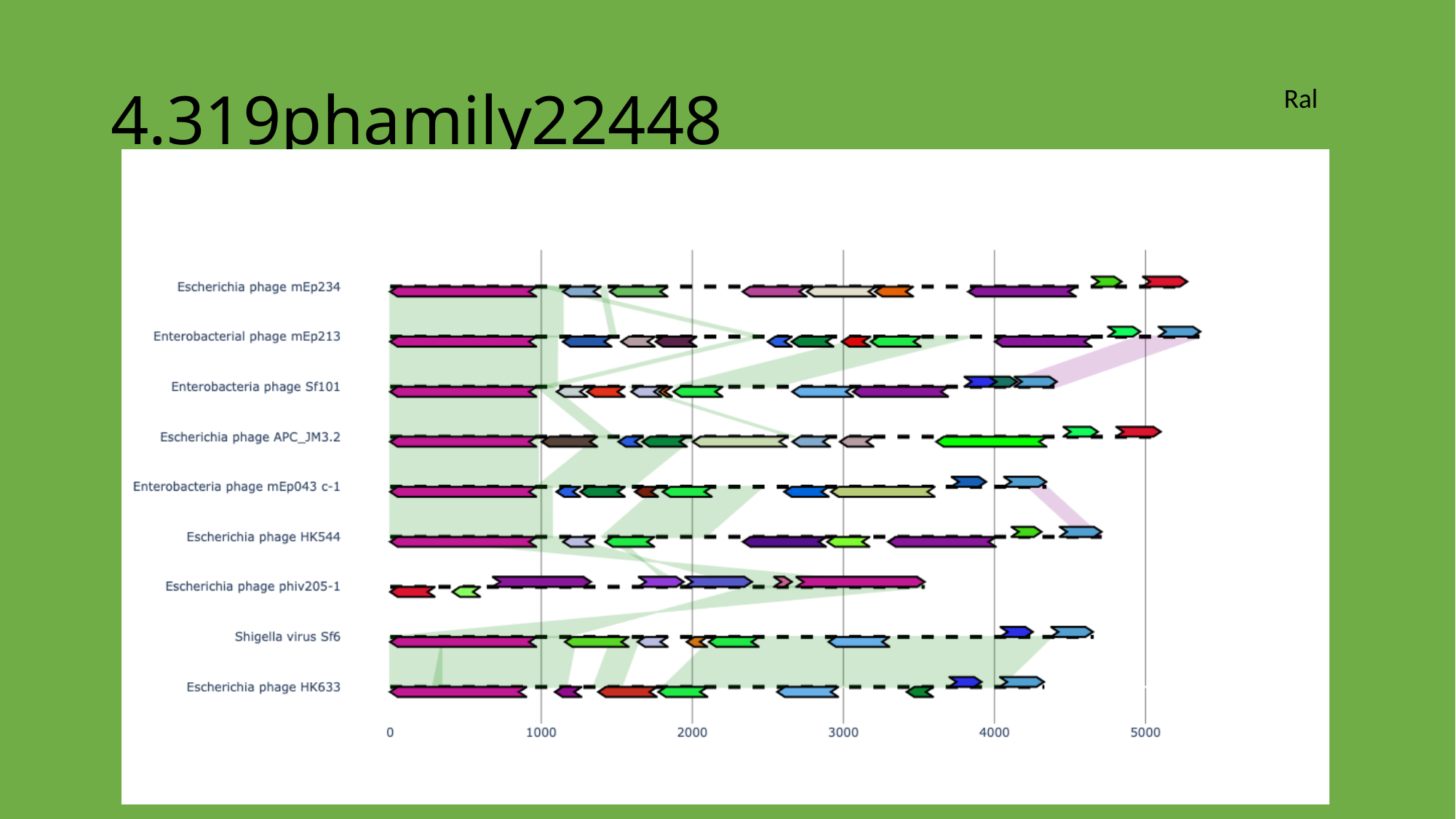

#### 4.319phamily22448
Ral

### Slide 3
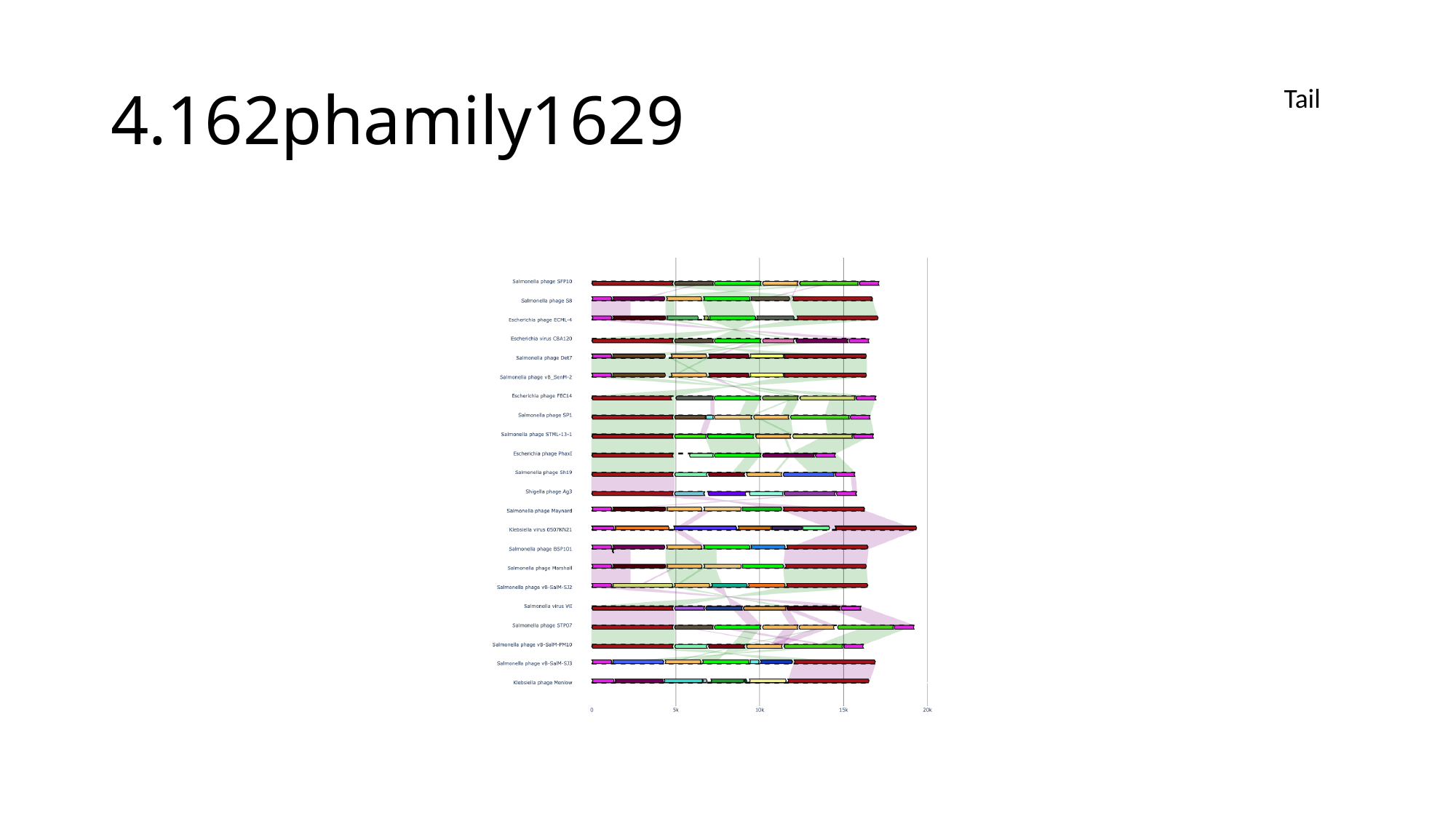

#### 4.162phamily1629
Tail

### Slide 4
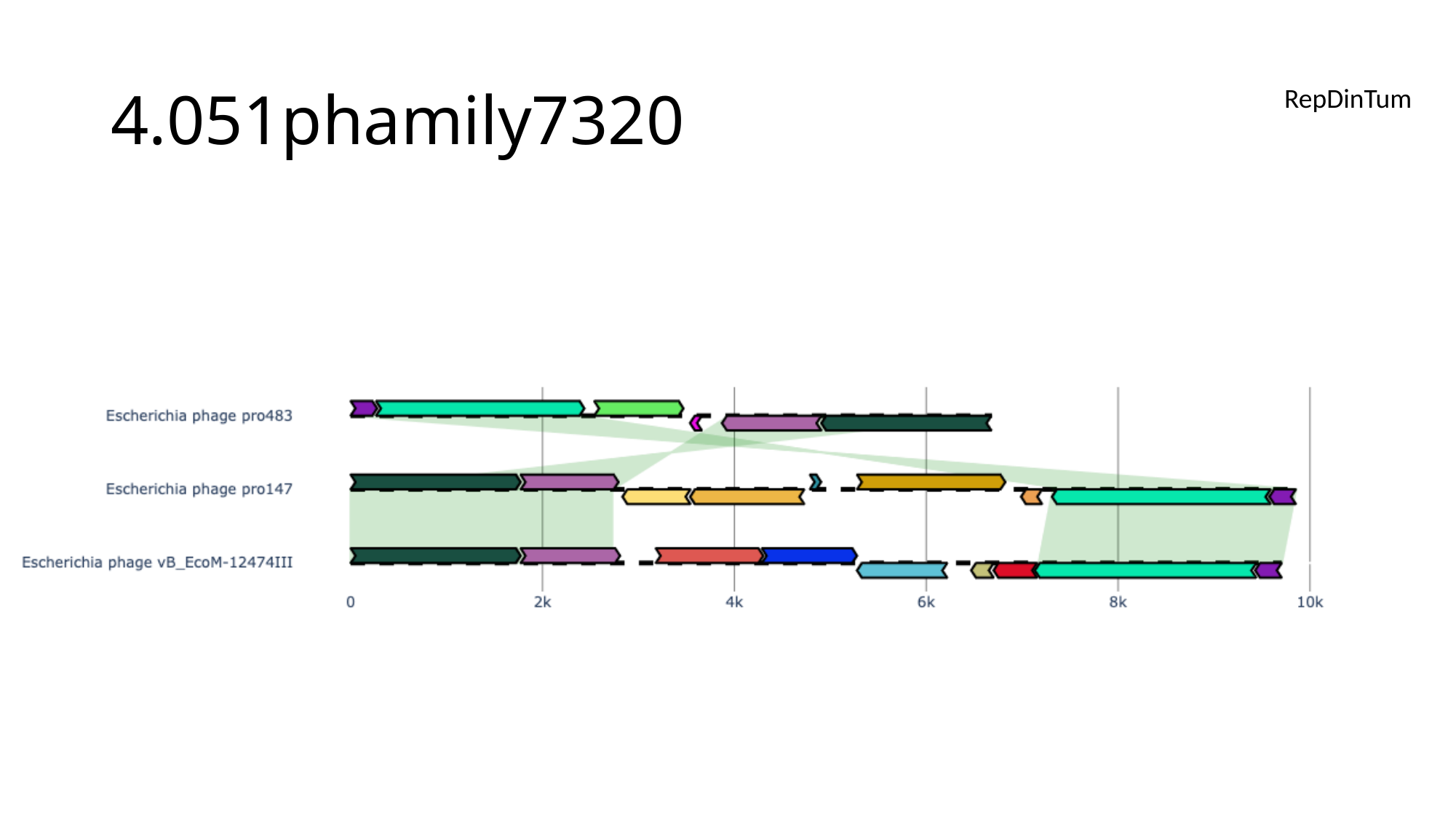

#### 4.051phamily7320
RepDinTum

### Slide 5
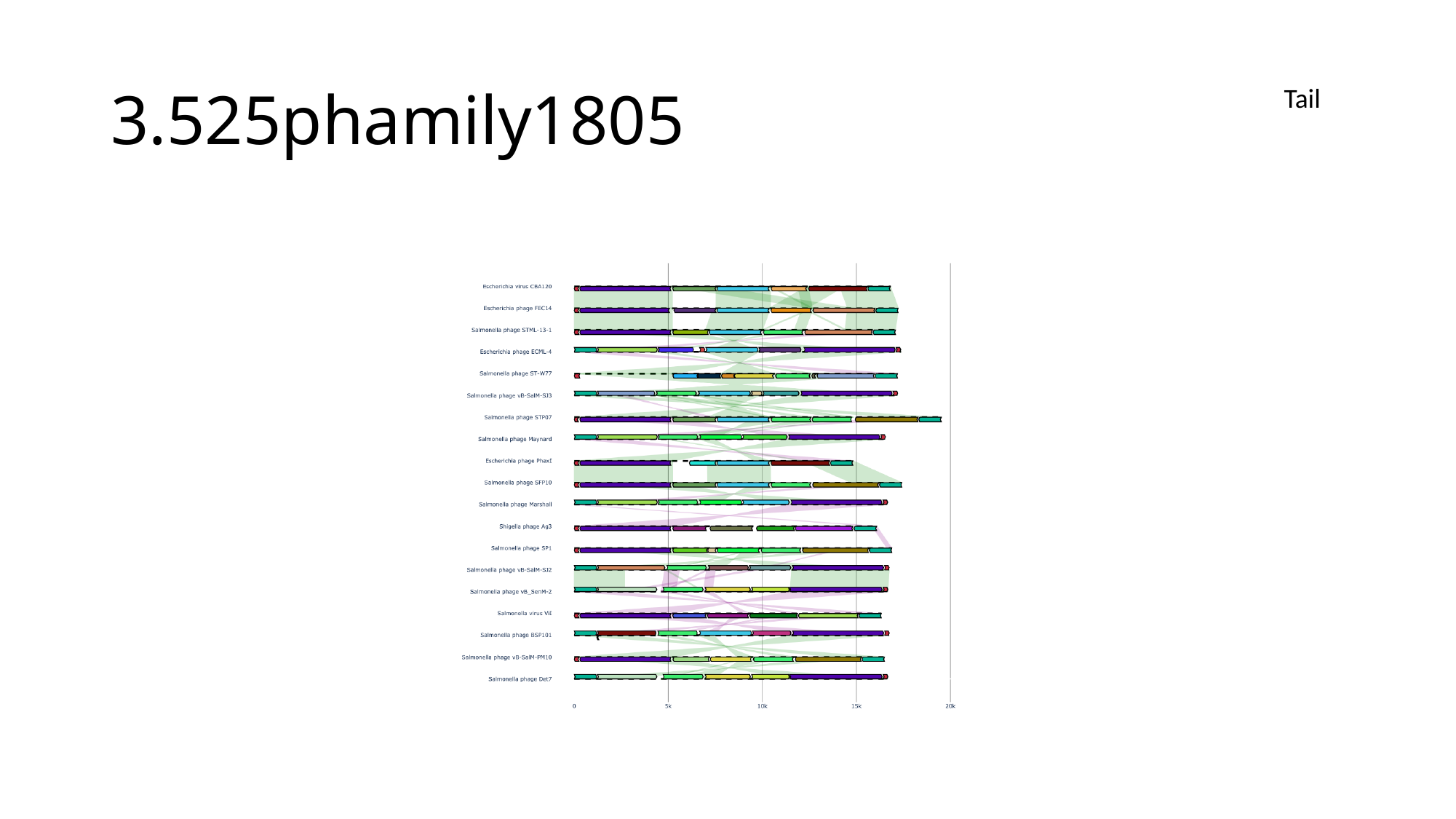

#### 3.525phamily1805
Tail

### Slide 6
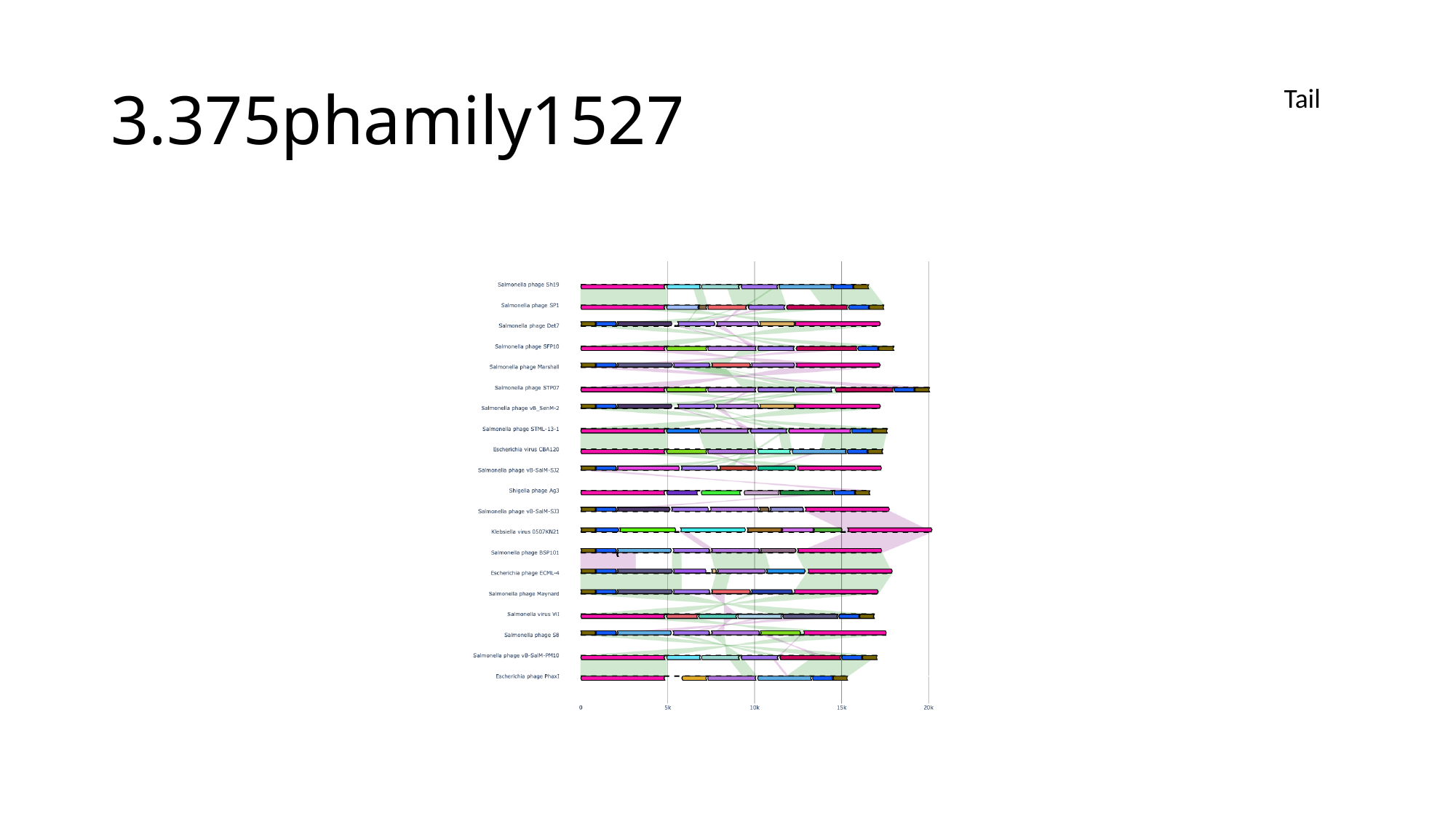

#### 3.375phamily1527
Tail

### Slide 7
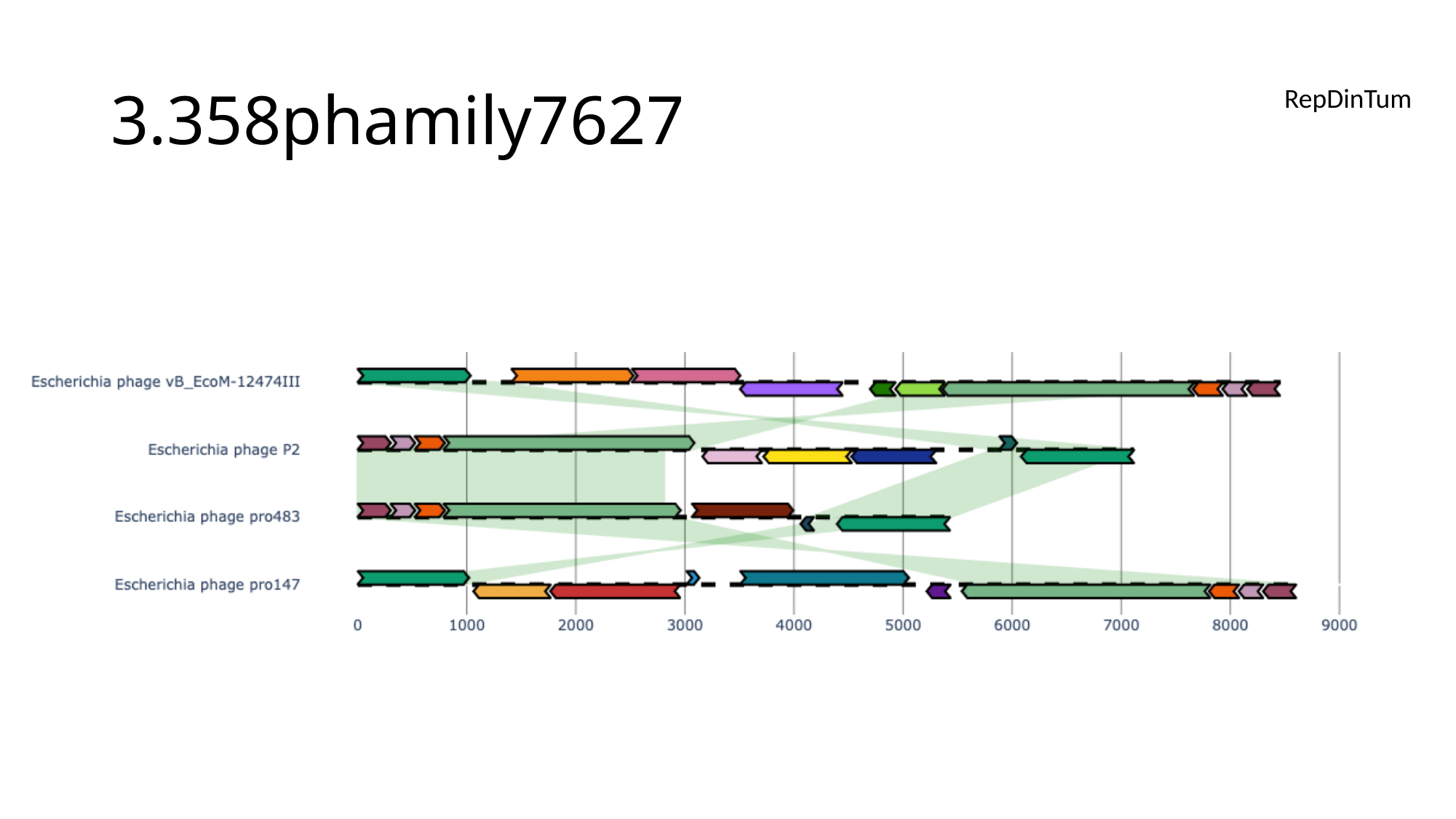

#### 3.358phamily7627
RepDinTum

### Slide 8
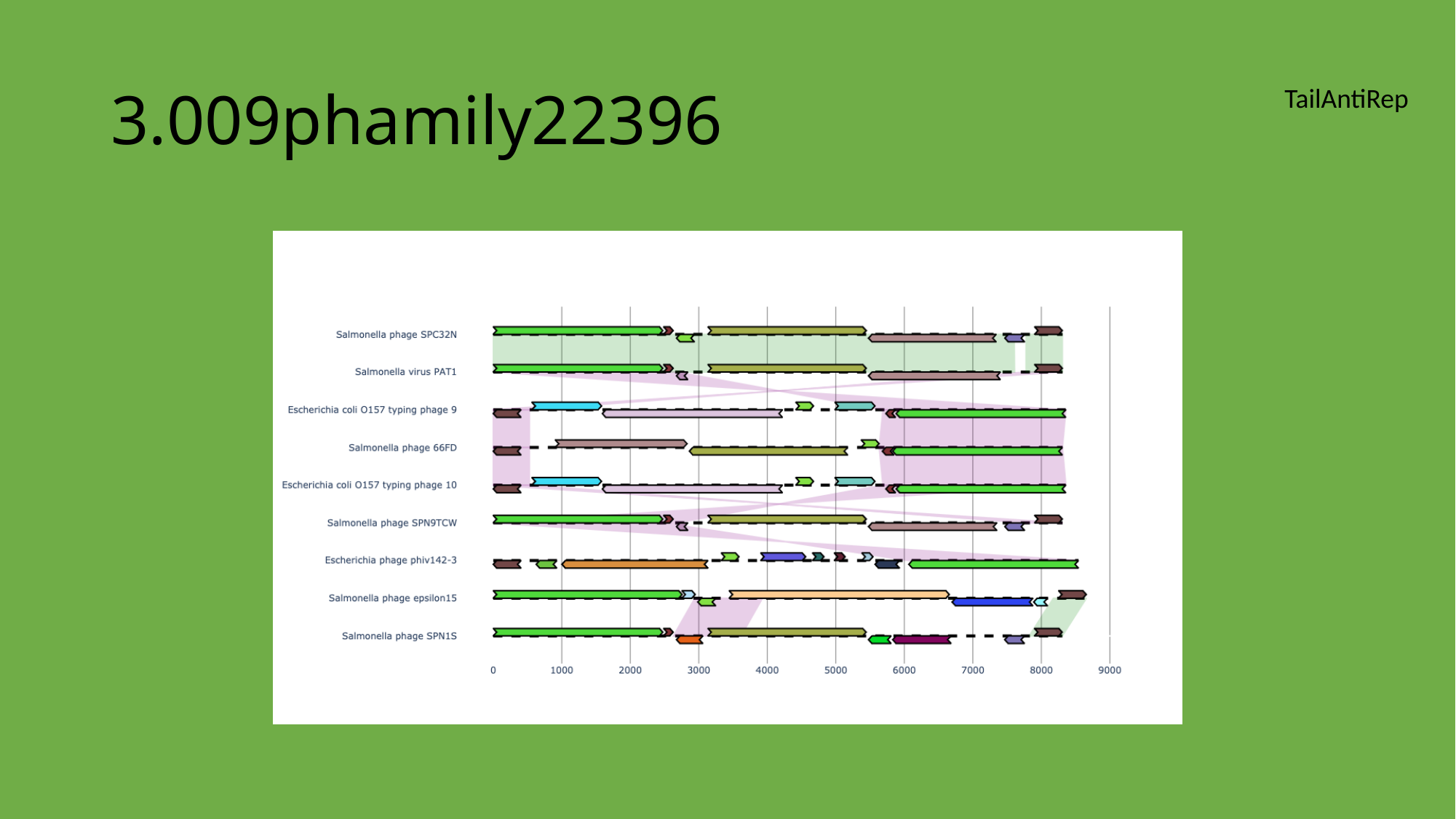

#### 3.009phamily22396
TailAntiRep

### Slide 9
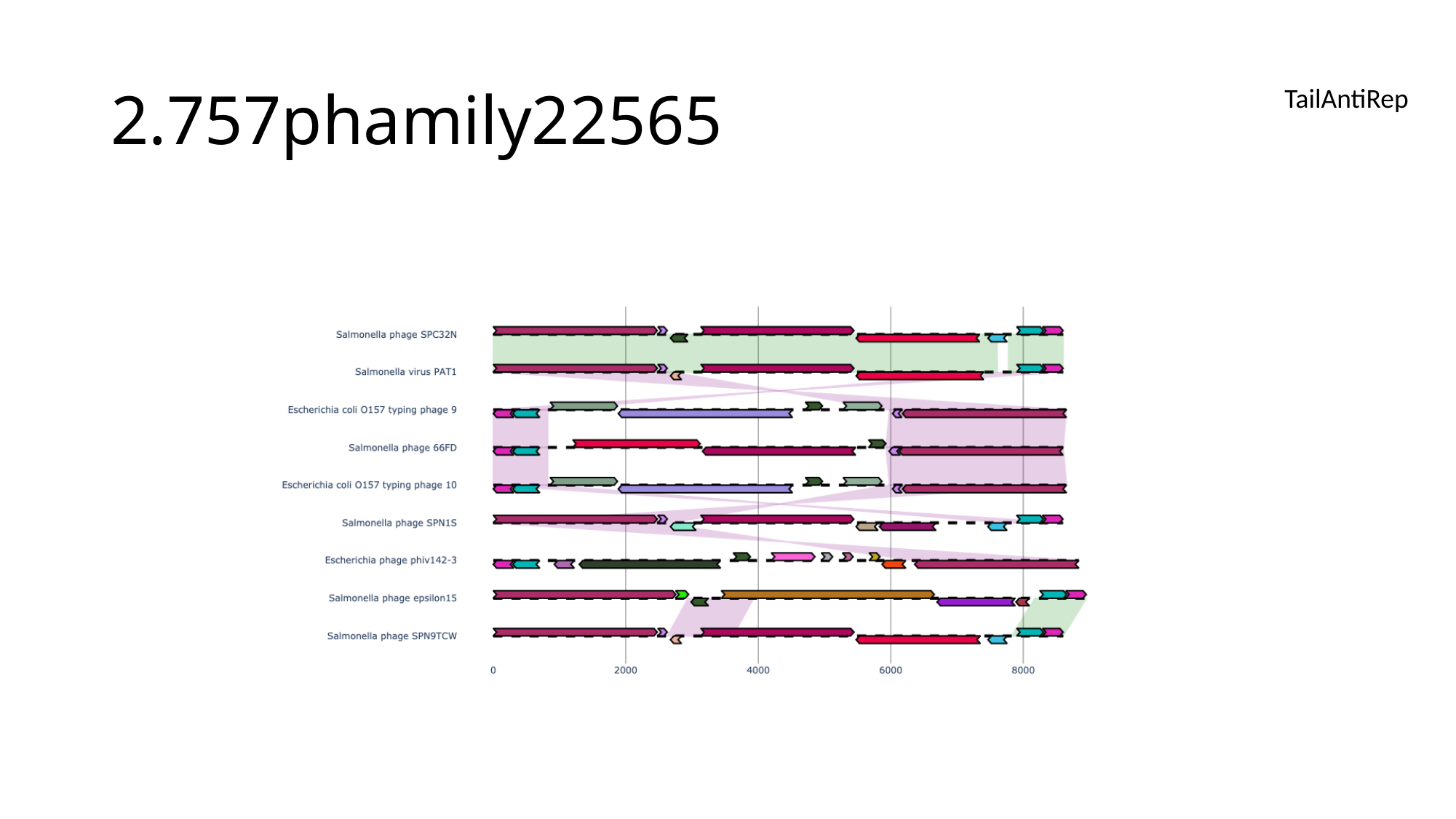

#### 2.757phamily22565
TailAntiRep

### Slide 10
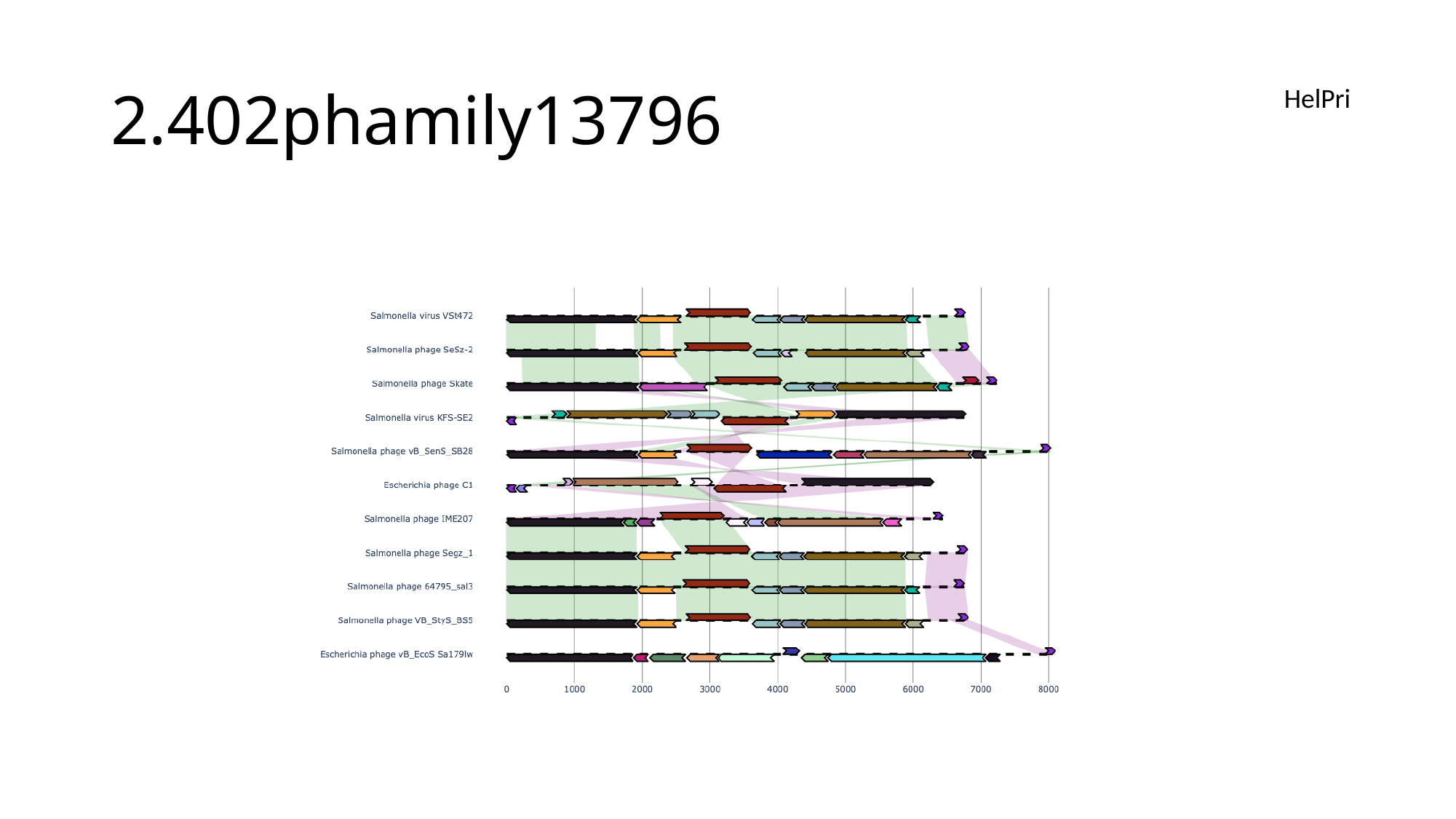

#### 2.402phamily13796
HelPri

### Slide 11
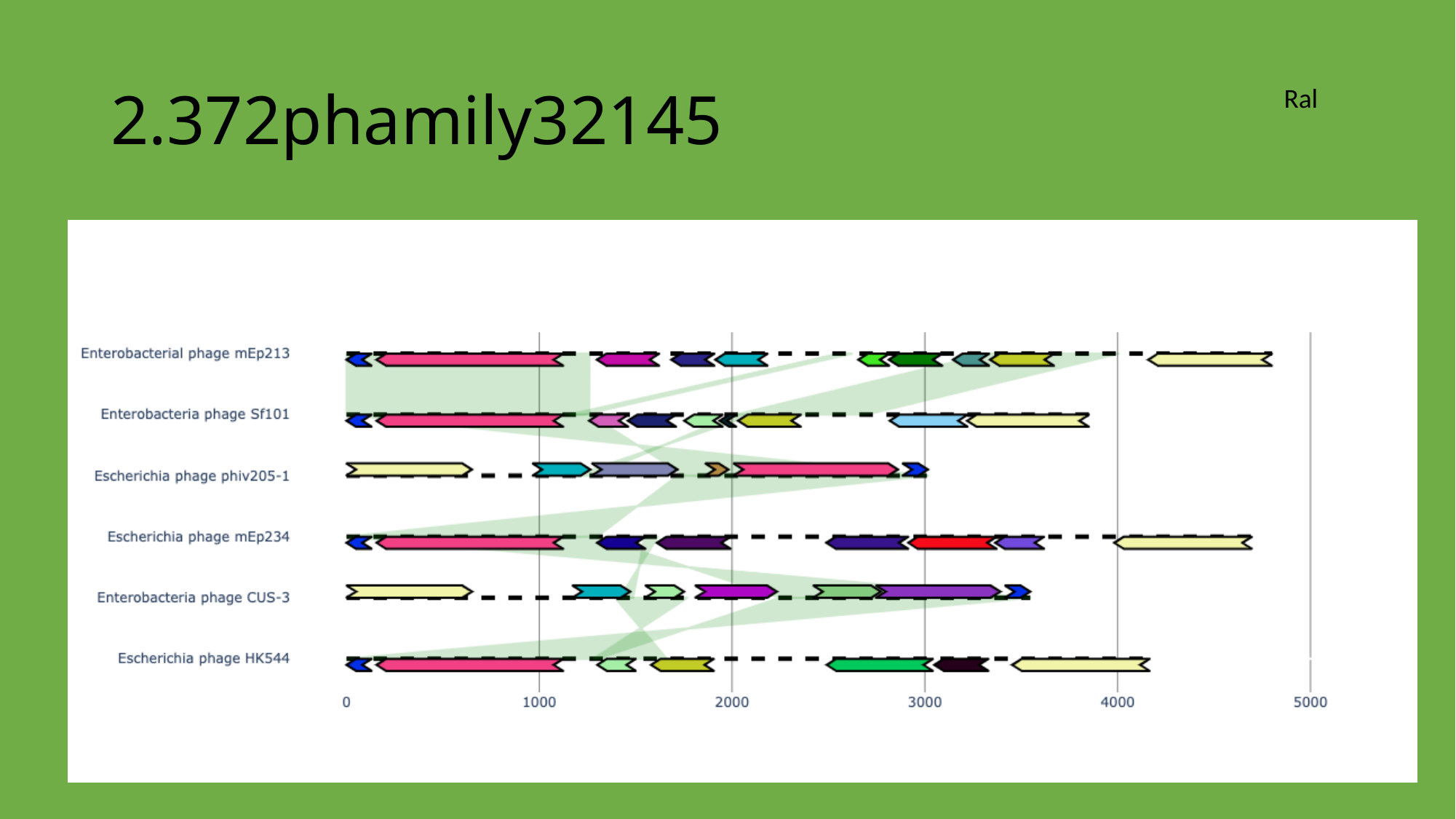

#### 2.372phamily32145
Ral

### Slide 12
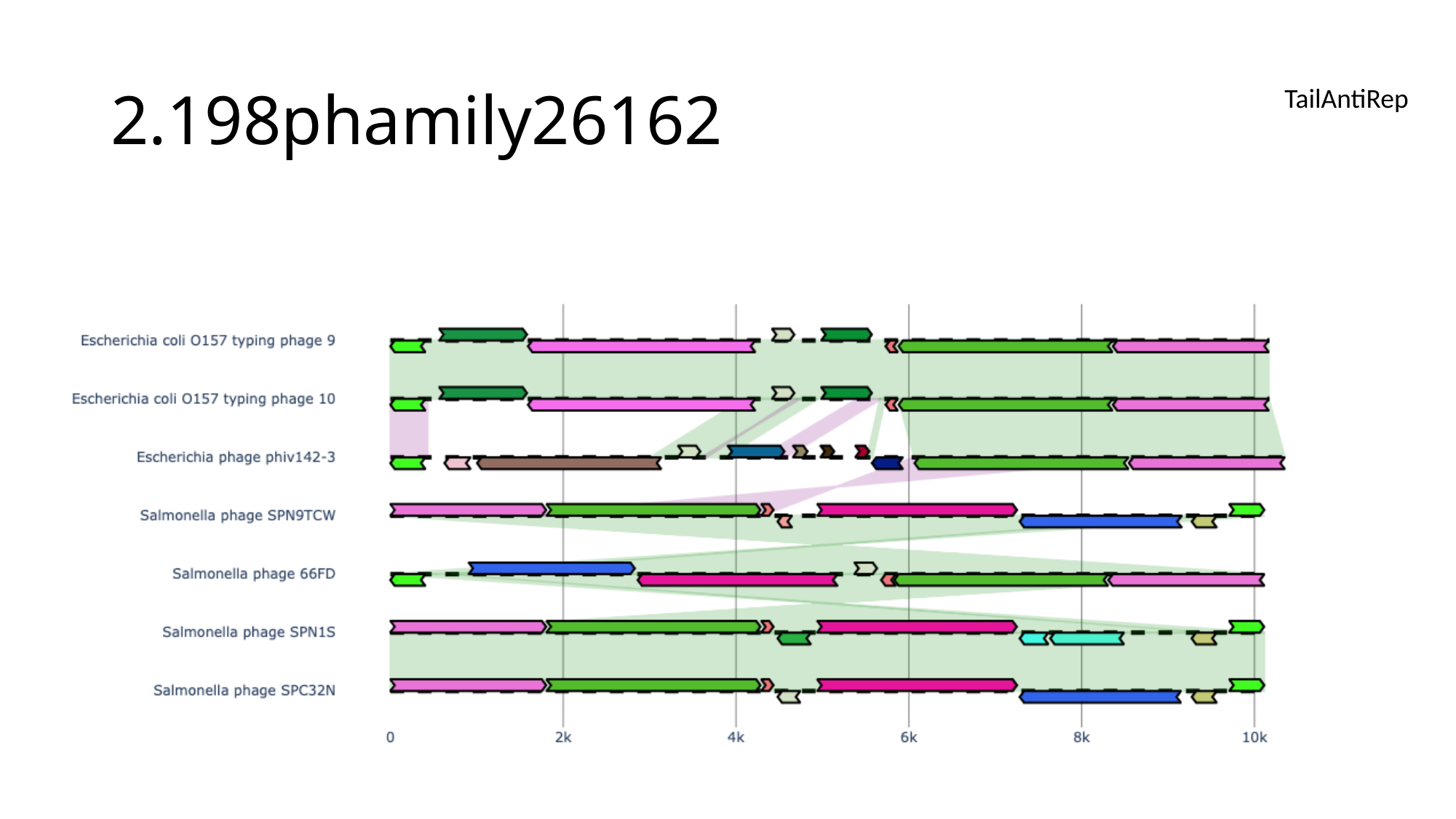

#### 2.198phamily26162
TailAntiRep

### Slide 13
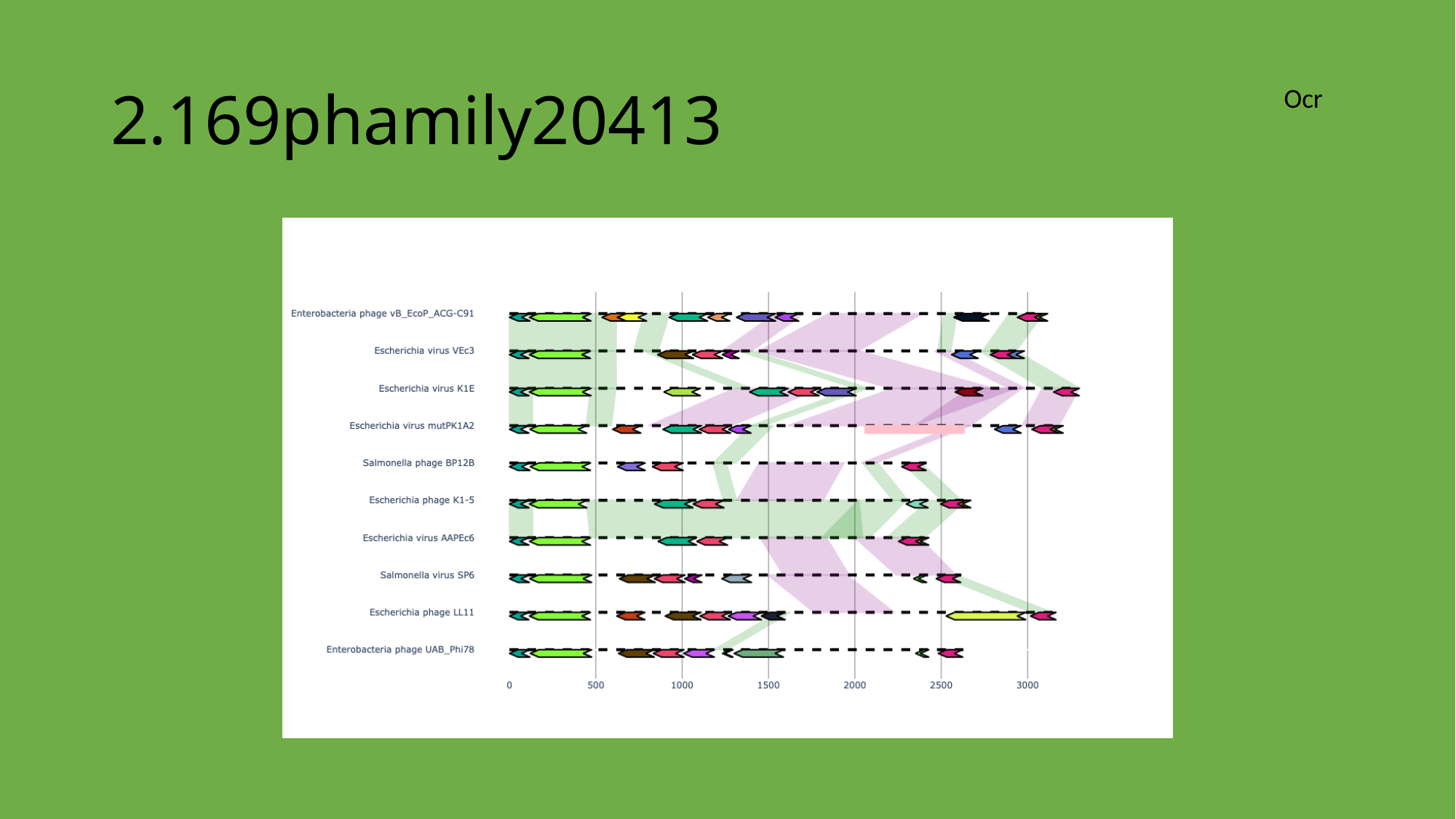

#### 2.169phamily20413
Ocr

### Slide 14
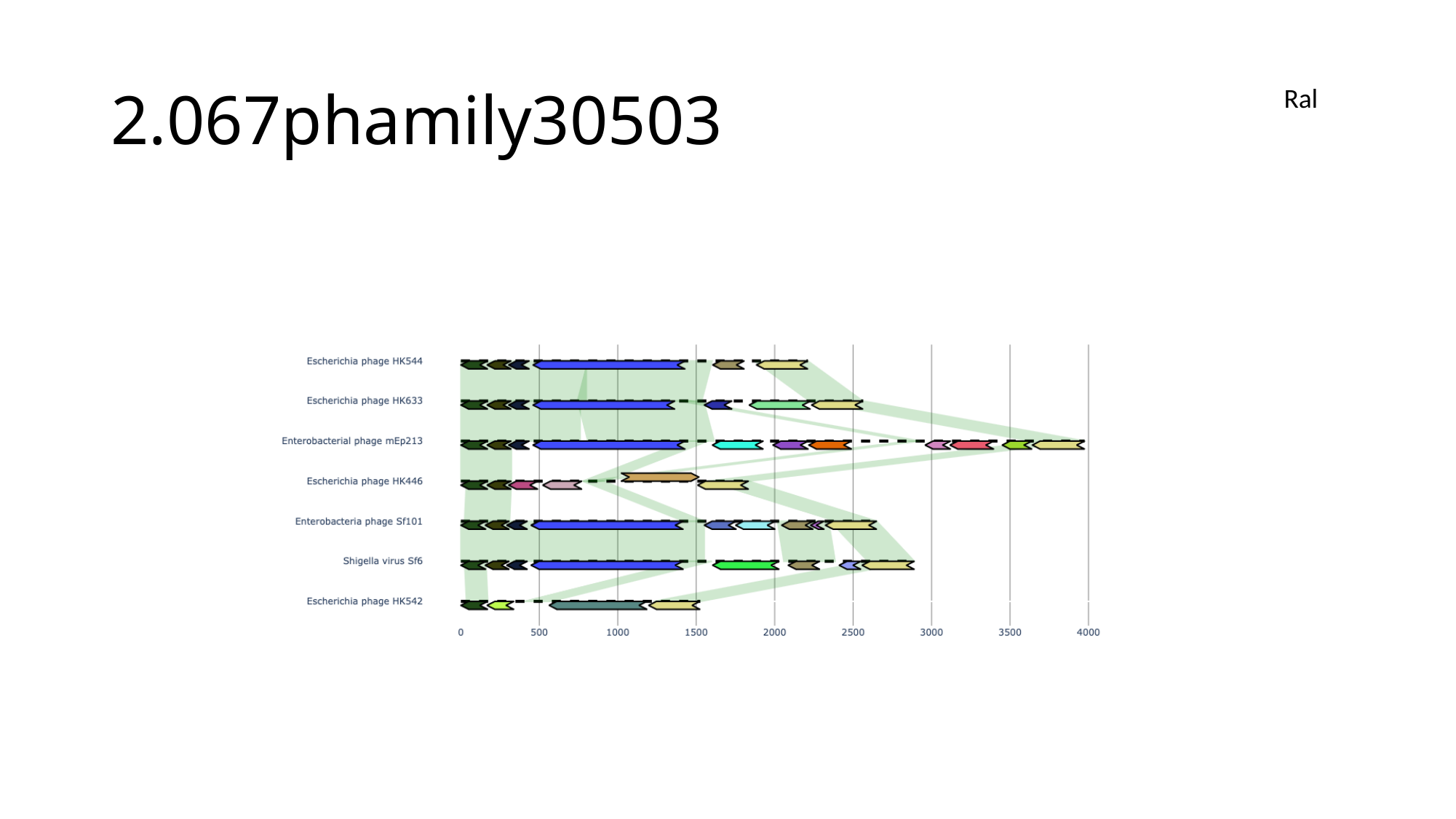

#### 2.067phamily30503
Ral

### Slide 15
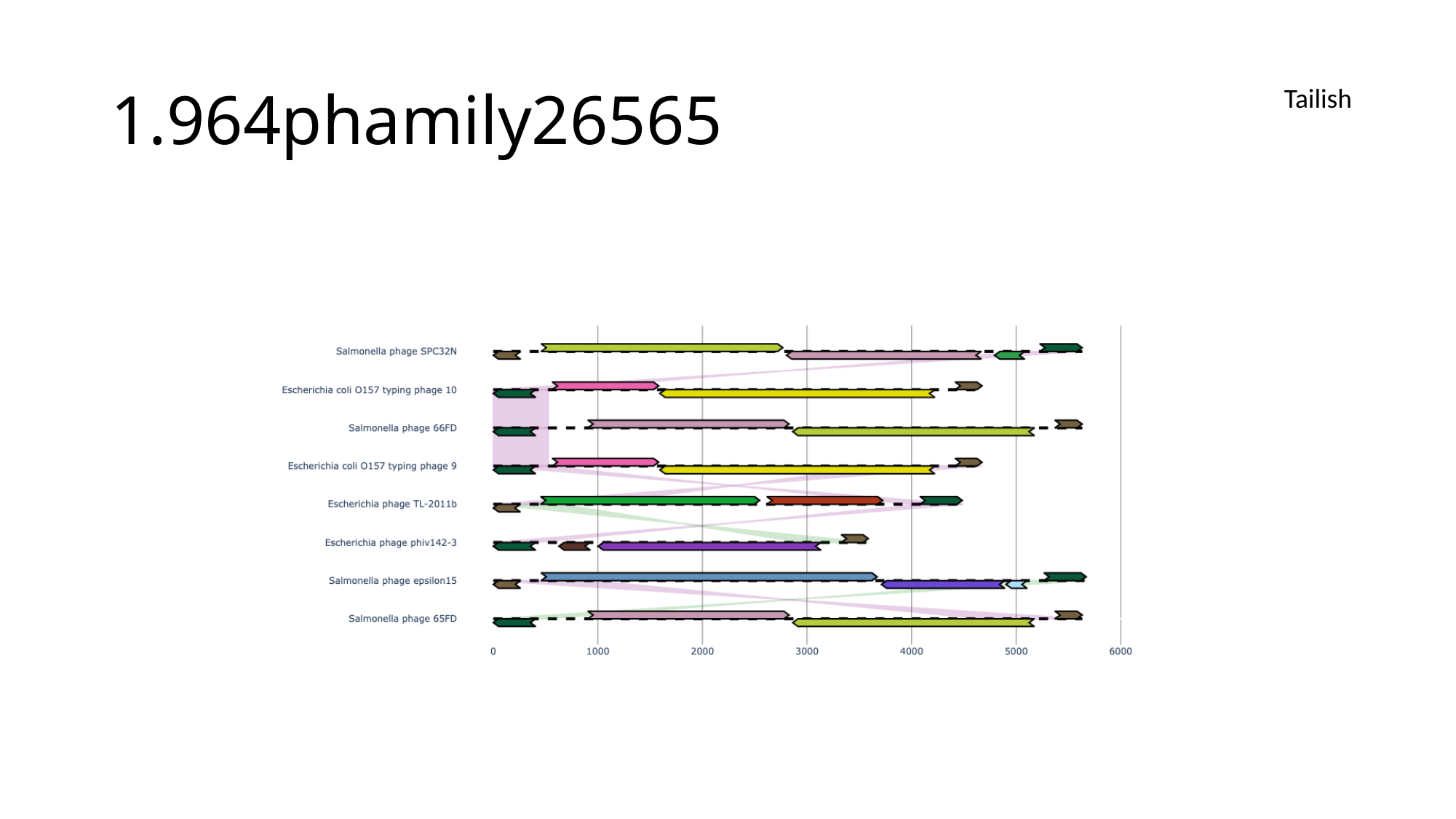

#### 1.964phamily26565
Tailish

### Slide 16
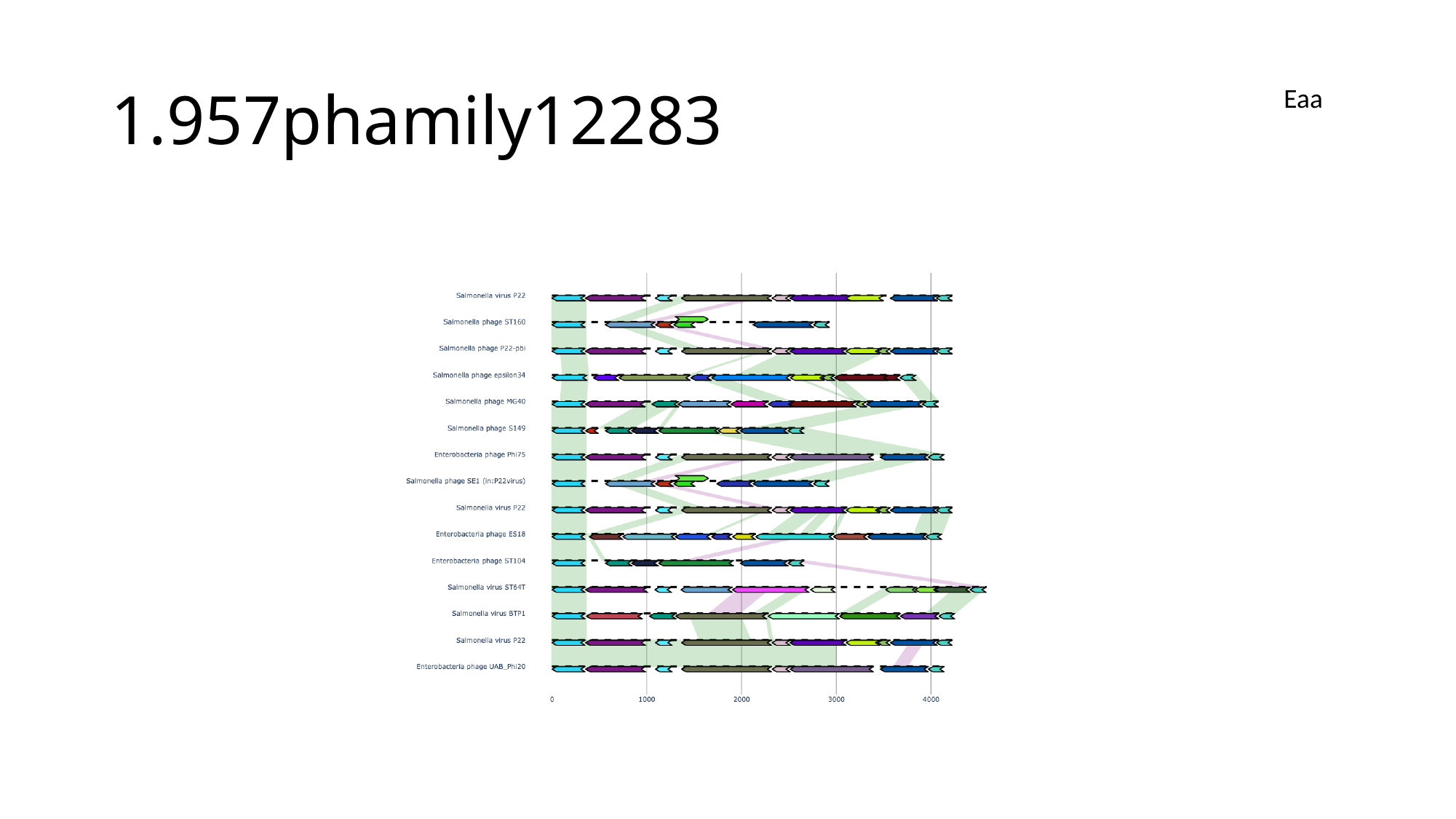

#### 1.957phamily12283
Eaa

### Slide 17
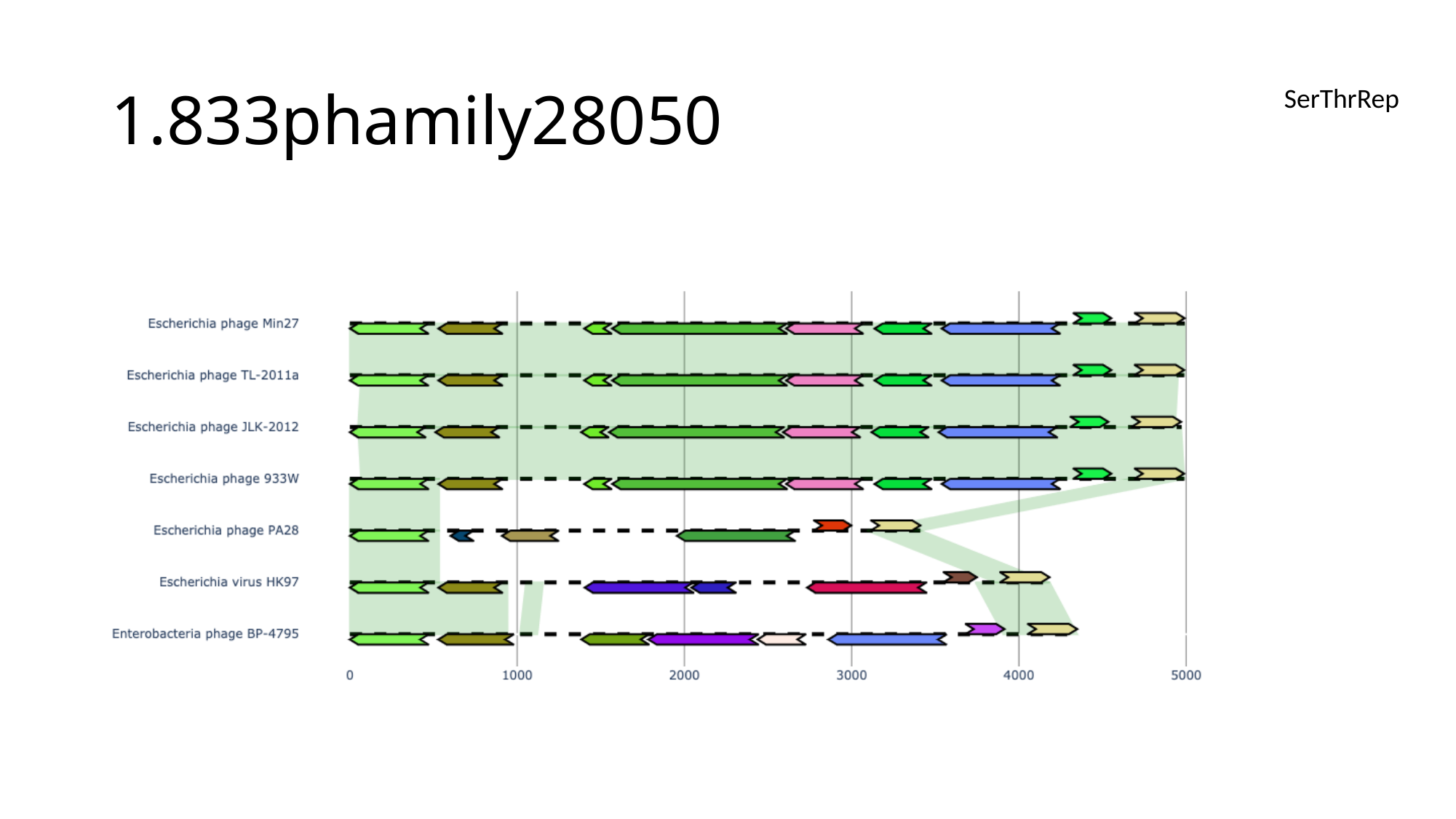

#### 1.833phamily28050
SerThrRep

### Slide 18
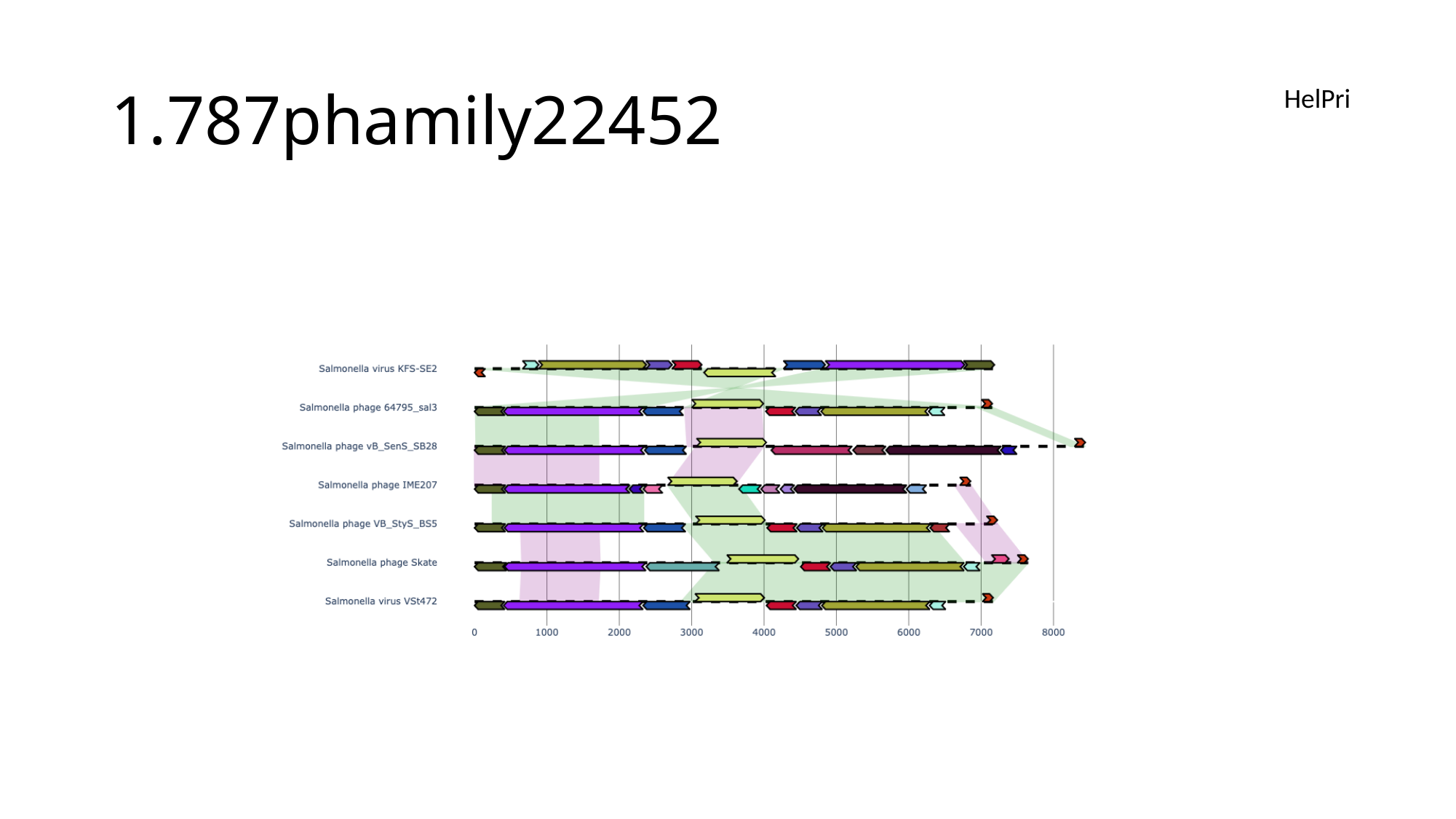

#### 1.787phamily22452
HelPri

### Slide 19
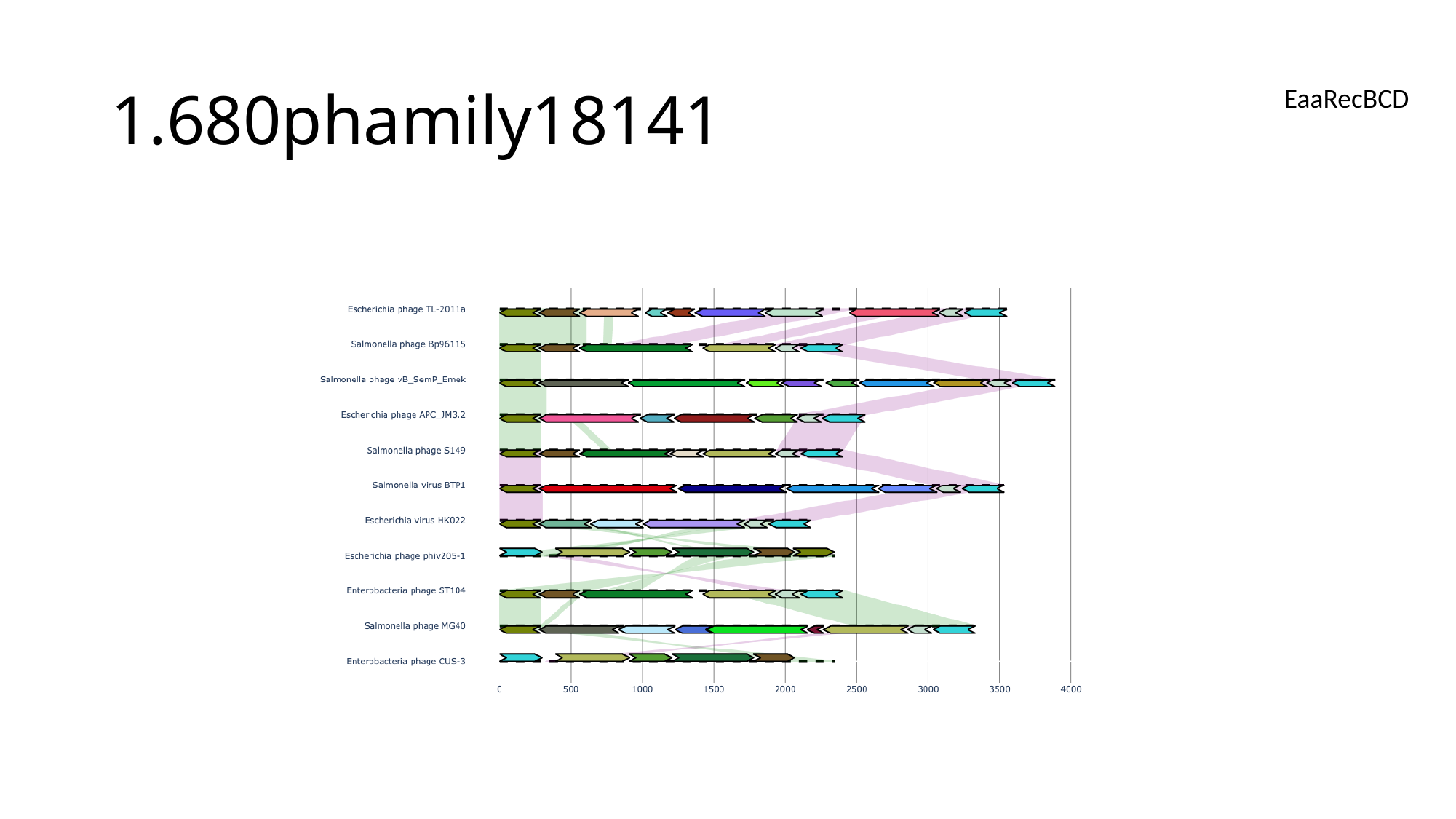

#### 1.680phamily18141
EaaRecBCD

### Slide 20
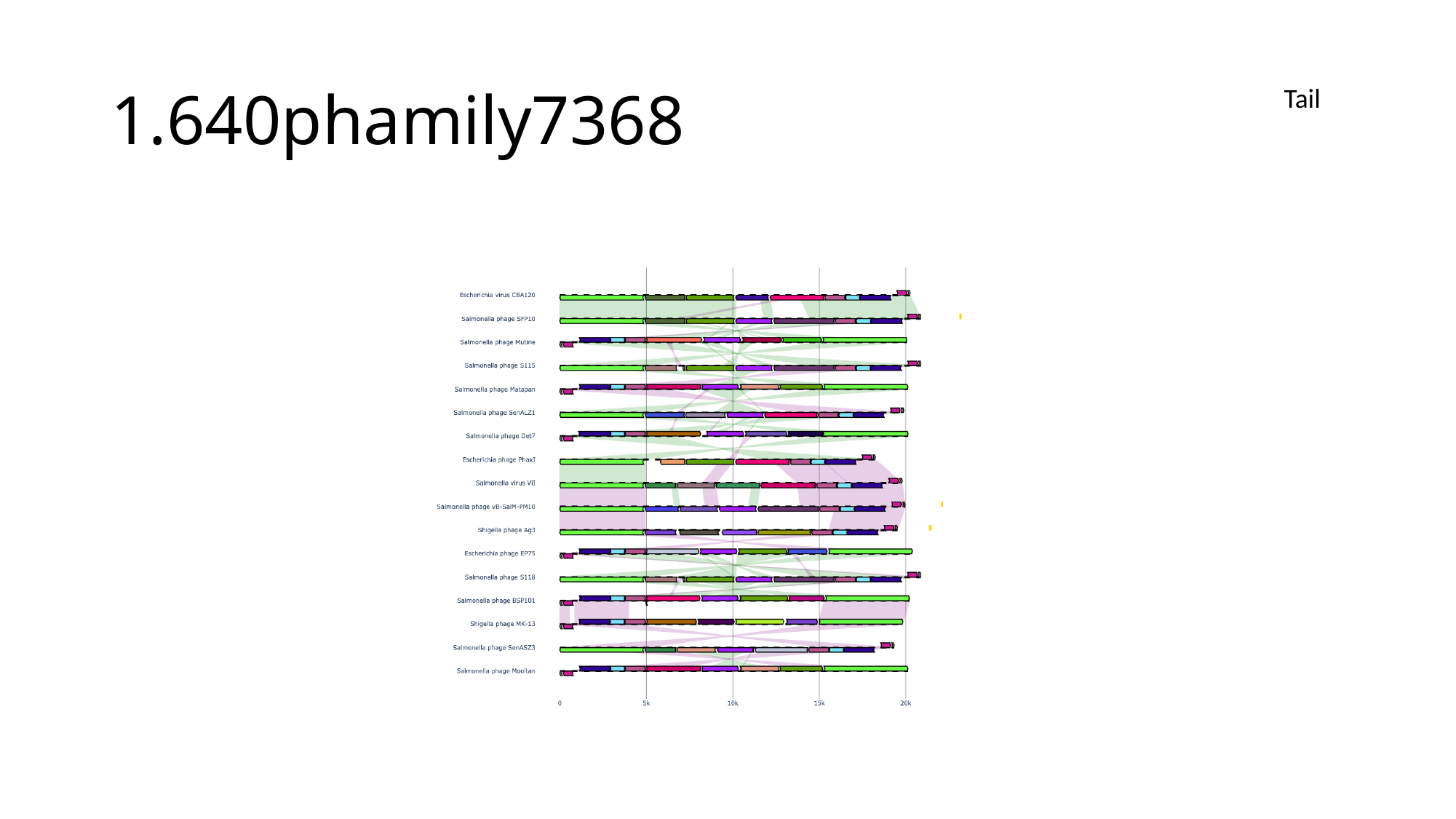

#### 1.640phamily7368
Tail

### Slide 21
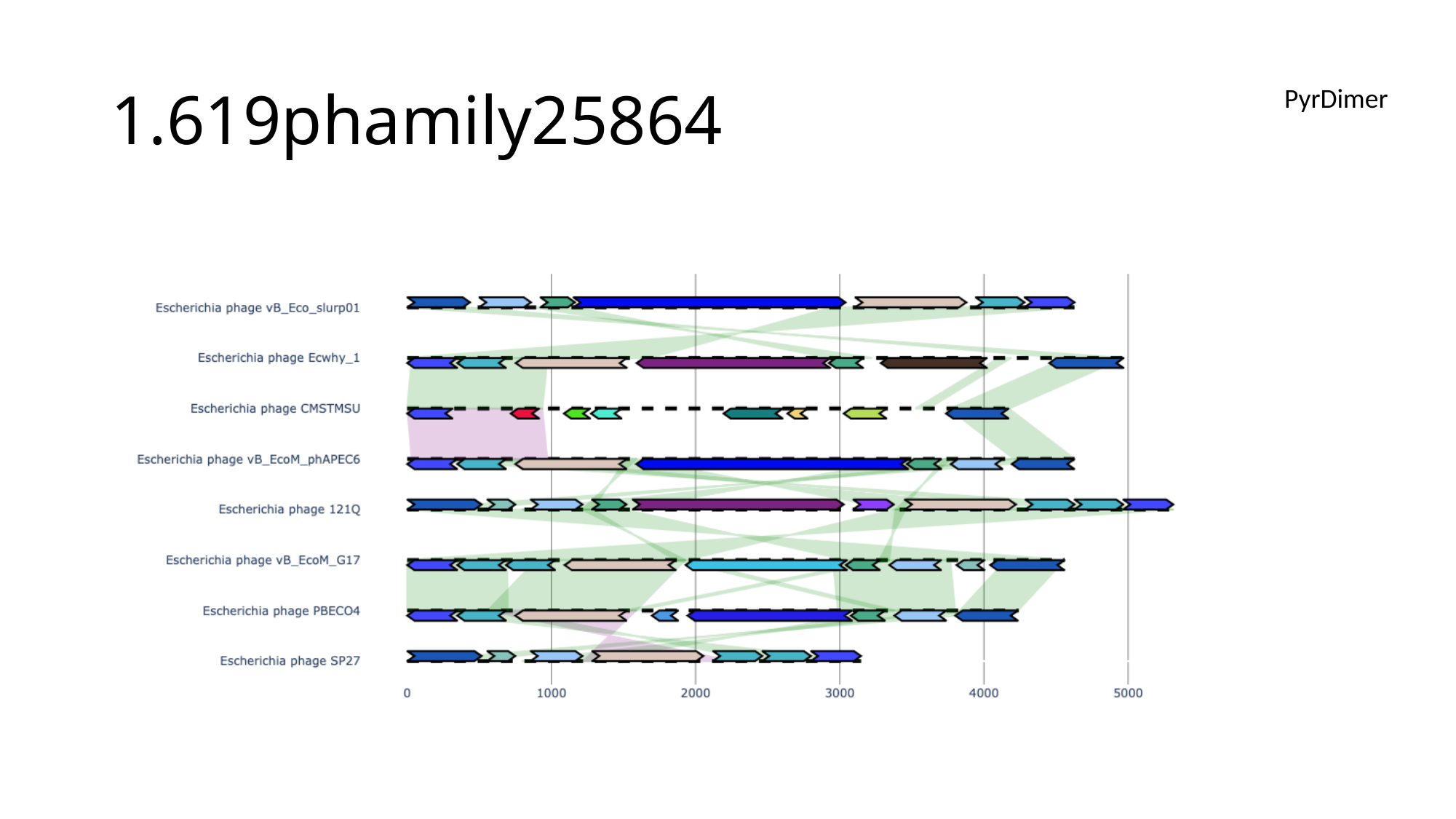

#### 1.619phamily25864
PyrDimer

### Slide 22
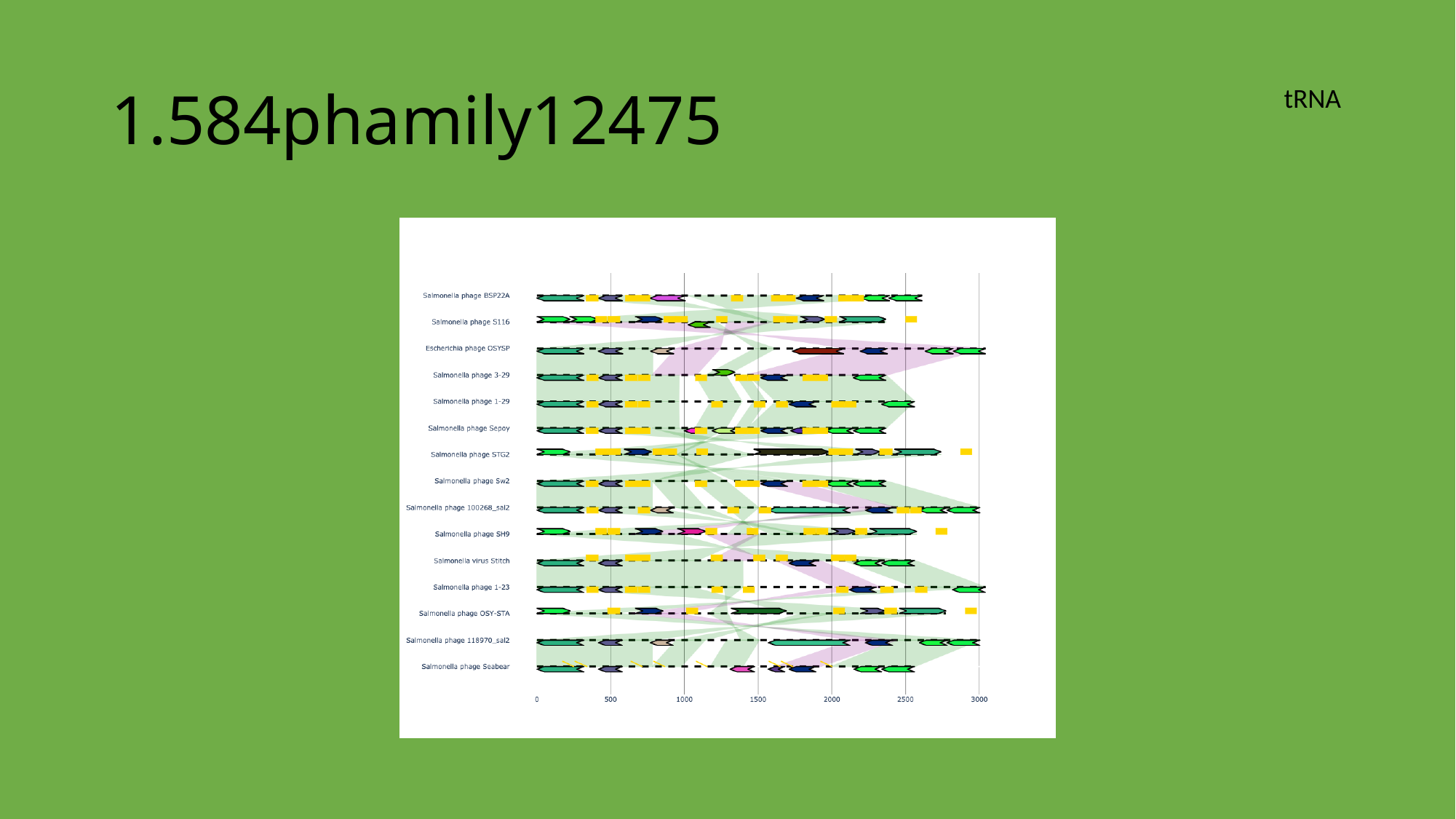

#### 1.584phamily12475
tRNA

### Slide 23
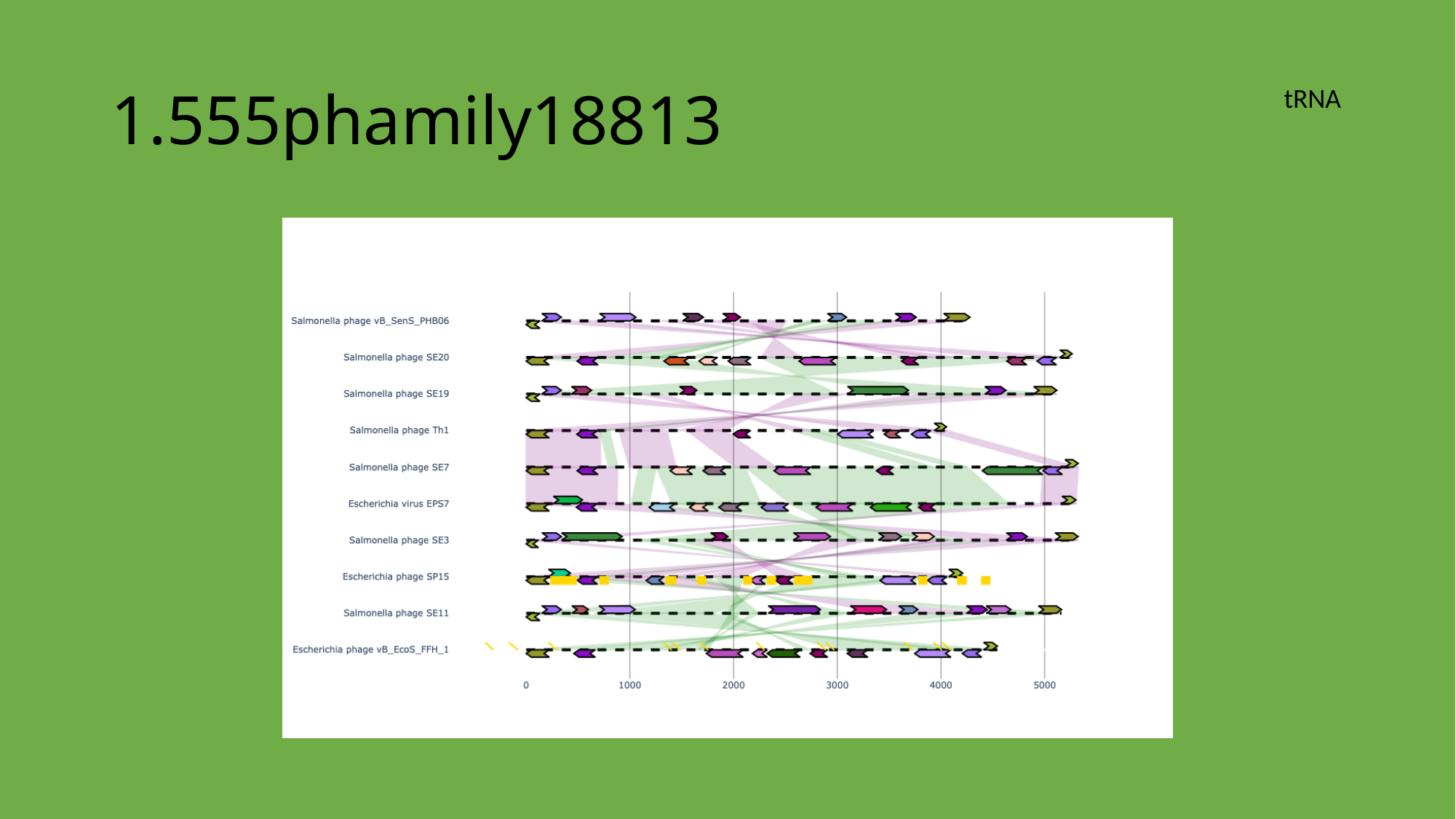

#### 1.555phamily18813
tRNA

### Slide 24
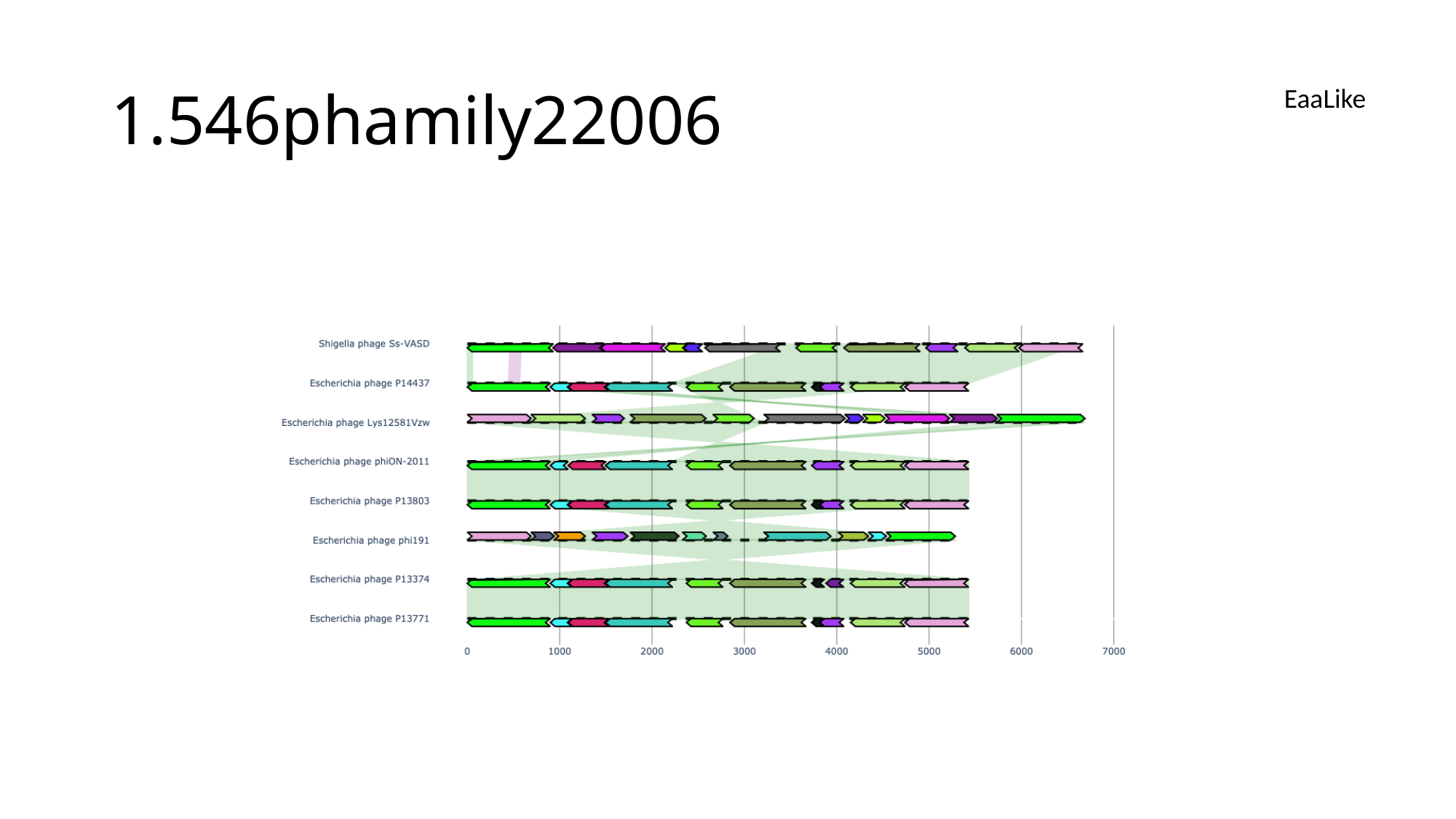

#### 1.546phamily22006
EaaLike

### Slide 25
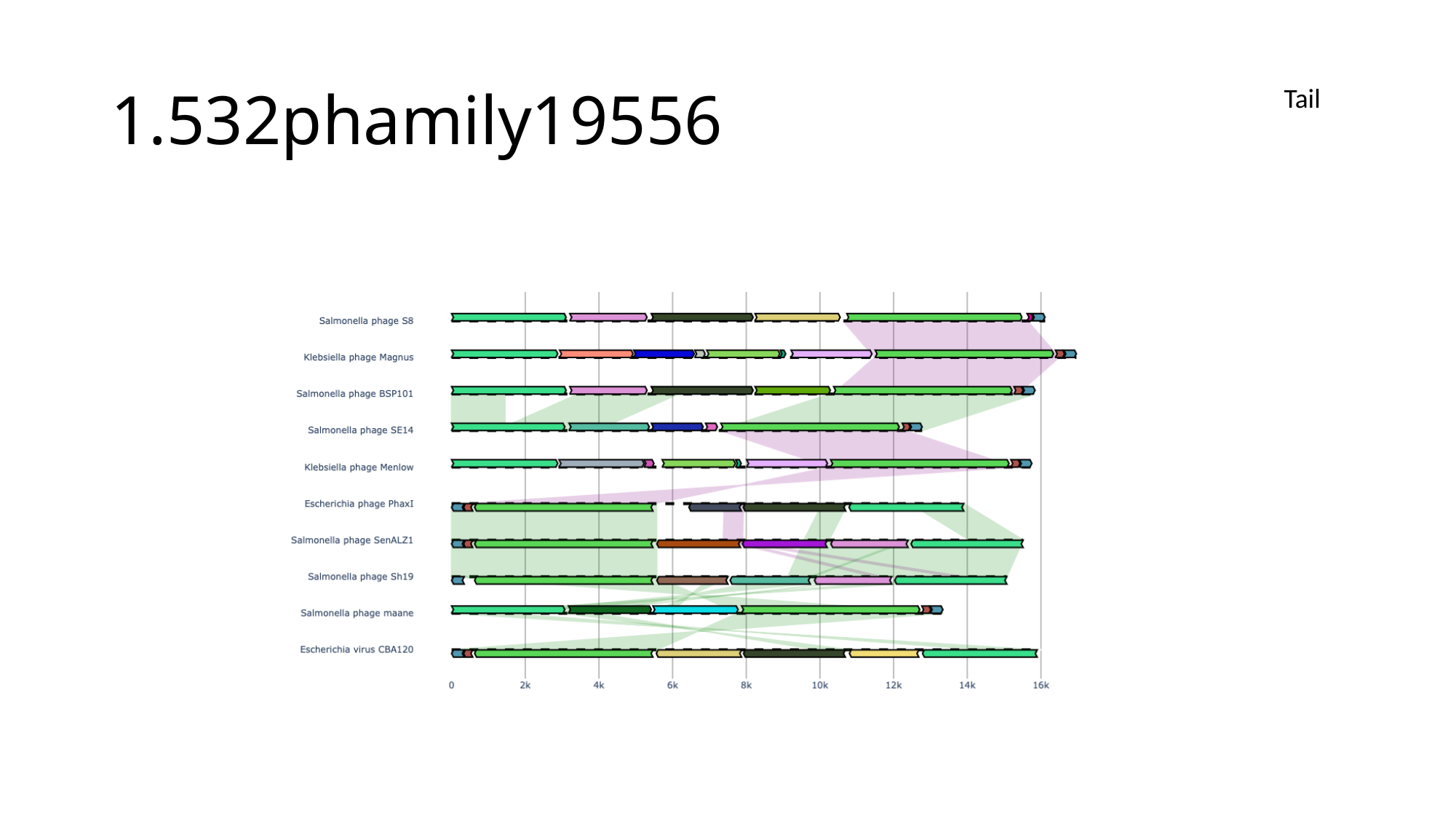

#### 1.532phamily19556
Tail

### Slide 26
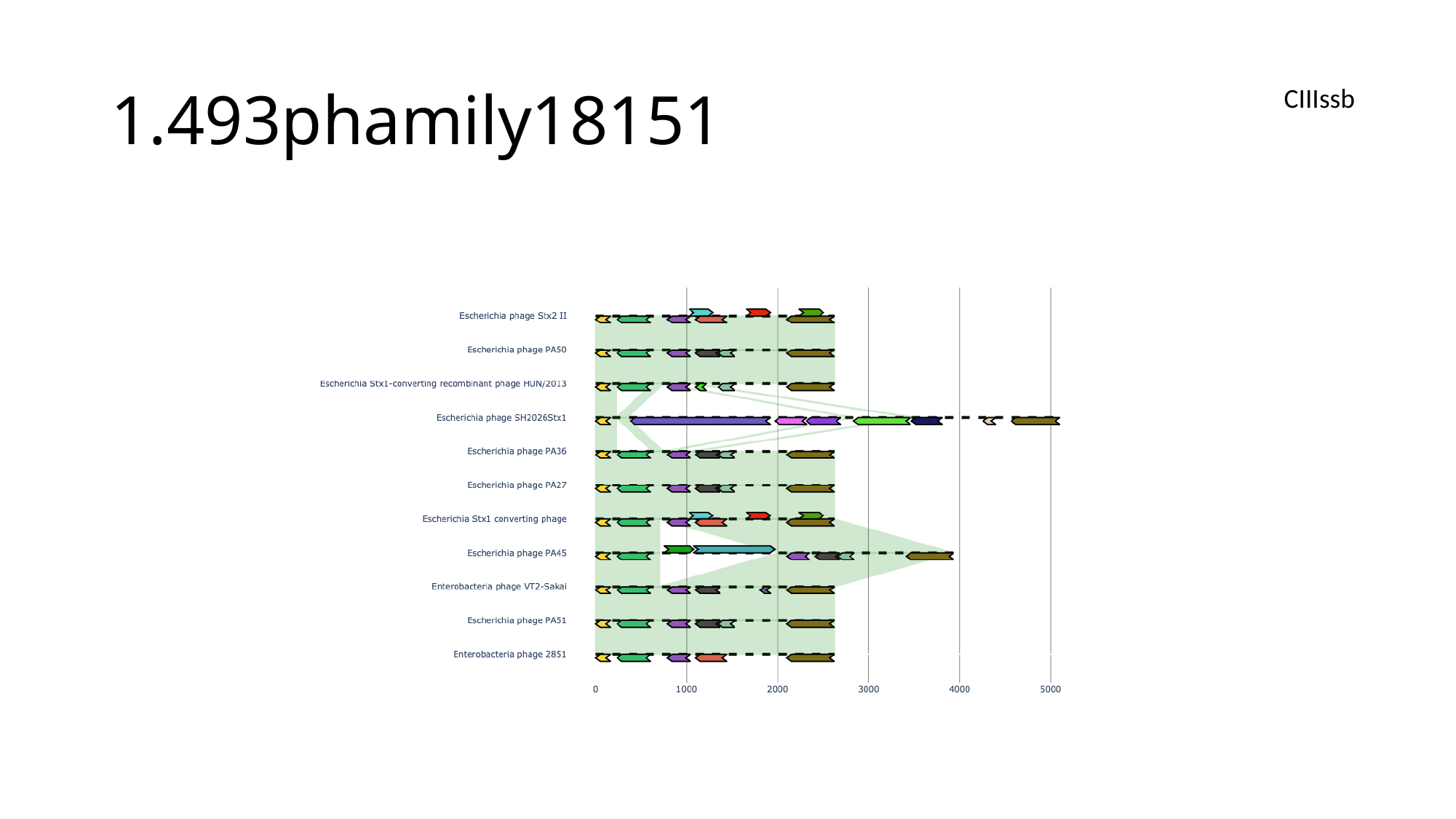

#### 1.493phamily18151
CIIIssb

### Slide 27
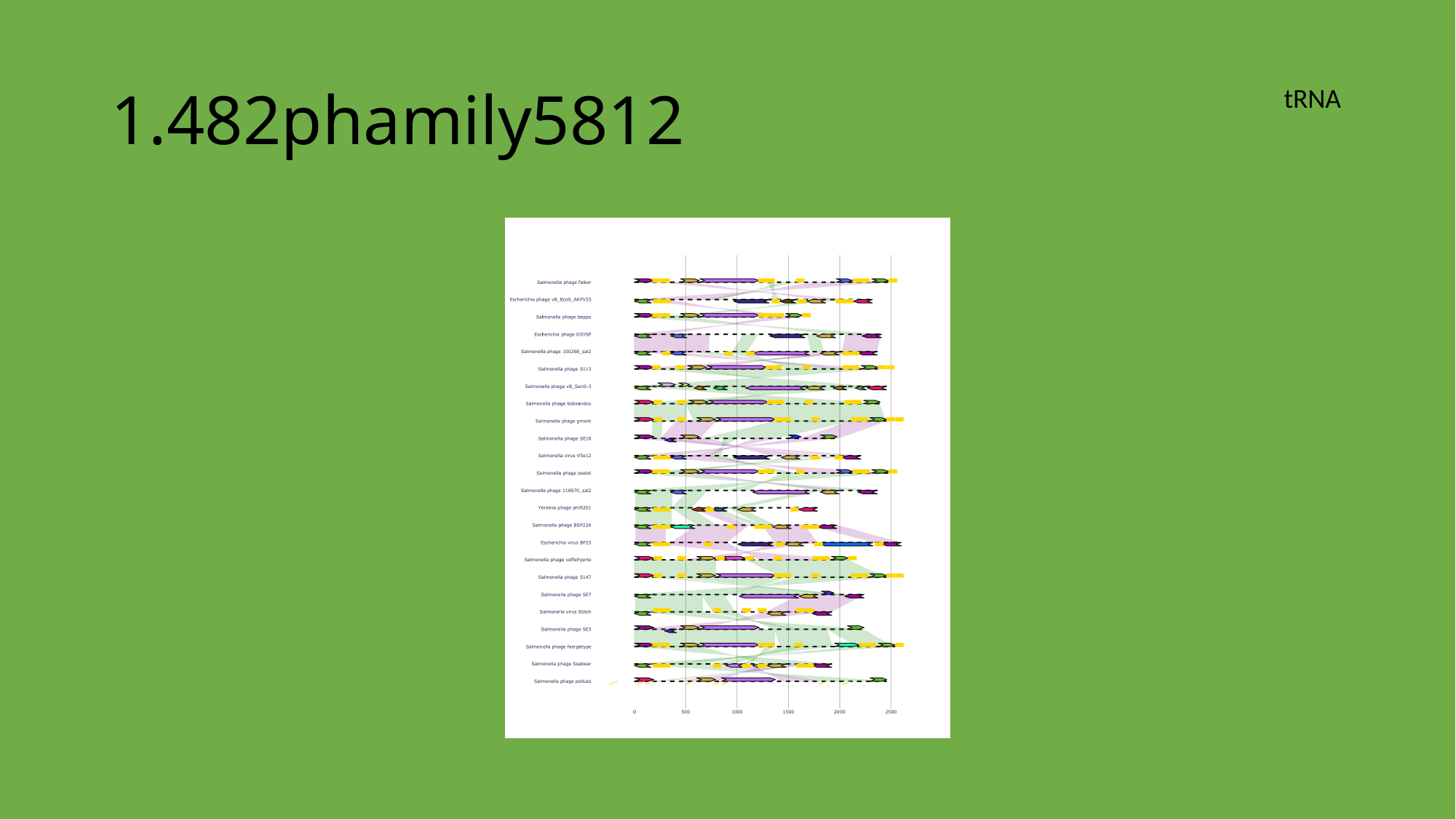

#### 1.482phamily5812
tRNA

### Slide 28
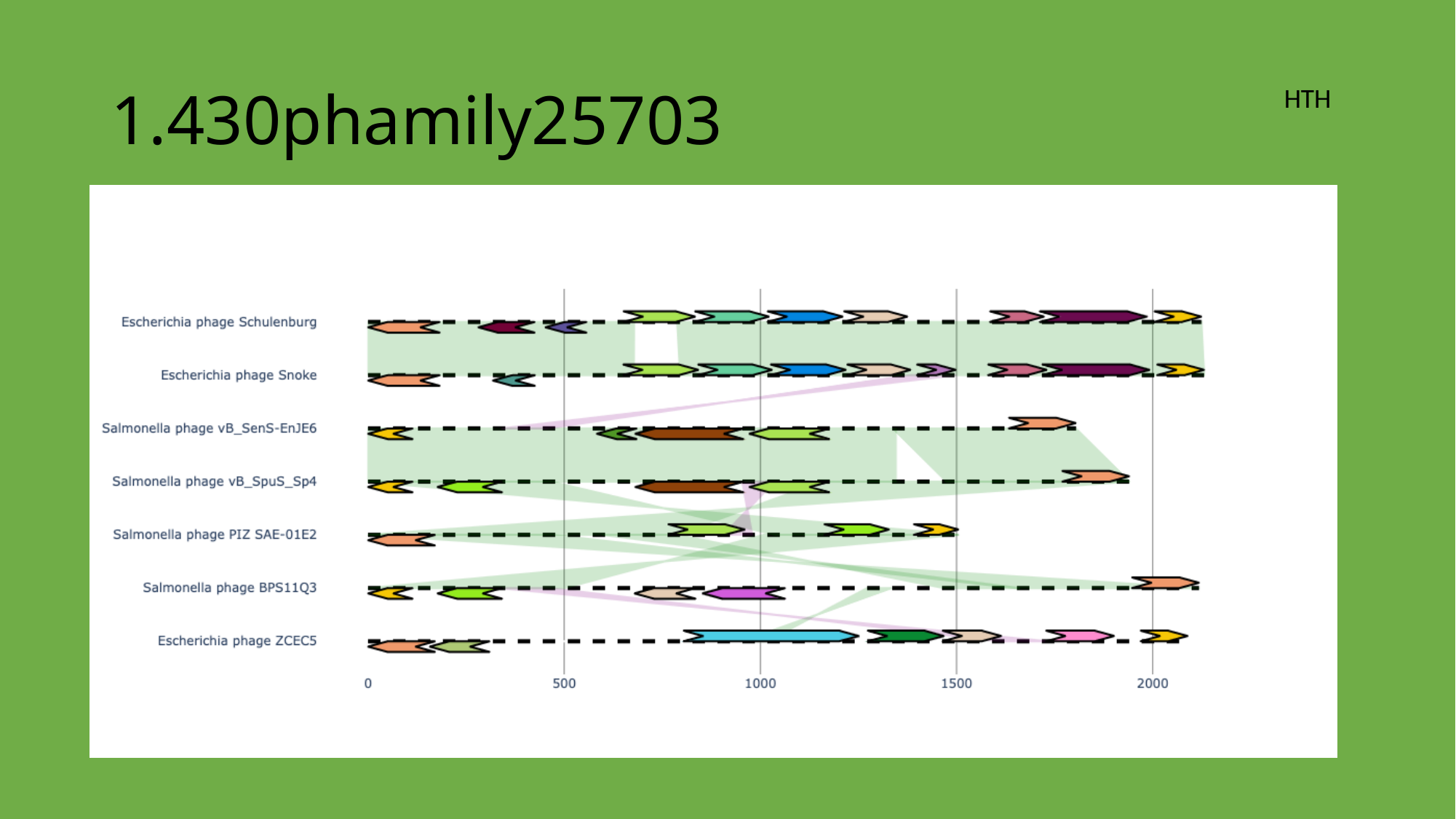

#### 1.430phamily25703
HTH

### Slide 29
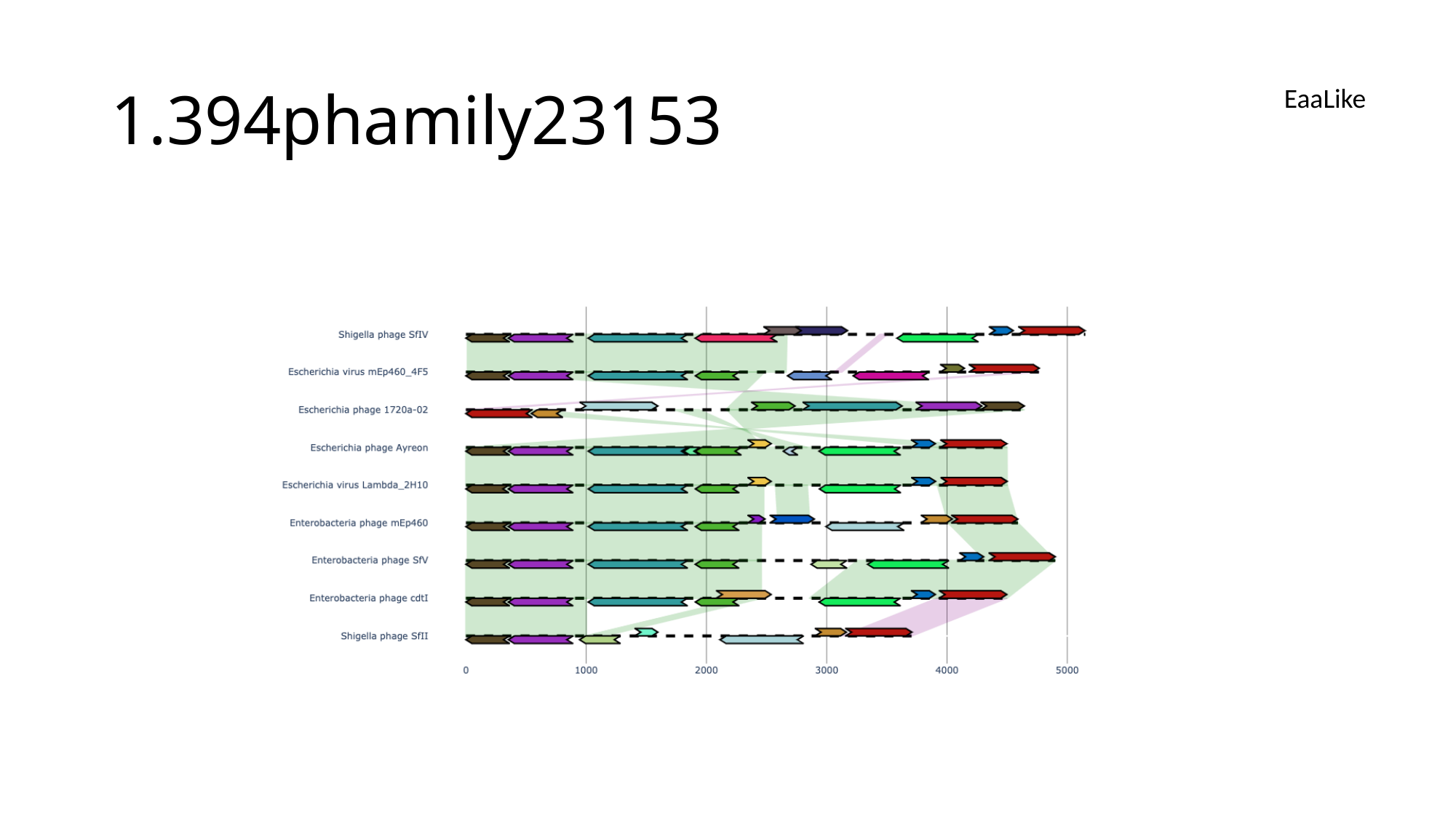

#### 1.394phamily23153
EaaLike

### Slide 30
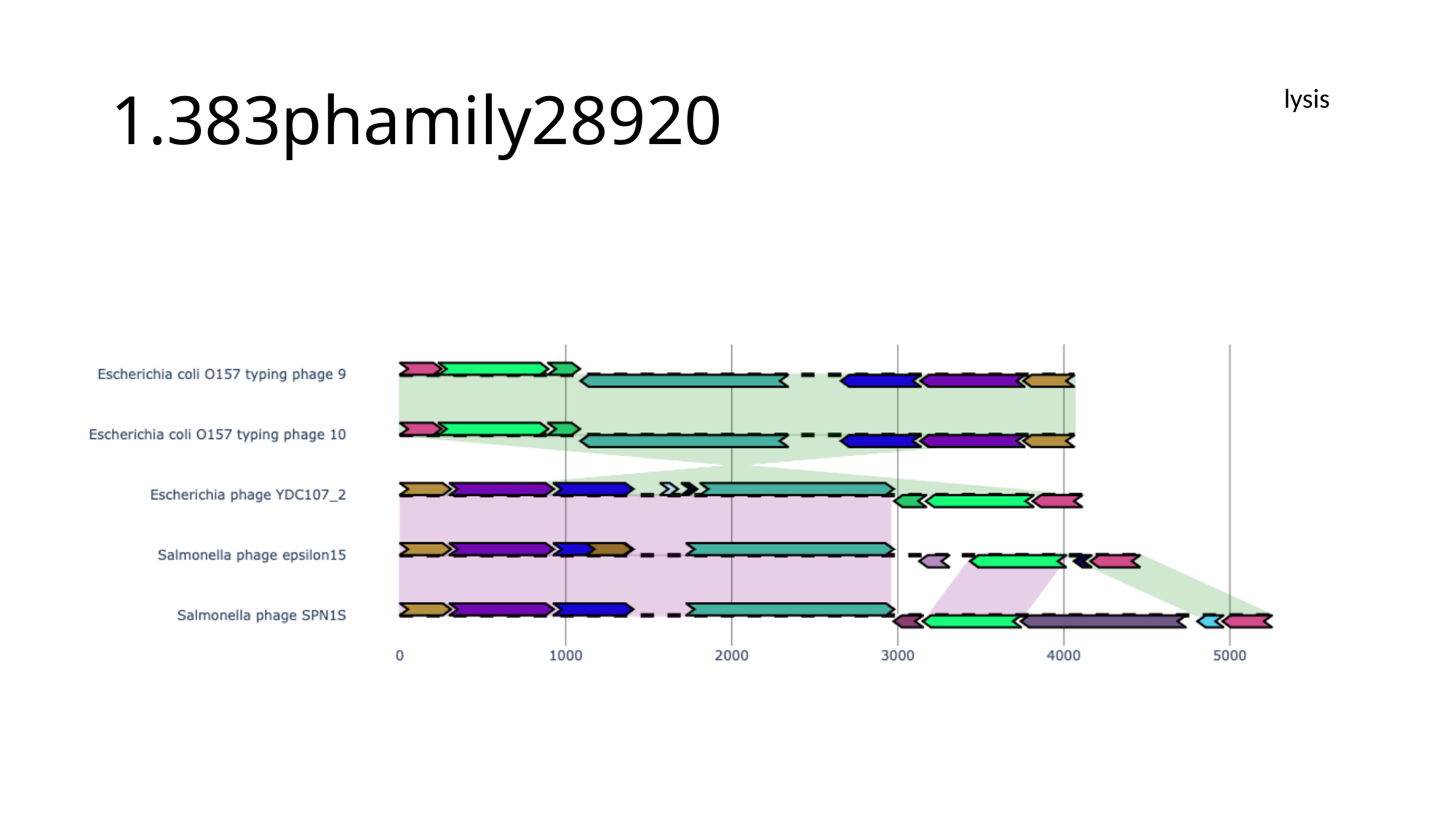

#### 1.383phamily28920
lysis

### Slide 31
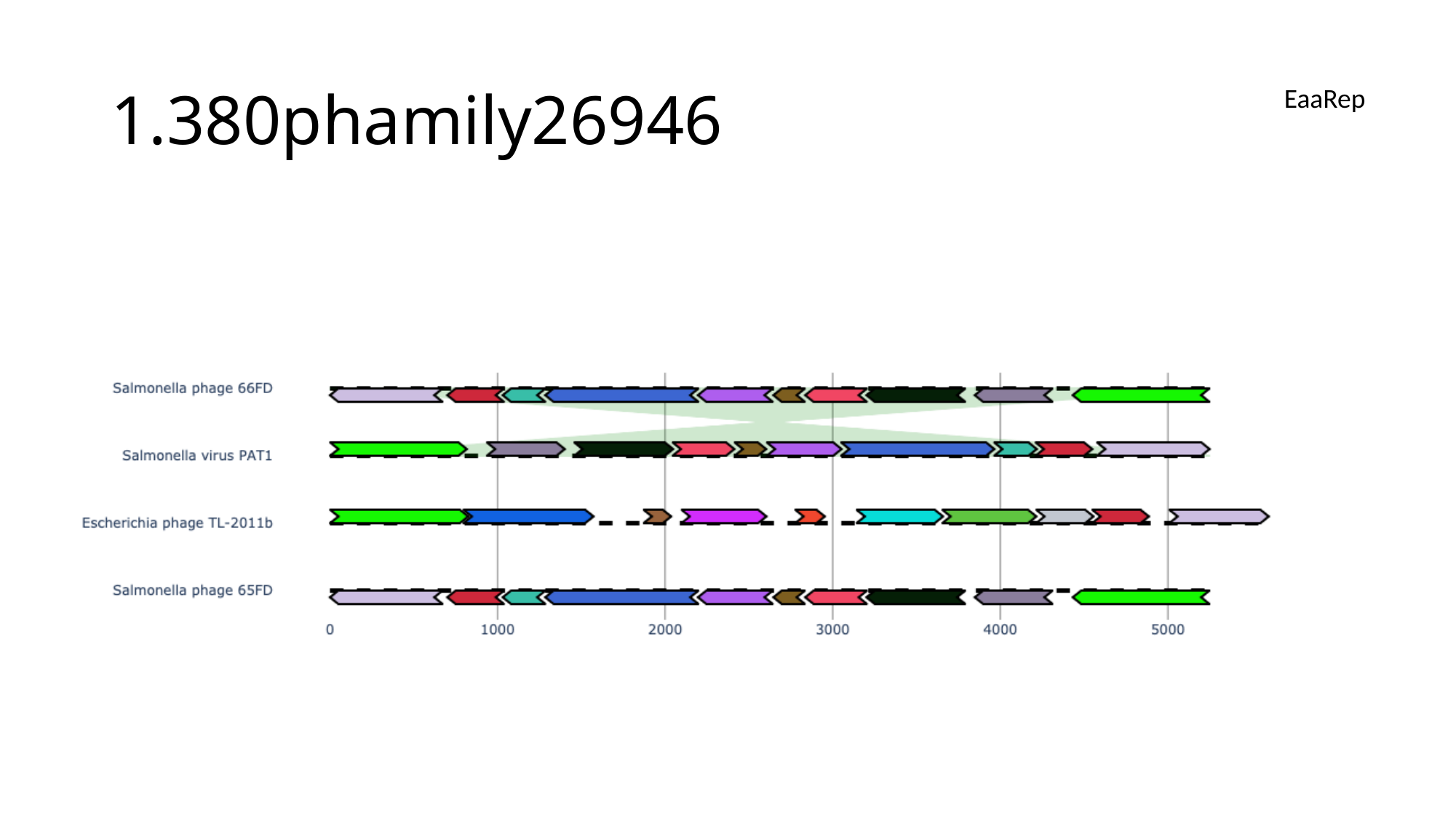

#### 1.380phamily26946
EaaRep

### Slide 32
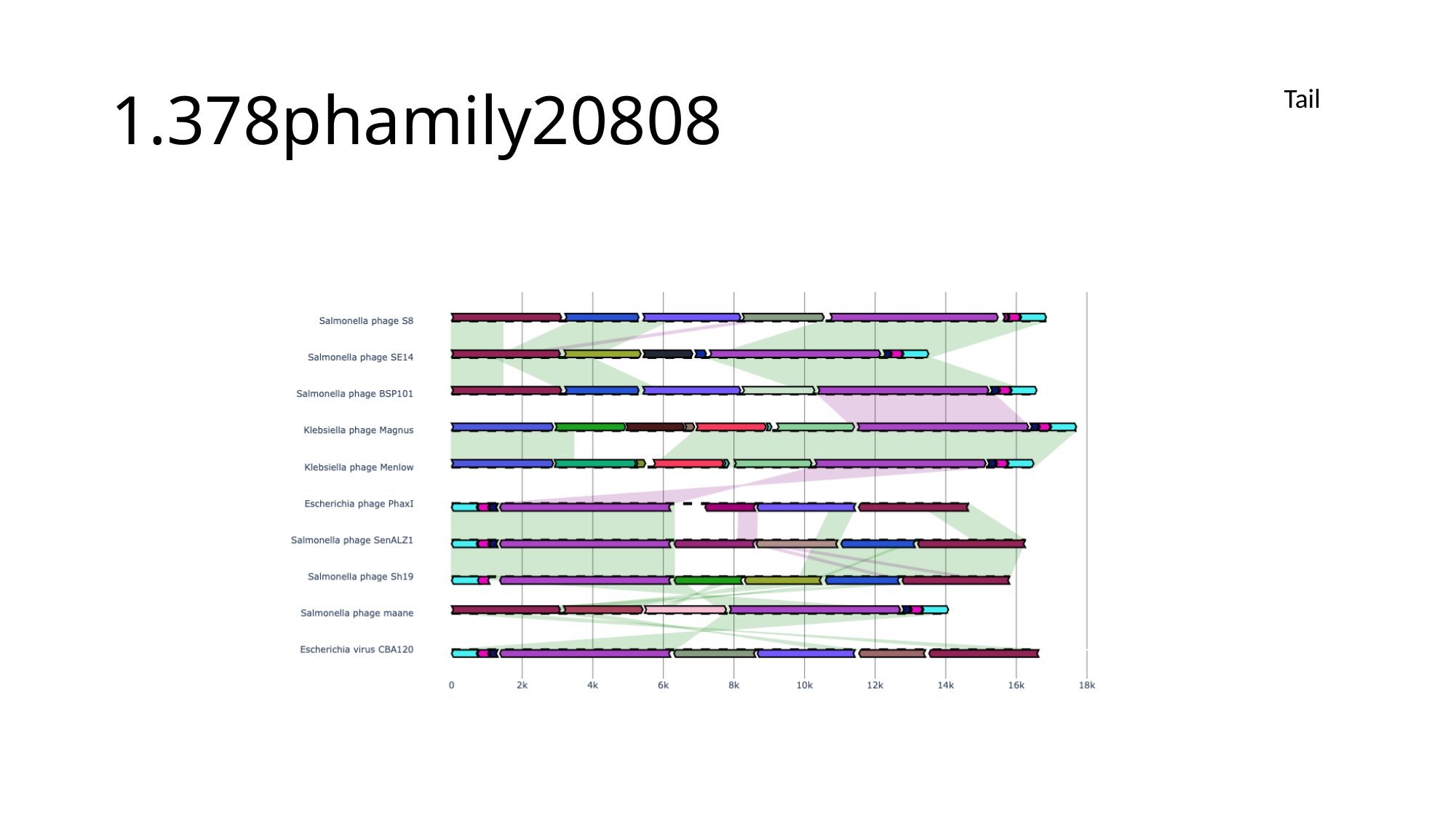

#### 1.378phamily20808
Tail

### Slide 33
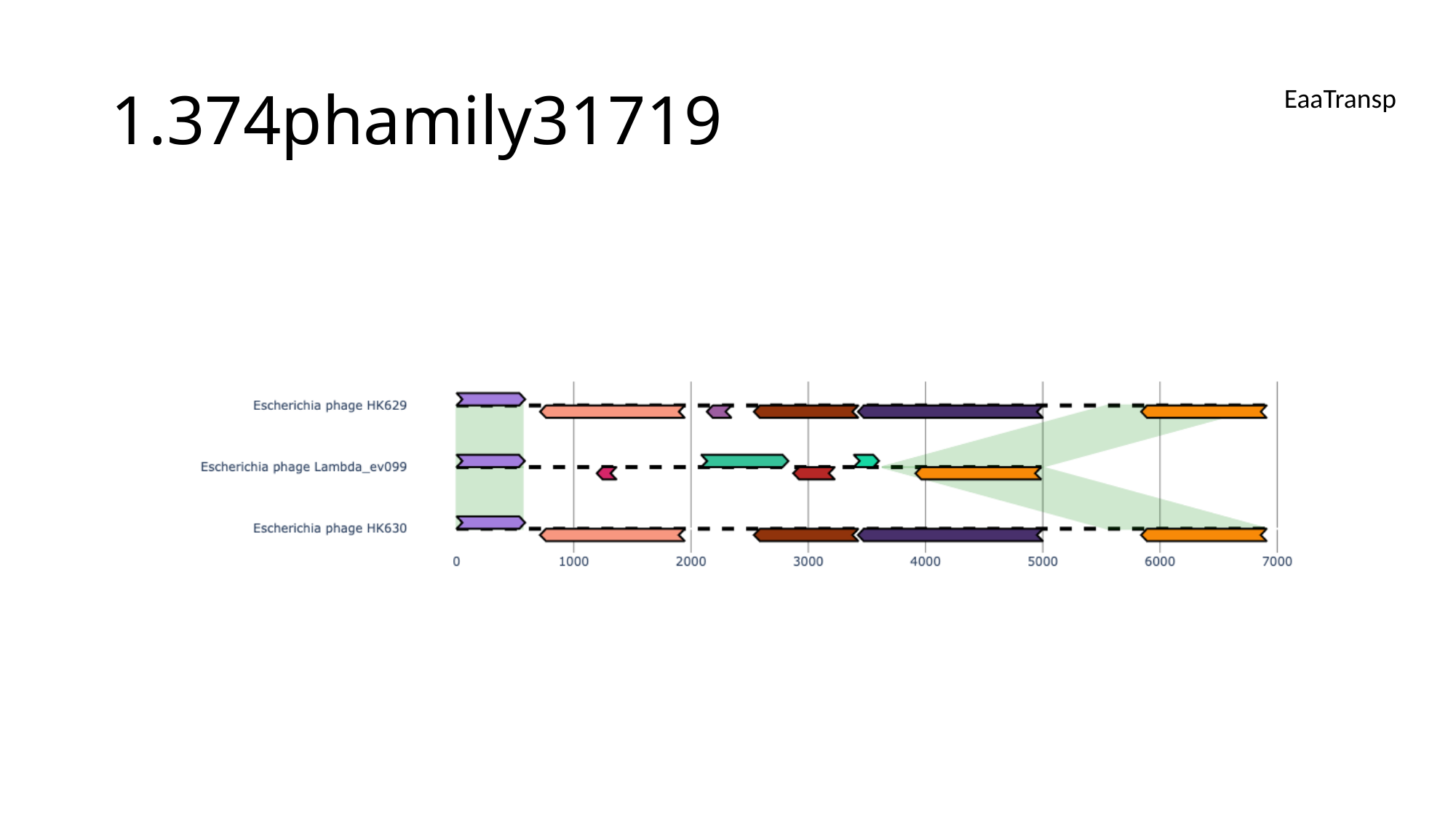

#### 1.374phamily31719
EaaTransp

### Slide 34
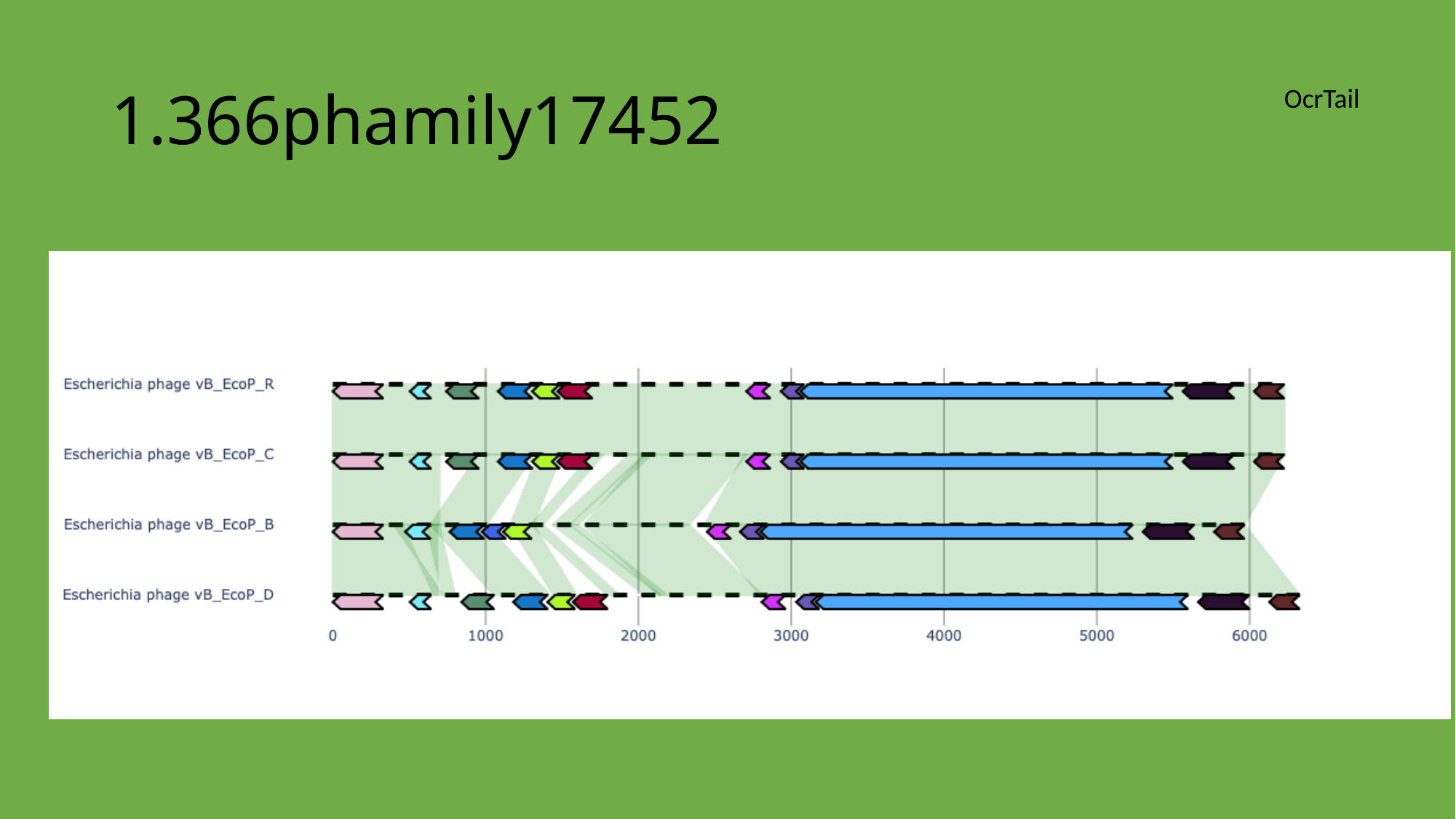

#### 1.366phamily17452
OcrTail

### Slide 35
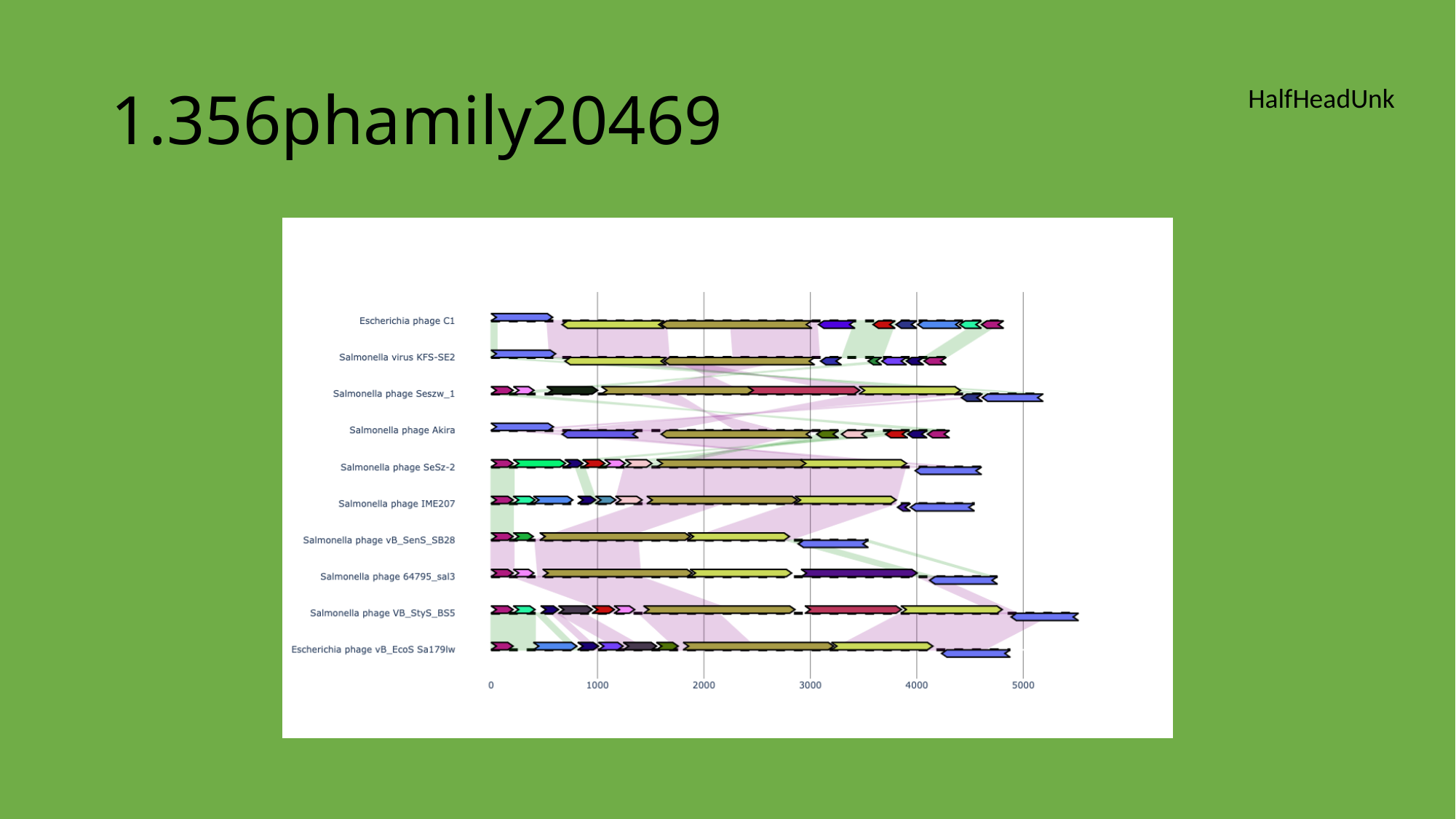

#### 1.356phamily20469
HalfHeadUnk

### Slide 36
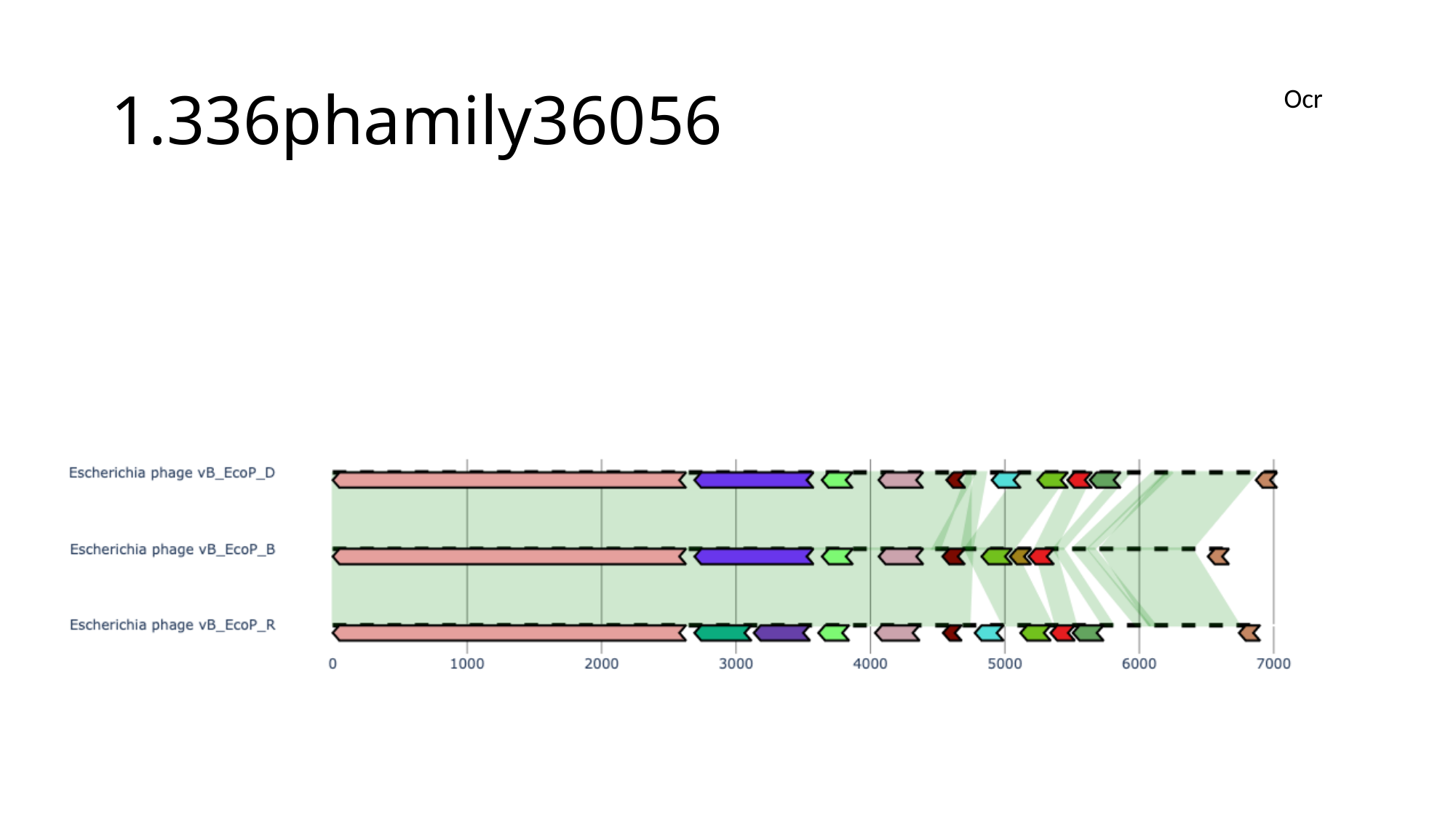

#### 1.336phamily36056
Ocr

### Slide 37
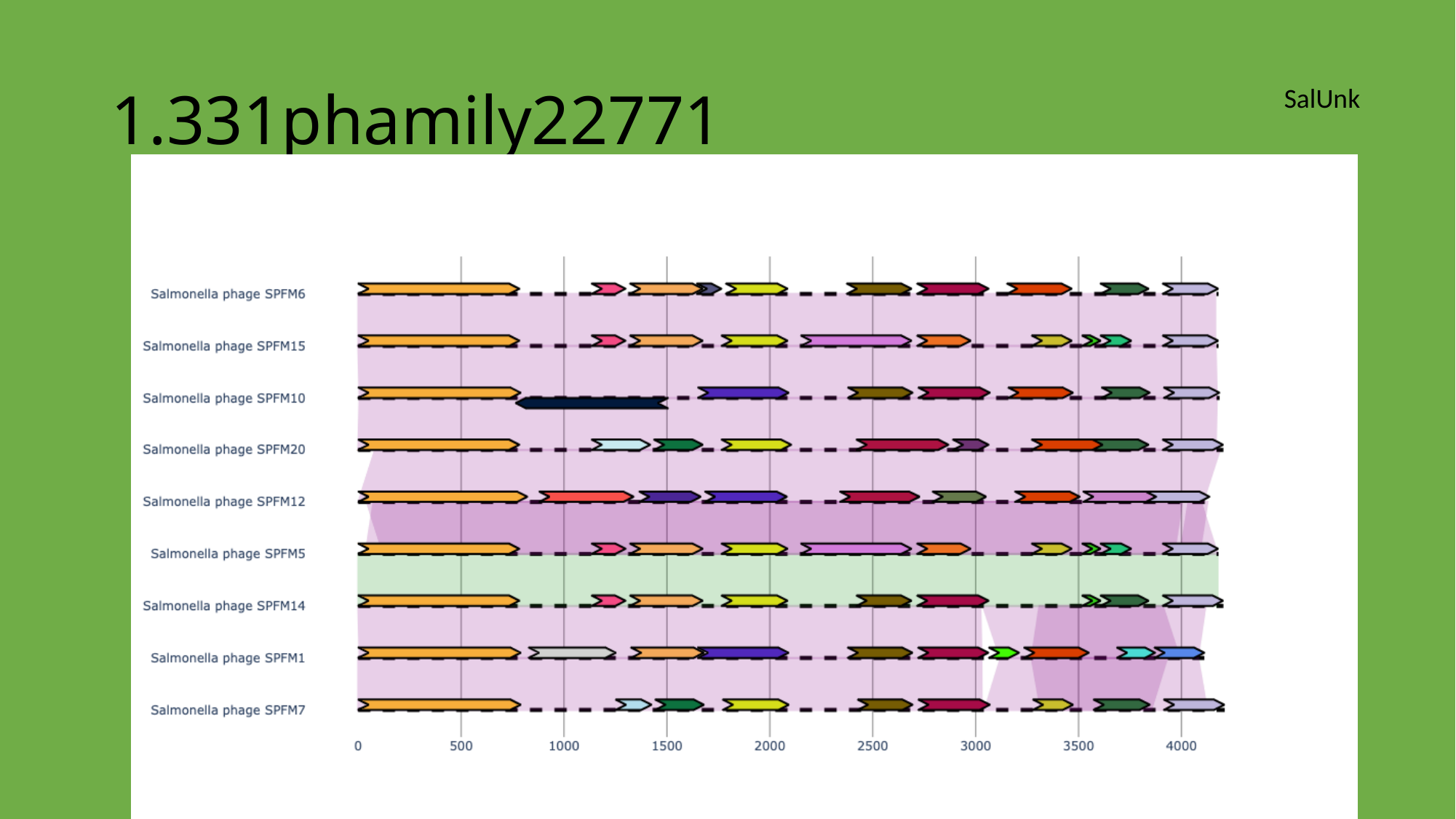

#### 1.331phamily22771
SalUnk

### Slide 38
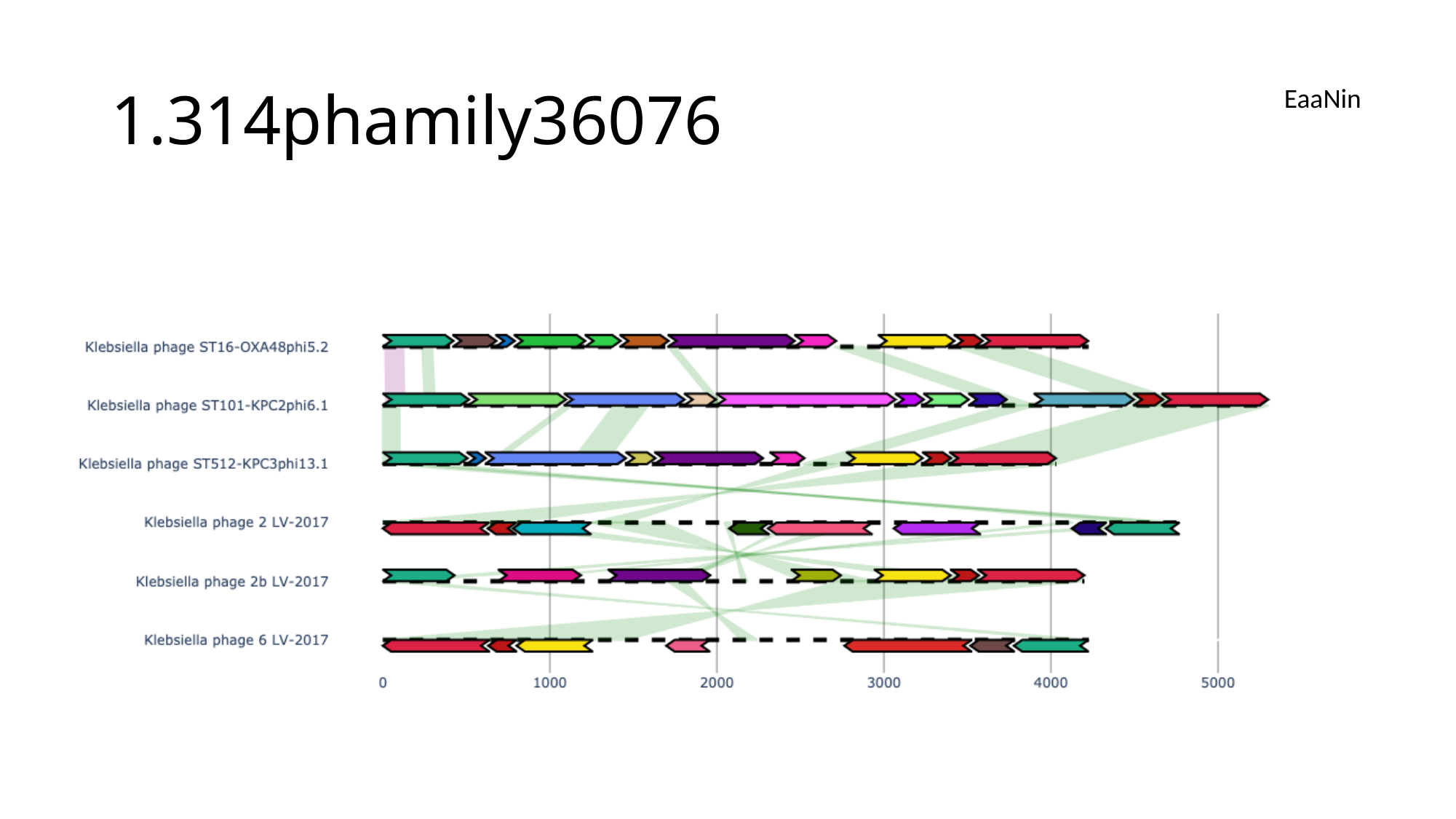

#### 1.314phamily36076
EaaNin

### Slide 39
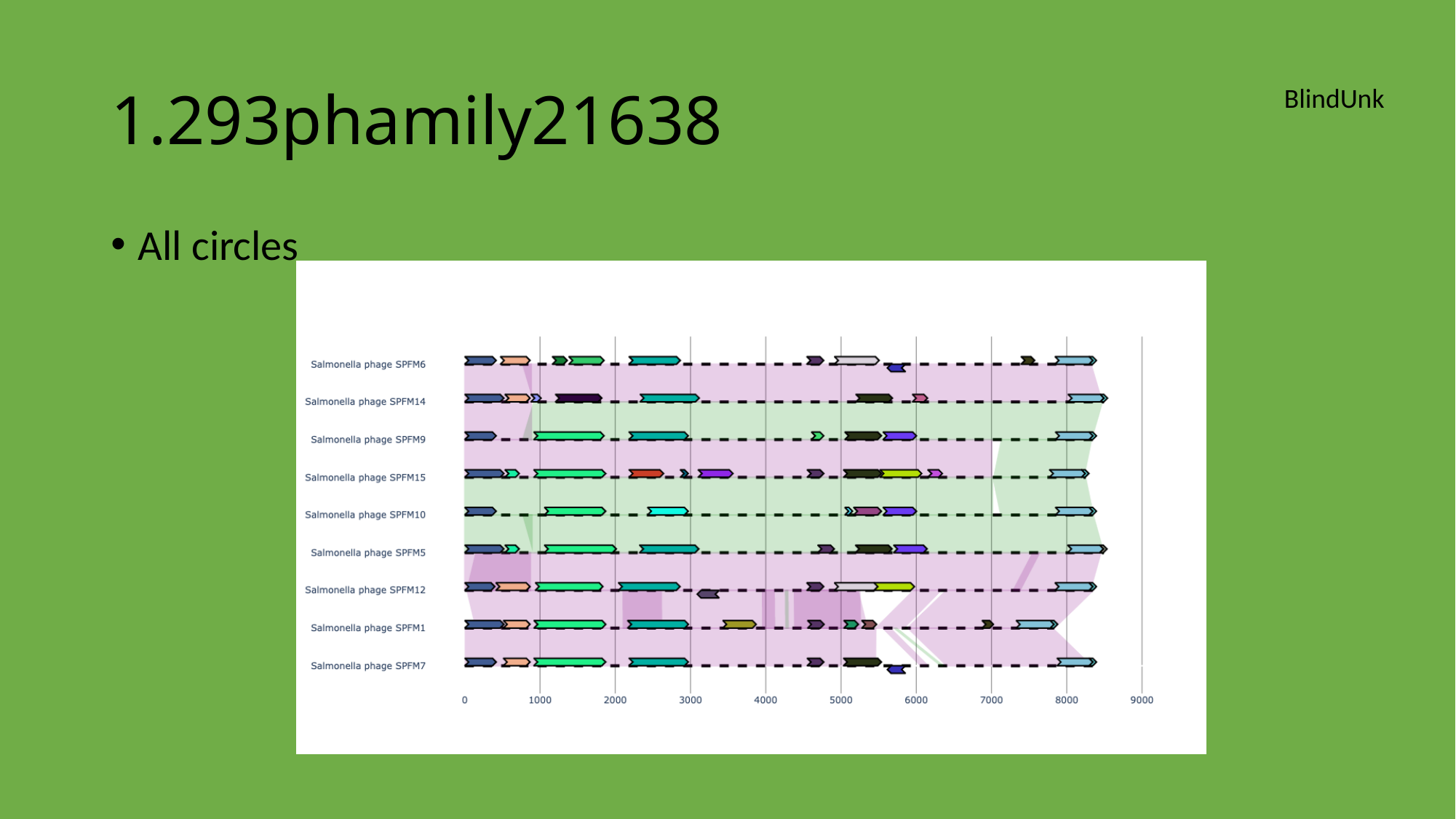

#### 1.293phamily21638
BlindUnk
All circles

### Slide 40
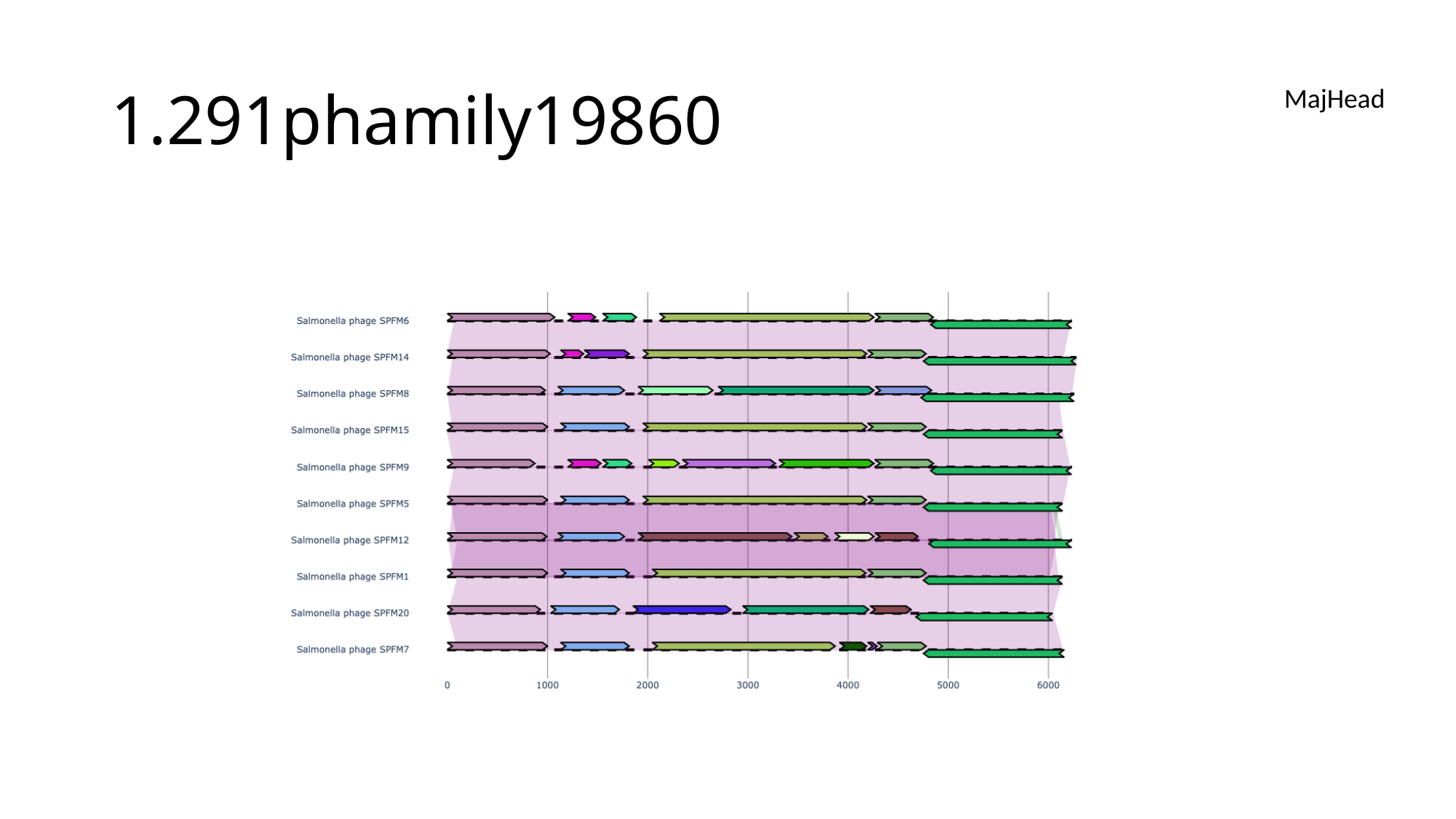

#### 1.291phamily19860
MajHead

### Slide 41
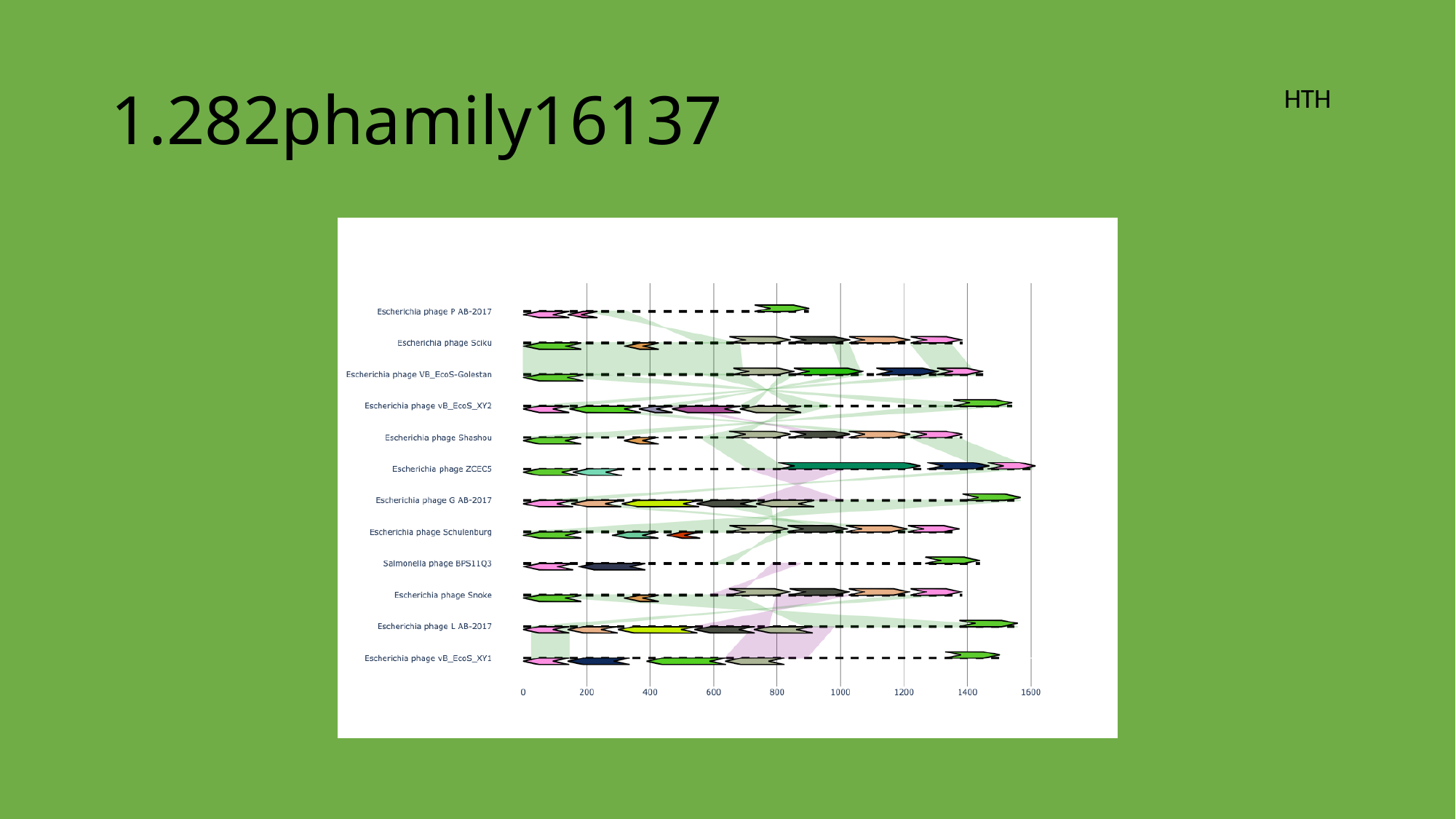

#### 1.282phamily16137
HTH

### Slide 42
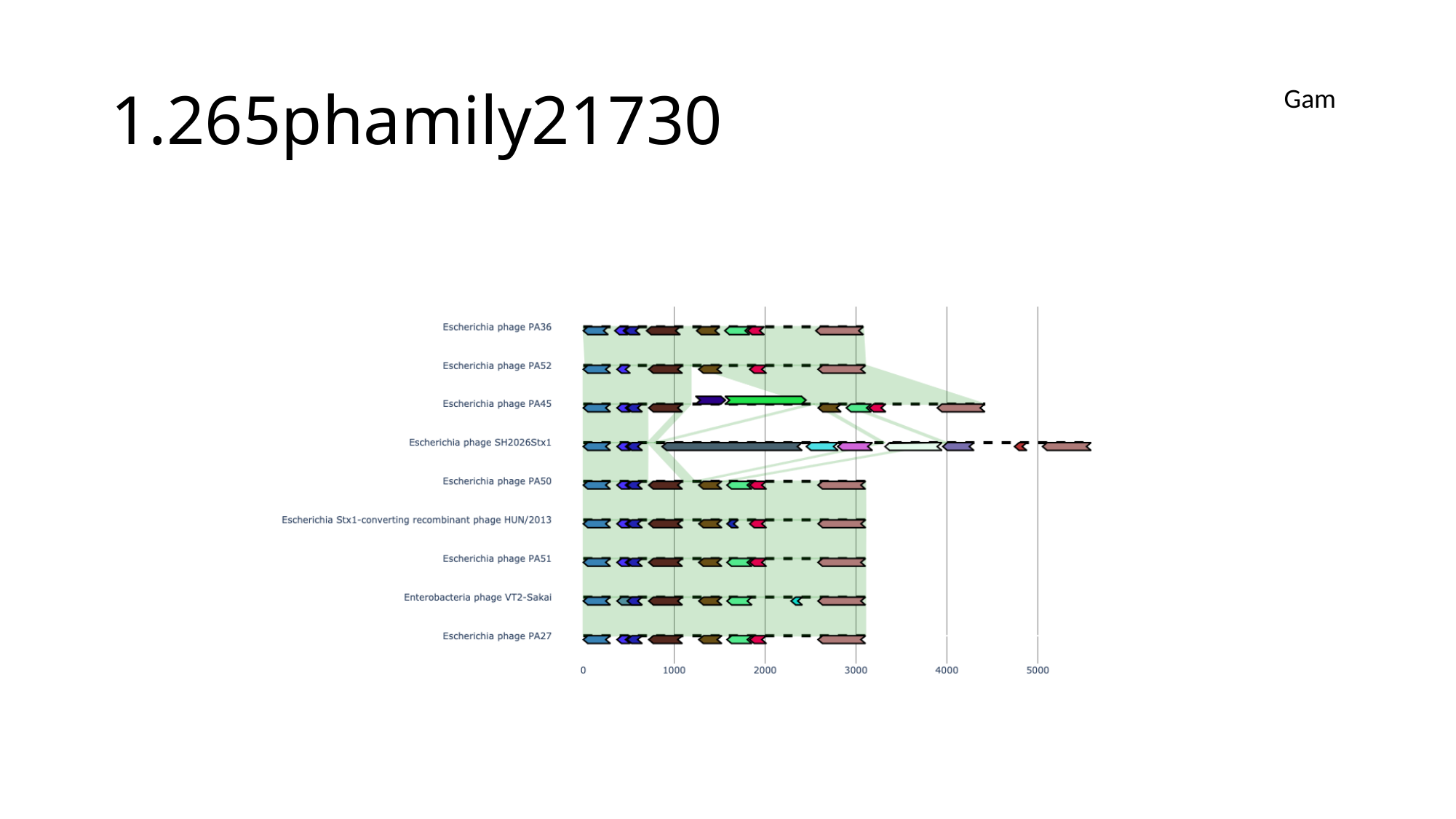

#### 1.265phamily21730
Gam

### Slide 43
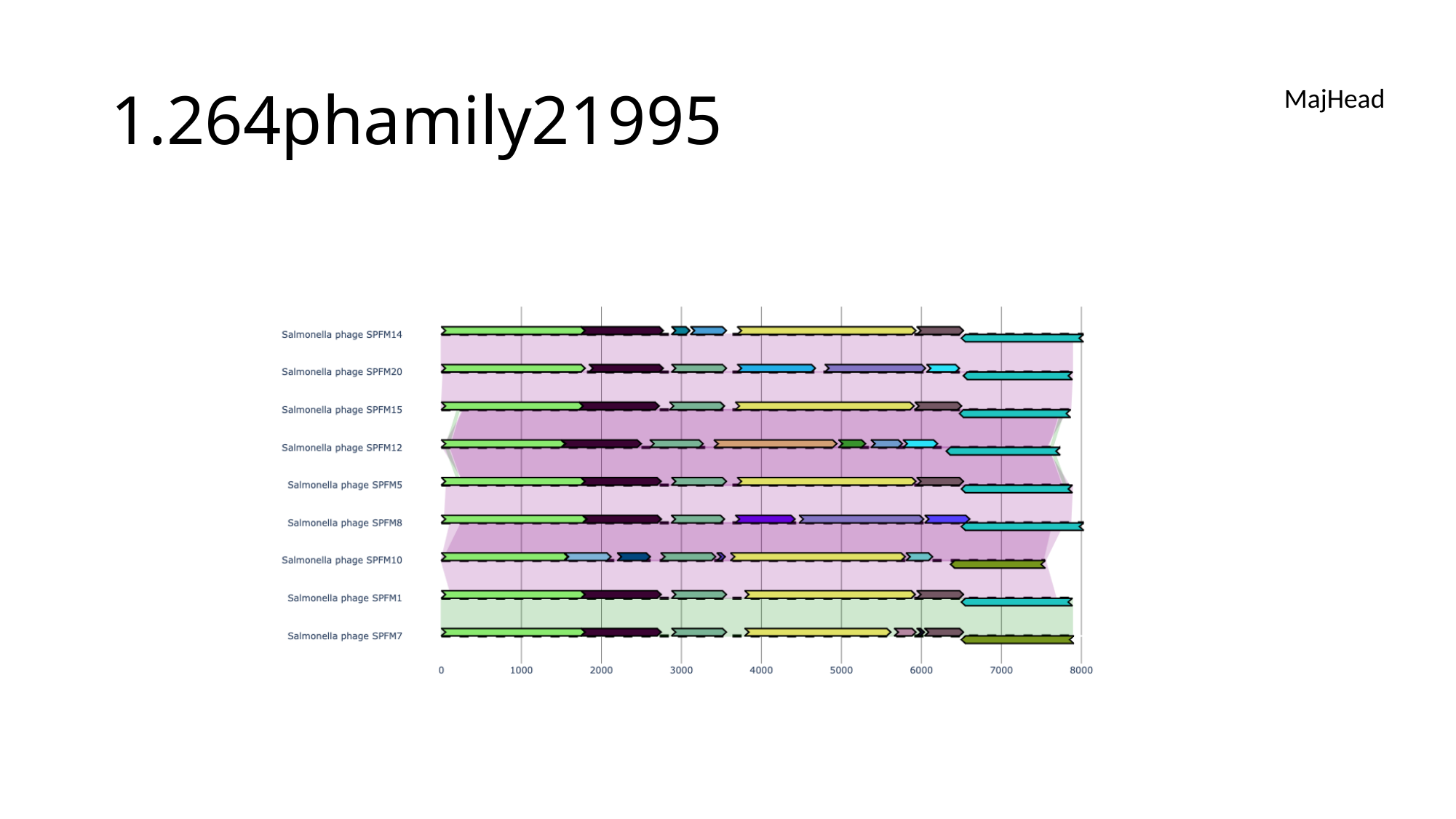

#### 1.264phamily21995
MajHead

### Slide 44
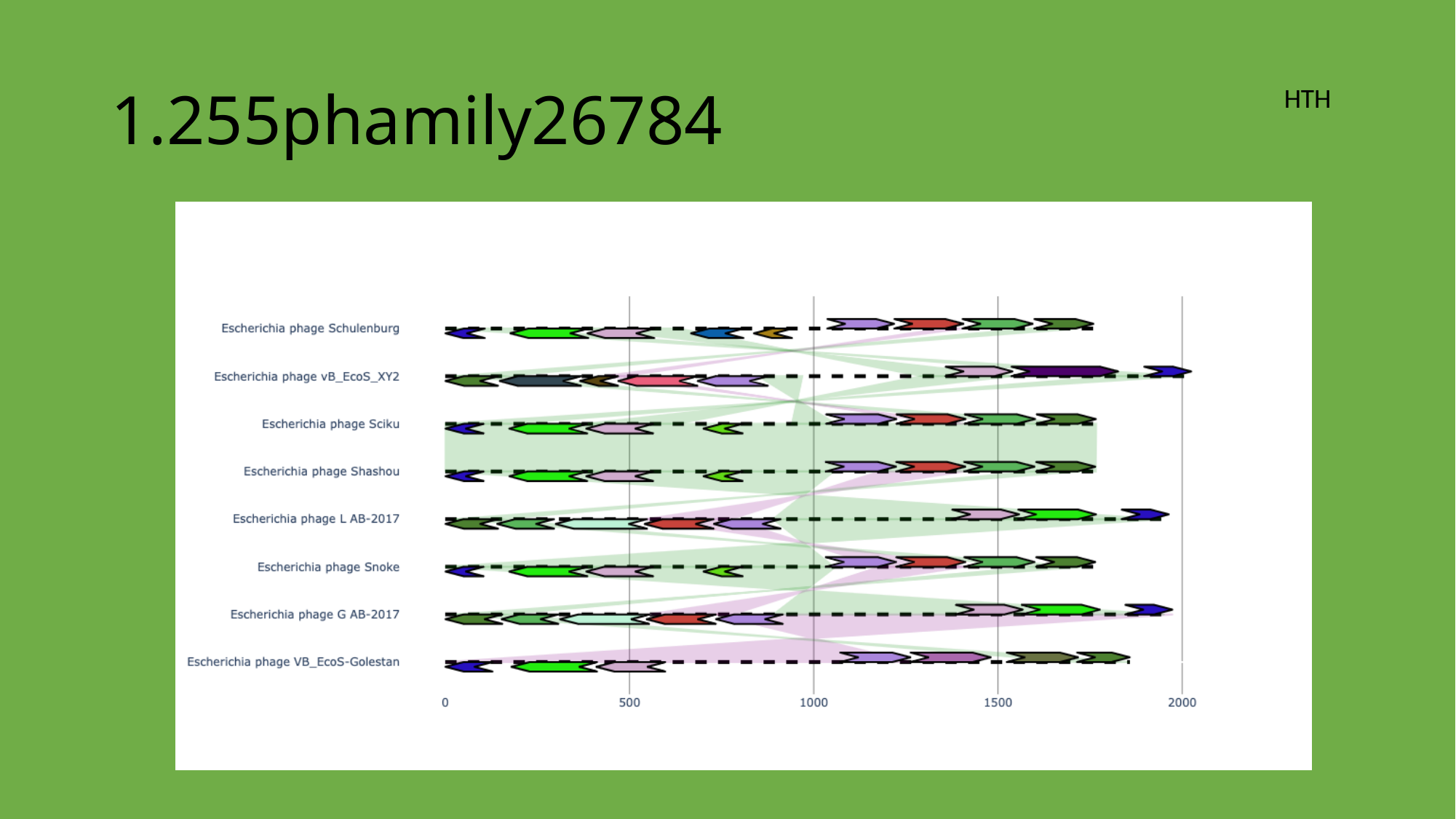

#### 1.255phamily26784
HTH

### Slide 45
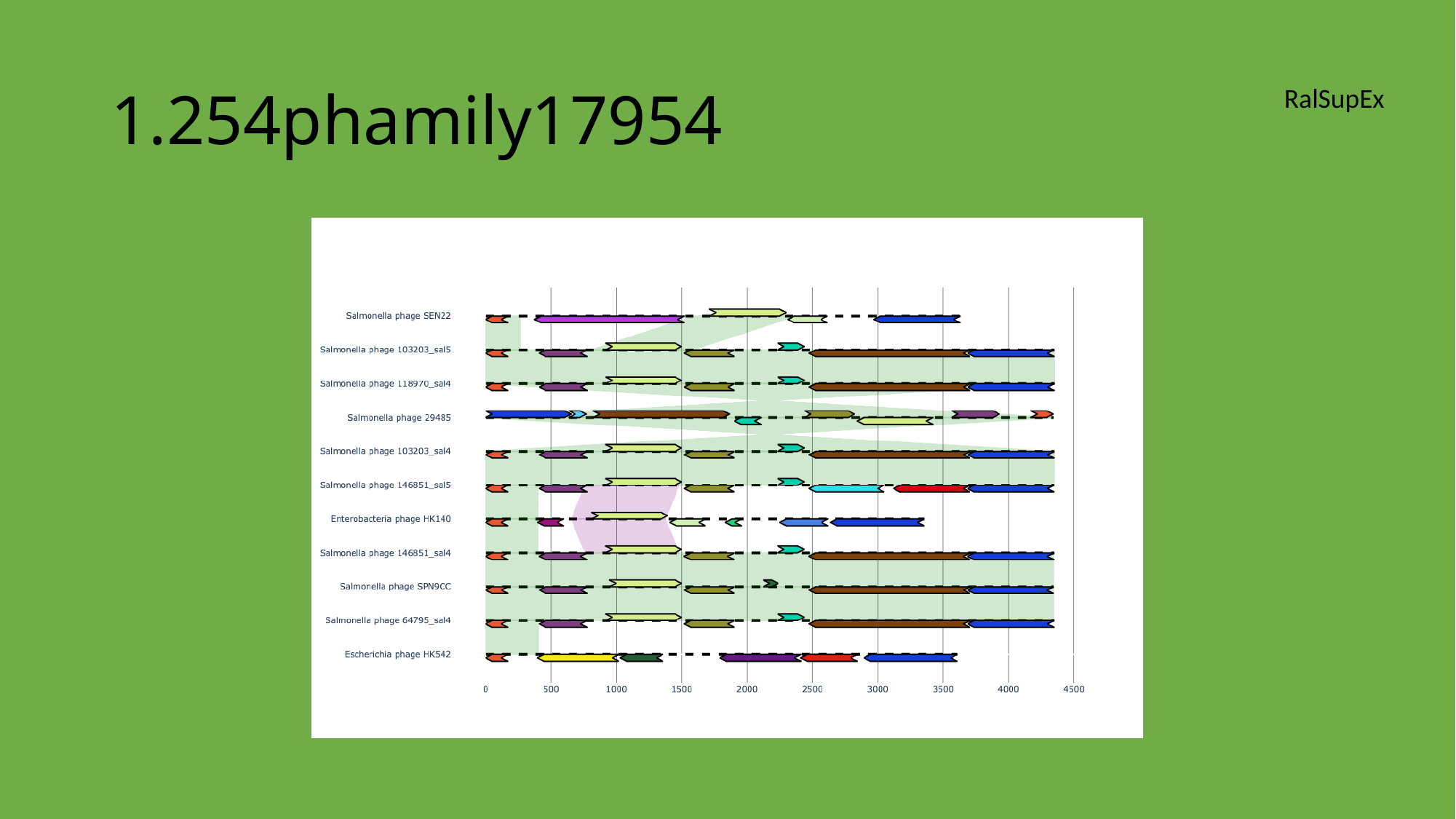

#### 1.254phamily17954
RalSupEx

### Slide 46
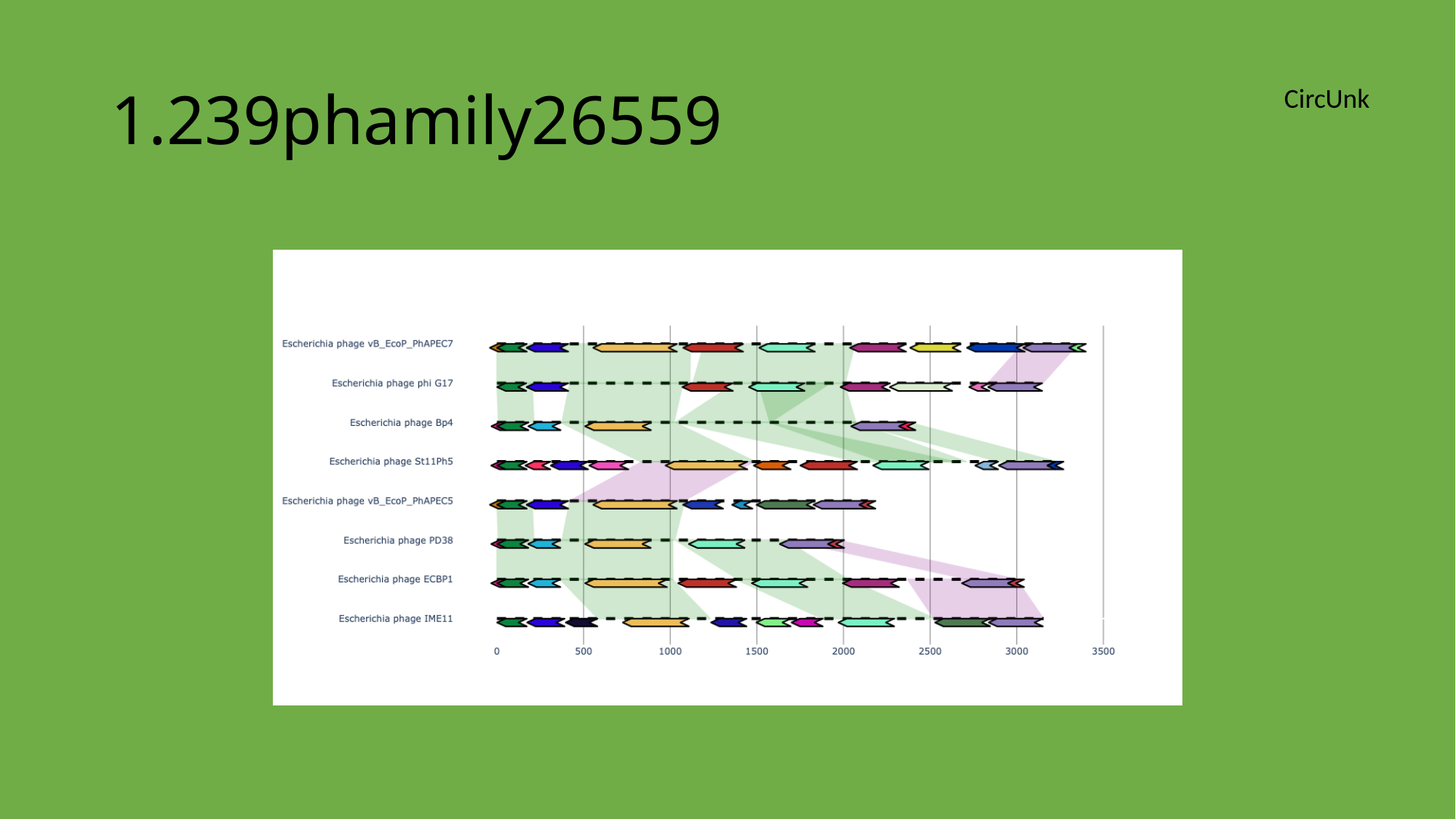

#### 1.239phamily26559
CircUnk

### Slide 47
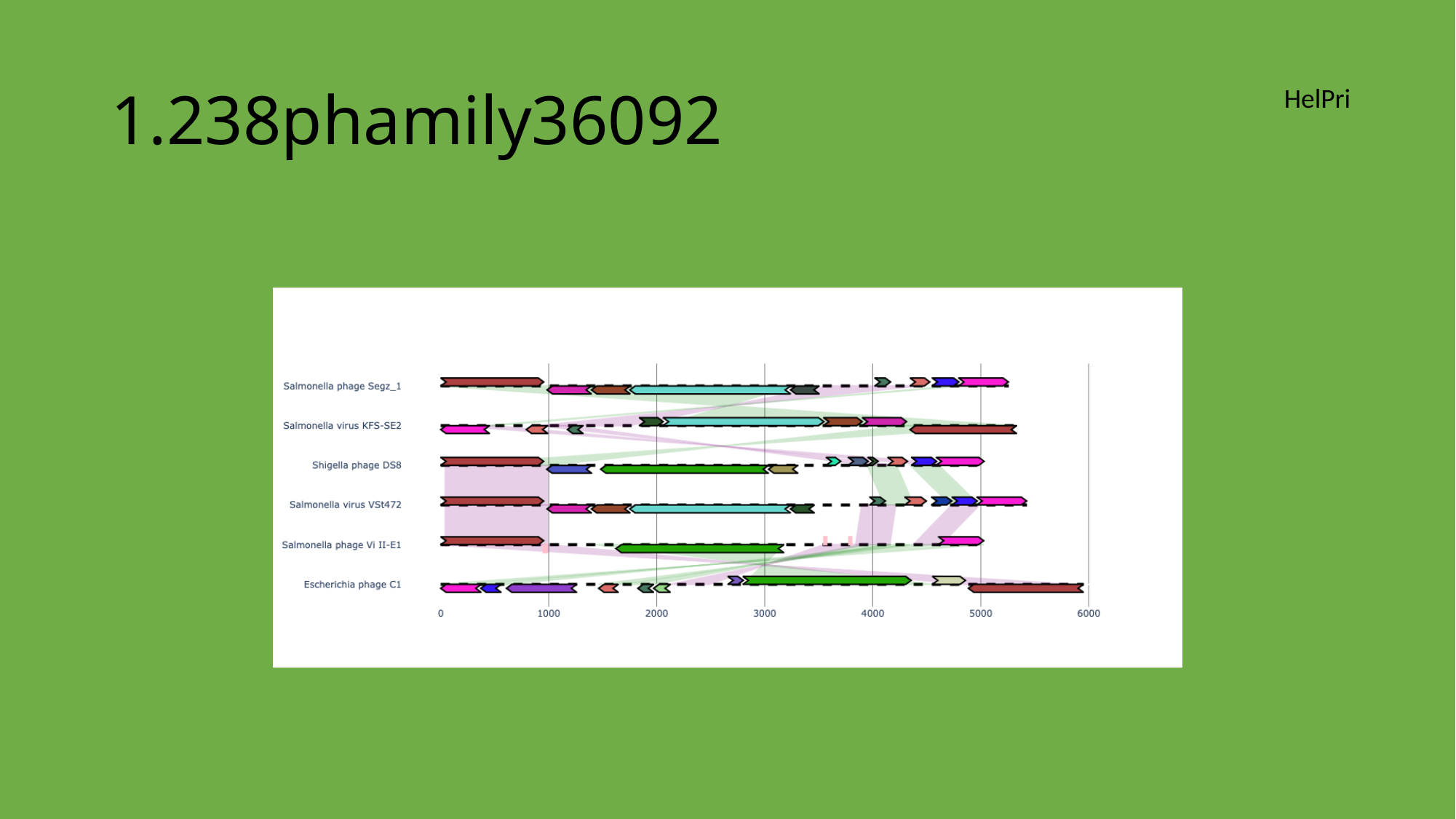

#### 1.238phamily36092
HelPri

### Slide 48
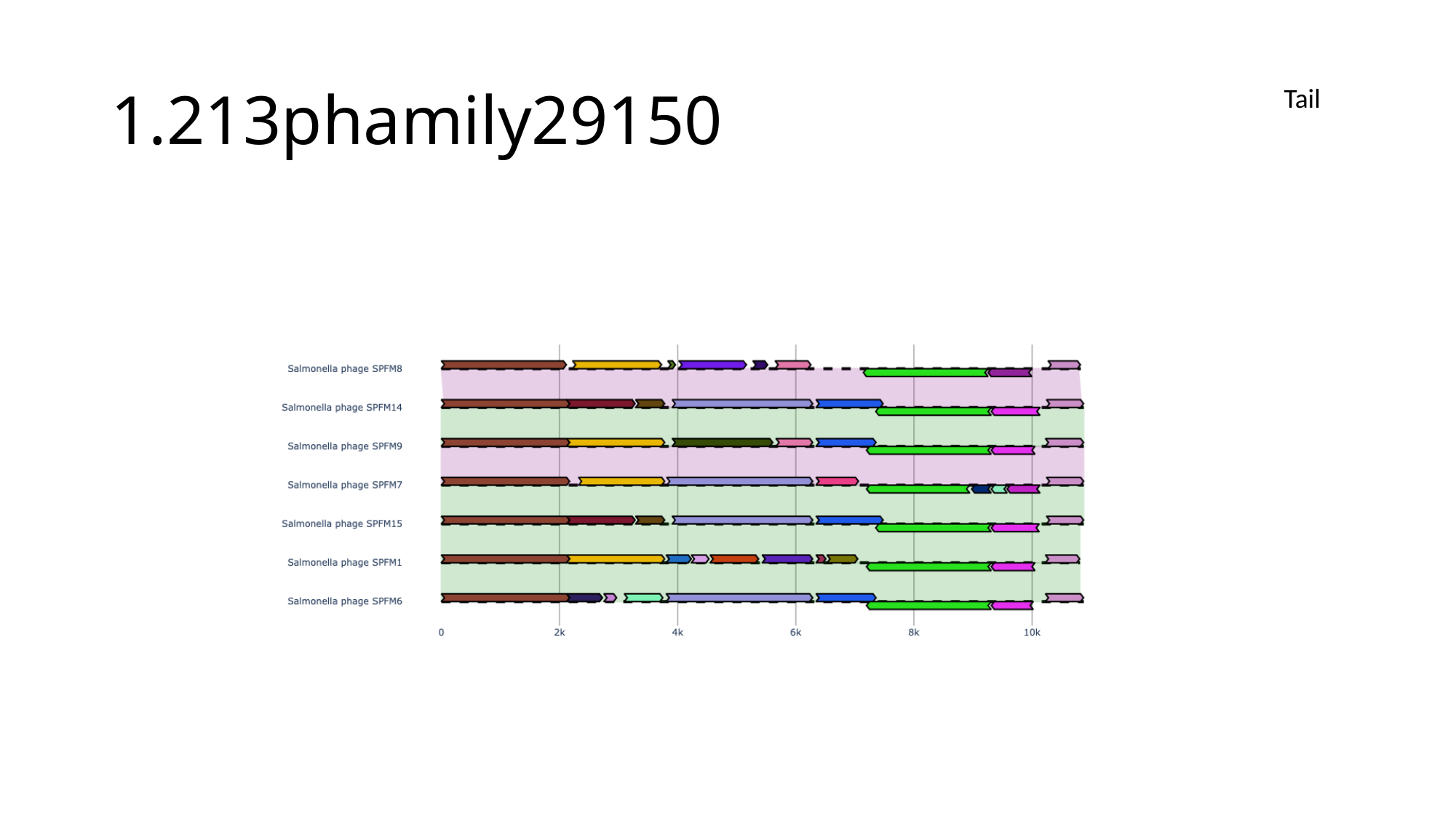

#### 1.213phamily29150
Tail

### Slide 49
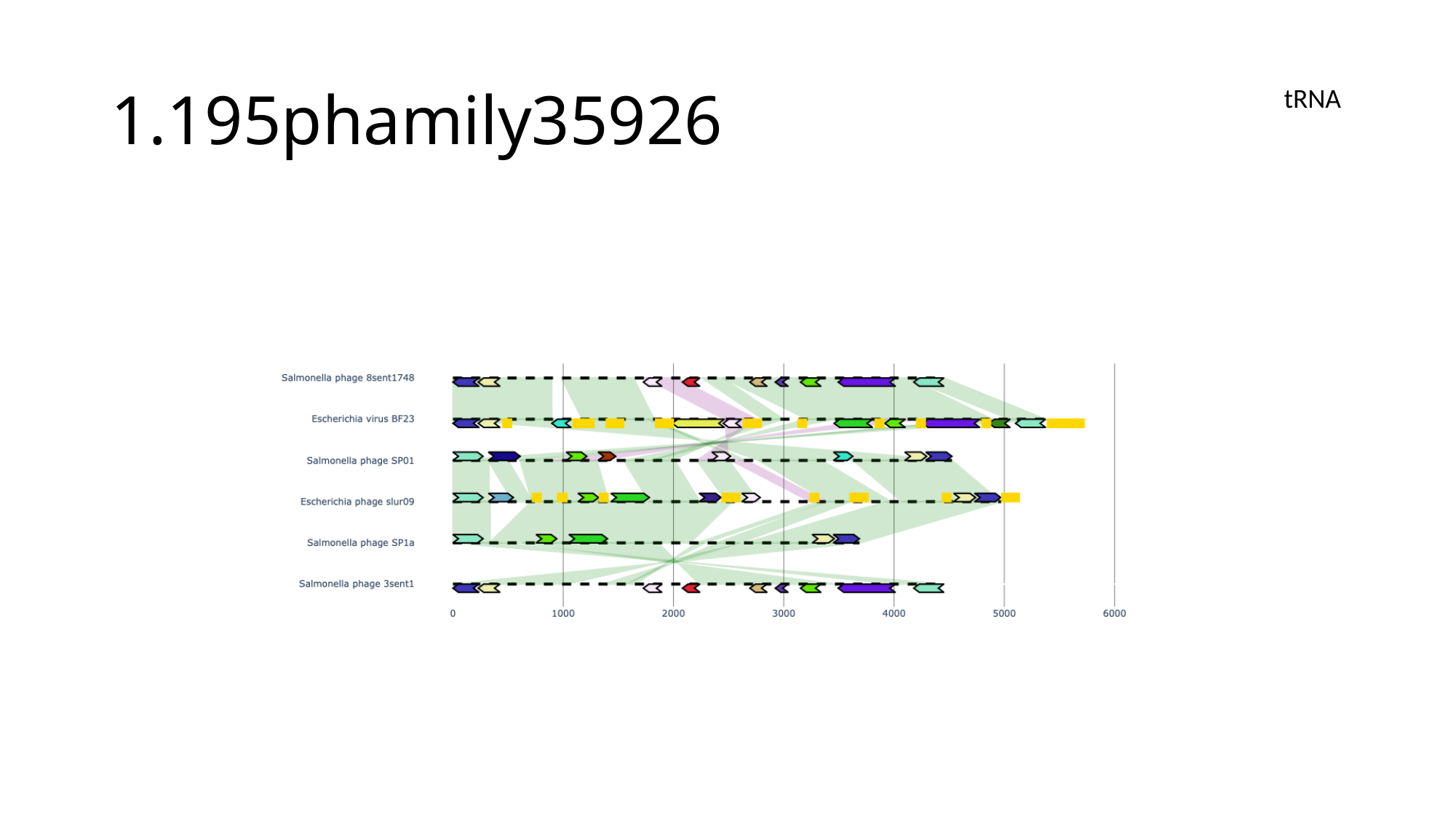

#### 1.195phamily35926
tRNA

### Slide 50
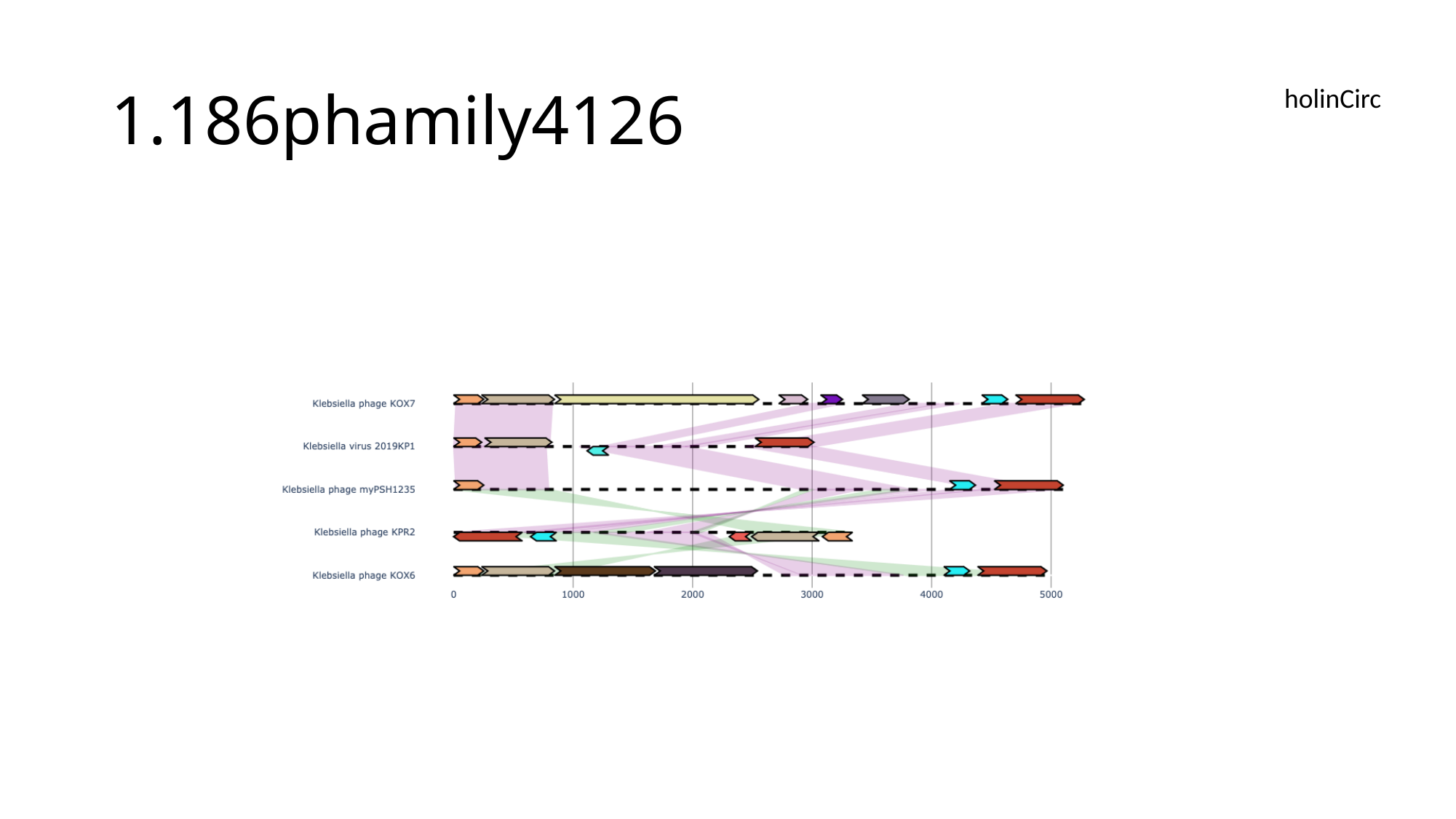

#### 1.186phamily4126
holinCirc

### Slide 51
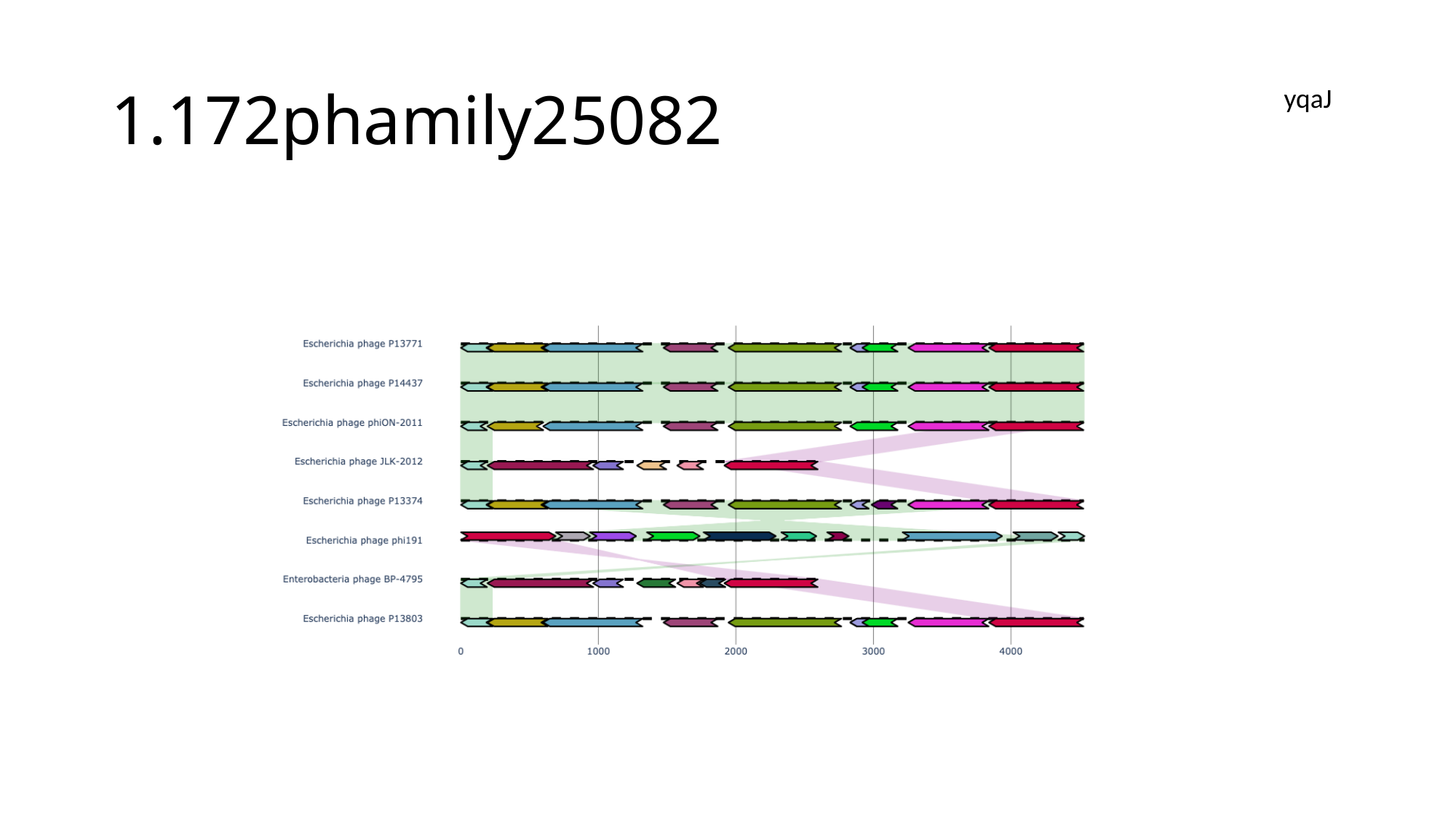

#### 1.172phamily25082
yqaJ

### Slide 52
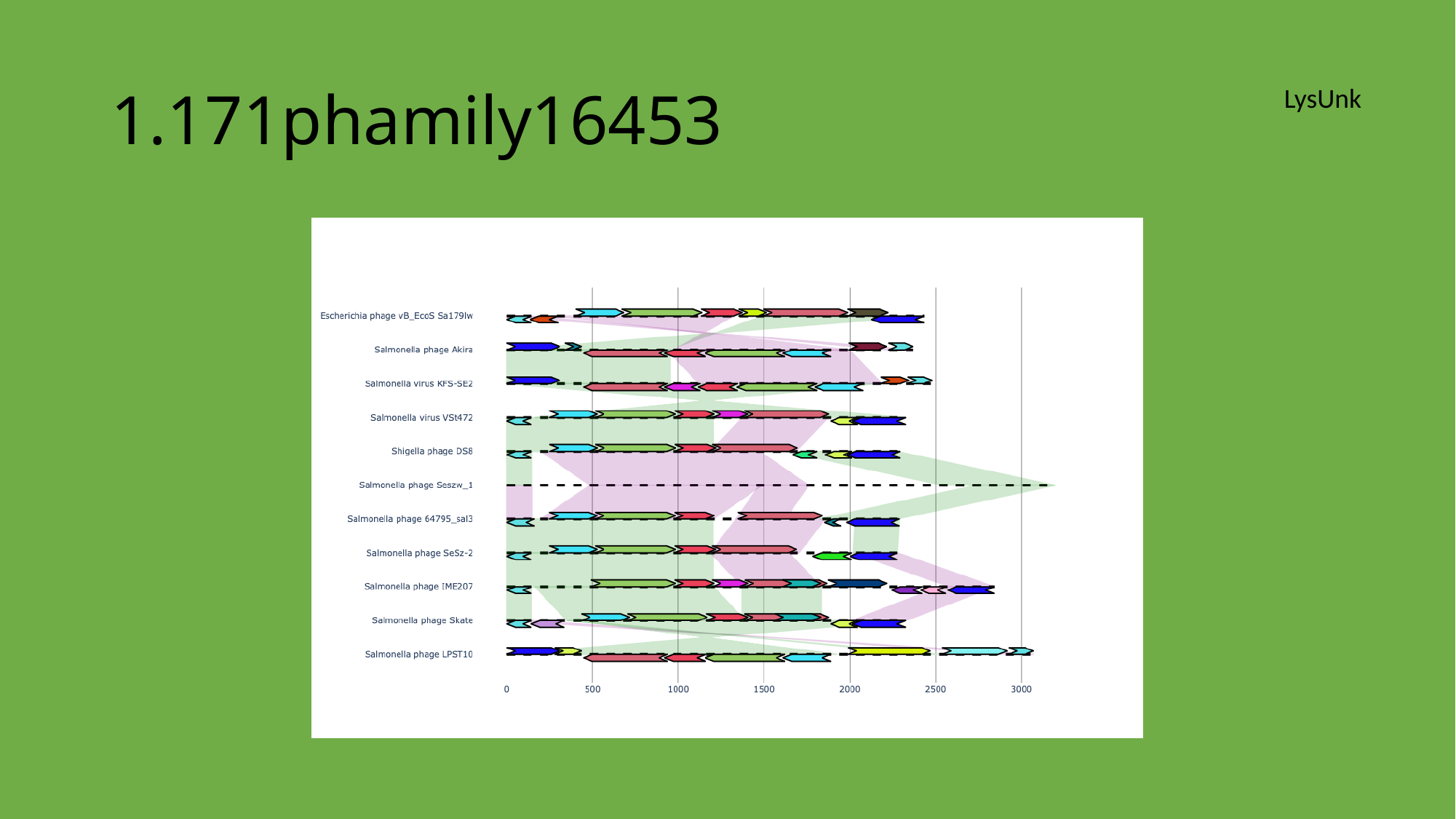

#### 1.171phamily16453
LysUnk

### Slide 53
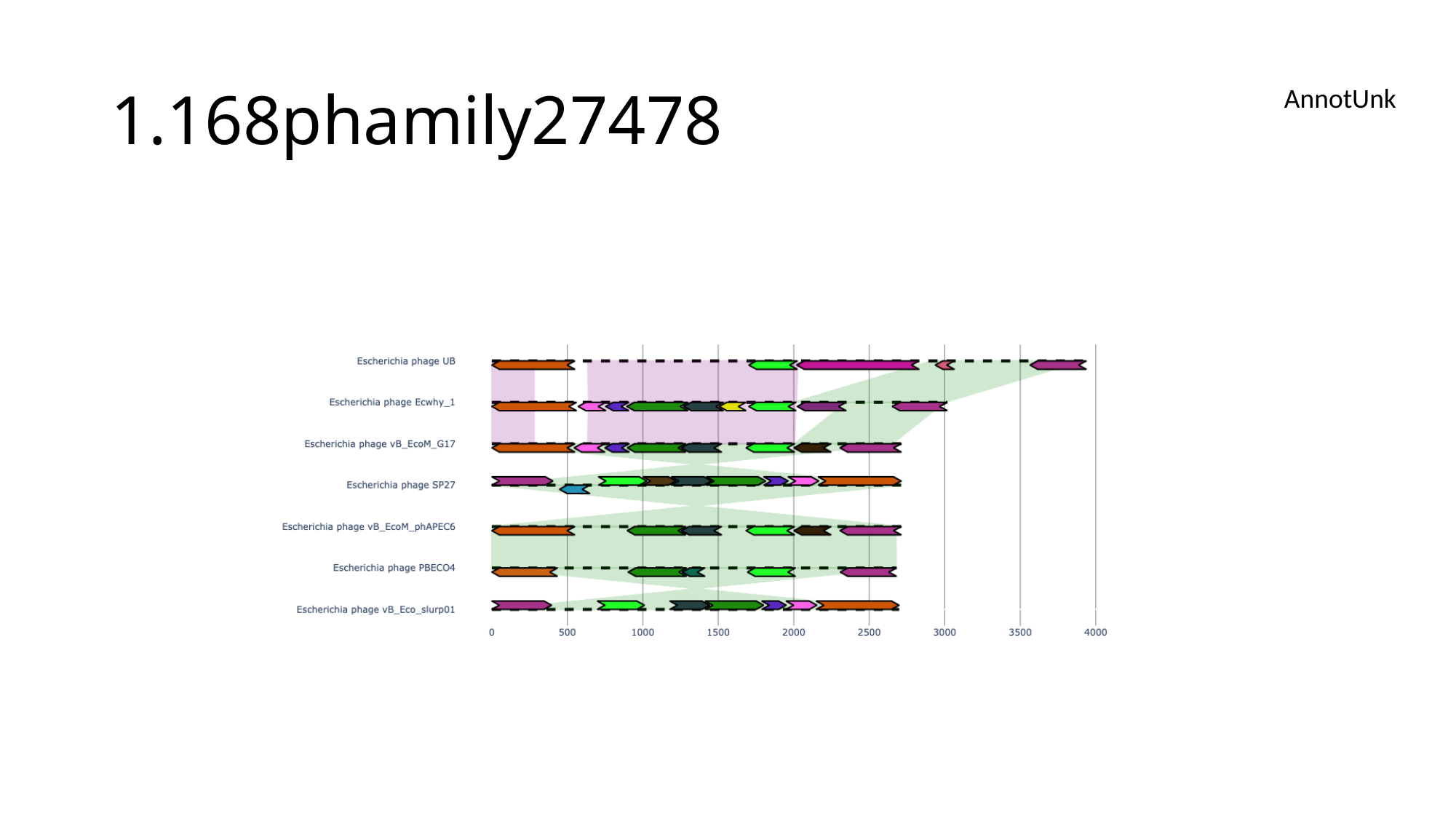

#### 1.168phamily27478
AnnotUnk

### Slide 54
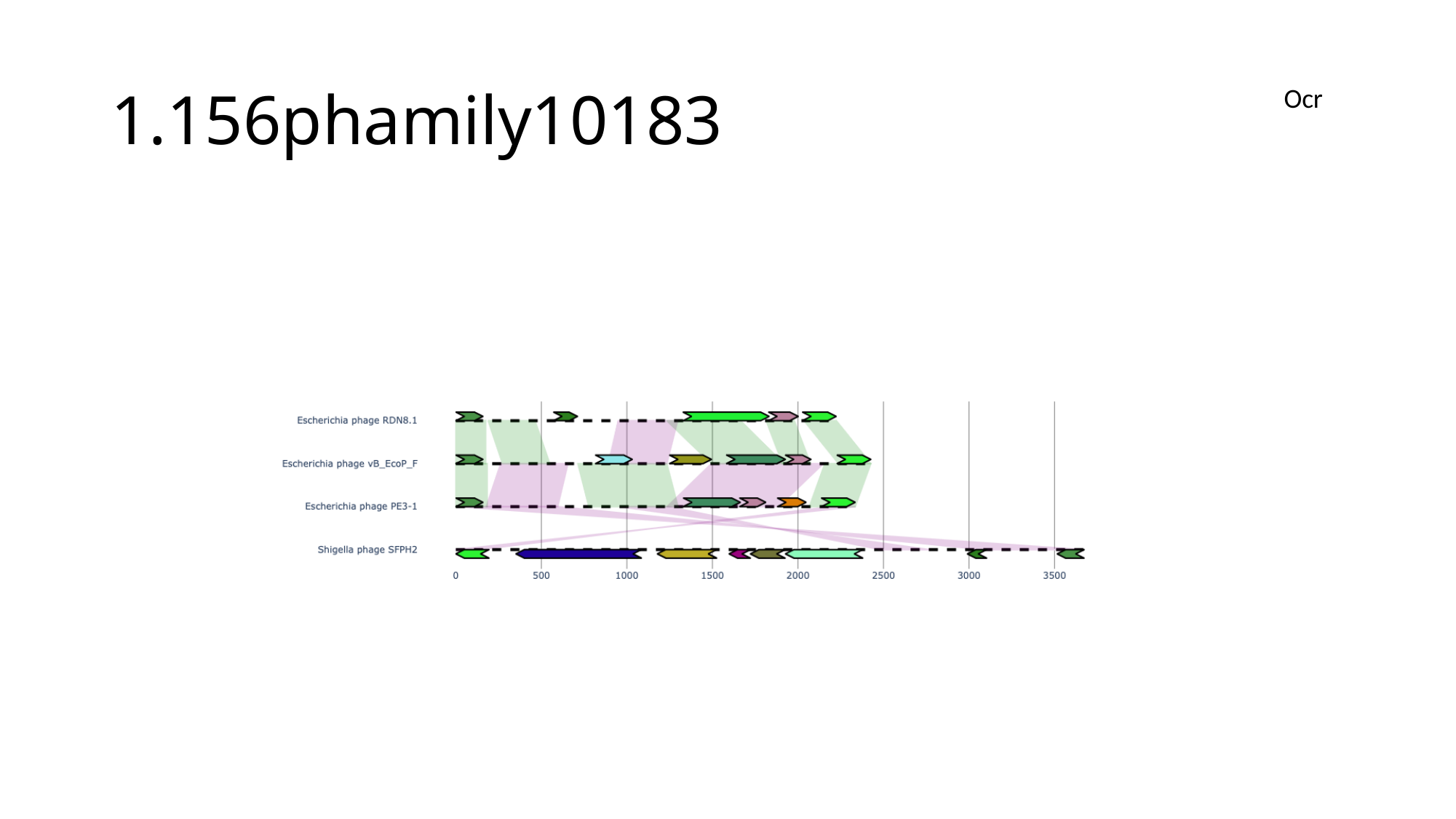

#### 1.156phamily10183
Ocr

### Slide 55
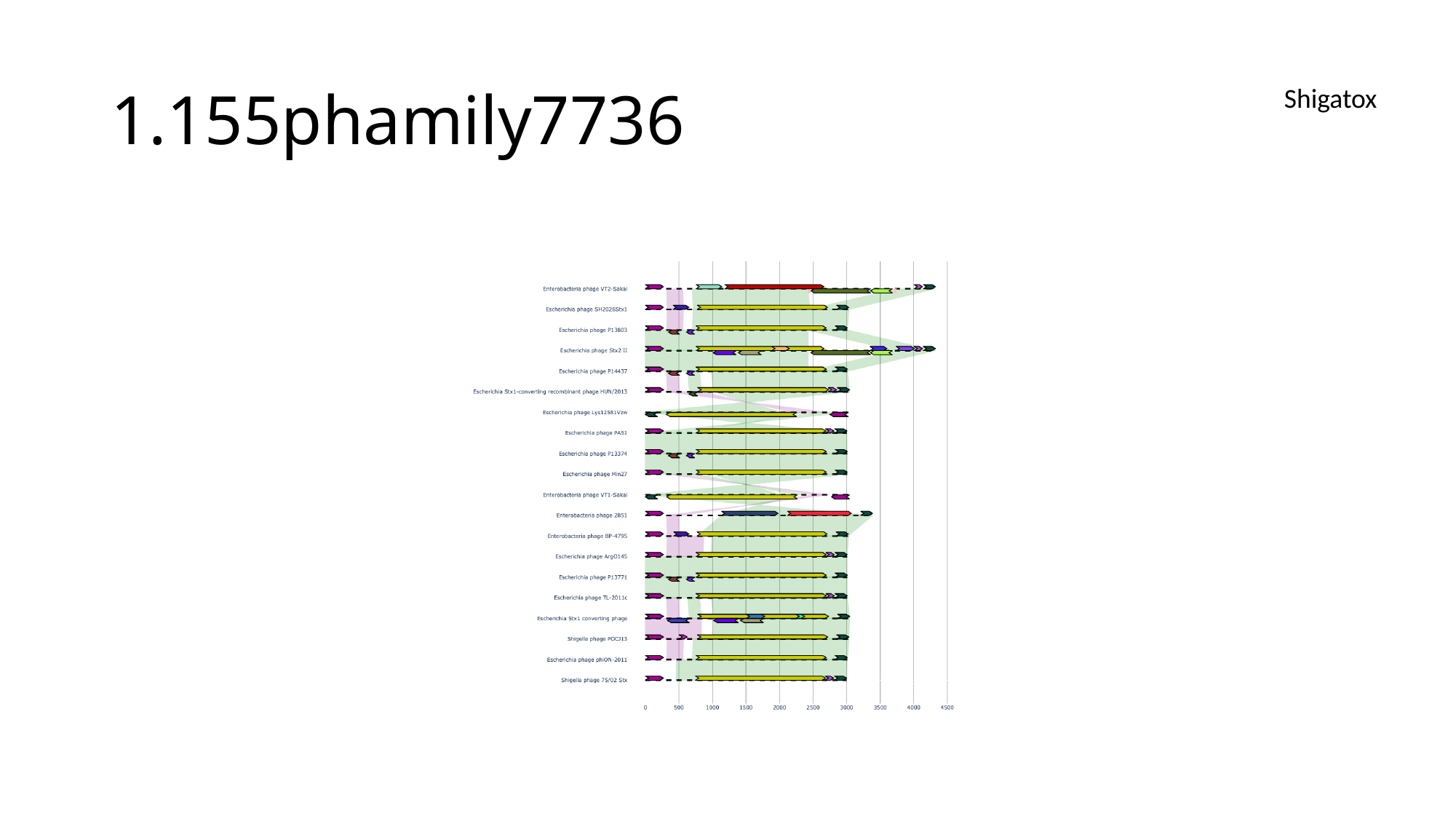

#### 1.155phamily7736
Shigatox

### Slide 56
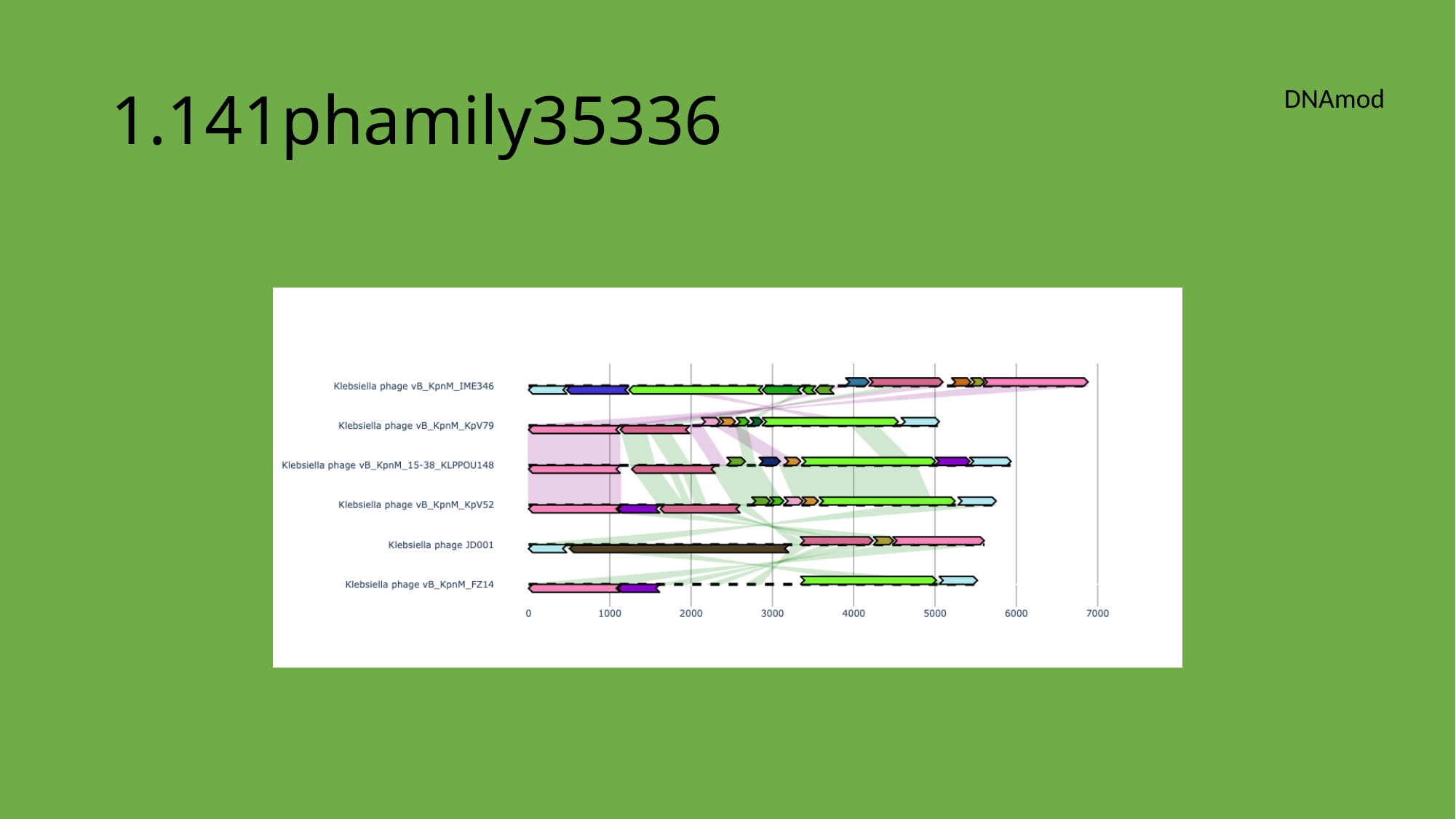

#### 1.141phamily35336
DNAmod

### Slide 57
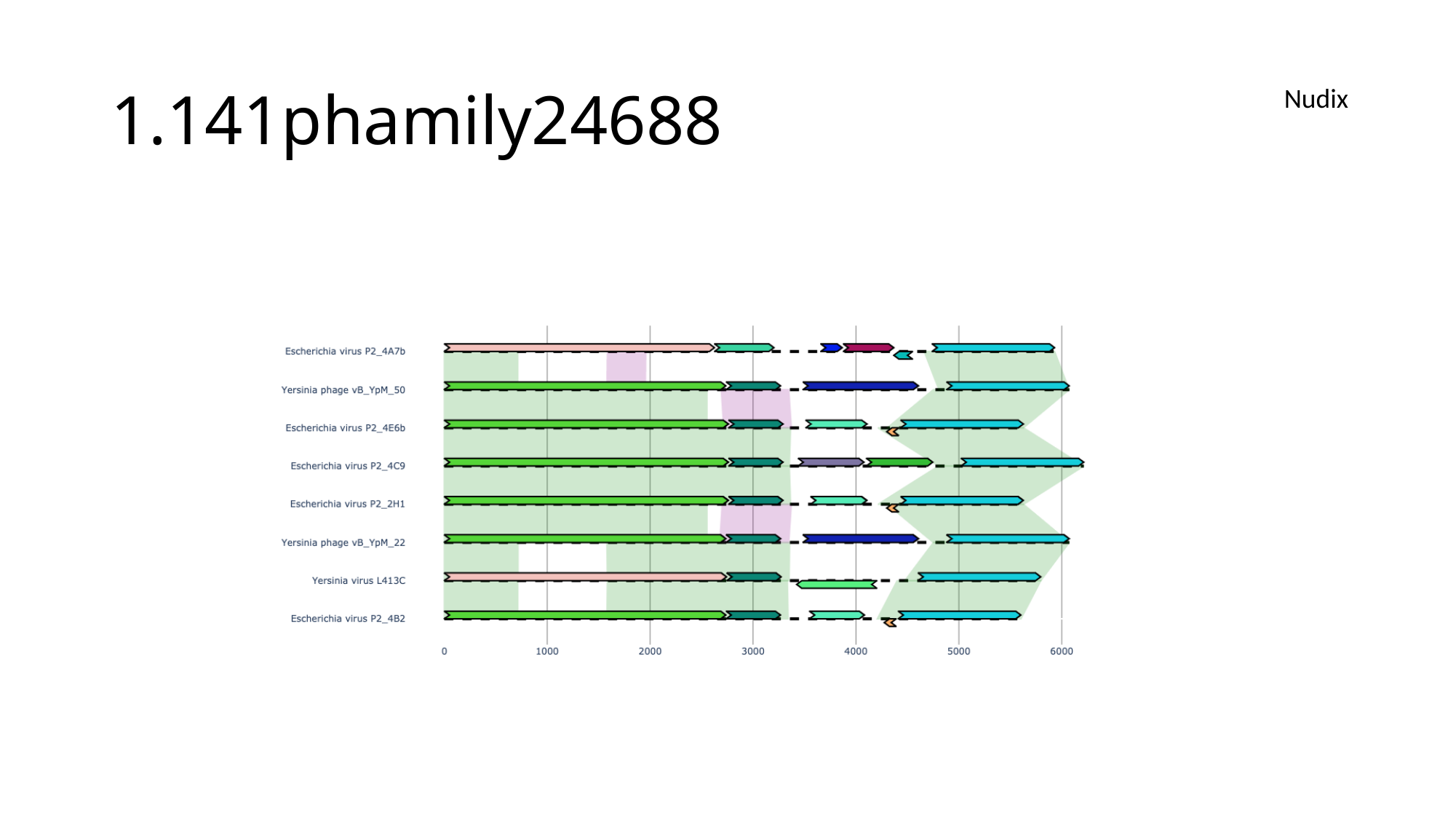

#### 1.141phamily24688
Nudix

### Slide 58
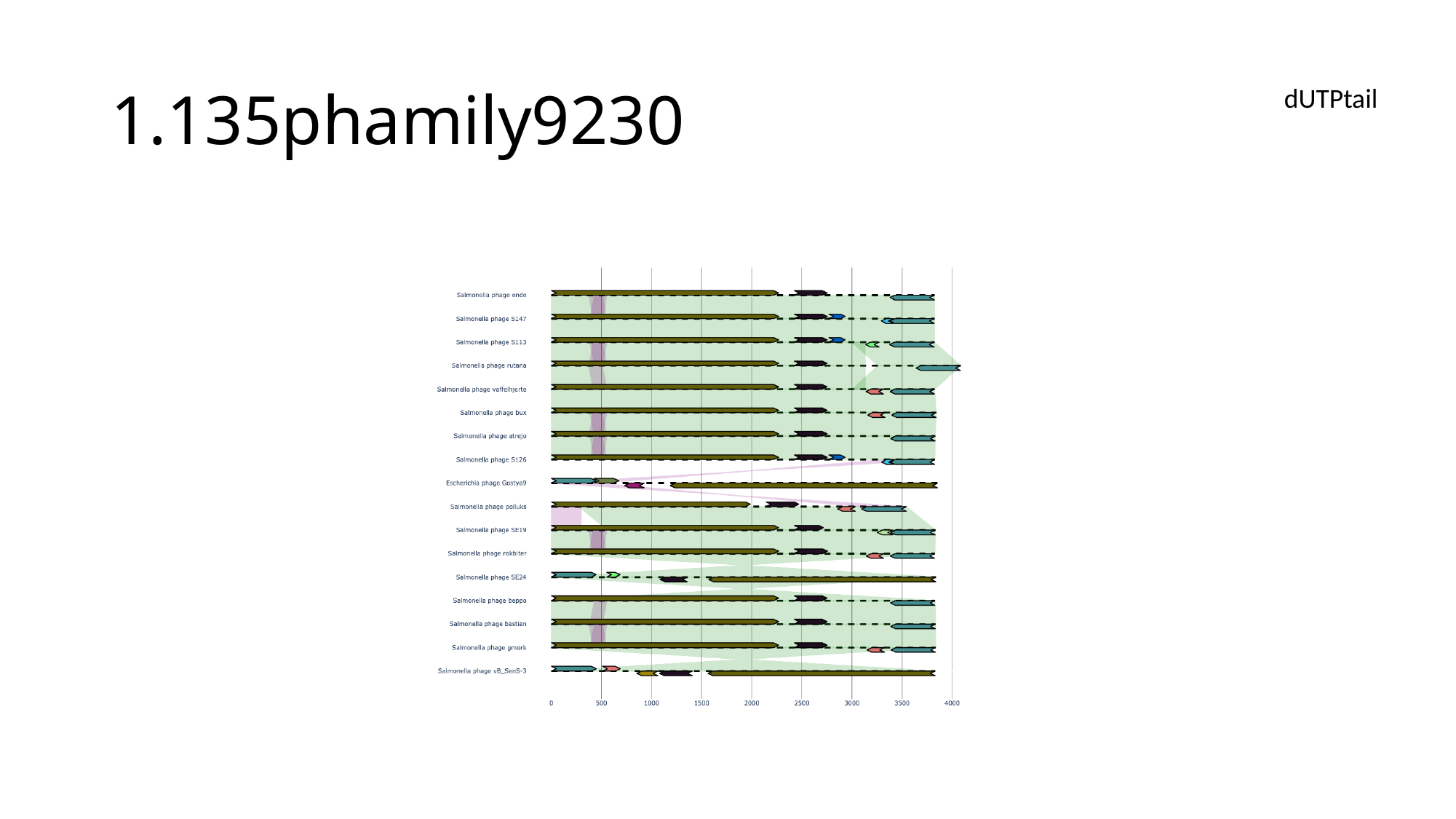

#### 1.135phamily9230
dUTPtail

### Slide 59
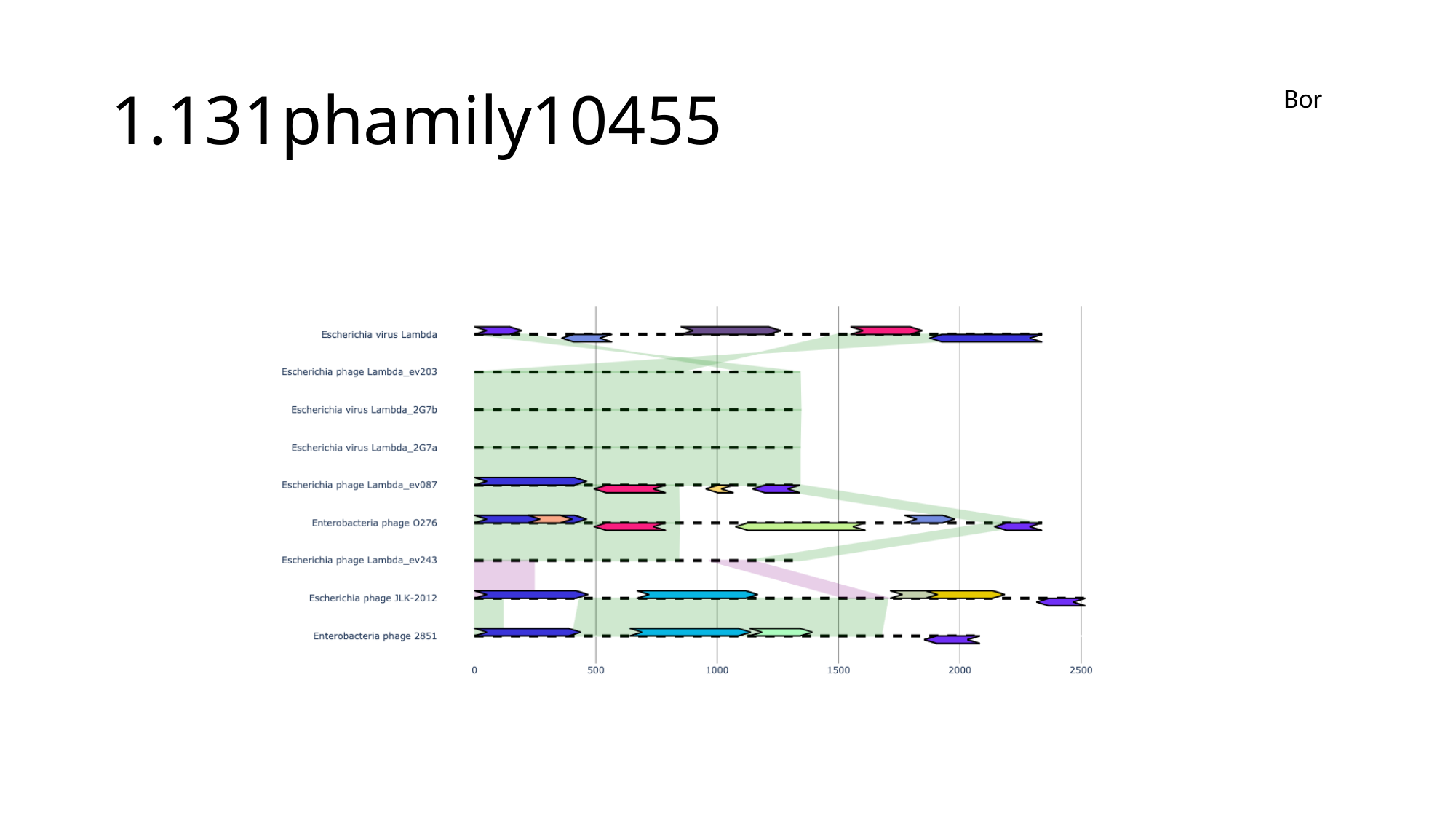

#### 1.131phamily10455
Bor

### Slide 60
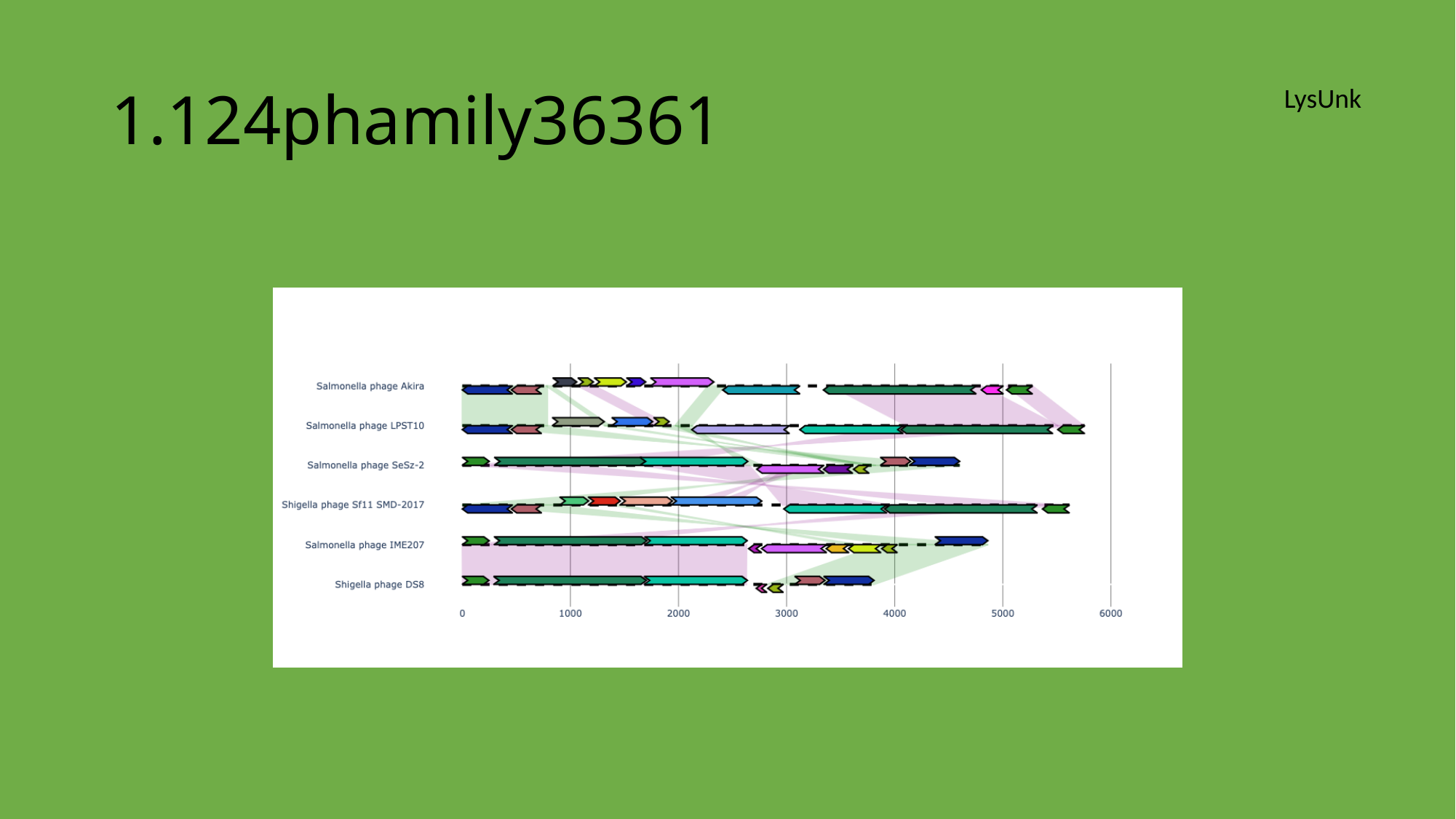

#### 1.124phamily36361
LysUnk

### Slide 61
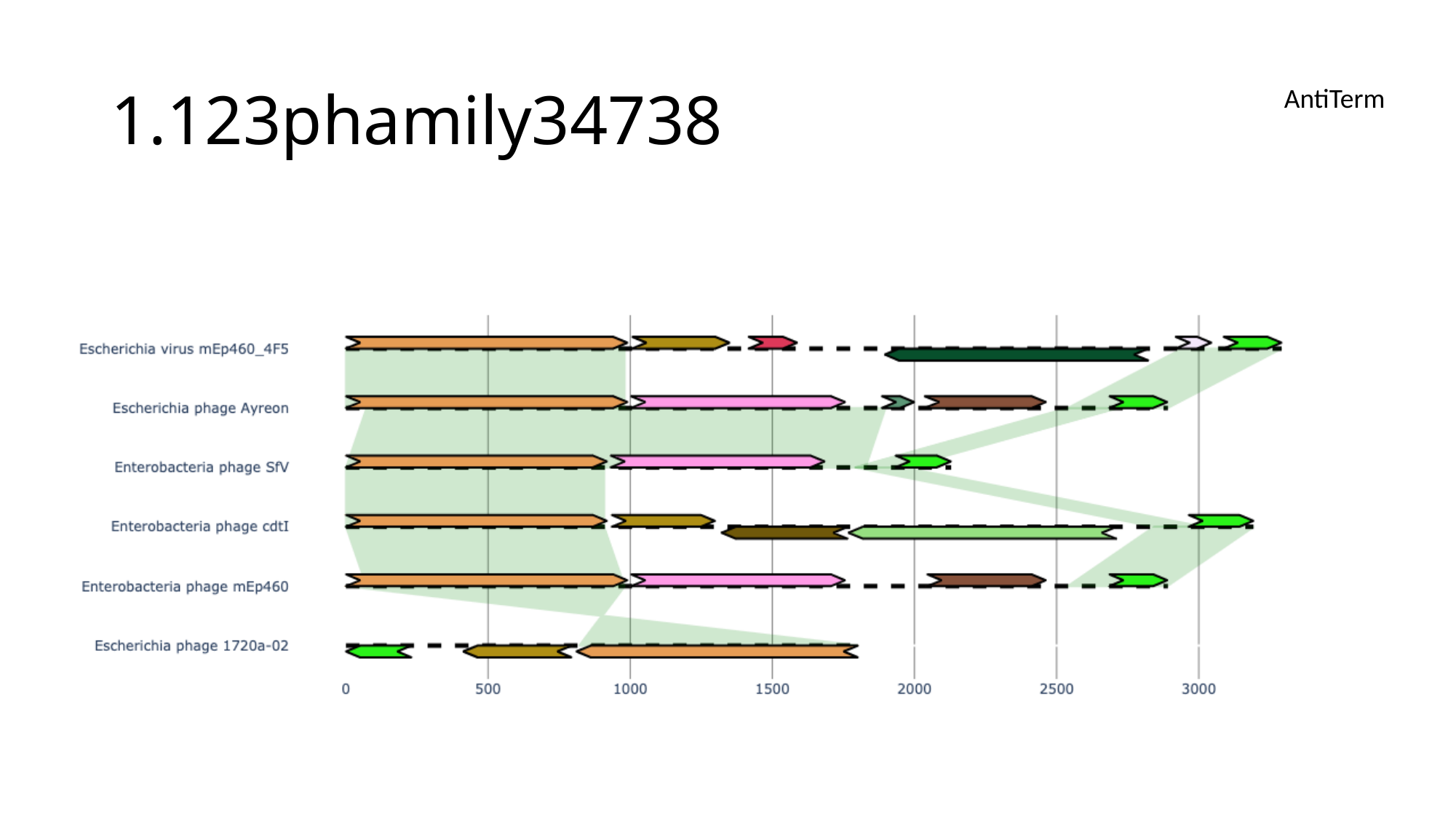

#### 1.123phamily34738
AntiTerm

### Slide 62
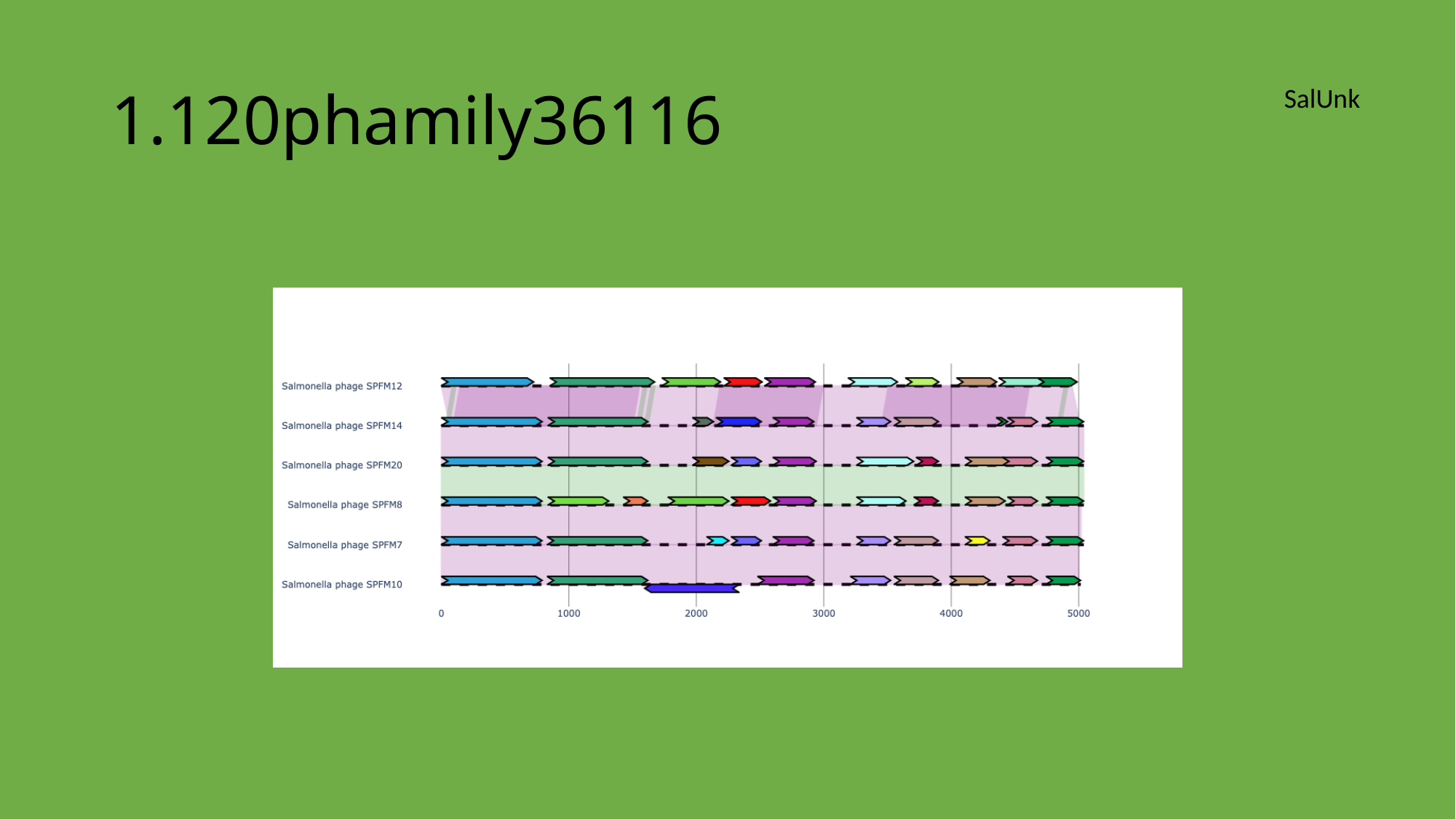

#### 1.120phamily36116
SalUnk

### Slide 63
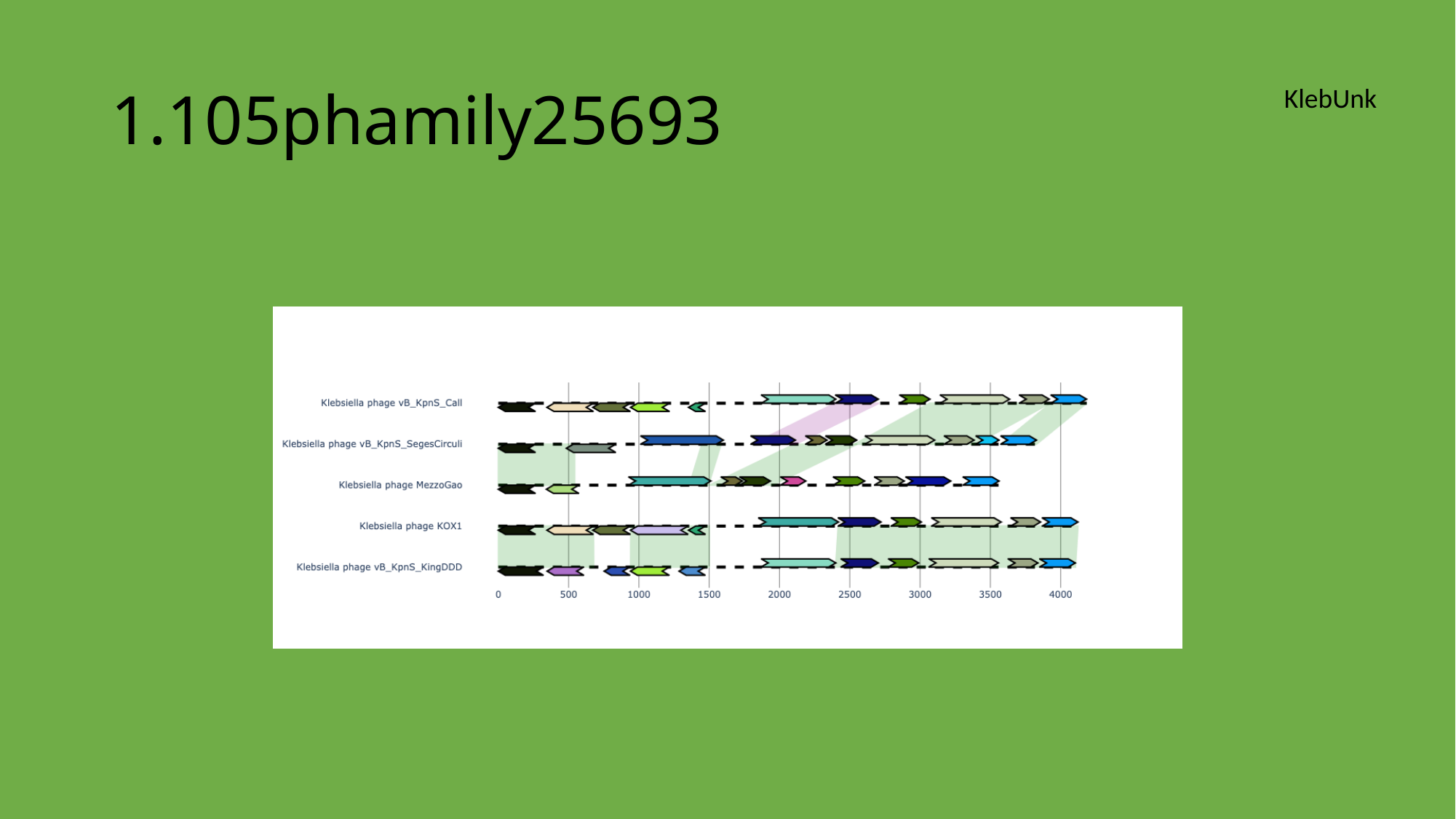

#### 1.105phamily25693
KlebUnk

### Slide 64
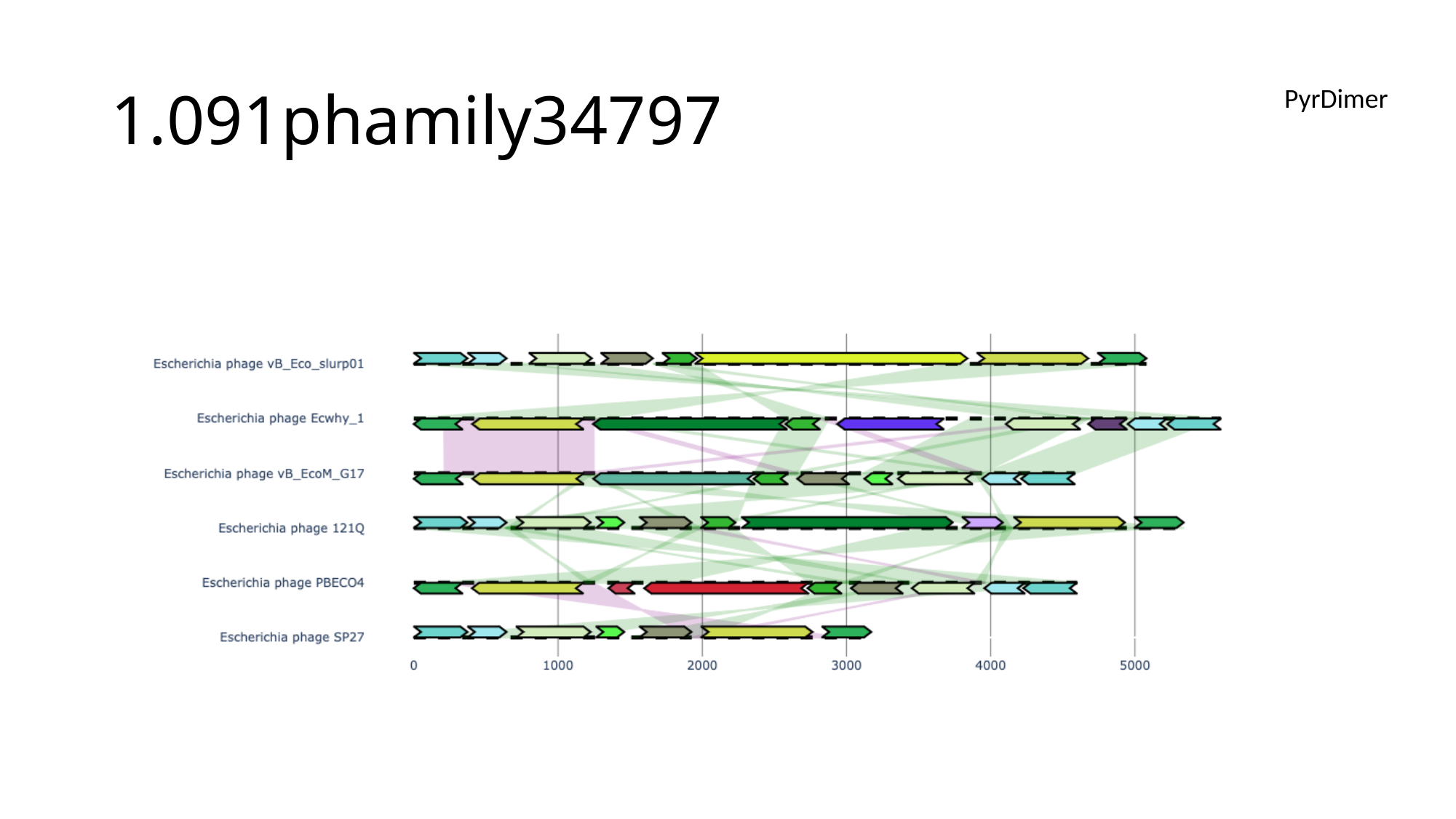

#### 1.091phamily34797
PyrDimer

### Slide 65
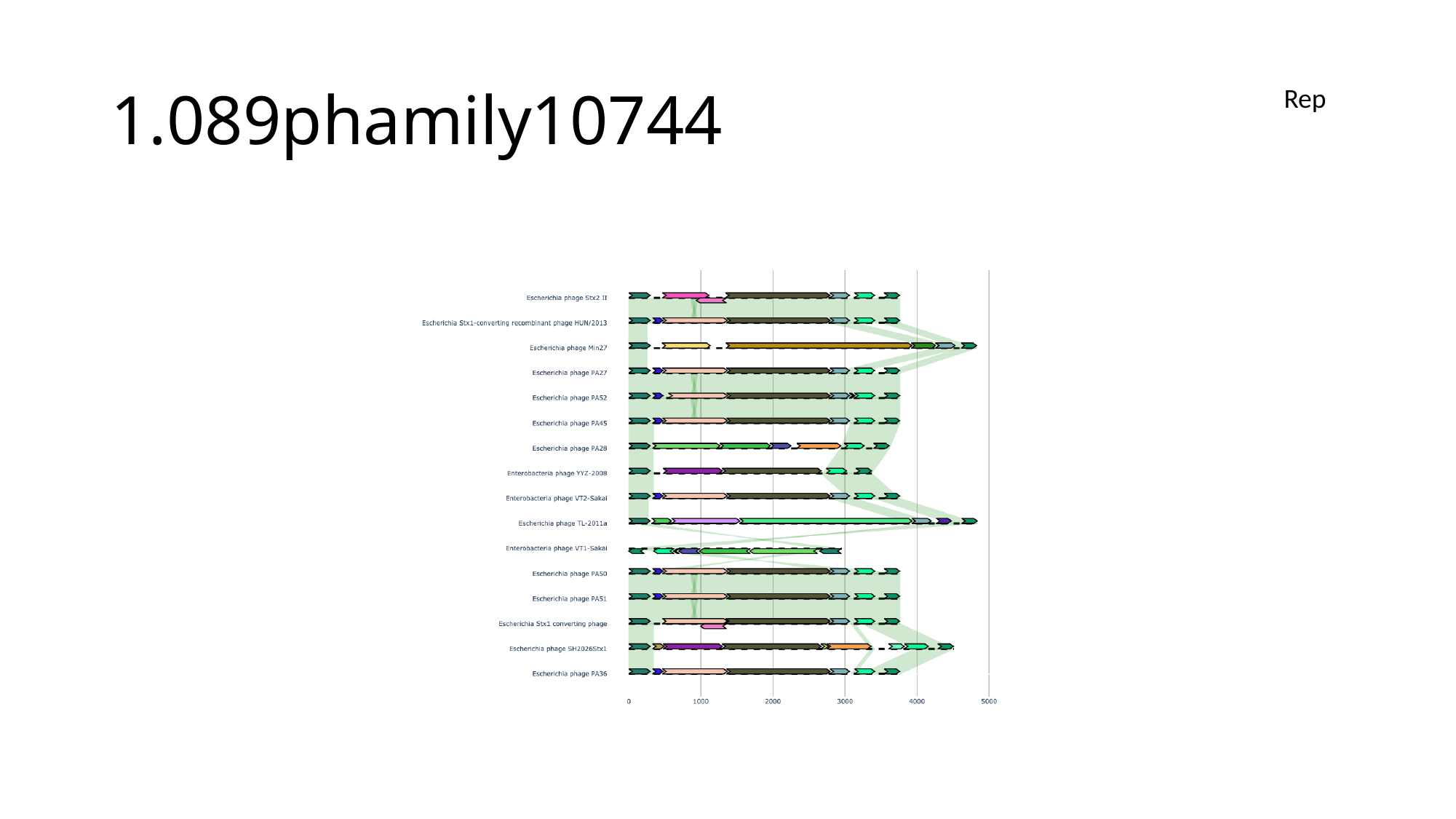

#### 1.089phamily10744
Rep

### Slide 66
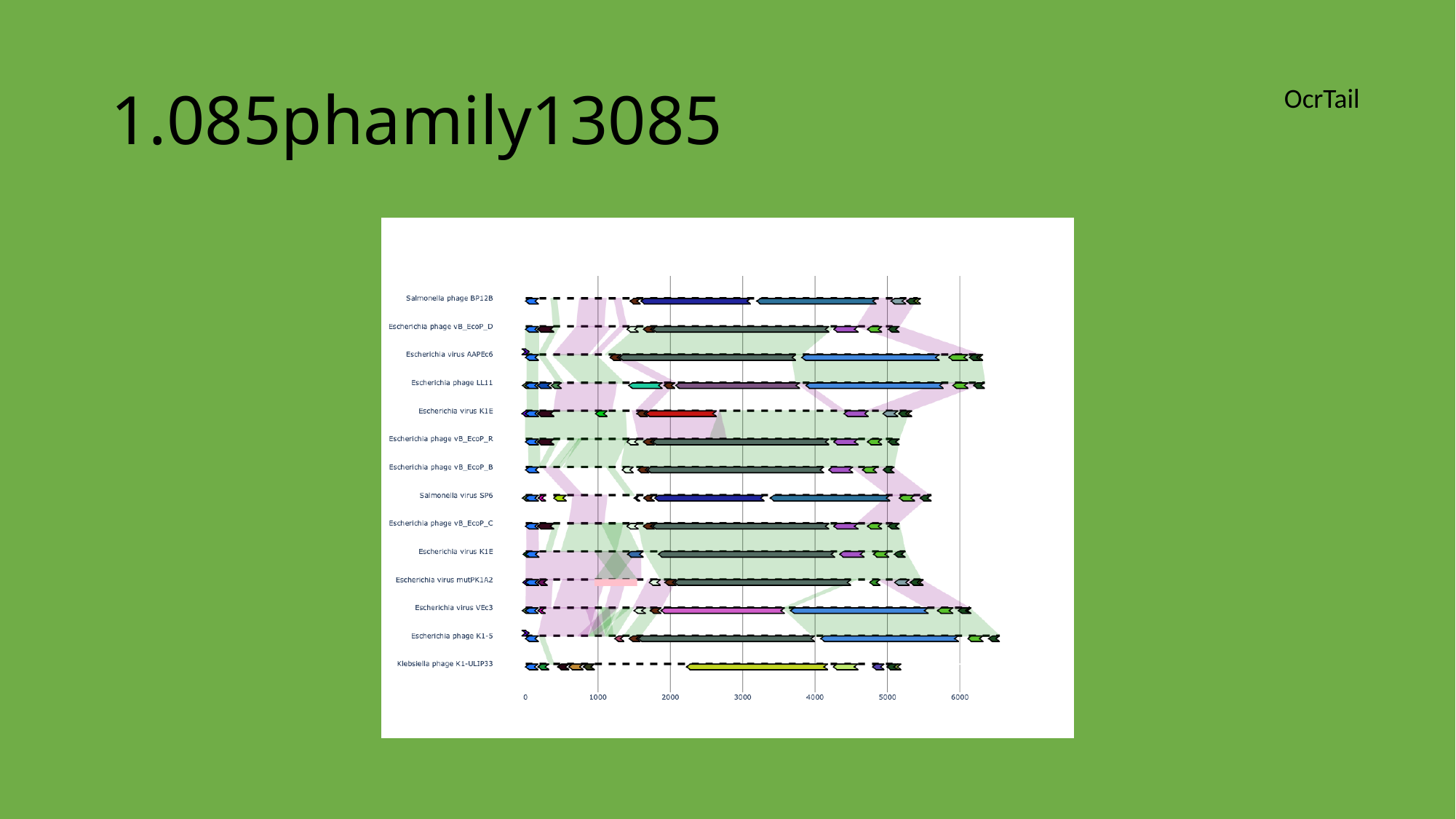

#### 1.085phamily13085
OcrTail

### Slide 67
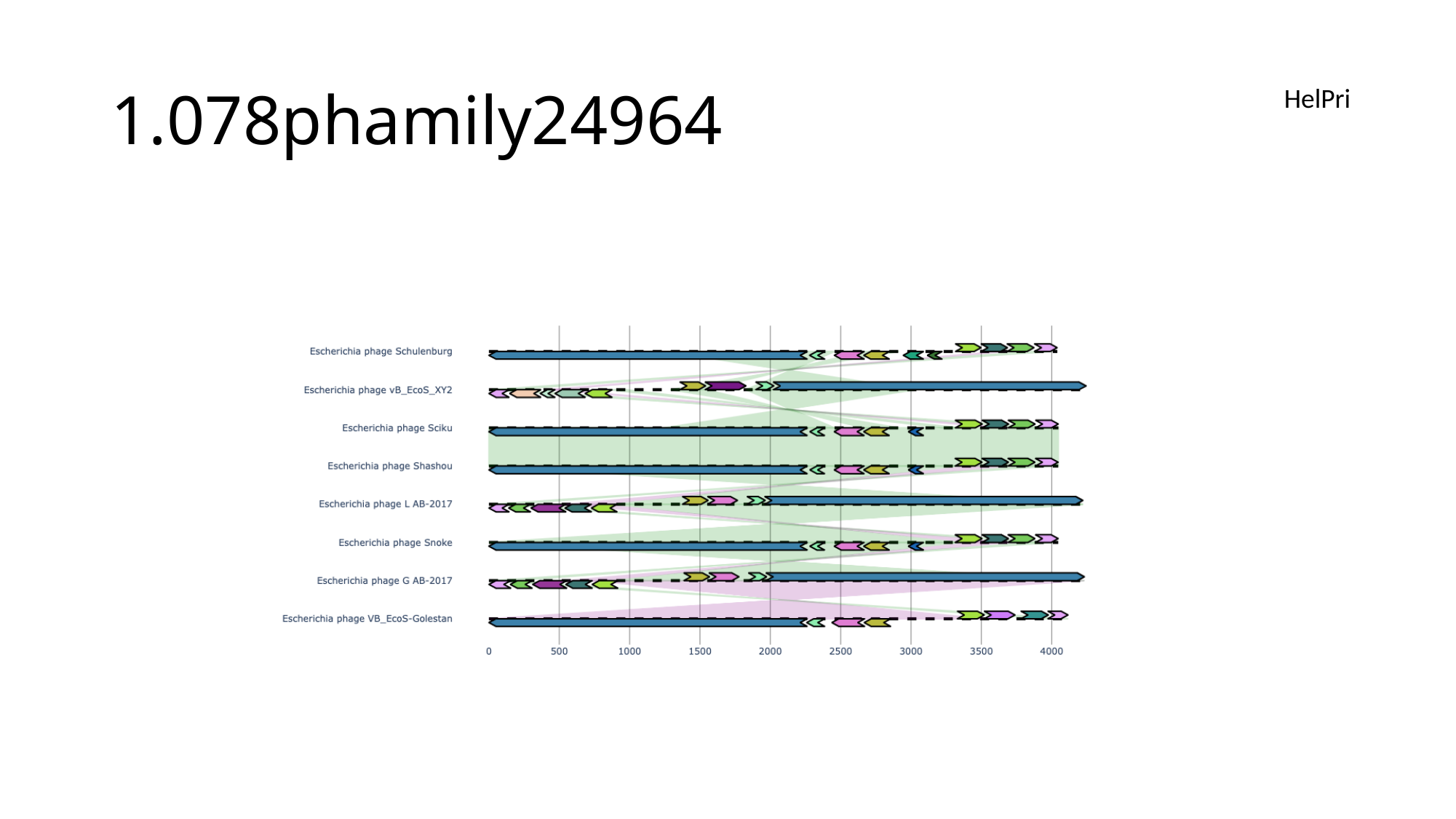

#### 1.078phamily24964
HelPri

### Slide 68
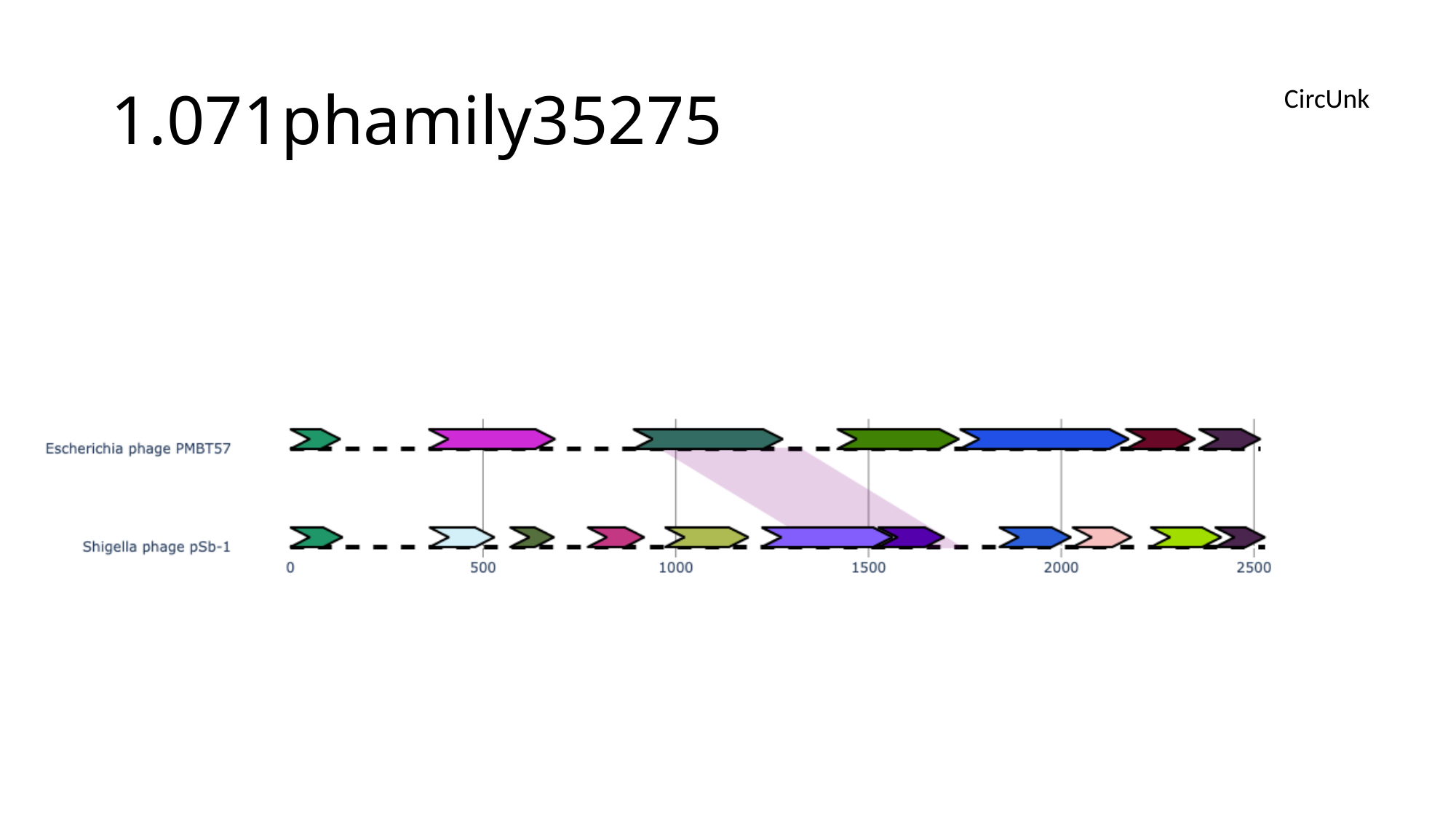

#### 1.071phamily35275
CircUnk

### Slide 69
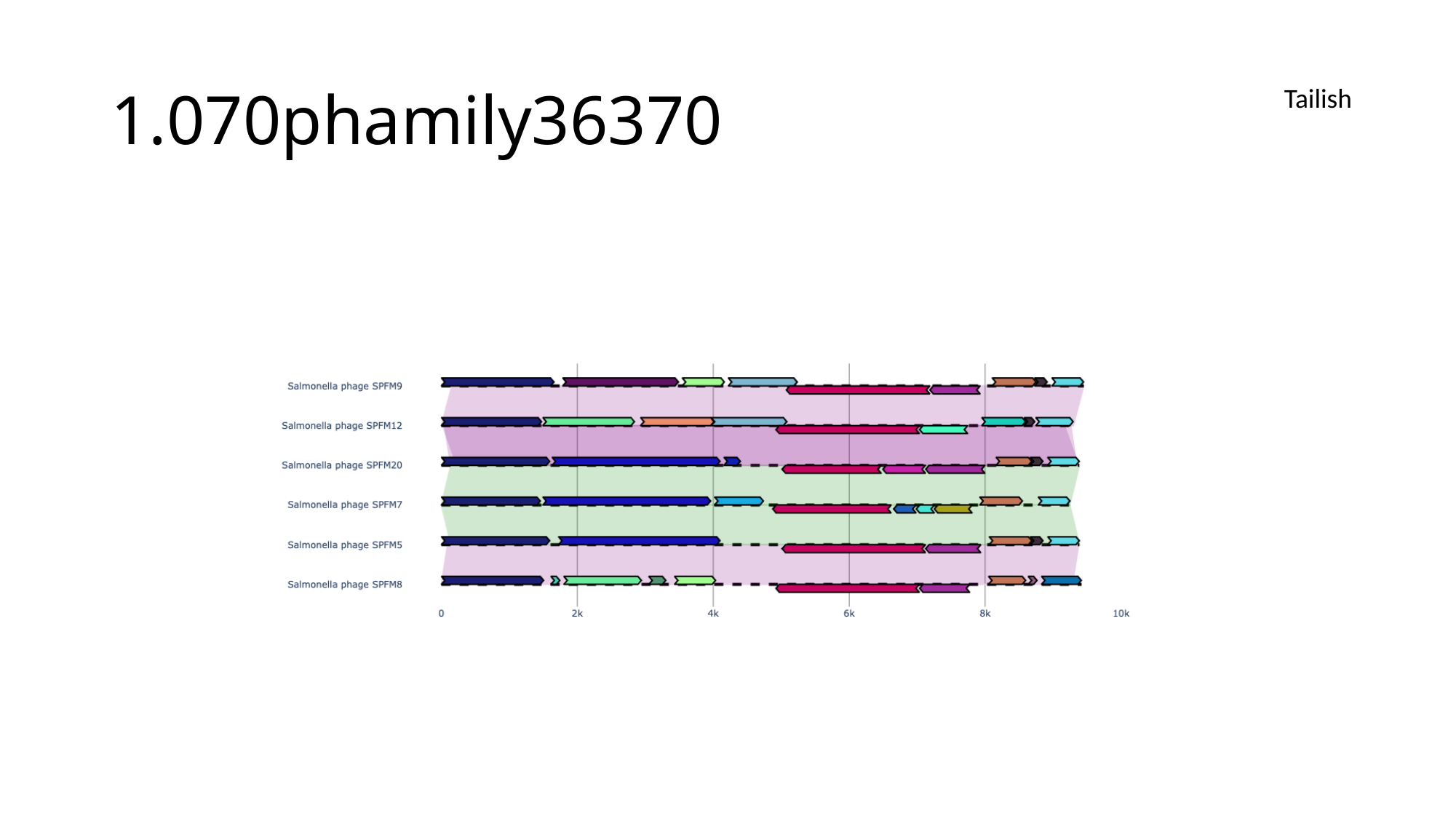

#### 1.070phamily36370
Tailish

### Slide 70
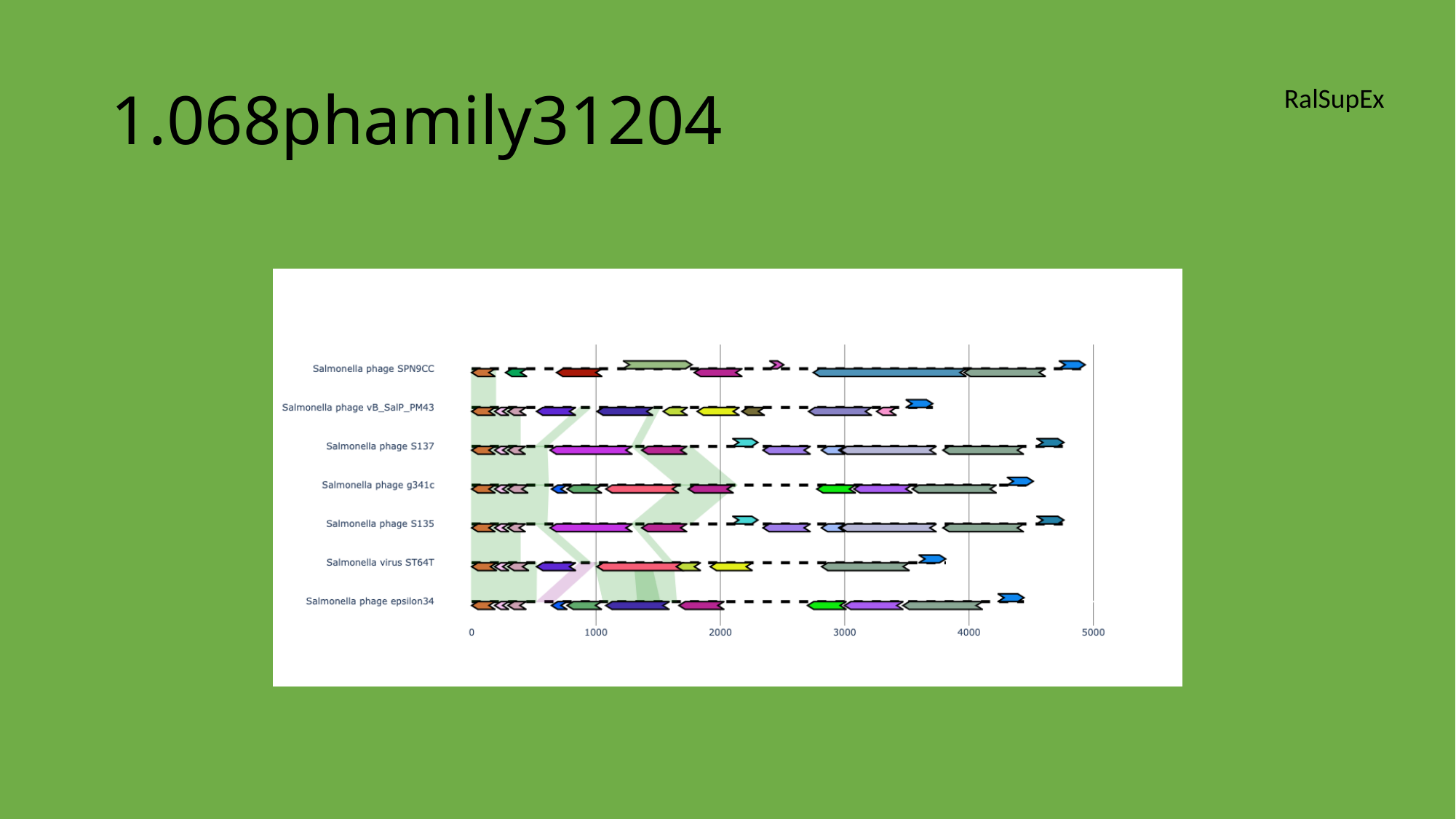

#### 1.068phamily31204
RalSupEx

### Slide 71
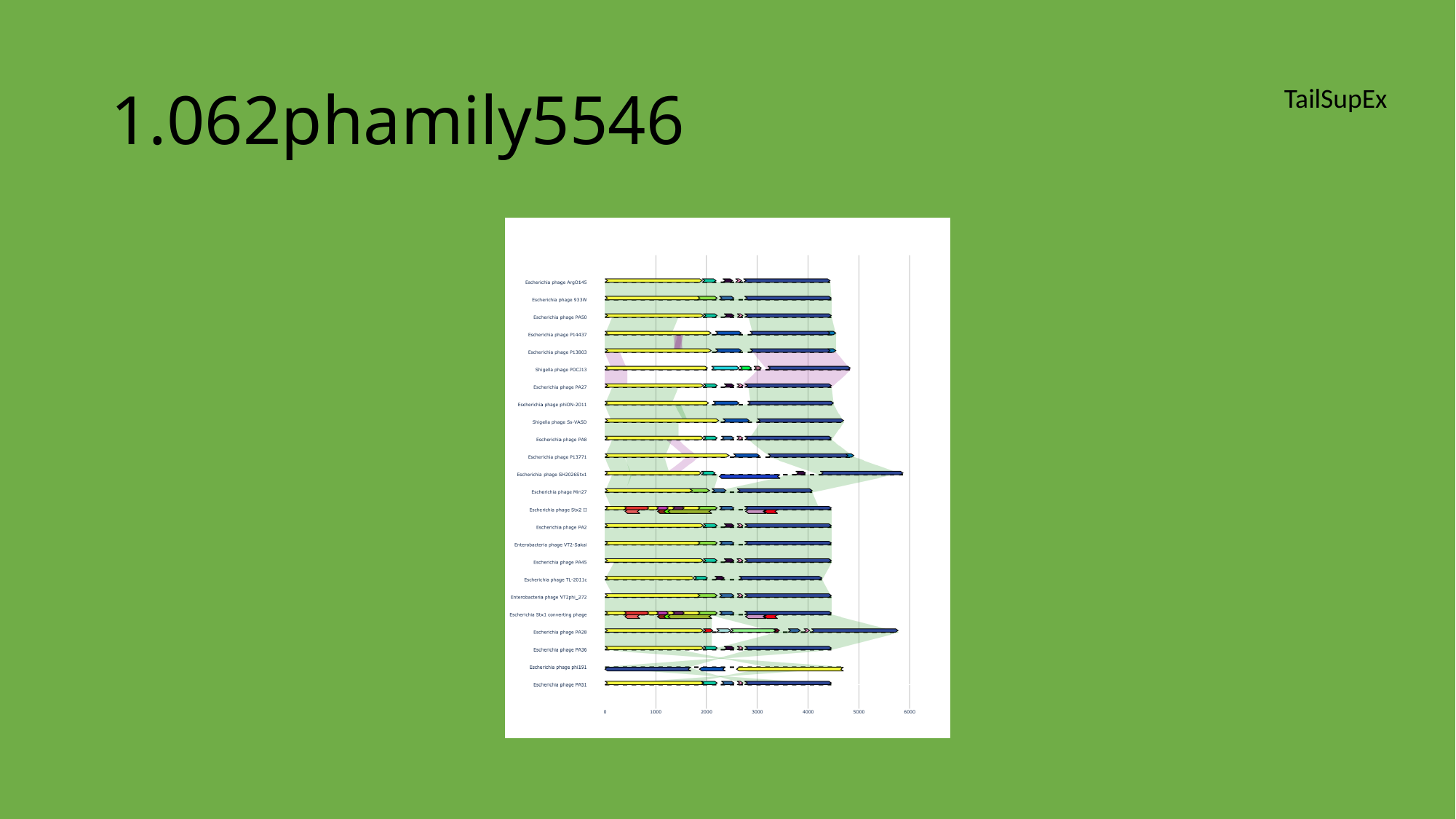

#### 1.062phamily5546
TailSupEx

### Slide 72
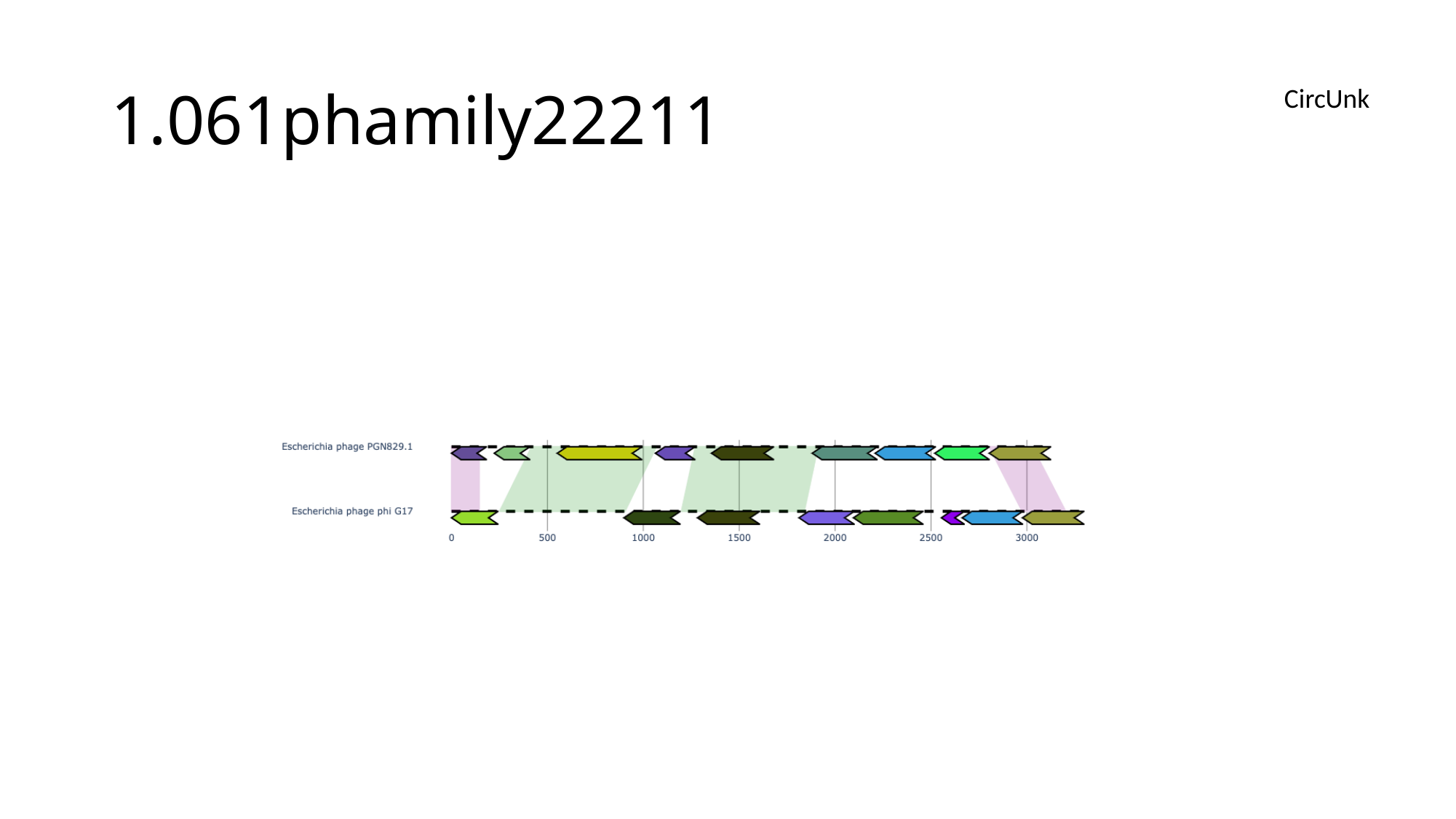

#### 1.061phamily22211
CircUnk

### Slide 73
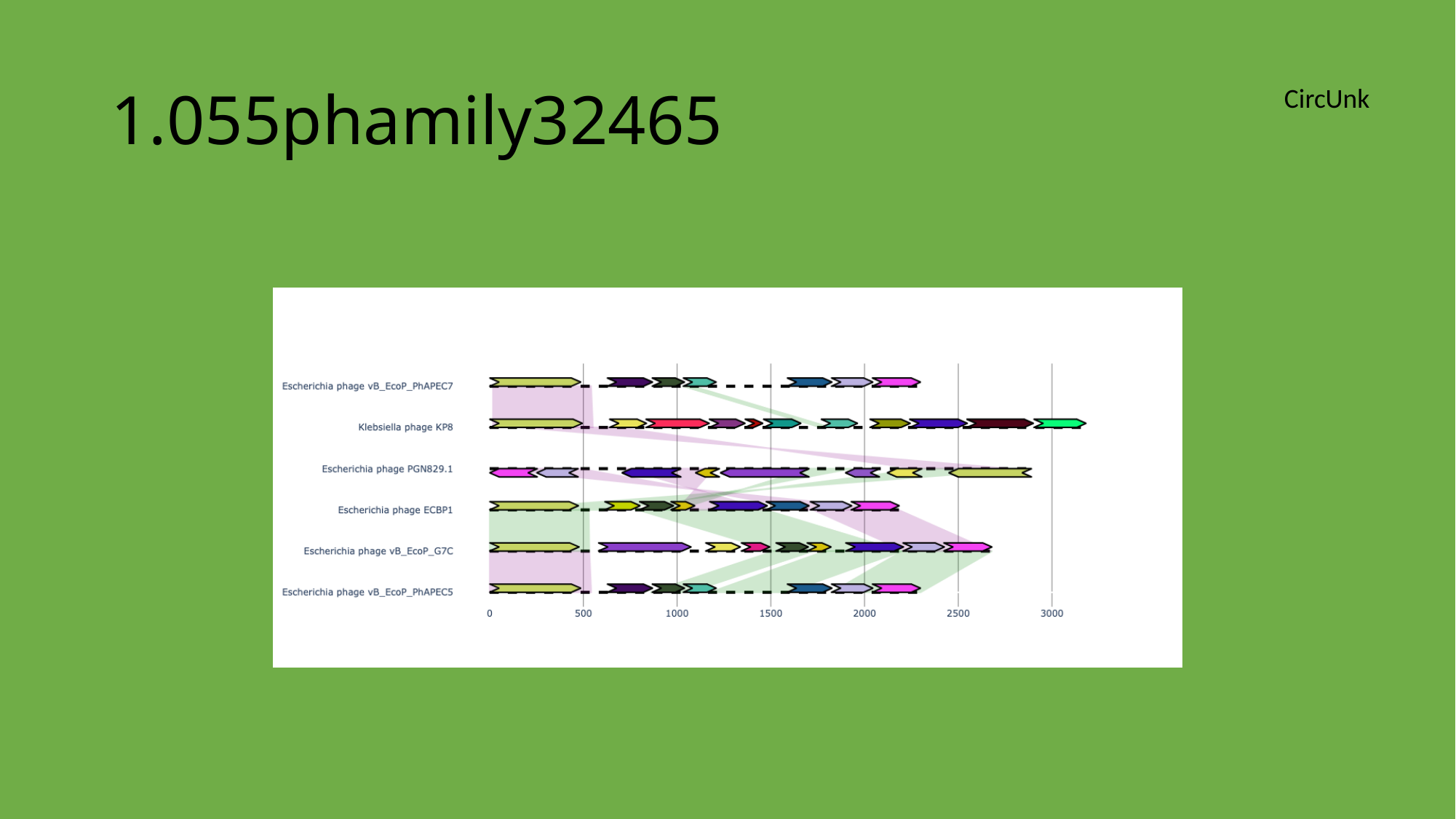

#### 1.055phamily32465
CircUnk

### Slide 74
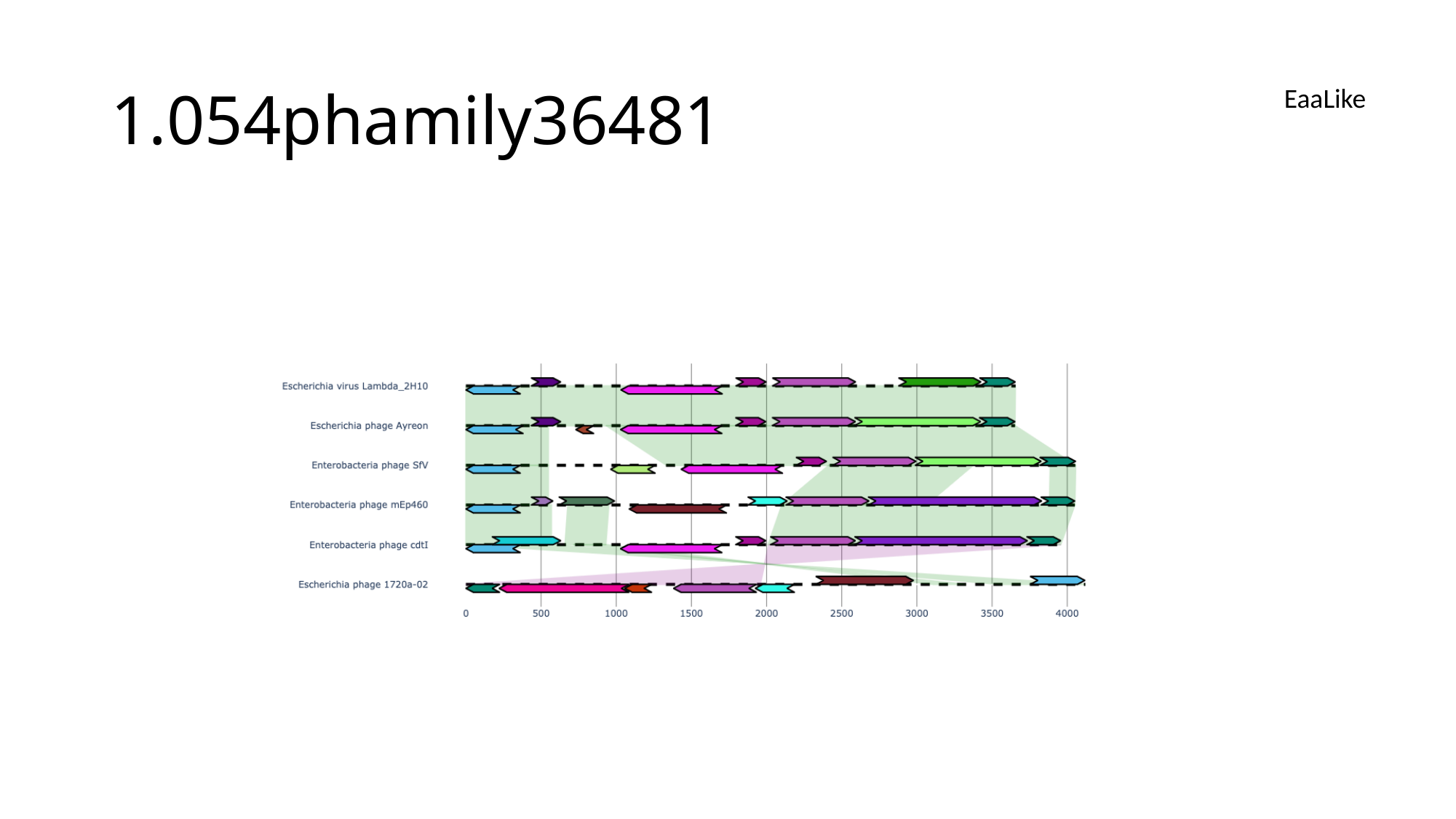

#### 1.054phamily36481
EaaLike

### Slide 75
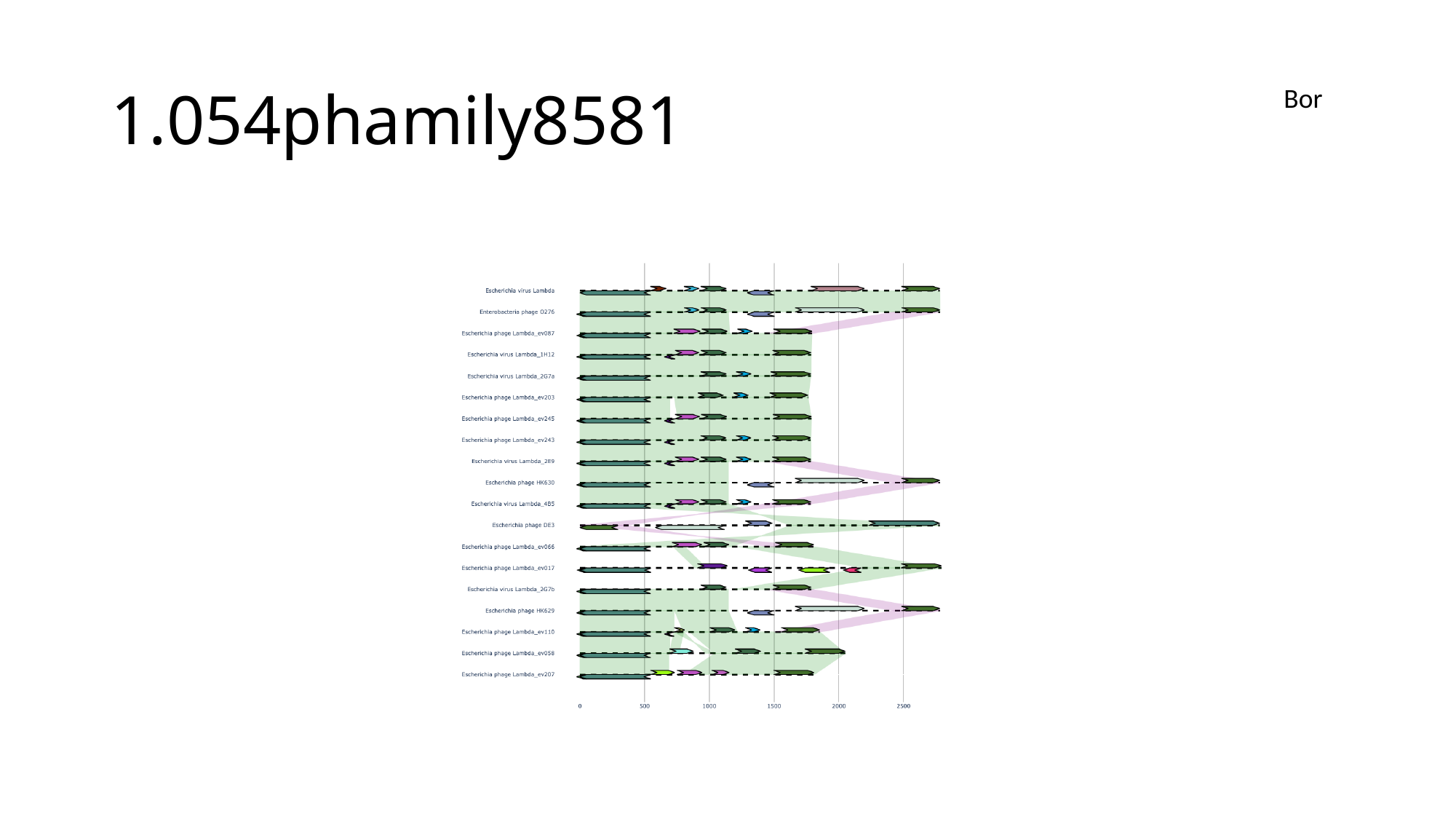

#### 1.054phamily8581
Bor

### Slide 76
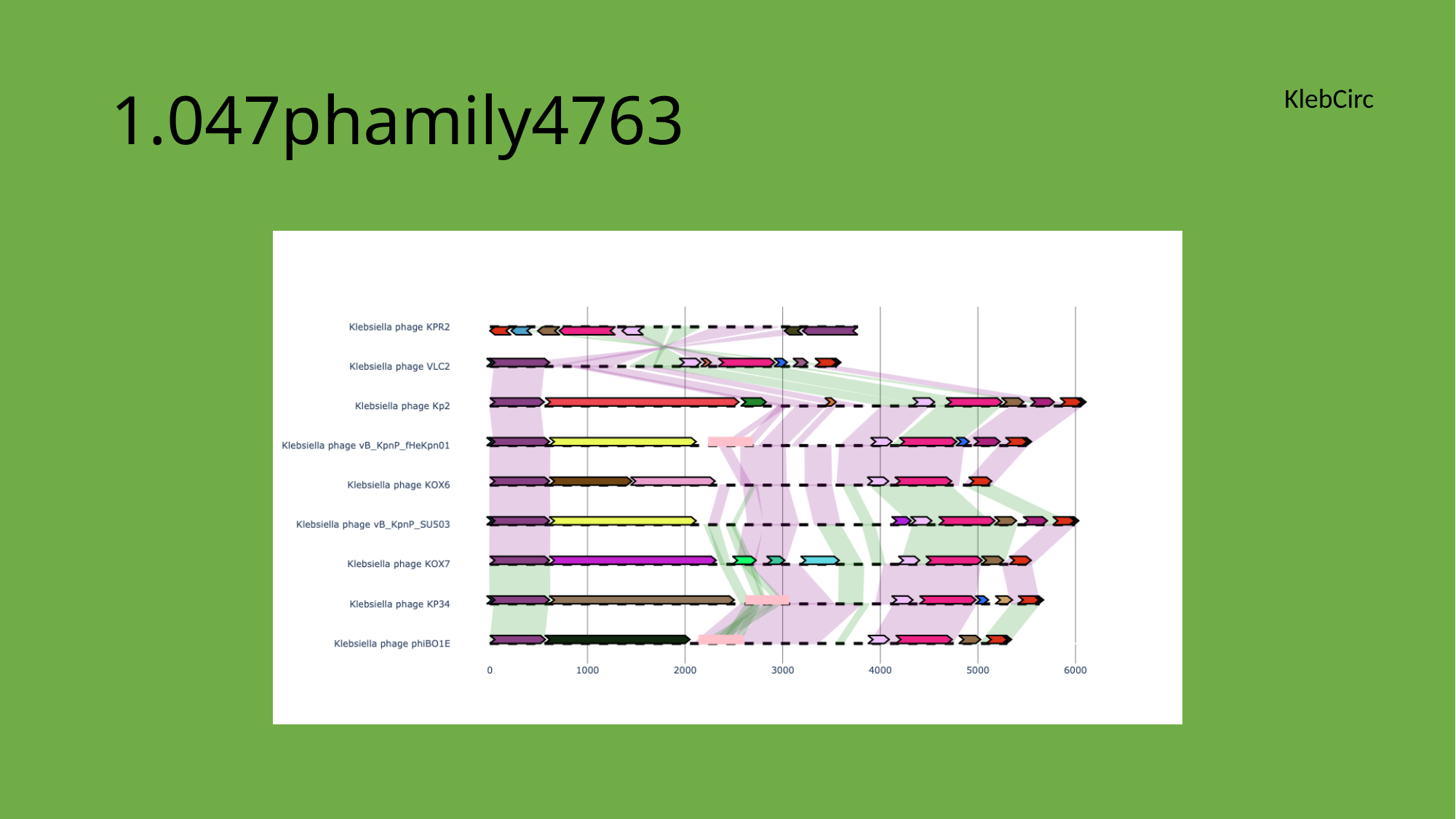

#### 1.047phamily4763
KlebCirc

### Slide 77
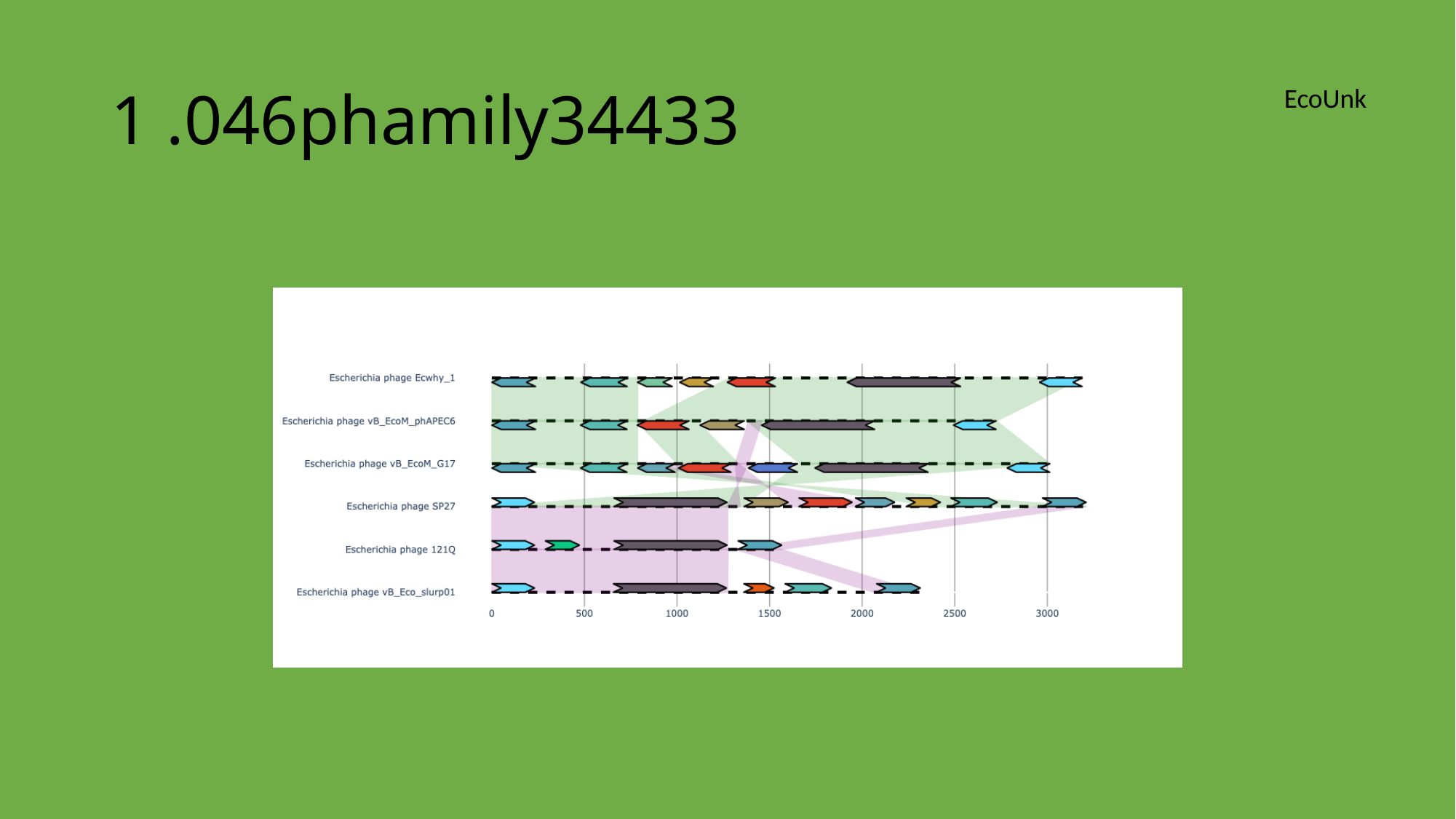

#### 1 .046phamily34433
EcoUnk

### Slide 78
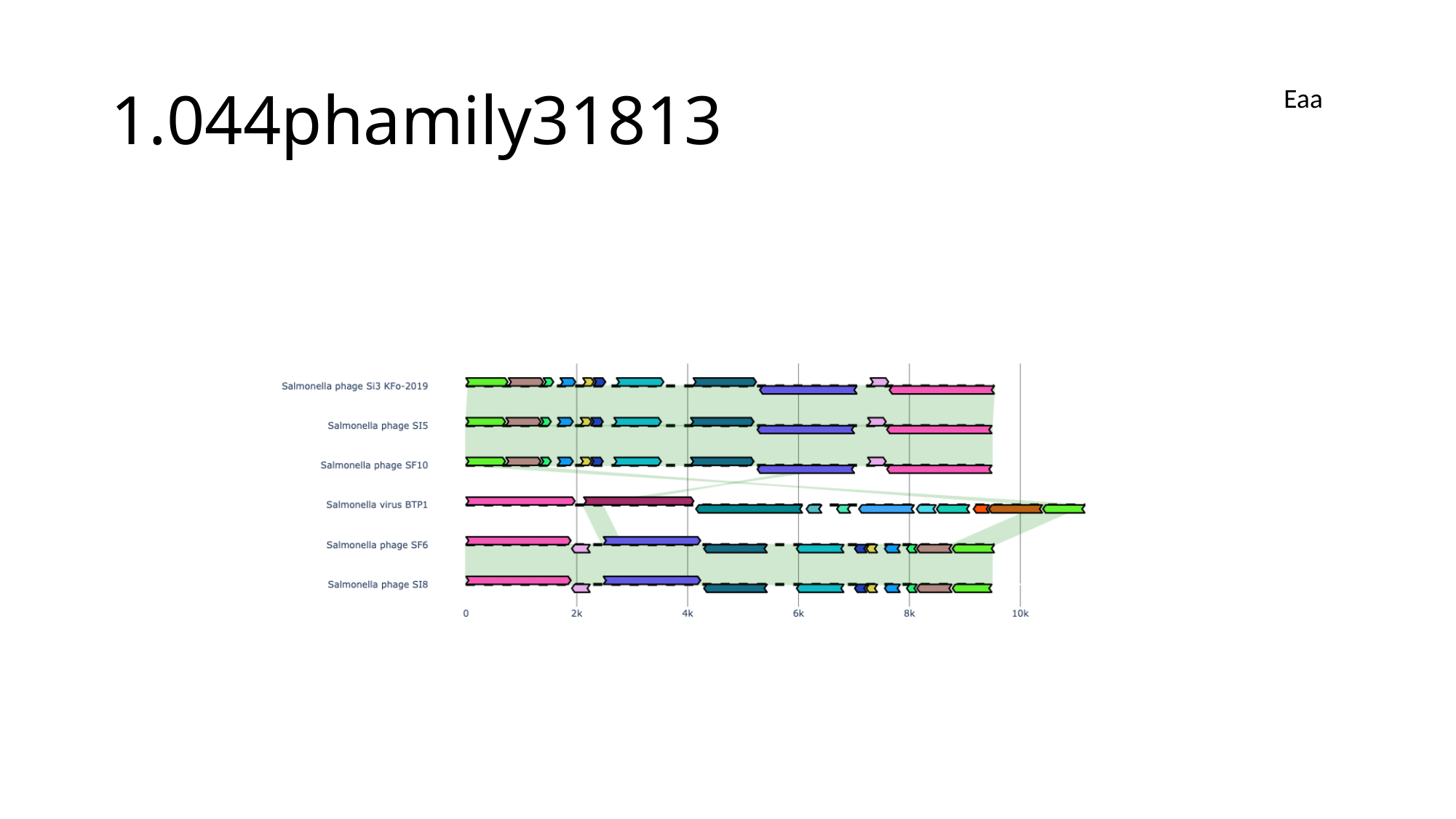

#### 1.044phamily31813
Eaa

### Slide 79
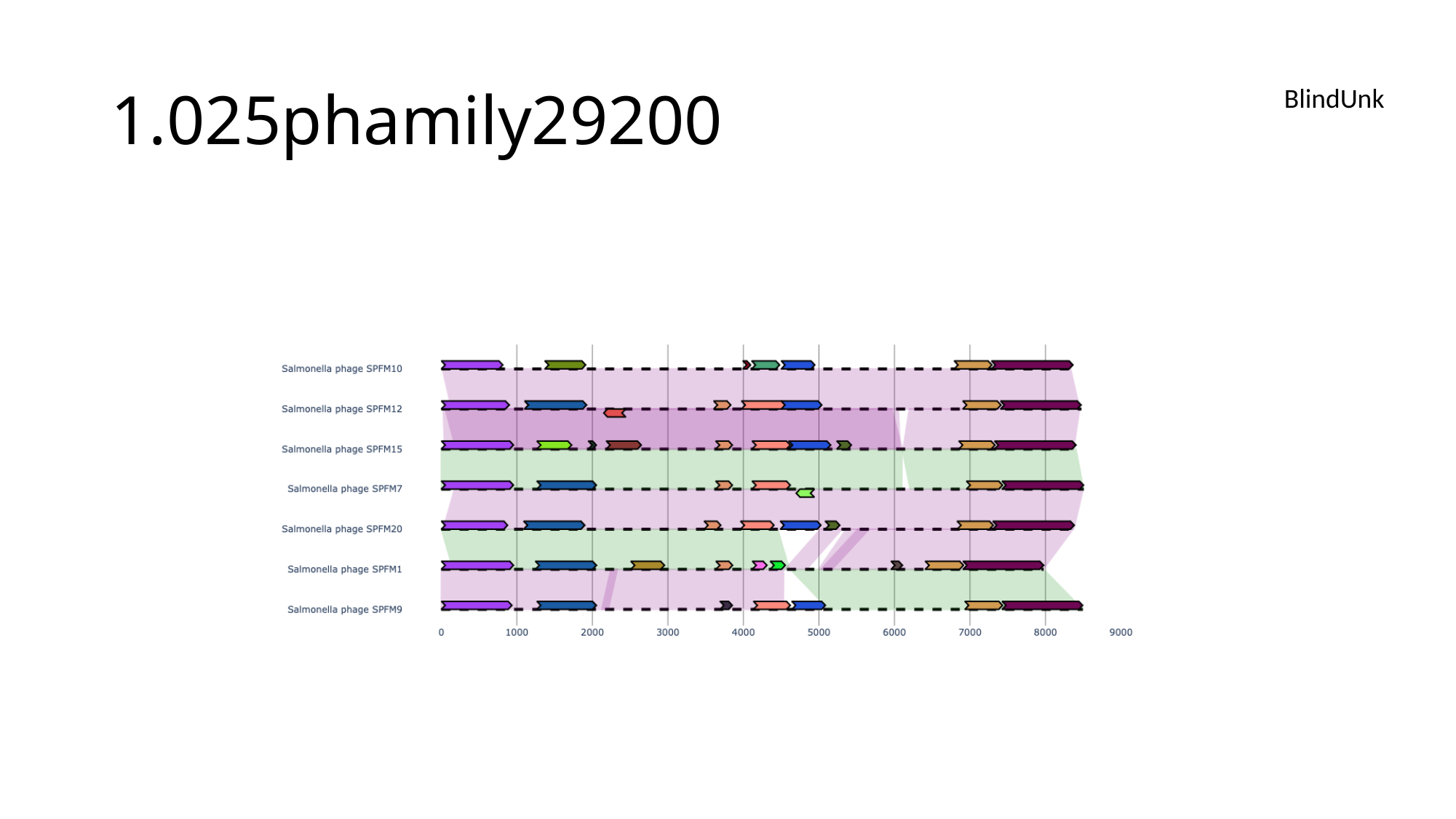

#### 1.025phamily29200
BlindUnk

### Slide 80
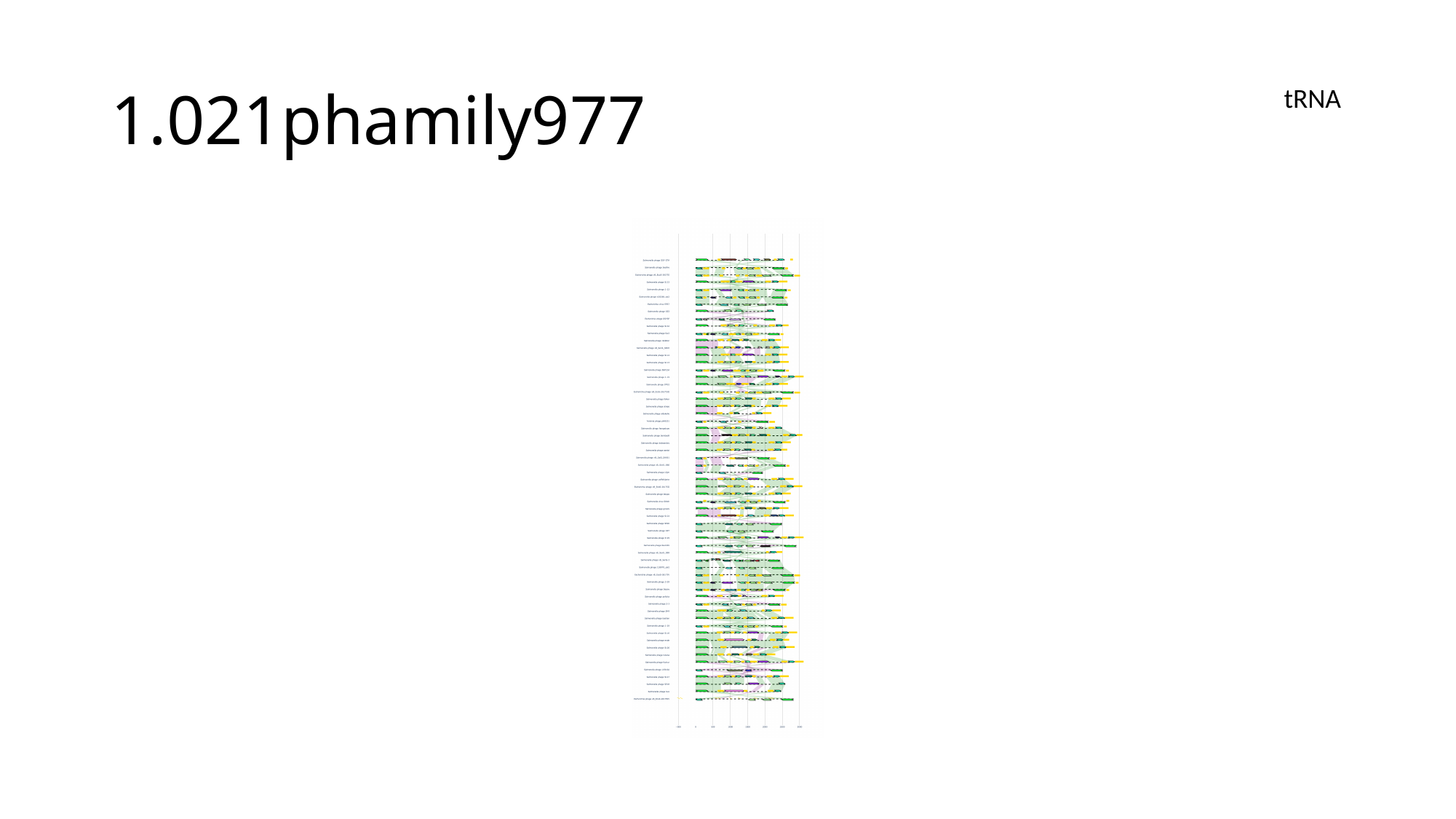

#### 1.021phamily977
tRNA

### Slide 81
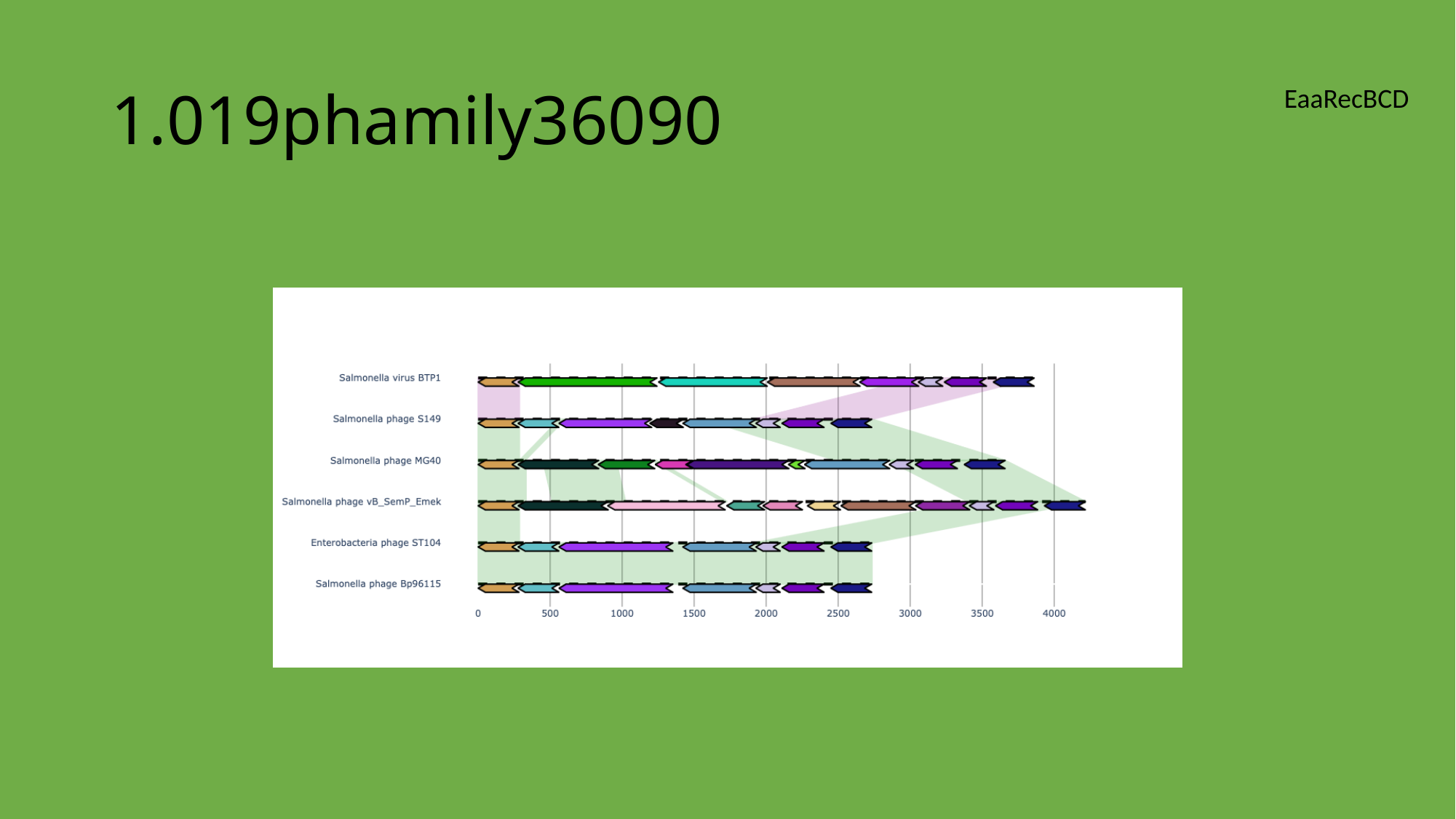

#### 1.019phamily36090
EaaRecBCD

### Slide 82
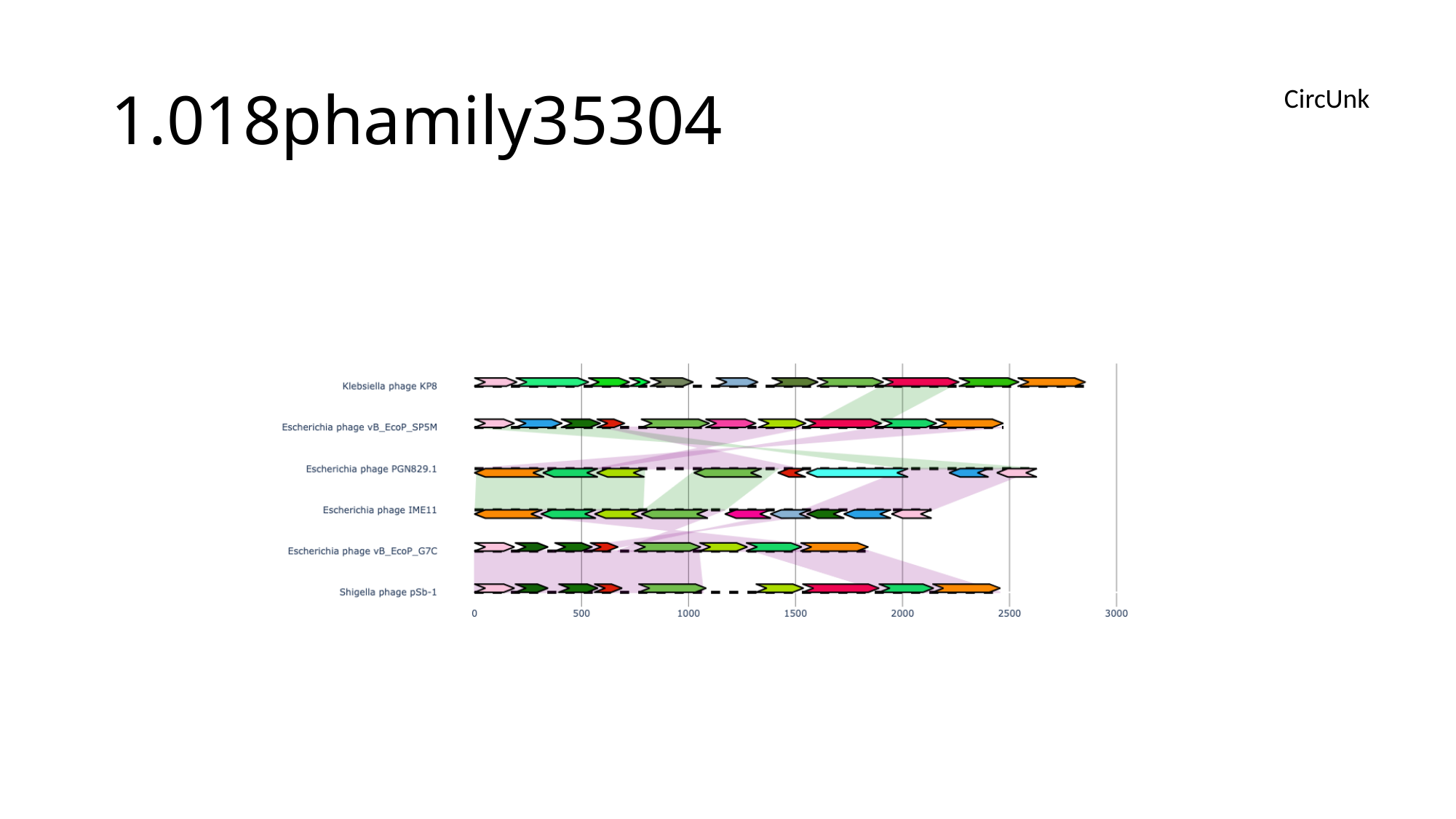

#### 1.018phamily35304
CircUnk

### Slide 83
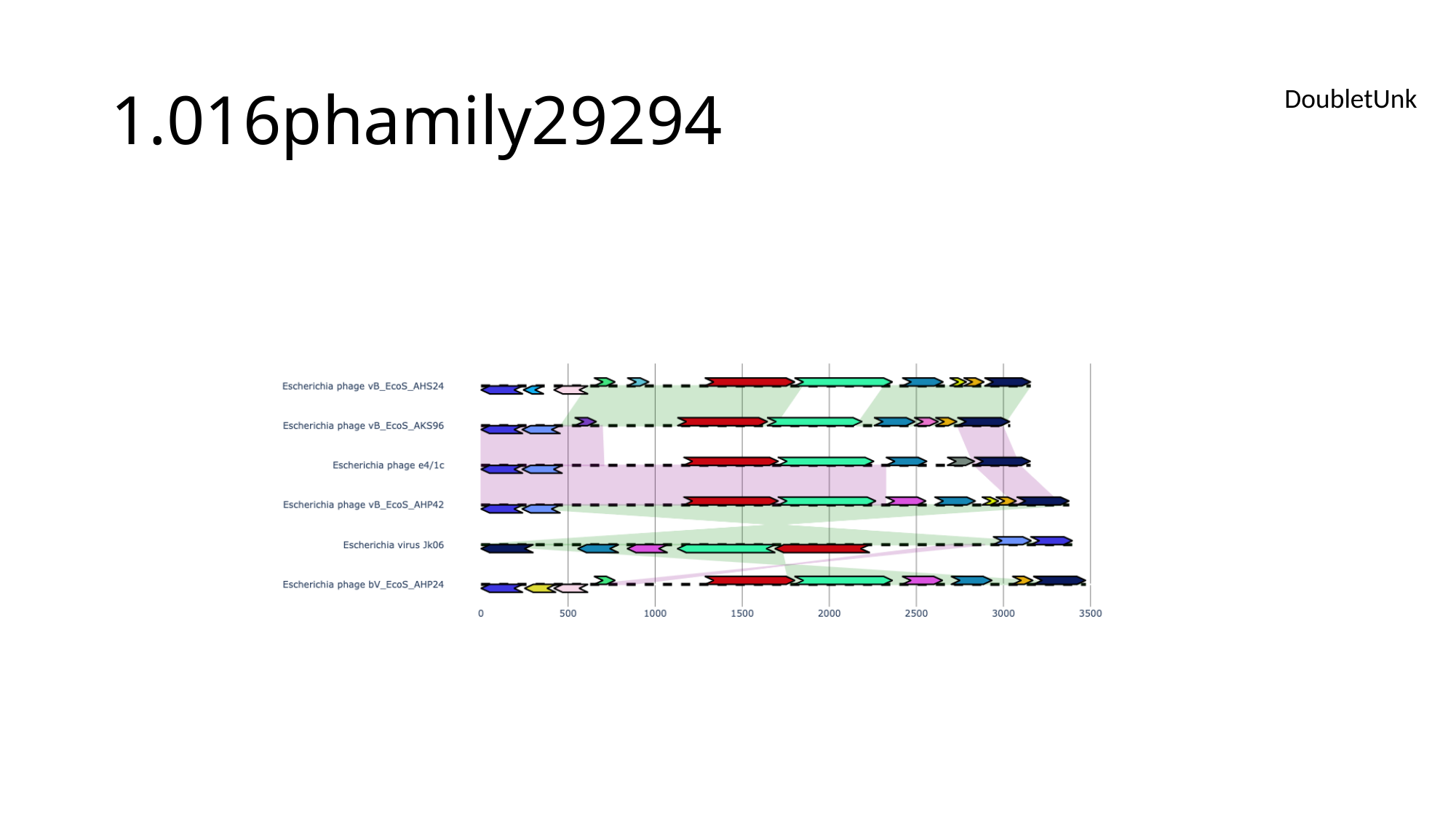

#### 1.016phamily29294
DoubletUnk

### Slide 84
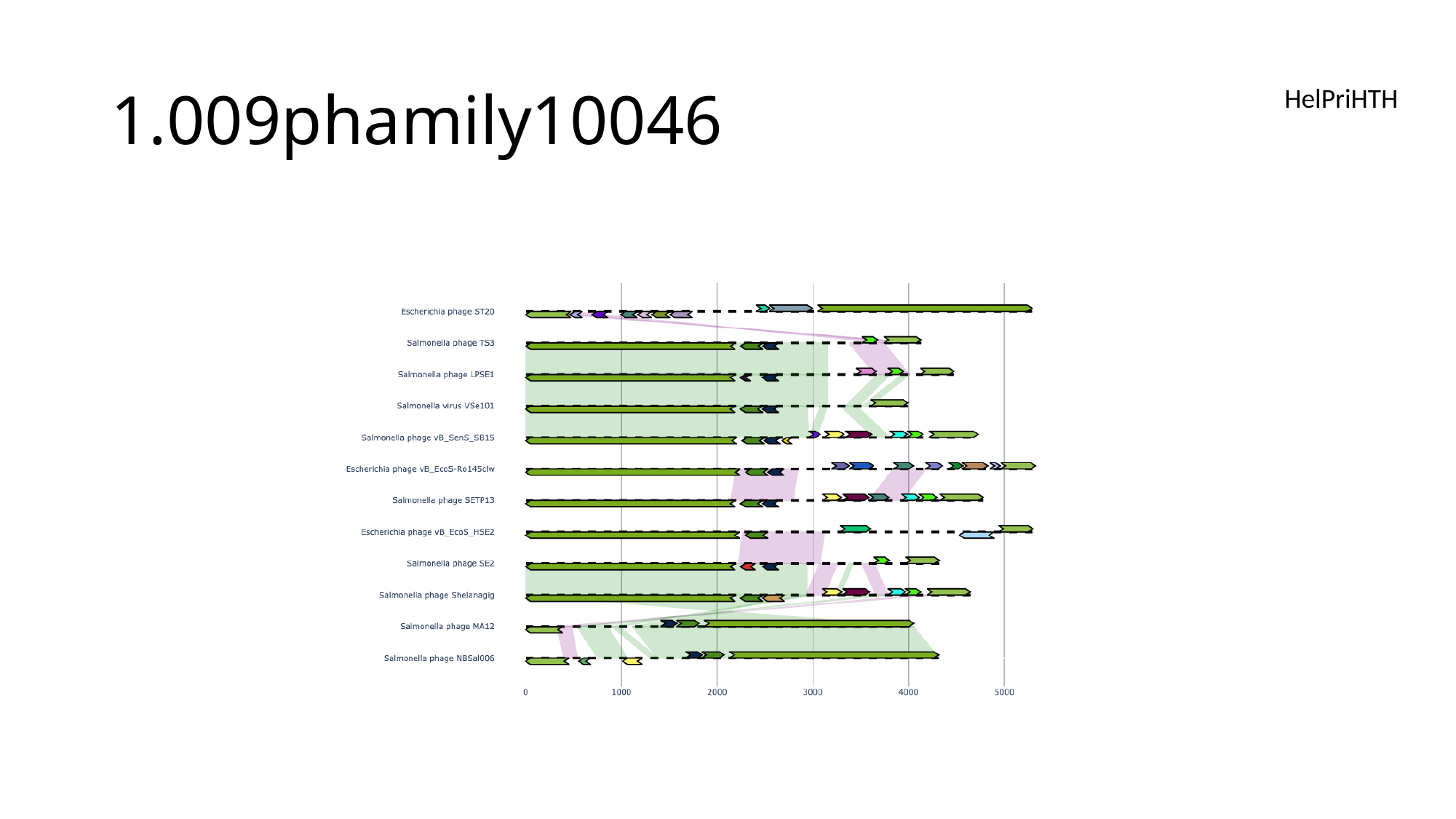

#### 1.009phamily10046
HelPriHTH

### Slide 85
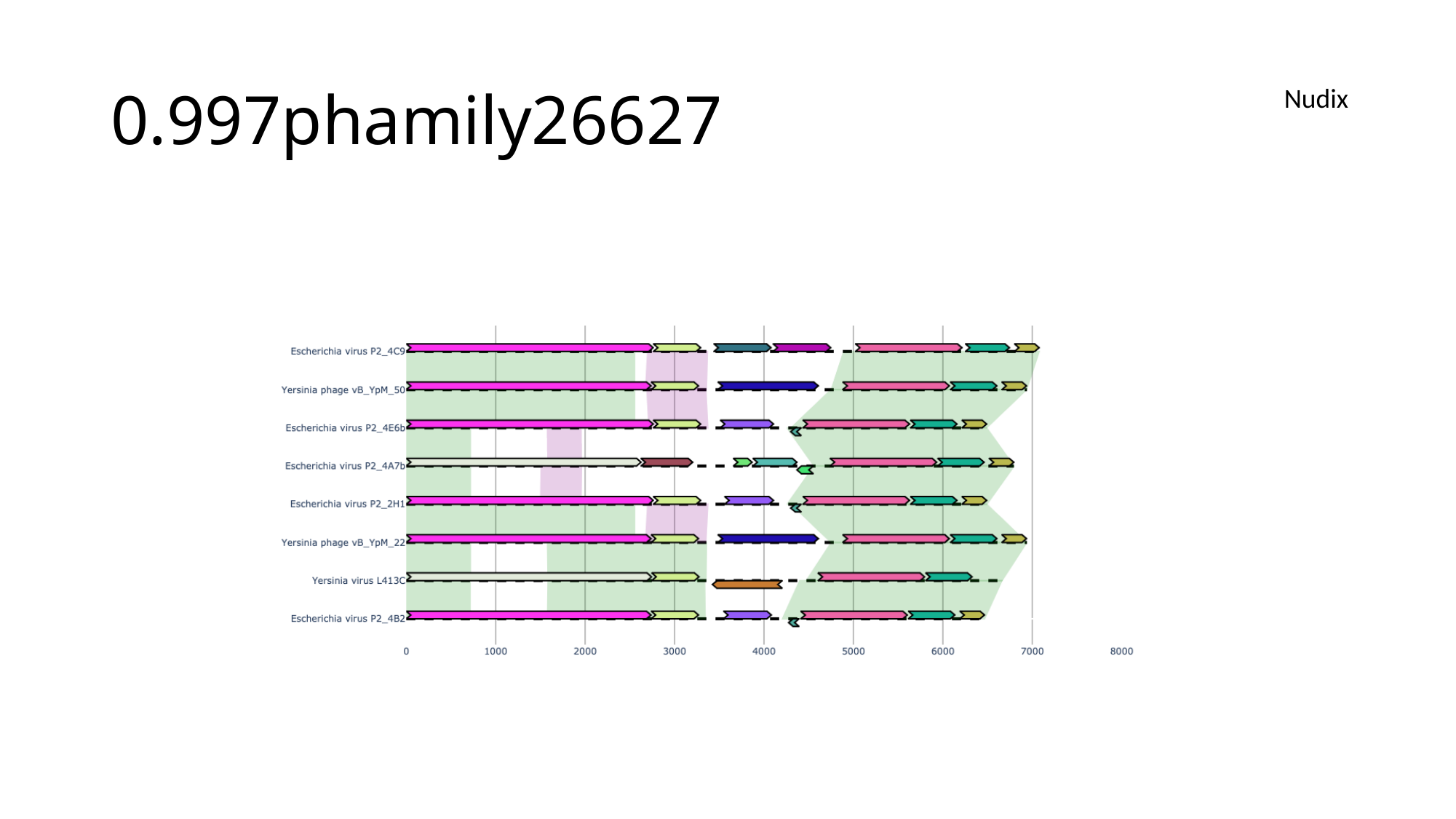

#### 0.997phamily26627
Nudix

### Slide 86
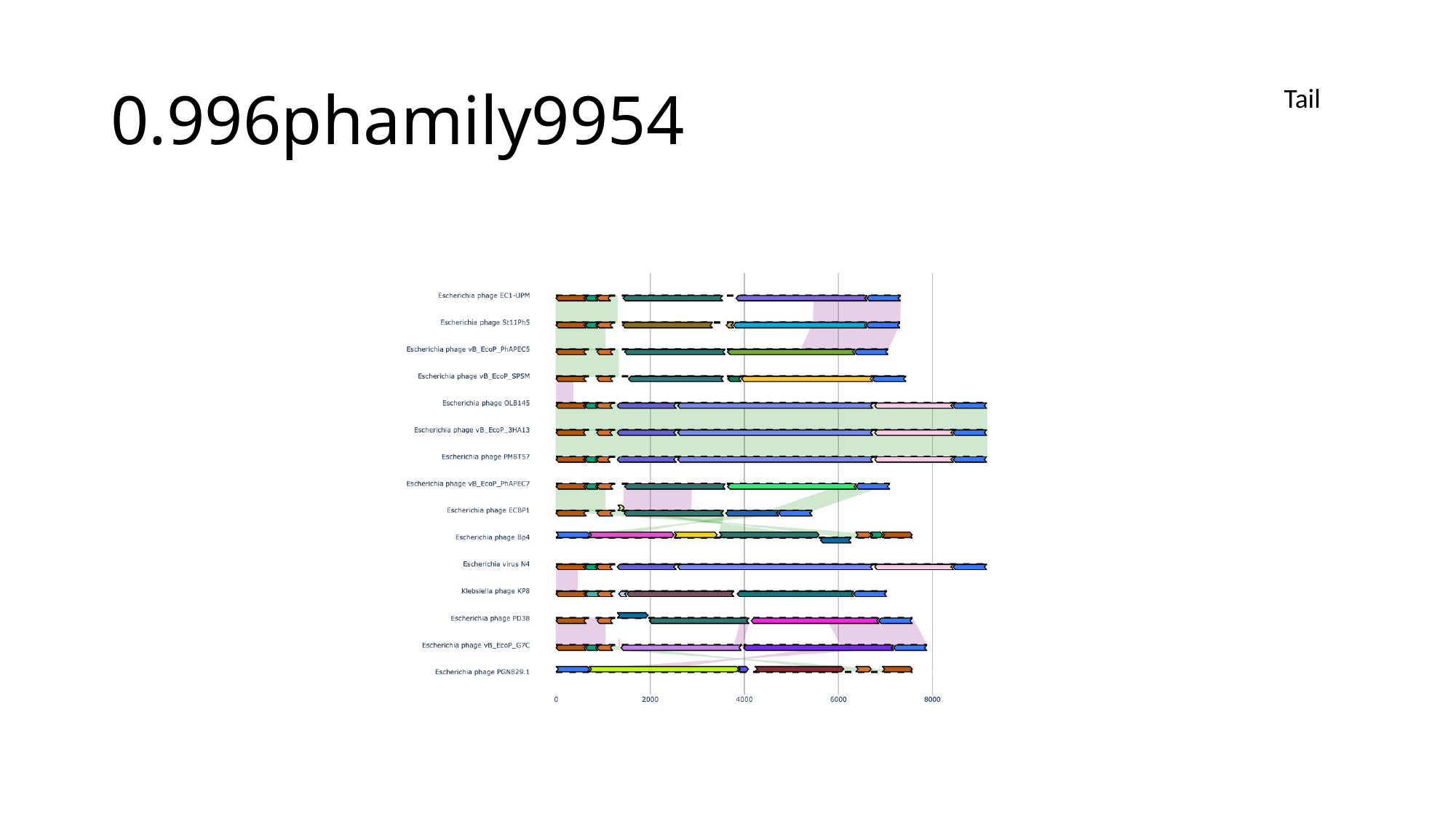

#### 0.996phamily9954
Tail

### Slide 87
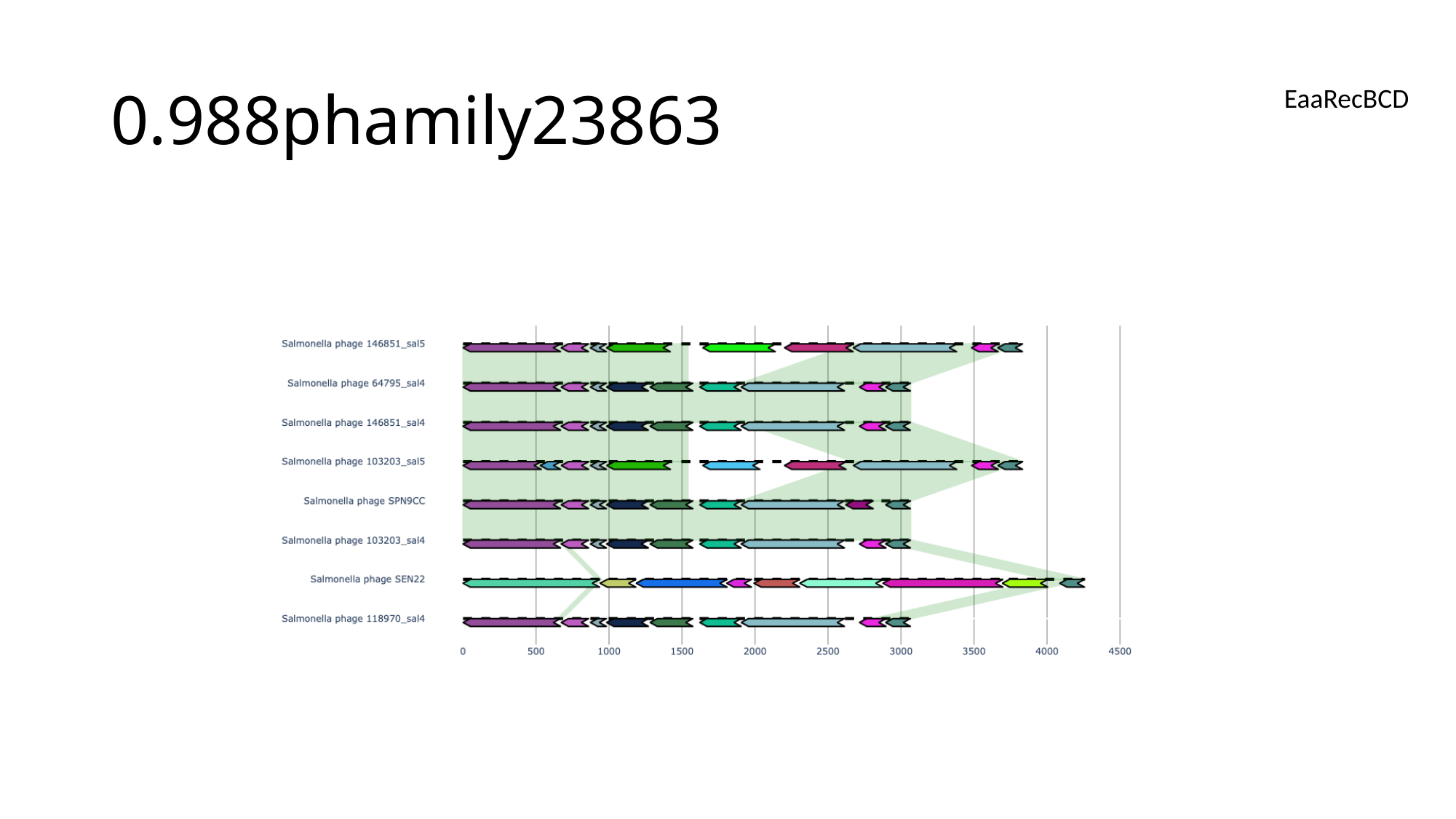

#### 0.988phamily23863
EaaRecBCD

### Slide 88
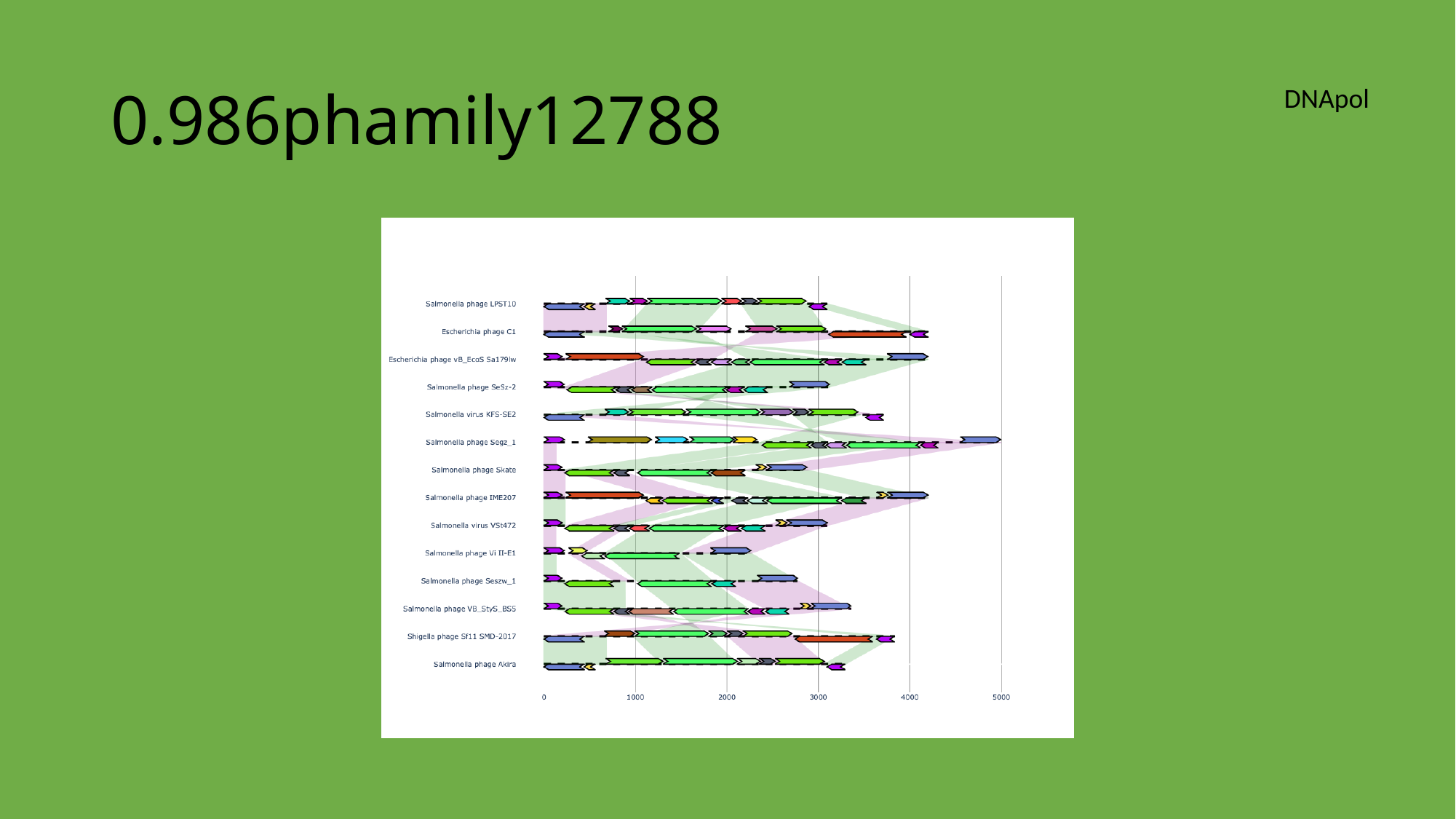

#### 0.986phamily12788
DNApol

### Slide 89
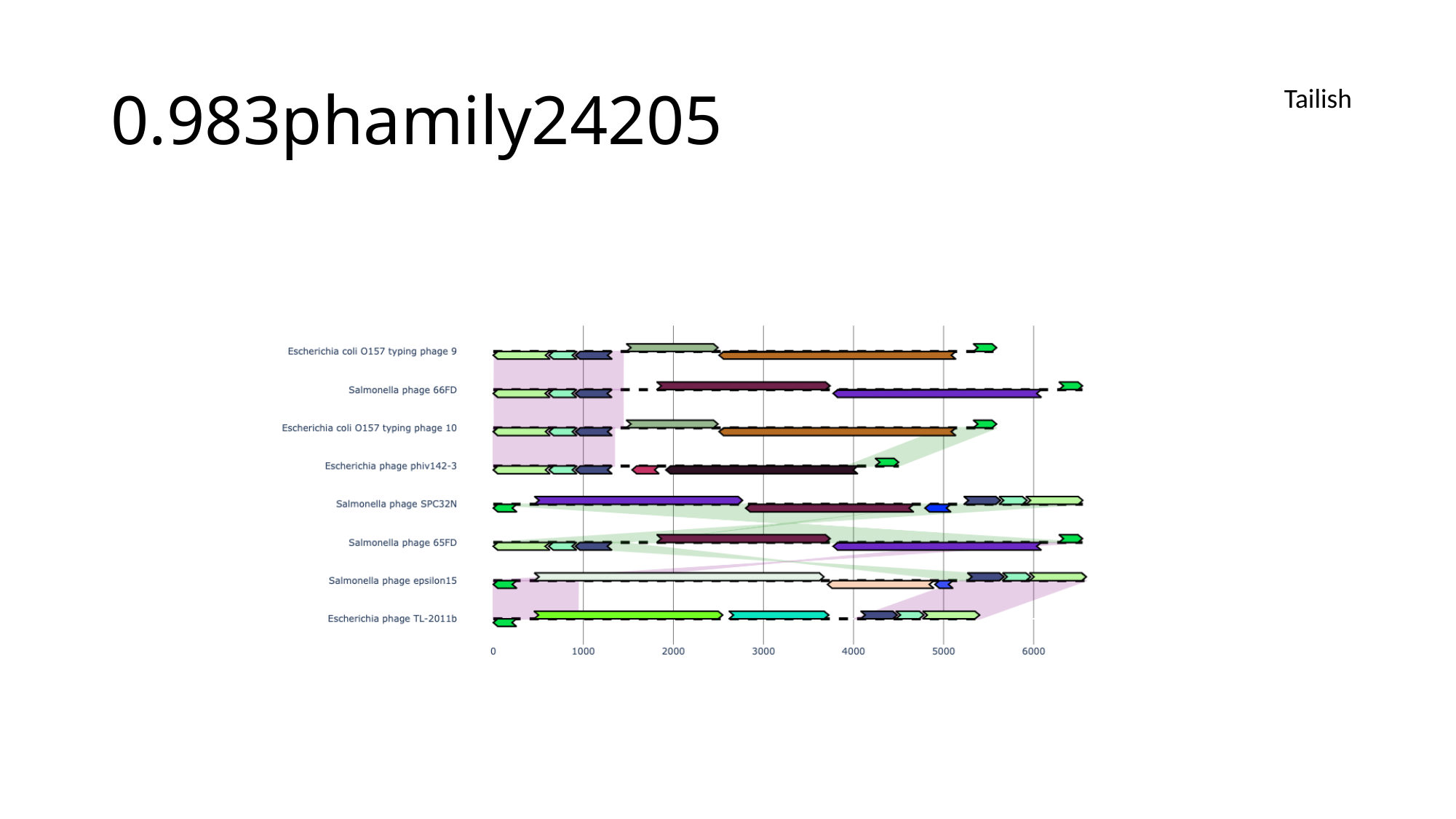

#### 0.983phamily24205
Tailish

### Slide 90
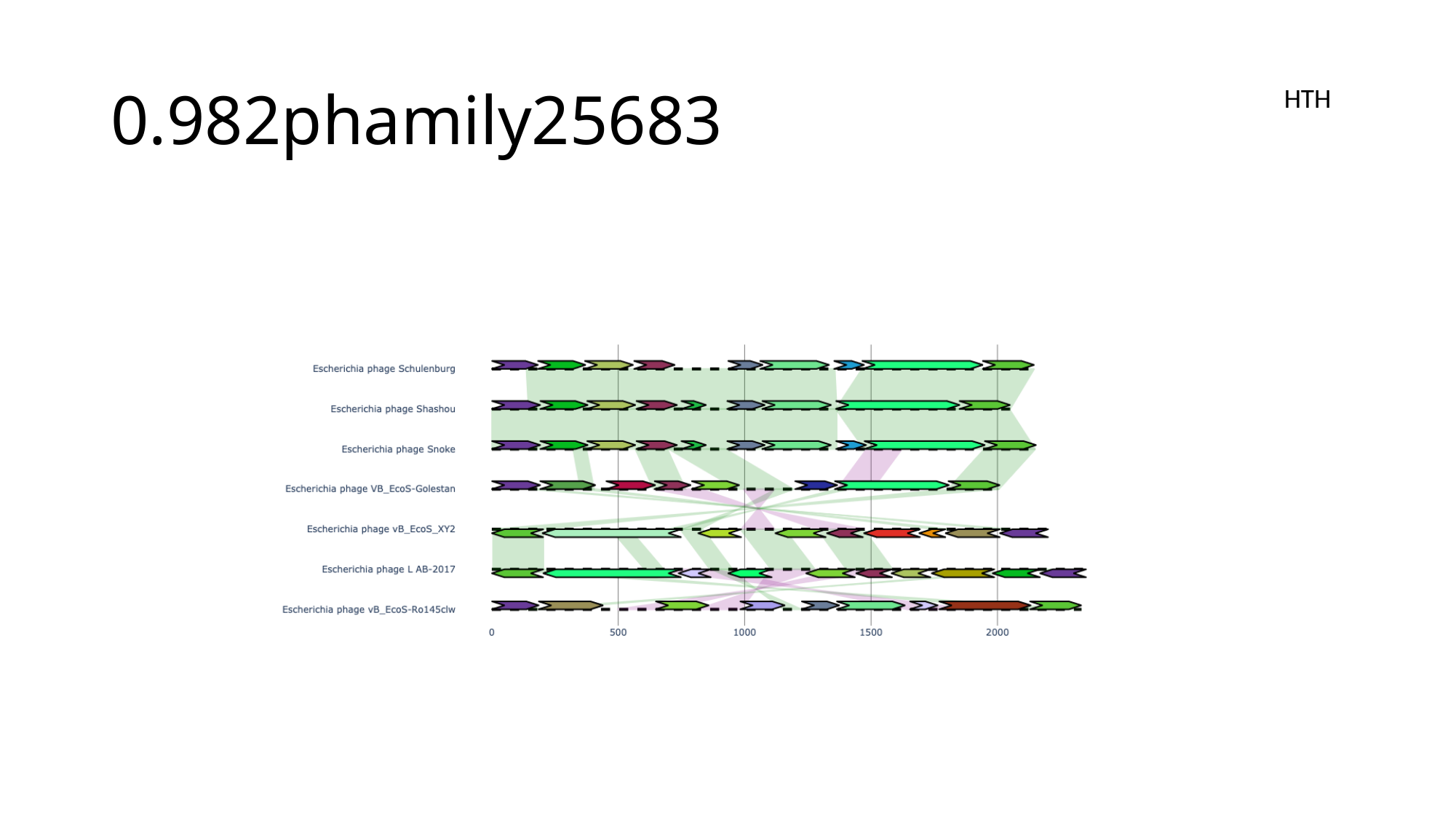

#### 0.982phamily25683
HTH

### Slide 91
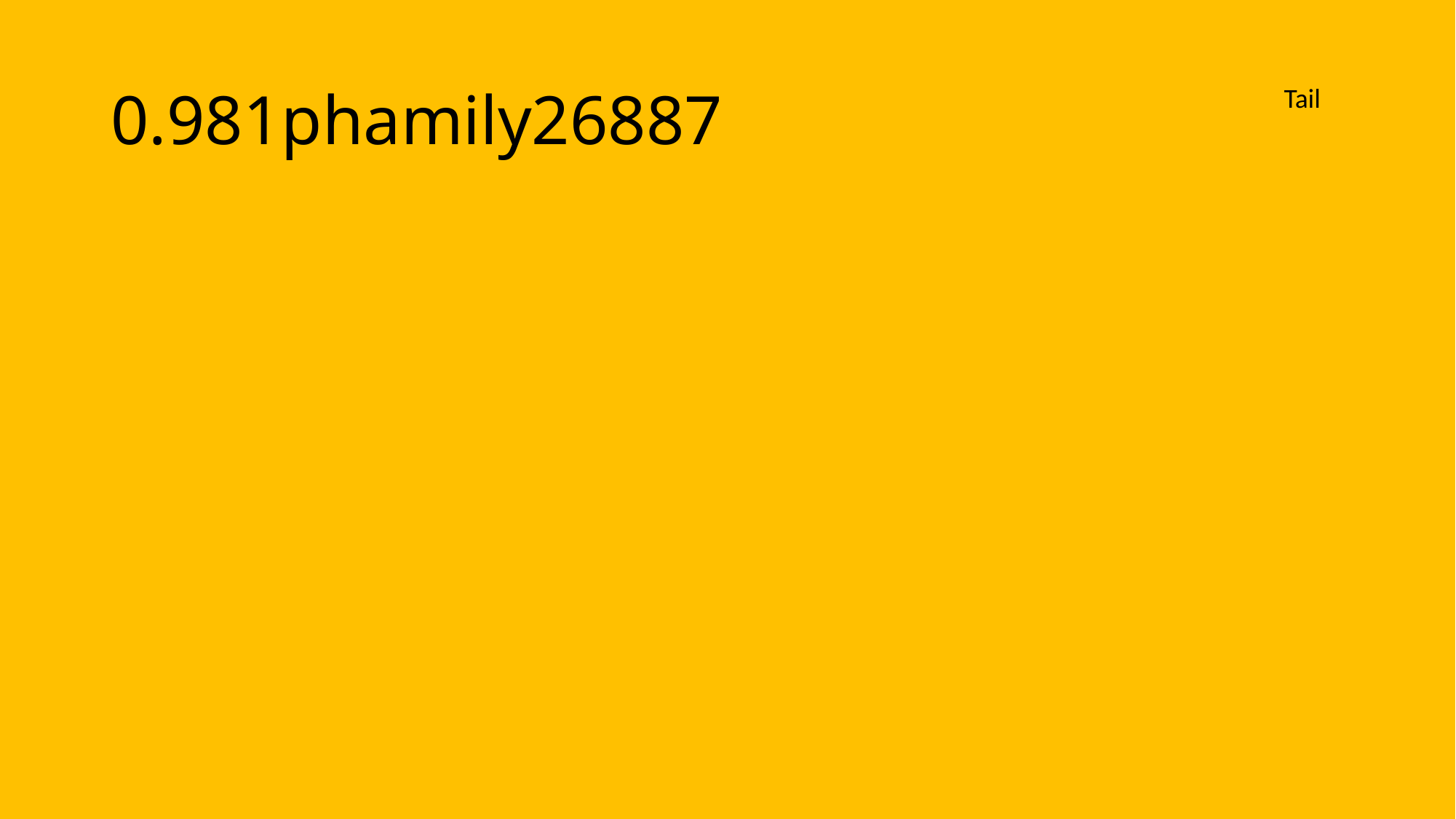

#### 0.981phamily26887
Tail

### Slide 92
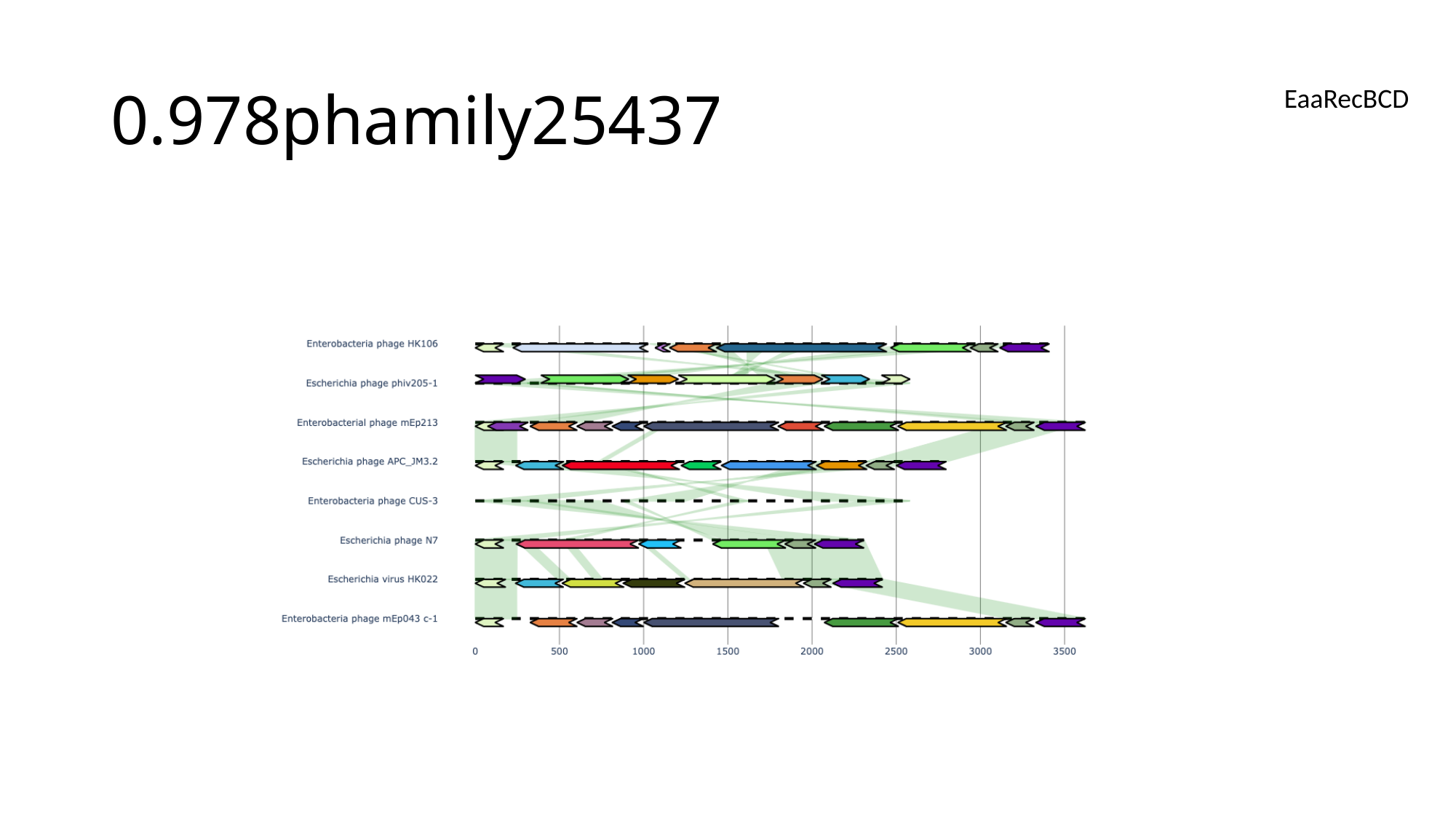

#### 0.978phamily25437
EaaRecBCD

### Slide 93
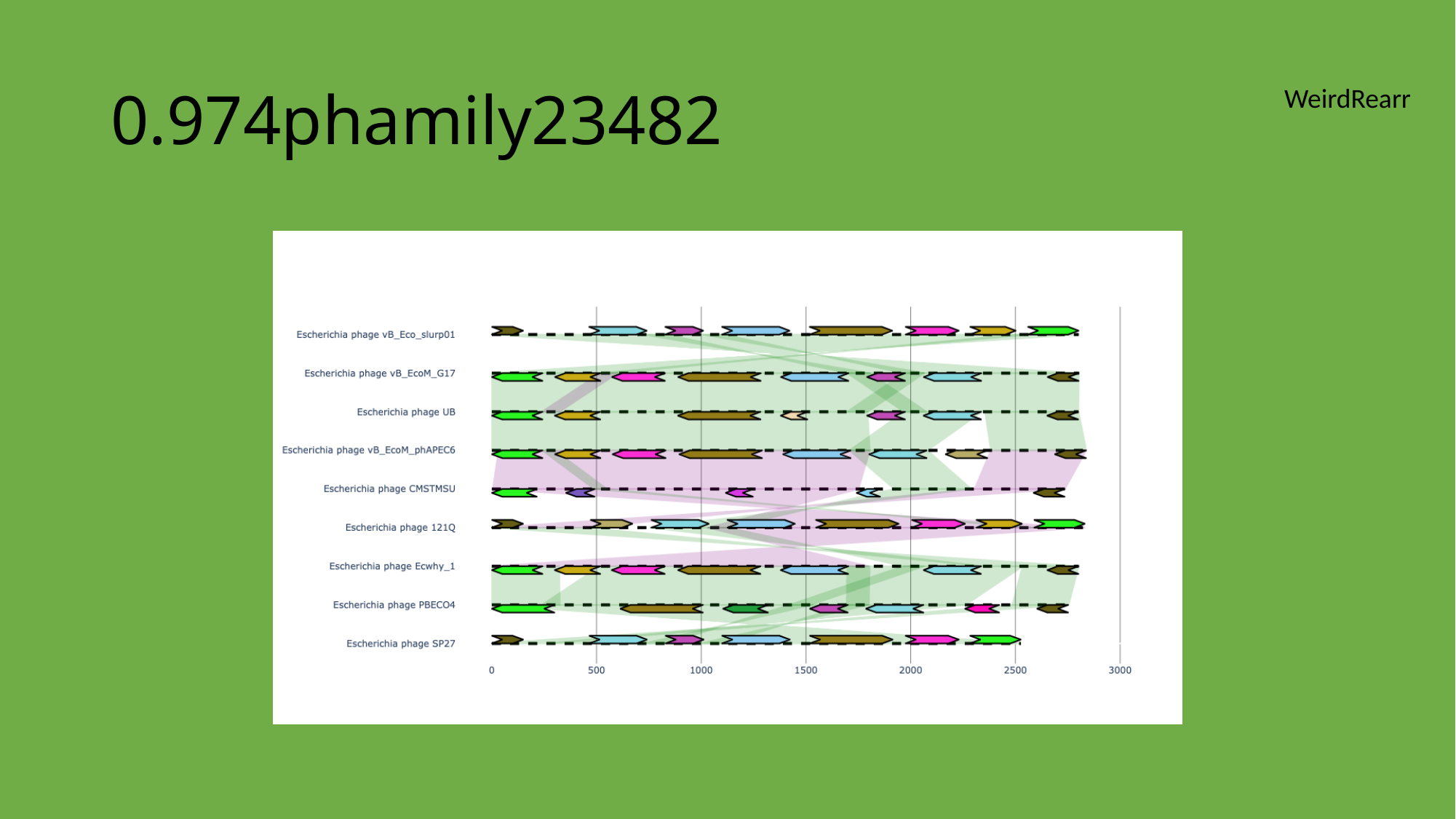

#### 0.974phamily23482
WeirdRearr

### Slide 94
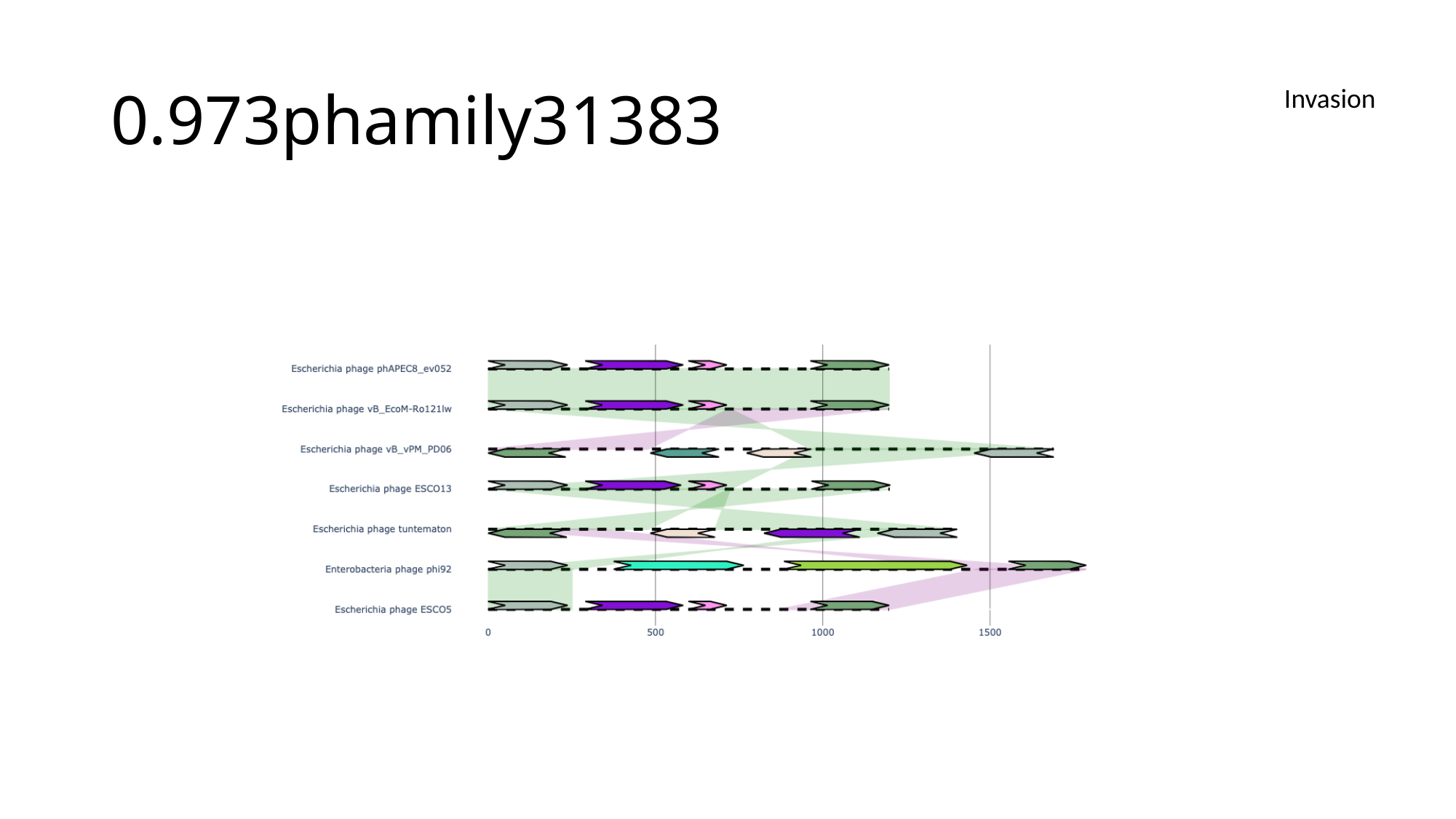

#### 0.973phamily31383
Invasion

### Slide 95
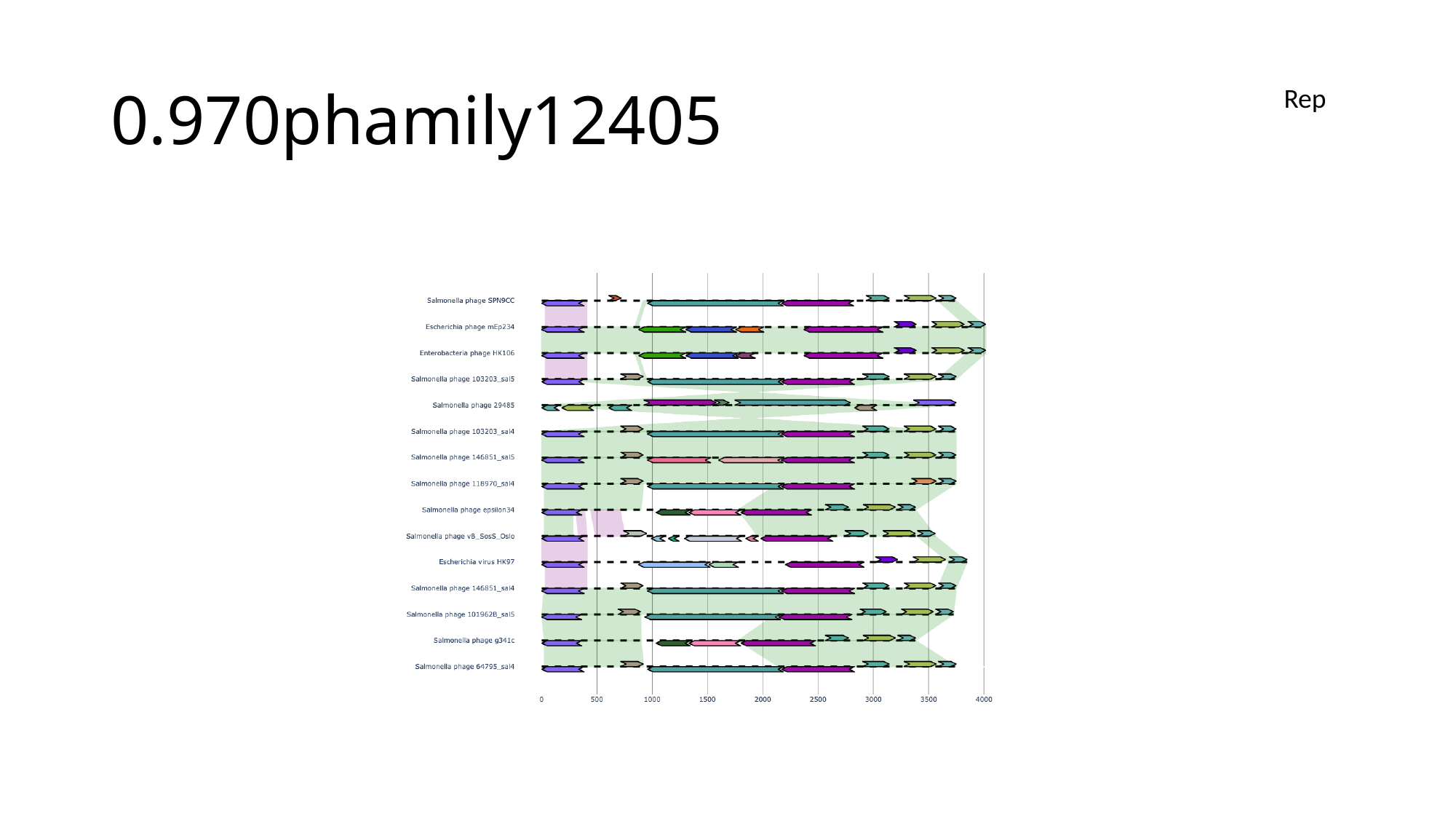

#### 0.970phamily12405
Rep

### Slide 96
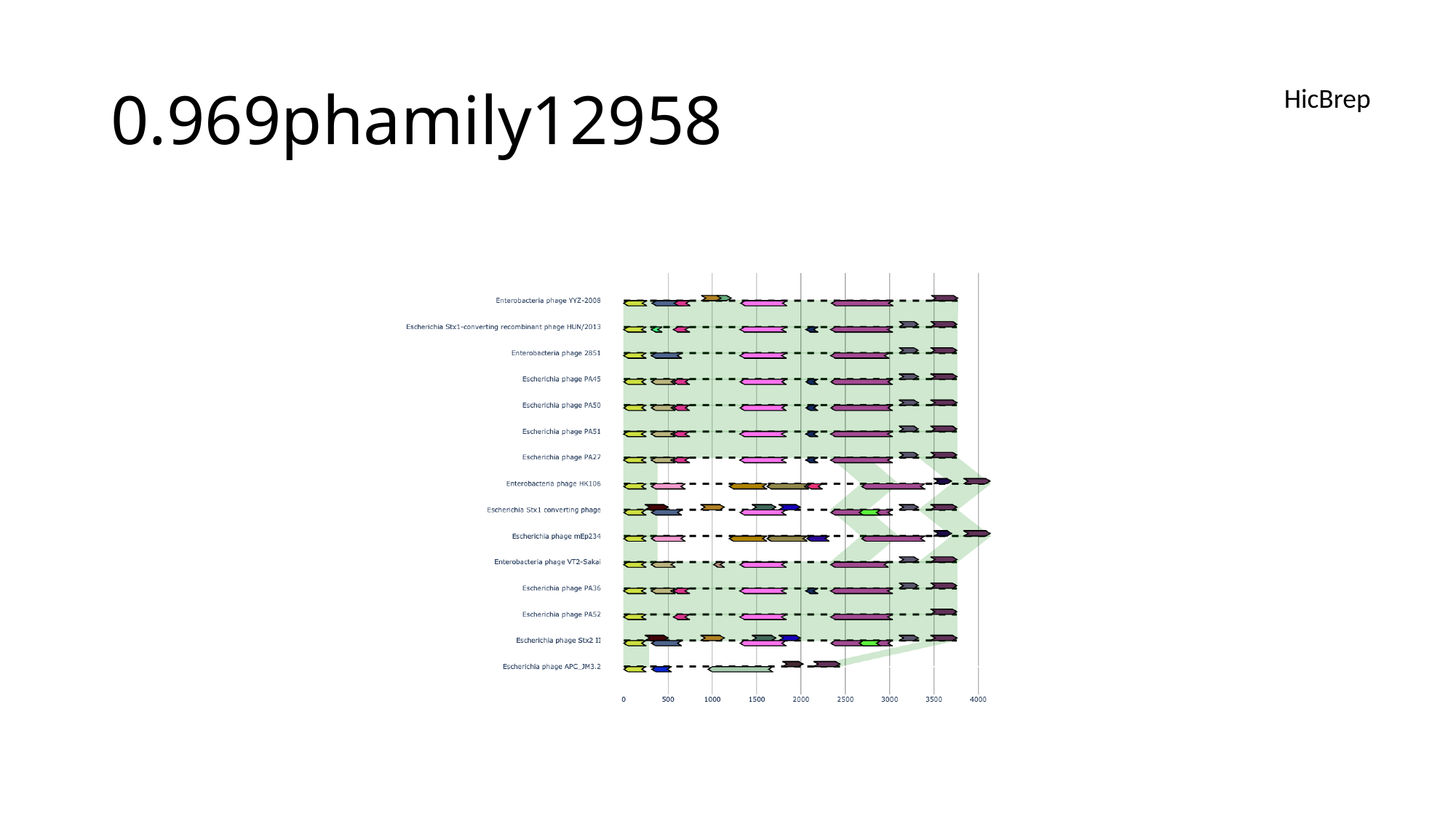

#### 0.969phamily12958
HicBrep

### Slide 97
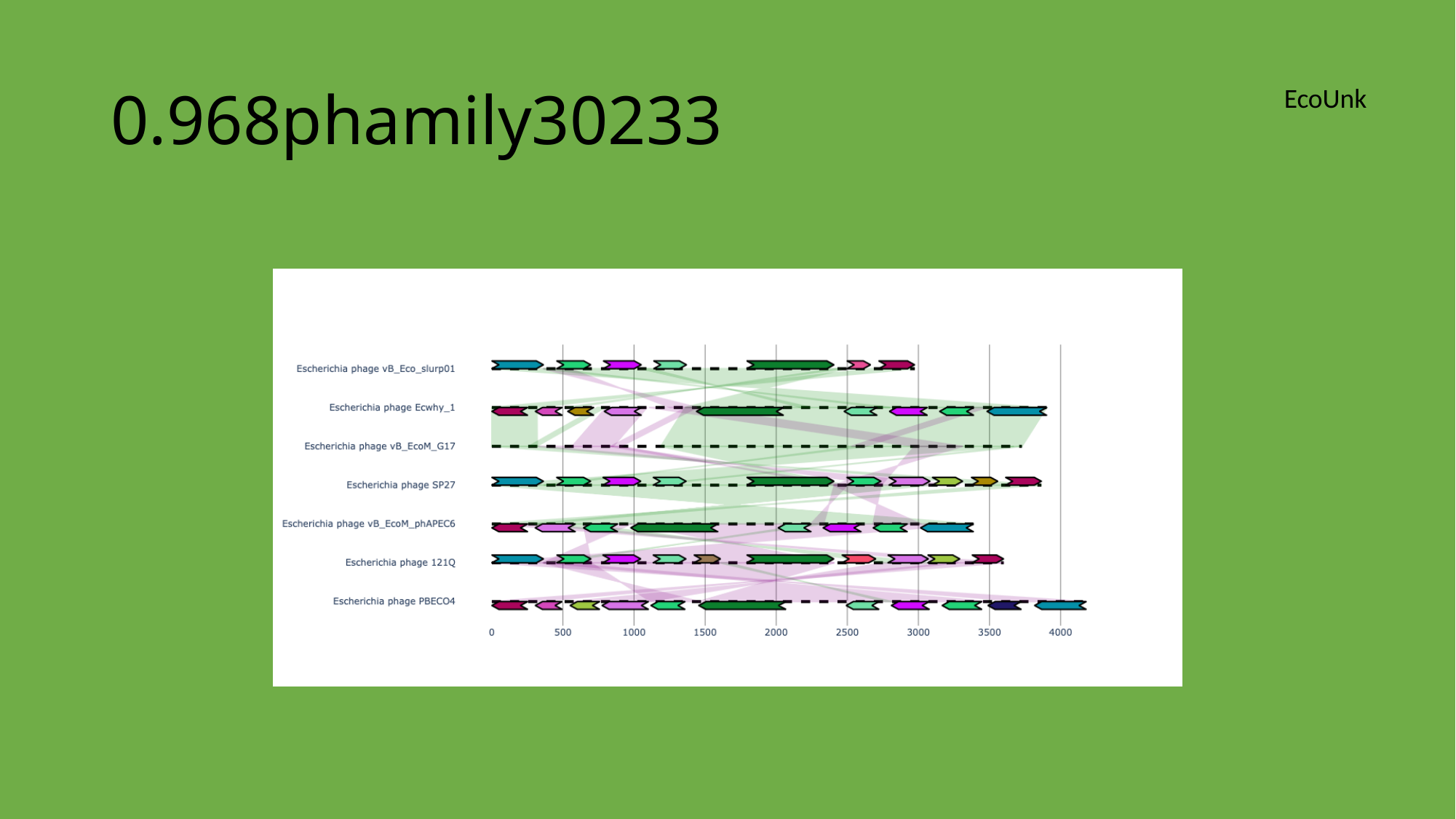

#### 0.968phamily30233
EcoUnk

### Slide 98
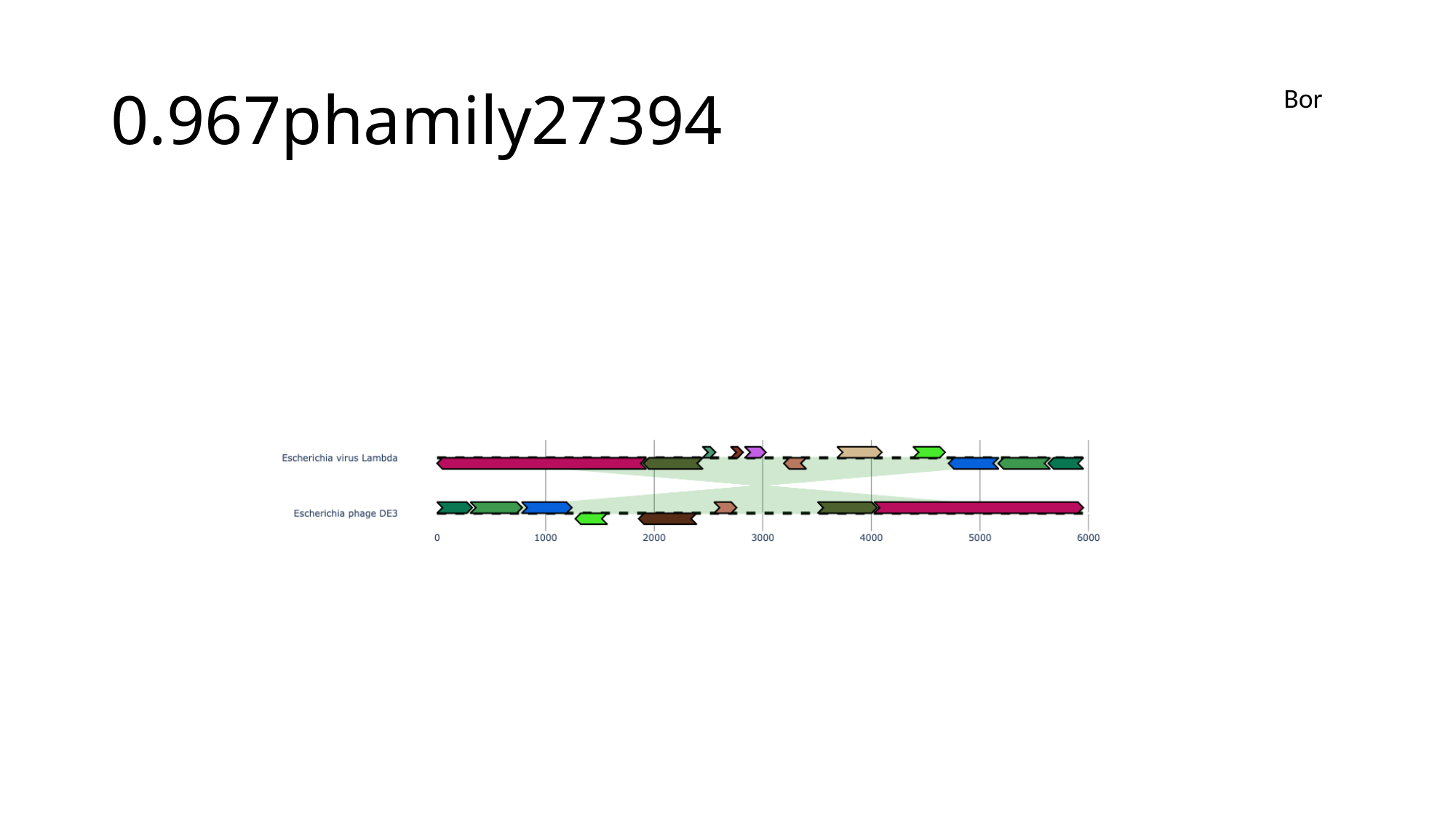

#### 0.967phamily27394
Bor

### Slide 99
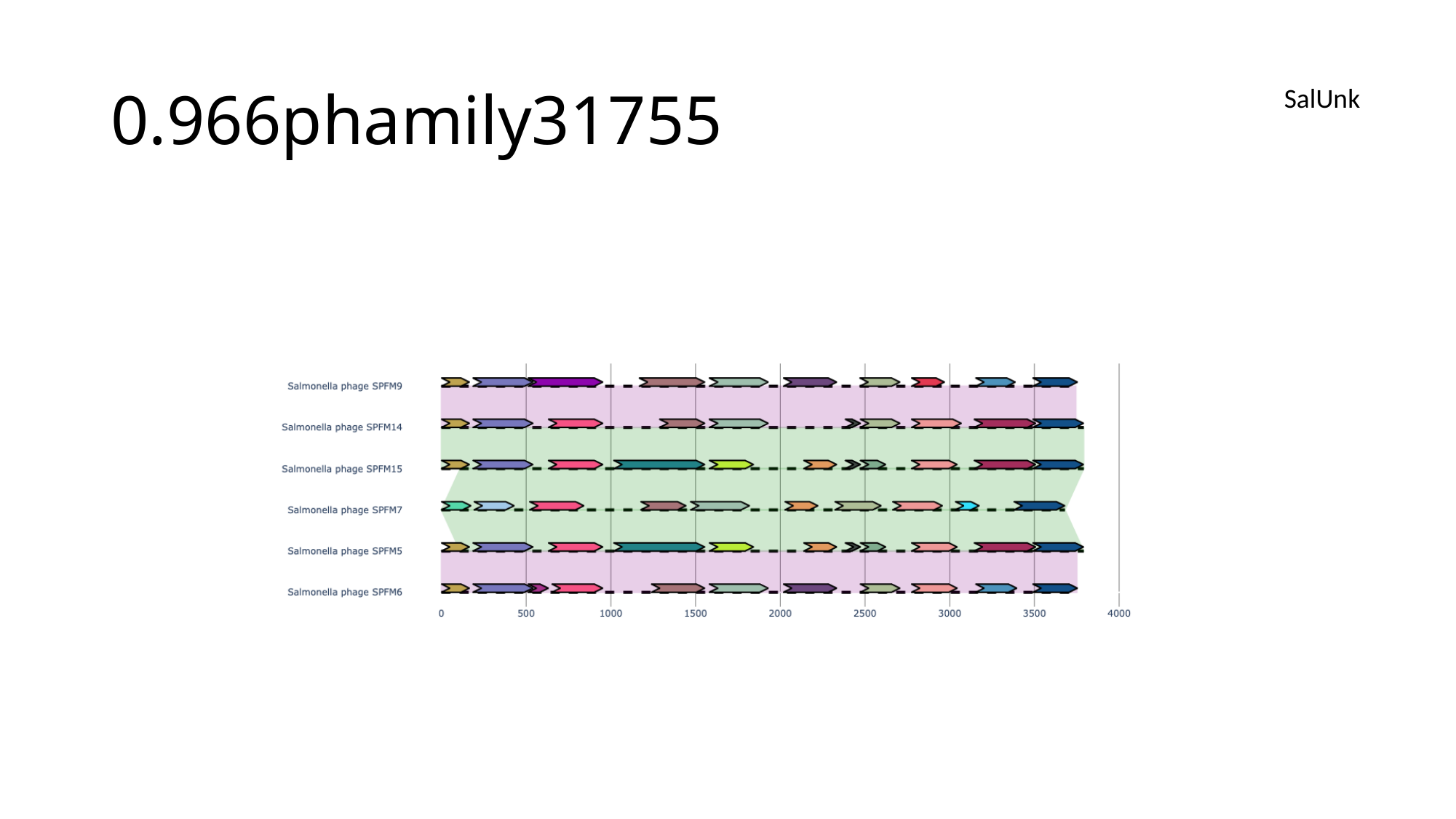

#### 0.966phamily31755
SalUnk

### Slide 100
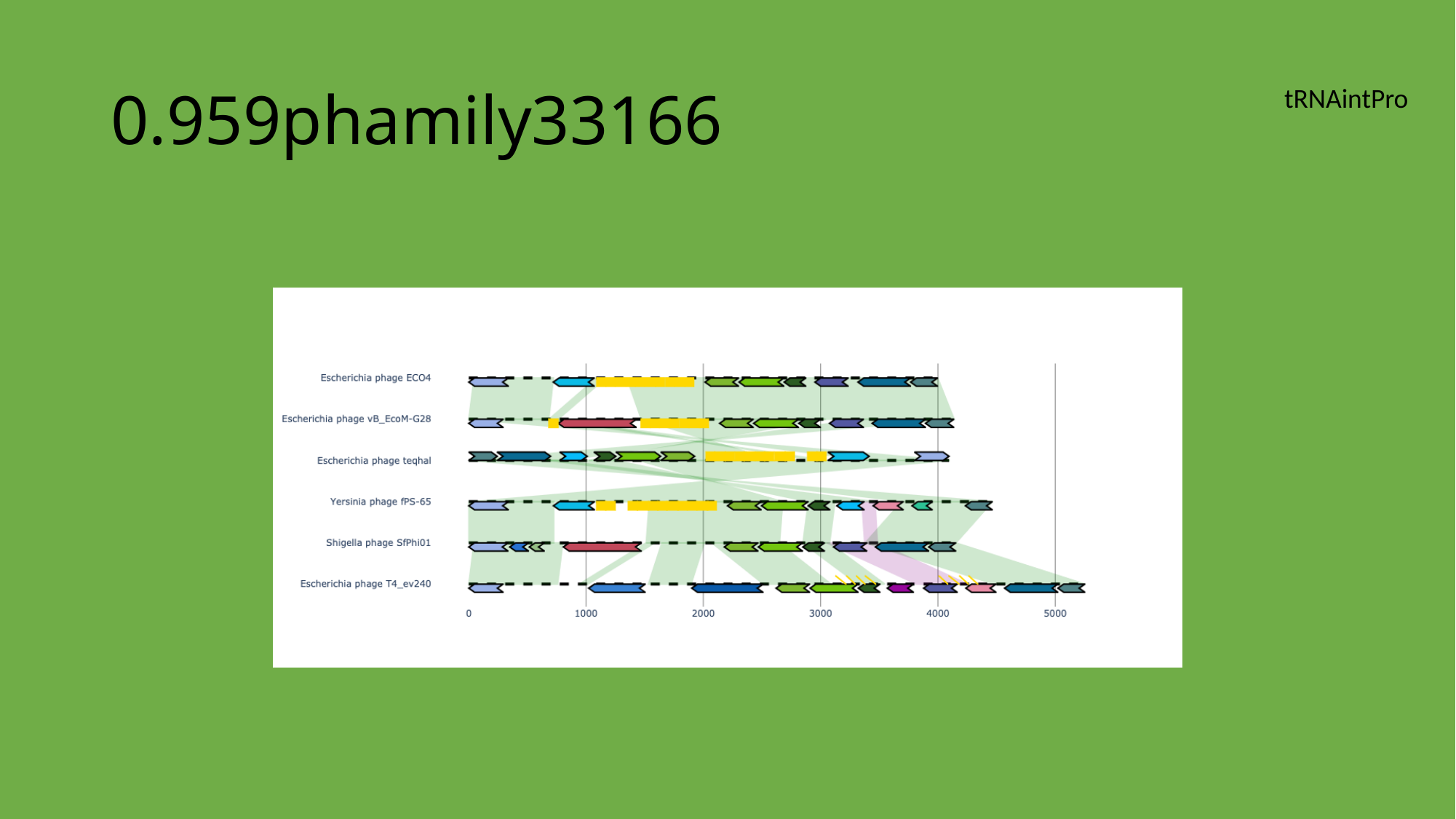

#### 0.959phamily33166
tRNAintPro

### Slide 101
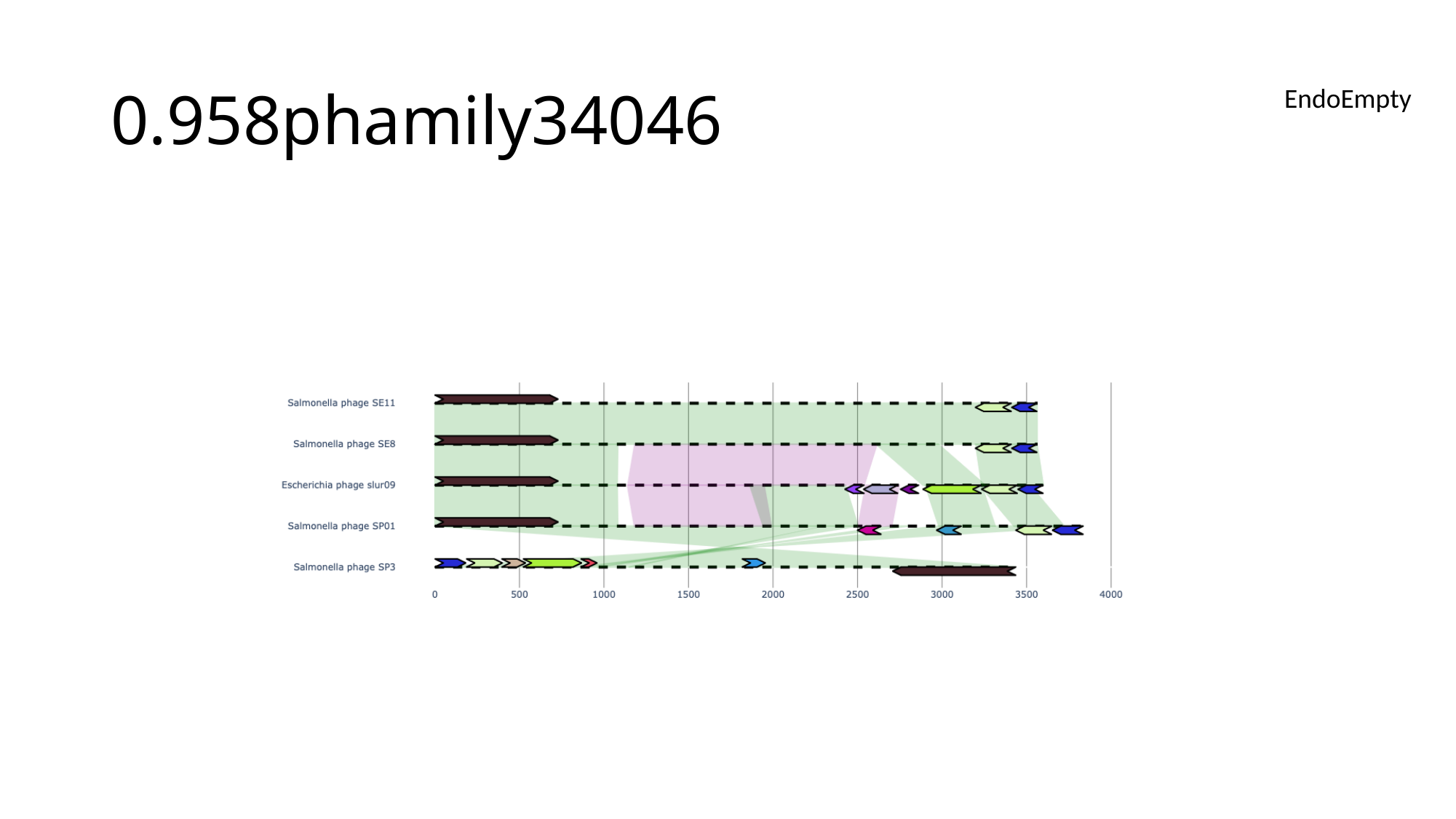

#### 0.958phamily34046
EndoEmpty

### Slide 102
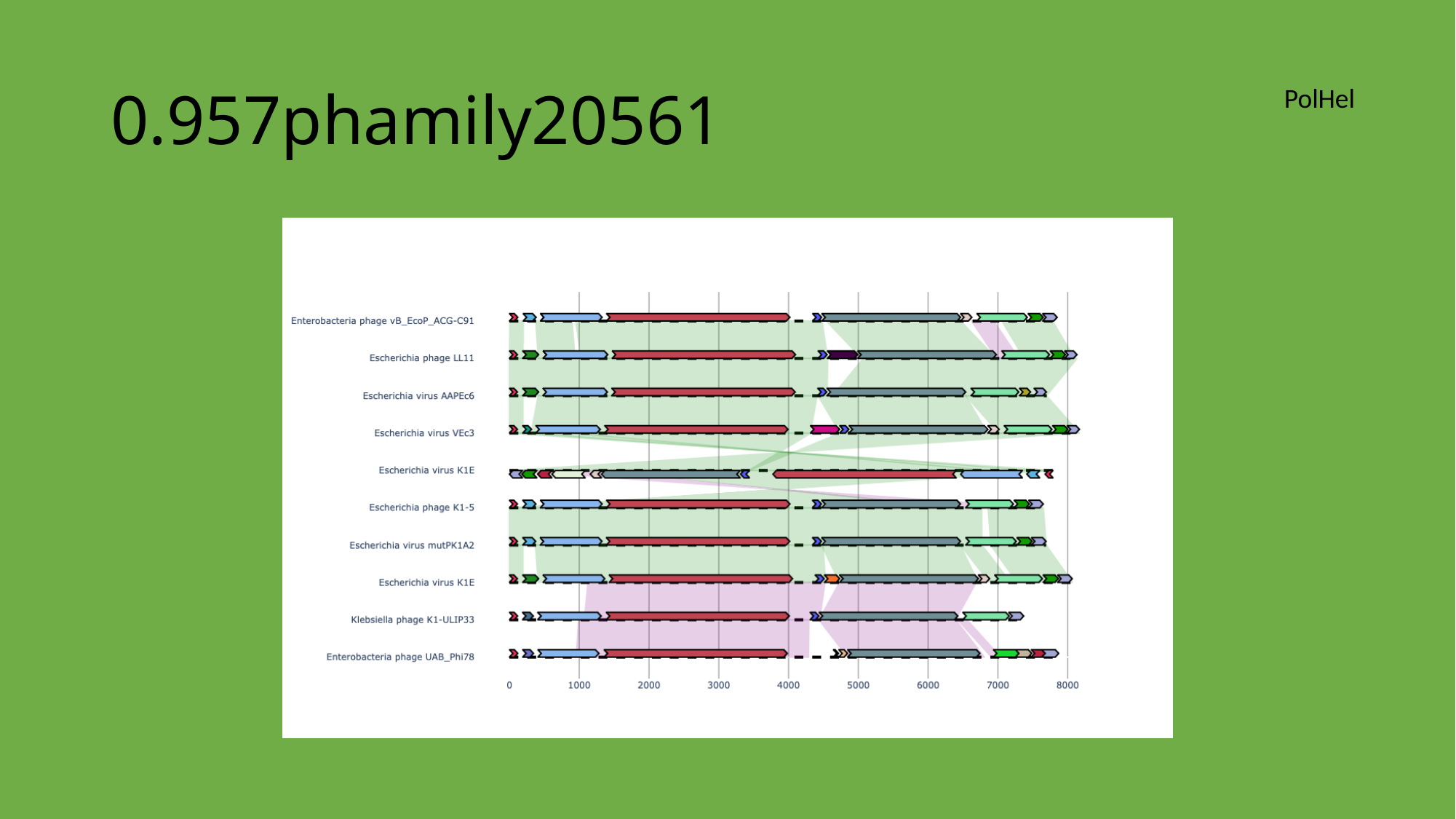

#### 0.957phamily20561
PolHel

### Slide 103
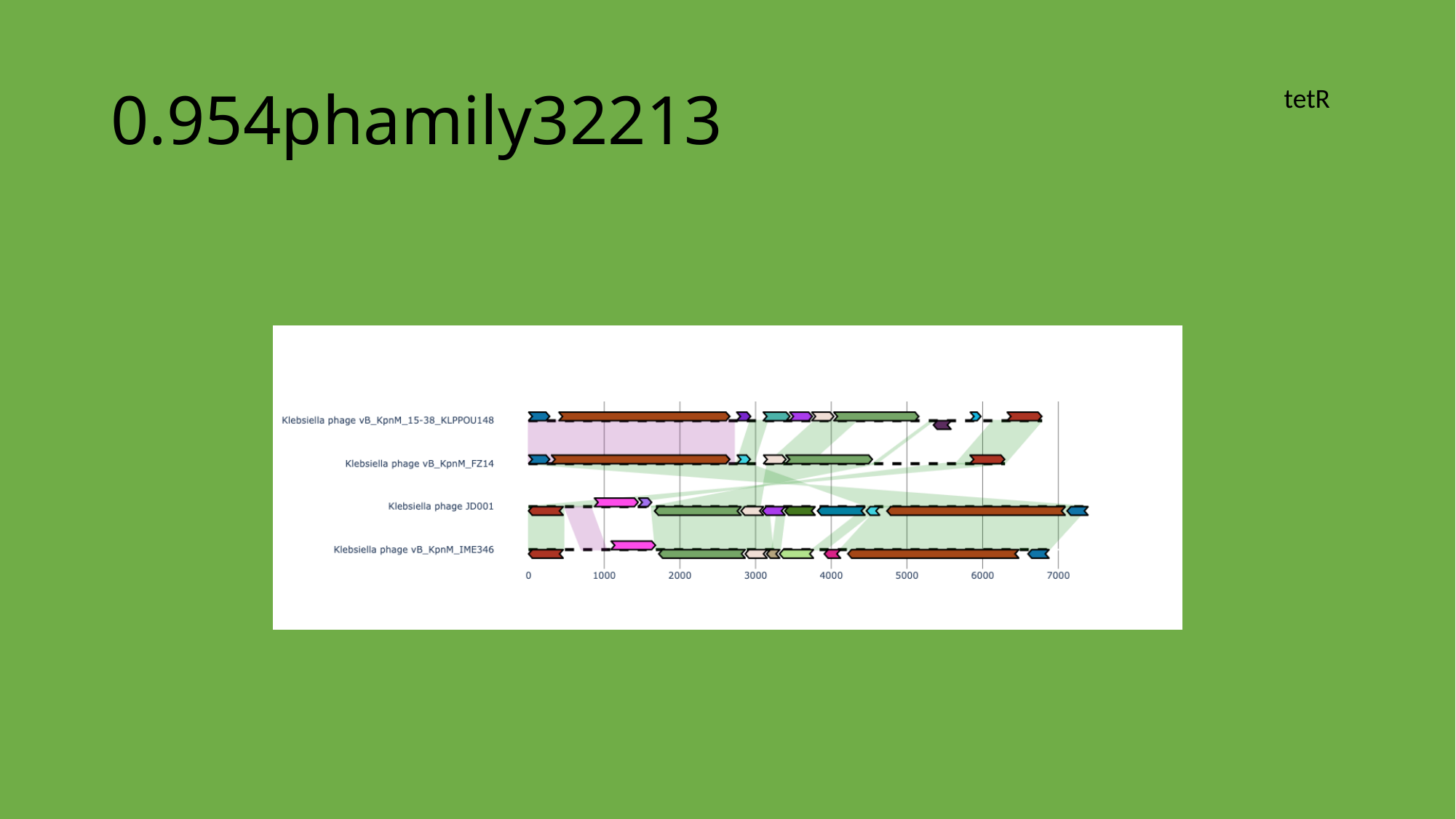

#### 0.954phamily32213
tetR

### Slide 104
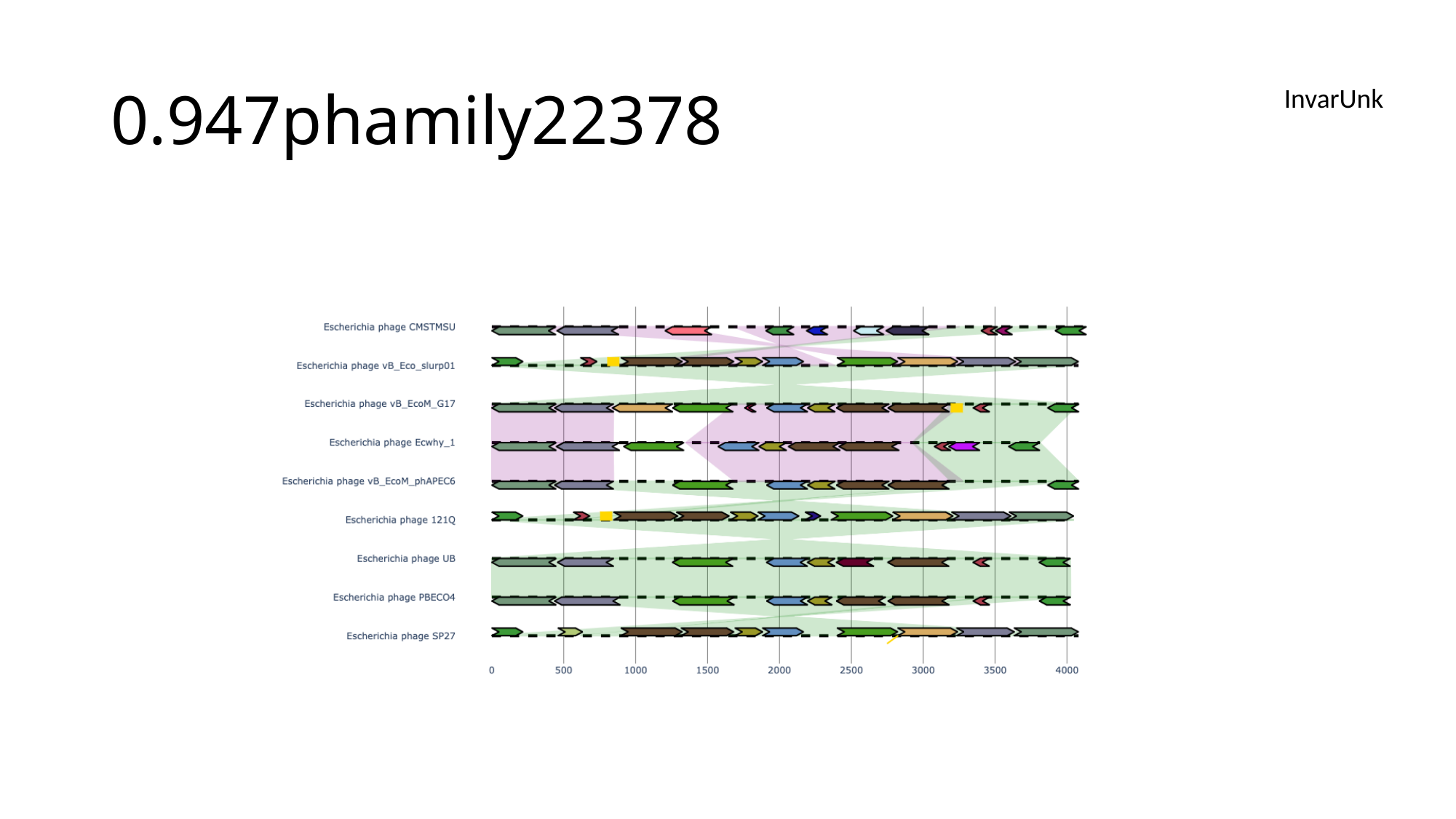

#### 0.947phamily22378
InvarUnk

### Slide 105
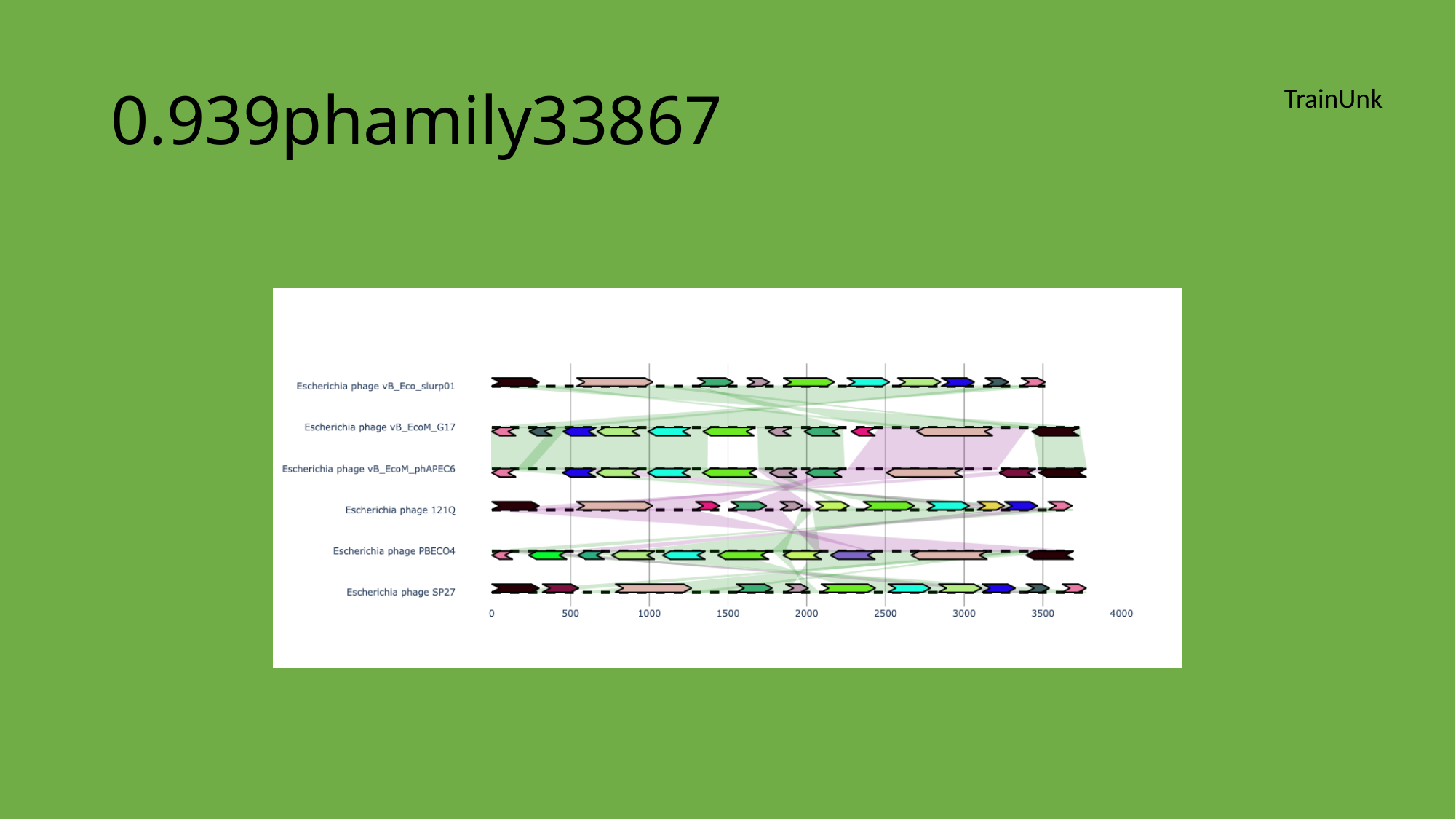

#### 0.939phamily33867
TrainUnk

### Slide 106
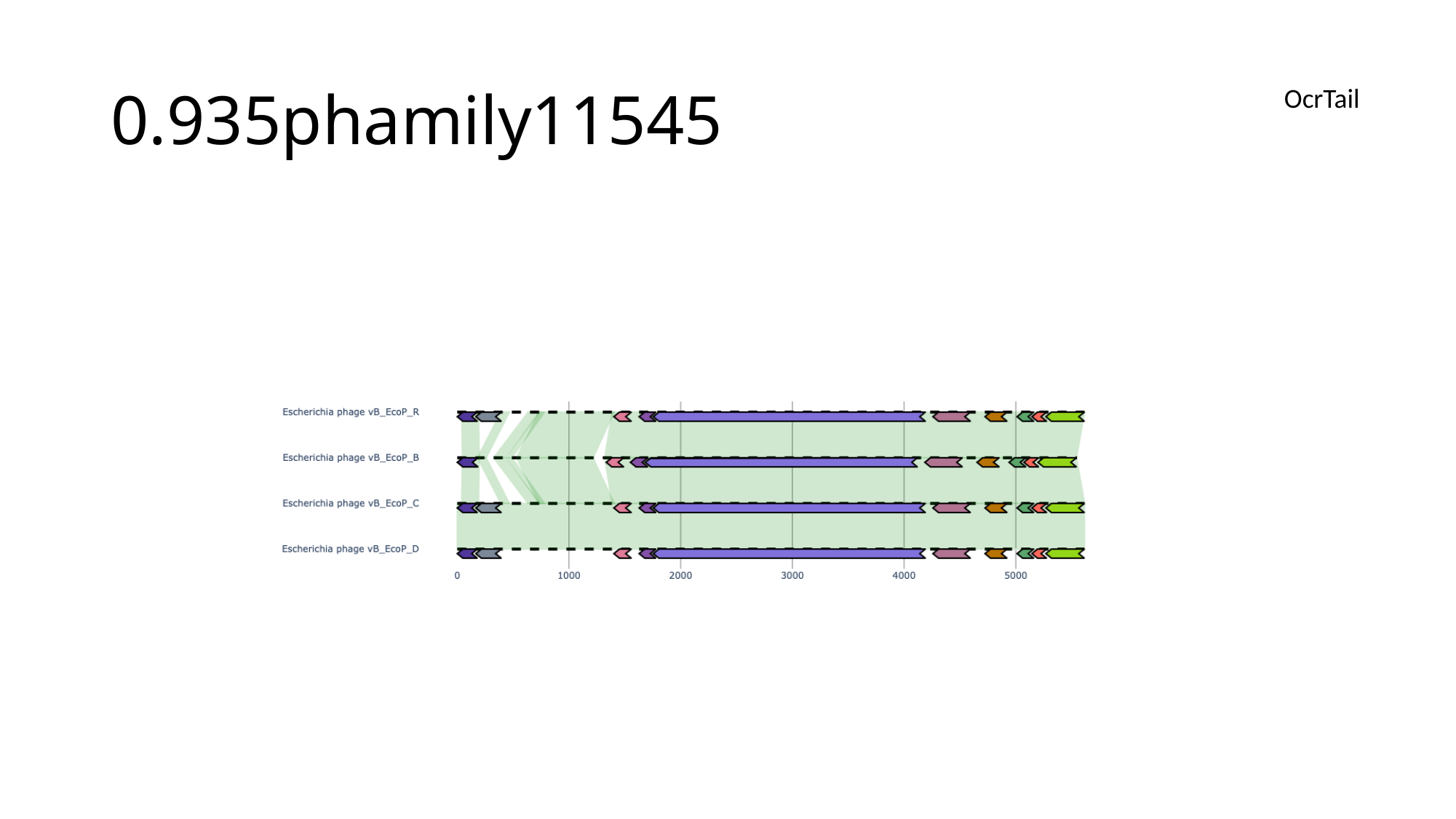

#### 0.935phamily11545
OcrTail

### Slide 107
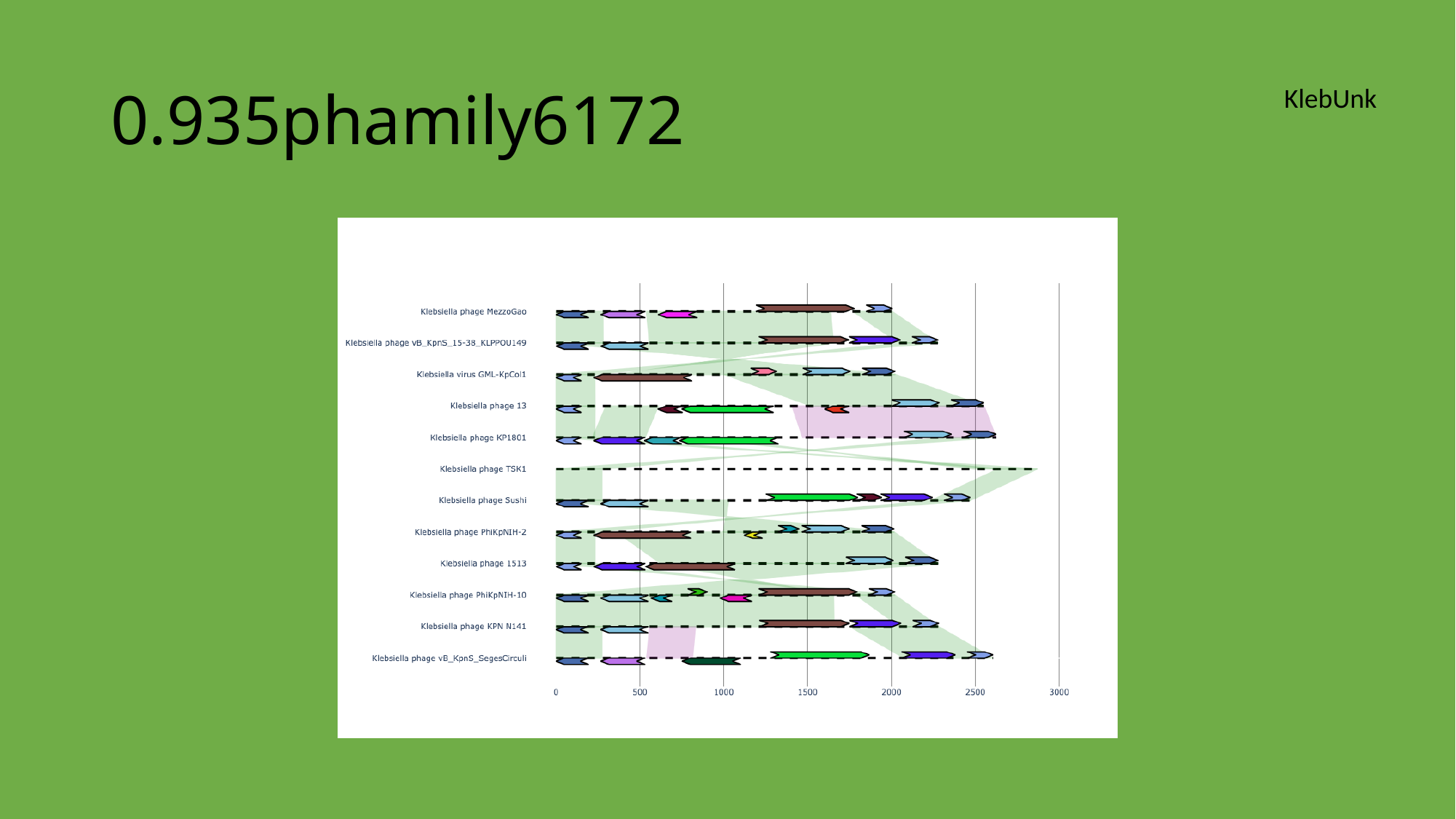

#### 0.935phamily6172
KlebUnk

### Slide 108
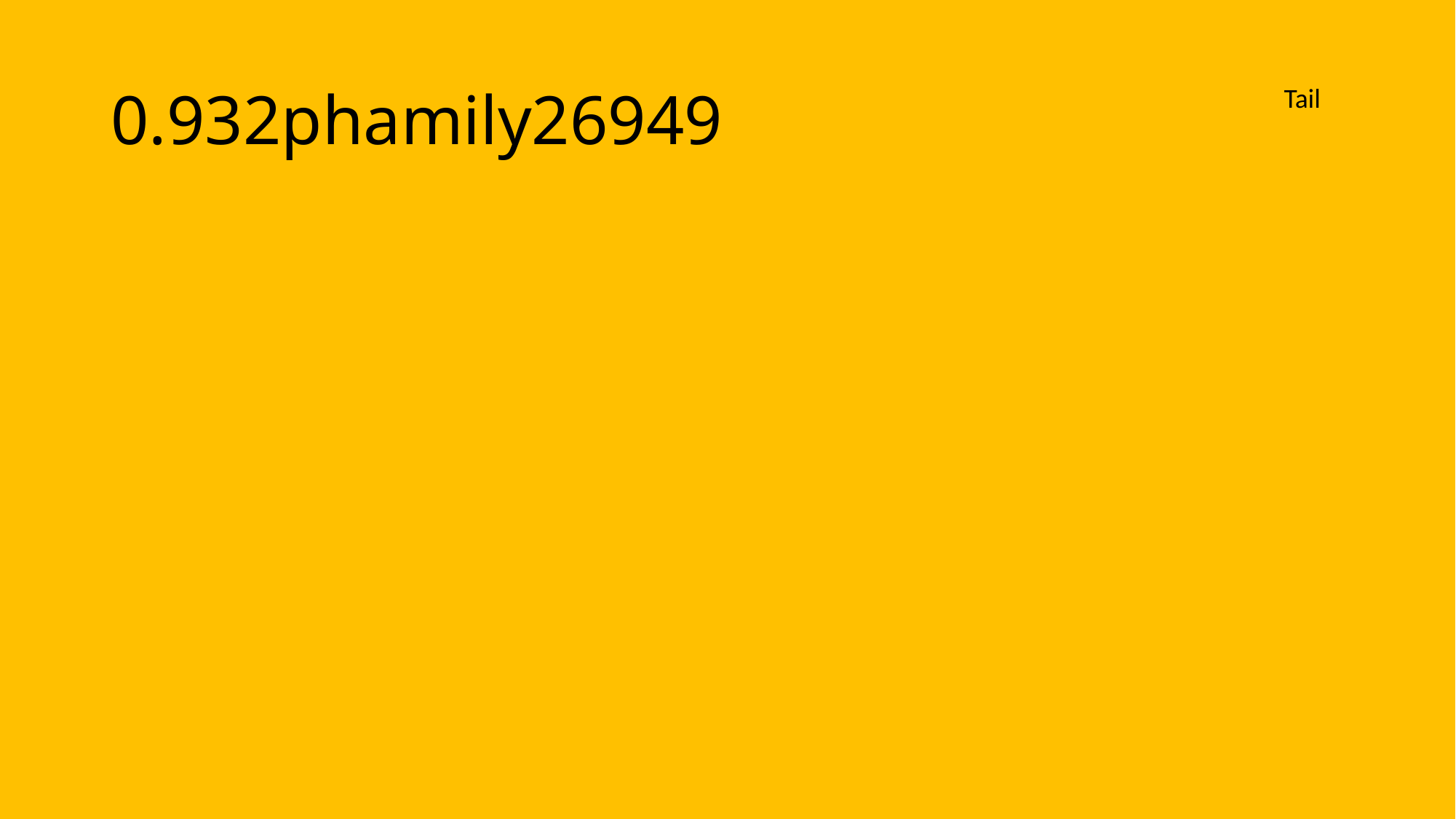

#### 0.932phamily26949
Tail

### Slide 109
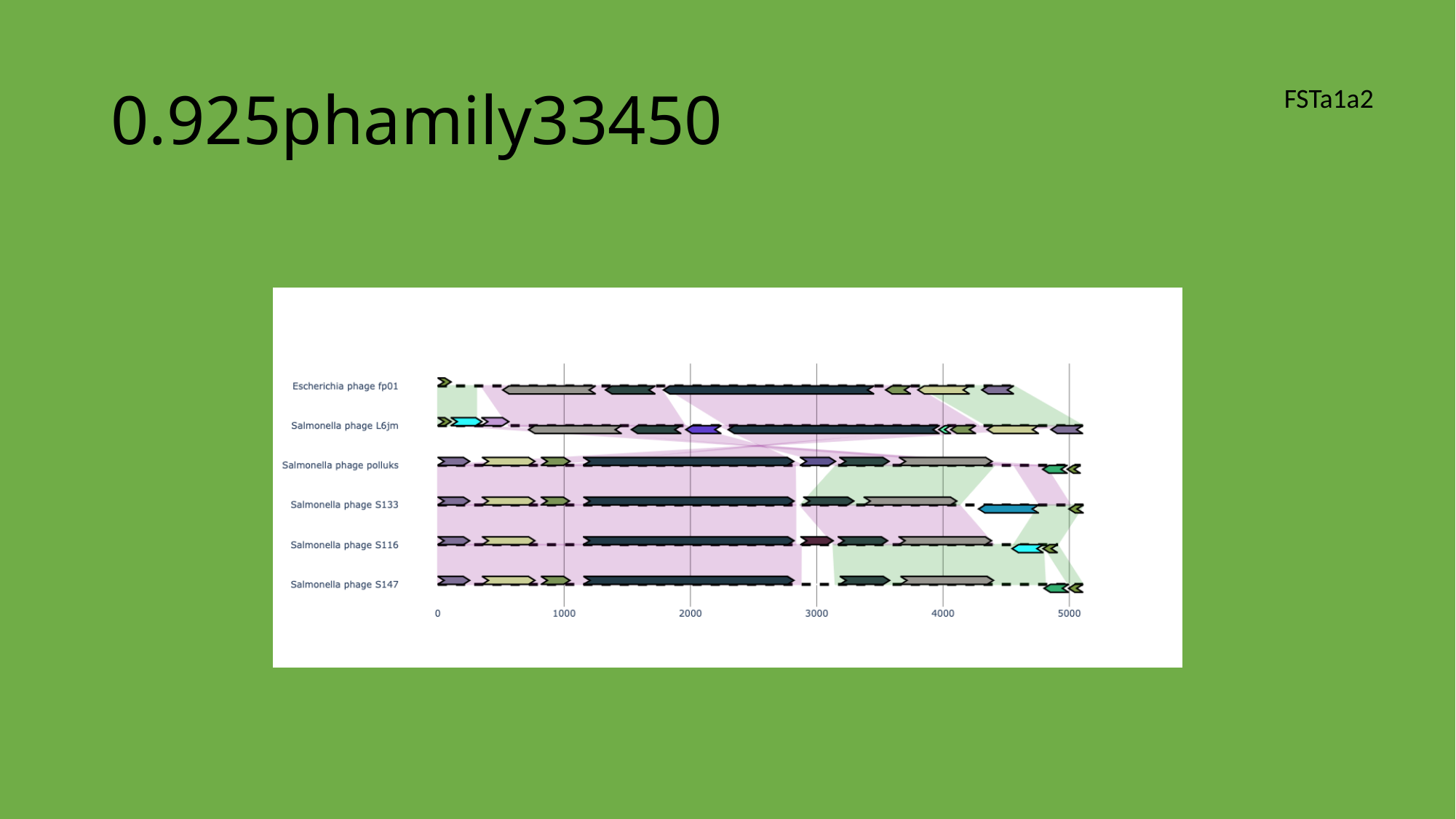

#### 0.925phamily33450
FSTa1a2

### Slide 110
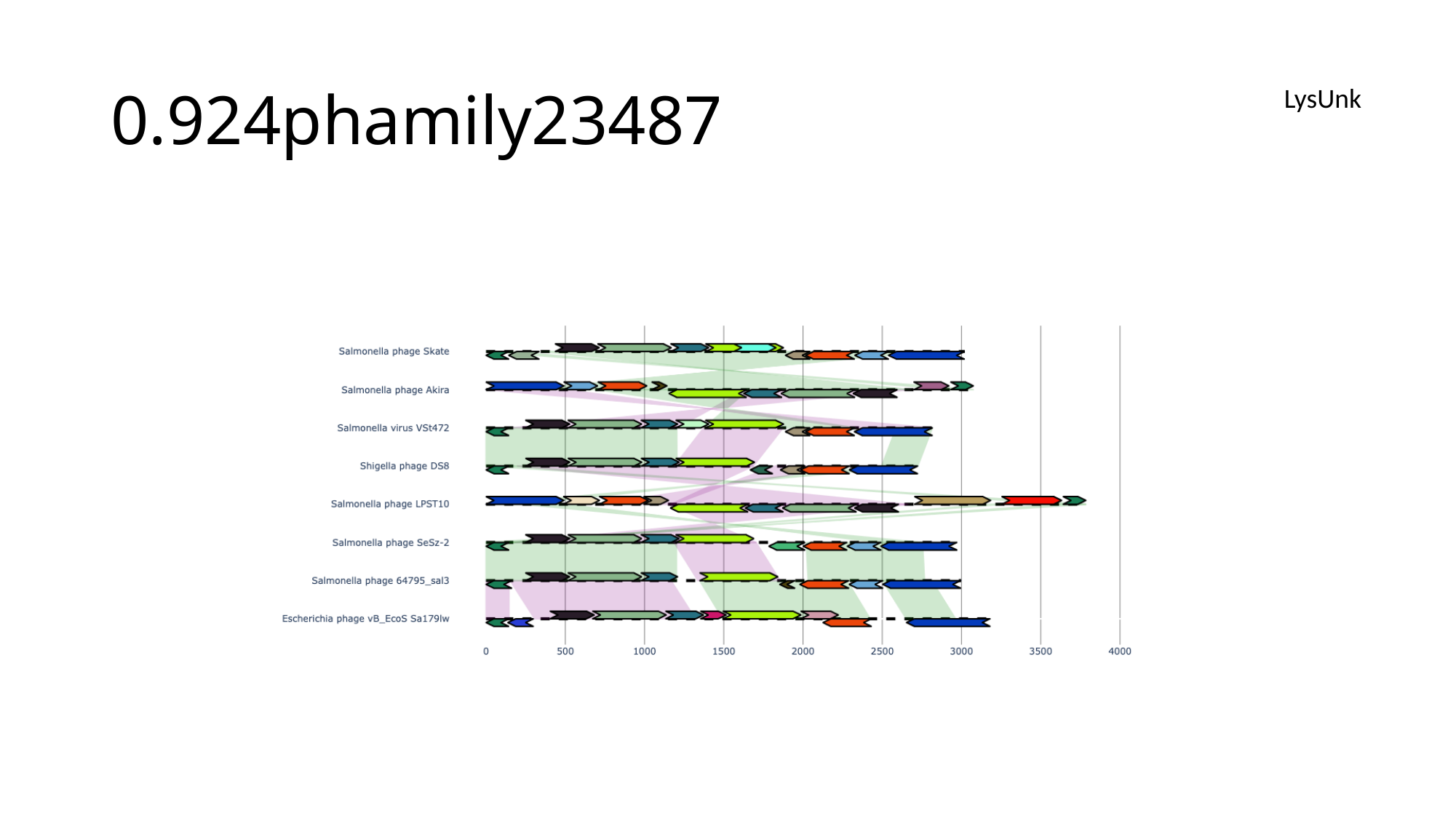

#### 0.924phamily23487
LysUnk

### Slide 111
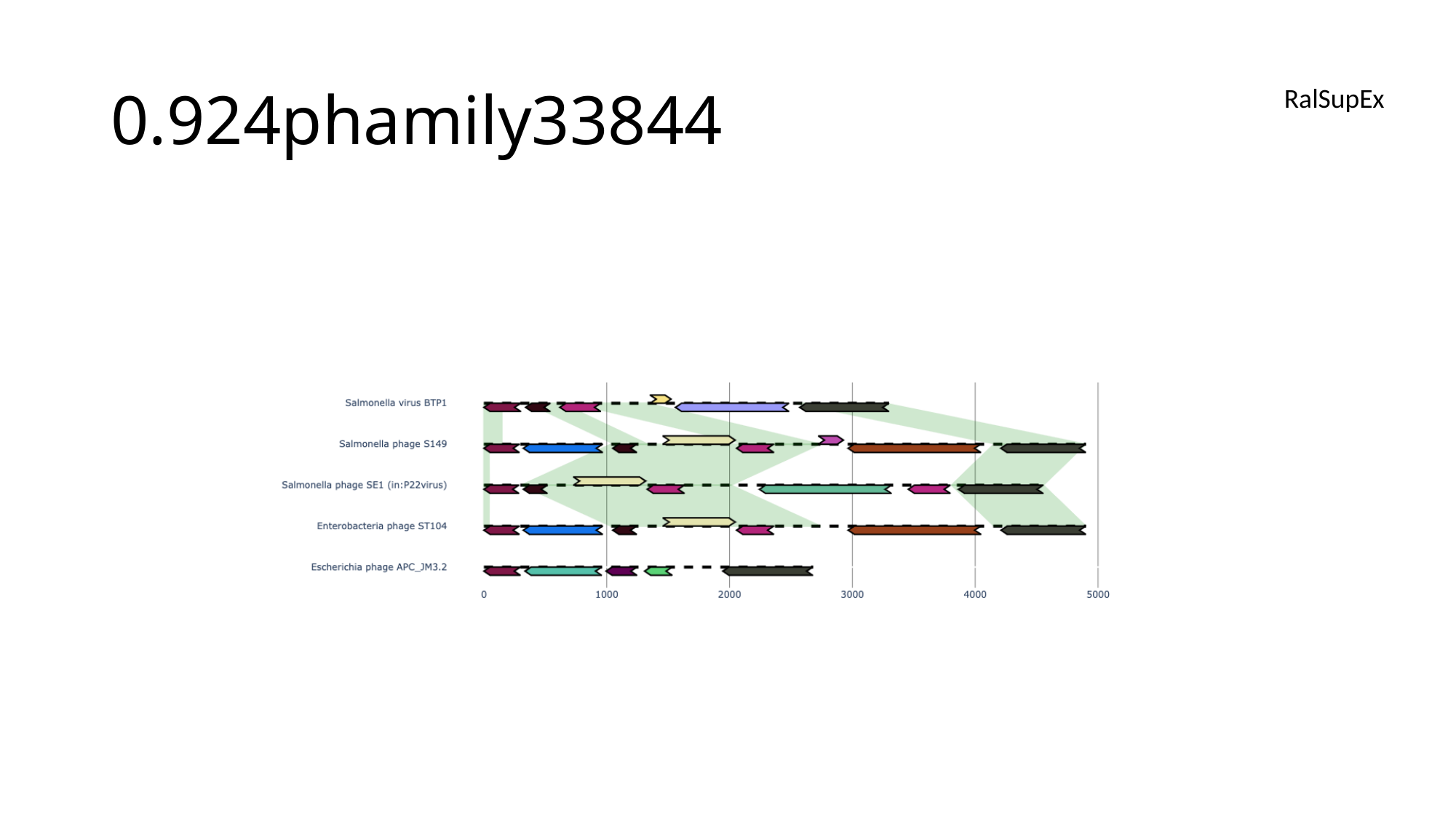

#### 0.924phamily33844
RalSupEx

### Slide 112
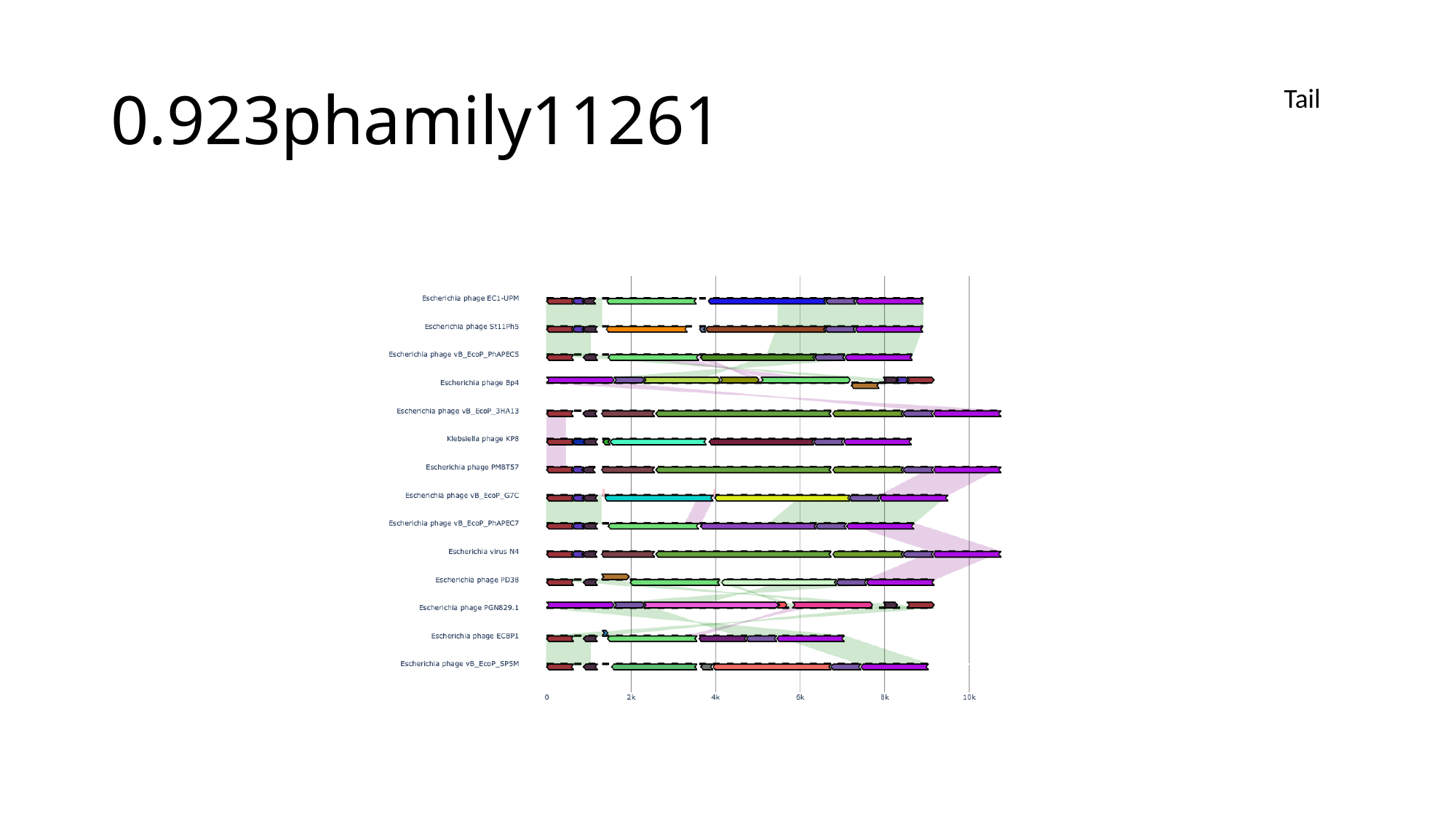

#### 0.923phamily11261
Tail

### Slide 113
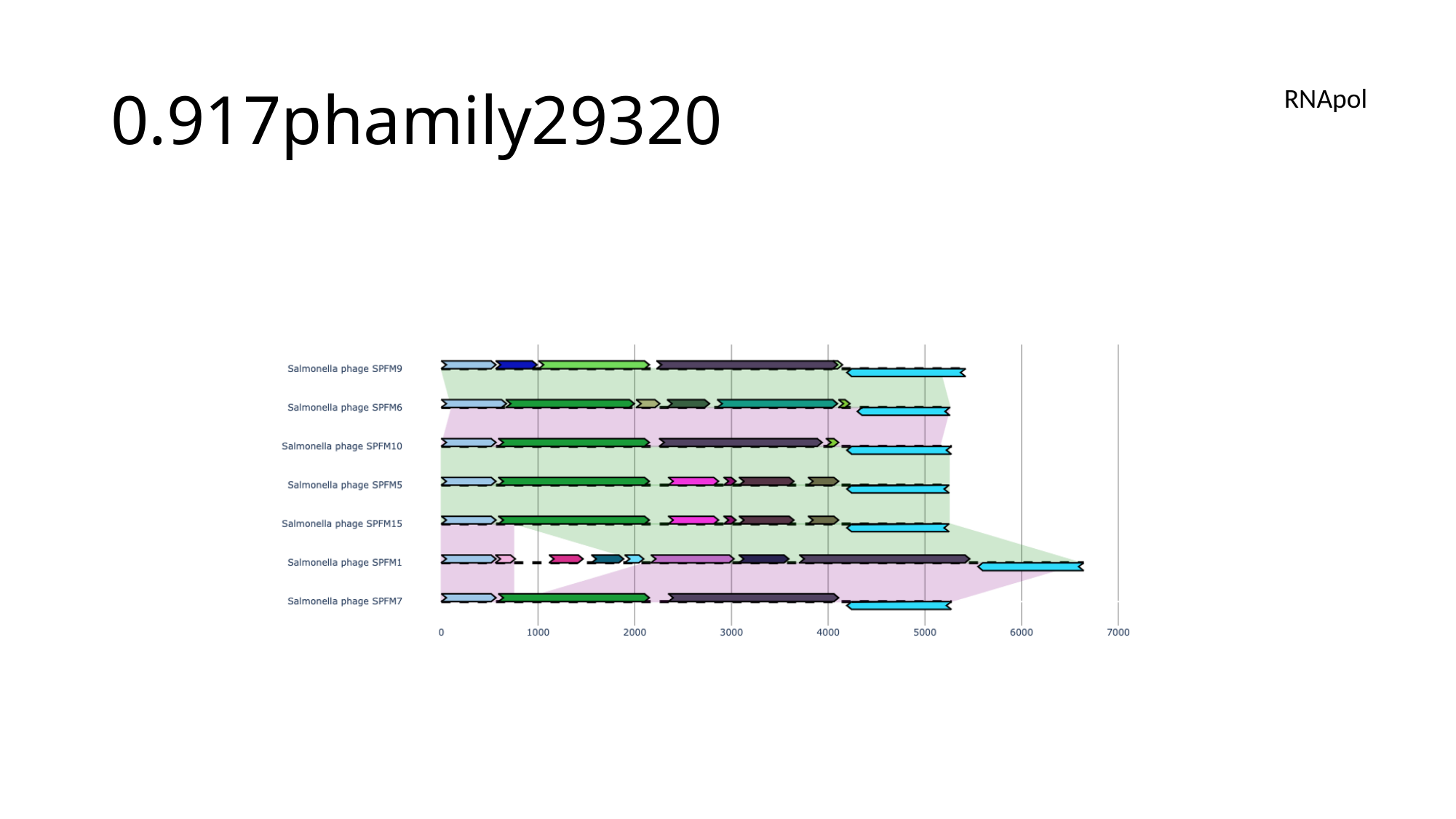

#### 0.917phamily29320
RNApol

### Slide 114
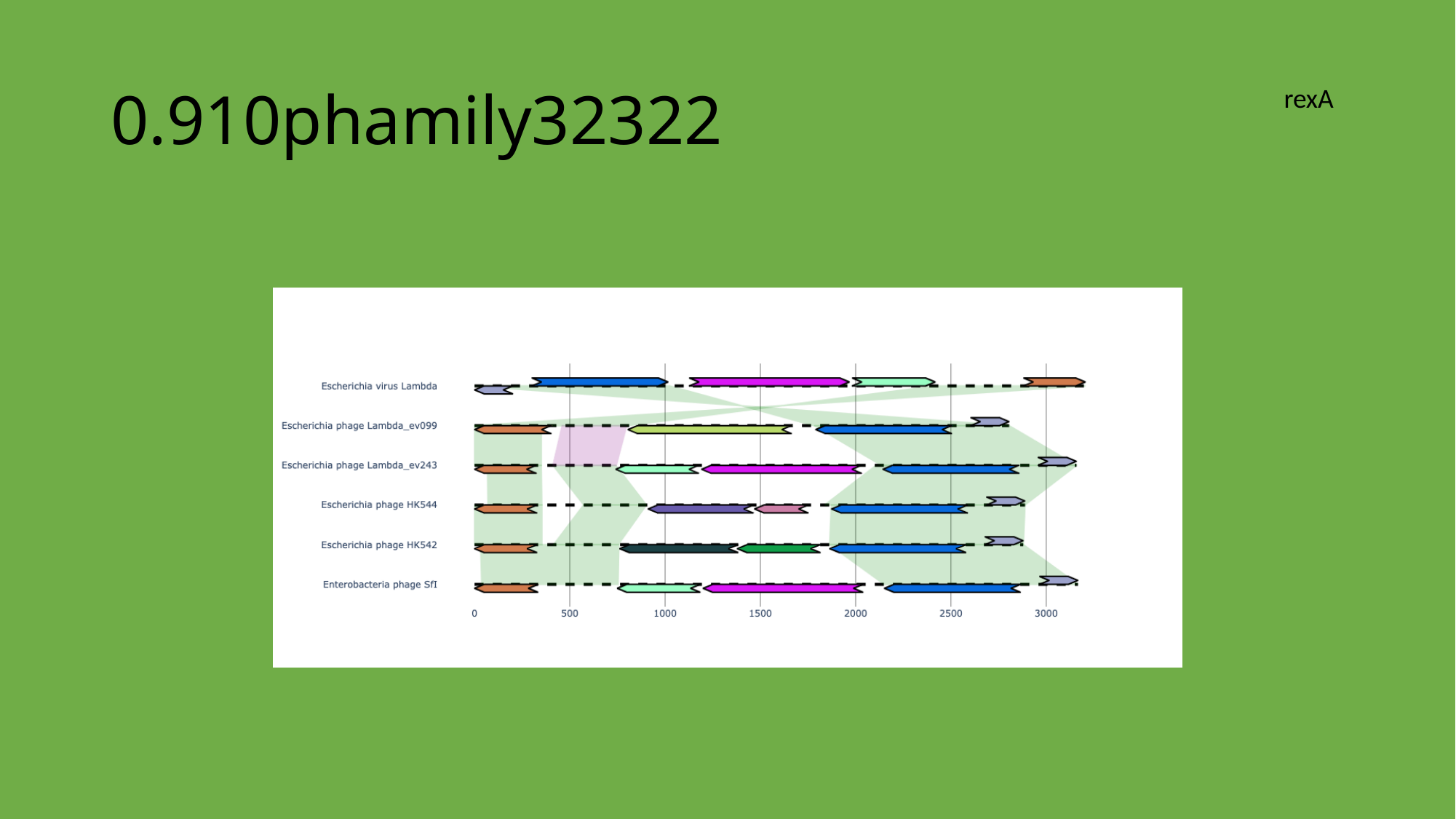

#### 0.910phamily32322
rexA

### Slide 115
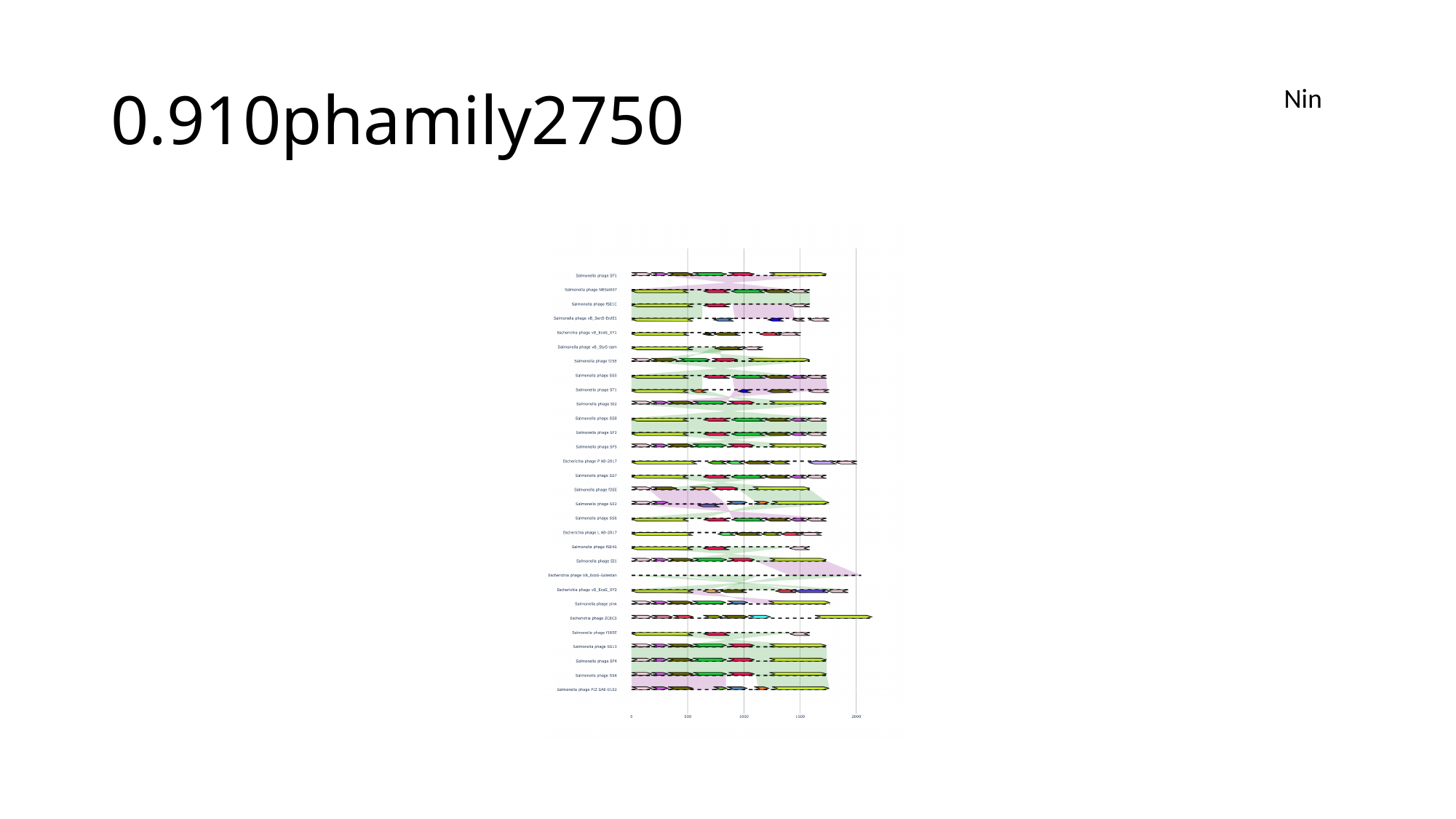

#### 0.910phamily2750
Nin

### Slide 116
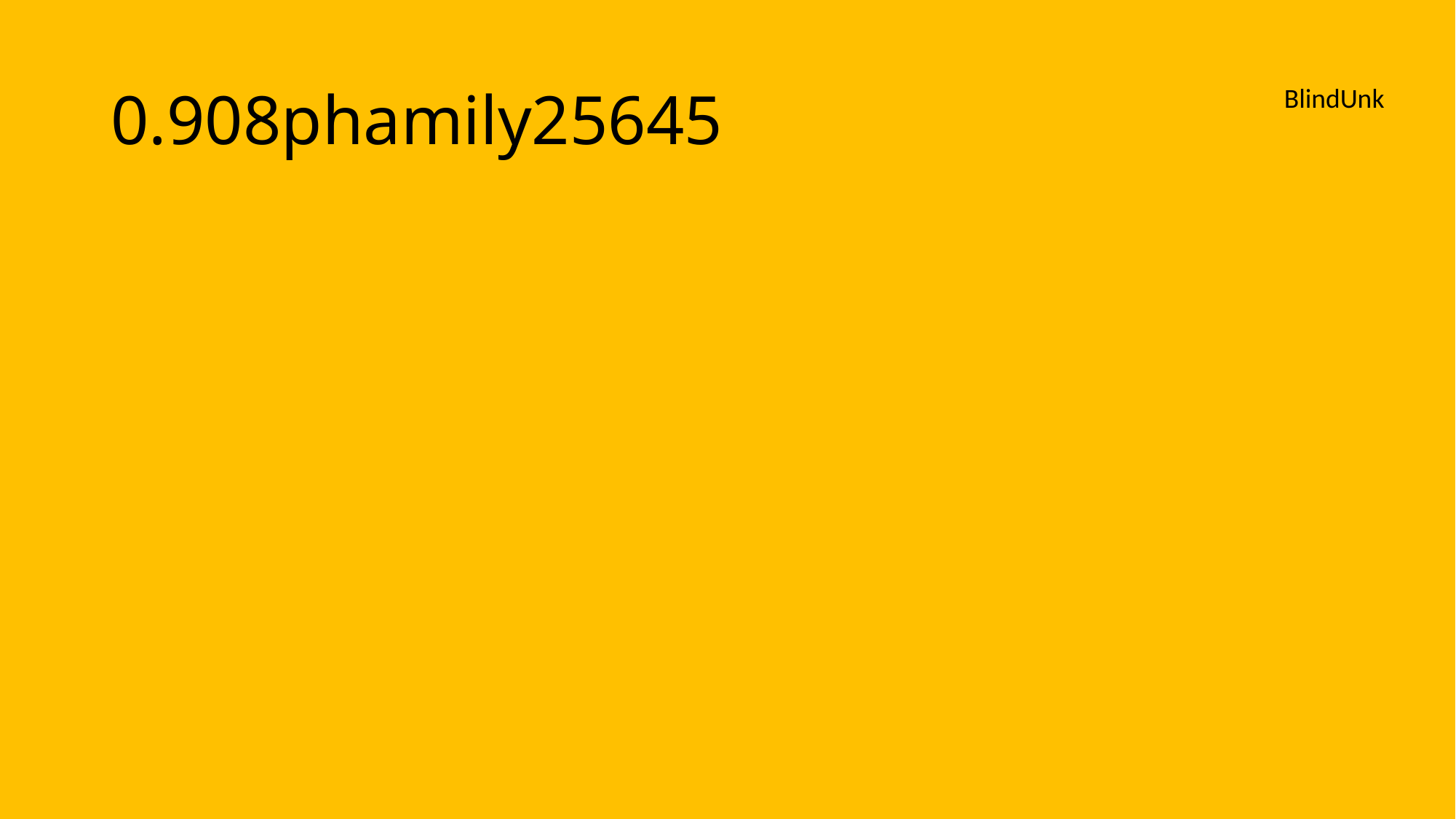

#### 0.908phamily25645
BlindUnk

### Slide 117
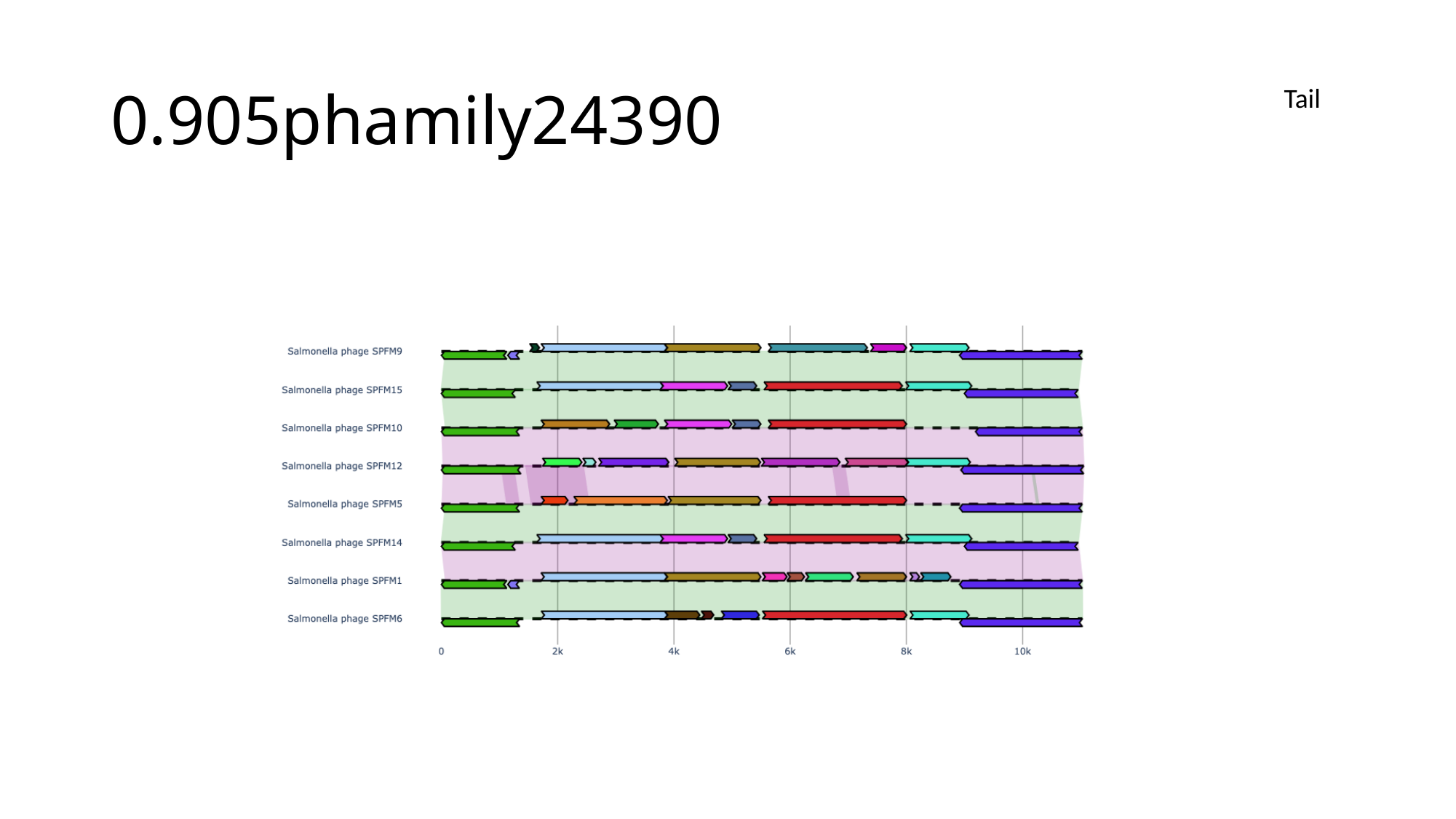

#### 0.905phamily24390
Tail

### Slide 118
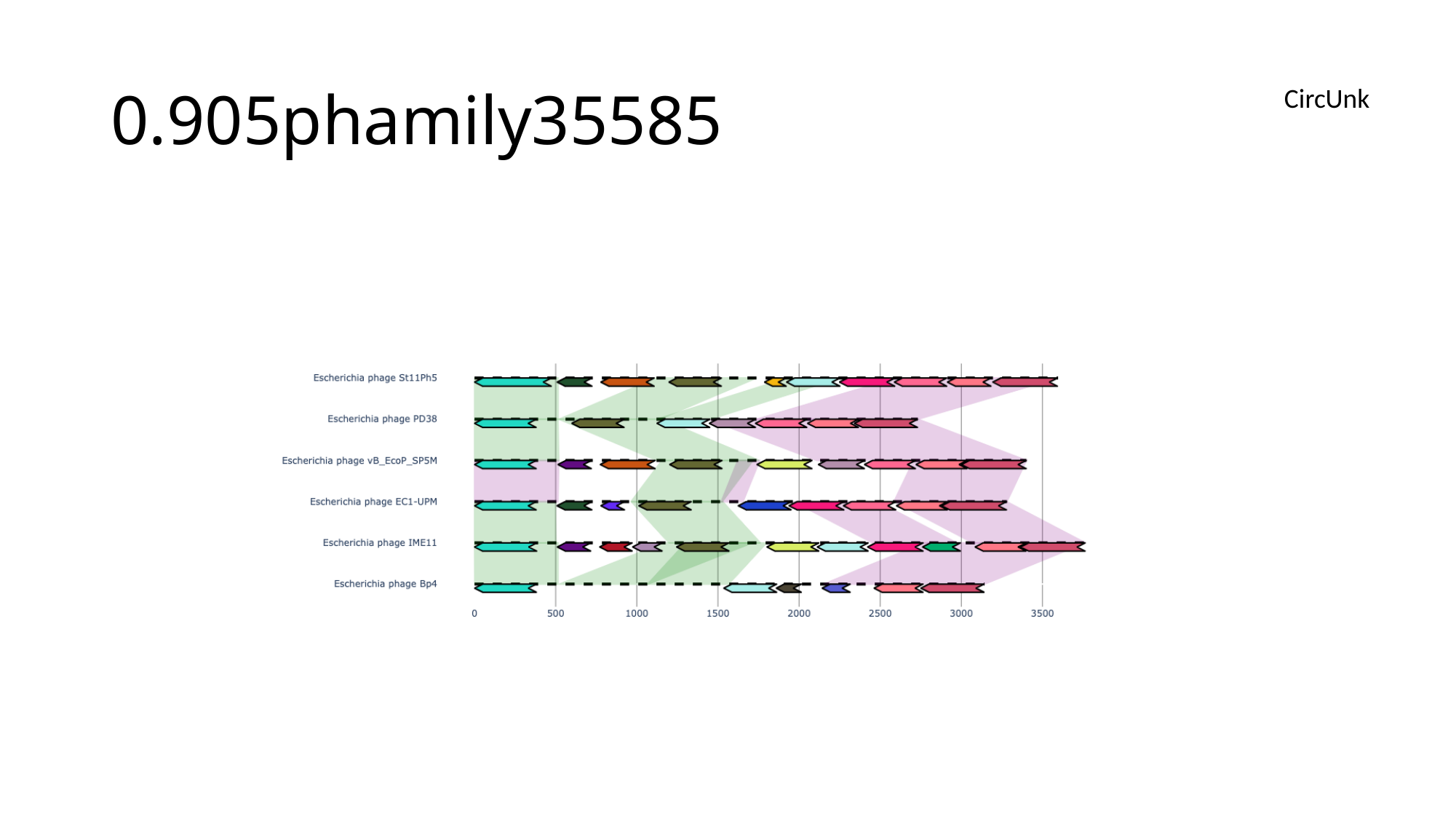

#### 0.905phamily35585
CircUnk

### Slide 119
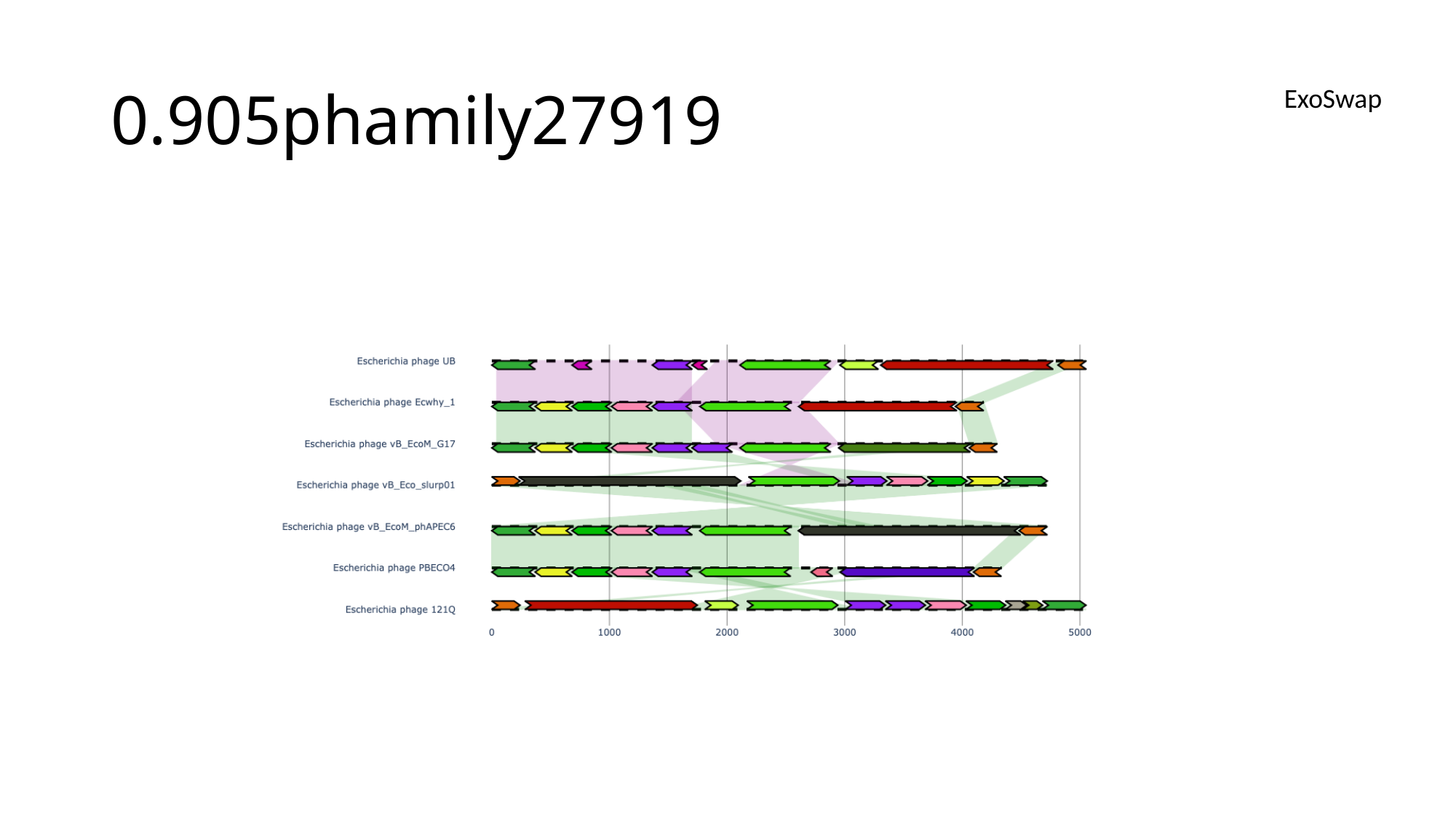

#### 0.905phamily27919
ExoSwap

### Slide 120
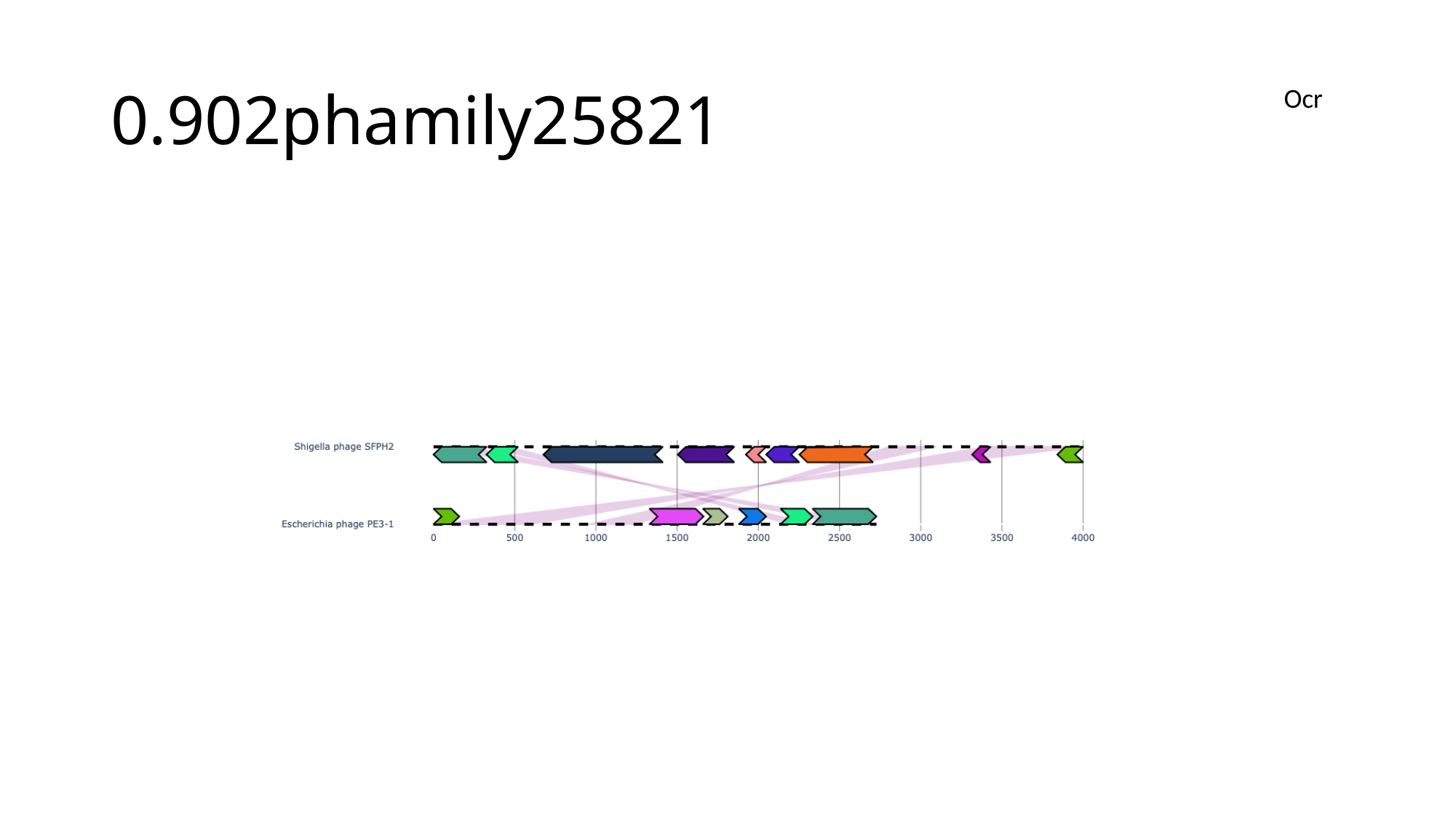

#### 0.902phamily25821
Ocr

### Slide 121
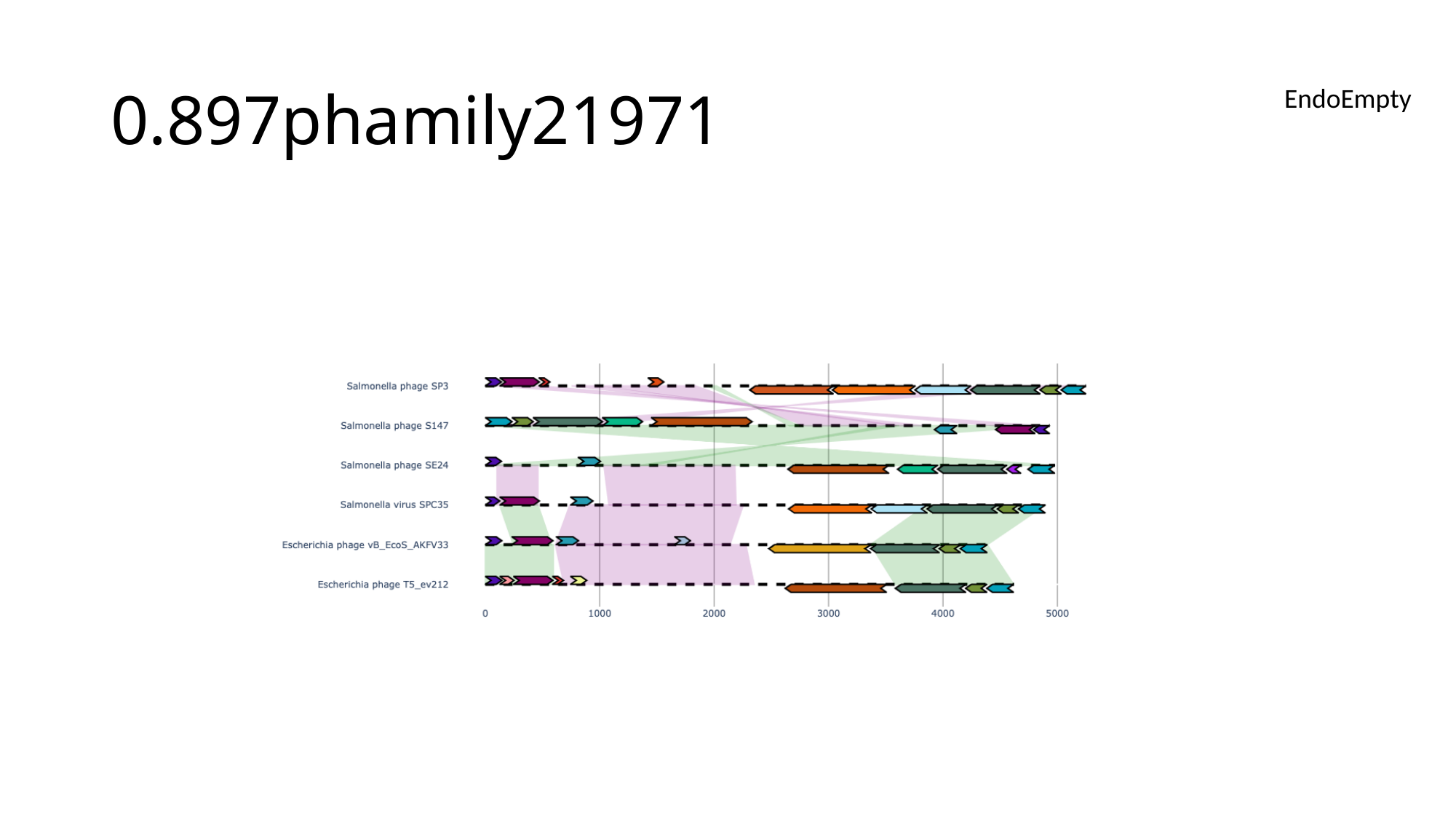

#### 0.897phamily21971
EndoEmpty

### Slide 122
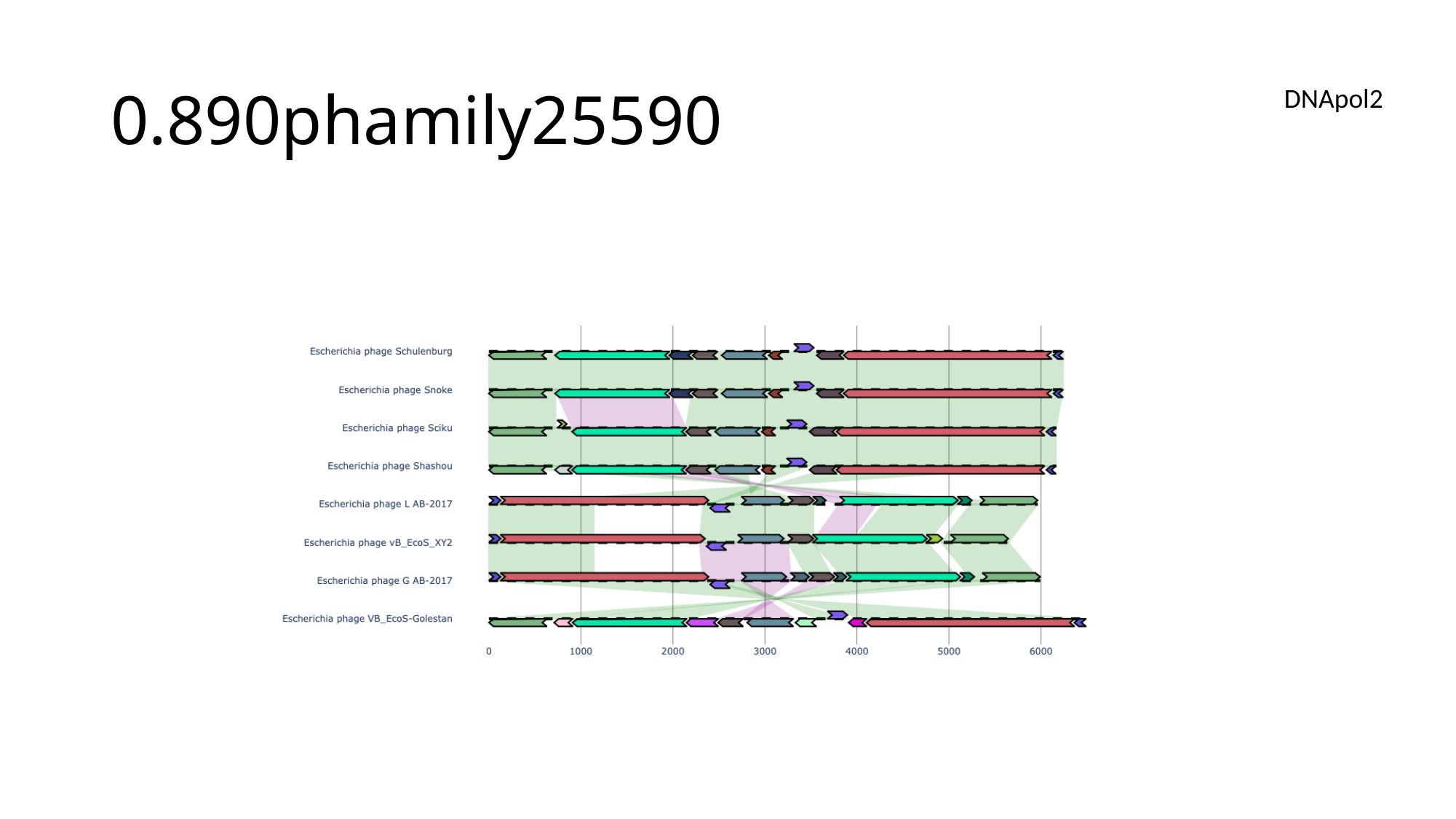

#### 0.890phamily25590
DNApol2

### Slide 123
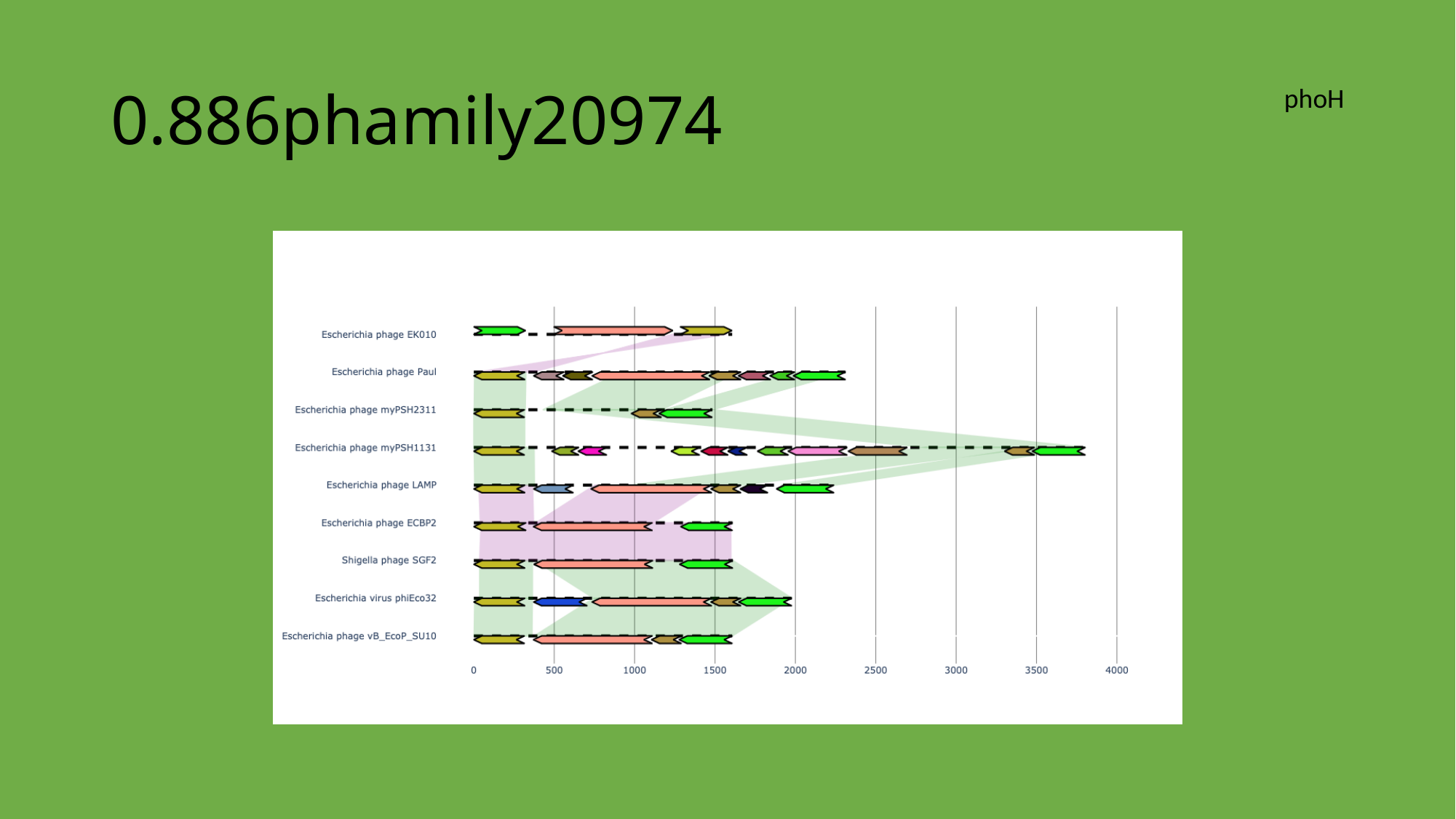

#### 0.886phamily20974
phoH

### Slide 124
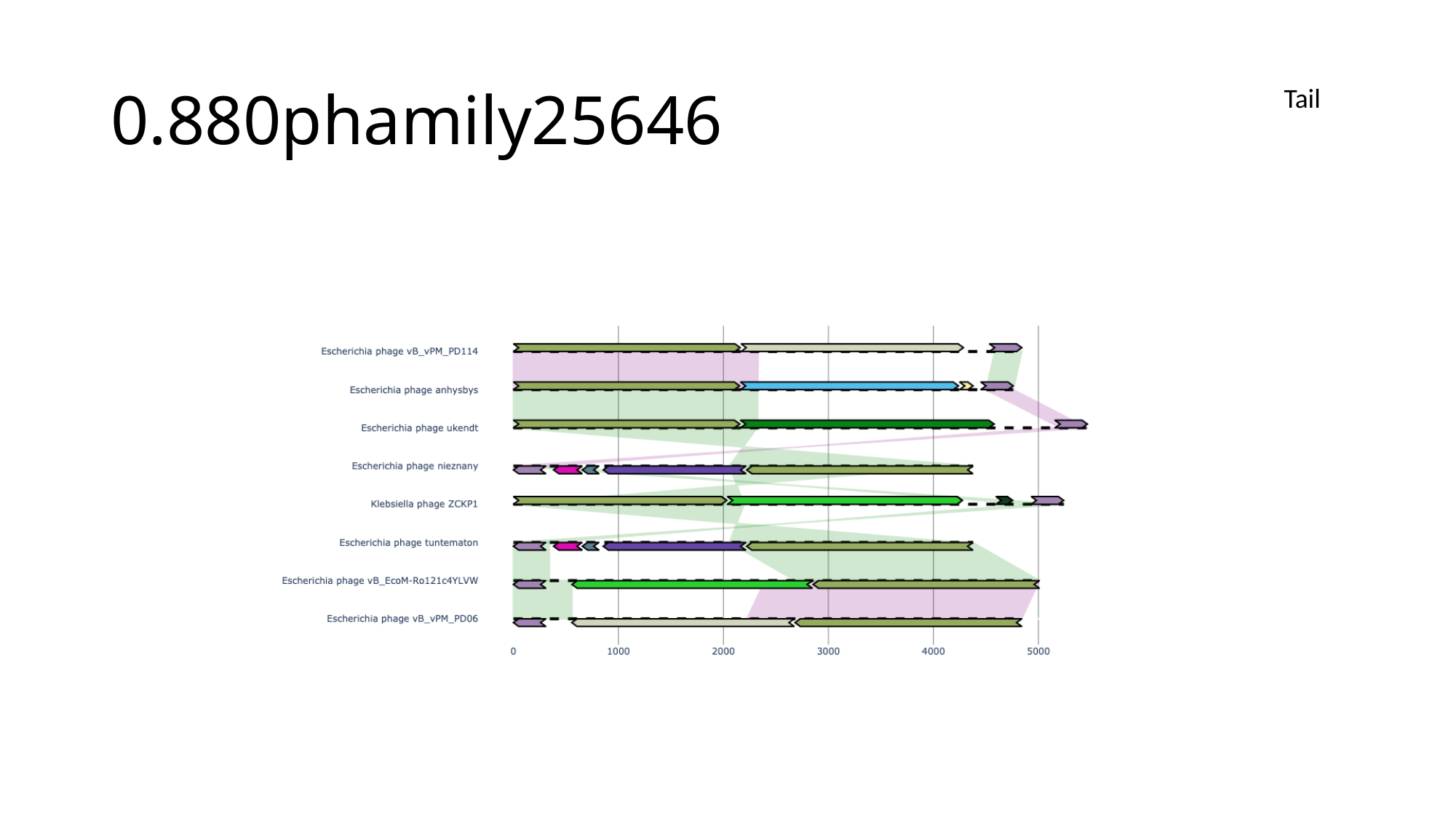

#### 0.880phamily25646
Tail

### Slide 125
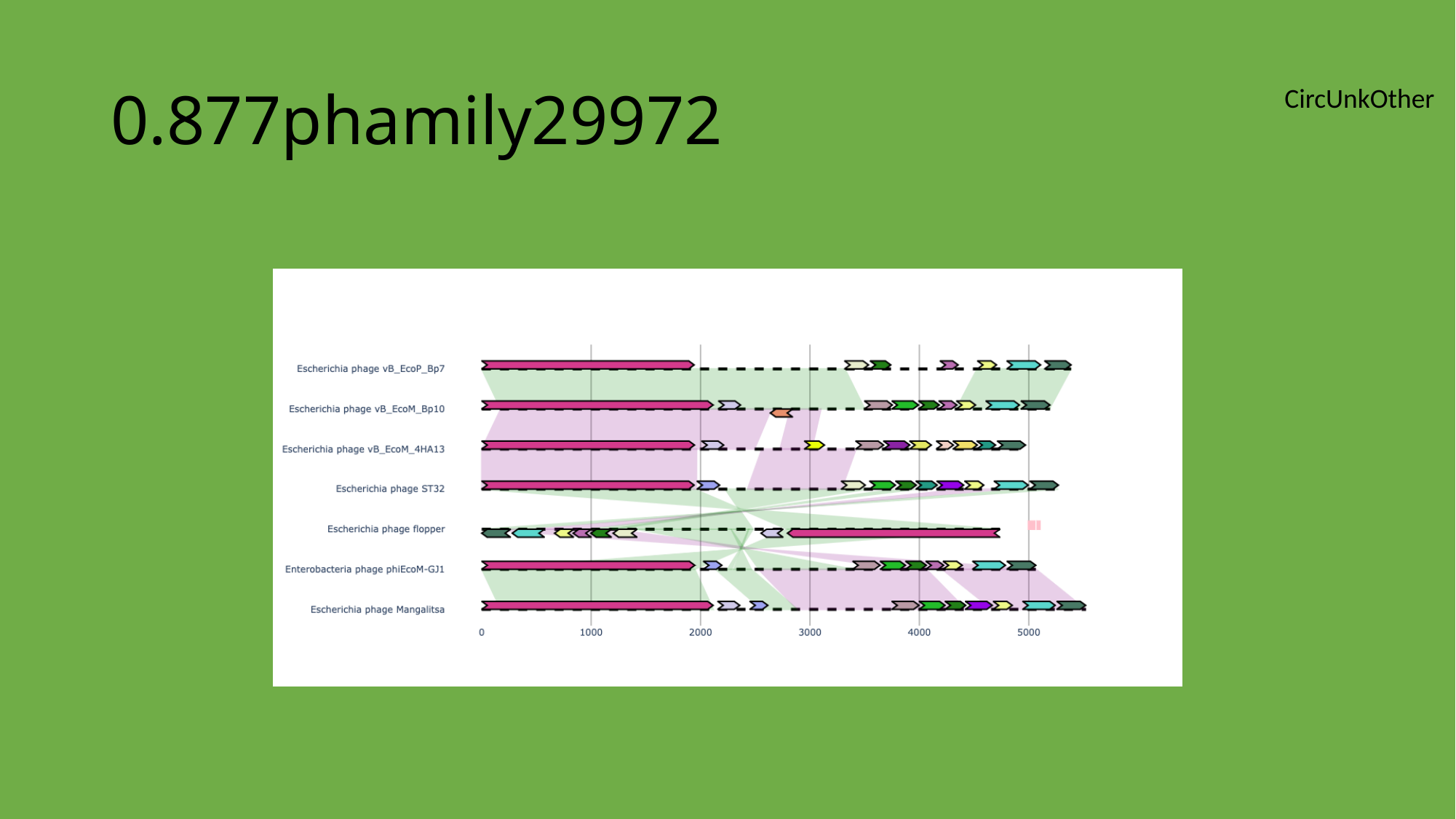

#### 0.877phamily29972
CircUnkOther

### Slide 126
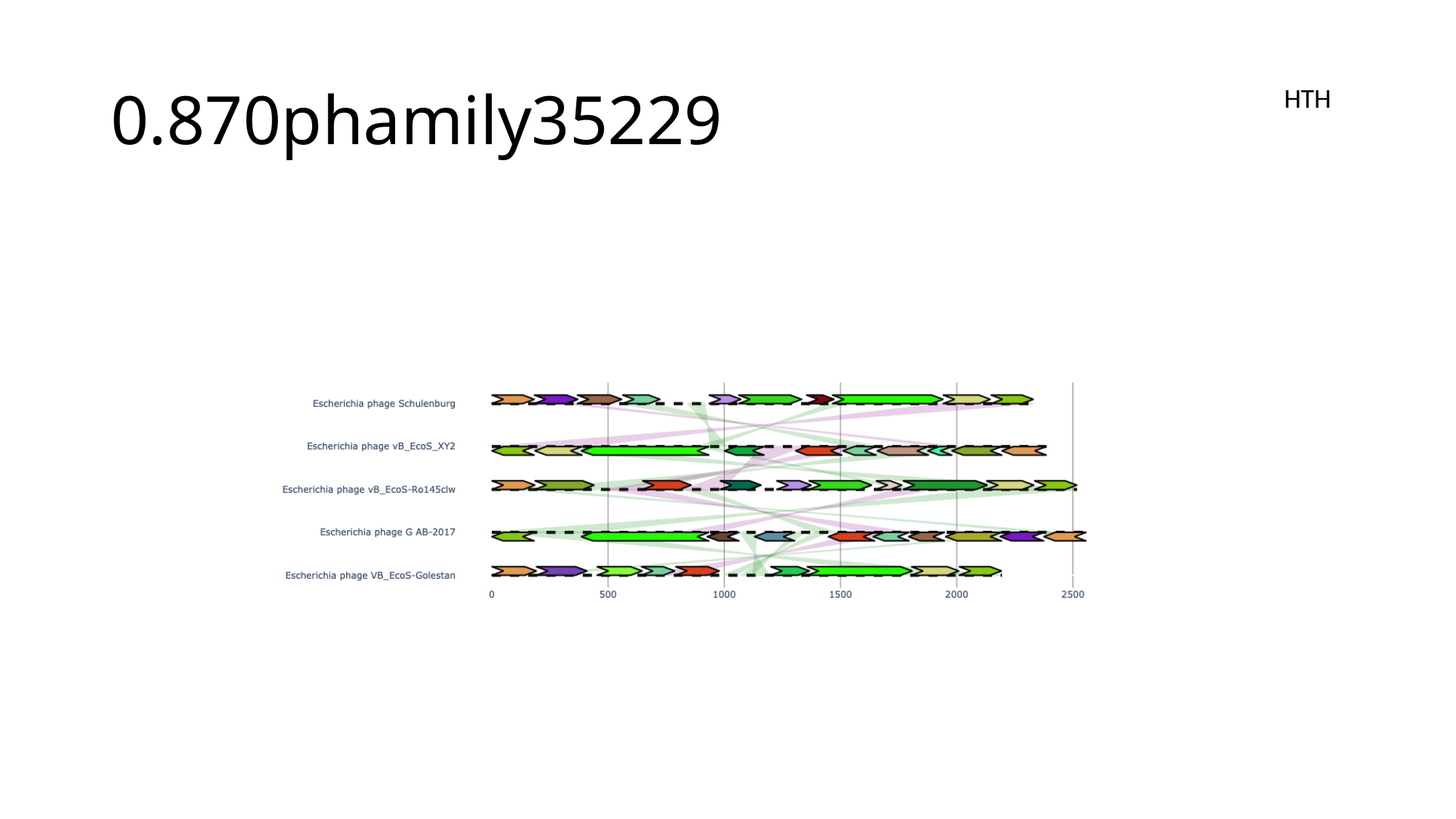

#### 0.870phamily35229
HTH

### Slide 127
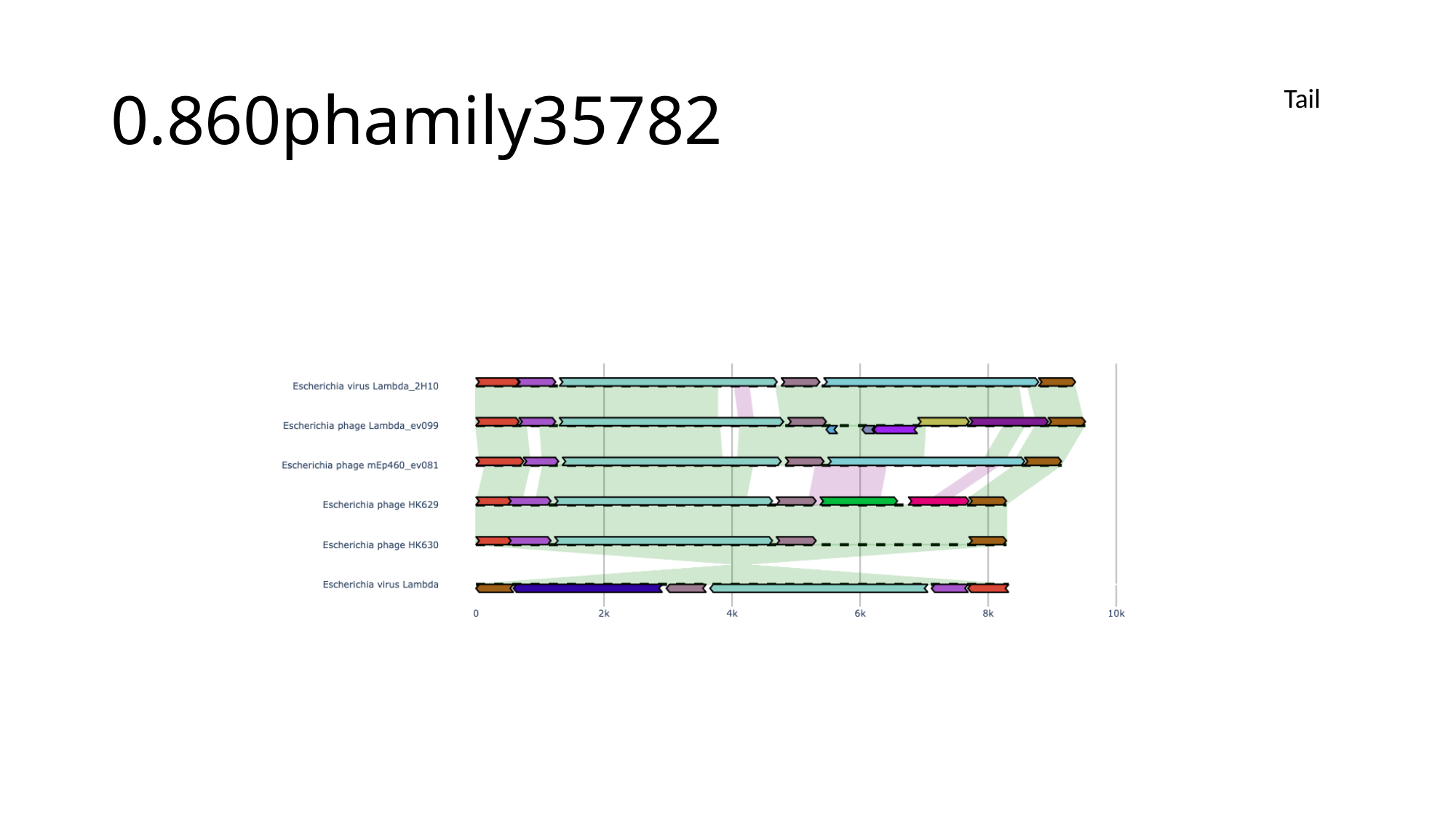

#### 0.860phamily35782
Tail

### Slide 128
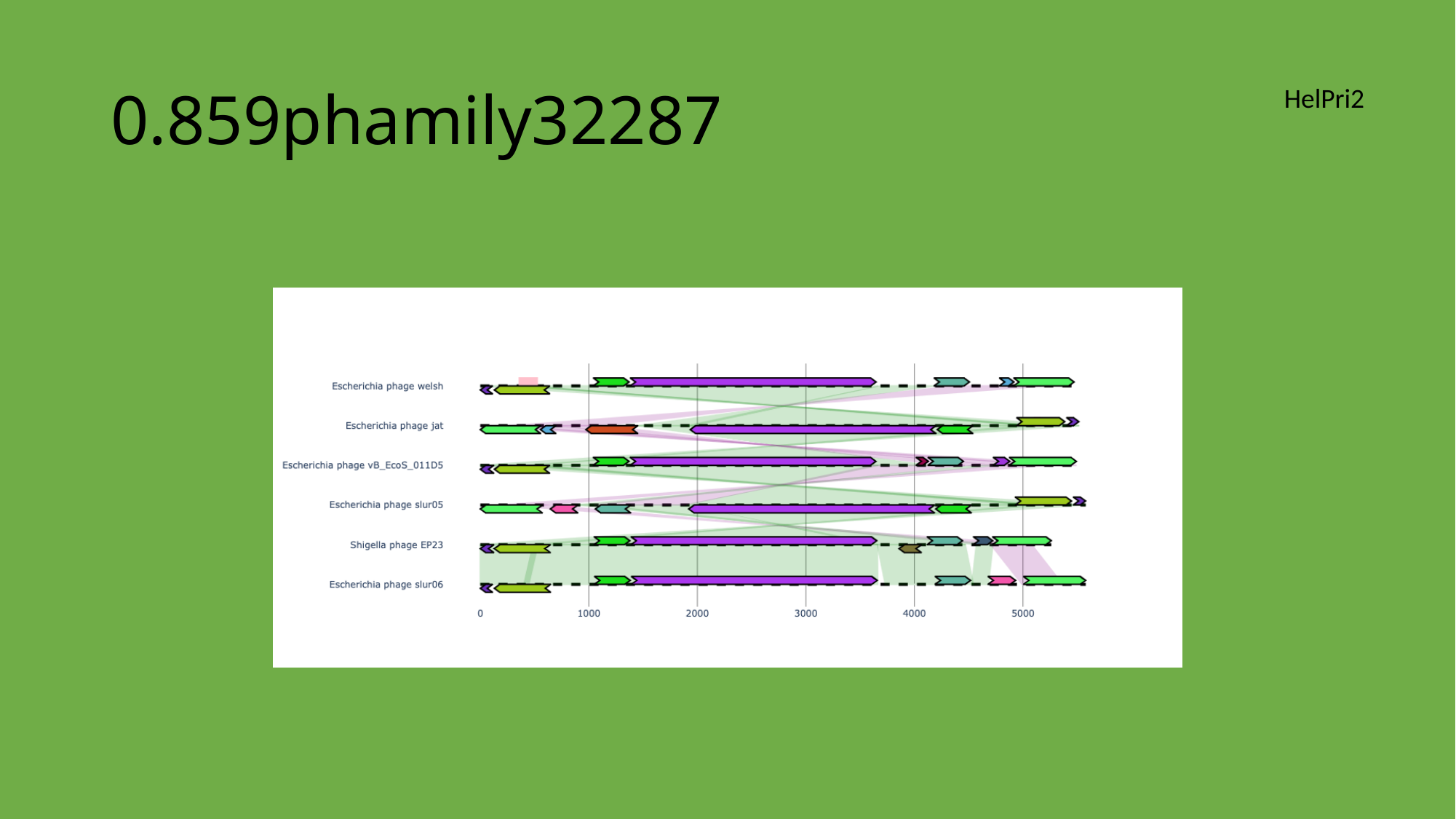

#### 0.859phamily32287
HelPri2

### Slide 129
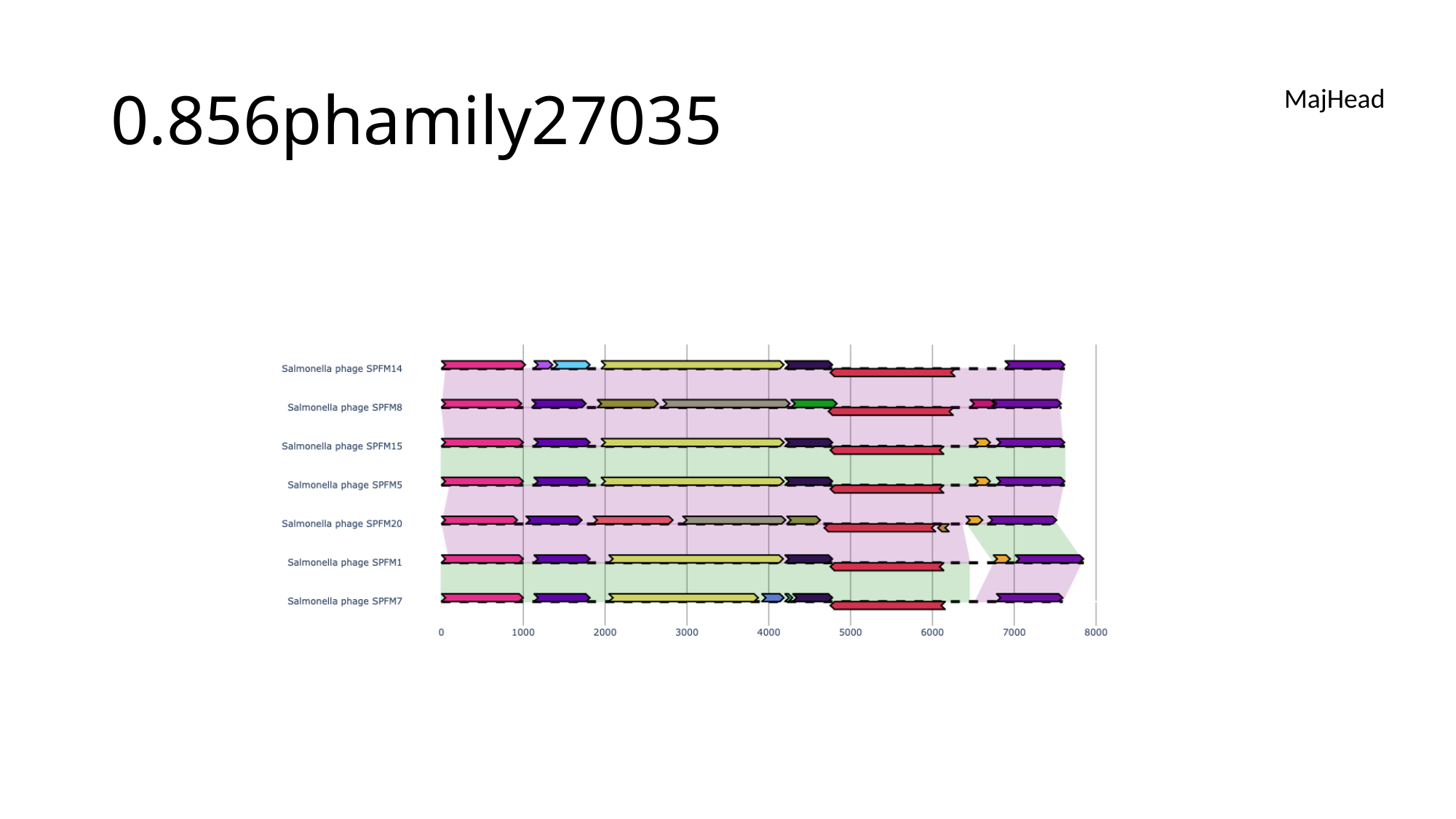

#### 0.856phamily27035
MajHead

### Slide 130
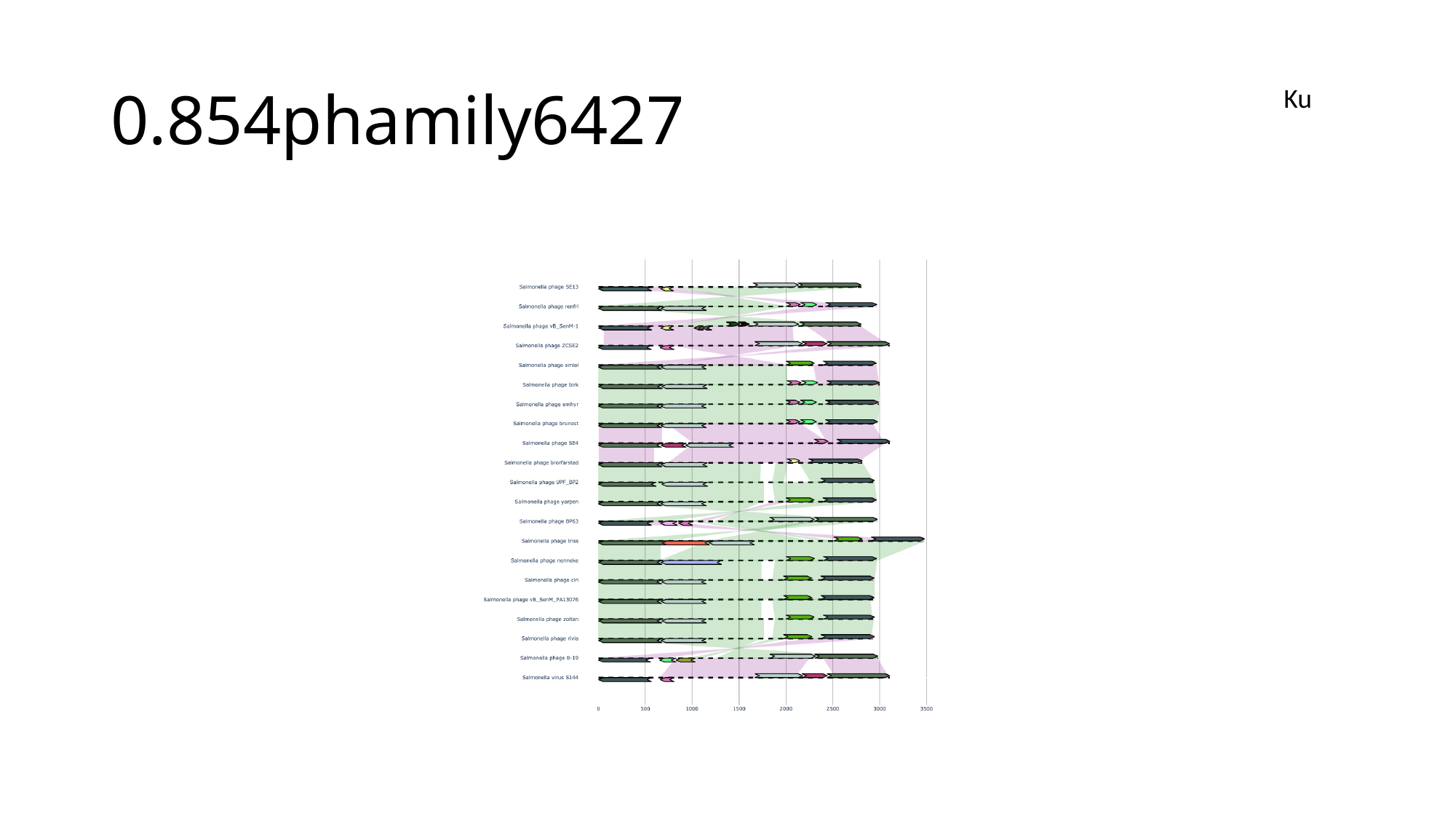

#### 0.854phamily6427
Ku

### Slide 131
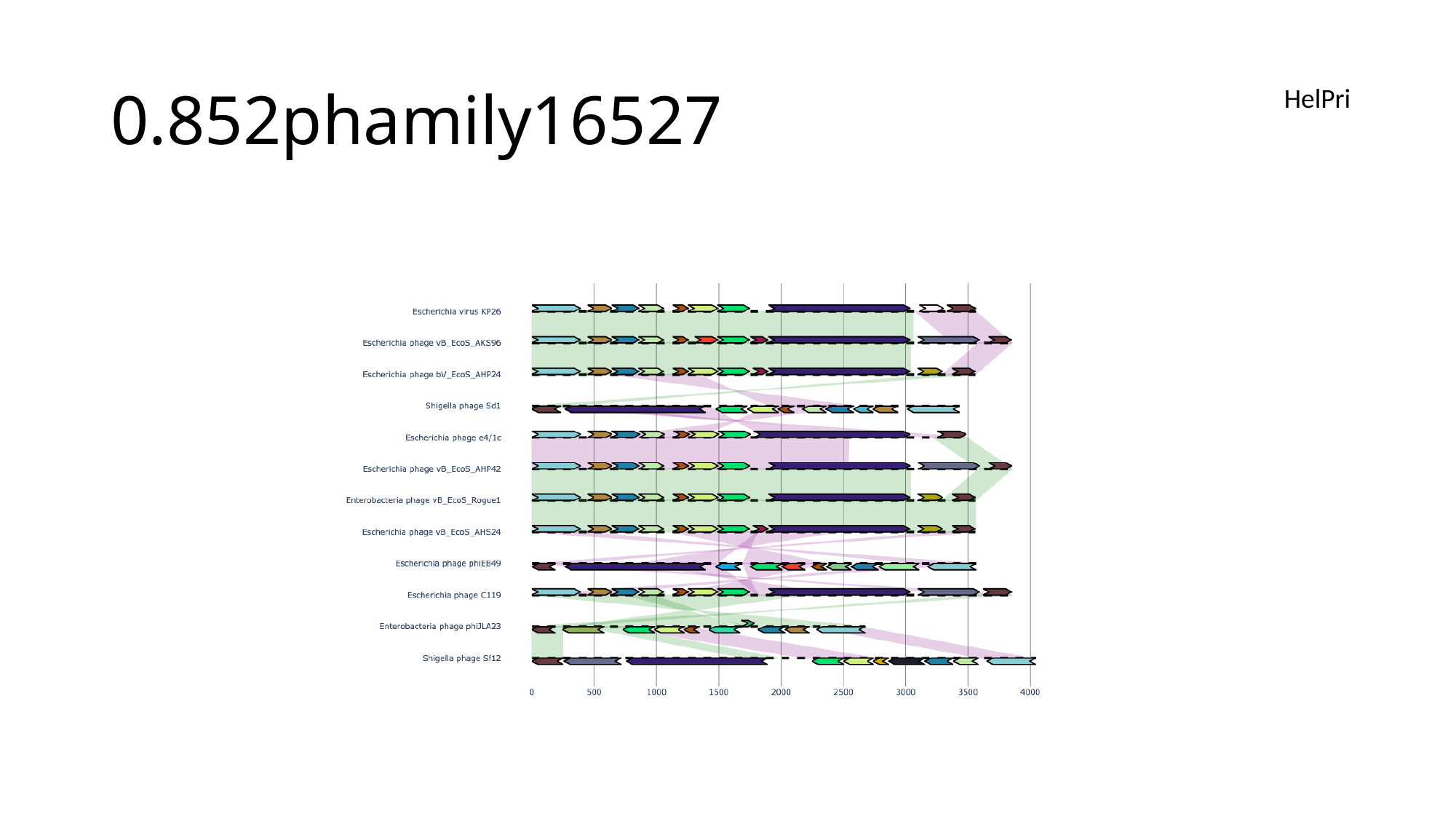

#### 0.852phamily16527
HelPri

### Slide 132
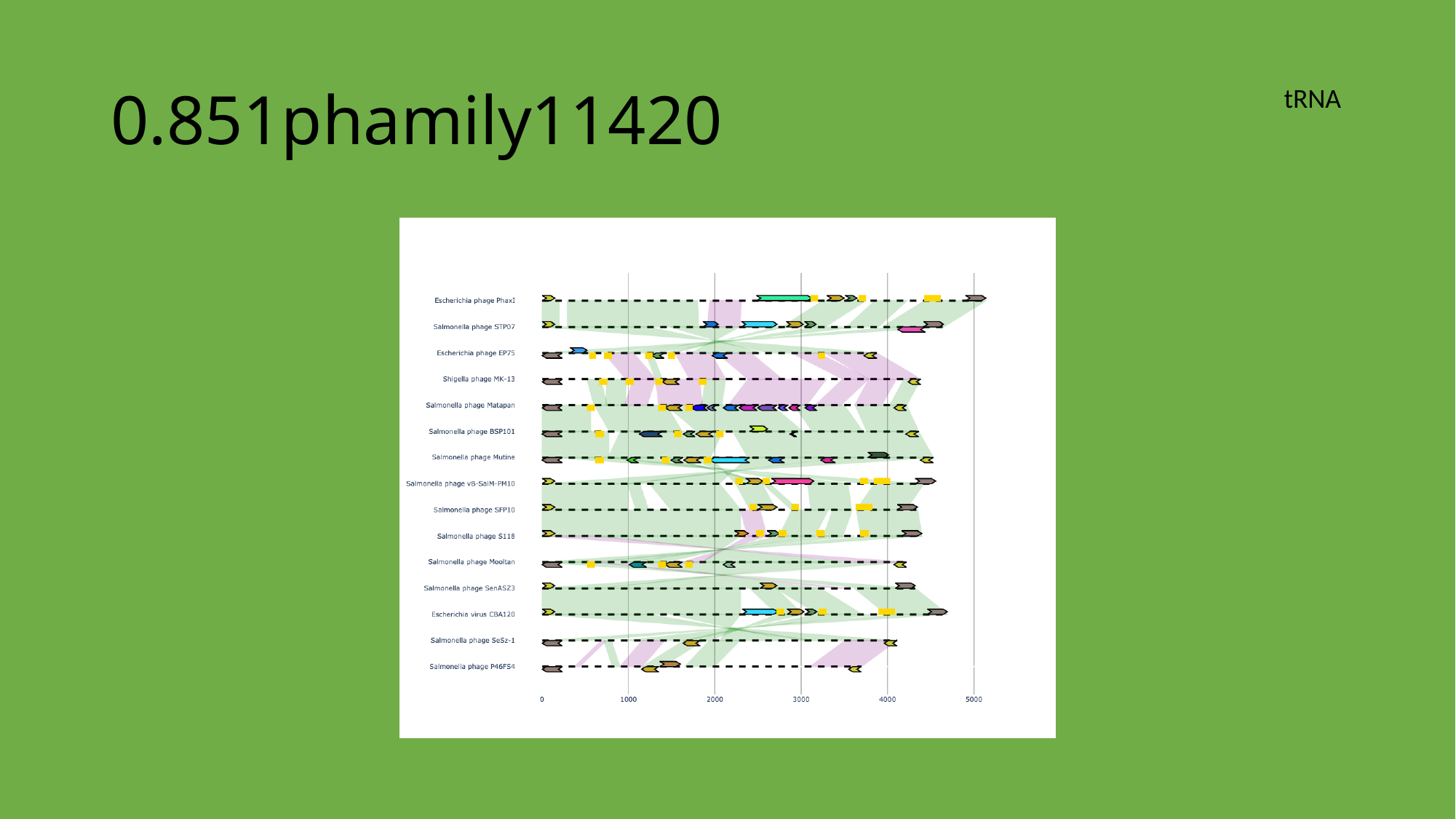

#### 0.851phamily11420
tRNA

### Slide 133
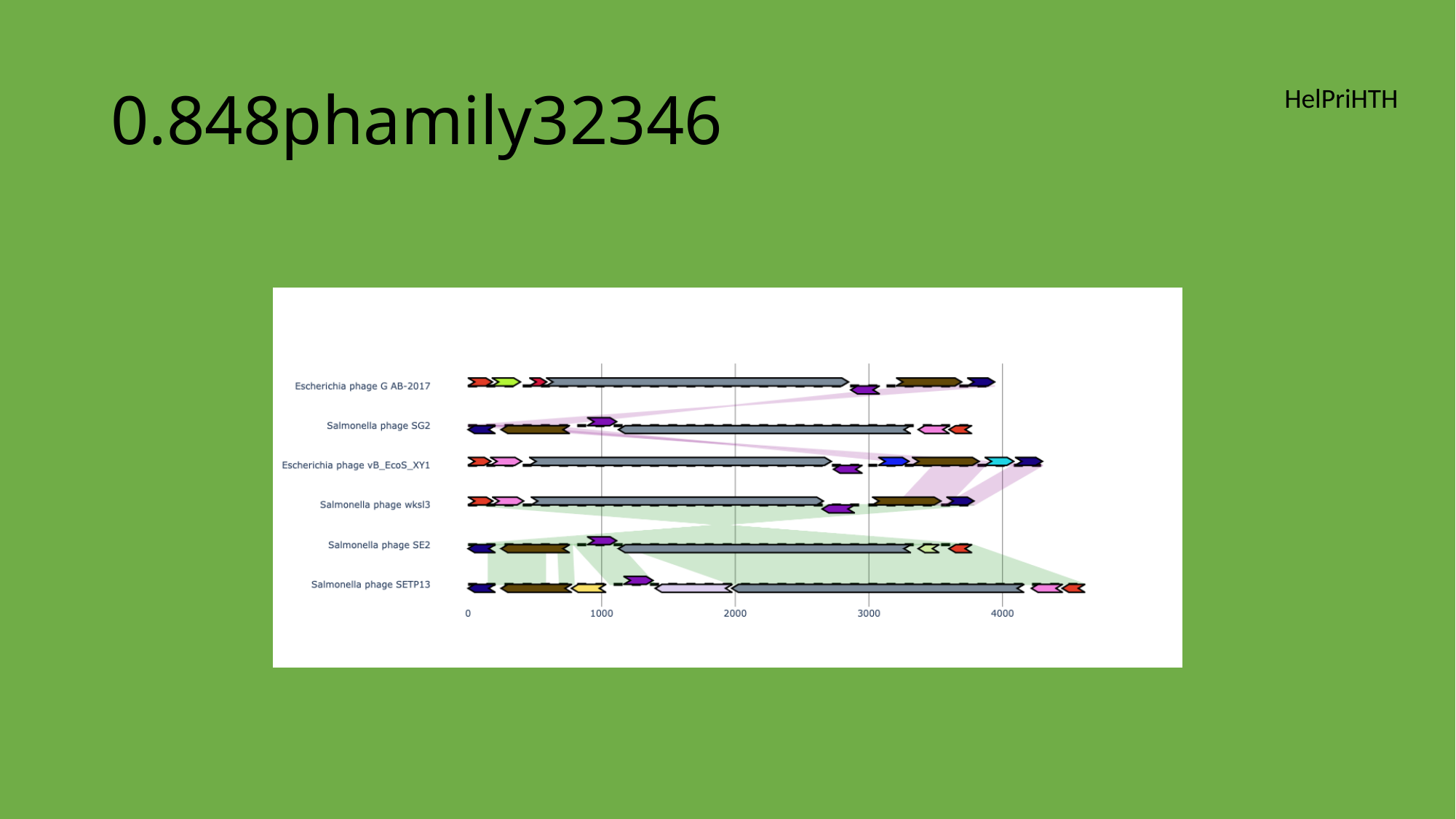

#### 0.848phamily32346
HelPriHTH

### Slide 134
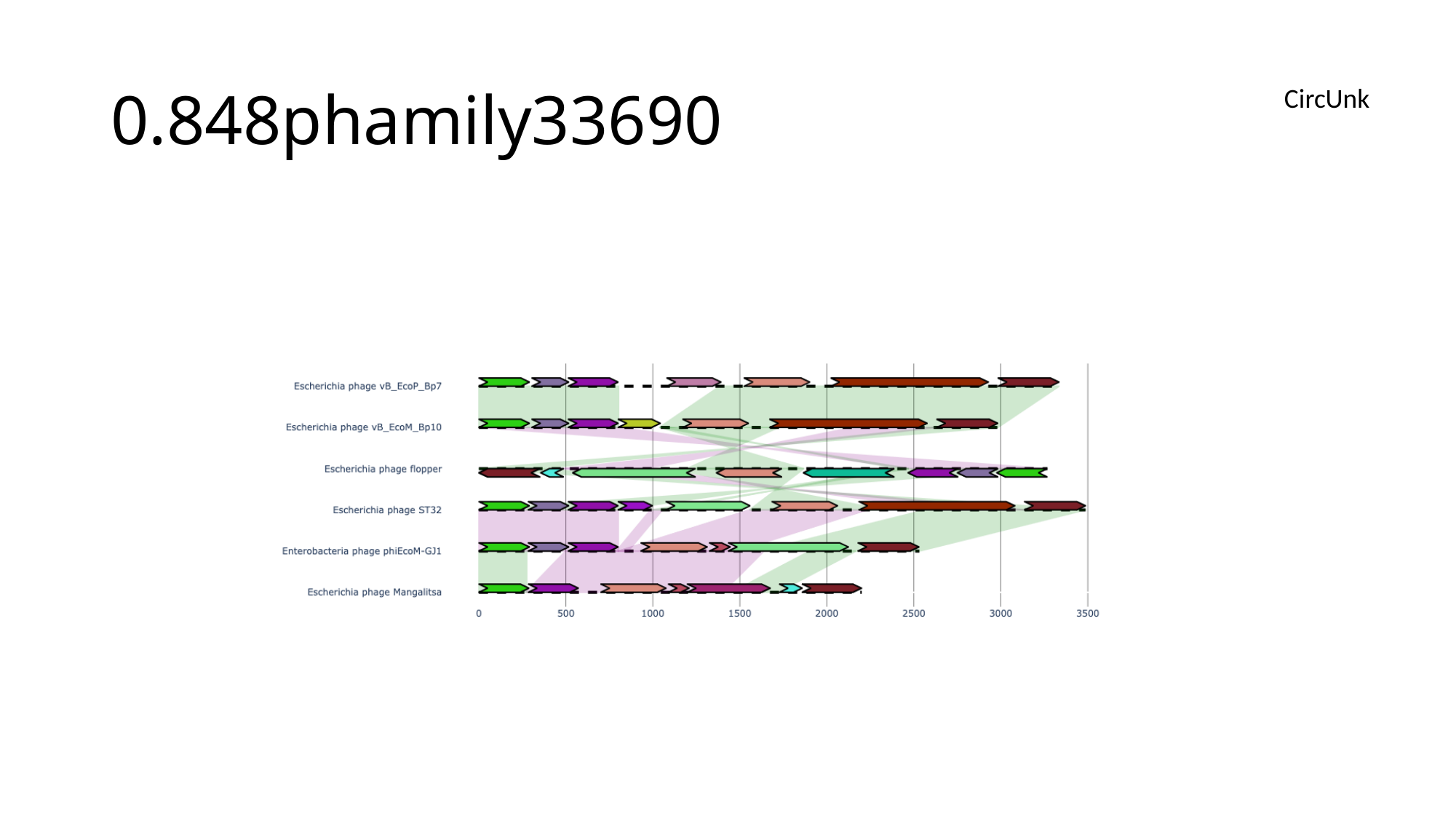

#### 0.848phamily33690
CircUnk

### Slide 135
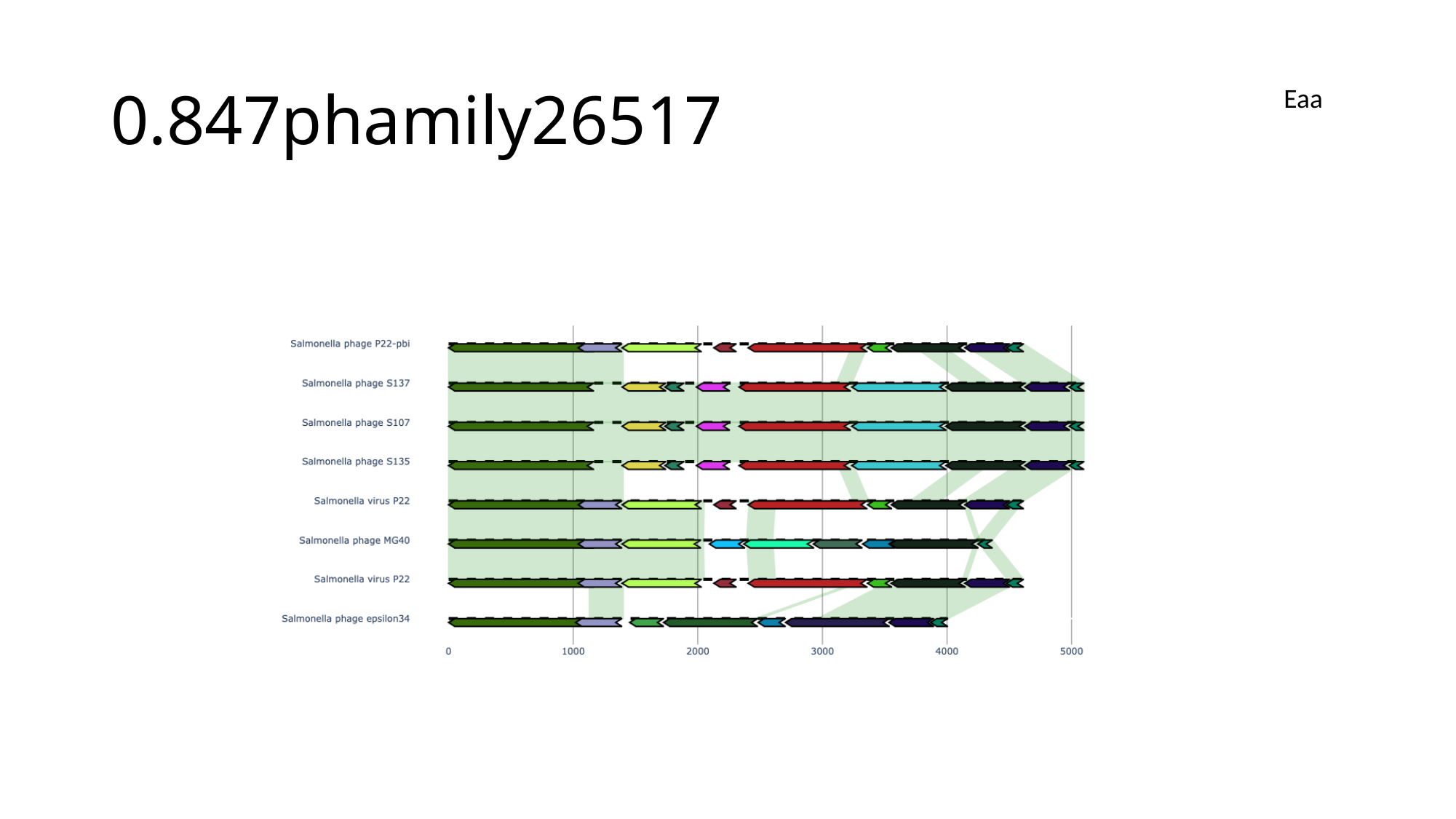

#### 0.847phamily26517
Eaa

### Slide 136
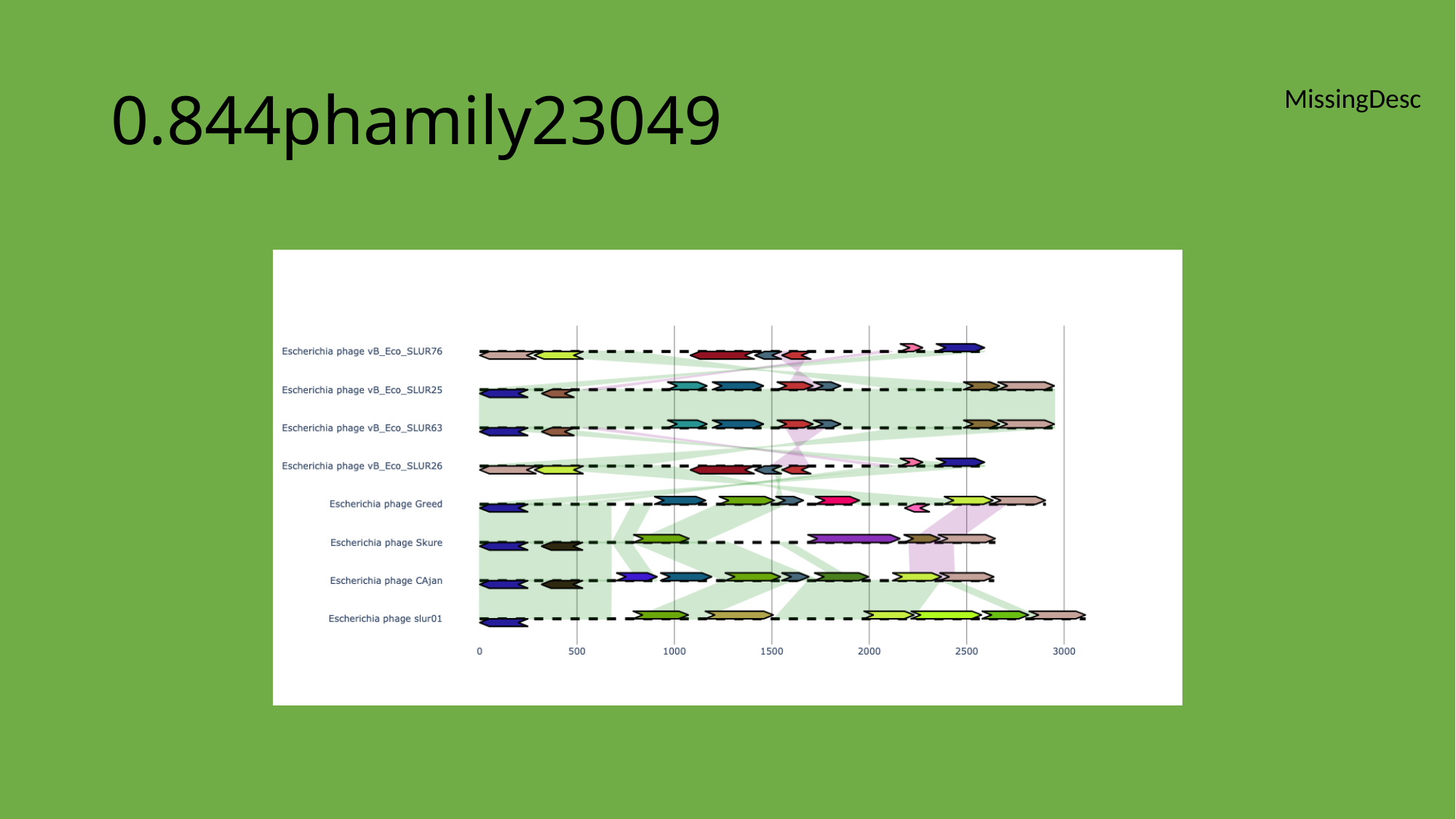

#### 0.844phamily23049
MissingDesc

### Slide 137
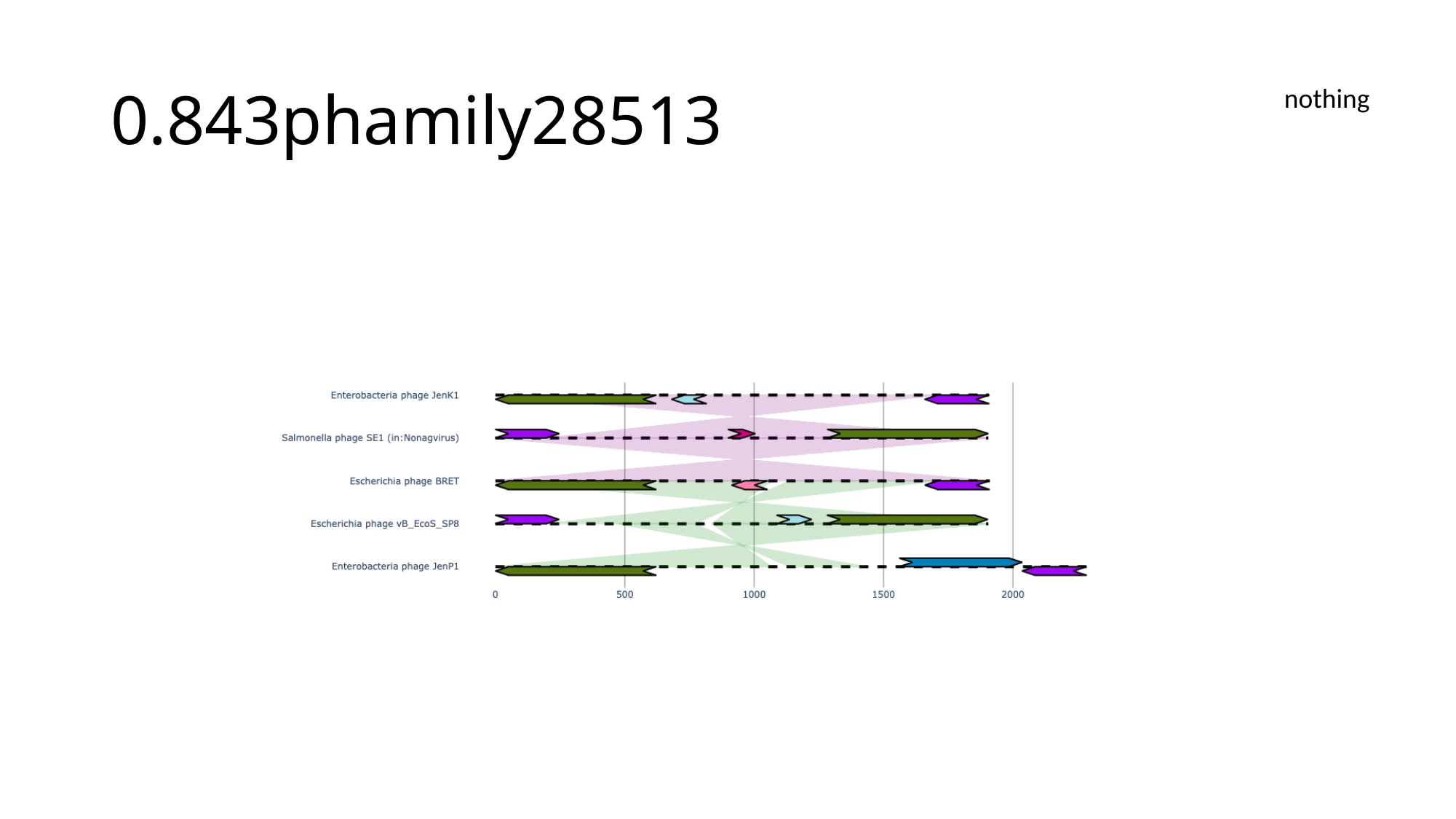

#### 0.843phamily28513
nothing

### Slide 138
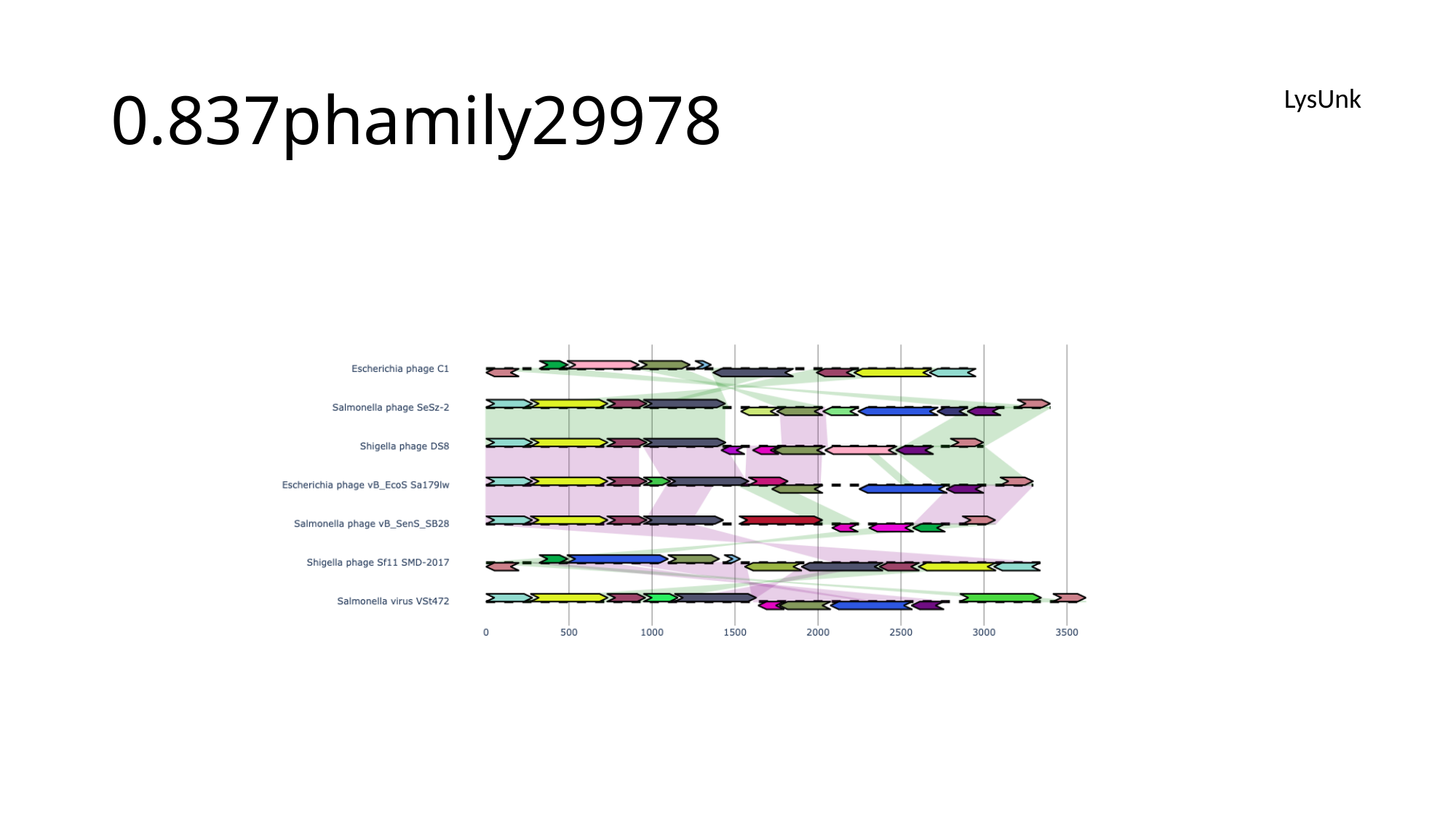

#### 0.837phamily29978
LysUnk

### Slide 139
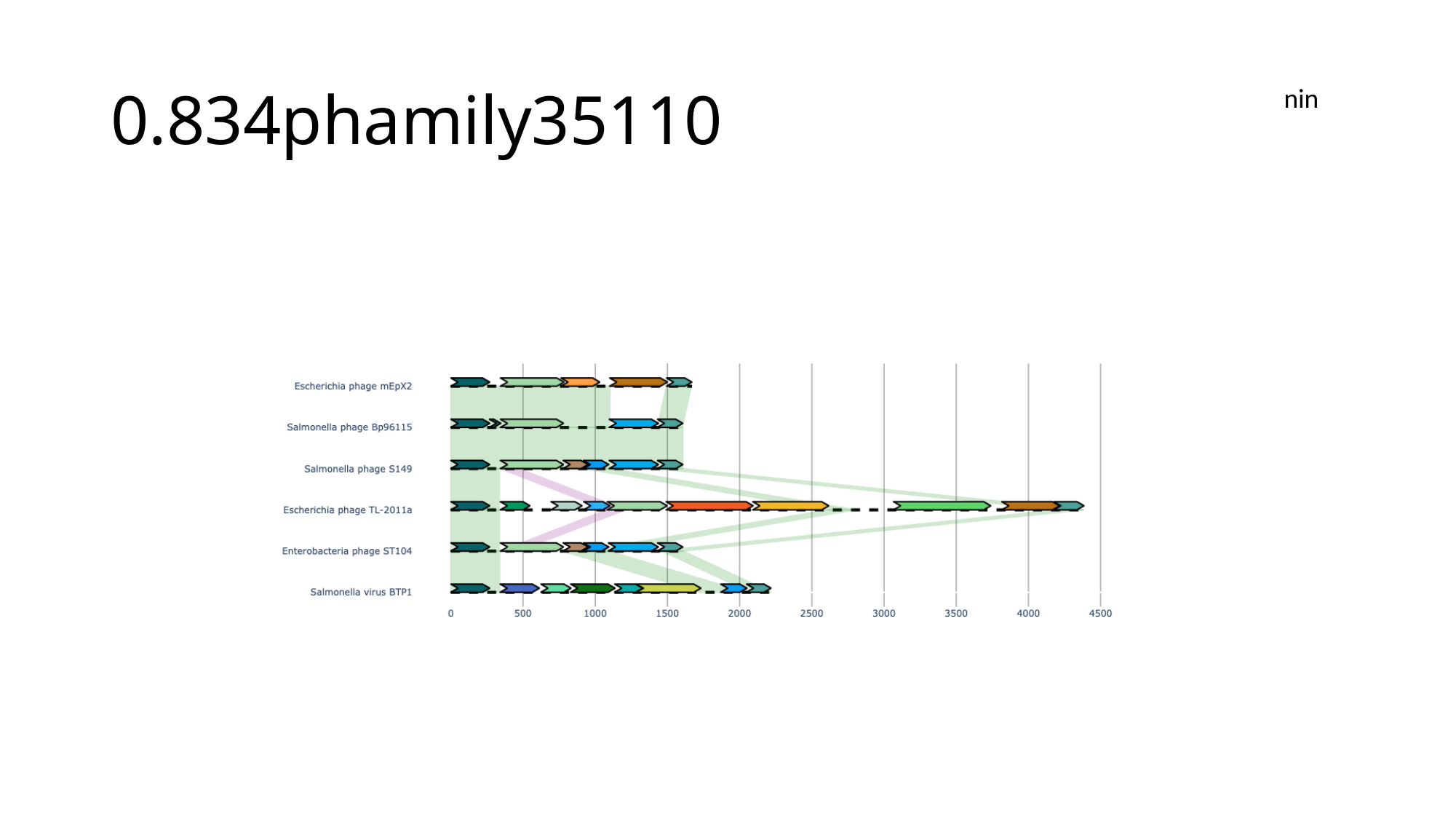

#### 0.834phamily35110
nin

### Slide 140
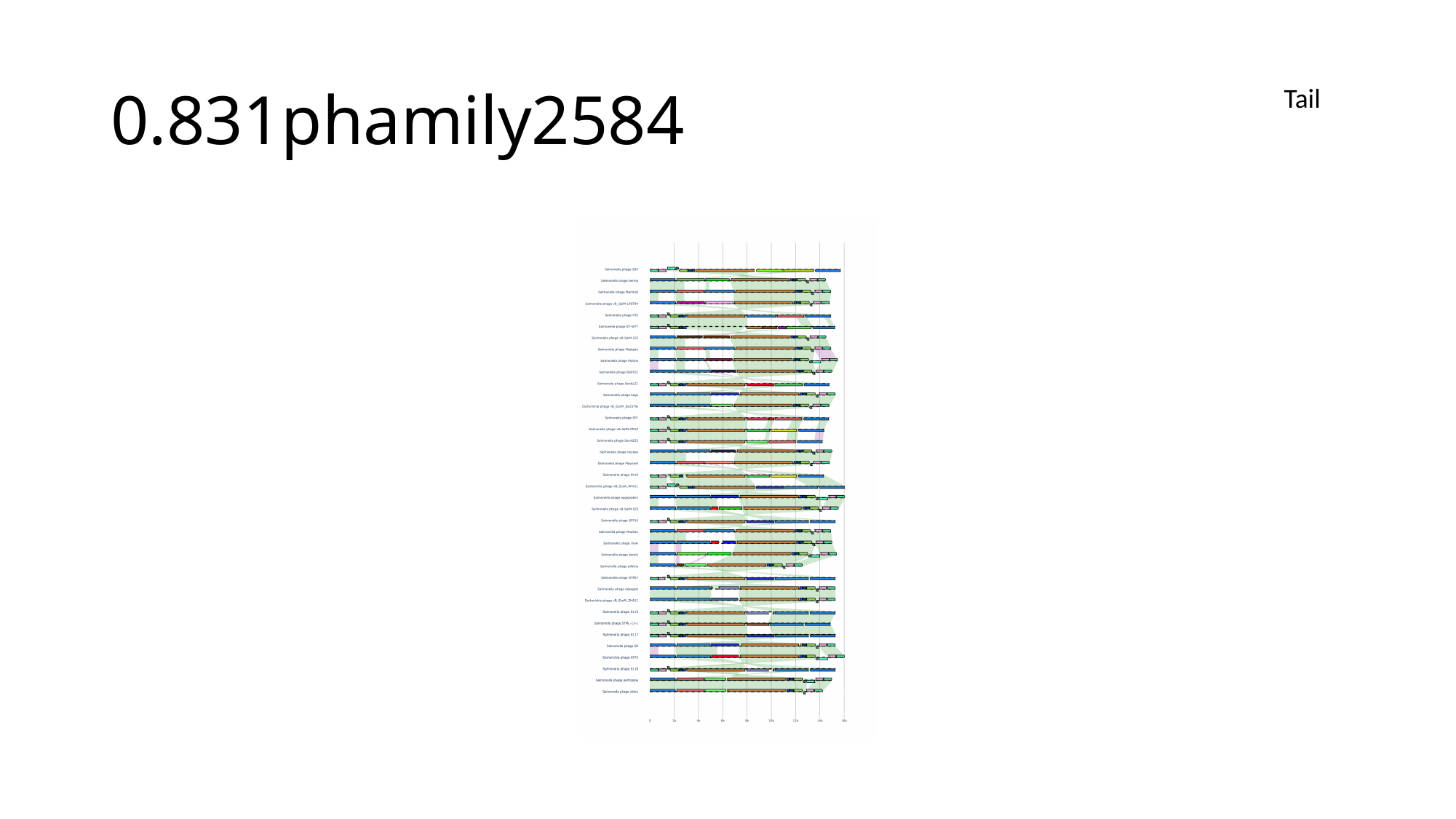

#### 0.831phamily2584
Tail

### Slide 141
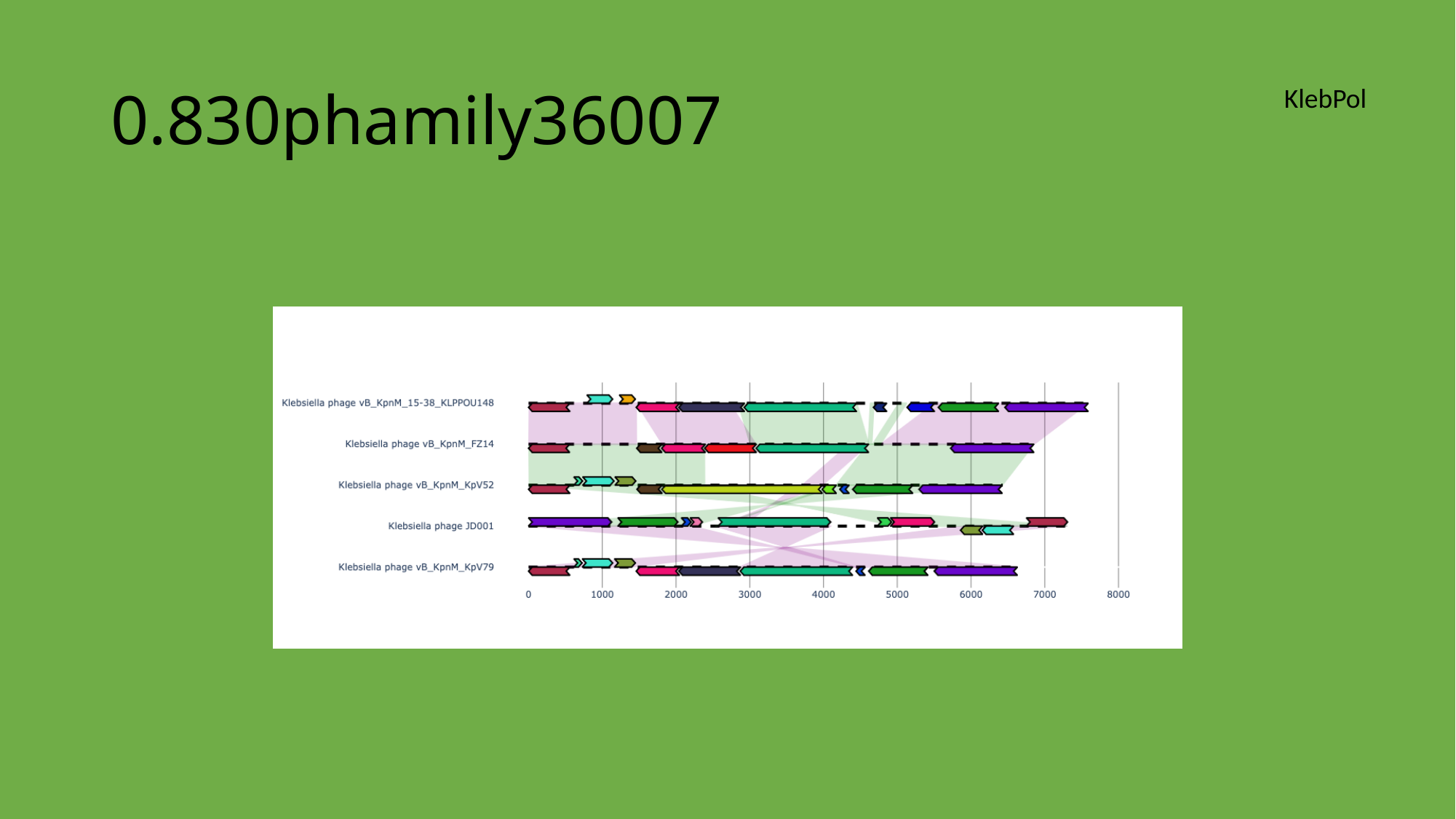

#### 0.830phamily36007
KlebPol

### Slide 142
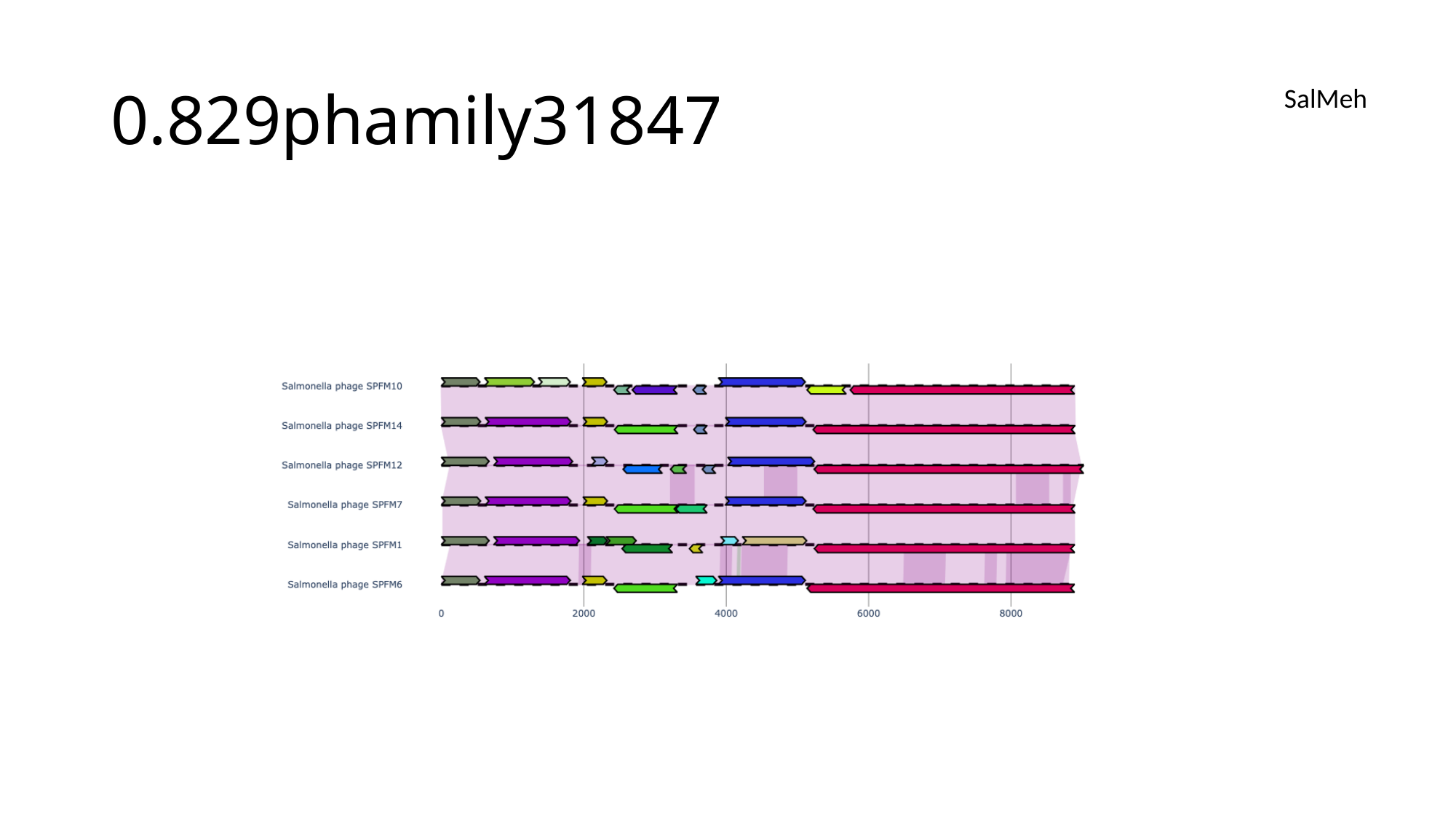

#### 0.829phamily31847
SalMeh

### Slide 143
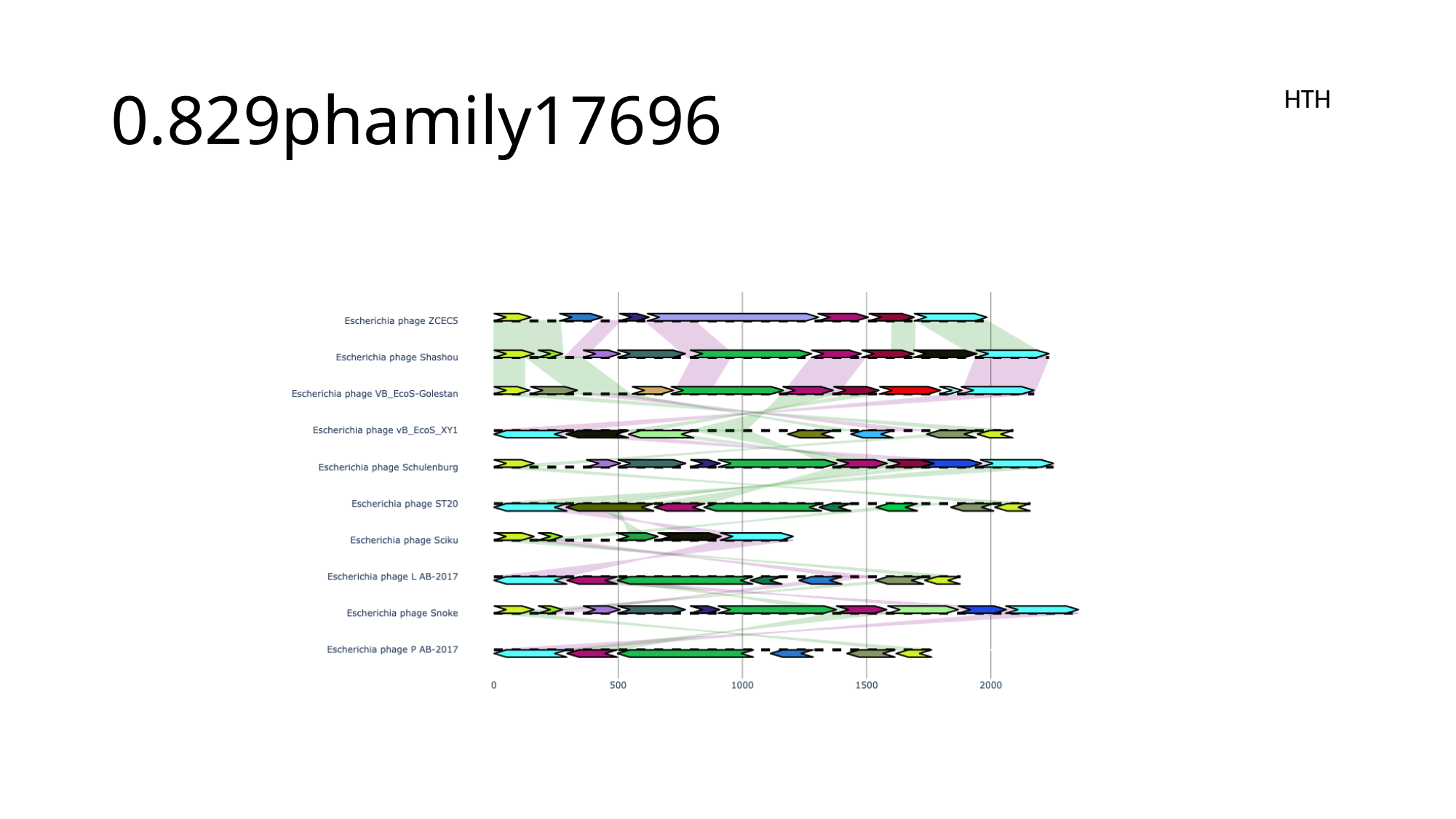

#### 0.829phamily17696
HTH

### Slide 144
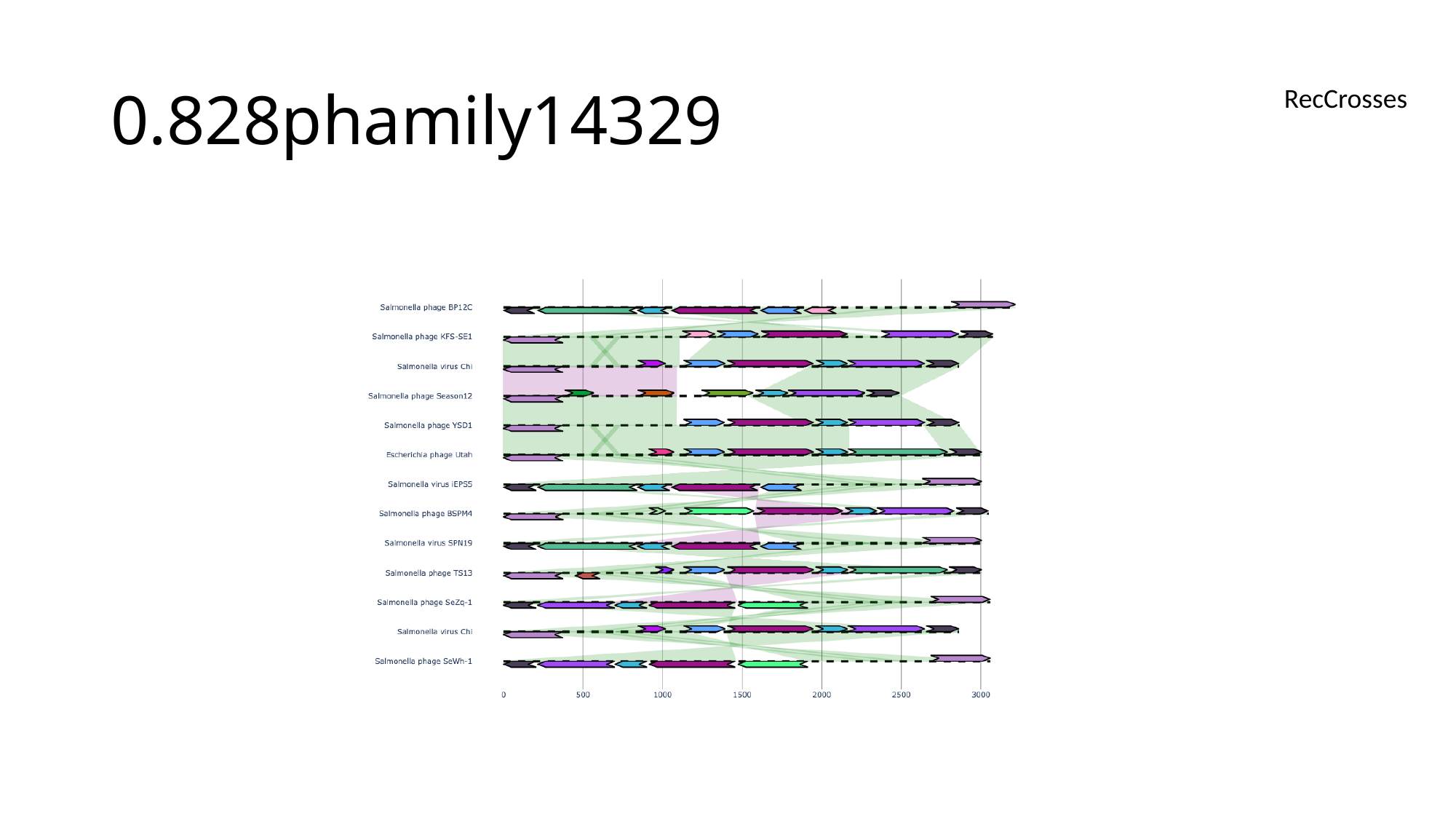

#### 0.828phamily14329
RecCrosses

### Slide 145
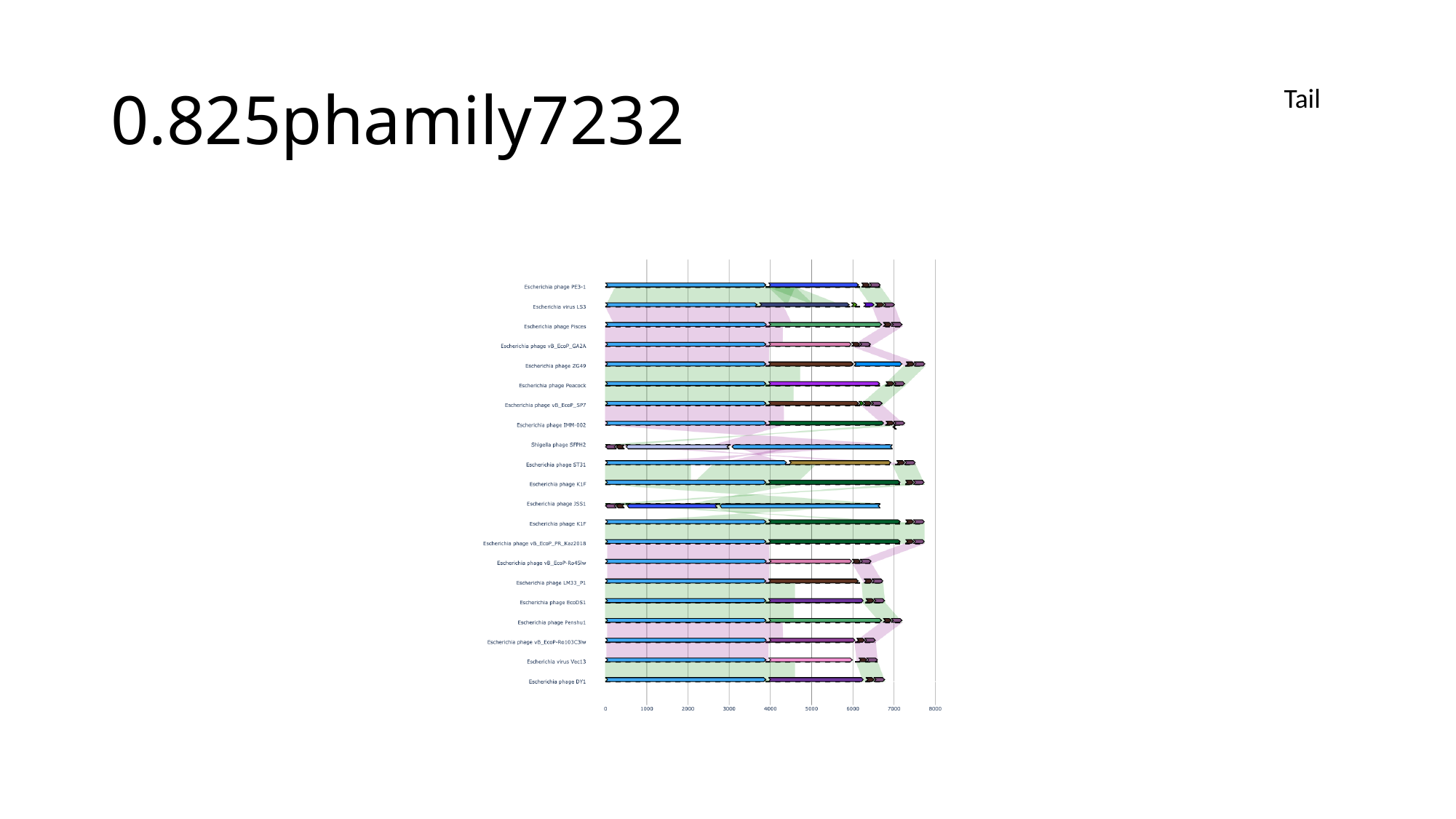

#### 0.825phamily7232
Tail

### Slide 146
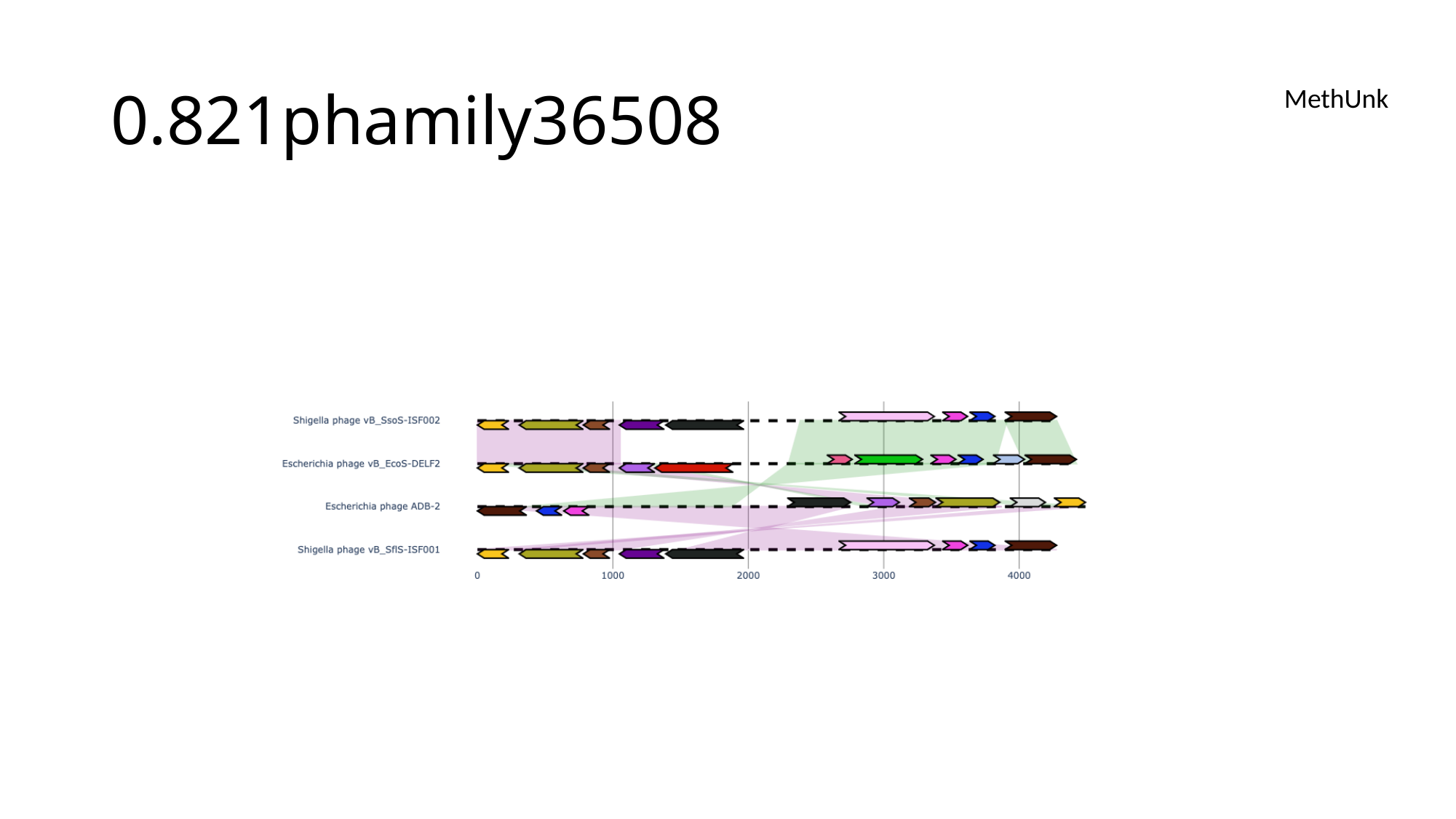

#### 0.821phamily36508
MethUnk

### Slide 147
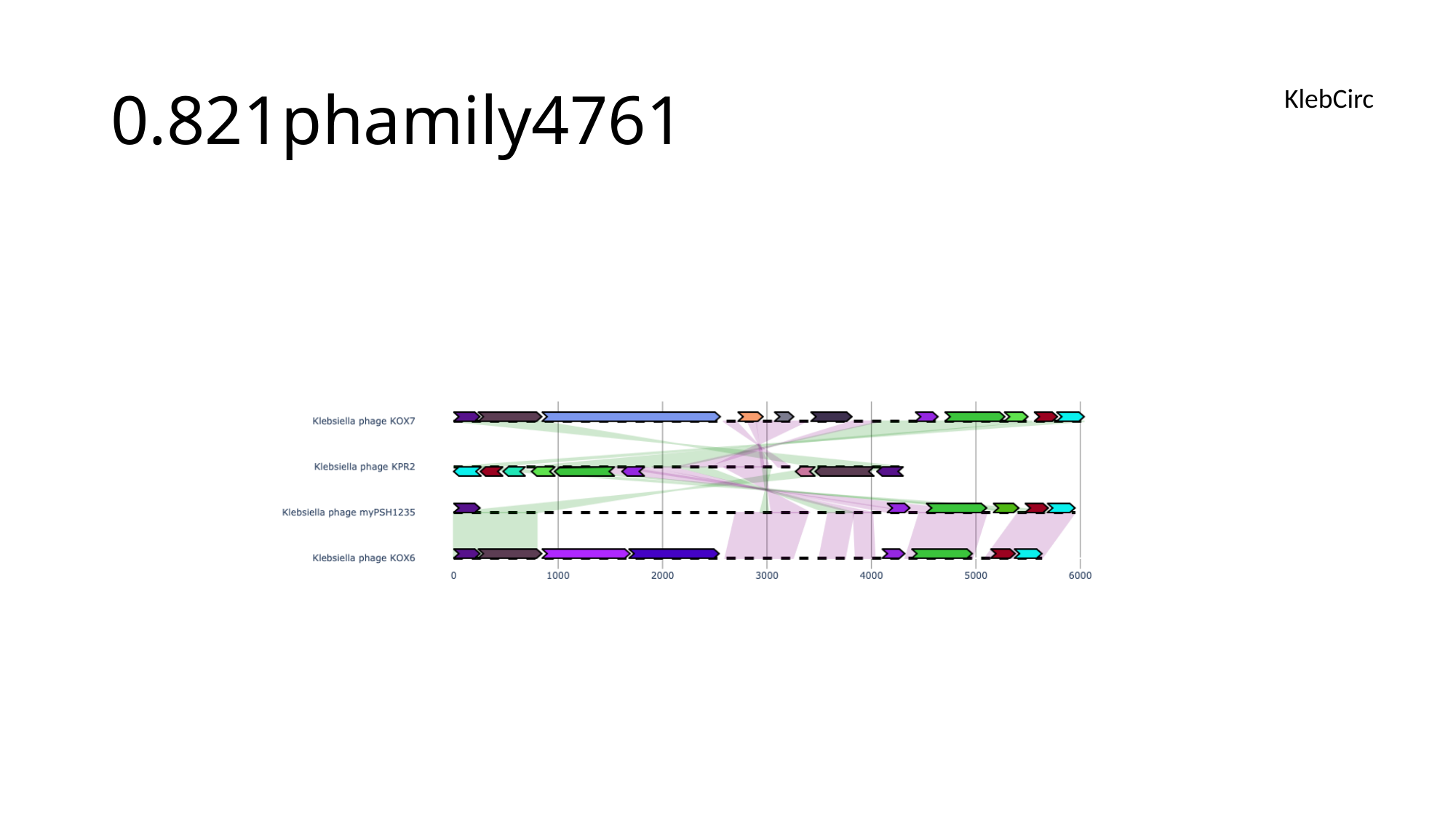

#### 0.821phamily4761
KlebCirc

### Slide 148
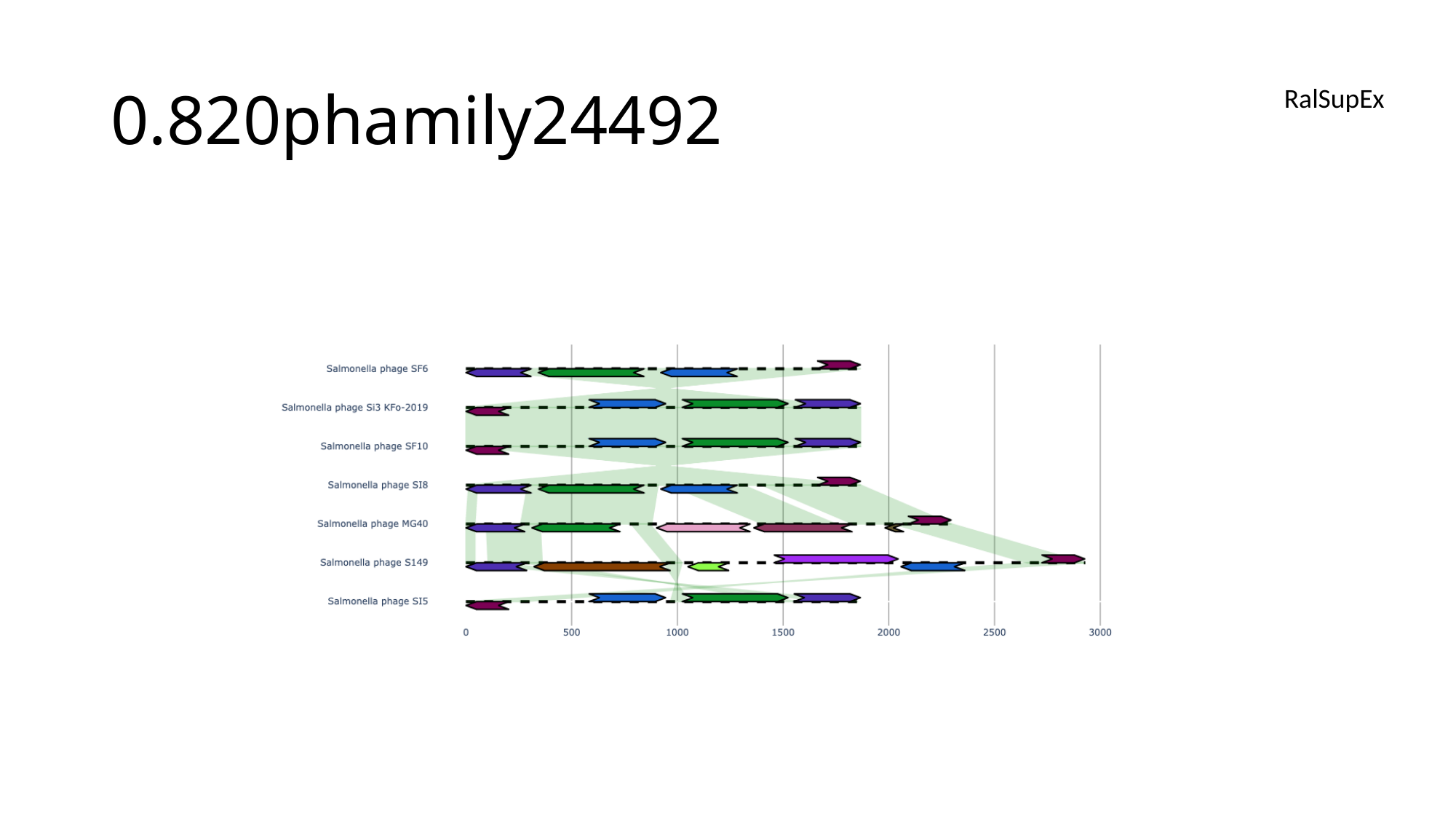

#### 0.820phamily24492
RalSupEx

### Slide 149
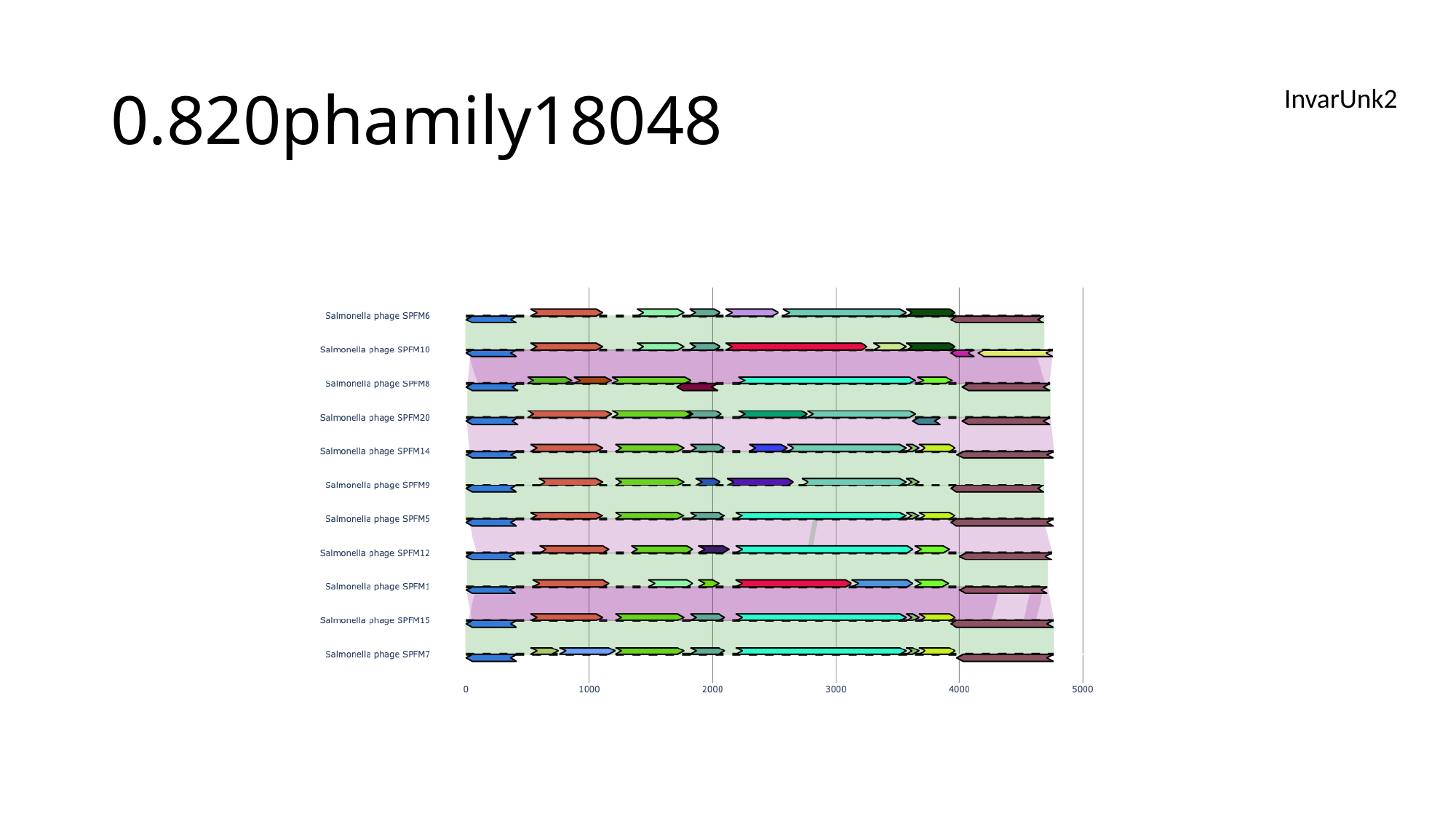

#### 0.820phamily18048
InvarUnk2

### Slide 150
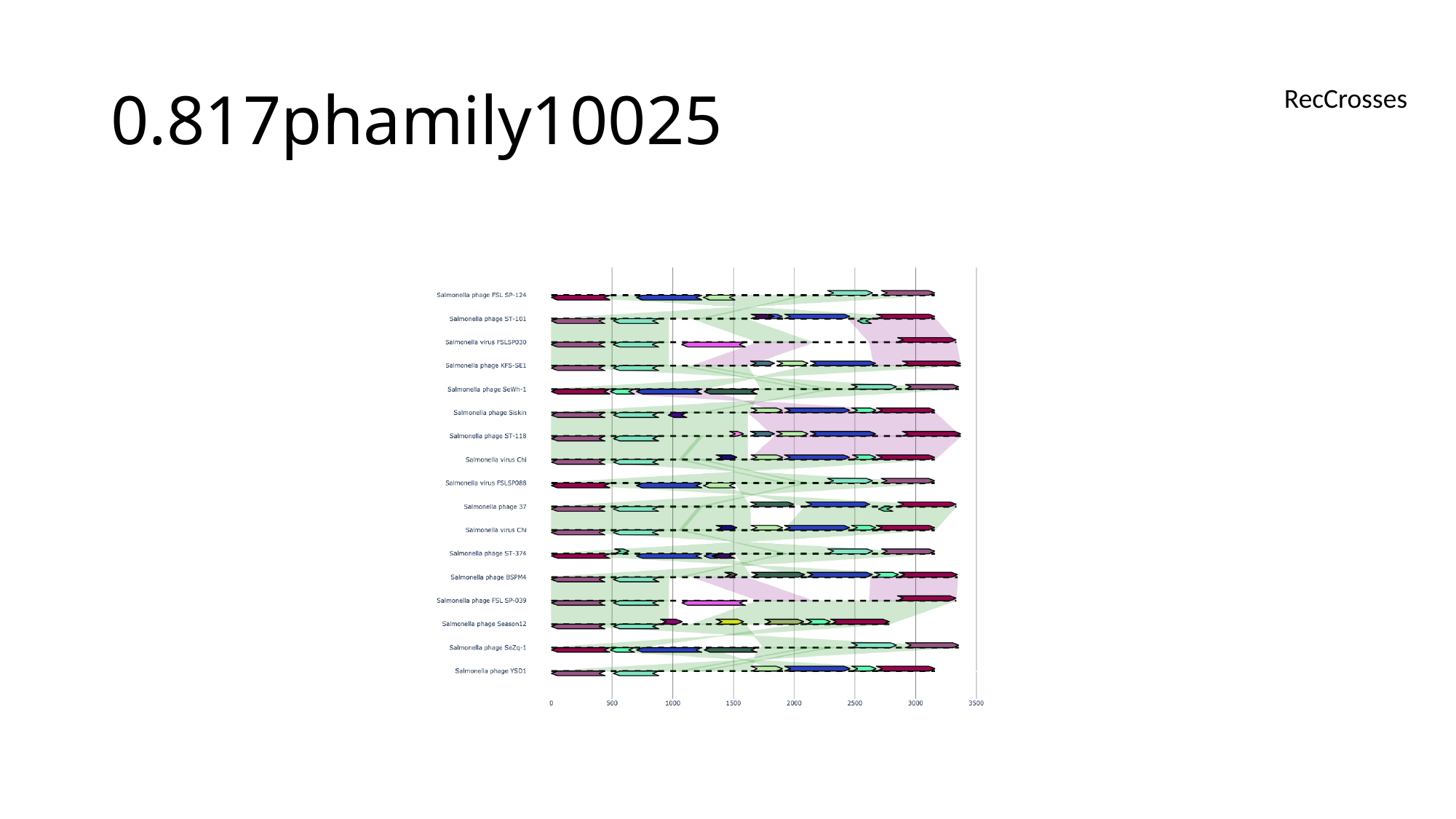

#### 0.817phamily10025
RecCrosses

### Slide 151
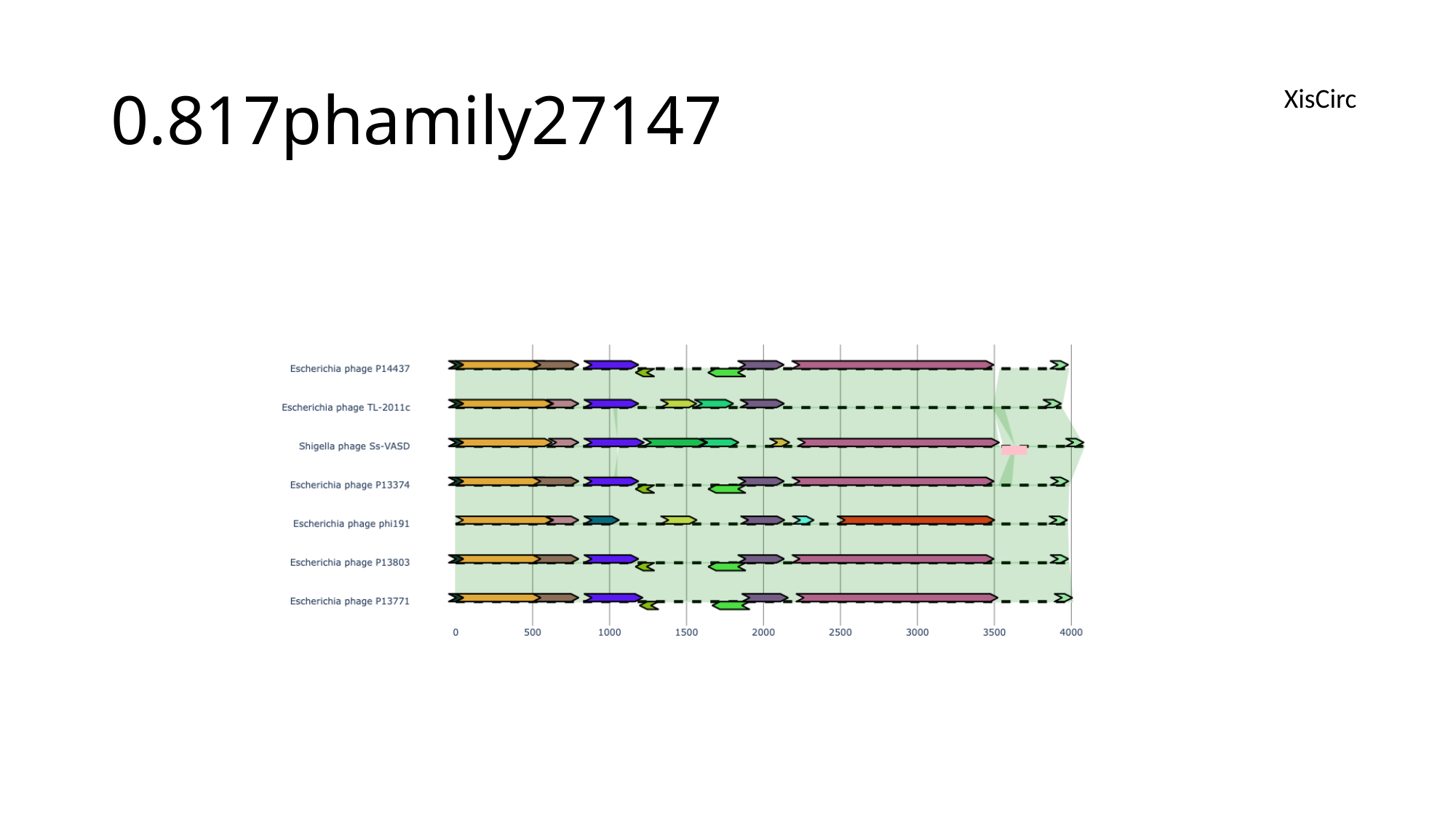

#### 0.817phamily27147
XisCirc

### Slide 152
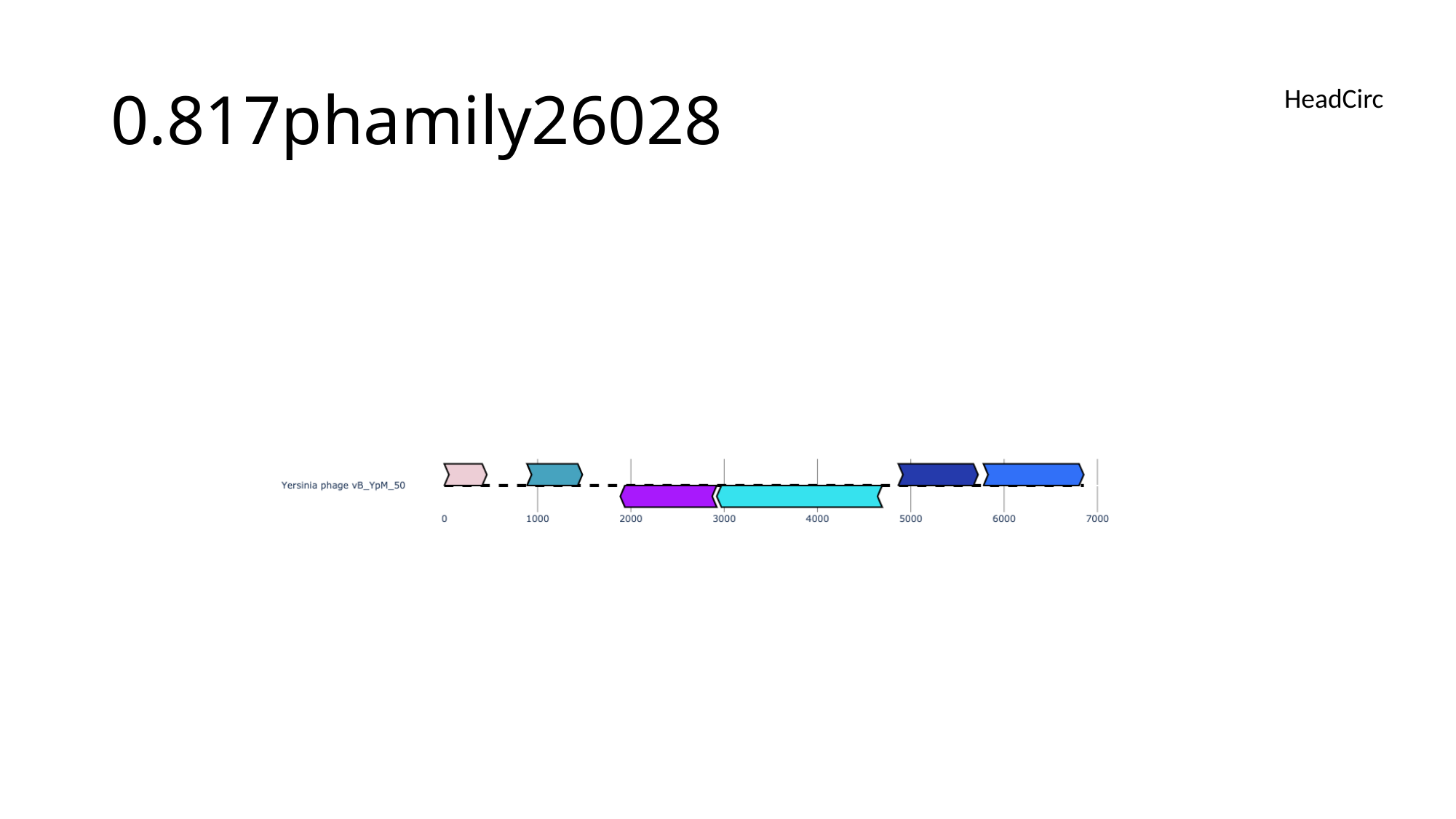

#### 0.817phamily26028
HeadCirc

### Slide 153
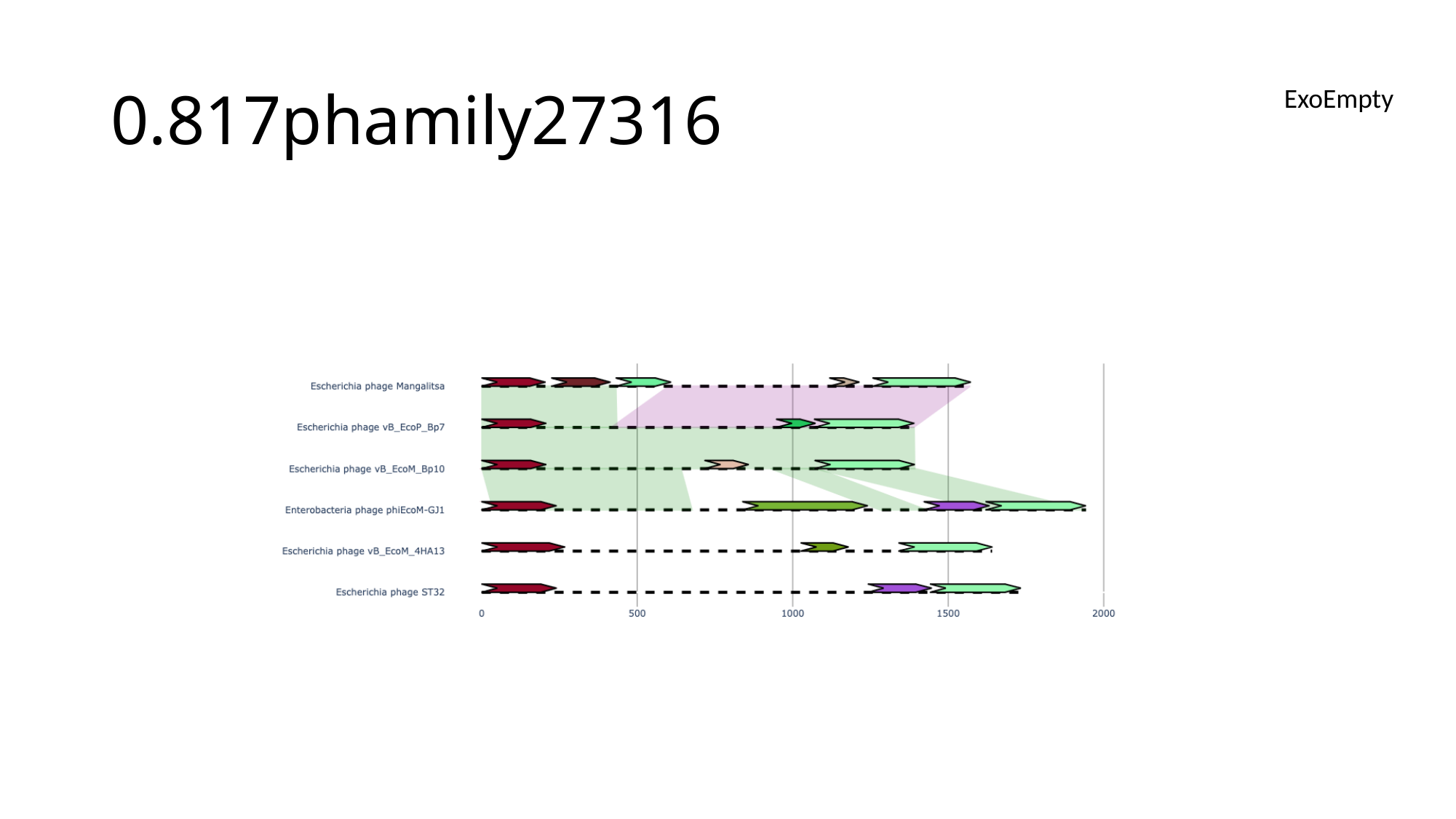

#### 0.817phamily27316
ExoEmpty

### Slide 154
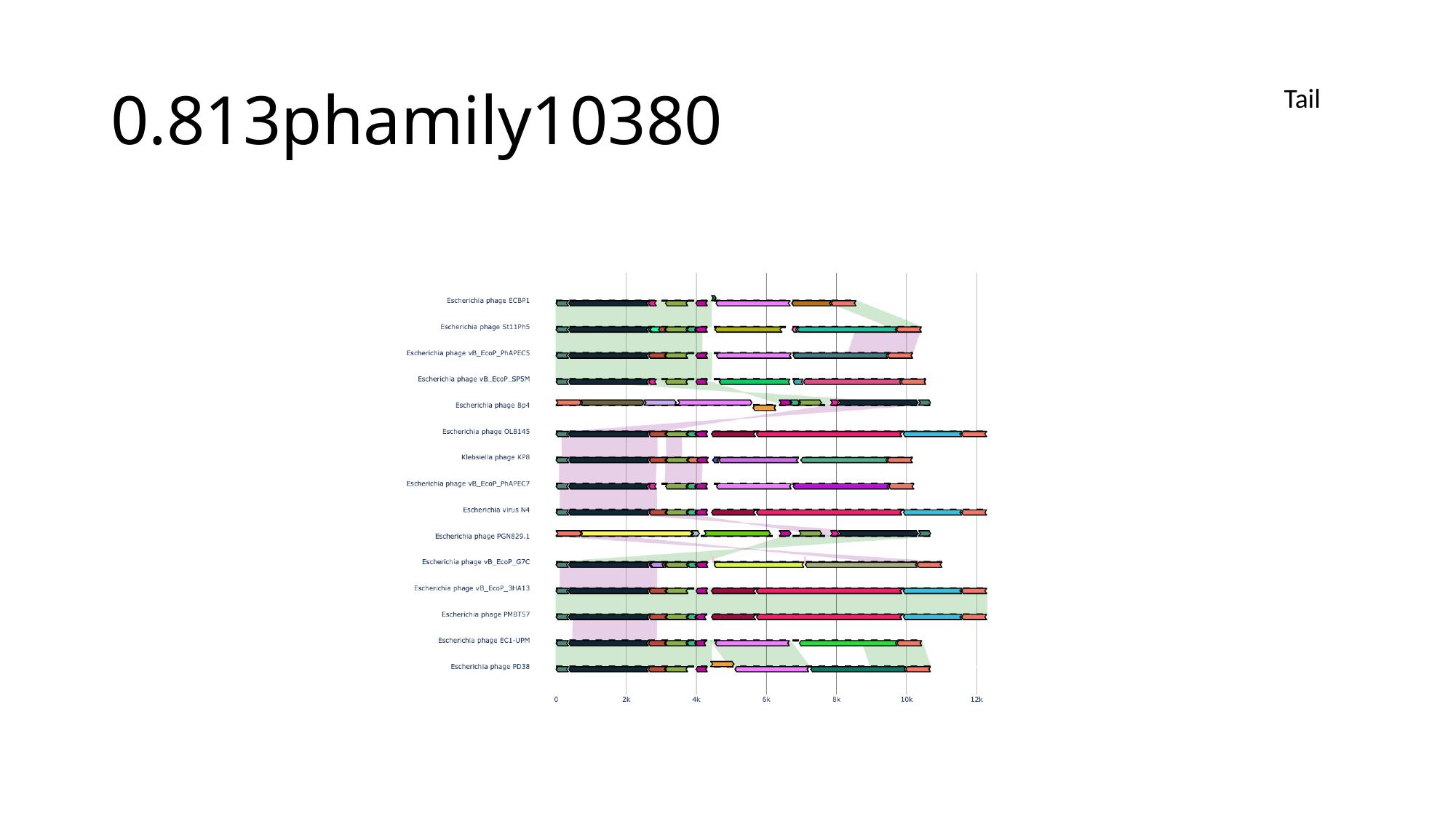

#### 0.813phamily10380
Tail

### Slide 155
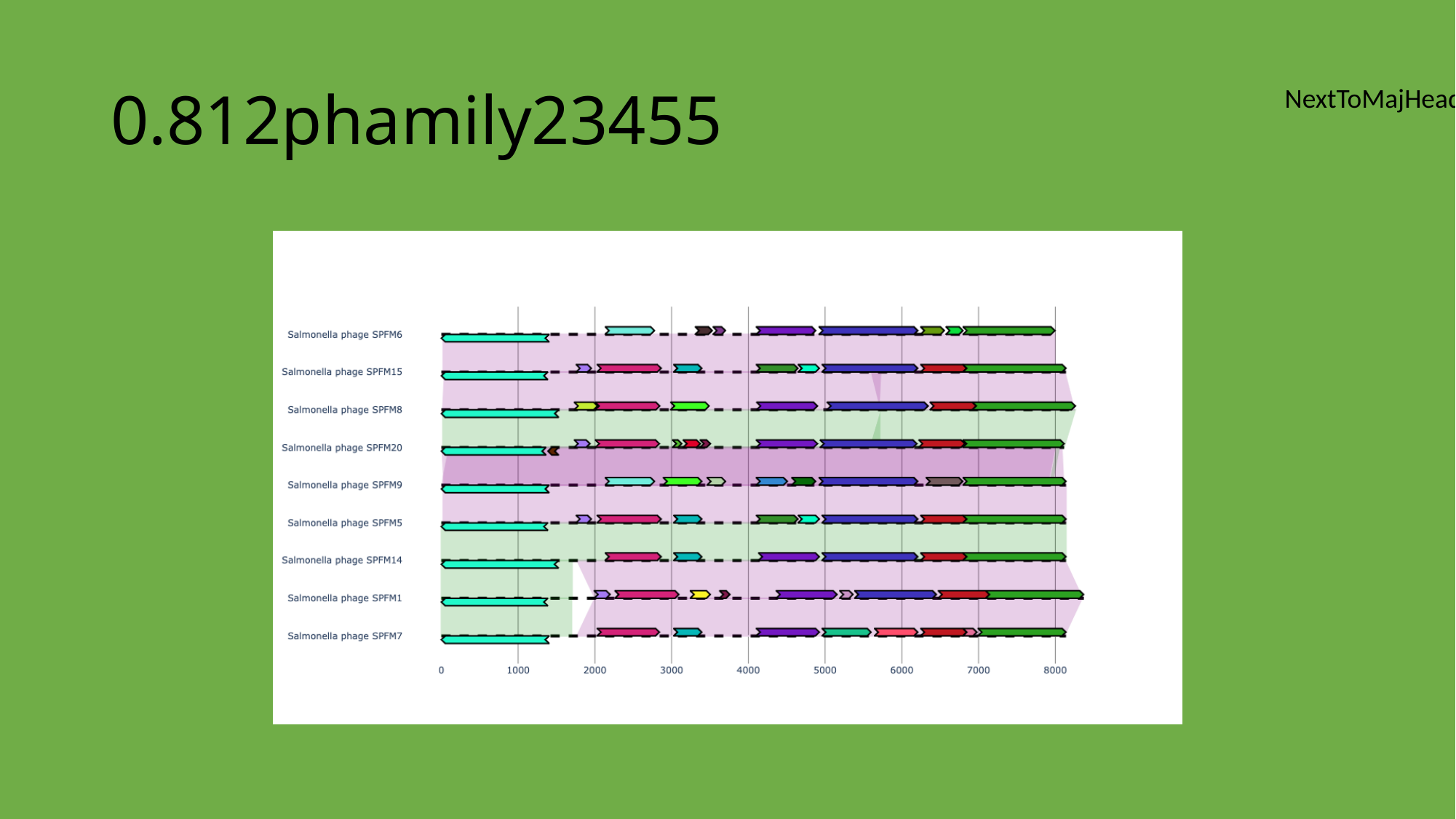

#### 0.812phamily23455
NextToMajHead

### Slide 156
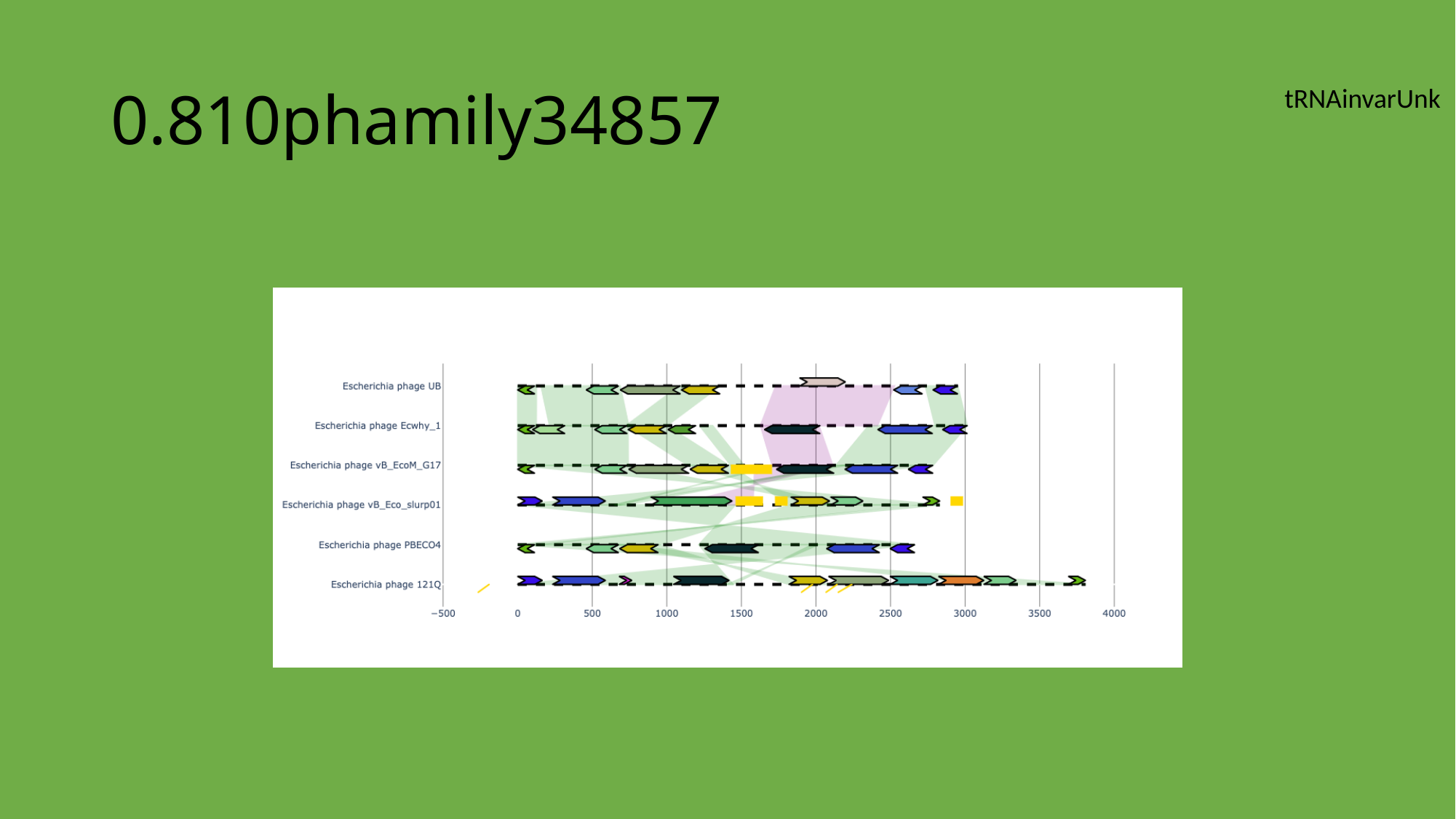

#### 0.810phamily34857
tRNAinvarUnk

### Slide 157
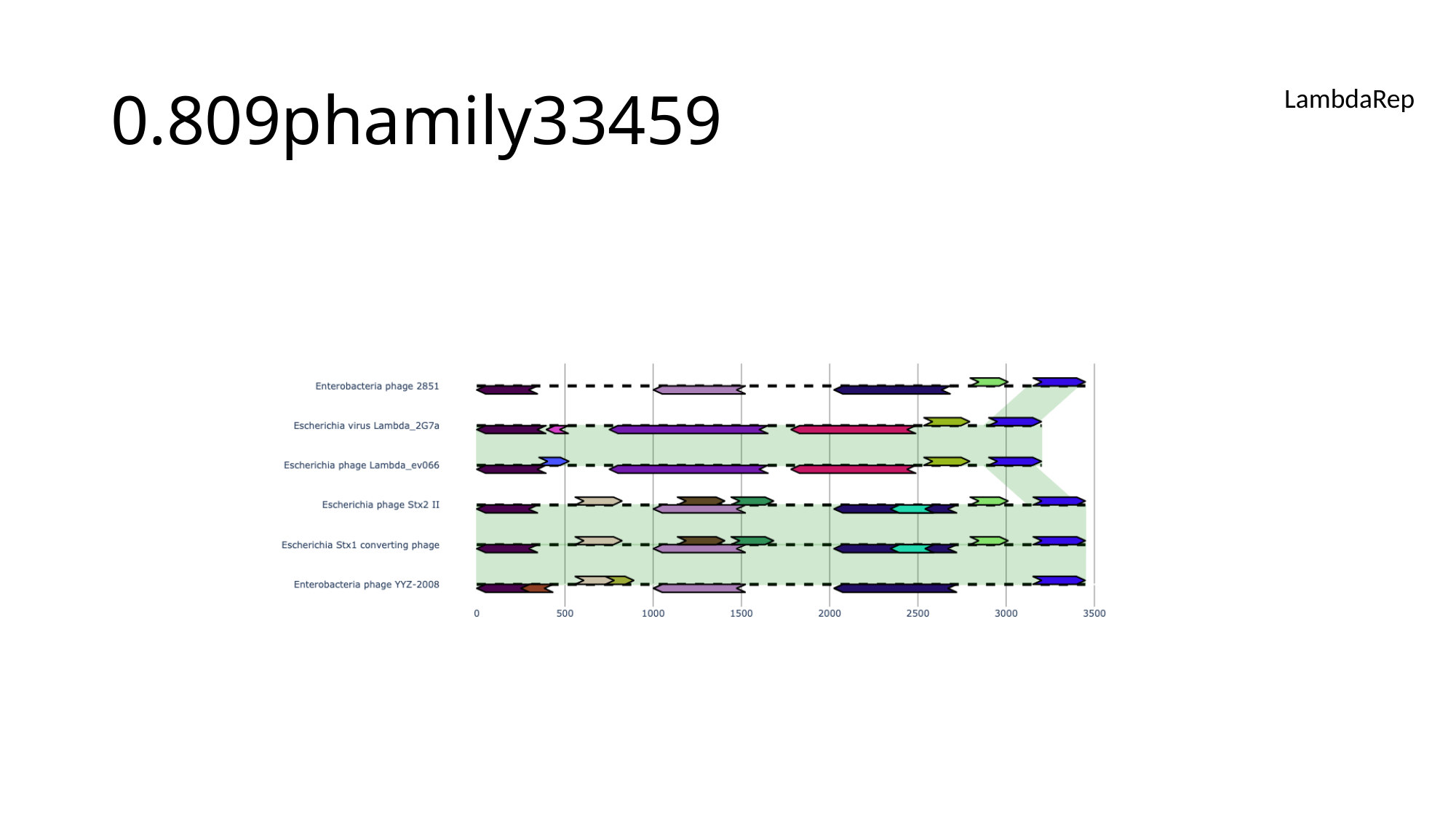

#### 0.809phamily33459
LambdaRep

### Slide 158
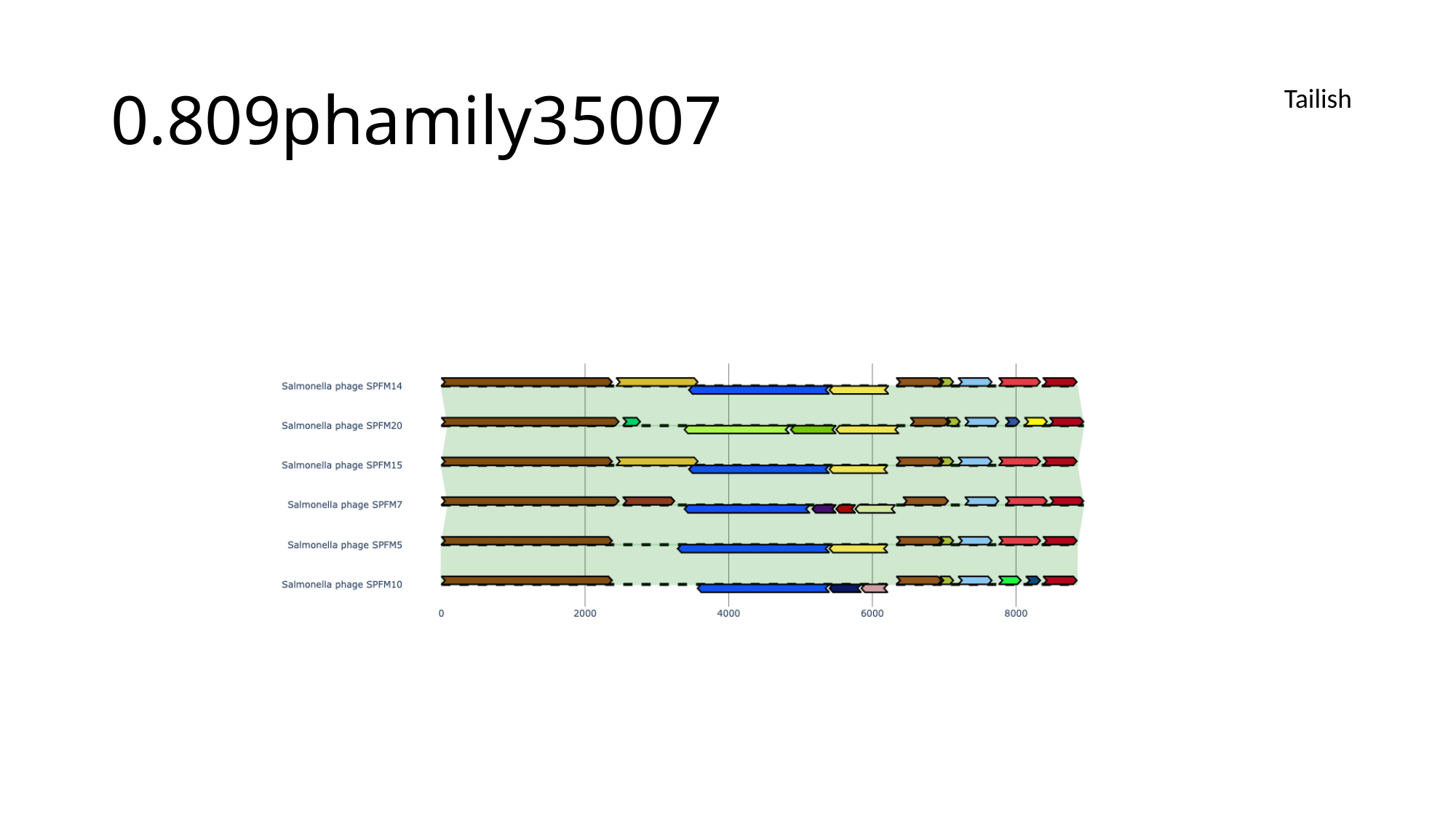

#### 0.809phamily35007
Tailish

### Slide 159
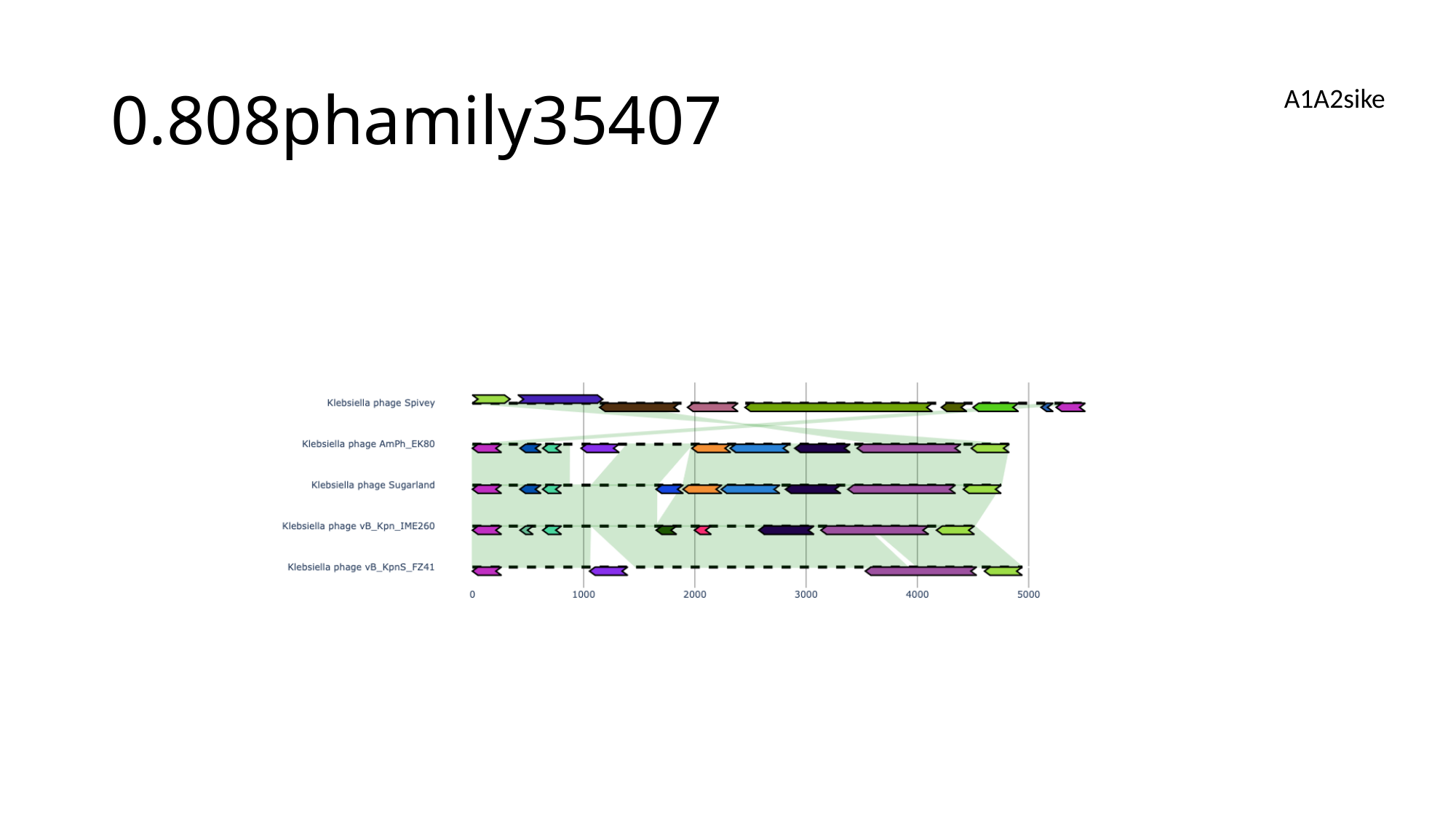

#### 0.808phamily35407
A1A2sike

### Slide 160
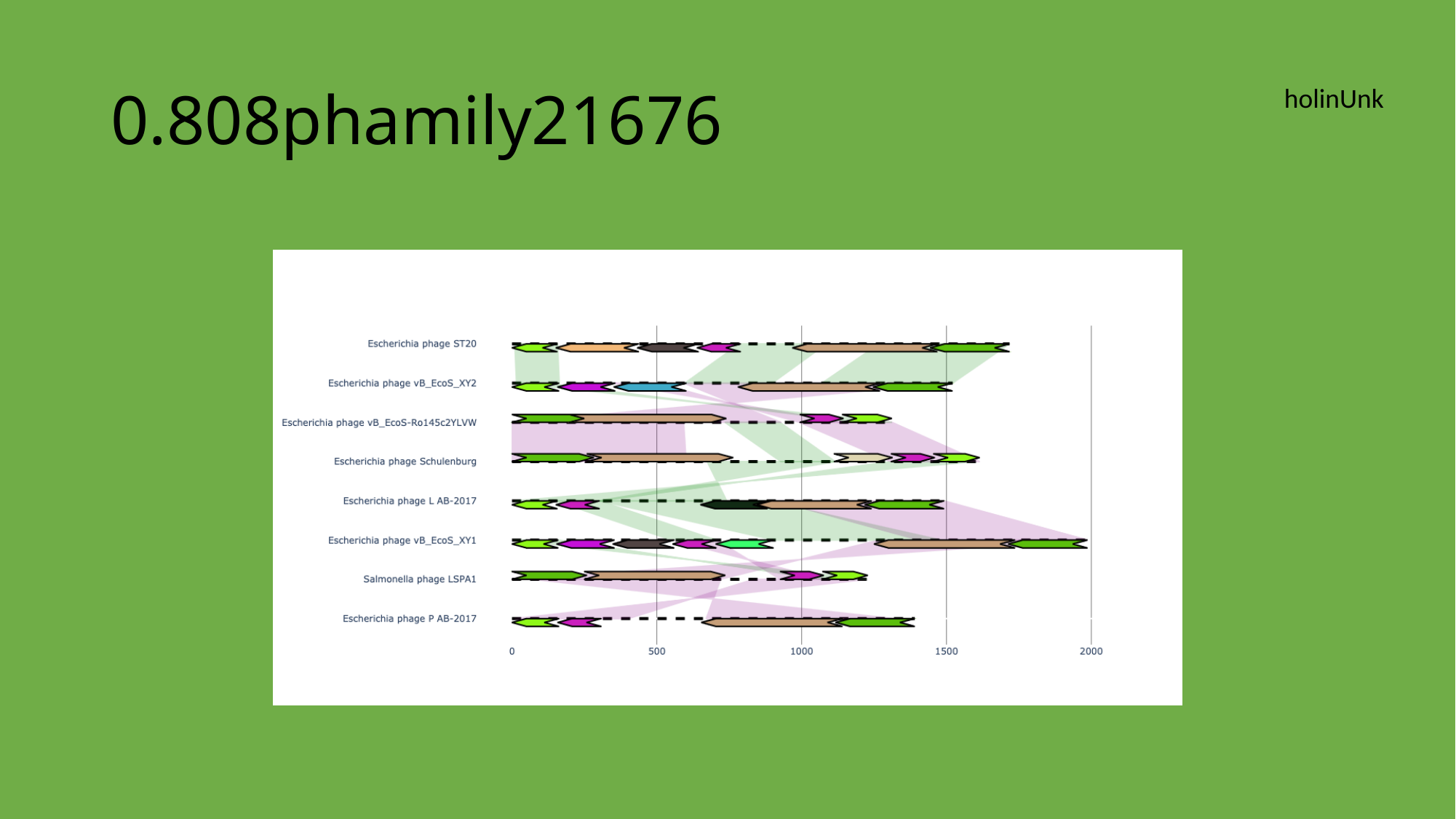

#### 0.808phamily21676
holinUnk

### Slide 161
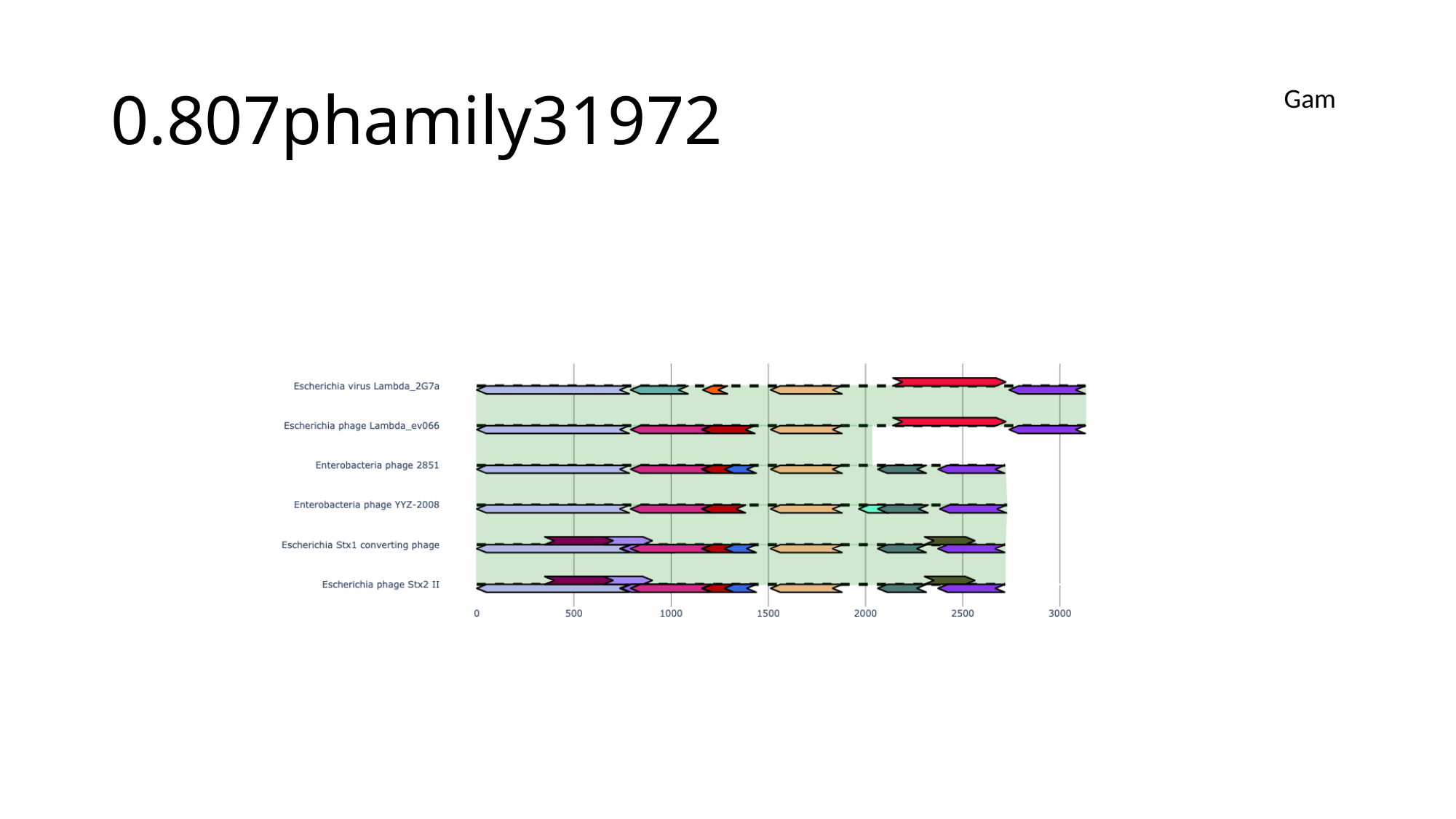

#### 0.807phamily31972
Gam

### Slide 162
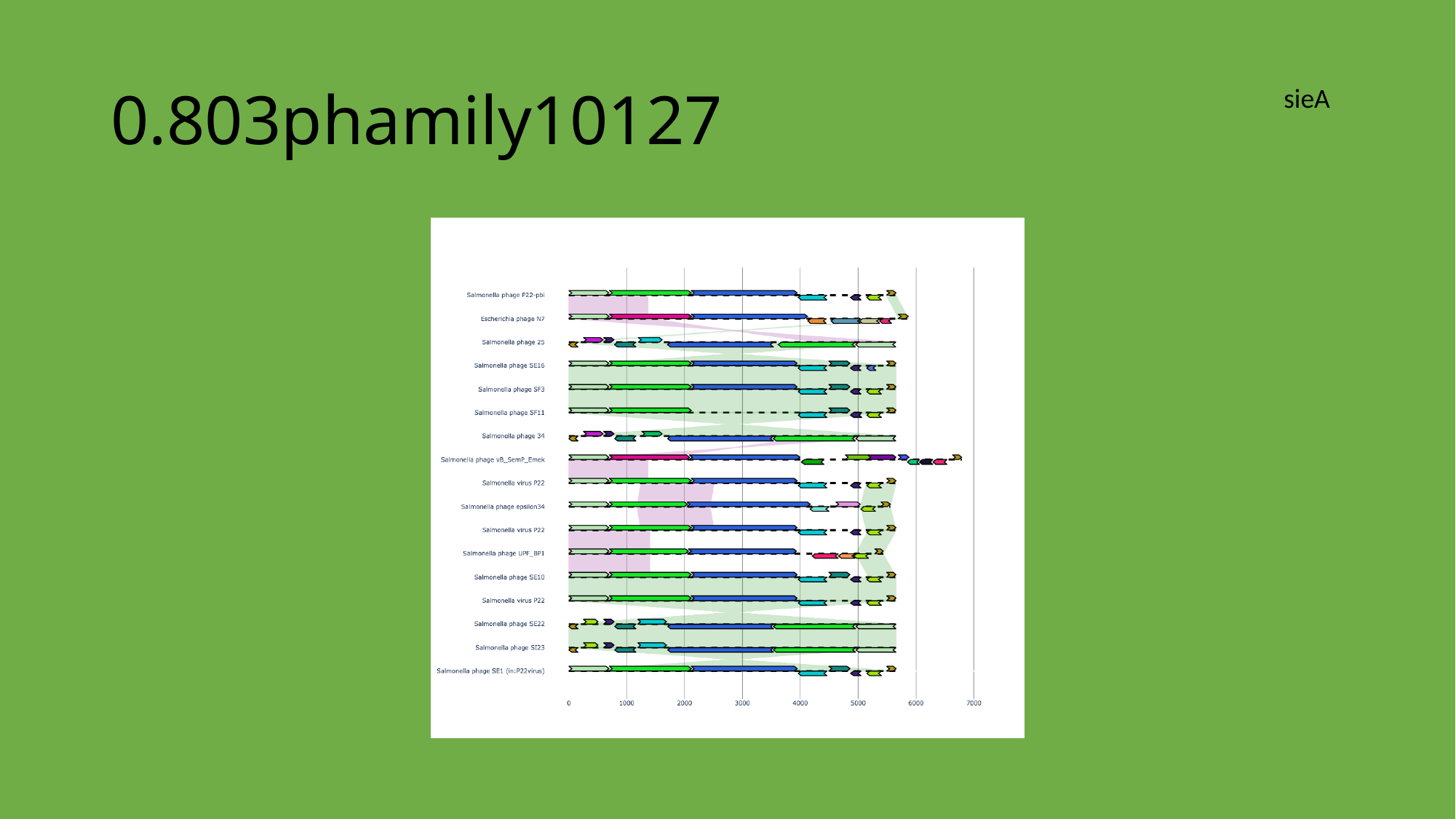

#### 0.803phamily10127
sieA

### Slide 163
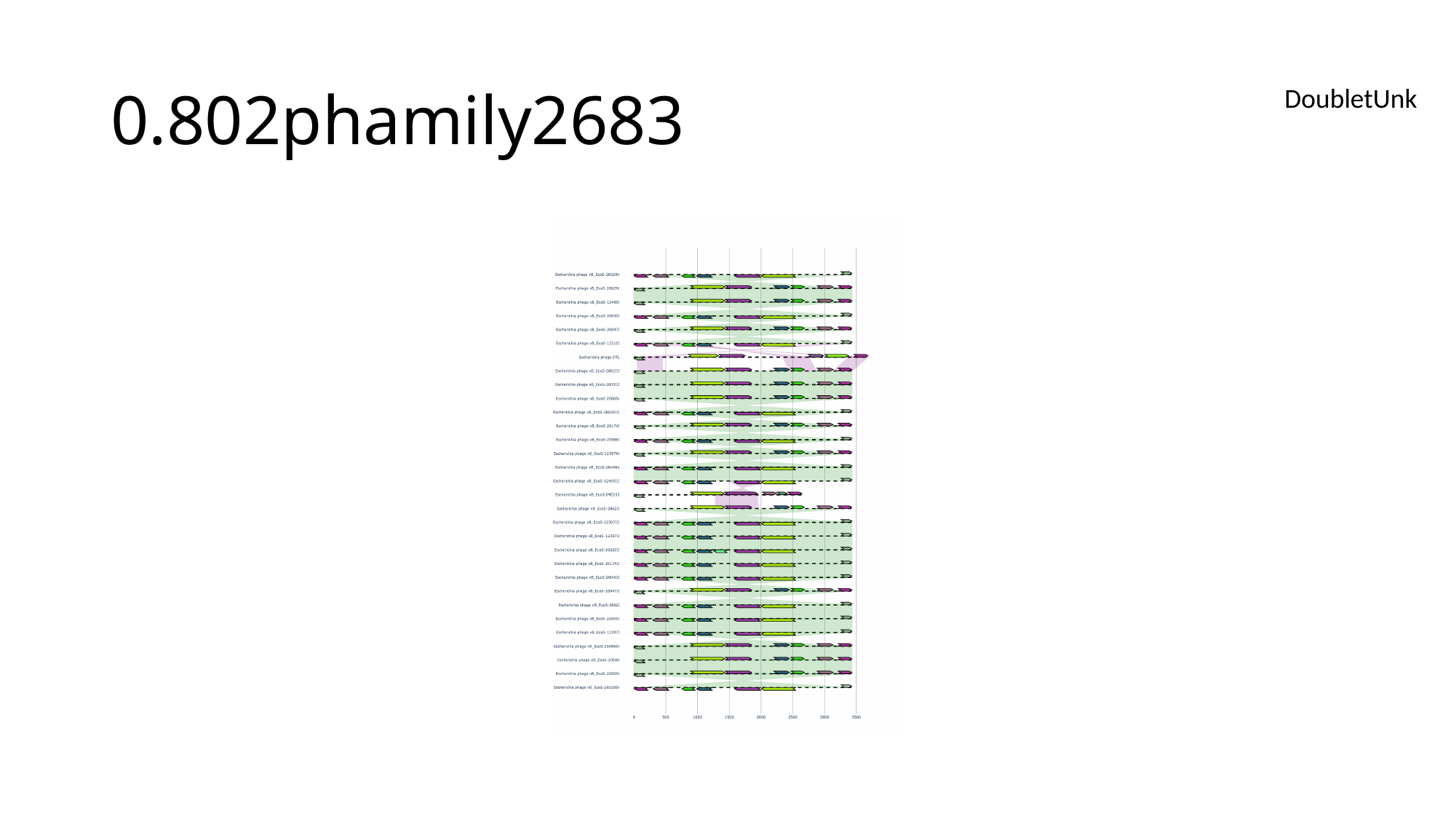

#### 0.802phamily2683
DoubletUnk

### Slide 164
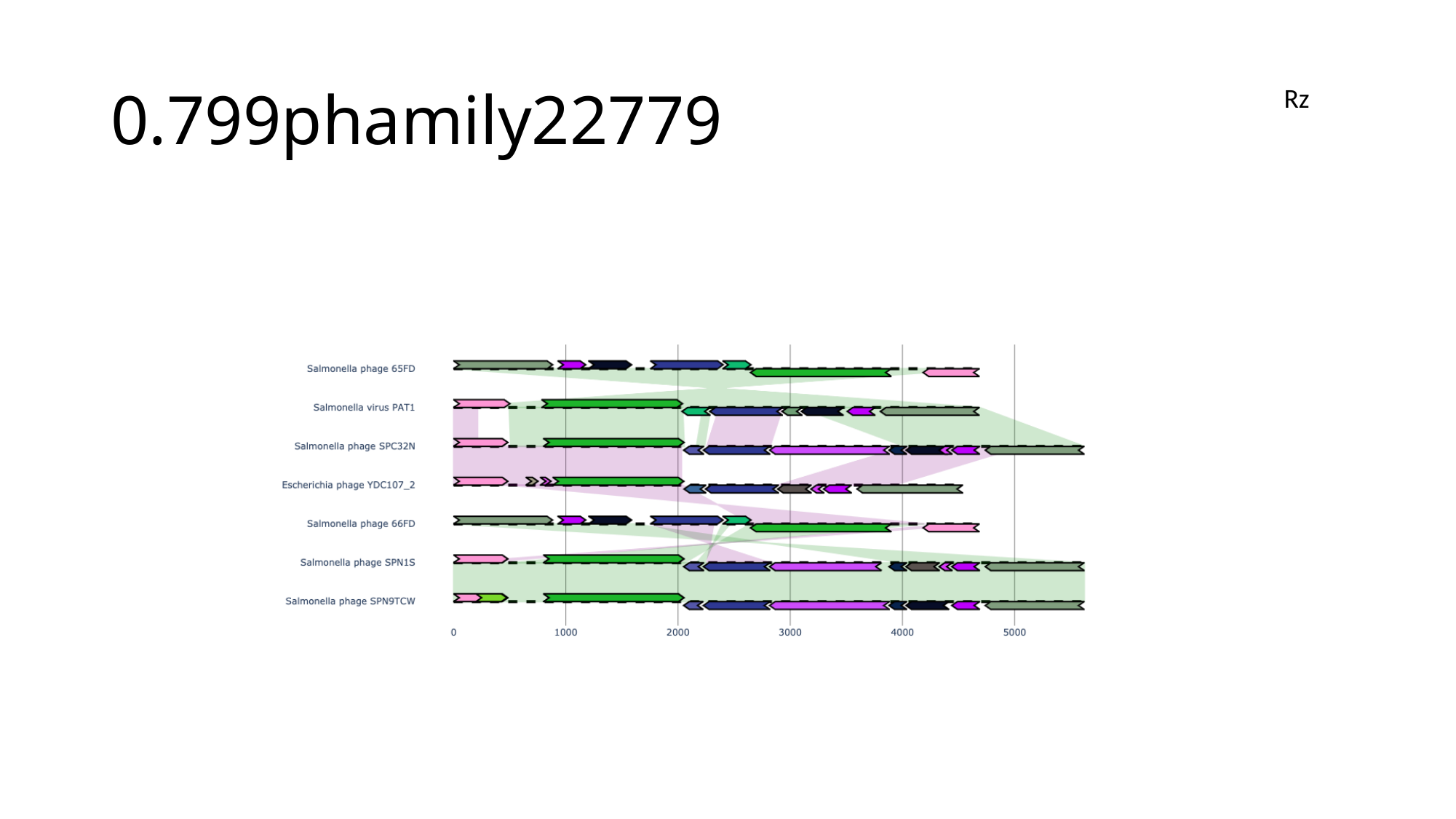

#### 0.799phamily22779
Rz

### Slide 165
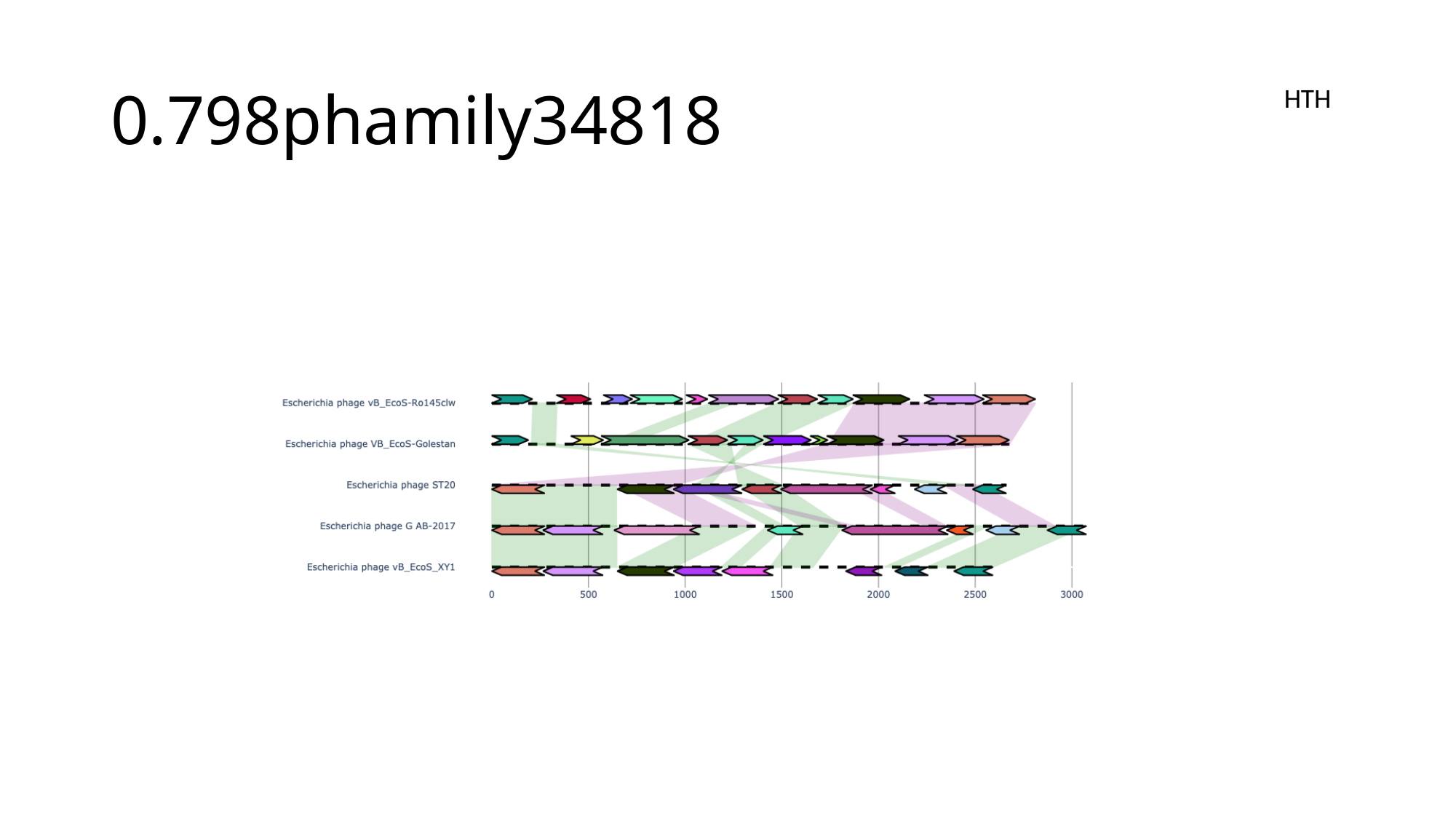

#### 0.798phamily34818
HTH

### Slide 166
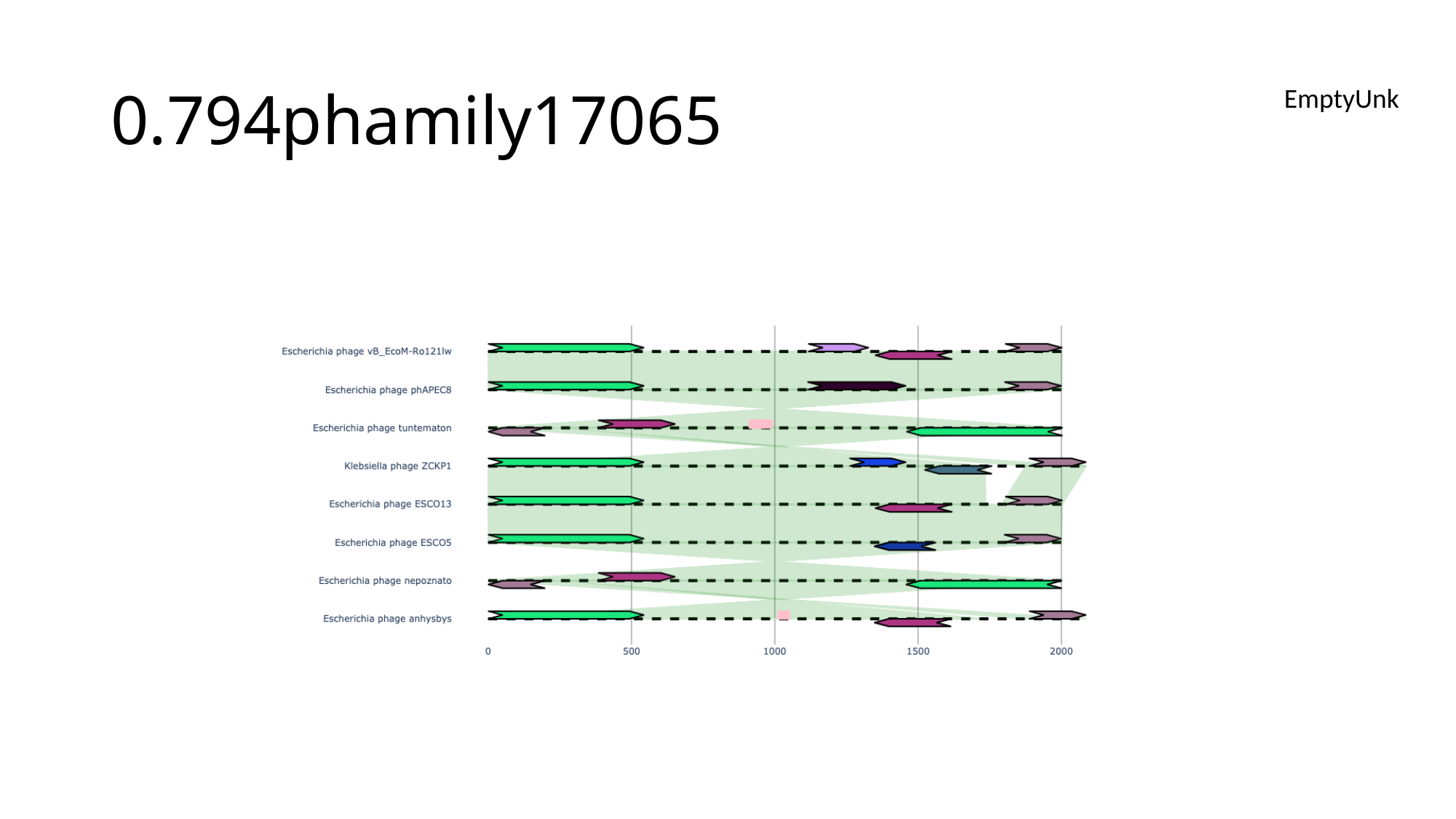

#### 0.794phamily17065
EmptyUnk

### Slide 167
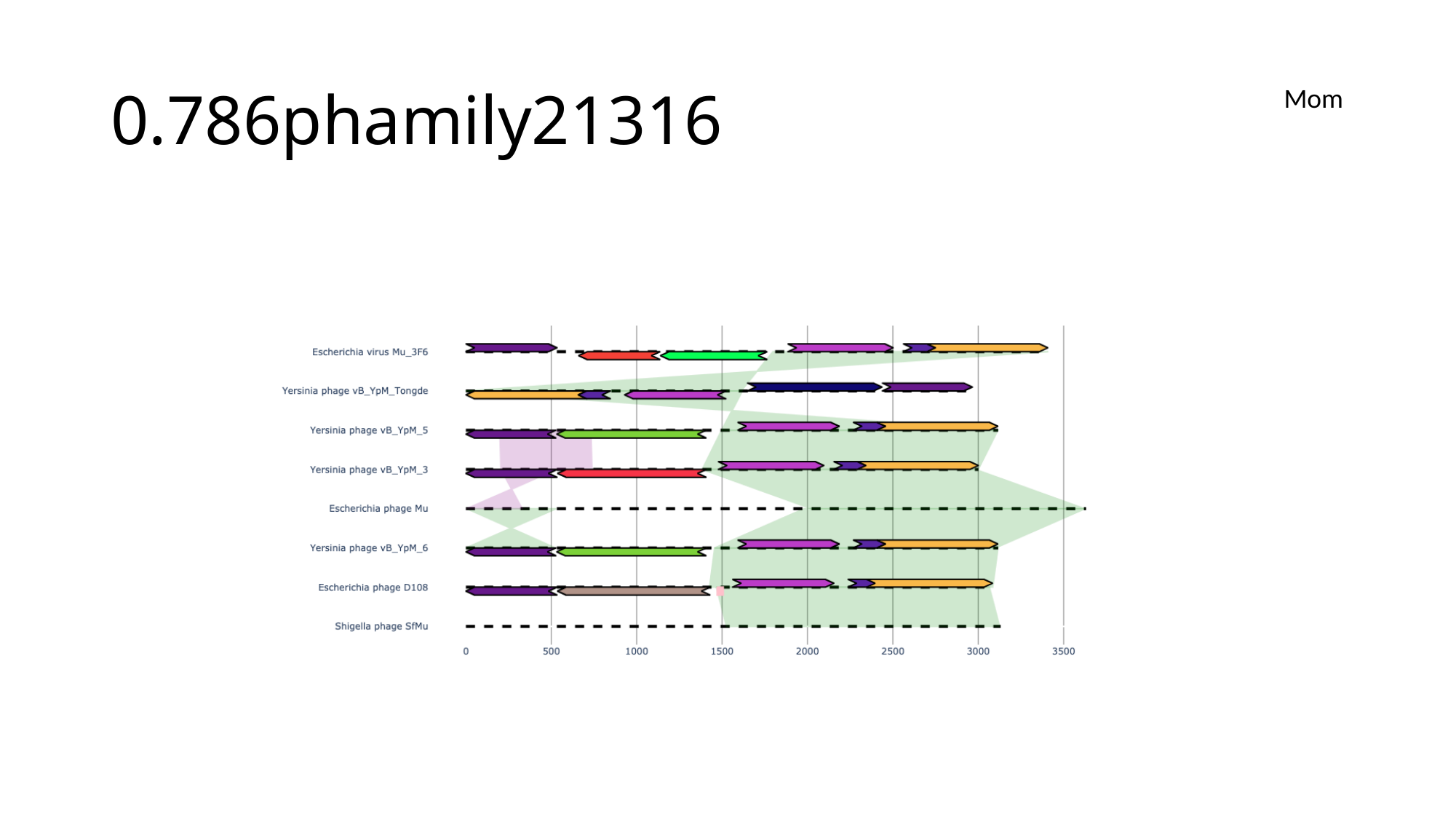

#### 0.786phamily21316
Mom

### Slide 168
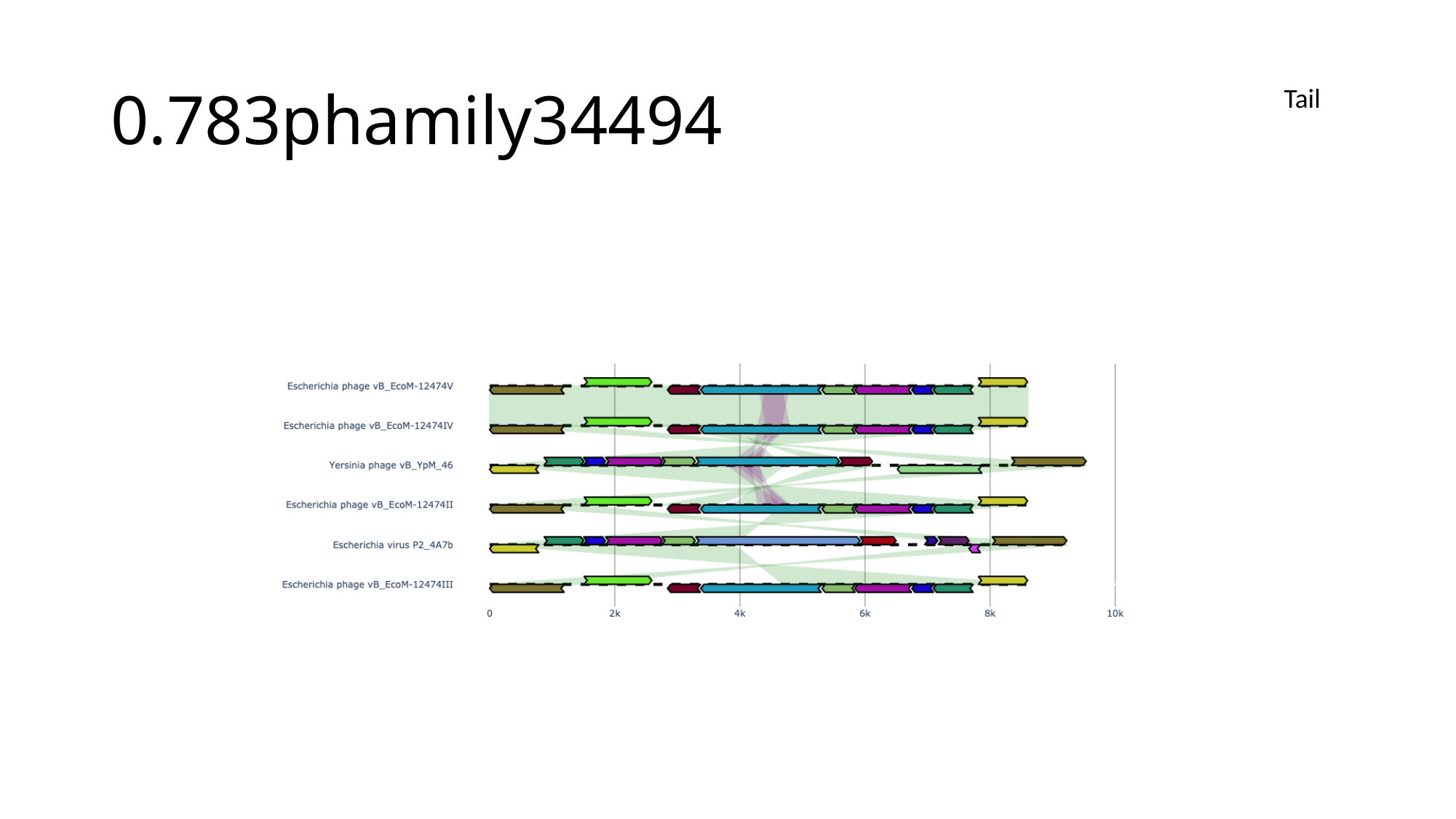

#### 0.783phamily34494
Tail

### Slide 169
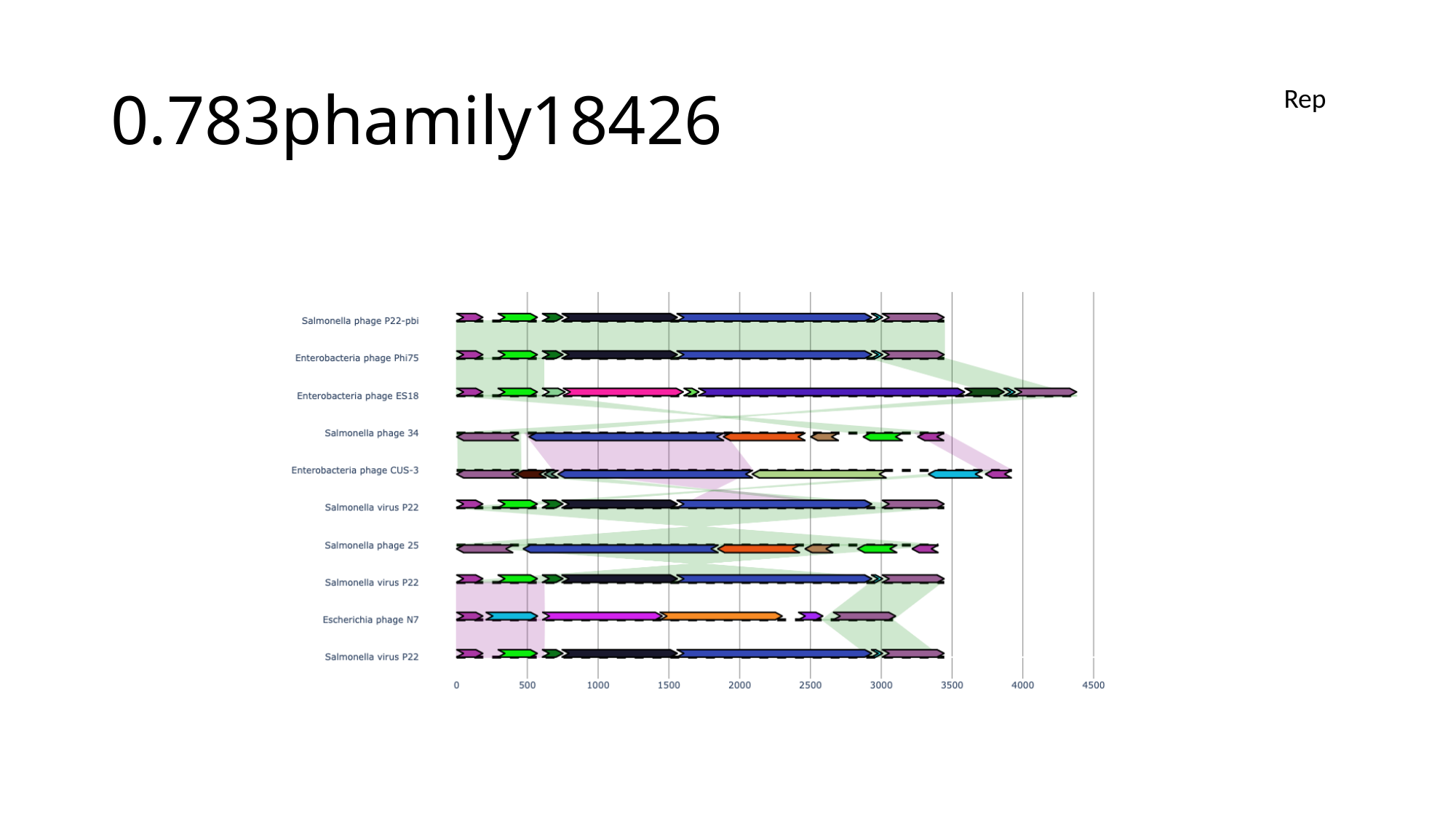

#### 0.783phamily18426
Rep

### Slide 170
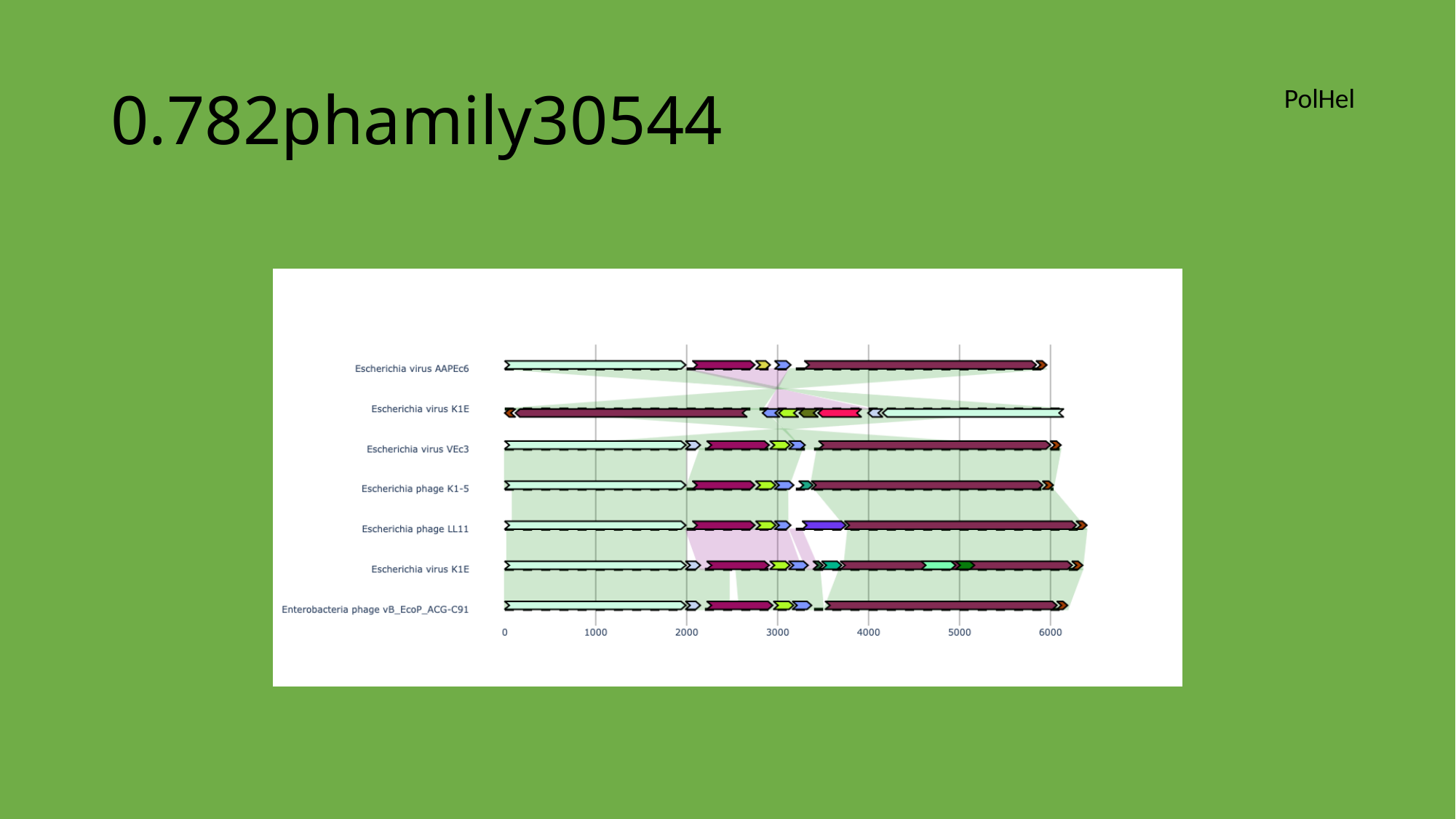

#### 0.782phamily30544
PolHel

### Slide 171
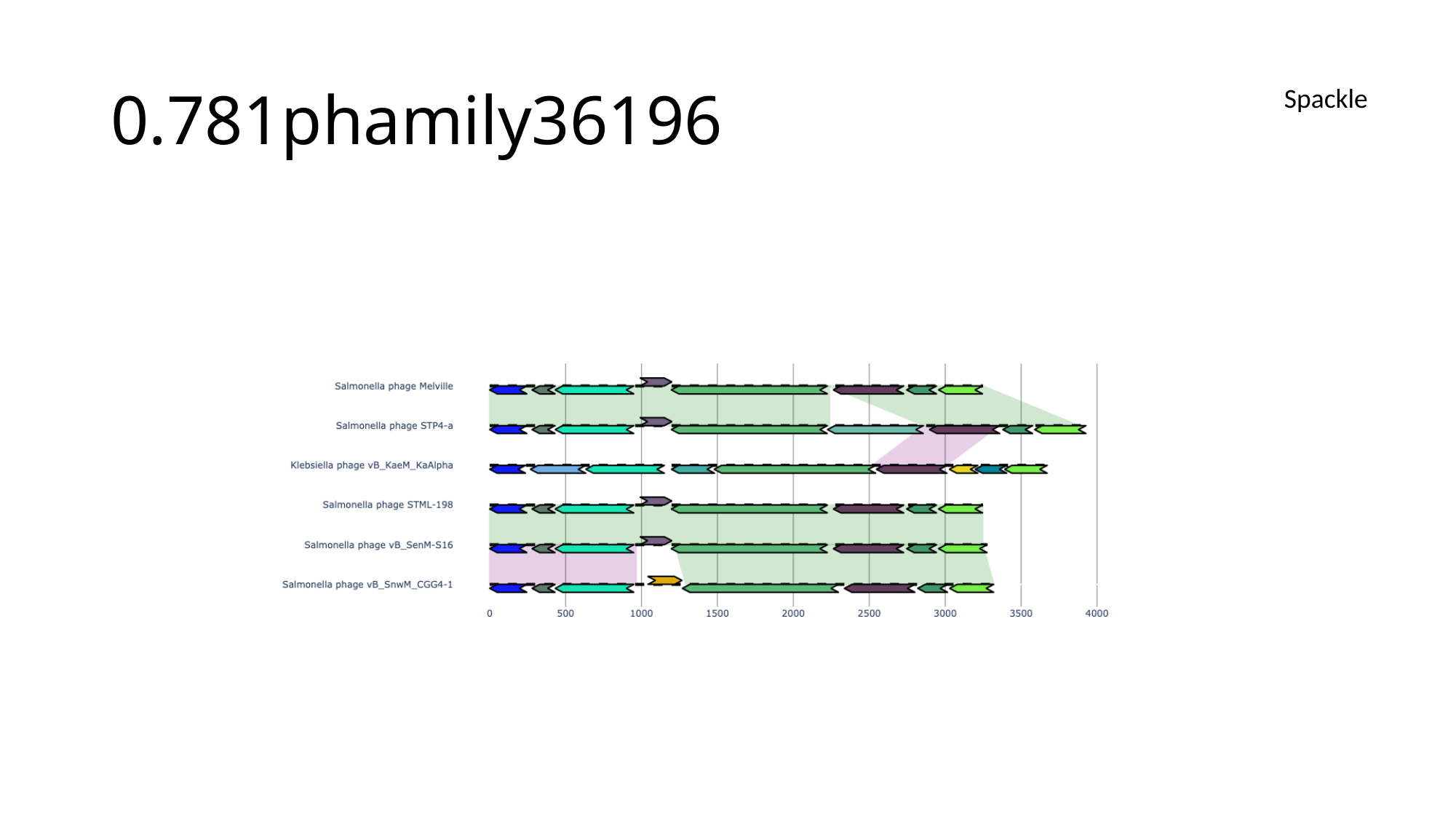

#### 0.781phamily36196
Spackle

### Slide 172
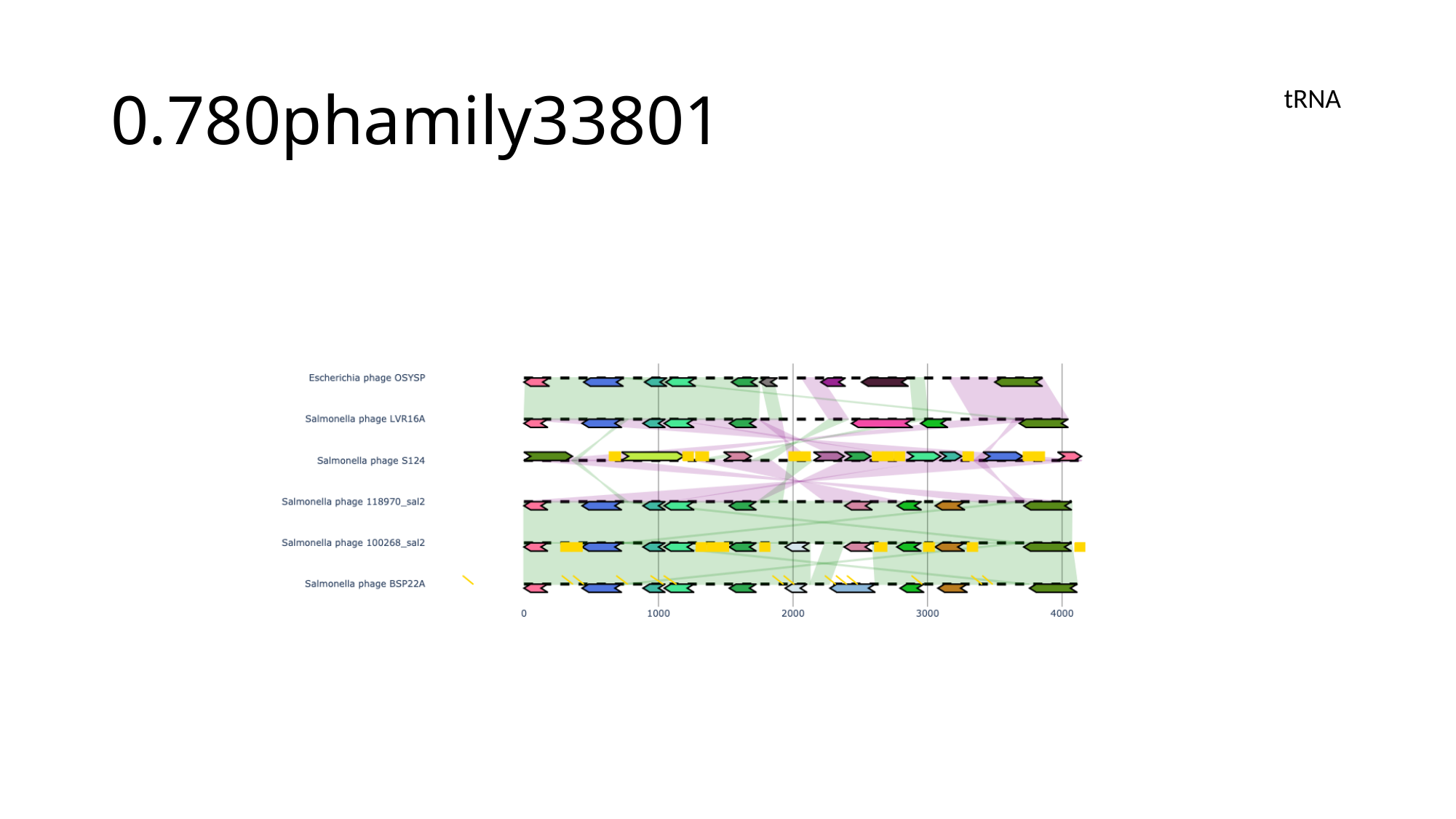

#### 0.780phamily33801
tRNA

### Slide 173
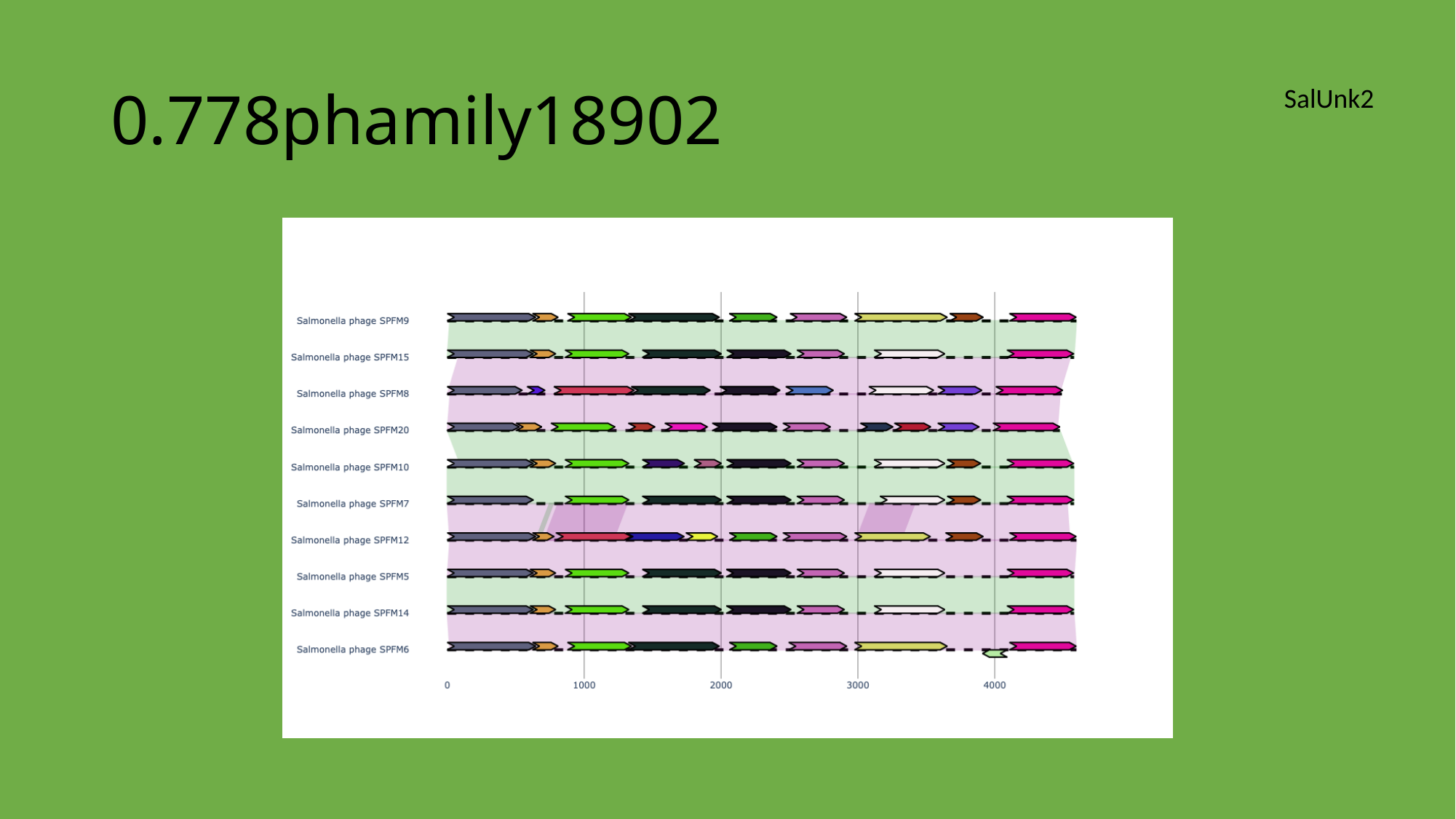

#### 0.778phamily18902
SalUnk2

### Slide 174
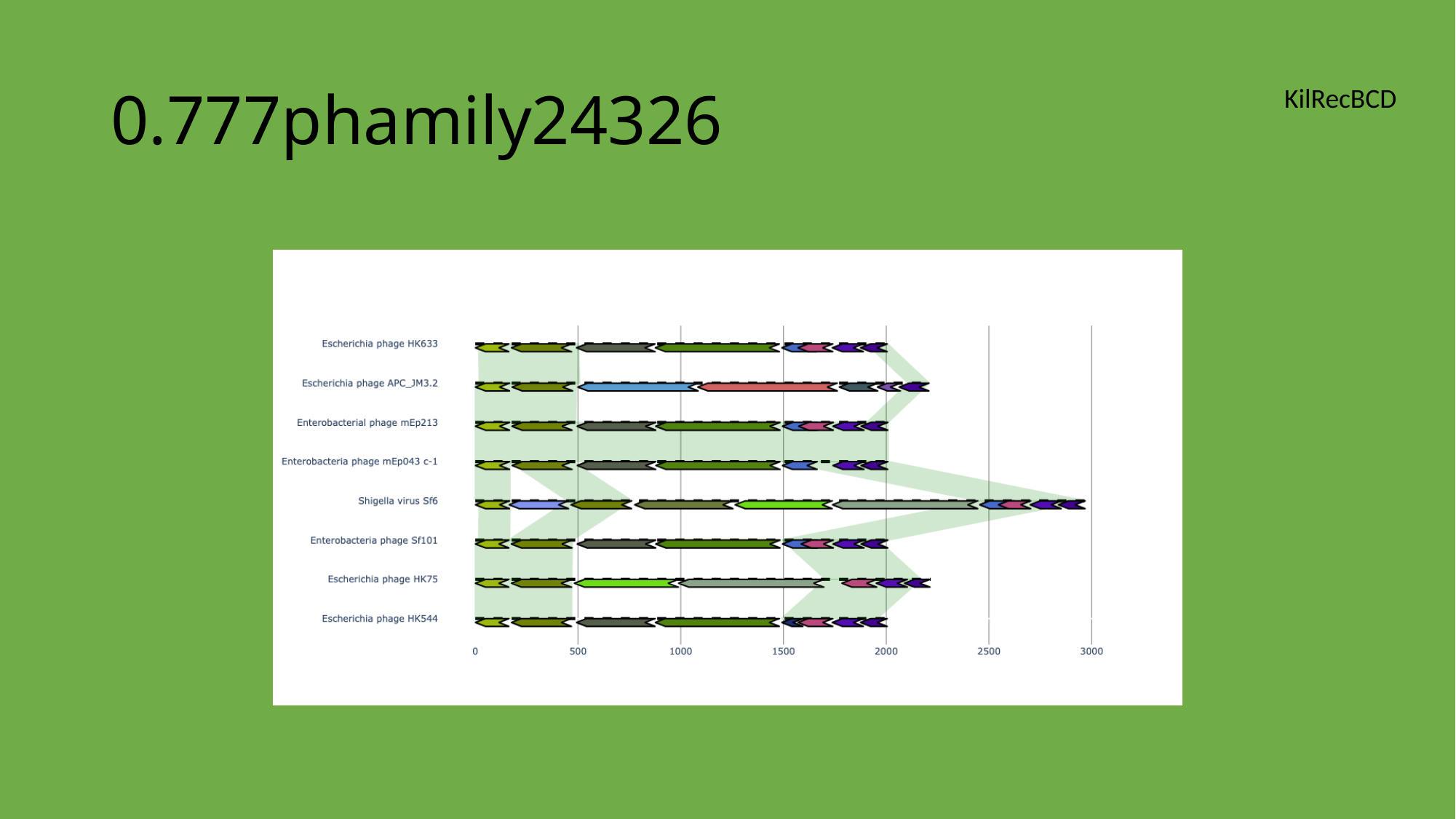

#### 0.777phamily24326
KilRecBCD

### Slide 175
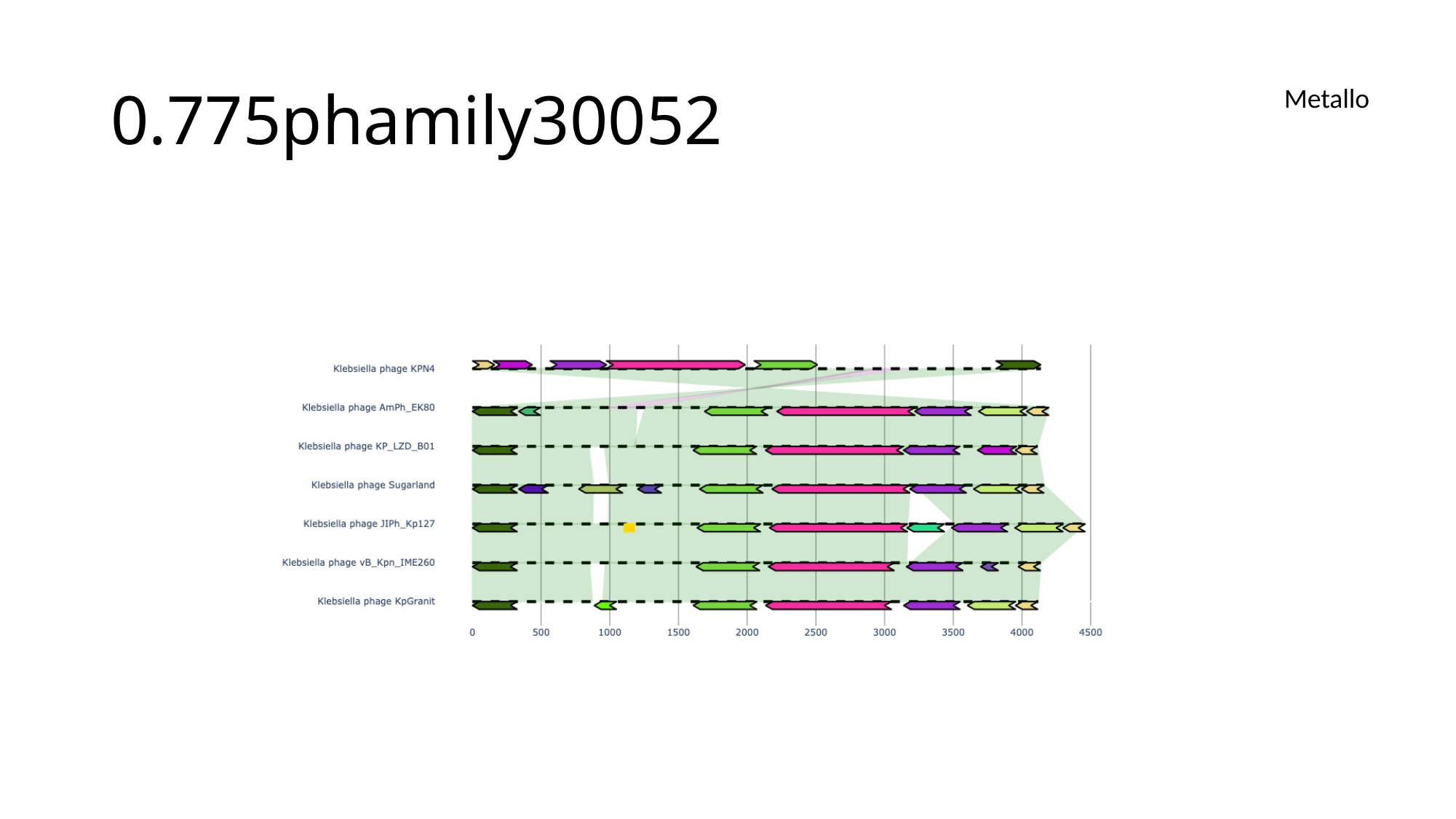

#### 0.775phamily30052
Metallo

### Slide 176
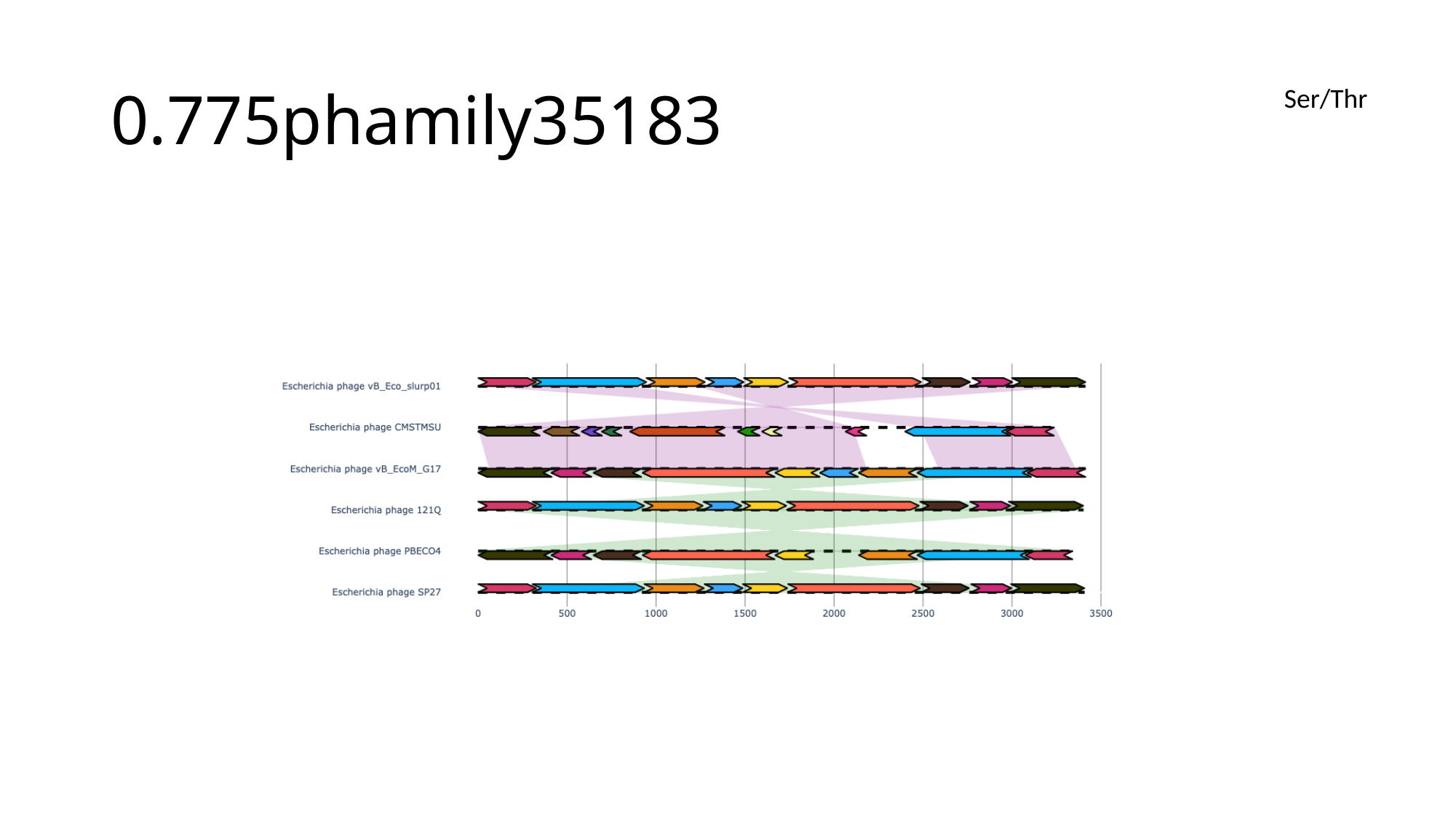

#### 0.775phamily35183
Ser/Thr

### Slide 177
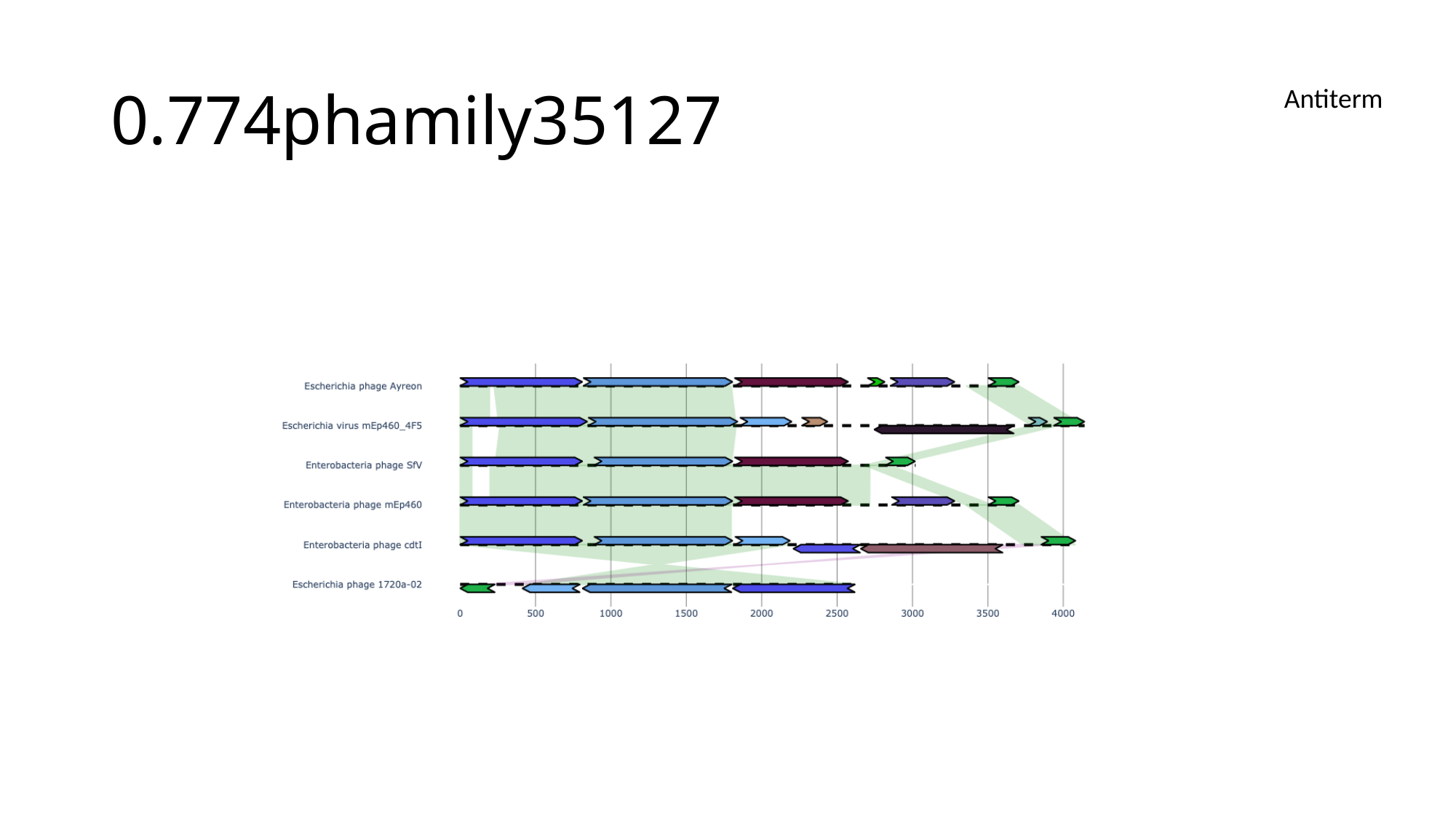

#### 0.774phamily35127
Antiterm

### Slide 178
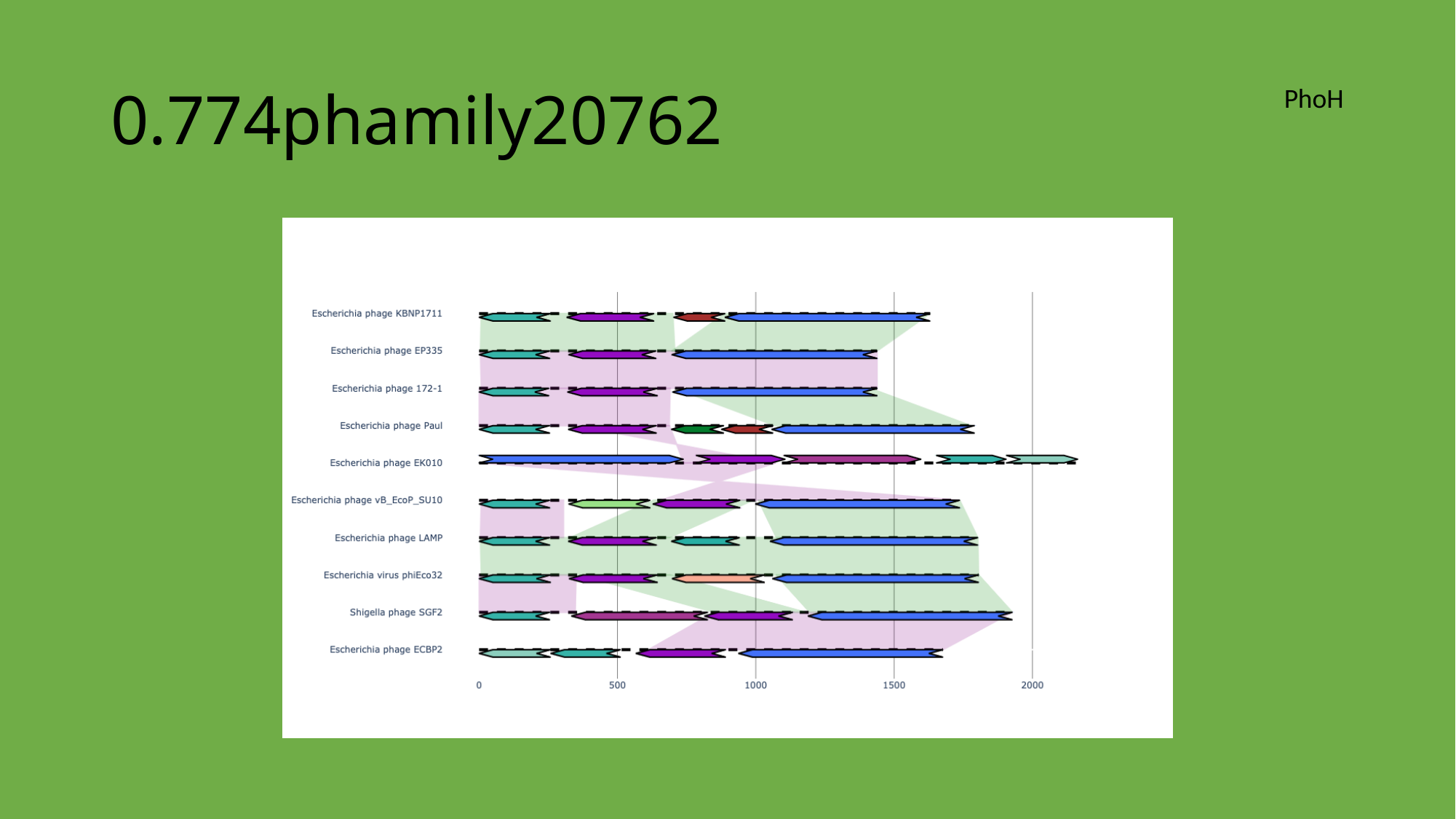

#### 0.774phamily20762
PhoH

### Slide 179
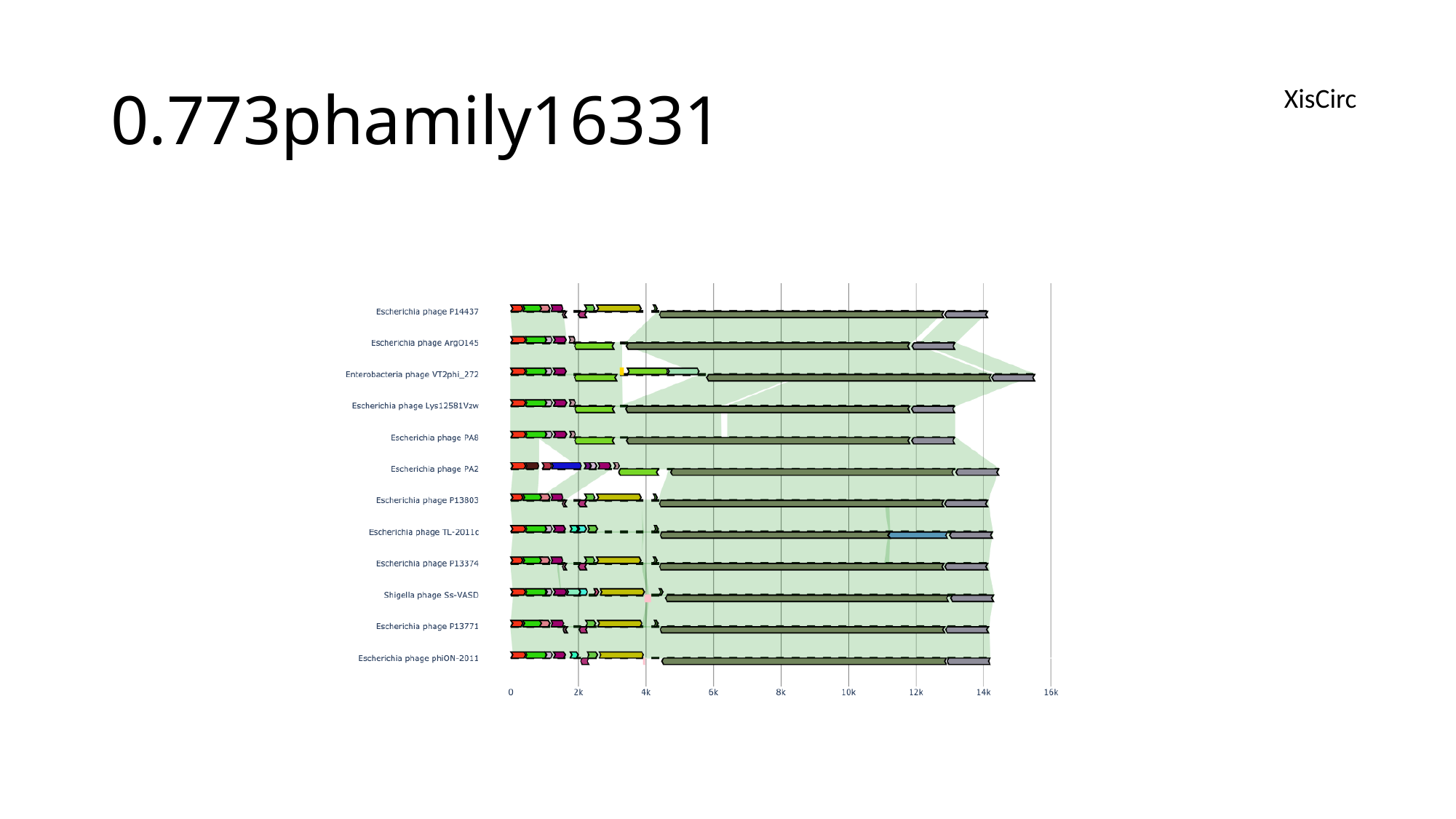

#### 0.773phamily16331
XisCirc

### Slide 180
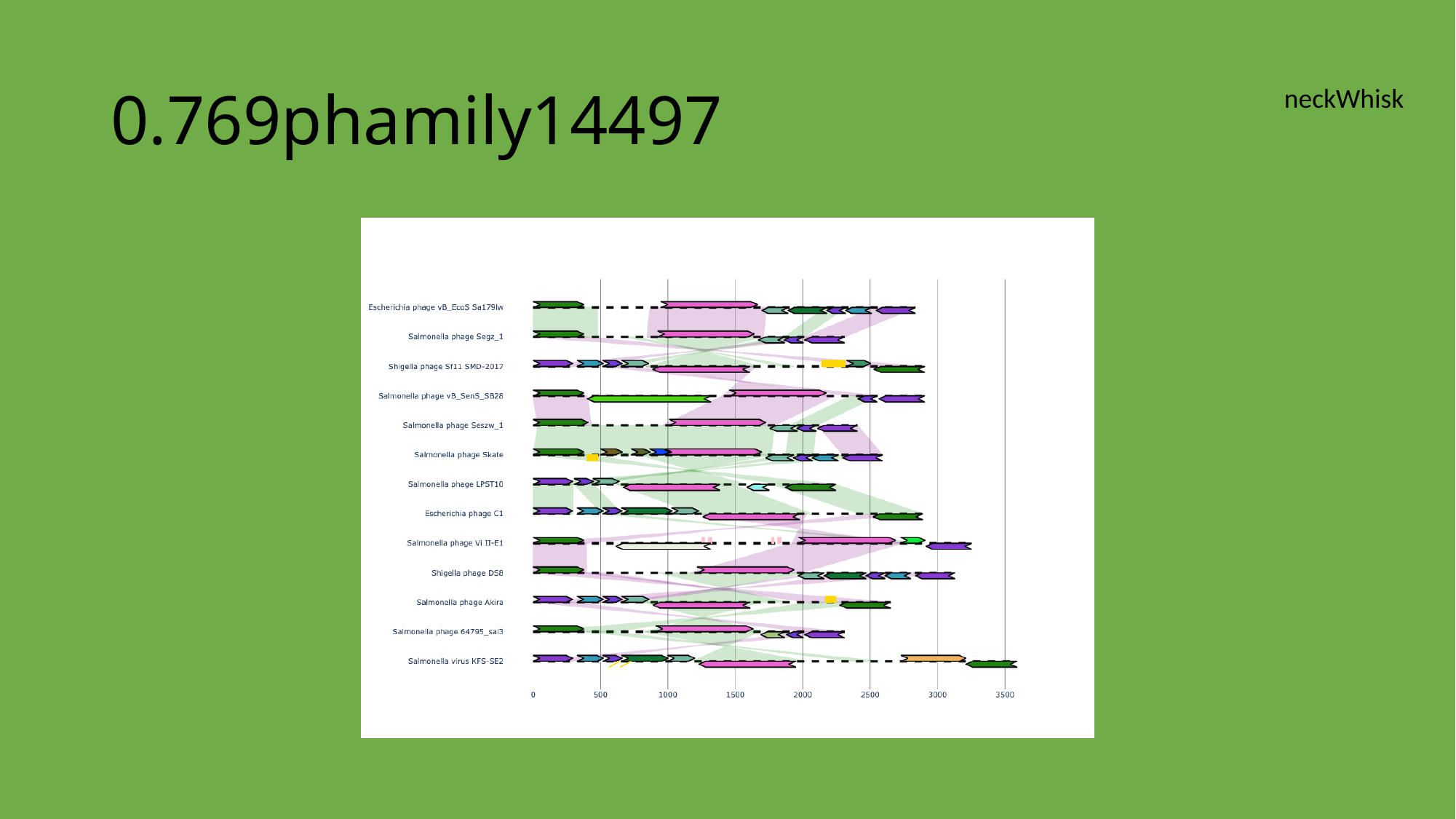

#### 0.769phamily14497
neckWhisk

### Slide 181
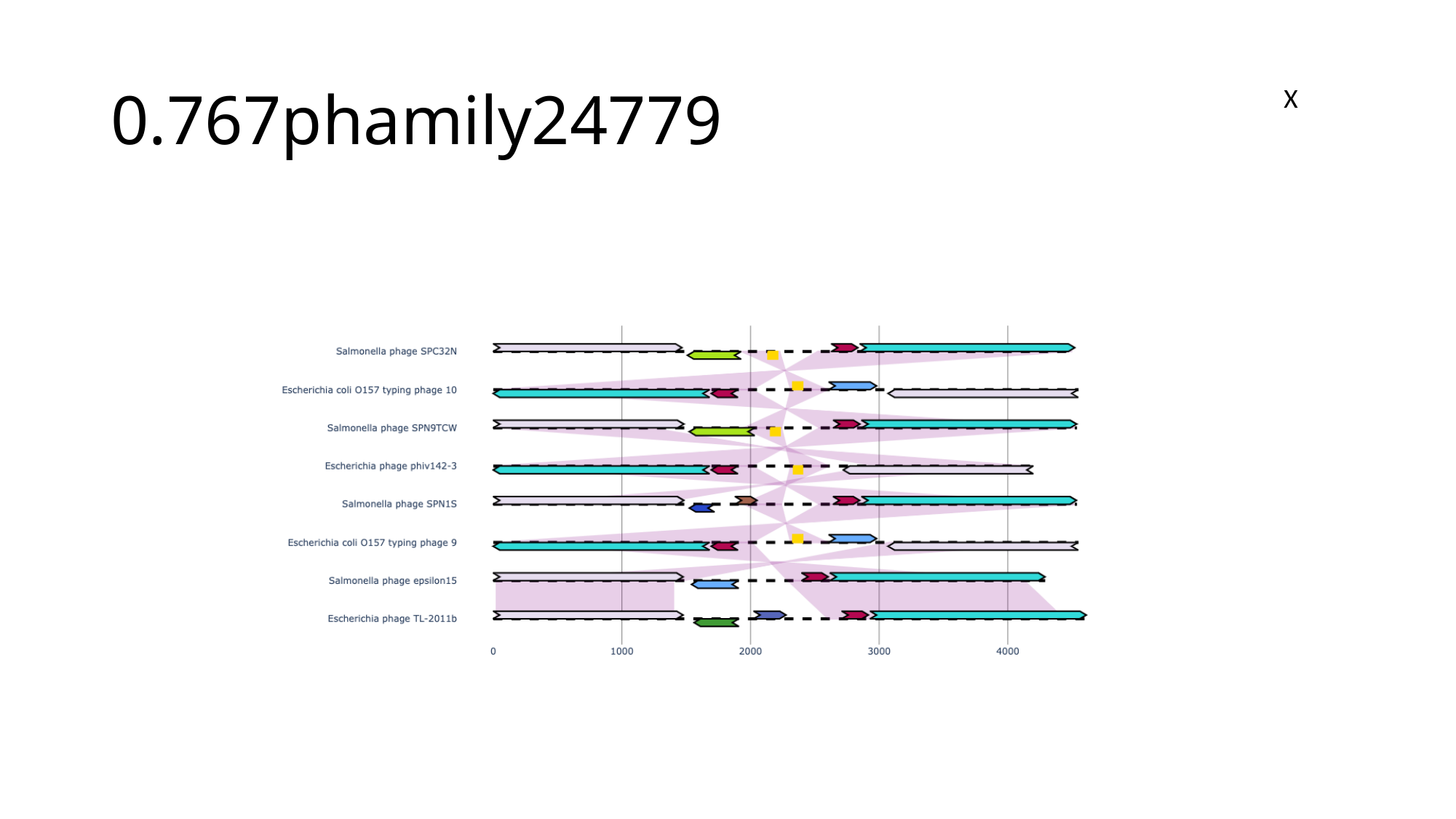

#### 0.767phamily24779
X

### Slide 182
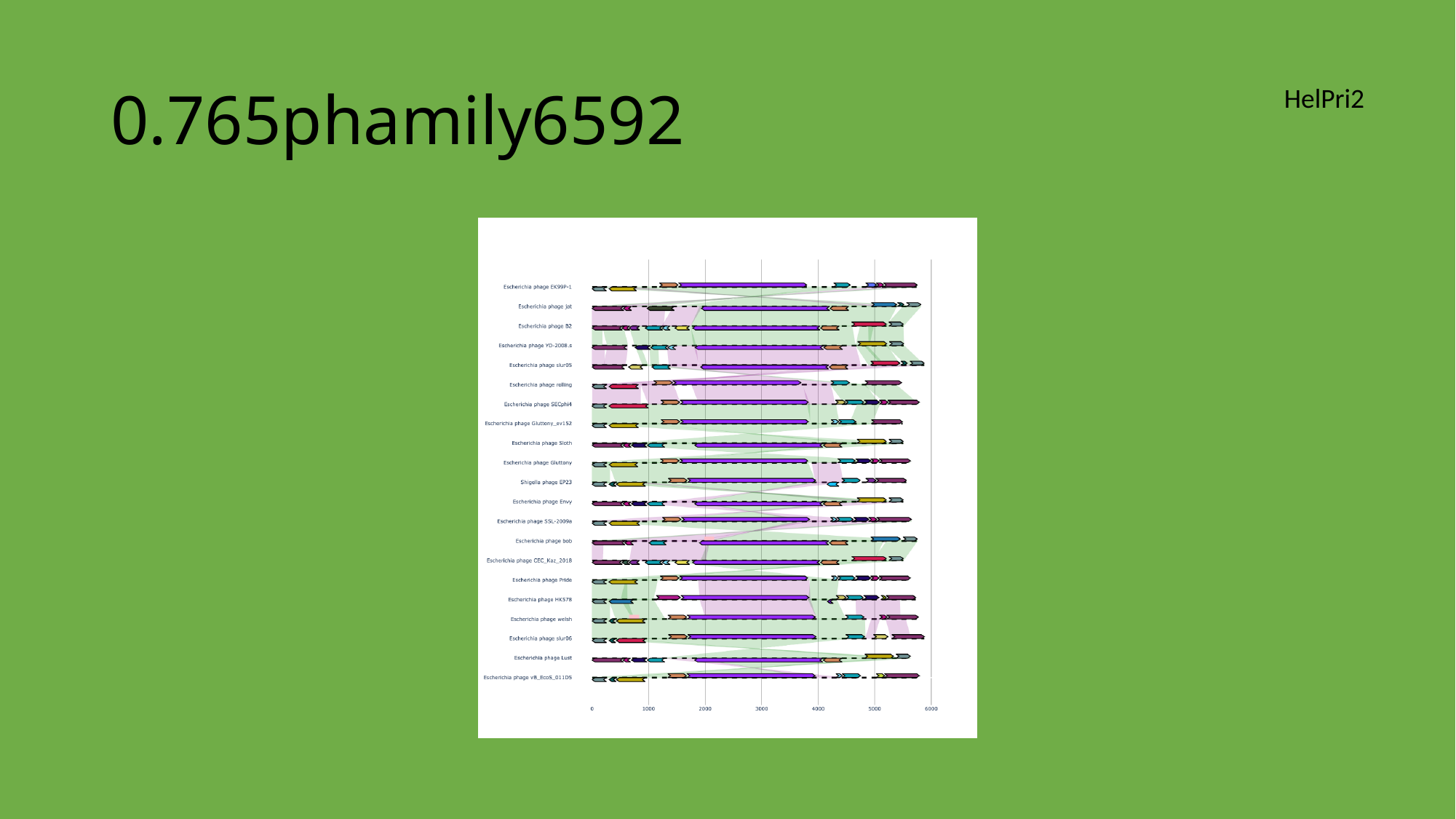

#### 0.765phamily6592
HelPri2

### Slide 183
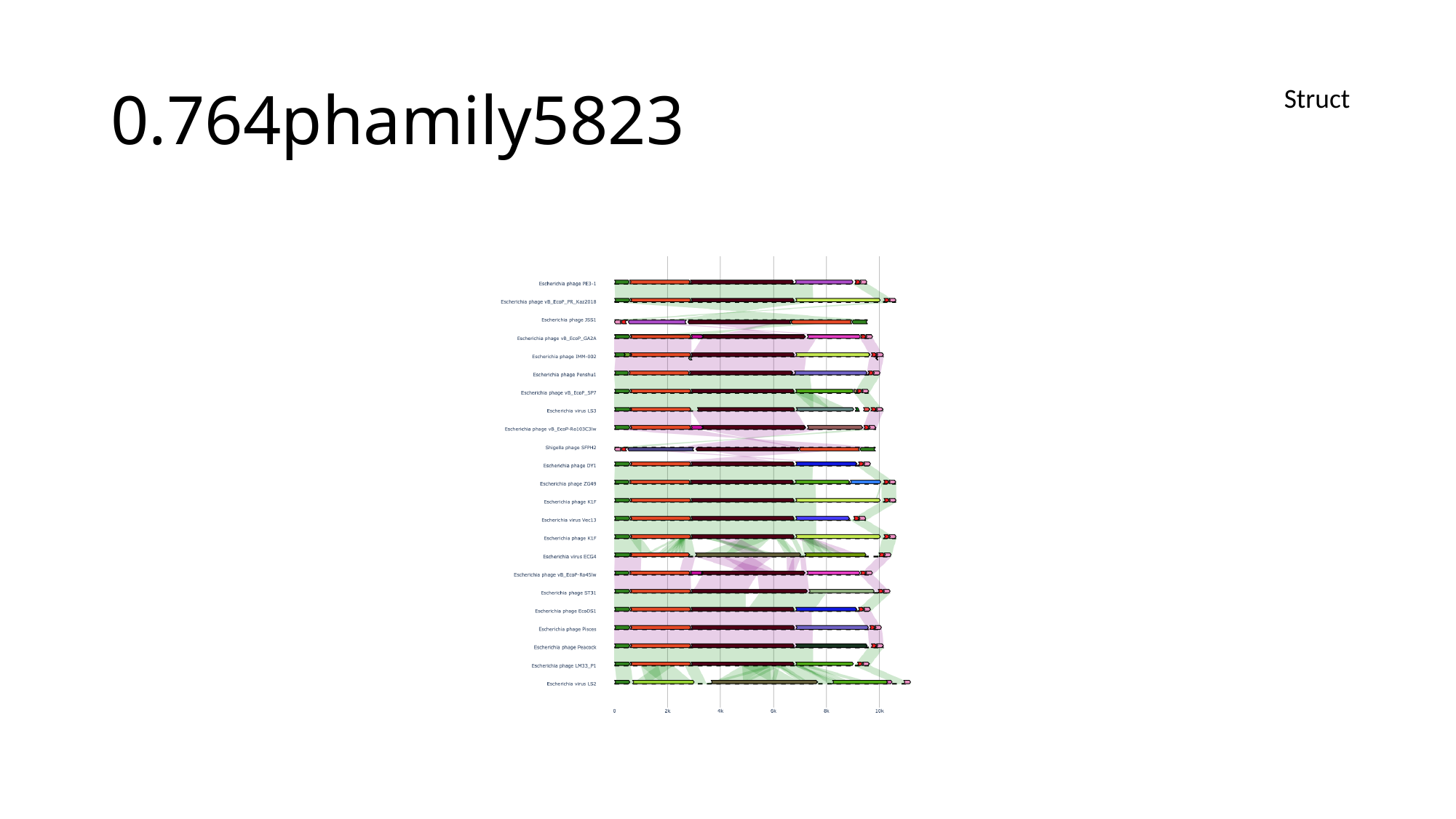

#### 0.764phamily5823
Struct

### Slide 184
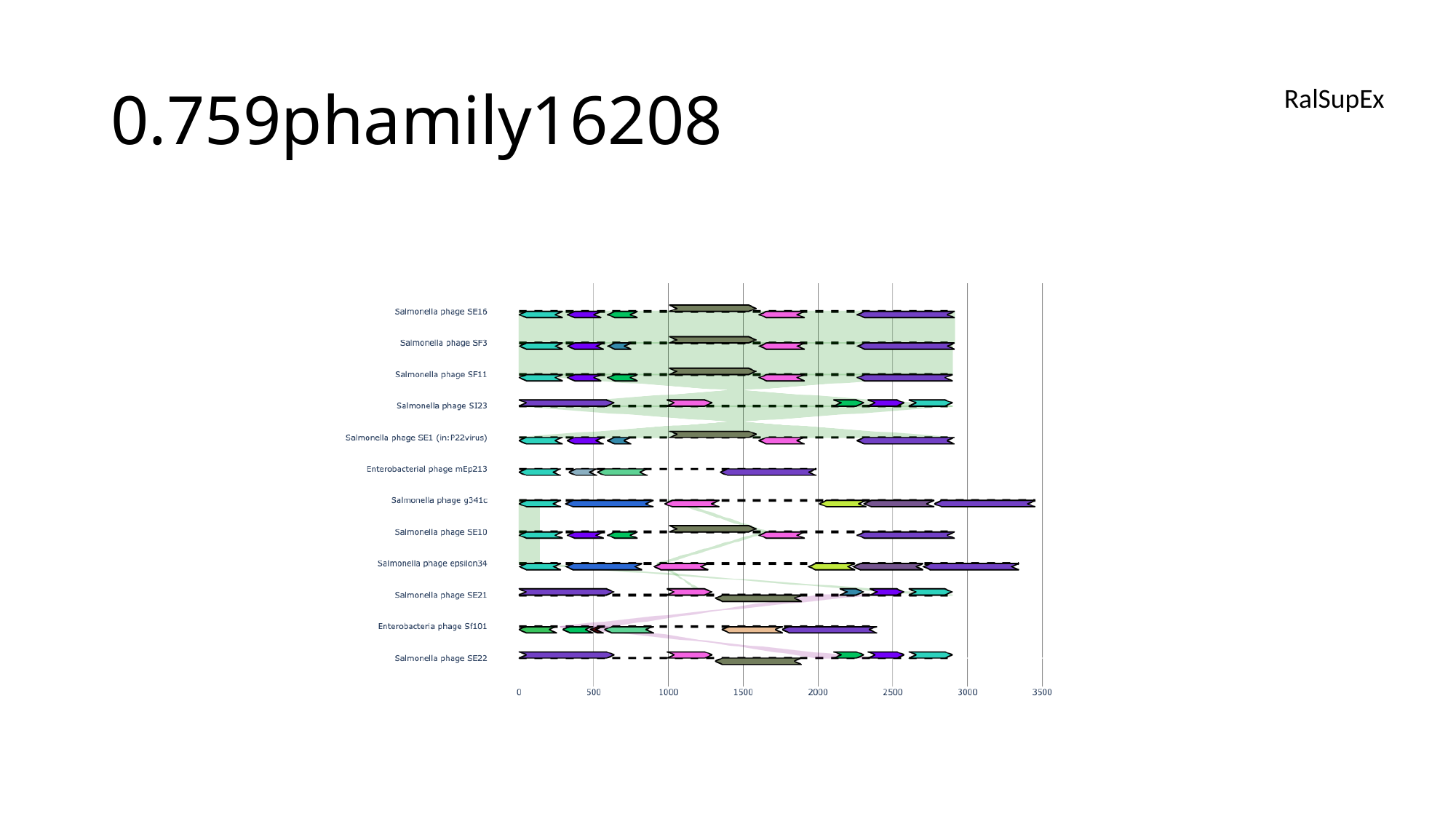

#### 0.759phamily16208
RalSupEx

### Slide 185
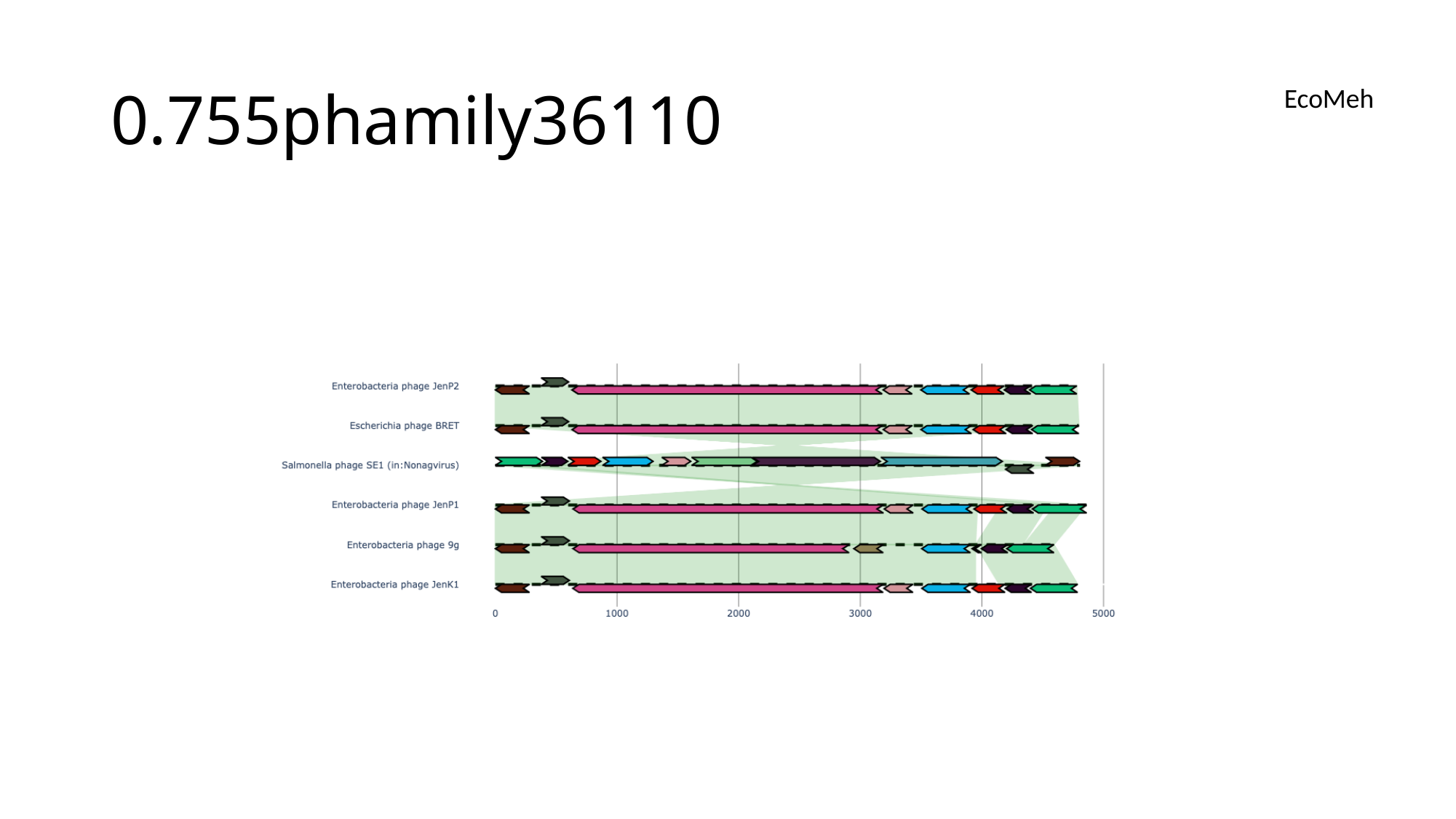

#### 0.755phamily36110
EcoMeh

### Slide 186
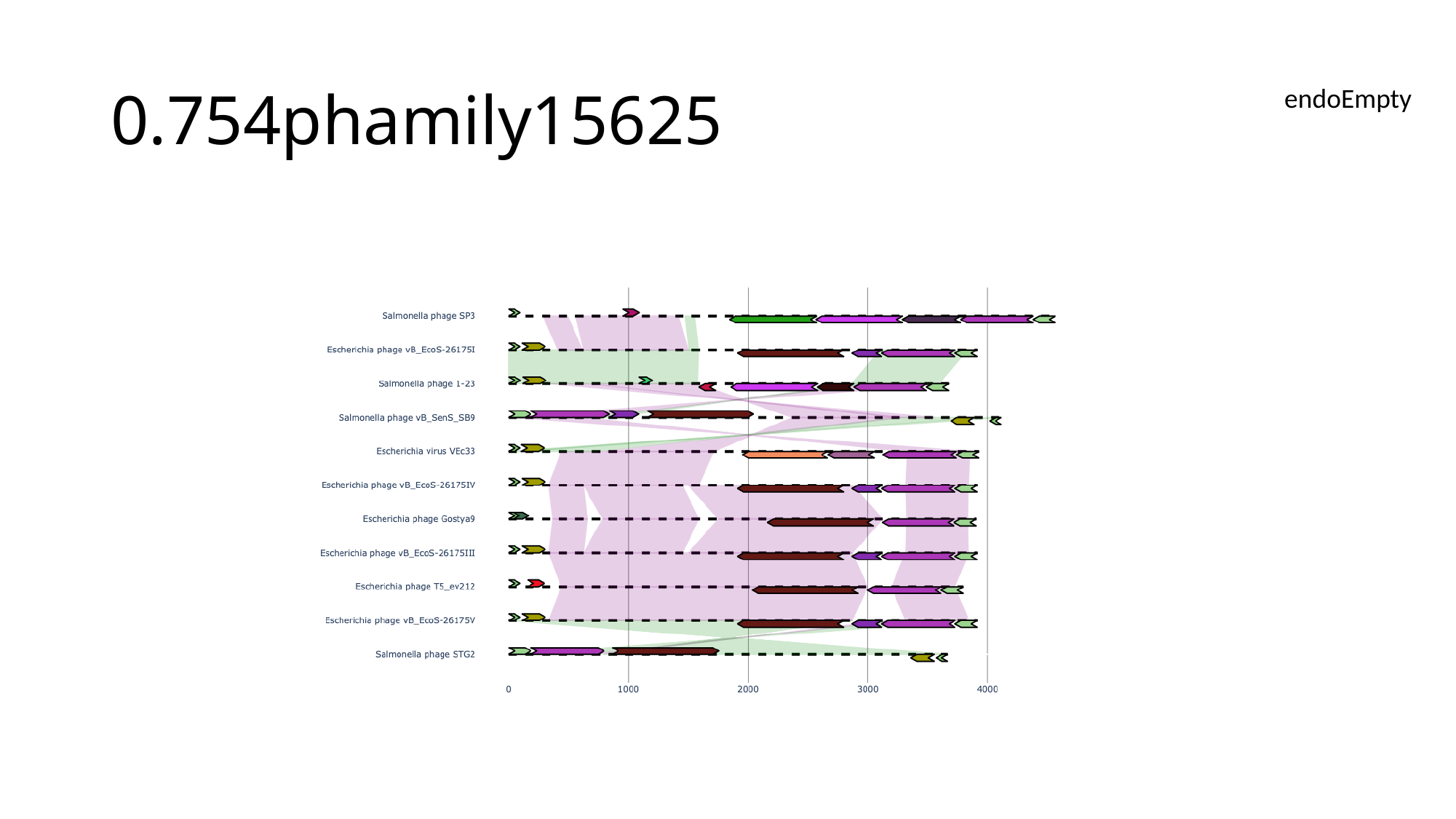

#### 0.754phamily15625
endoEmpty

### Slide 187
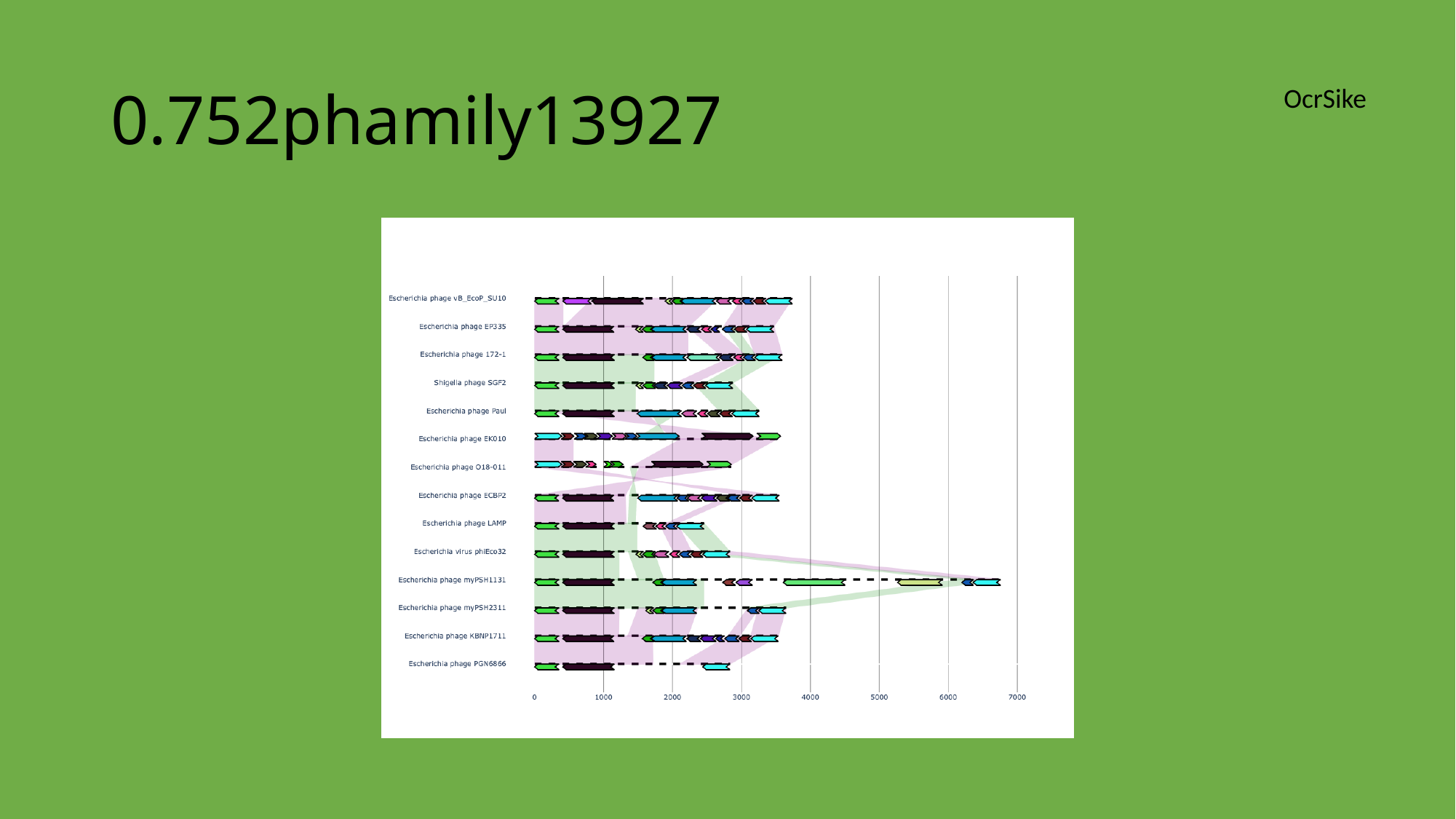

#### 0.752phamily13927
OcrSike

### Slide 188
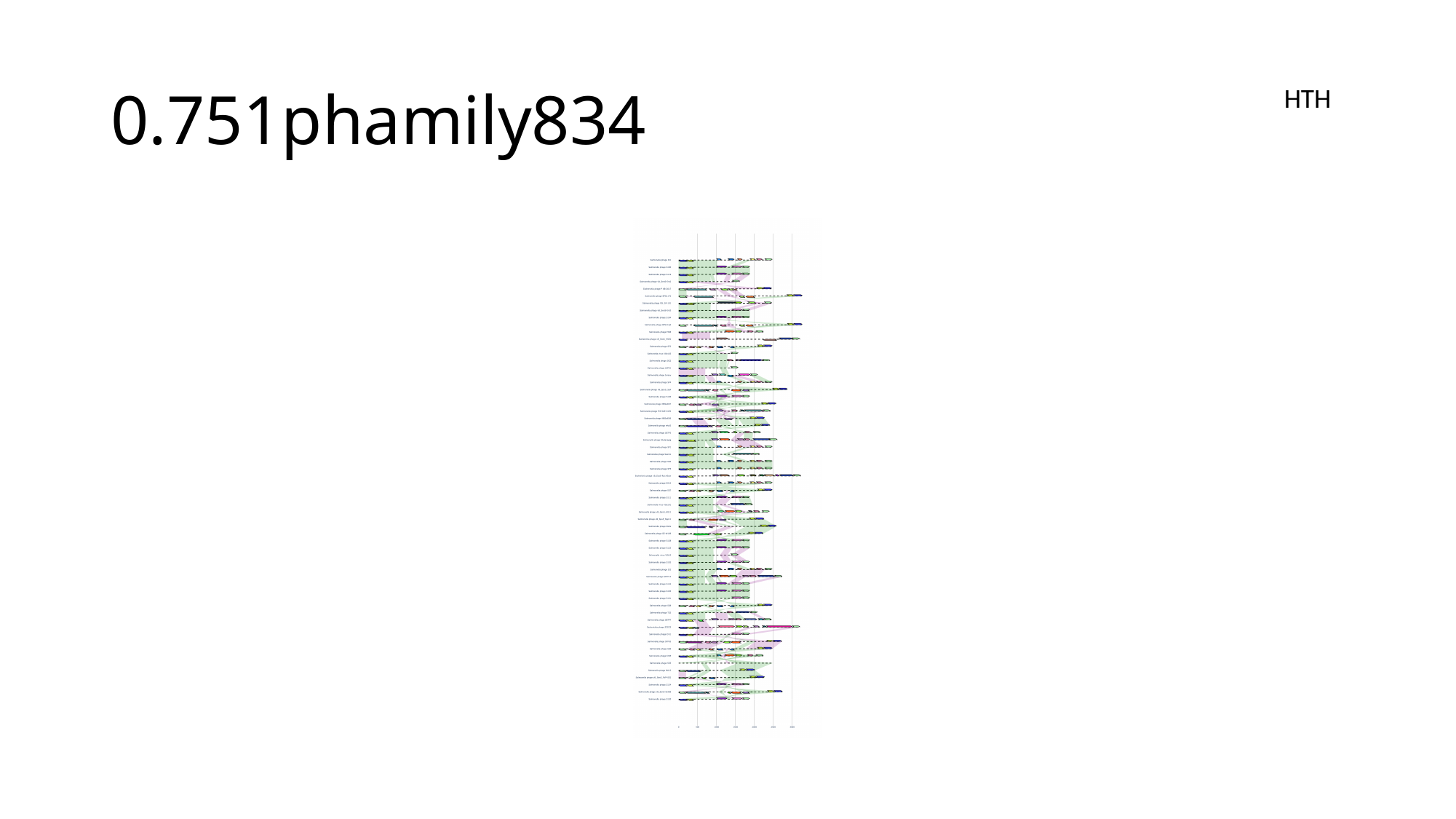

#### 0.751phamily834
HTH

### Slide 189
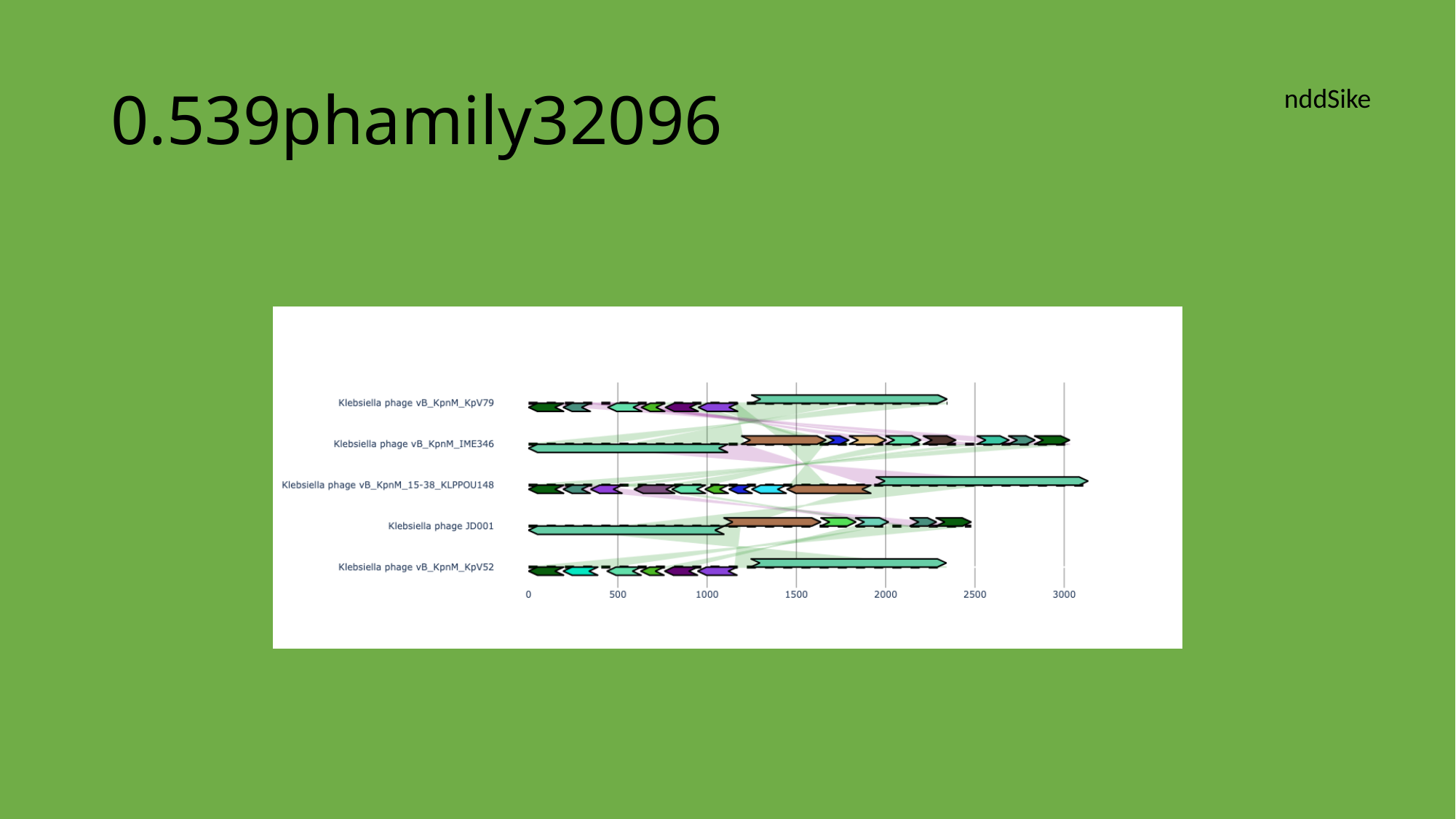

#### 0.539phamily32096
nddSike

### Slide 190
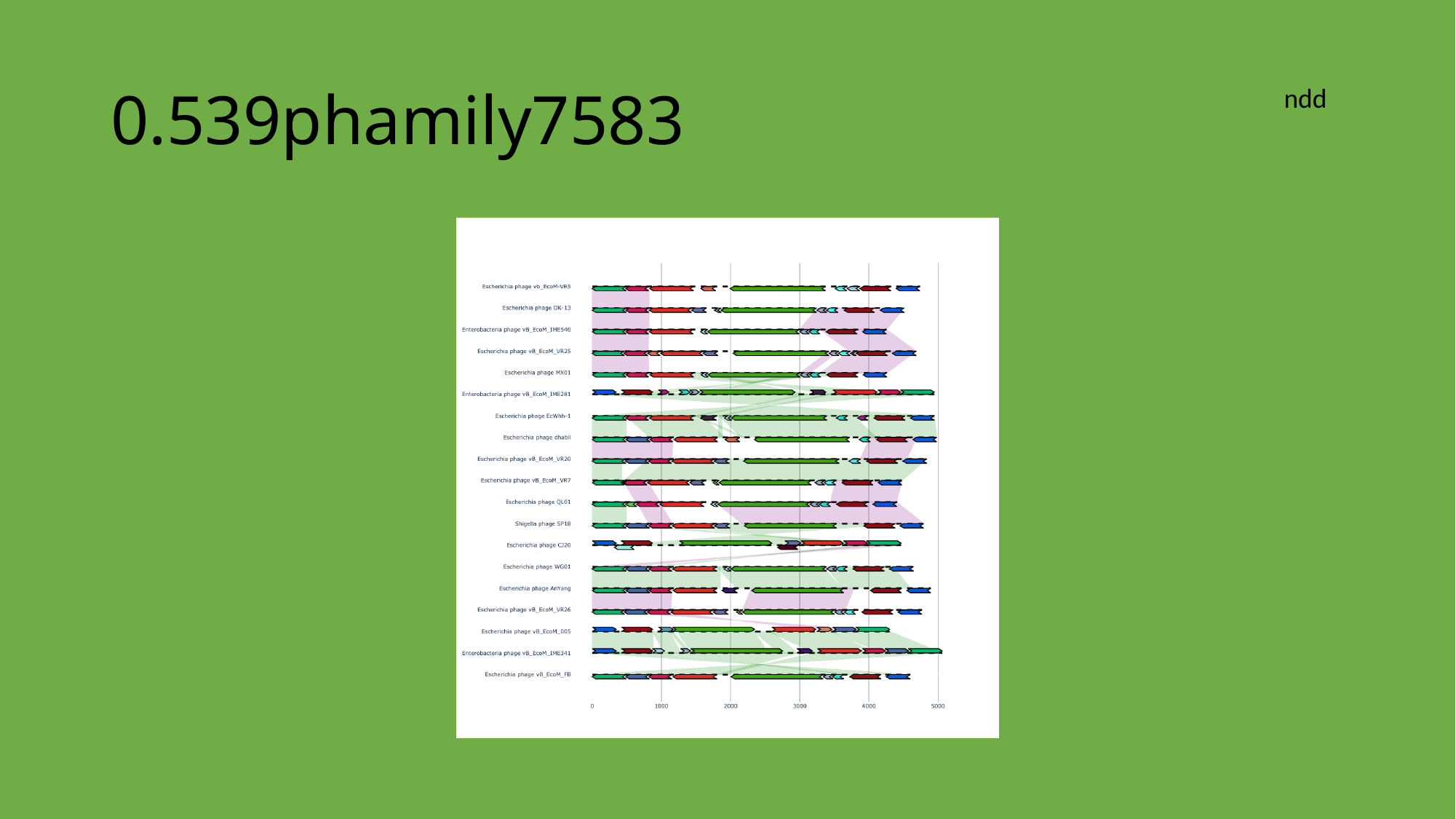

#### 0.539phamily7583
ndd

### Slide 191
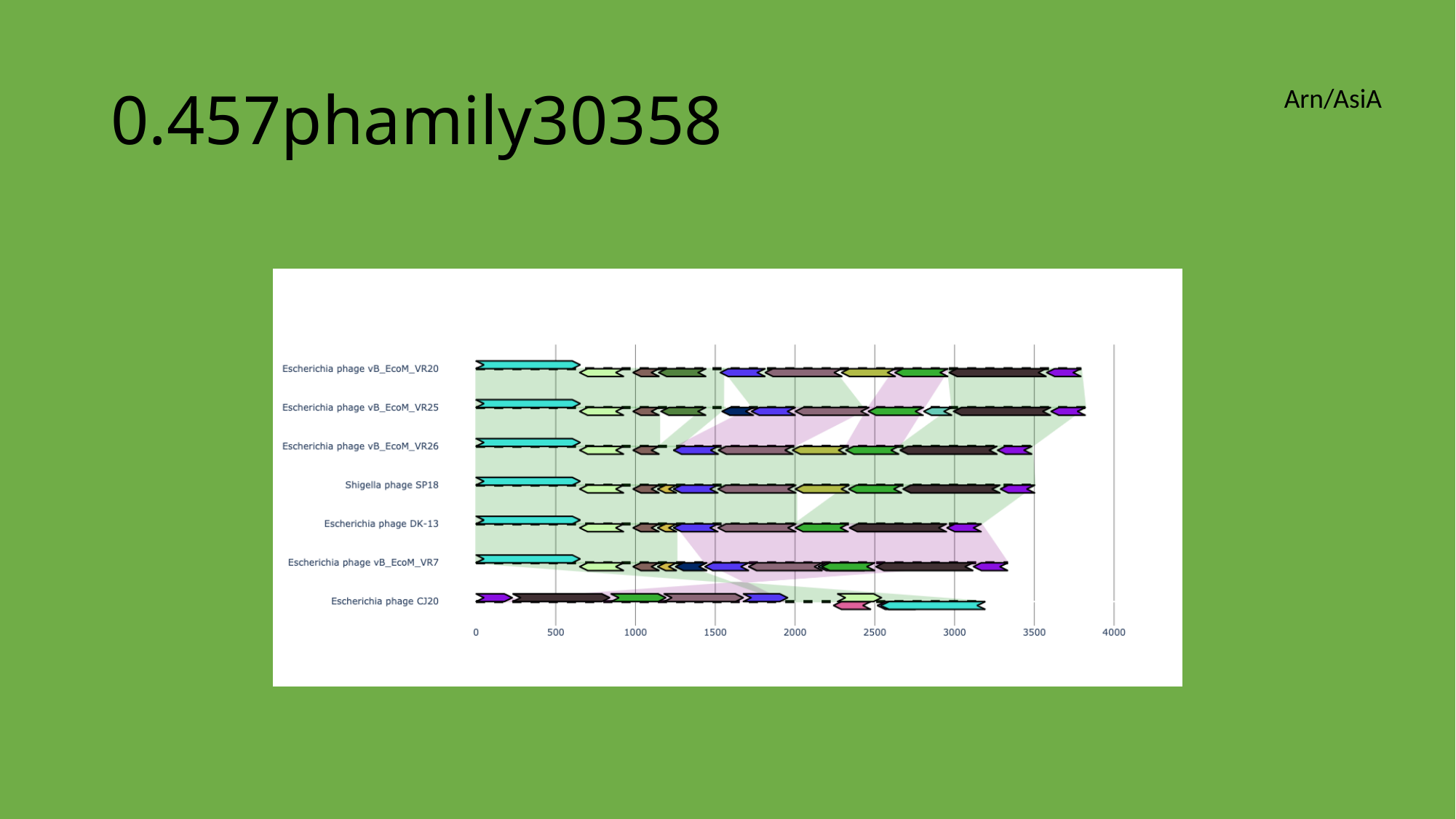

#### 0.457phamily30358
Arn/AsiA

### Slide 192
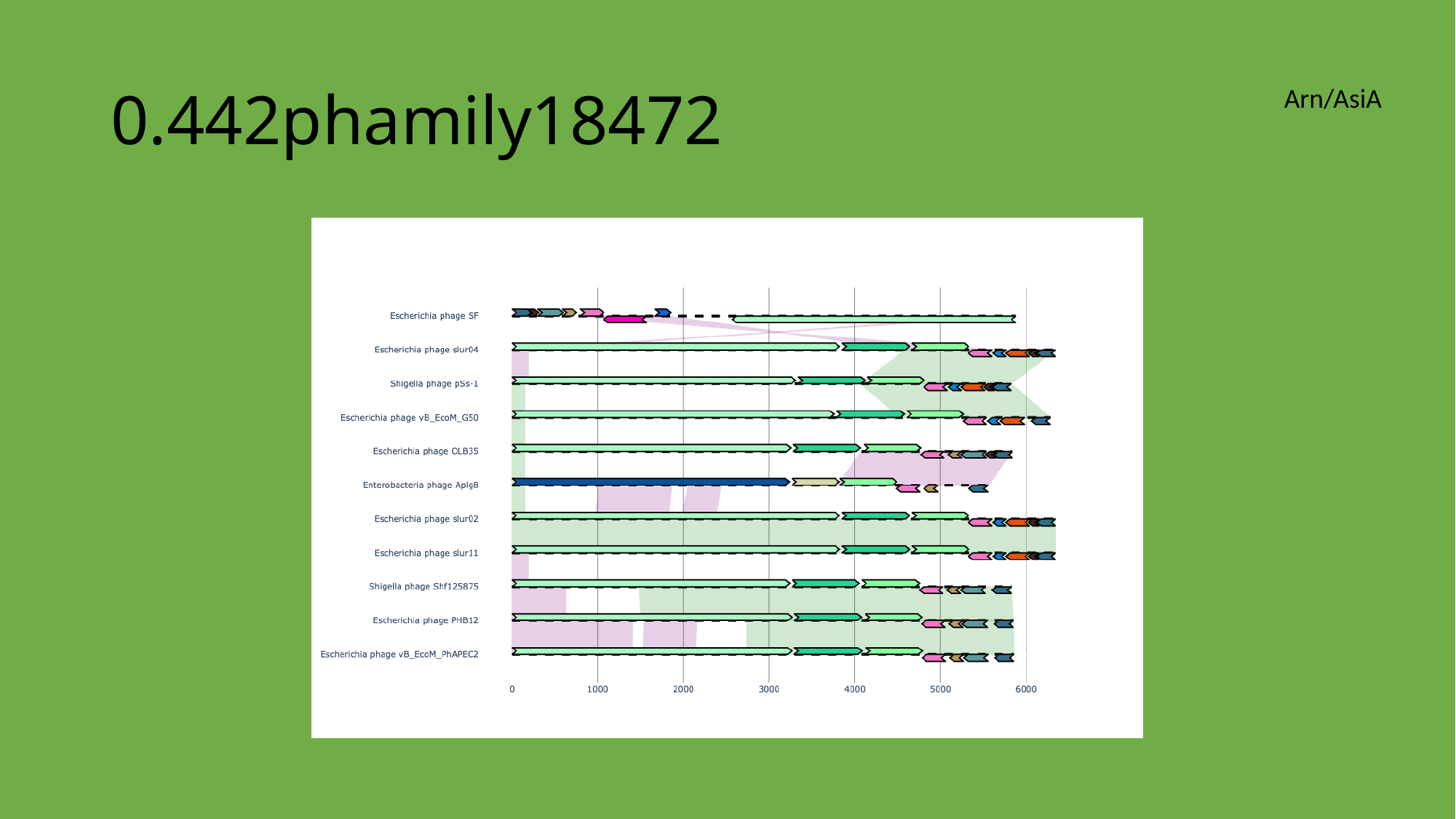

#### 0.442phamily18472
Arn/AsiA

### Slide 193
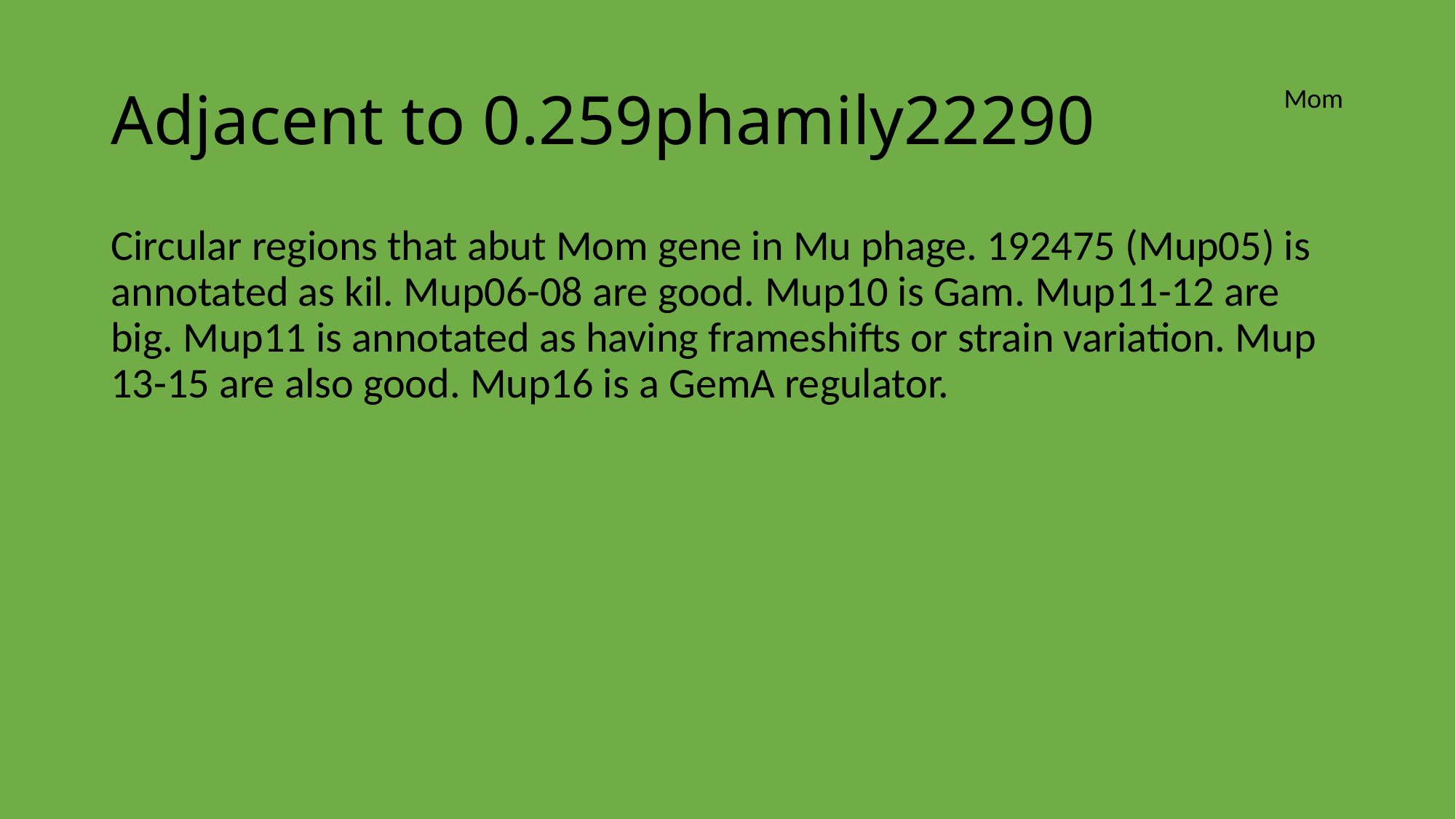

#### Adjacent to 0.259phamily22290
Mom
Circular regions that abut Mom gene in Mu phage. 192475 (Mup05) is annotated as kil. Mup06-08 are good. Mup10 is Gam. Mup11-12 are big. Mup11 is annotated as having frameshifts or strain variation. Mup 13-15 are also good. Mup16 is a GemA regulator.
