## Supplementary File 10 for "Activation of programmed cell death and counter-defense functions of phage accessory genes"

### Slide 1
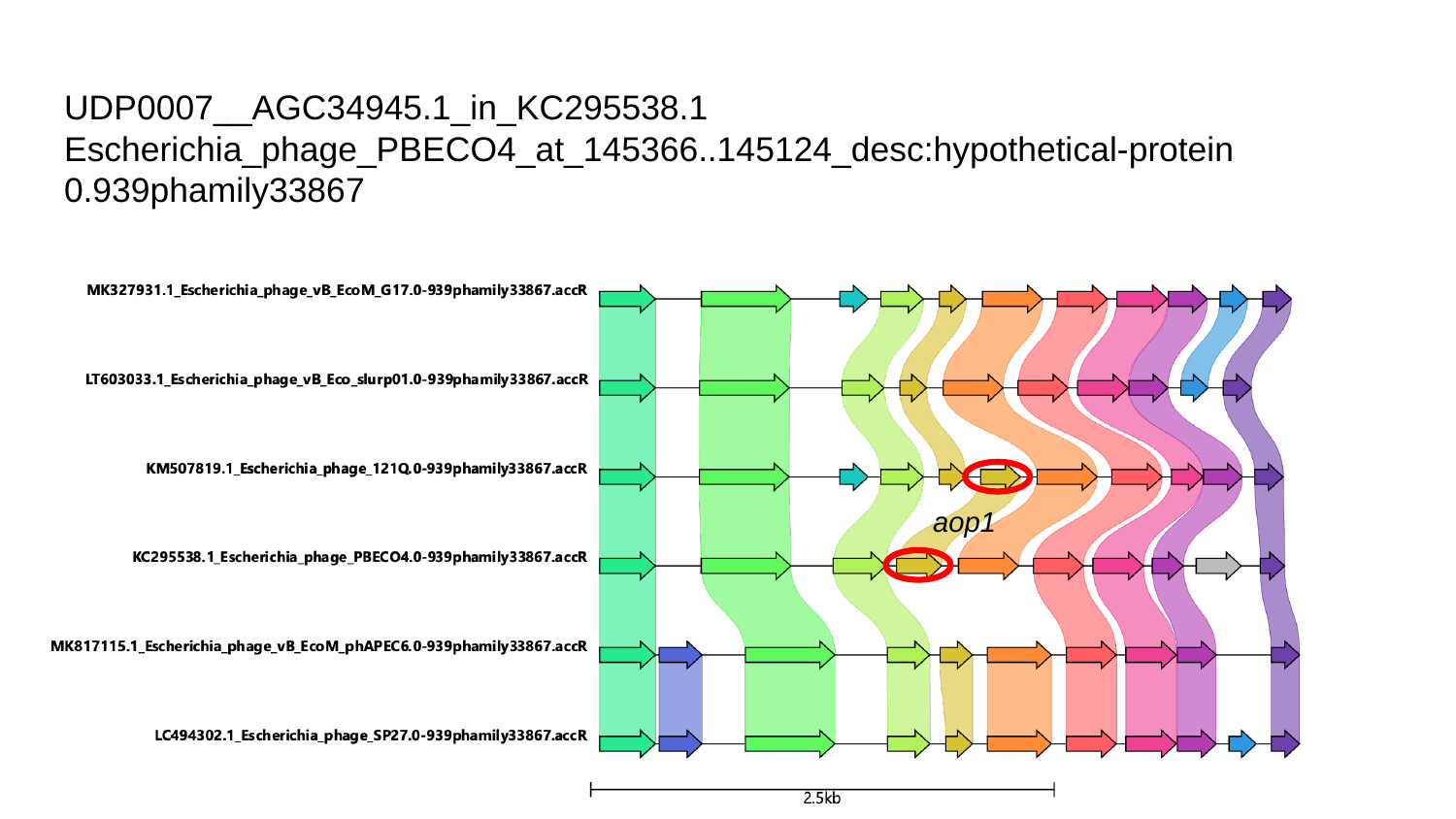

#### UDP0007__AGC34945.1_in_KC295538.1
Escherichia_phage_PBECO4_at_145366..145124_desc:hypothetical-protein
0.939phamily33867
aop1

### Slide 2
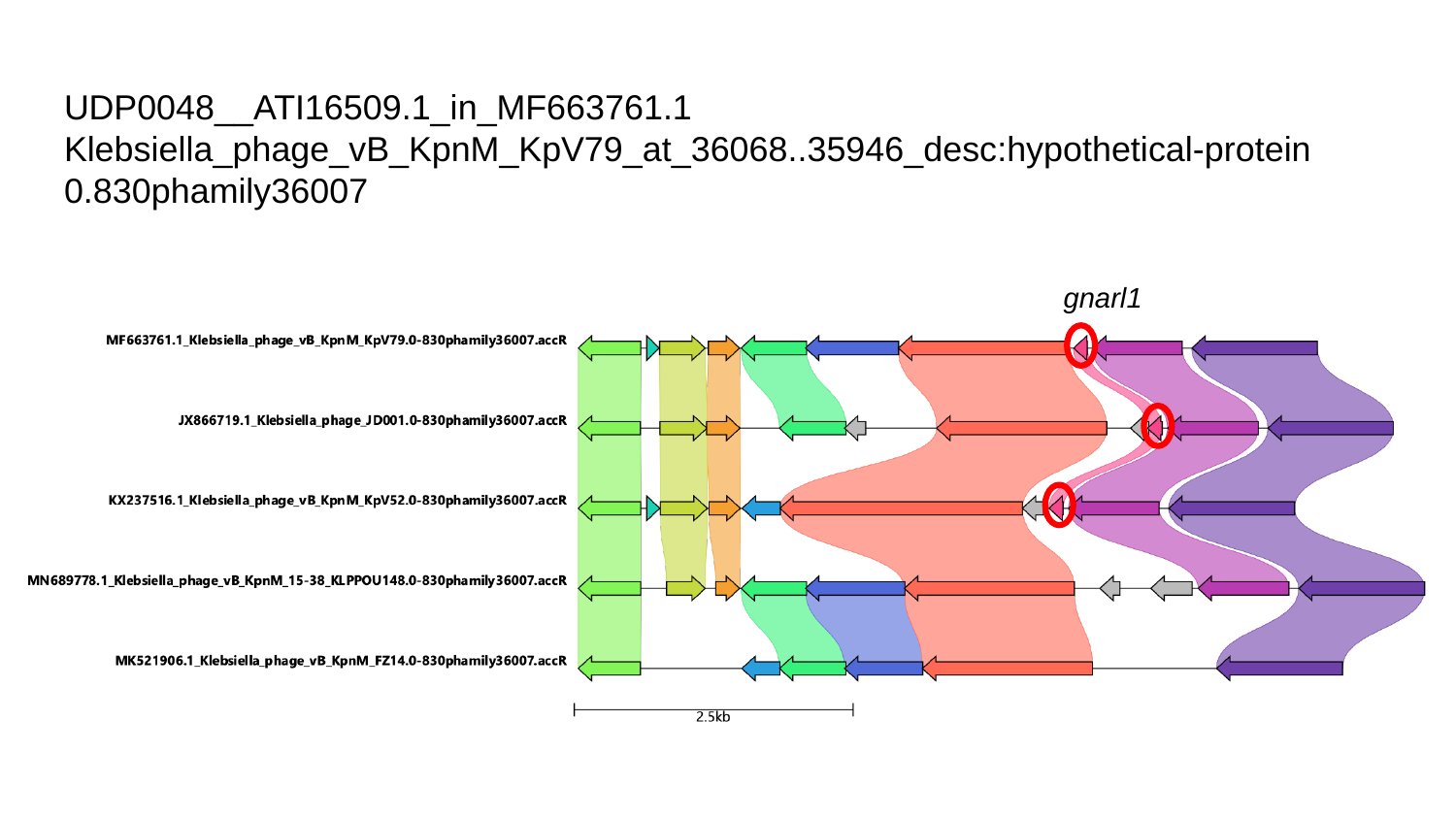

#### UDP0048__ATI16509.1_in_MF663761.1
Klebsiella_phage_vB_KpnM_KpV79_at_36068..35946_desc:hypothetical-protein
0.830phamily36007
gnarl1

### Slide 3
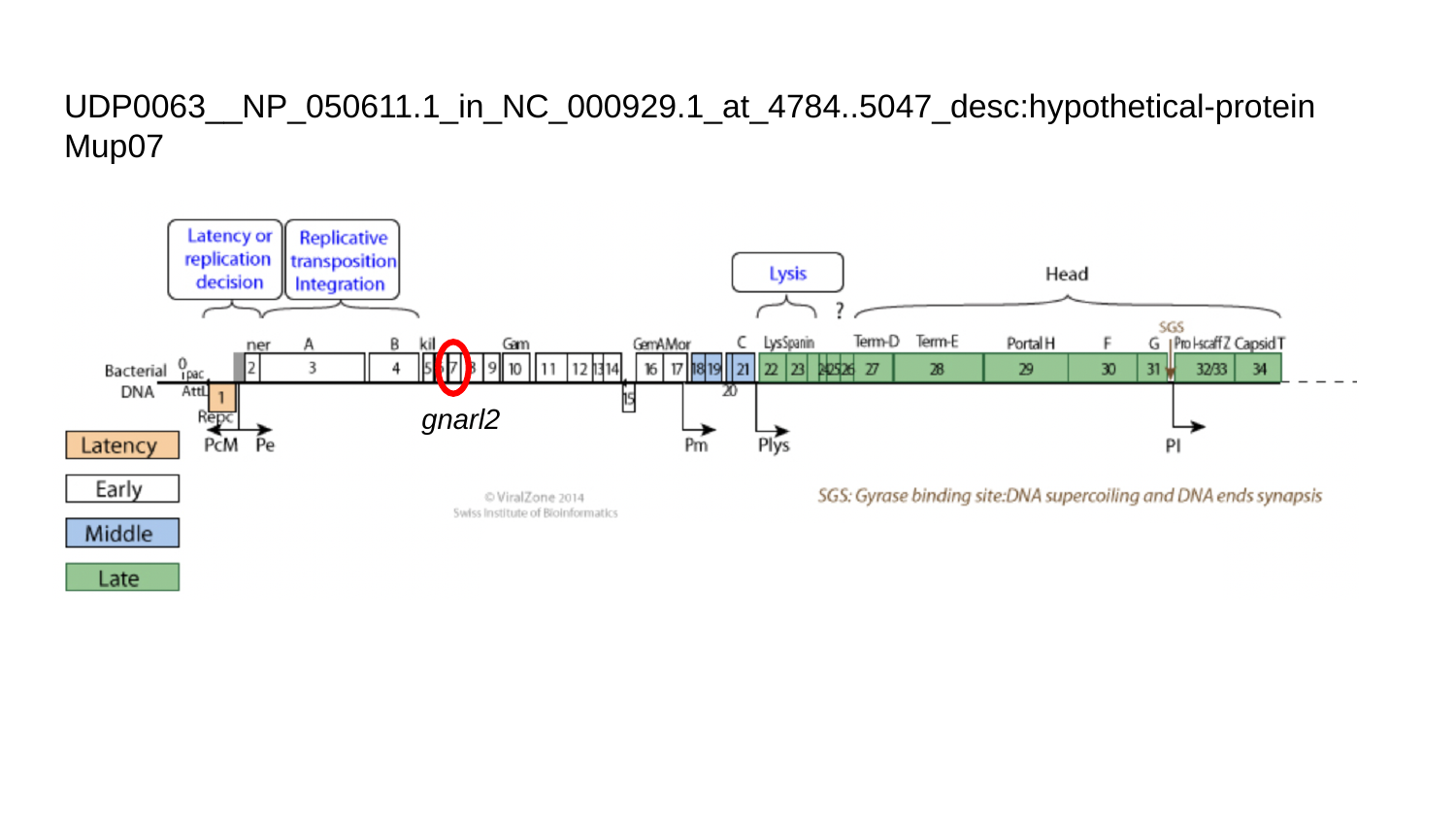

#### UDP0063__NP_050611.1_in_NC_000929.1_at_4784..5047_desc:hypothetical-protein
Mup07
gnarl2

### Slide 4
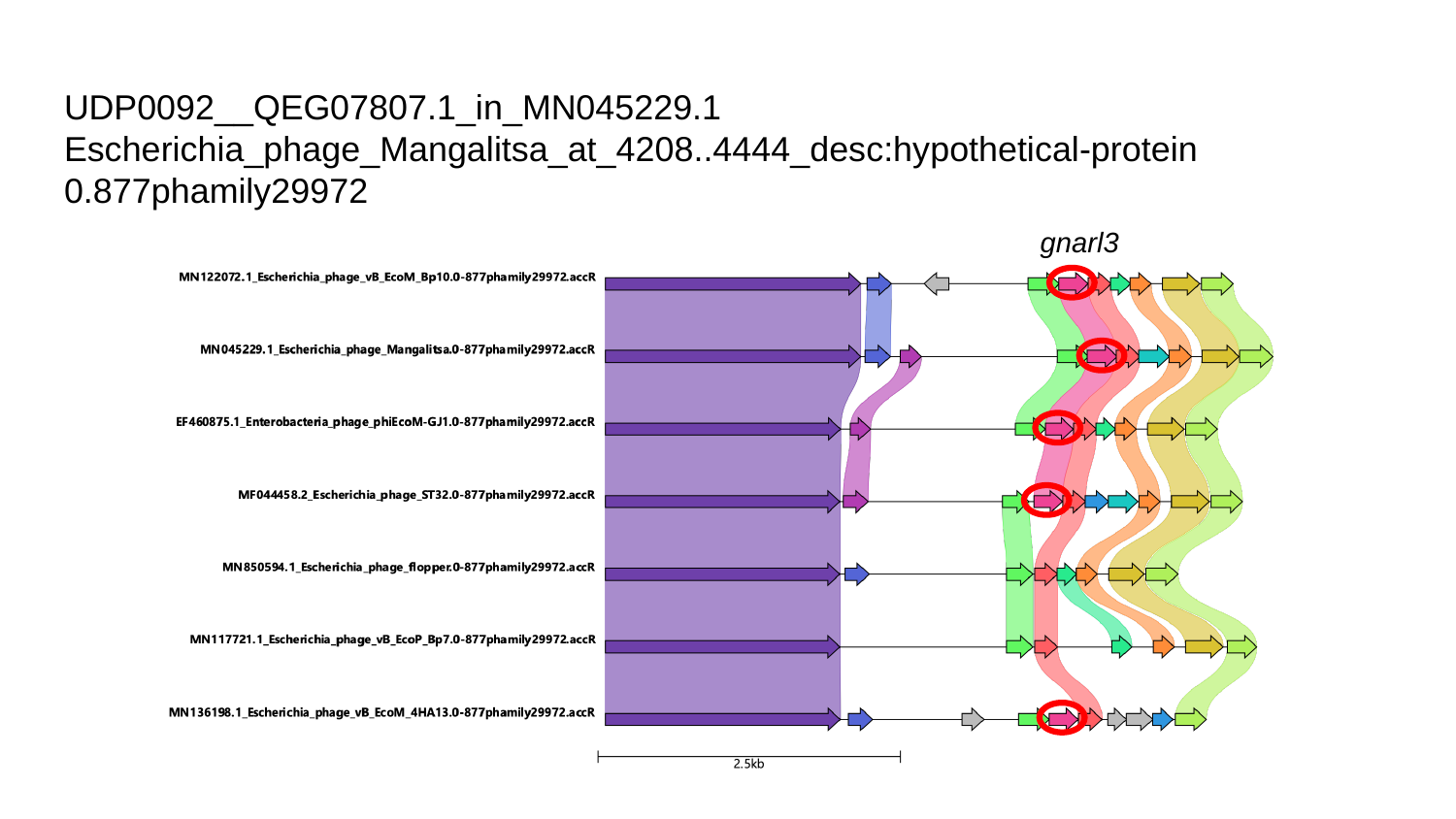

#### UDP0092__QEG07807.1_in_MN045229.1
Escherichia_phage_Mangalitsa_at_4208..4444_desc:hypothetical-protein
0.877phamily29972
gnarl3

### Slide 5
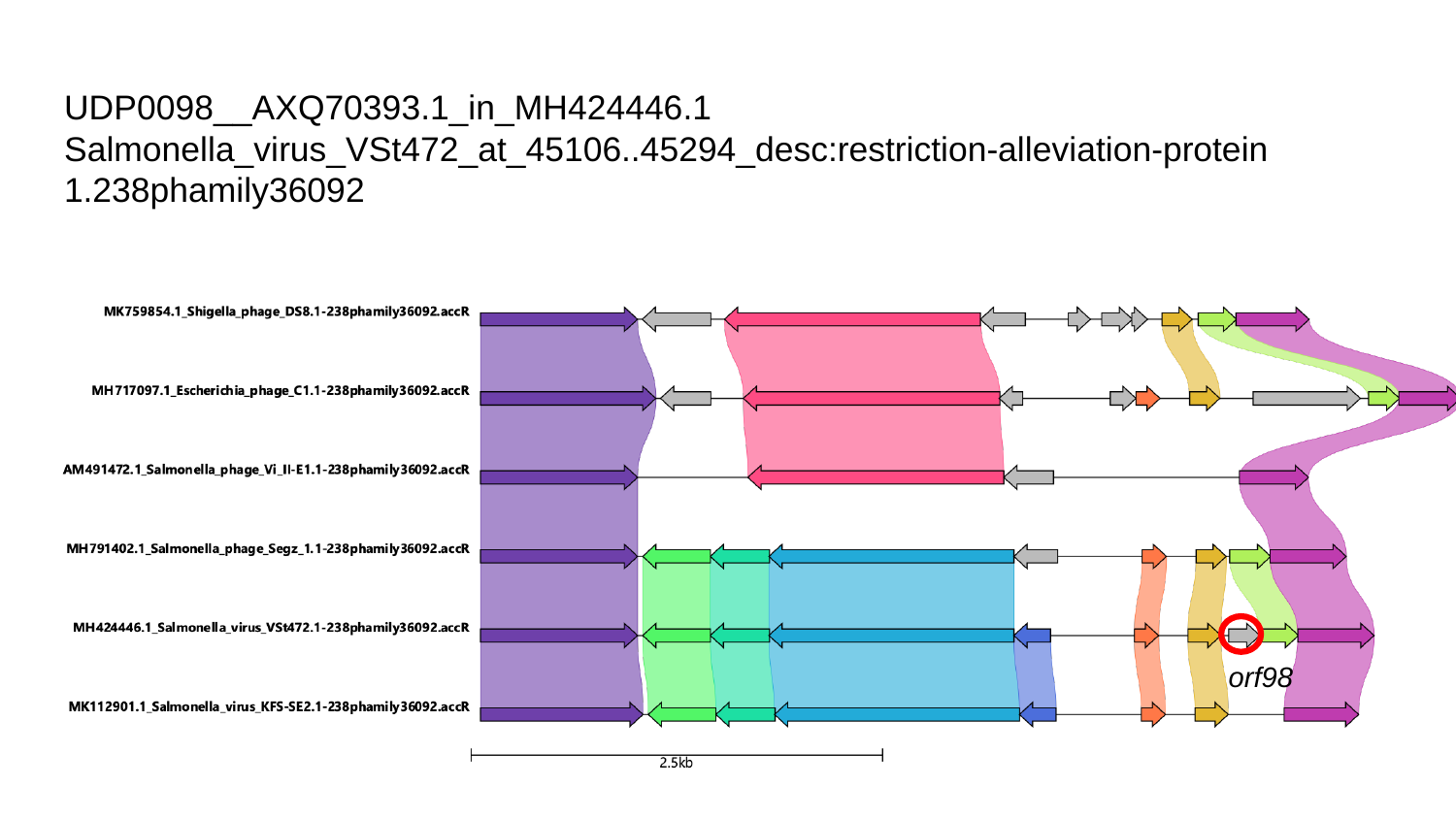

#### UDP0098__AXQ70393.1_in_MH424446.1
Salmonella_virus_VSt472_at_45106..45294_desc:restriction-alleviation-protein
1.238phamily36092
orf98

### Slide 6
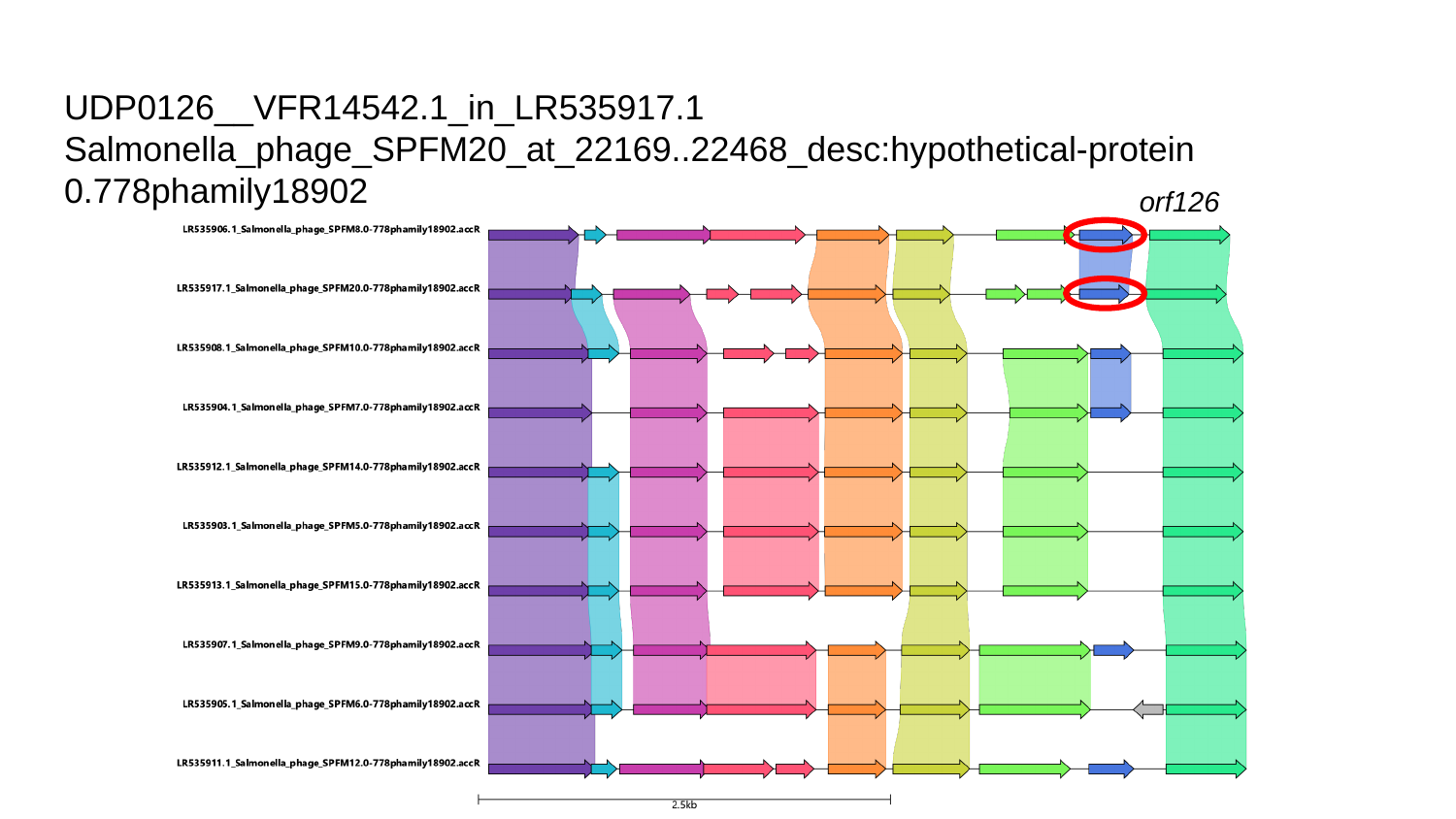

#### UDP0126__VFR14542.1_in_LR535917.1
Salmonella_phage_SPFM20_at_22169..22468_desc:hypothetical-protein
0.778phamily18902
orf126

### Slide 7
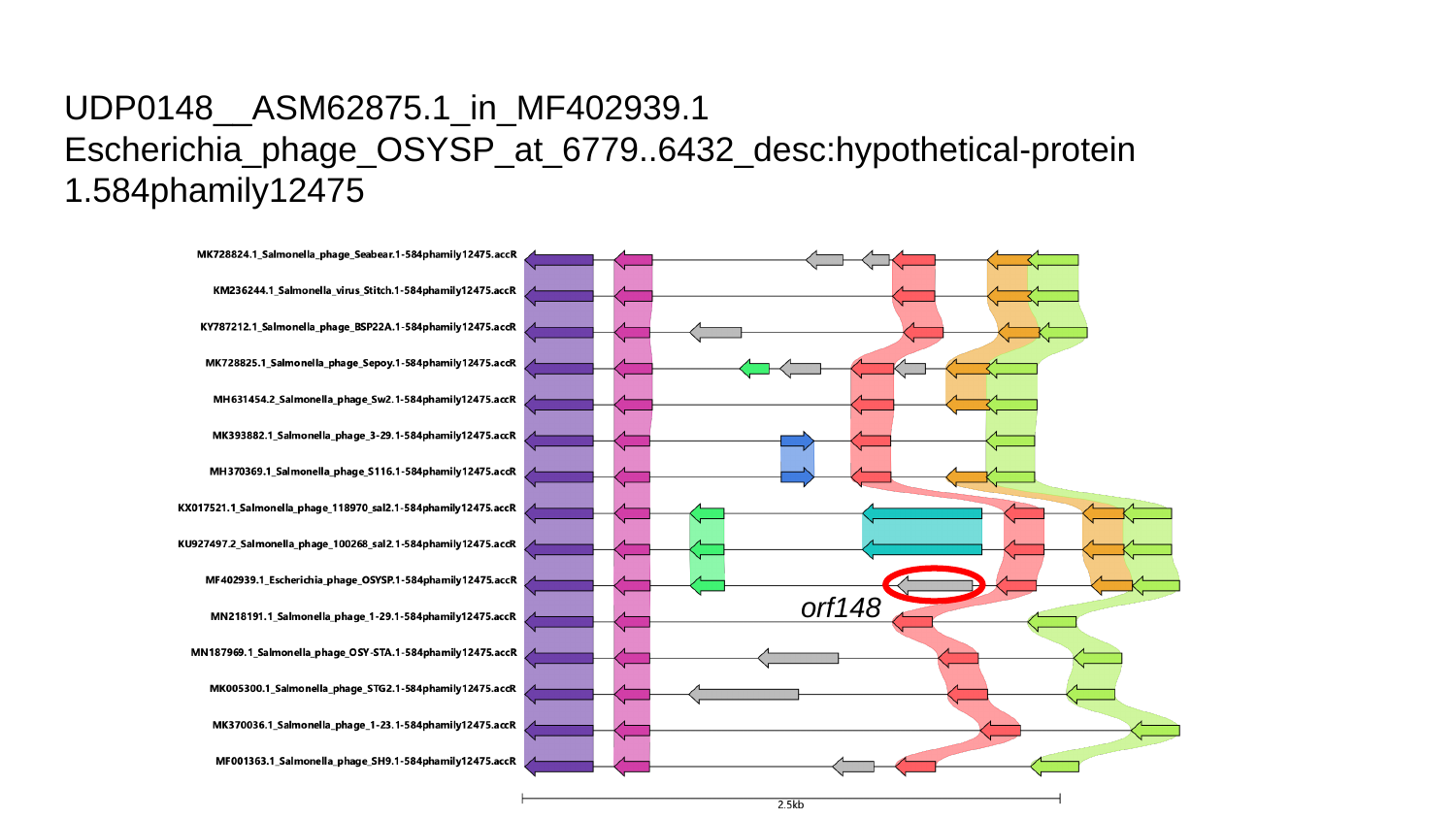

#### UDP0148__ASM62875.1_in_MF402939.1
Escherichia_phage_OSYSP_at_6779..6432_desc:hypothetical-protein
1.584phamily12475
orf148
